# An alternative HIV-1 non-nucleoside Reverse Transcriptase inhibition mechanism: Targeting the p51 subunit

**DOI:** 10.1101/699470

**Authors:** Kwok-Fong Chan, Chinh Tran-To Su, Alexander Krah, Ser-Xian Phua, Peter J. Bond, Samuel Ken-En Gan

**Author notes:** Both authors contributed equally to the work (Kwok-Fong Chan, Chinh Tran-To Su). Corresponding author (S. K-E. G.).

## Abstract

HIV drug resistance continues to demand for alternative drug targets. Since Reverse Transcriptase (RT) is unique and critical for the virus life cycle, it is a rational target that is likely to have less off-target effects in humans. Serendipitously, we found two chemical compound scaffolds from the NCI Diversity Set V that inhibited the HIV1- RT catalytic activity. Computational structural analyses and subsequent experimental testing demonstrated that one of the two chemical scaffolds binds to a novel location in the HIV-1 RT p51 subunit, interacting with residue Y183 that has no known association with previously reported drug resistance. This finding leads to the notion of a novel druggable site on p51 for a new class of non-nucleoside RT Inhibitors that may inhibit HIV-1 RT allosterically. Although inhibitory activity was shown experimentally only to be in the hundreds micromolar range, the scaffolds serve as a proof-of-concept of targeting HIV RT p51, with the possibility for medical chemistry methods to be applied to improve the inhibitory activity, towards a functioning drug.

## INTRODUCTION

Human immunodeficiency virus (HIV) remains one of the major global pandemics with millions infected worldwide [1]. Current therapies including pre-exposure prophylaxis (PrEP) for prevention and cocktail antiretroviral therapy (ART) for treatment target stages of the HIV life cycle, encompasses of: Protease Inhibitors (PIs) to interfere with Protease binding to its substrate Gag and Gag-Pol during viral maturation [2]; Reverse Transcriptase Inhibitors (RTIs) to inhibit viral DNA production [3]; Integrase Inhibitors to block viral DNA insertion into the host genome [4]; fusion inhibitors [5], chemokine receptor antagonists [6], and attachment inhibitors [7].

While integrase inhibitors are currently widely accepted and recommended by WHO [8] as the first line of treatment, there are safety issues for pregnant women. On the other hand, the RTI drug class, comprising of both the nucleoside (NRTI) and non-nucleoside (NNRTI) inhibitors, are safer and more affordable. RTIs target the Reverse Transcriptase (RT) enzyme that is unique to viruses, making it a safer target and more suitable than integrase inhibitors for children, pregnant women, or individuals being treated for tuberculosis with rifampicin. Recently, a two-drug switch regimen [9] that involves a combination of NNRTI (Rilpivirine) and PI (cobicistat-boosted darunavir [9]) has been shown to be effective, thus RTIs remain a recommended component [8] in the standard triple-inhibitor ART.

Both NRTI and NNRTI are used in ART, with each inhibiting via different modus operandi. NRTIs are nucleotide analogues and bind to the nucleotide binding site to competitively block reverse transcription [10], leading to termination of the DNA elongation process [11]. In contrast, NNRTIs bind to a site located near to the polymerase active site, disrupting the structural alignment of the deoxyribonucleotide triphosphates (dNTPs) and template/primer substrates at the RT “primer grip”, thus inducing effects on the polymerase active site indirectly and non-competitively [12] to inhibit reverse transcription [13]. Unlike NRTIs, which mimic nucleotides to directly interfere with cellular replication machinery [14], NNRTIs can vary in structure, generally do not exhibit off-target effects, and have been found to non-competitively induce conformational changes in the RT structure [15]. Therefore, NNRTI drug resistant mutations typically occur at the less conserved drug-binding pocket [16].

HIV-1 RT is a heterodimer of the p66 and p51 subunits. The larger p66 subunit contains five subdomains: fingers, palm, thumb, connection, and RNase H. Despite having an identical sequence to that of p66 excluding the RNase H subdomain, the p51 subunit forms a more compact structure [17]. Despite targeting hot spots on the p51 domain [18, 19], no clinical inhibitors to p51 have been found to date.

In our search for alternative NNRTIs against HIV-1 RT, we serendipitously found two inhibitory chemical scaffolds that exhibited RT inhibition. Despite inhibiting only at hundred micromolar range, one of the compounds elicited the possibility of non-canonical targeting at the RT p51 subunit. We therefore attempted to investigate the underlying mechanism for their interactions on the RT, using computational docking and molecular dynamics (MD) simulations. The inhibition thus demonstrated a possible alternative HIV-1 RT inhibition mechanism via the RT p51.

## Materials & Methods

### RNA Extraction

RNA was extracted from EXPI293F (Invitrogen, Cat no. A14527) cells using TRIzol Reagent (ThermoFisher Scientific, Cat. No.: 15596018) according to the manufacturer’s recommendations. The RNA was treated with RNase-free DNase I (Roche, Cat. No.: 04716728001) and stored at −80 °C.

### Library Reagents Preparation

NCI Diversity Set V chemical compounds from the National Cancer Institute (NCI) Developmental Therapeutic Program’s Open Compound Repository, NIH (http://dtp.cancer.gov) were reconstituted into 1 mM concentrations in dimethyl sulfoxide (DMSO) and stored at −20 °C. The chemicals were initially screened at 300 μM, and potential hit compounds were subsequently diluted to lower concentrations: 150 μM, 30 μM, 3 μM and 0.3 μM.

### cDNA synthesis Reverse Transcription (HIV-1)

All the library compounds were first warmed to 37 °C for 10 min for better solubility prior to addition into the RT reaction mix. For HIV-1 RT, SinoBiological HIV-1 Reverse Transcriptase p51 (isolate HXB2, Cat No.: 40244-V07E) and p66 (isolate HXB2, Cat No.: 40244-V07E1) subunits were both used to form the RT complex for cDNA synthesis. The 20 μL cDNA synthesis mix is as follows: 0.5 μL of random hexamers (ThermoFisher Scientific, Cat No.: SO142, 100μM), 0.5 μL of Oligo d(T)s (ThermoFisher Scientific, Cat No.: SO131, 100 μM), 1.0 μL of Ribolock RNase Inhibitor (ThermoFisher Scientific, Cat No.: EO0381, 40 U/μL), 1.0 μL of dNTP mixture (1st Base, Cat. No.: BIO-5120, 10 mM of each dNTP), 4.0 μL of 5 × RT Buffer (250 mM Tris-HCl pH8.3, 375mM KCl, 15mM MgCl_2_, 0.1M DTT), 2 μL of DEPC-Treated Water, 0.5 μL of RT p51 (0.17 μg/μL), 1.0 μL of RT p66 (0.33 μg/μL), 6.0 μL of candidate inhibitors and 3.0 μg of DNase-treated RNA. The reaction mixes were prepared on ice, followed by incubation in ProFlex 3x 32-Well PCR System (Applied Biosystems) at 25 °C for 18 min, followed by 37 °C for 1 hour, and RT termination at 85 °C for 5 min. Reaction mixtures containing either water or DMSO only without RNA were used as negative controls.

### Polymerase Chain Reaction (PCR) Gene Amplification

The GAPDH PCR reactions were setup in 27.0 μL reactions, containing 13.5 μL of GoTaq Green Master Mix (Promega, Cat No.: M7123), 1.5 μL of Human GAPDH Forward Primer (5’– AGAAGGCTGGGGCTCATTTG–3’), 1.5 μL of Human GAPDH Reverse Primer (5’– AGGGGCCATCCACAGTCTTC–3’) and 2 μL of cDNA template. Thermocycler conditions were set at 95 °C (2 min), 20 cycles of 95 °C (30 s), 50 °C (30 s), and 72 °C (1 min 30 s). The final extension step was performed at 72 °C for 7 min. Agarose gel electrophoresis analysis were carried out on the PCR products using 2% agar gels and GAPDH amplified products. Band sizes were confirmed using GelApp [20].

### Structural Docking

The chemical structures of the two compounds (NSC48443 and NSC127133 from the NCI/DTP Diversity Set V) were retrieved from PubChem library, CID: 241217 and 278037 (particularly from the ligand code 6B3 in this entry) for compounds 1 and 2, respectively. Wild type HIV-1 RT structure (PDB: 3T19) was processed as previously described [21].

We performed rigid blind docking using the program Autodock Vina [22] to find potential binding sites in the HIV-1 RT structure for the two identified compounds. We next performed focused docking with flexible treatment of side chains for residues identified as part of the binding sites (as receptor) derived from the initial blind docking. The compound bound conformations that were significantly different from others and had favorable docking scores were selected for further analysis and subsequent MD simulations.

### MD simulations of HIV-1 RT complexes

We simulated the whole HIV-1 RT p66/p51 with the selected compound conformations using GROMACS (version 2018) [23], with the CHARMM36m [24] forcefield for the protein, the CHARMM general force field (CGenFF) [25] for the ligand, and the TIP3P [26] water model. Nosé-Hoover [27] and Parrinello-Rahman [28] thermostat and barostat were used to maintain constant temperature and pressure at 300 K and 1 bar, respectively. The particle mesh Ewald (PME) method was used for electrostatics, with a real-space cut-off of 12 Å. A 12 Å cut-off was used to calculate the van der Waal’s potential, switching the energy function after 10 Å. Bonds involving hydrogen atoms were restrained using the LINCS [29] algorithm. We carried out a 20 ns equilibration, restraining the protein backbone atoms, followed by unrestrained simulations for 100 ns in triplicate.

### Conventional MD simulation analysis

Hydrophobic contacts were calculated using a 4.0 Å cut-off between the ligand heavy atoms and protein carbon atoms. The number of water molecules around the ligand was determined using a cut-off of 3.5 Å between the ligand heavy atoms and water oxygen atoms. Hydrogen bonds were calculated between donor and acceptors of the ligand and the protein using a cut-off of 3.5 Å, with a maximum angle between donor – hydrogen atom and acceptor of 30°. To determine stable positions of the ligand in each binding site, we calculated distances between the centre of mass of the ligand and of the binding site residues, which were defined via the LigPlot+ program [30].

### Binding energy calculations

The molecular mechanics Poisson-Boltzmann Surface Area (MM-PBSA) method was applied to calculate the binding energies of the compounds and their corresponding binding sites on the HIV-1 RT subunits, using *g_mmpbsa* [31]. Various internal dielectric constants of the solute [32–34] (ε_in_), were used to estimate the polarizability of the binding sites, e.g. ε_in_ = 2 (hydrophobic), ε_in_ = 8 (polar), or ε_in_ = 20 (highly charged). The external dielectric constant ε_out_ was set to 80 (for water). The calculations were performed using the last 50 ns of each MD simulation trajectory for all independent replicas.

### Analysis of structural conservation among RT proteins

A multiple sequence alignment of the HIV-1 RT sequences with 152 retrovirus RT sequences (retrieved from NCBI RefSeq Databases) was first performed using ClustalW [35]. The resulting alignment was then subjected to the ConSurf server [36, 37] with default parameters to study the conservation of the regions of interest across the RT families.

## RESULTS

### Two compound scaffolds that inhibit HIV-1 RT activity

From 40 compounds of the NCI/DTP Diversity Set V, we serendipitously identified two chemical compounds NSC48443 and NSC127133 (referred to as compounds 1 and 2, respectively) that exhibited inhibition on HIV-1 RT activity using the RT-PCR assay. In these assays, the GAPDH housekeeping gene was used for cDNA synthesis control given its constitutive expression [38].

Compound 1 inhibited the HIV-1 RT cDNA synthesis fully at a concentration of 300 μM and partially at 150 μM, whereas compound 2 was found to partially inhibit HIV-1 RT activity at 300 μM (Figure 1). The PCR band intensities were observed to decrease at higher concentrations for both the compounds. The results were obtained via independent triplicates.

**Figure 1.**
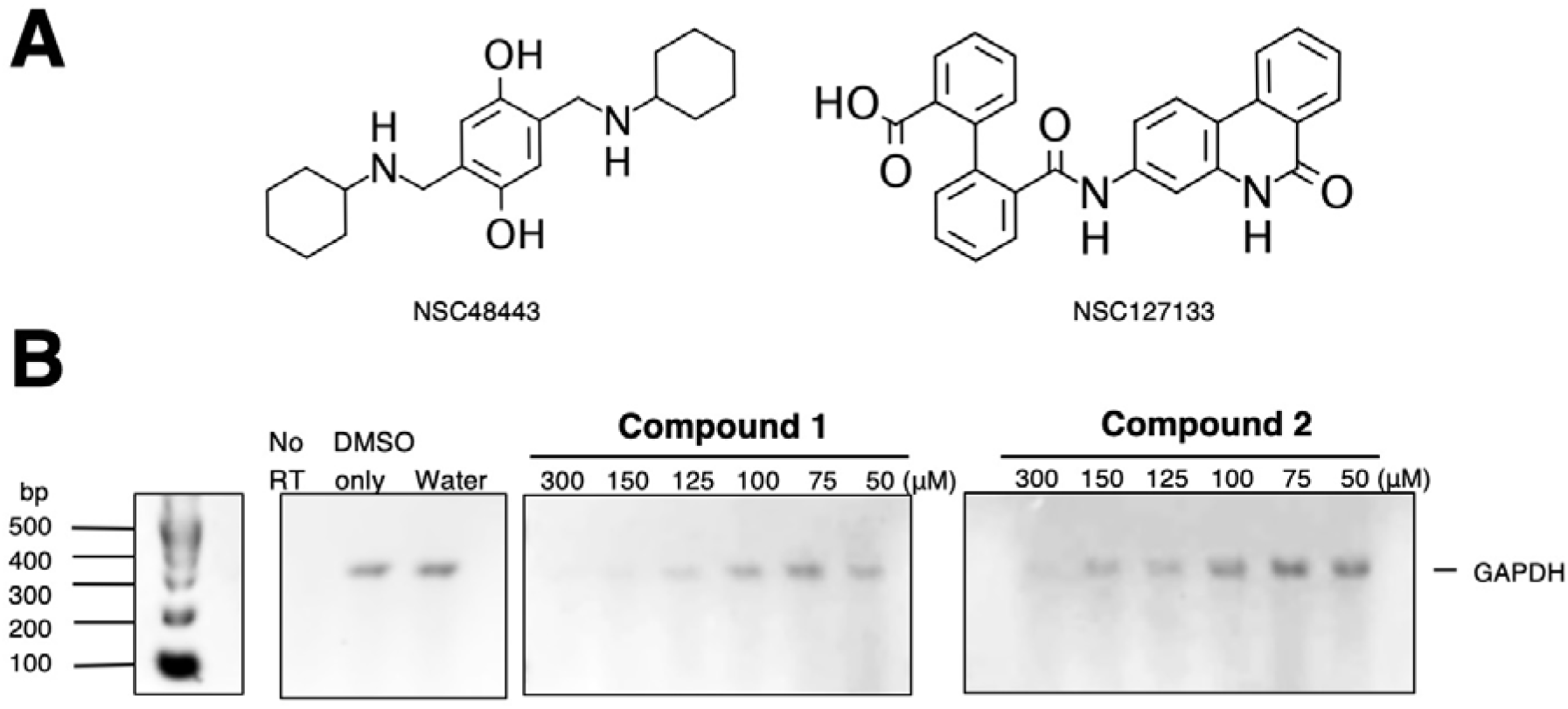
The two compounds that inhibited HIV-1 RT function. (**A**) 2D chemical structures of compounds 1 (NSC48443) and 2 (NSC127133) from the NCI/DTP Diversity Set V. (**B**) RT-PCR using HIV-1 RT assay, representative results of triplicate experiments on GAPDH gene with the two identified compounds. The experiments were performed at various concentrations (in micromolar, μM) from 50 μM to 300 μM of all the compounds.

### Binding sites of the two compounds on HIV-1 RT structure

To identify the binding sites of compounds 1 and 2 on the HIV-1 RT structure heterodimer (including p66 and p51 subunits), blind docking experiments were performed using Autodock Vina [22], followed by MD simulations with various initial bound conformations of each compound as inputs (Supplementary Figure S1). Compound 1 bound to two different sites on the HIV-1 RT (blue in Figure 2A), with one site on the p66 subunit close to a loop located on p51 comprising of residues S134 to P140, and another site on the p51 subunit, whereas compound 2 bound only to one site on the p51 subunit (red in Figure 2A). The interacting residues of the binding sites on p66 (to compound 1) and on p51 (to compound 2) are shown in Figure 2B.

**Figure 2.**
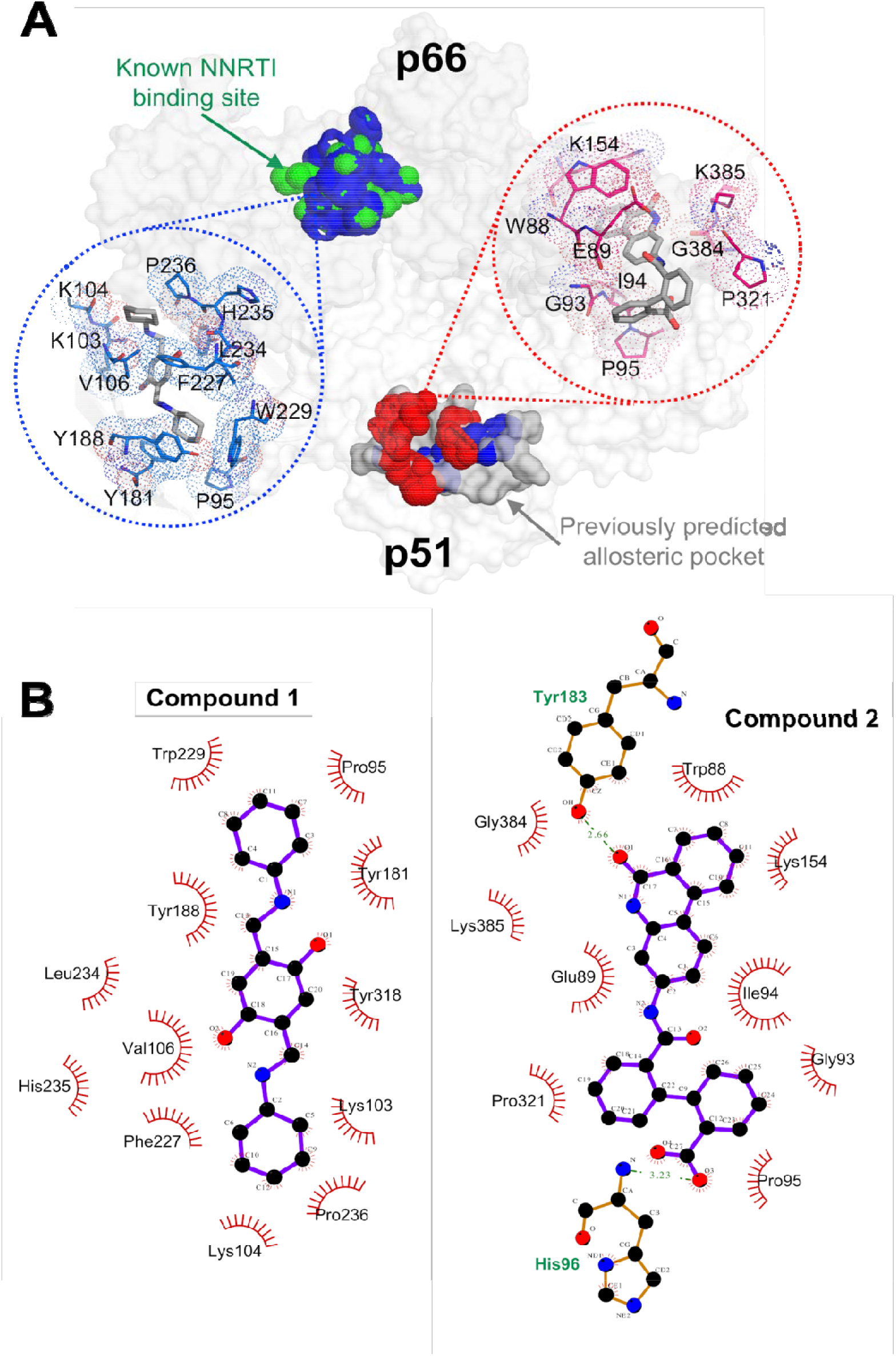
Structural analyses of HIV-1 RT interaction with the two inhibitory compounds. (A) Binding sites of compound 1 (blue surface and lines) and compound 2 (red surface and lines) on HIV-1 RT. One of the two binding sites for compound 1 overlaps with the known NNRTI binding pocket (green, on p66), whereas the other overlaps with our previously predicted allosteric pocket [21] (grey surface on p51), at which compound 2 was also found to bind (red). The two chemical compounds are presented in grey sticks within the binding pockets. (B) 2D presentation of the interactions between the compounds and their binding sites, constructed using LigPlot+ [30].

Compound 1 occupied the site on p66 with a more favourable predicted binding energies than the binding site on p51 (Supplementary Table S1), and with a more stable bound conformational state at the p66 site than at the p51 site (Figure 3A, showing the more stable center-of-mass distances between the binding site and the compound during simulations). Given that more hydrophobic contacts but fewer hydrogen bonds were detected between compound 1 and its p66 binding site than the p51 binding site (Supplementary Table S2), hydrophobic interactions are the dominant binding force for association with p66. Hydrophobic contacts were observed between L100, K103, V106, Y181, Y188, P225, F227, L234, P236 and Y318, along with less prominent interactions between P95, S105 and W229. Unstable/weak hydrogen bonds were detected between compound 1 and residue K101 (Supplementary Figure S2).

**Figure 3.**
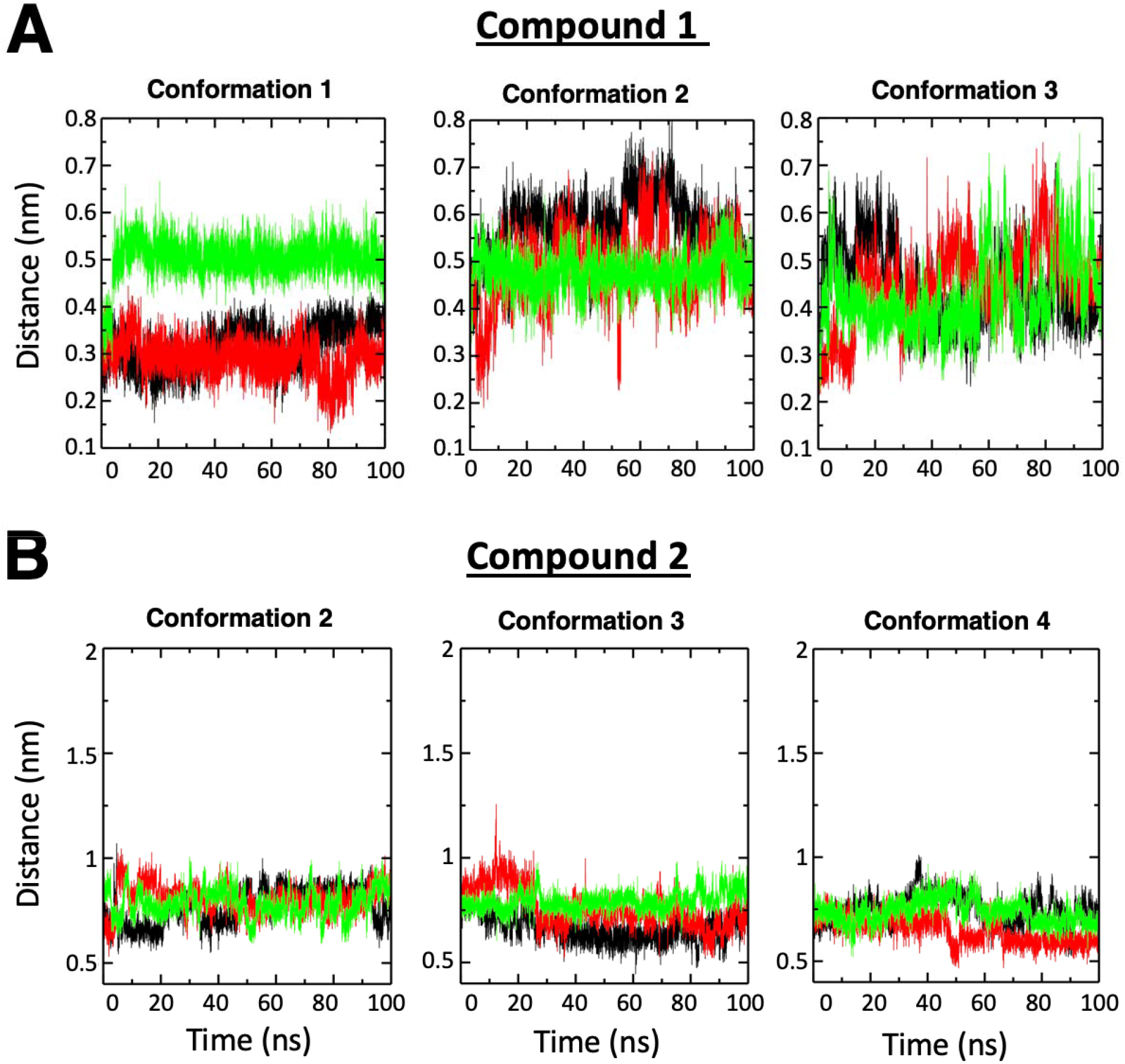
The center-of-mass distances between ligand and the respective binding site on RT during MD simulations. Data are shown for (A) compound 1 and (B) compound 2, calculated using independent triplicates (black, green and red) of 100 ns trajectories. For compound 2, only the results of the three setups that exhibited stable ligand binding (i.e. conformation 2, 3, and 4) are shown. The remaining data are shown in Supplementary Figure S2.

On the other hand, compound 2 maintained stable binding to the p51 subunit in three out of six successful setups for the MD simulations – each setup had a unique initial compound 2 conformation (Figure 2 and Supplementary Figure S1). In the three stable conformations, compound 2 exhibited low binding energies to the same site on p51, with the lowest in conformation 3 (Supplementary Table S3); thus the binding site for conformation 3 was selected as the most likely binding mode. Compared to compound 1 when binding to the p66 subunit, the binding site of compound 2 on p51 is more solvated and polarized due to several polar/charged nearby residues such as H96, K154, and K385. Consistent trends in energy contributions (with increasing interior dielectric constants, shown in Supplementary Table S3) showed that the binding may be dominated by electrostatic interactions. Y183 established strong hydrogen bonds with compound 2, whereas some others (W88, E89, G93, I94, P95, K154, P157 and K183) made hydrophobic contacts with compound 2 (Supplementary Figure S2).

### Experimental inhibition on separate HIV-1 RT p66 and p51 subunits

Our RT-PCR assays using the p51 and p66 subunits separately did not show detectable bands for p51 alone (Figure 4A). On the other hand, bands were detected for the p66-only reactions in the case of compound 2 and the control. There was no band detected in the case of compound 1 added to the p66-only experiment (Figure 4A). We affirmed that inhibition did not occur during the GAPDH PCR step, as the PCR bands were still observed when the compounds were added only after the cDNA synthesis step (Figure 4B). These findings highlighted that compound 1, but not compound 2, inhibited the RT p66 subunits.

**Figure 4.**
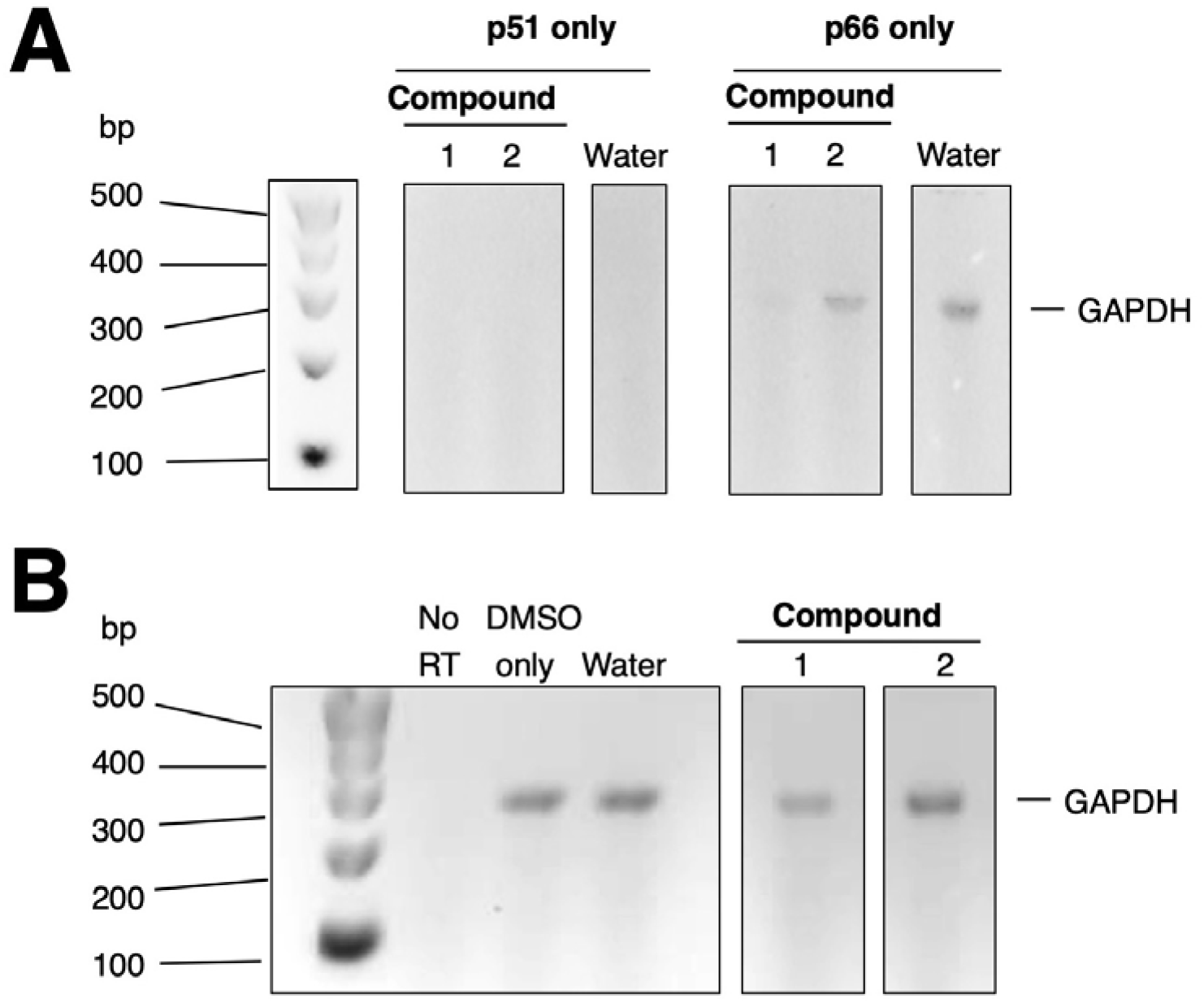
Agarose gel electrophoresis results of RT-PCR on GAPDH gene products on a 2% agar gel. Water and DMSO incubated reaction mixture were used as negative controls. (A) RT-PCR results of compounds on HIV-1 RT p51 or p66 subunits separately. The experiments were performed at 300 μM of all the compounds. (B) Results of post-cDNA synthesis analysis in the cases of the two compound and the control. All agarose gel electrophoresis results were analysed with GelApp [20] and bands were found to be within the expected size of ~300bp.

## DISCUSSION

We set out to determine if p51 could be a potential druggable target for novel RTIs and the required binding mechanisms. We did this by experimentally screening 40 compounds of the NCI Diversity Set V, and serendipitously finding two compounds that could inhibited the HIV-1 RT in RT-PCR. Given that RT is unique to viruses and absent in mammalian cells, the potential inhibitors are unlikely to cause side effects (unpublished limited mice experiments showed no toxicity effects at dosages up to 7 mg/kg for both compounds).

Our initial blind docking results identified two potential binding sites (on p66 and p51 subunits) for compound 1 (Figure 2). Structural analyses demonstrated that compound 1 bound energetically favourably to the binding site on p66, which was also supported by our experimental functional assay when compound 1 inhibited the activity of p66 homodimers (Figure 4A). Compound 1 was found to bind to the p66 site predominantly via hydrophobic interactions. On the other hand, compound 2 bound to the p51 subunit, driven predominantly by electrostatic interactions with its more polar pocket (Supplementary Table S2).

Computational analyses showed that both compounds interact with the RT p51 subunit (with less pronounced propensity for compound 1 at its second binding site in p51), distant from the polymerase site located on p66 (Figure 2A), suggesting an underlying allosteric mechanism. Interestingly, compound 2 established stable hydrogen bonds with residue Y183 of p51 (Figure 2B). To date, there are no known clinical mutations reported to occur at this residue of p51 [39] nor in selection-free *in vitro* mutagenesis studies [40], suggesting conservation of this residue and the possibility for compound 2 to be used against resistant HIV-1 variants.

It should be noted that compound 2 also interacted weakly (e.g. via conformations 5 and 6, as shown in Supplementary Figure S2) with another site on the p51 subunit in a similar binding mode (data not shown) to that of the other dominant site; nonetheless, this affirmed that compound 2 would target p51 rather than p66.

We found the first binding site on p51 for compound 2 overlaps with our previously predicted allosteric pocket [21]. Since the structural fold of p51 differs from that of p66 (e.g. overall RMSD ~4.9 Å, excluding the RNase H region), there is support that the main inhibition site of compound 2 may be located on p51. Previous studies [41, 42] showed that p66 or p51 homodimers of HIV-1 RT retain RT activity, albeit at a reduced level but we were unable to detect p51 homodimer activity. Nonetheless, given the lack of inhibition on p66 homodimers, there is additional support for the main modus operandi of inhibition to be on p51 that is in agreement with our computational analyses (Figure 4A). Since the DNA polymerase active site is on the p66 subunit, compound 2 is a potential novel type of NNRTI acting via p51, thereby requiring high concentrations of 300 μM compared to the direct action of compound 1 on p66 at 150 μM, albeit also a relatively high concentration.

With the high concentrations required, the two compounds are not likely to be effective therapeutic inhibitors in themselves. However, they provide a proof-of-concept of inhibiting RT via p51 and serve as possible scaffolds with a structural rationale for the key interactions required to guide the modification and development of new inhibitors. Most importantly, compound 2 confirmed the possibility of targeting the p51 subunit at a novel site that we had previously identified computationally [21].

In addition, our structural conservation analysis revealed that the compound 2 binding region on HIV-1 p51 is conserved across multiple RTs (Figure 5 and Supplementary Data 1), highlighting its significance. Since viruses can develop cross-resistance to multiple drugs [21, 43], and Y183 has yet to have a reported clinical mutation, compound 2 serves as a potentially important guide for targeting drug-resistant HIV-1 variants, and with the conservation of the site across RTs – a possible broad-spectrum RTI as well. By taking a holistic approach [44] in studying the whole structure of the target proteins and looking for common sites and how resistance develops, as demonstrated for HIV-1 Gag [45], HIV-1 Protease [43], and antibodies [46–51], the search for common druggable conserved sites across protein families can add to the success of developing broad-spectrum therapeutics.

**Figure 5.**
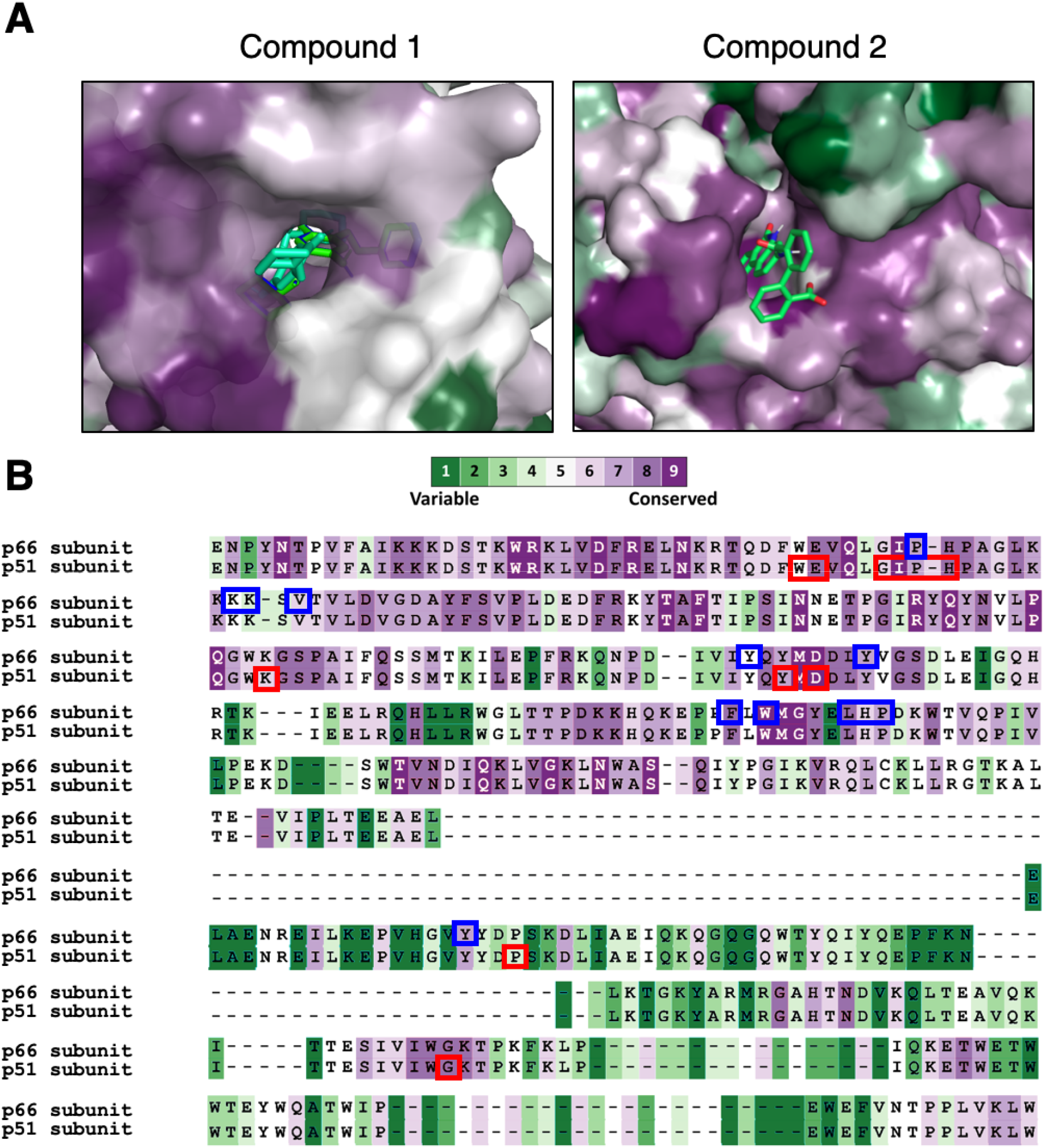
The two binding sites on HIV-1 RT that are highly conserved. (A) The binding sites on p66 and p51 complexed with compound 1 and 2, respectively, are shown in surface representation (coloured according to the level of conservation among the RT families). The two potential inhibitory compounds are shown as sticks. (B) Sequence alignment of the binding sites on p66 and p51, for compound 1 (blue boxes) and compound 2 (red boxes). The conservation colour scheme is the same as in (A). An augmented reality presentation of the binding pockets may be viewed using our “APD AR Holistic Review” App available on both Google and Apple app stores (for more details, see Poh et al. [52, 53]).

In conclusion, we have demonstrated the feasibility of inhibiting HIV RT via the p51 subunit on a previously identified druggable pock and serendipitously found two and characterized possible RTI scaffolds and the necessary structural engagements. Despite inhibiting only at high concentrations, the scaffolds are a promising starting point for further development of new HIV-1 RT lead candidates and even potential broad-spectrum RTIs.

## ACCESSION CODES

PDB code for HIV-1 Reverse Transcriptase in Figure 5 is 3T19.

## AUTHOR CONTRIBUTIONS

K.F.C. and S.X.P. performed the RT-PCR experiments. A.K. performed the molecular simulations. K.F.C., C.T.T.S., A.K., P.J.B, and S.K.E.G. analysed the results and wrote the manuscript. C.T.T.S. and S.K.E.G. conceived and supervised the study. All authors read and approved the final version of the manuscript.

## SUPPLEMENTARY MATERIALS

Supplementary Information and Data are provided online.

## ACKNOWLEDGEMENTS

This research was funded by A*STAR core fund. The forty compounds tested were provided by National Cancer Institute (NCI) Developmental Therapeutic Program’s Open Compound Repository, NIH. We thank Ling Wei Li, Lua Wai Heng, Poh Jun Jie and Yeo Joshua for their help in performing independent replicates of the RT-PCR experiments.

