## Supplementary for "An alternative HIV-1 non-nucleoside Reverse Transcriptase inhibition mechanism: Targeting the p51 subunit"

**SUPPLEMENTARY INFORMATION**

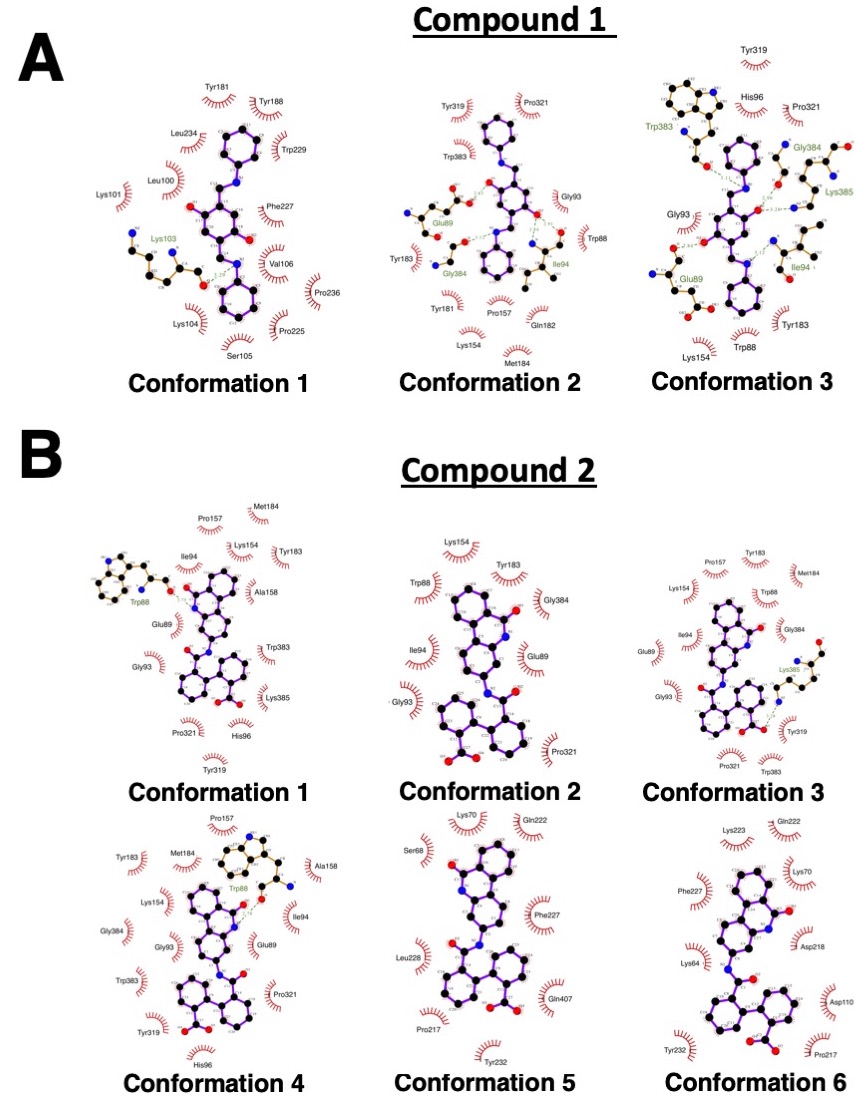

**Supplementary Figure S1**. Initial ligand bound conformations used for different setups of the MD simulations.

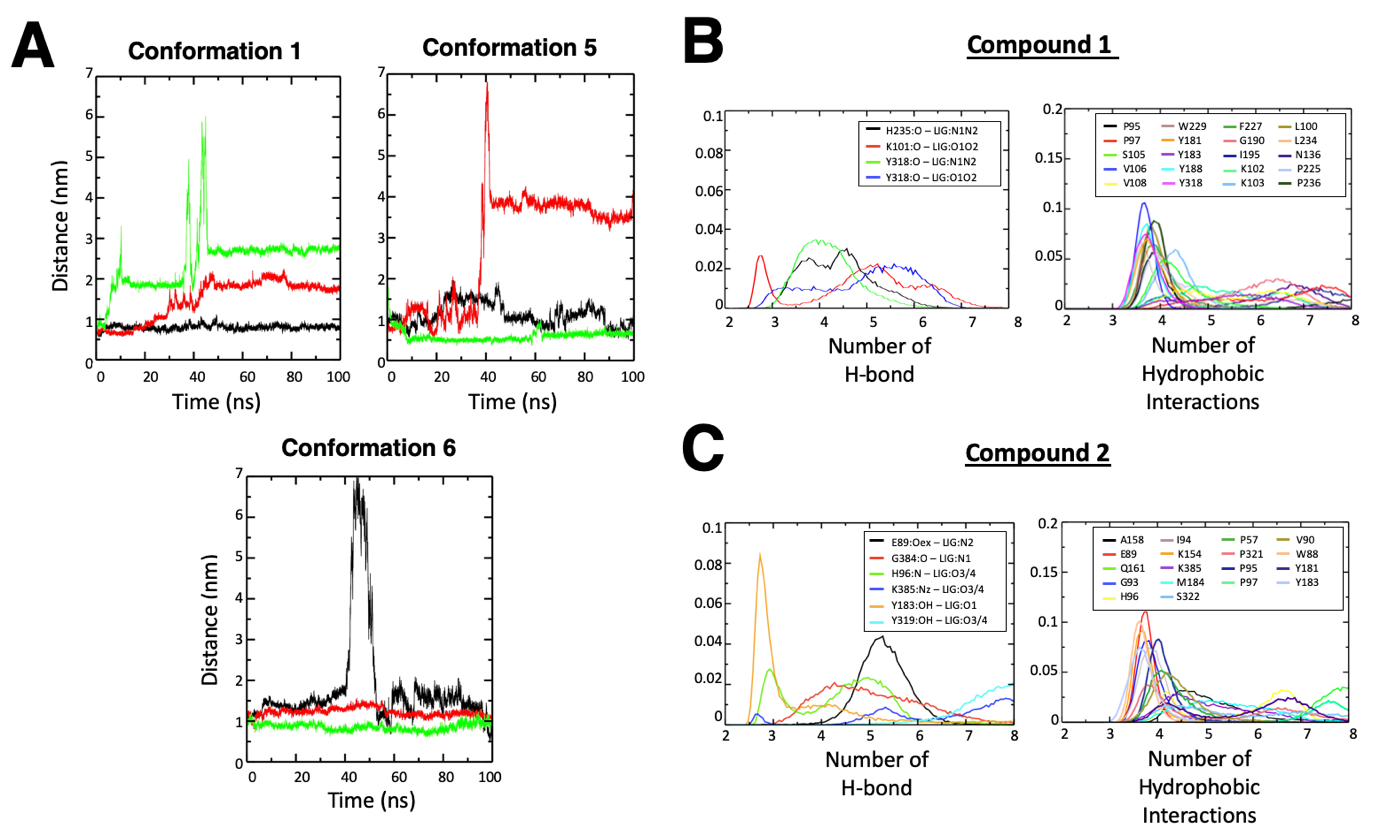

**Supplementary Figure S2.** Molecular simulation analyses of the binding sites for compounds 1 and 2. (A) The center-of-mass distances between the binding sites and compound 2 for three initial conformations that exhibited unstable ligand bound positions, calculated using independent triplicates (black, green and red) of 100 ns trajectories. Density plots of interactions with their respective binding sites are shown for (B) compound 1 and (C) compound 2.

**Supplementary Table S1**. Decomposition of binding energies for compound 1 in the binding sites on RT p66 and p51 subunits. MM-PBSA calculations were carried out with the program *g_mmpbsa*^1^ for the last 50 ns of each trajectory, using various internal dielectric constants ε_in_ for the solute^2-4^. Analyses were performed across independent triplicate simulations, using mean and standard deviation.

| Compound 1 | ε_int_ = 2 | | |  | ε_int_ = 8 | | |  | ε_int_ = 20 | | |
| --- | --- | --- | --- | --- | --- | --- | --- | --- | --- | --- | --- |
| (kcal/mol) | Conform. 1 (p66) | Conform. 2 (p51)^#^ | Conform. 3 (p51)^#^ |  | Conform. 1 (p66) | Conform. 2 (p51)^#^ | Conform. 3 (p51)^#^ |  | Conform. 1 (p66) | Conform. 2 (p51)^#^ | Conform. 3 (p51)^#^ |
| Total binding energy | -31.5 ± 3.3 | -16.4 ± 2.3 | -14.2 ± 1.7 |  | -39.3 ± 3.9 | -21.7 ± 2.2 | -19.7 ± 2.5 |  | -43.5 ± 4.1 | -27.8 ± 2.0 | -26.0 ± 2.3 |
| van der Waal | -44.2 ± 3.6 | -33.8 ± 1.9 | -33.1 ± 1.3 |  | -44.2 ± 3.6 | -33.8 ± 1.9 | -33.1 ± 1.3 |  | -44.2 ± 3.6 | -33.8 ± 1.9 | -33.1 ± 1.3 |
| Electrostatics | -3.1 ± 0.9 | -3.8 ± 1.0 | -5.6 ± 1.3 |  | -0.8 ± 0.2 | -1.0 ± 0.2 | -1.4 ± 0.3 |  | -0.3 ± 0.1 | -0.4 ± 0.1 | -0.6 ± 0.1 |
| Polar solvation | 20.7 ± 2.0 | 25.2 ± 3.2 | 28.6 ± 1.9 |  | 10.6 ± 0.8 | 17.0 ± 1.8 | 18.9 ± 1.2 |  | 5.9 ± 0.4 | 10.4 ± 1.0 | 11.7 ± 0.9 |
| Non-polar solvation | -4.9 ± 0.4 | -4.0 ± 0.2 | -4.1 ± 0.2 |  | -4.9 ± 0.4 | -4.0 ± 0.2 | -4.1 ± 0.2 |  | -4.9 ± 0.4 | -4.0 ± 0.2 | -4.1 ± 0.2 |

\# In both conformations 2 and 3, compound 1 was found to bind to the same site on the RT p51 subunit.

**Supplementary Table S2**. Additional binding analyses for the two compounds in their binding sites on the RT p66 and p51 subunits. Hydrophobic contacts between protein and ligand were calculated using a 4.0 Å cut-off. Hydrogen bonds between protein and ligand were calculated using a cut-off of 3.5 Å and maximum angle between donor – hydrogen atom and acceptor of 30°. The number of water molecules around the ligand was determined using a cut-off of 3.5 Å. Analyses were performed across independent triplicate simulations, using mean and standard deviation.

| Compound 1 | | | |  | Compound 2 | | | |
| --- | --- | --- | --- | --- | --- | --- | --- | --- |
|  | Number of water molecules | Number of hydrogen bonds | Number of hydrophobic contacts |  |  | Number of water molecules | Number of hydrogen bonds | Number of hydrophobic contacts |
| Conform. 1 (p66) | 2.8 ± 1.5 | 0.6 ±0.6 | 20.4 ± 4.8 |  | Conform. 1 (p51) | 15.2 ± 3.5 | 1.8 ± 1.3 | 13.6 ± 5.1 |
| Conform. 2 (p51) ^a^ | 8.2 ± 2.2 | 1.3 ± 0.8 | 12.2 ± 3.2 |  | Conform. 2 (p51) | 13.3 ± 2.6 | 3.0 ± 1.0 | 16.3 ± 3.6 |
| Conform. 3 (p51) ^a^ | 7.5 ± 2.2 | 1.3 ± 0.8 | 12.2 ± 3.5 |  | Conform. 3 (p51) | 15.4 ± 2.7 | 1.5 ± 1.1 | 15.0 ± 4.0 |
|  |  |  |  |  | Conform. 4 (p51) | 14.1 ± 2.5 | 2.1 ± 1.3 | 15.7 ± 3.6 |
|  |  |  |  |  | Conform. 5 (p51) ^b^ | 15.9 ± 3.6 | 1.7 ± 1.1 | 12.3 ± 5.6 |
|  |  |  |  |  | Conform. 6 (p51) ^b^ | 15.2 ± 3.9 | 2.2 ± 1.2 | 12.8 ± 5.7 |

^a^ In both conformations 2 and 3, compound 1 was found to bind to the same site on the RT p51 subunit.

^b^ Secondary binding site (weaker and unsteady) of compound 2 on the p51 subunit.

**Supplementary Table S3**. Decomposition of binding energies for compound 2 in the binding sites on RT p51 subunits. MM-PBSA calculations were carried out with the program *g_mmpbsa*^1^ for the last 50 ns of each trajectory, using various internal dielectric constants ε_in_ for the solute. Analyses were performed across independent triplicate simulations. Only results for the three setups that exhibited stable ligand binding (i.e. conformation 2, 3, and 4) are shown. Analyses were performed across independent triplicate simulations, using mean and standard deviation.

| Compound 2 | ε_int_ = 2 | | |  | ε_int_ = 8 | | |  | ε_int_ = 20 | | |
| --- | --- | --- | --- | --- | --- | --- | --- | --- | --- | --- | --- |
| (kcal/mol) | Conform. 2 (p51) | Conform. 3 (p51) | Conform. 4 (p51) |  | Conform. 2 (p51) | Conform. 3 (p51) | Conform. 4 (p51) |  | Conform. 2 (p51) | Conform. 3 (p51) | Conform. 4 (p51) |
| Total binding energy | -43.7 ± 2.0 | -49.9 ± 3.0 | -45.5 ± 2.2 |  | -13.4 ± 3.6 | -20.1 ± 2.6 | -12.2 ± 5.9 |  | -16.5 ± 2.2 | -17.2 ± 6.3 | -15.1 ± 4.4 |
| van der Waal | -38.7 ± 0.9 | -38.7 ± 0.3 | -38.2 ± 1.1 |  | -38.7 ± 0.9 | -38.7 ± 0.3 | -38.2 ± 1.1 |  | -38.7 ± 0.9 | -38.7 ± 0.3 | -38.2 ± 1.1 |
| Electrostatics | -58.4 ± 8.1 | -57.7 ± 2.8 | -70.3 ± 13.1 |  | -14.0 ± 2.0 | -14.4 ± 0.7 | -17.6 ± 3.3 |  | -5.8 ± 0.8 | -5.8 ± 0.3 | -7.0 ± 1.1 |
| Polar solvation | 57.9 ± 9.9 | 50.9 ± 5.3 | 67.4 ± 13.5 |  | 43.9 ± 5.1 | 37.1 ± 3.6 | 48.0 ± 8.0 |  | 32.5 ± 2.8 | 28.0 ± 2.4 | 34.5 ± 4.6 |
| Non-polar solvation | -4.5 ± 0.1 | -4.1 ± 0.2 | -4.3 ± 0.1 |  | -4.5 ± 0.1 | -4.1 ± 0.2 | -4.3 ± 0.1 |  | -4.5 ± 0.1 | -4.1 ± 0.2 | -4.3 ± 0.1 |
