## Supplementary Data for "An alternative HIV-1 non-nucleoside Reverse Transcriptase inhibition mechanism: Targeting the p51 subunit"

### ConSurf Color-Coded MSA

```
001 YP_004442836.1    --------------------------------------------------
002 NP_056728.1       --------------------------------------------------
003 YP_009508537.1    --------------------------------------------------
004 YP_004732983.2    --------------------------------------------------
005 YP_009508407.1    --------------------------------------------------
006 NP_040939.1       --------------------------------------------------
007 YP_003864102.1    --------------------------------------------------
008 YP_009002585.1    --------------------------------------------------
009 YP_003284237.1    --------------------------------------------------
010 YP_009268869.1    --------------------------------------------------
011 YP_009506251.1    --------------------------------------------------
012 NP_955611.1       --------------------------------------------------
013 YP_008567619.1    --------------------------------------------------
014 NP_149413.1       --------------------------------------------------
015 YP_004442824.1    --------------------------------------------------
016 YP_001931967.1    --------------------------------------------------
017 YP_002455786.1    --------------------------------------------------
018 YP_009109694.1    --------------------------------------------------
019 NP_789739.1       --------------------------------------------------
020 NP_955619.1       --------------------------------------------------
021 YP_224289.1       --------------------------------------------------
022 YP_009182100.1    --------------------------------------------------
023 NP_705927.1       --------------------------------------------------
024 YP_009508577.1    --------------------------------------------------
025 YP_004442830.1    --------------------------------------------------
026 NP_758887.1       --------------------------------------------------
027 YP_009508406.1    --------------------------------------------------
028 YP_610965.1       --------------------------------------------------
029 YP_001931961.1    --------------------------------------------------
030 YP_241114.1       --------------------------------------------------
031 YP_006732334.1    --------------------------------------------------
032 YP_009041481.1    --------------------------------------------------
033 YP_001036293.1    --------------------------------------------------
034 YP_605811.1       --------------------------------------------------
035 NP_955577.1       --------------------------------------------------
036 YP_003208050.1    --------------------------------------------------
037 NP_789740.1       --------------------------------------------------
038 YP_004442839.1    --------------------------------------------------
039 YP_009507791.1    --------------------------------------------------
040 YP_004581513.1    --------------------------------------------------
041 NP_043933.1       --------------------------------------------------
042 YP_009507248.1    --------------------------------------------------
043 YP_002117531.1    --------------------------------------------------
044 YP_004347415.1    --------------------------------------------------
045 NP_056891.1       -------------------------------------------------M
046 YP_009513242.1    --------------------------------------------------
047 YP_009352866.1    --------------------------------------------------
048 YP_009165750.1    --------------------------------------------------
049 NP_040840.1       --------------------------------------------------
050 YP_004222728.1    --------------------------------------------------
051 YP_009506264.1    --------------------------------------------------
052 YP_567050.1       --------------------------------------------------
053 YP_009508571.1    --------------------------------------------------
054 NP_056803.1       --------------------------------------------------
055 NP_955564.1       --------------------------------------------------
056 YP_009508411.1    --------------------------------------------------
057 YP_009140788.1    --------------------------------------------------
058 YP_009508410.1    --------------------------------------------------
059 NP_040333.1       --------------------------------------------------
060 YP_008992013.1    --------------------------------------------------
061 NP_041734.1       --------------------------------------------------
062 NP_056895.1       --------------------------------------------------
063 YP_009109689.1    --------------------------------------------------
064 YP_002519387.1    --------------------------------------------------
065 YP_595725.1       --------------------------------------------------
066 YP_009508551.1    --------------------------------------------------
067 YP_001497148.1    --------------------------------------------------
068 NP_041186.1       --------------------------------------------------
069 NP_663784.1       --------------------------------------------------
```

```
070 YP_009508443.1    --------------------------------------------------
071 YP_003987465.1    --------------------------------------------------
072 NP_861410.1       --------------------------------------------------
073 YP_009345071.1    --------------------------------------------------
074 YP_009508582.1    --------------------------------------------------
075 NP_569141.1       --------------------------------------------------
076 YP_002916057.1    --------------------------------------------------
077 NP_777384.2       --------------------------------------------------
078 YP_233110.1       --------------------------------------------------
079 NP_542258.1       --------------------------------------------------
080 YP_009130664.1    --------------------------------------------------
081 YP_009508566.1    --------------------------------------------------
082 YP_006607892.1    --------------------------------------------------
083 YP_009506270.1    --------------------------------------------------
084 NP_042513.1       --------------------------------------------------
085 YP_006907834.1    --------------------------------------------------
086 YP_004442827.1    --------------------------------------------------
087 YP_009408594.1    --------------------------------------------------
088 YP_009508561.1    --------------------------------------------------
089 NP_056907.1       --------------------------------------------------
090 YP_002308474.1    --------------------------------------------------
091 YP_007761644.1    --------------------------------------------------
092 NP_040550.1       --------------------------------------------------
093 NP_758808.1       --------------------------------------------------
094 YP_009345075.1    --------------------------------------------------
095 NP_068729.1       --------------------------------------------------
096 NP_047255.1       MSGASSGTAIGAHLFGVSPEYRVLIGDEGAGPSKSLSEVSFSVWYRSRAA
097 YP_006273075.1    --------------------------------------------------
098 YP_009508888.1    --------------------------------------------------
099 NP_395469.1       --------------------------------------------------
100 YP_009508546.1    --------------------------------------------------
101 NP_056762.1       --------------------------------------------------
102 YP_009508556.1    --------------------------------------------------
103 NP_569153.1       --------------------------------------------------
104 NP_043924.1       --------------------------------------------------
105 YP_009243641.1    --------------------------------------------------
106 YP_004300274.1    --------------------------------------------------
107 NP_056848.1       --------------------------------------------------
108 YP_006495799.1    --------------------------------------------------
109 YP_009508409.1    --------------------------------------------------
110 NP_687035.1       --------------------------------------------------
111 NP_045937.2       --------------------------------------------------
112 NP_612577.1       --------------------------------------------------
113 NP_619548.1       --------------------------------------------------
114 NP_056880.1       ------------------------------------------------MG
115 YP_009121747.1    --------------------------------------------------
116 YP_009113237.2    --------------------------------------------------
117 YP_009508408.1    --------------------------------------------------
118 NP_955579.1       --------------------------------------------------
119 YP_009229919.1    --------------------------------------------------
120 YP_004442833.1    --------------------------------------------------
121 NP_862833.2       --------------------------------------------------
122 NP_040563.1       --------------------------------------------------
123 YP_443922.1       -----MASSQPLKMKFSEYLDQRMQDRAGGKVQWKDHKKVWERLKLRWTQ
124 NP_057933.2       --------------------------------------------------
125 YP_009116631.1    --------------------------------------------------
126 YP_009513249.1    --------------------------------------------------
127 YP_223871.1       --------------------------------------------------
128 NP_127504.1       --------------------------------------------------
129 NP_044929.1       --------------------------------------------------
130 NP_057860.1       --------------------------------------------------
131 NP_040973.1       --------------------------------------------------
132 NP_777317.1       --------------------------------------------------
133 NP_054716.1       --------------------------------------------------
134 YP_001856242.1    --------------------------------------------------
135 NP_954565.2       -------------------------------------------------M
136 NP_569150.1       --------------------------------------------------
137 NP_659397.1       --------------------------------------------------
138 YP_009513211.1    --------------------------------------------------
139 YP_233107.1       --------------------------------------------------
140 YP_009109692.1    --------------------------------------------------
141 NP_057849.4       --------------------------------------------------
142 NP_049560.1       --------------------------------------------------
```

```
143 YP_009506267.1   --------------------------------------------------
144 NP_039820.1      --------------------------------------------------
145 NP_056790.1      --------------------------------------------------
146 YP_002321513.1   --------------------------------------------------
147 NP_955591.1      --------------------------------------------------
148 YP_001956722.2   --------------------------------------------------
149 NP_056902.1      --------------------------------------------------
150 NP_056886.1      --------------------------------------------------
151 NP_041261.1      --------------------------------------------------


001 YP_004442836.1   --------------------------------------------------
002 NP_056728.1      --------------------------------------------------
003 YP_009508537.1   --------------------------------------------------
004 YP_004732983.2   --------------------------------------------------
005 YP_009508407.1   --------------------------------------------------
006 NP_040939.1      --------------------------------------------------
007 YP_003864102.1   --------------------------------------------------
008 YP_009002585.1   --------------------------------------------------
009 YP_003284237.1   --------------------------------------------------
010 YP_009268869.1   --------------------------------------------------
011 YP_009506251.1   --------------------------------------------------
012 NP_955611.1      --------------------------------------------------
013 YP_008567619.1   --------------------------------------------------
014 NP_149413.1      --------------------------------------------------
015 YP_004442824.1   --------------------------------------------------
016 YP_001931967.1   --------------------------------------------------
017 YP_002455786.1   --------------------------------------------------
018 YP_009109694.1   --------------------------------------------------
019 NP_789739.1      --------------------------------------------------
020 NP_955619.1      --------------------------------------------------
021 YP_224289.1      --------------------------------------------------
022 YP_009182100.1   --------------------------------------------------
023 NP_705927.1      --------------------------------------------------
024 YP_009508577.1   --------------------------------------------------
025 YP_004442830.1   --------------------------------------------------
026 NP_758887.1      --------------------------------------------------
027 YP_009508406.1   --------------------------------------------------
028 YP_610965.1      --------------------------------------------------
029 YP_001931961.1   --------------------------------------------------
030 YP_241114.1      --------------------------------------------------
031 YP_006732334.1   --------------------------------------------------
032 YP_009041481.1   --------------------------------------------------
033 YP_001036293.1   --------------------------------------------------
034 YP_605811.1      --------------------------------------------------
035 NP_955577.1      --------------------------------------------------
036 YP_003208050.1   --------------------------------------------------
037 NP_789740.1      --------------------------------------------------
038 YP_004442839.1   --------------------------------------------------
039 YP_009507791.1   --------------------------------------------------
040 YP_004581513.1   --------------------------------------------------
041 NP_043933.1      --------------------------------------------------
042 YP_009507248.1   --------------------------------------------------
043 YP_002117531.1   --------------------------------------------------
044 YP_004347415.1   --------------------------------------------------
045 NP_056891.1      GQELSQHERYVEQLKQALKTRGVKVKYADLLKFFDFVKDTCPWFPQEGTI
046 YP_009513242.1   --------------------------------------------------
047 YP_009352866.1   --------------------------------------------------
048 YP_009165750.1   --------------------------------------------------
049 NP_040840.1      --------------------------------------------------
050 YP_004222728.1   --------------------------------------------------
051 YP_009506264.1   --------------------------------------------------
052 YP_567050.1      --------------------------------------------------
053 YP_009508571.1   --------------------------------------------------
054 NP_056803.1      --------------------------------------------------
055 NP_955564.1      --------------------------------------------------
056 YP_009508411.1   --------------------------------------------------
057 YP_009140788.1   --------------------------------------------------
058 YP_009508410.1   --------------------------------------------------
059 NP_040333.1      --------------------------------------------------
060 YP_008992013.1   --------------------------------------------------
061 NP_041734.1      --------------------------------------------------
```

```
062 NP_056895.1       --------------------------------------------------
063 YP_009109689.1    ------------------------MGQTLTTPLSLTLTHFSDVRARAHNL
064 YP_002519387.1    --------------------------------------------------
065 YP_595725.1       --------------------------------------------------
066 YP_009508551.1    --------------------------------------------------
067 YP_001497148.1    ------------------------MGQTLTTPLTLTLTHFSDVQARAHNL
068 NP_041186.1       --------------------------------------------------
069 NP_663784.1       --------------------------------------------------
070 YP_009508443.1    --------------------------------------------------
071 YP_003987465.1    --------------------------------------------------
072 NP_861410.1       --------------------------------------------------
073 YP_009345071.1    --------------------------------------------------
074 YP_009508582.1    --------------------------------------------------
075 NP_569141.1       --------------------------------------------------
076 YP_002916057.1    --------------------------------------------------
077 NP_777384.2       --------------------------------------------------
078 YP_233110.1       --------------------------------------------------
079 NP_542258.1       --------------------------------------------------
080 YP_009130664.1    --------------------------------------------------
081 YP_009508566.1    --------------------------------------------------
082 YP_006607892.1    --------------------------------------------------
083 YP_009506270.1    --------------------------------------------------
084 NP_042513.1       --------------------------------------------------
085 YP_006907834.1    --------------------------------------------------
086 YP_004442827.1    --------------------------------------------------
087 YP_009408594.1    --------------------------------------------------
088 YP_009508561.1    --------------------------------------------------
089 NP_056907.1       --------------------------------------------------
090 YP_002308474.1    --------------------------------------------------
091 YP_007761644.1    --------------------------------------------------
092 NP_040550.1       --------------------------------------------------
093 NP_758808.1       --------------------------------------------------
094 YP_009345075.1    --------------------------------------------------
095 NP_068729.1       --------------------------------------------------
096 NP_047255.1       RLVIFCLVASFLVPCLTFLIAETVMGQTITTPLSLTLDHWSEVRARAHNQ
097 YP_006273075.1    --------------------------------------------------
098 YP_009508888.1    --------------------------------------------------
099 NP_395469.1       --------------------------------------------------
100 YP_009508546.1    --------------------------------------------------
101 NP_056762.1       --------------------------------------------------
102 YP_009508556.1    --------------------------------------------------
103 NP_569153.1       --------------------------------------------------
104 NP_043924.1       --------------------------------------------------
105 YP_009243641.1    --------------------------------------------------
106 YP_004300274.1    --------------------------------------------------
107 NP_056848.1       --------------------------------------------------
108 YP_006495799.1    --------------------------------------------------
109 YP_009508409.1    --------------------------------------------------
110 NP_687035.1       --------------------------------------------------
111 NP_045937.2       --------------------------------------------------
112 NP_612577.1       --------------------------------------------------
113 NP_619548.1       --------------------------------------------------
114 NP_056880.1       VSGSKGQKLFVSVLQRLLSERGLHVKESSAIEFYQFLIKVSPWFPEEGGL
115 YP_009121747.1    --------------------------------------------------
116 YP_009113237.2    --------------------------------------------------
117 YP_009508408.1    --------------------------------------------------
118 NP_955579.1       --------------------------------------------------
119 YP_009229919.1    --------------------------------------------------
120 YP_004442833.1    --------------------------------------------------
121 NP_862833.2       --------------------------------------------------
122 NP_040563.1       --------------------------------------------------
123 YP_443922.1       EGYLAGGAPTRVQLSTMEGELREGKQRAKDSEVDLNRQHVFRKEGTKQRE
124 NP_057933.2       ------------------------MGQTVTTPLSLTLGHWKDVERIAHNQ
125 YP_009116631.1    --------------------------------------------------
126 YP_009513249.1    --------------------------------------------------
127 YP_223871.1       --------------------------------------------------
128 NP_127504.1       --------------------------------------------------
129 NP_044929.1       --------------------------------------------------
130 NP_057860.1       --------------------------------------------------
131 NP_040973.1       --------------------------------------------------
132 NP_777317.1       --------------------------------------------------
133 NP_054716.1       --------------------------------------------------
134 YP_001856242.1    --------------------------------------------------
```

```
135 NP_954565.2         G Q E L S Q H E R Y V E Q L K Q A L K T R G V K V K Y A D L L K F F D F V K D T C P W F P Q E G T I
136 NP_569150.1         - - - - - - - - - - - - - - - - - - - - - - - - - - - - - - - - - - - - - - - - - - - - - - - - - -
137 NP_659397.1         - - - - - - - - - - - - - - - - - - - - - - - - - - - - - - - - - - - - - - - - - - - - - - - - - -
138 YP_009513211.1      - - - - - - - - - - - - - - - - - - - - - - - - - - - - - - - - - - - - - - - - - - - - - - - - - -
139 YP_233107.1         - - - - - - - - - - - - - - - - - - - - - - - - - - - - - - - - - - - - - - - - - - - - - - - - - -
140 YP_009109692.1      - - - - - - - - - - - - - - - - - - - - - - - - - - - - - - - - - - - - - - - - - - - - - - - - - -
141 NP_057849.4         - - - - - - - - - - - - - - - - - - - - - - - - - - - - - - - - - - - - - - - - - - - - - - - - - -
142 NP_049560.1         - - - - - - - - - - - - - - - - - - - - - - - - - - - - - - - - - - - - - - - - - - - - - - - - - -
143 YP_009506267.1      - - - - - - - - - - - - - - - - - - - - - - - - - - - - - - - - - - - - - - - - - - - - - - - - - -
144 NP_039820.1         - - - - - - - - - - - - - - - - - - - - - - - - - - - - - - - - - - - - - - - - - - - - - - - - - -
145 NP_056790.1         - - - - - - - - - - - - - - - - - - - - - - - - - - - - - - - - - - - - - - - - - - - - - - - - - -
146 YP_002321513.1      - - - - - - - - - - - - - - - - - - - - - - - - - - - - - - - - - - - - - - - - - - - - - - - - - -
147 NP_955591.1         - - - - - - - - - - - - - - - - - - - - - - - - - - - - - - - - - - - - - - - - - - - - - - - - - -
148 YP_001956722.2      - - - - - - - - - - - - - - - - - - - - - - - - - - - - - - - - - - - - - - - - - - - - - - - - - -
149 NP_056902.1         - - - - - - - - - - - - - - - - - - - - - - - - - - - - - - - - - - - - - - - - - - - - - - - - - -
150 NP_056886.1         - - - - - - - - - - - - - - - - - - - - - - - - - - - - - - - - - - - - - - - - - - - - - - - - - -
151 NP_041261.1         - - - - - - - - - - - - - - - - - - - - - - - - - - - - - - - - - - - - - - - - - - - - - - - - - -
```

```
001 YP_004442836.1      - - - - - - - - - - - - - - - - - - - - - - - - - - - - - - - - - - - - - - - - - - - - - - - - - -
002 NP_056728.1         - - - - - - - - - - - - - - - - - - - - - - - - - - - - - - - - - - - - - - - - - - - - - - - - - -
003 YP_009508537.1      - - - - - - - - - - - - - - - - - - - - - - - - - - - - - - - - - - - - - - - - - - - - - - - - - -
004 YP_004732983.2      - - - - - - - - - - - - - - - - - - - - - - - - - - - - - - - - - - - - - - - - - - - - - - - - - -
005 YP_009508407.1      - - - - - - - - - - - - - - - - - - - - - - - - - - - - - - - - - - - - - - - - - - - - - - - - - -
006 NP_040939.1         - - - - - - - - - - - - - - - - - - - - - - - - - - - - - - - - - - - - - - - - - - - - - - - - - -
007 YP_003864102.1      - - - - - - - - - - - - - - - - - - - - - - - - - - - - - - - - - - - - - - - - - - - - - - - - - -
008 YP_009002585.1      - - - - - - - - - - - - - - - - - - - - - - - - - - - - - - - - - - - - - - - - - - - - - - - - - -
009 YP_003284237.1      - - - - - - - - - - - - - - - - - - - - - - - - - - - - - - - - - - - - - - - - - - - - - - - - - -
010 YP_009268869.1      - - - - - - - - - - - - - - - - - - - - - - - - - - - - - - - - - - - - - - - - - - - - - - - - - -
011 YP_009506251.1      - - - - - - - - - - - - - - - - - - - - - - - - - - - - - - - - - - - - - - - - - - - - - - - - - -
012 NP_955611.1         - - - - - - - - - - - - - - - - - - - - - - - - - - - - - - - - - - - - - - - - - - - - - - - - - -
013 YP_008567619.1      - - - - - - - - - - - - - - - - - - - - - - - - - - - - - - - - - - - - - - - - - - - - - - - - - -
014 NP_149413.1         - - - - - - - - - - - - - - - - - - - - - - - - - - - - - - - - - - - - - - - - - - - - - - - - - -
015 YP_004442824.1      - - - - - - - - - - - - - - - - - - - - - - - - - - - - - - - - - - - - - - - - - - - - - - - - - -
016 YP_001931967.1      - - - - - - - - - - - - - - - - - - - - - - - - - - - - - - - - - - - - - - - - - - - - - - - - - -
017 YP_002455786.1      - - - - - - - - - - - - - - - - - - - - - - - - - - - - - - - - - - - - - - - - - - - - - - - - - -
018 YP_009109694.1      - - - - - - - - - - - - - - - - - - - - - - - - - - - - - - - - - - - - - - - - - - - - - - - - - -
019 NP_789739.1         - - - - - - - - - - - - - - - - - - - - - - - - - - - - - - - - - - - - - - - - - - - - - - - - - -
020 NP_955619.1         - - - - - - - - - - - - - - - - - - - - - - - - - - - - - - - - - - - - - - - - - - - - - - - - - -
021 YP_224289.1         - - - - - - - - - - - - - - - - - - - - - - - - - - - - - - - - - - - - - - - - - - - - - - - - - -
022 YP_009182100.1      - - - - - - - - - - - - - - - - - - - - - - - - - - - - - - - - - - - - - - - - - - - - - - - - - -
023 NP_705927.1         - - - - - - - - - - - - - - - - - - - - - - - - - - - - - - - - - - - - - - - - - - - - - - - - - -
024 YP_009508577.1      - - - - - - - - - - - - - - - - - - - - - - - - - - - - - - - - - - - - - - - - - - - - - - - - - -
025 YP_004442830.1      - - - - - - - - - - - - - - - - - - - - - - - - - - - - - - - - - - - - - - - - - - - - - - - - - -
026 NP_758887.1         - - - - - - - - - - - - - - - - - - - - - - - - - - - - - - - - - - - - - - - - - - - - - - - - - -
027 YP_009508406.1      - - - - - - - - - - - - - - - - - - - - - - - - - - - - - - - - - - - - - - - - - - - - - - - - - -
028 YP_610965.1         - - - - - - - - - - - - - - - - - - - - - - - - - - - - - - - - - - - - - - - - - - - - - - - - - -
029 YP_001931961.1      - - - - - - - - - - - - - - - - - - - - - - - - - - - - - - - - - - - - - - - - - - - - - - - - - -
030 YP_241114.1         - - - - - - - - - - - - - - - - - - - - - - - - - - - - - - - - - - - - - - - - - - - - - - - - - -
031 YP_006732334.1      - - - - - - - - - - - - - - - - - - - - - - - - - - - - - - - - - - - - - - - - - - - - - - - - - -
032 YP_009041481.1      - - - - - - - - - - - - - - - - - - - - - - - - - - - - - - - - - - - - - - - - - - - - - - - - - -
033 YP_001036293.1      - - - - - - - - - - - - - - - - - - - - - - - - - - - - - - - - - - - - - - - - - - - - - - - - - -
034 YP_605811.1         - - - - - - - - - - - - - - - - - - - - - - - - - - - - - - - - - - - - - - - - - - - - - - - - - -
035 NP_955577.1         - - - - - - - - - - - - - - - - - - - - - - - - - - - - - - - - - - - - - - - - - - - - - - - - - -
036 YP_003208050.1      - - - - - - - - - - - - - - - - - - - - - - - - - - - - - - - - - - - - - - - - - - - - - - - - - -
037 NP_789740.1         - - - - - - - - - - - - - - - - - - - - - - - - - - - - - - - - - - - - - - - - - - - - - - - - - -
038 YP_004442839.1      - - - - - - - - - - - - - - - - - - - - - - - - - - - - - - - - - - - - - - - - - - - - - - - - - -
039 YP_009507791.1      - - - - - - - - - - - - - - - - - - - - - - - - - - - - - - - - - - - - - - - - - - - - - - - - - -
040 YP_004581513.1      - - - - - - - - - - - - - - - - - - - - - - - - - - - - - - - - - - - - - - - - - - - - - - - - - -
041 NP_043933.1         - - - - - - - - - - - - - - - - - - - - - - - - - - - - - - - - - - - - - - - - - - - - - - - - - -
042 YP_009507248.1      - - - - - - - - - - - - - - - - - - - - - - - - - - - - - - - - - - - - - - - - - - - - - - - - - -
043 YP_002117531.1      - - - - - - - - - - - - - - - - - - - - - - - - - - - - - - - - - - - - - - - - - - - - - - - - - -
044 YP_004347415.1      - - - - - - - - - - - - - - - - - - - - - - - - - - - - - - - - - - - - - - - - - - - - - - - - - -
045 NP_056891.1         D I K R W R R V G D C F Q D Y Y N T F G P E K V P V T A F S Y W N L I K E L I D K K E V N P Q V M A
046 YP_009513242.1      - - - - - - - - - - - - - - - - - - - - - - - - - - - - - - - - - - - - - - - - - - - - - - - - - -
047 YP_009352866.1      - - - - - - - - - - - - - - - - - - - - - - - - - - - - - - - - - - - - - - - - - - - - - - - - - -
048 YP_009165750.1      - - - - - - - - - - - - - - - - - - - - - - - - - - - - - - - - - - - - - - - - - - - - - - - - - -
049 NP_040840.1         - - - - - - - - - - - - - - - - - - - - - - - - - - - - - - - - - - - - - - - - - - - - - - - - - -
050 YP_004222728.1      - - - - - - - - - - - - - - - - - - - - - - - - - - - - - - - - - - - - - - - - - - - - - - - - - -
051 YP_009506264.1      - - - - - - - - - - - - - - - - - - - - - - - - - - - - - - - - - - - - - - - - - - - - - - - - - -
052 YP_567050.1         - - - - - - - - - - - - - - - - - - - - - - - - - - - - - - - - - - - - - - - - - - - - - - - - - -
053 YP_009508571.1      - - - - - - - - - - - - - - - - - - - - - - - - - - - - - - - - - - - - - - - - - - - - - - - - - -
```

```
054 NP_056803.1      --------------------------------------------------
055 NP_955564.1      --------------------------------------------------
056 YP_009508411.1   --------------------------------------------------
057 YP_009140788.1   --------------------------------------------------
058 YP_009508410.1   --------------------------------------------------
059 NP_040333.1      --------------------------------------------------
060 YP_008992013.1   --------------------------------------------------
061 NP_041734.1      --------------------------------------------------
062 NP_056895.1      --------------------------------------------------
063 YP_009109689.1   SVGVRKGRWQTFCSSEWPTLHVGWPRDGTFDLSVILQVKTKVMDPGPHGH
064 YP_002519387.1   --------------------------------------------------
065 YP_595725.1      --------------------------------------------------
066 YP_009508551.1   --------------------------------------------------
067 YP_001497148.1   SLEVRKGRWRTYCSSEWPTLSVGWPRDGTFDLSIILQVKTKVMDPGPRGH
068 NP_041186.1      --------------------------------------------------
069 NP_663784.1      --------------------------------------------------
070 YP_009508443.1   --------------------------------------------------
071 YP_003987465.1   --------------------------------------------------
072 NP_861410.1      --------------------------------------------------
073 YP_009345071.1   --------------------------------------------------
074 YP_009508582.1   --------------------------------------------------
075 NP_569141.1      --------------------------------------------------
076 YP_002916057.1   --------------------------------------------------
077 NP_777384.2      --------------------------------------------------
078 YP_233110.1      --------------------------------------------------
079 NP_542258.1      --------------------------------------------------
080 YP_009130664.1   --------------------------------------------------
081 YP_009508566.1   --------------------------------------------------
082 YP_006607892.1   --------------------------------------------------
083 YP_009506270.1   --------------------------------------------------
084 NP_042513.1      --------------------------------------------------
085 YP_006907834.1   --------------------------------------------------
086 YP_004442827.1   --------------------------------------------------
087 YP_009408594.1   --------------------------------------------------
088 YP_009508561.1   --------------------------------------------------
089 NP_056907.1      --------------------------------------------------
090 YP_002308474.1   --------------------------------------------------
091 YP_007761644.1   --------------------------------------------------
092 NP_040550.1      --------------------------------------------------
093 NP_758808.1      --------------------------------------------------
094 YP_009345075.1   --------------------------------------------------
095 NP_068729.1      --------------------------------------------------
096 NP_047255.1      GVEVRKKKWITLCEAEWVMMNVGWPREGTFSLDNISQVEKKIFAPGPYGH
097 YP_006273075.1   --------------------------------------------------
098 YP_009508888.1   --------------------------------------------------
099 NP_395469.1      --------------------------------------------------
100 YP_009508546.1   --------------------------------------------------
101 NP_056762.1      --------------------------------------------------
102 YP_009508556.1   --------------------------------------------------
103 NP_569153.1      --------------------------------------------------
104 NP_043924.1      --------------------------------------------------
105 YP_009243641.1   --------------------------------------------------
106 YP_004300274.1   --------------------------------------------------
107 NP_056848.1      --------------------------------------------------
108 YP_006495799.1   --------------------------------------------------
109 YP_009508409.1   --------------------------------------------------
110 NP_687035.1      --------------------------------------------------
111 NP_045937.2      --------------------------------------------------
112 NP_612577.1      --------------------------------------------------
113 NP_619548.1      --------------------------------------------------
114 NP_056880.1      NLQDWKRVGREMKRYAAEHGTDSIPKQAYPIWLQLREILTEQSDLVLLSA
115 YP_009121747.1   --------------------------------------------------
116 YP_009113237.2   --------------------------------------------------
117 YP_009508408.1   --------------------------------------------------
118 NP_955579.1      --------------------------------------------------
119 YP_009229919.1   --------------------------------------------------
120 YP_004442833.1   --------------------------------------------------
121 NP_862833.2      --------------------------------------------------
122 NP_040563.1      --------------------------------------------------
123 YP_443922.1      KAEEELRIGTWAIEETRRFIMLRTPAVMGIATDMKEGGVPTISPATPAEN
124 NP_057933.2      SVDVKKRRWVTFCSAEWPTFNVGWPRDGTFNRDLITQVKIKVFSPGPHGH
125 YP_009116631.1   --------------------------------------------------
126 YP_009513249.1   --------------------------------------------------
```

```
127 YP_223871.1        --------------------------------------------------
128 NP_127504.1        --------------------------------------------------
129 NP_044929.1        --------------------------------------------------
130 NP_057860.1        --------------------------------------------------
131 NP_040973.1        --------------------------------------------------
132 NP_777317.1        --------------------------------------------------
133 NP_054716.1        --------------------------------------------------
134 YP_001856242.1     --------------------------------------------------
135 NP_954565.2        DIKRWRRVGDCFQDYYNTFGPEKVPVTAFSYWNLIKELIDKKEVNPQVMA
136 NP_569150.1        --------------------------------------------------
137 NP_659397.1        --------------------------------------------------
138 YP_009513211.1     --------------------------------------------------
139 YP_233107.1        --------------------------------------------------
140 YP_009109692.1     --------------------------------------------------
141 NP_057849.4        --------------------------------------------------
142 NP_049560.1        --------------------------------------------------
143 YP_009506267.1     --------------------------------------------------
144 NP_039820.1        --------------------------------------------------
145 NP_056790.1        --------------------------------------------------
146 YP_002321513.1     --------------------------------------------------
147 NP_955591.1        --------------------------------------------------
148 YP_001956722.2     --------------------------------------------------
149 NP_056902.1        --------------------------------------------------
150 NP_056886.1        --------------------------------------------------
151 NP_041261.1        --------------------------------------------------


001 YP_004442836.1     --------------------------------------------------
002 NP_056728.1        --------------------------------------------------
003 YP_009508537.1     --------------------------------------------------
004 YP_004732983.2     --------------------------------------------------
005 YP_009508407.1     --------------------------------------------------
006 NP_040939.1        --------------------------------------------------
007 YP_003864102.1     --------------------------------------------------
008 YP_009002585.1     --------------------------------------------------
009 YP_003284237.1     --------------------------------------------------
010 YP_009268869.1     --------------------------------------------------
011 YP_009506251.1     --------------------------------------------------
012 NP_955611.1        --------------------------------------------------
013 YP_008567619.1     --------------------------------------------------
014 NP_149413.1        --------------------------------------------------
015 YP_004442824.1     --------------------------------------------------
016 YP_001931967.1     --------------------------------------------------
017 YP_002455786.1     --------------------------------------------------
018 YP_009109694.1     --------------------------------------------------
019 NP_789739.1        --------------------------------------------------
020 NP_955619.1        --------------------------------------------------
021 YP_224289.1        --------------------------------------------------
022 YP_009182100.1     --------------------------------------------------
023 NP_705927.1        --------------------------------------------------
024 YP_009508577.1     --------------------------------------------------
025 YP_004442830.1     --------------------------------------------------
026 NP_758887.1        --------------------------------------------------
027 YP_009508406.1     --------------------------------------------------
028 YP_610965.1        --------------------------------------------------
029 YP_001931961.1     --------------------------------------------------
030 YP_241114.1        --------------------------------------------------
031 YP_006732334.1     --------------------------------------------------
032 YP_009041481.1     --------------------------------------------------
033 YP_001036293.1     --------------------------------------------------
034 YP_605811.1        --------------------------------------------------
035 NP_955577.1        --------------------------------------------------
036 YP_003208050.1     --------------------------------------------------
037 NP_789740.1        --------------------------------------------------
038 YP_004442839.1     --------------------------------------------------
039 YP_009507791.1     --------------------------------------------------
040 YP_004581513.1     --------------------------------------------------
041 NP_043933.1        --------------------------------------------------
042 YP_009507248.1     --------------------------------------------------
043 YP_002117531.1     --------------------------------------------------
044 YP_004347415.1     --------------------------------------------------
045 NP_056891.1        AVAQTEEILKSNSQTDLTKTSQNPDLDLISLDSDDEGAKSSSLQDKGLSS
```

```
046 YP_009513242.1    --------------------------------------------------
047 YP_009352866.1    --------------------------------------------------
048 YP_009165750.1    --------------------------------------------------
049 NP_040840.1       --------------------------------------------------
050 YP_004222728.1    --------------------------------------------------
051 YP_009506264.1    --------------------------------------------------
052 YP_567050.1       --------------------------------------------------
053 YP_009508571.1    --------------------------------------------------
054 NP_056803.1       --------------------------------------------------
055 NP_955564.1       --------------------------------------------------
056 YP_009508411.1    --------------------------------------------------
057 YP_009140788.1    --------------------------------------------------
058 YP_009508410.1    --------------------------------------------------
059 NP_040333.1       --------------------------------------------------
060 YP_008992013.1    --------------------------------------------------
061 NP_041734.1       --------------------------------------------------
062 NP_056895.1       --------------------------------------------------
063 YP_009109689.1    PDQVAYIITWEDLVRNPPPWVKPFLHTPSTSKSTLLALEVPKNRT-----
064 YP_002519387.1    --------------------------------------------------
065 YP_595725.1       --------------------------------------------------
066 YP_009508551.1    --------------------------------------------------
067 YP_001497148.1    PDQIAYILTWEDLIRNPPAWVKPFLPSCPLSQSTLLPLKTSKDRASTQPS
068 NP_041186.1       --------------------------------------------------
069 NP_663784.1       --------------------------------------------------
070 YP_009508443.1    --------------------------------------------------
071 YP_003987465.1    --------------------------------------------------
072 NP_861410.1       --------------------------------------------------
073 YP_009345071.1    --------------------------------------------------
074 YP_009508582.1    --------------------------------------------------
075 NP_569141.1       --------------------------------------------------
076 YP_002916057.1    --------------------------------------------------
077 NP_777384.2       --------------------------------------------------
078 YP_233110.1       --------------------------------------------------
079 NP_542258.1       --------------------------------------------------
080 YP_009130664.1    --------------------------------------------------
081 YP_009508566.1    --------------------------------------------------
082 YP_006607892.1    --------------------------------------------------
083 YP_009506270.1    --------------------------------------------------
084 NP_042513.1       --------------------------------------------------
085 YP_006907834.1    --------------------------------------------------
086 YP_004442827.1    --------------------------------------------------
087 YP_009408594.1    --------------------------------------------------
088 YP_009508561.1    --------------------------------------------------
089 NP_056907.1       --------------------------------------------------
090 YP_002308474.1    --------------------------------------------------
091 YP_007761644.1    --------------------------------------------------
092 NP_040550.1       --------------------------------------------------
093 NP_758808.1       --------------------------------------------------
094 YP_009345075.1    --------------------------------------------------
095 NP_068729.1       --------------------------------------------------
096 NP_047255.1       PDQVPYITTWRSLATDPPSWVRPFLPPPKPPTP-----------------
097 YP_006273075.1    --------------------------------------------------
098 YP_009508888.1    --------------------------------------------------
099 NP_395469.1       --------------------------------------------------
100 YP_009508546.1    --------------------------------------------------
101 NP_056762.1       --------------------------------------------------
102 YP_009508556.1    --------------------------------------------------
103 NP_569153.1       --------------------------------------------------
104 NP_043924.1       --------------------------------------------------
105 YP_009243641.1    --------------------------------------------------
106 YP_004300274.1    --------------------------------------------------
107 NP_056848.1       --------------------------------------------------
108 YP_006495799.1    --------------------------------------------------
109 YP_009508409.1    --------------------------------------------------
110 NP_687035.1       --------------------------------------------------
111 NP_045937.2       --------------------------------------------------
112 NP_612577.1       --------------------------------------------------
113 NP_619548.1       --------------------------------------------------
114 NP_056880.1       EAKSVTEEELEEGLTGLLSTSSQEKT----YGTRGTAYAEIDTEVDKLSE
115 YP_009121747.1    --------------------------------------------------
116 YP_009113237.2    --------------------------------------------------
117 YP_009508408.1    --------------------------------------------------
118 NP_955579.1       --------------------------------------------------
```

```
119 YP_009229919.1    --------------------------------------------------
120 YP_004442833.1    --------------------------------------------------
121 NP_862833.2       --------------------------------------------------
122 NP_040563.1       --------------------------------------------------
123 YP_443922.1       QTPTQRLYPDLPEDSRPPPYSPSASPSPTVQAPVLTVRGGVIKGEVHLVE
124 NP_057933.2       PDQVPYIVTWEALAFDPPPWVKPFVHPKPPPPL-----------------
125 YP_009116631.1    --------------------------------------------------
126 YP_009513249.1    --------------------------------------------------
127 YP_223871.1       --------------------------------------------------
128 NP_127504.1       --------------------------------------------------
129 NP_044929.1       --------------------------------------------------
130 NP_057860.1       --------------------------------------------------
131 NP_040973.1       --------------------------------------------------
132 NP_777317.1       --------------------------------------------------
133 NP_054716.1       --------------------------------------------------
134 YP_001856242.1    --------------------------------------------------
135 NP_954565.2       AVAQTEEILKSNSQTDLTKTSQNPDLDLISLDSDDEGAKSSSLQDKGLSS
136 NP_569150.1       --------------------------------------------------
137 NP_659397.1       --------------------------------------------------
138 YP_009513211.1    --------------------------------------------------
139 YP_233107.1       --------------------------------------------------
140 YP_009109692.1    --------------------------------------------------
141 NP_057849.4       --------------------------------------------------
142 NP_049560.1       --------------------------------------------------
143 YP_009506267.1    --------------------------------------------------
144 NP_039820.1       --------------------------------------------------
145 NP_056790.1       --------------------------------------------------
146 YP_002321513.1    --------------------------------------------------
147 NP_955591.1       --------------------------------------------------
148 YP_001956722.2    --------------------------------------------------
149 NP_056902.1       --------------------------------------------------
150 NP_056886.1       --------------------------------------------------
151 NP_041261.1       --------------------------------------------------
```

```
001 YP_004442836.1    --------------------------------------------------
002 NP_056728.1       --------------------------------------------------
003 YP_009508537.1    --------------------------------------------------
004 YP_004732983.2    --------------------------------------------------
005 YP_009508407.1    --------------------------------------------------
006 NP_040939.1       --------------------------------------------------
007 YP_003864102.1    --------------------------------------------------
008 YP_009002585.1    --------------------------------------------------
009 YP_003284237.1    --------------------------------------------------
010 YP_009268869.1    --------------------------------------------------
011 YP_009506251.1    --------------------------------------------------
012 NP_955611.1       --------------------------------------------------
013 YP_008567619.1    --------------------------------------------------
014 NP_149413.1       --------------------------------------------------
015 YP_004442824.1    --------------------------------------------------
016 YP_001931967.1    --------------------------------------------------
017 YP_002455786.1    --------------------------------------------------
018 YP_009109694.1    --------------------------------------------------
019 NP_789739.1       --------------------------------------------------
020 NP_955619.1       --------------------------------------------------
021 YP_224289.1       --------------------------------------------------
022 YP_009182100.1    --------------------------------------------------
023 NP_705927.1       --------------------------------------------------
024 YP_009508577.1    --------------------------------------------------
025 YP_004442830.1    --------------------------------------------------
026 NP_758887.1       --------------------------------------------------
027 YP_009508406.1    --------------------------------------------------
028 YP_610965.1       --------------------------------------------------
029 YP_001931961.1    --------------------------------------------------
030 YP_241114.1       --------------------------------------------------
031 YP_006732334.1    --------------------------------------------------
032 YP_009041481.1    --------------------------------------------------
033 YP_001036293.1    --------------------------------------------------
034 YP_605811.1       --------------------------------------------------
035 NP_955577.1       --------------------------------------------------
036 YP_003208050.1    --------------------------------------------------
037 NP_789740.1       --------------------------------------------------
```

```
038 YP_004442839.1  --------------------------------------------------
039 YP_009507791.1  --------------------------------------------------
040 YP_004581513.1  --------------------------------------------------
041 NP_043933.1     --------------------------------------------------
042 YP_009507248.1  --------------------------------------------------
043 YP_002117531.1  --------------------------------------------------
044 YP_004347415.1  --------------------------------------------------
045 NP_056891.1     TKKPKRFPVLLTAQTSKDPEDPNPSEVDWDGLEDEAAKYHNPDWPPFLTR
046 YP_009513242.1  --------------------------------------------------
047 YP_009352866.1  --------------------------------------------------
048 YP_009165750.1  --------------------------------------------------
049 NP_040840.1     --------------------------------------------------
050 YP_004222728.1  --------------------------------------------------
051 YP_009506264.1  --------------------------------------------------
052 YP_567050.1     --------------------------------------------------
053 YP_009508571.1  --------------------------------------------------
054 NP_056803.1     --------------------------------------------------
055 NP_955564.1     --------------------------------------------------
056 YP_009508411.1  --------------------------------------------------
057 YP_009140788.1  --------------------------------------------------
058 YP_009508410.1  --------------------------------------------------
059 NP_040333.1     --------------------------------------------------
060 YP_008992013.1  --------------------------------------------------
061 NP_041734.1     --------------------------------------------------
062 NP_056895.1     --------------------------------------------------
063 YP_009109689.1  --LDPPKPVLPDESQQDLLFQDPLPHPPHNPLLEPPPYNSPSPPVLSPVS
064 YP_002519387.1  --------------------------------------------------
065 YP_595725.1     --------------------------------------------------
066 YP_009508551.1  --------------------------------------------------
067 YP_001497148.1  APPKPPKPVLPDESQKDPPLLDALSSPPHNPLLQPPPYNPPLAPALTPVG
068 NP_041186.1     --------------------------------------------------
069 NP_663784.1     --------------------------------------------------
070 YP_009508443.1  --------------------------------------------------
071 YP_003987465.1  --------------------------------------------------
072 NP_861410.1     --------------------------------------------------
073 YP_009345071.1  --------------------------------------------------
074 YP_009508582.1  --------------------------------------------------
075 NP_569141.1     --------------------------------------------------
076 YP_002916057.1  --------------------------------------------------
077 NP_777384.2     --------------------------------------------------
078 YP_233110.1     --------------------------------------------------
079 NP_542258.1     --------------------------------------------------
080 YP_009130664.1  --------------------------------------------------
081 YP_009508566.1  --------------------------------------------------
082 YP_006607892.1  --------------------------------------------------
083 YP_009506270.1  --------------------------------------------------
084 NP_042513.1     --------------------------------------------------
085 YP_006907834.1  --------------------------------------------------
086 YP_004442827.1  --------------------------------------------------
087 YP_009408594.1  --------------------------------------------------
088 YP_009508561.1  --------------------------------------------------
089 NP_056907.1     --------------------------------------------------
090 YP_002308474.1  --------------------------------------------------
091 YP_007761644.1  --------------------------------------------------
092 NP_040550.1     --------------------------------------------------
093 NP_758808.1     --------------------------------------------------
094 YP_009345075.1  --------------------------------------------------
095 NP_068729.1     --------------------------------------------------
096 NP_047255.1     ------LPQP-----LSPQPSAPLTSSLYPVLPKSDPPKPPVLPPDPSSP
097 YP_006273075.1  --------------------------------------------------
098 YP_009508888.1  --------------------------------------------------
099 NP_395469.1     --------------------------------------------------
100 YP_009508546.1  --------------------------------------------------
101 NP_056762.1     --------------------------------------------------
102 YP_009508556.1  --------------------------------------------------
103 NP_569153.1     --------------------------------------------------
104 NP_043924.1     --------------------------------------------------
105 YP_009243641.1  --------------------------------------------------
106 YP_004300274.1  --------------------------------------------------
107 NP_056848.1     --------------------------------------------------
108 YP_006495799.1  --------------------------------------------------
109 YP_009508409.1  --------------------------------------------------
110 NP_687035.1     --------------------------------------------------
```

```
111 NP_045937.2       --------------------------------------------------
112 NP_612577.1       --------------------------------------------------
113 NP_619548.1       --------------------------------------------------
114 NP_056880.1       HIYDEPYEEKEKADKNEEKDHVRKIKKVVQRKENSEGKRKEKDSKAFLAT
115 YP_009121747.1    --------------------------------------------------
116 YP_009113237.2    --------------------------------------------------
117 YP_009508408.1    --------------------------------------------------
118 NP_955579.1       --------------------------------------------------
119 YP_009229919.1    --------------------------------------------------
120 YP_004442833.1    --------------------------------------------------
121 NP_862833.2       --------------------------------------------------
122 NP_040563.1       --------------------------------------------------
123 YP_443922.1       GTIDVENSEPPQRSMDTDDLGPAGGRSTRGPDSLEEGSMRTQSSKRTRNP
124 NP_057933.2       ------PPSAPSLPLEPPRSTPPRSSLYPALTPSLGAKPKPQVLSDSGGP
125 YP_009116631.1    --------------------------------------------------
126 YP_009513249.1    --------------------------------------------------
127 YP_223871.1       --------------------------------------------------
128 NP_127504.1       --------------------------------------------------
129 NP_044929.1       --------------------------------------------------
130 NP_057860.1       --------------------------------------------------
131 NP_040973.1       --------------------------------------------------
132 NP_777317.1       --------------------------------------------------
133 NP_054716.1       --------------------------------------------------
134 YP_001856242.1    --------------------------------------------------
135 NP_954565.2       TKKPKRFPVLLTAQTSKDPEDPNPSEVDWDGLEDEAAKYHNPDWPPFLTR
136 NP_569150.1       --------------------------------------------------
137 NP_659397.1       --------------------------------------------------
138 YP_009513211.1    --------------------------------------------------
139 YP_233107.1       --------------------------------------------------
140 YP_009109692.1    --------------------------------------------------
141 NP_057849.4       --------------------------------------------------
142 NP_049560.1       --------------------------------------------------
143 YP_009506267.1    --------------------------------------------------
144 NP_039820.1       --------------------------------------------------
145 NP_056790.1       --------------------------------------------------
146 YP_002321513.1    --------------------------------------------------
147 NP_955591.1       --------------------------------------------------
148 YP_001956722.2    --------------------------------------------------
149 NP_056902.1       --------------------------------------------------
150 NP_056886.1       --------------------------------------MEAVIKVISSAC
151 NP_041261.1       --------------------------------------------------

001 YP_004442836.1    --------------------------------------------------
002 NP_056728.1       --------------------------------------------------
003 YP_009508537.1    --------------------------------------------------
004 YP_004732983.2    --------------------------------------------------
005 YP_009508407.1    --------------------------------------------------
006 NP_040939.1       --------------------------------------------------
007 YP_003864102.1    --------------------------------------------------
008 YP_009002585.1    --------------------------------------------------
009 YP_003284237.1    --------------------------------------------------
010 YP_009268869.1    --------------------------------------------------
011 YP_009506251.1    --------------------------------------------------
012 NP_955611.1       --------------------------------------------------
013 YP_008567619.1    --------------------------------------------------
014 NP_149413.1       --------------------------------------------------
015 YP_004442824.1    --------------------------------------------------
016 YP_001931967.1    --------------------------------------------------
017 YP_002455786.1    --------------------------------------------------
018 YP_009109694.1    --------------------------------------------------
019 NP_789739.1       --------------------------------------------------
020 NP_955619.1       --------------------------------------------------
021 YP_224289.1       --------------------------------------------------
022 YP_009182100.1    --------------------------------------------------
023 NP_705927.1       --------------------------------------------------
024 YP_009508577.1    --------------------------------------------------
025 YP_004442830.1    --------------------------------------------------
026 NP_758887.1       --------------------------------------------------
027 YP_009508406.1    --------------------------------------------------
028 YP_610965.1       --------------------------------------------------
029 YP_001931961.1    --------------------------------------------------
```

```
030 YP_241114.1      --------------------------------------------------
031 YP_006732334.1   --------------------------------------------------
032 YP_009041481.1   --------------------------------------------------
033 YP_001036293.1   --------------------------------------------------
034 YP_605811.1      --------------------------------------------------
035 NP_955577.1      --------------------------------------------------
036 YP_003208050.1   --------------------------------------------------
037 NP_789740.1      --------------------------------------------------
038 YP_004442839.1   --------------------------------------------------
039 YP_009507791.1   --------------------------------------------------
040 YP_004581513.1   --------------------------------------------------
041 NP_043933.1      --------------------------------------------------
042 YP_009507248.1   --------------------------------------------------
043 YP_002117531.1   --------------------------------------------------
044 YP_004347415.1   --------------------------------------------------
045 NP_056891.1      PPPYNKATPSAPTVMAVVNPKEELKEKIAQLEEQIKLEELHQALISKLQK
046 YP_009513242.1   --------------------------------------------------
047 YP_009352866.1   --------------------------------------------------
048 YP_009165750.1   --------------------------------------------------
049 NP_040840.1      --------------------------------------------------
050 YP_004222728.1   --------------------------------------------------
051 YP_009506264.1   --------------------------------------------------
052 YP_567050.1      --------------------------------------------------
053 YP_009508571.1   --------------------------------------------------
054 NP_056803.1      --------------------------------------------------
055 NP_955564.1      --------------------------------------------------
056 YP_009508411.1   --------------------------------------------------
057 YP_009140788.1   --------------------------------------------------
058 YP_009508410.1   --------------------------------------------------
059 NP_040333.1      --------------------------------------------------
060 YP_008992013.1   --------------------------------------------------
061 NP_041734.1      --------------------------------------------------
062 NP_056895.1      --------------------------------------------------
063 YP_009109689.1   PTTPSAPTPSSLVSSST------PPSSPAPPELTPRTPPQTPRLRLRRAE
064 YP_002519387.1   --------------------------------------------------
065 YP_595725.1      --------------------------------------------------
066 YP_009508551.1   --------------------------------------------------
067 YP_001497148.1   PTSPPASSSSSLSPTSSPACTSAPSSTPAPPDLTPQTPPQTPRLRLRRLD
068 NP_041186.1      --------------------------------------------------
069 NP_663784.1      --------------------------------------------------
070 YP_009508443.1   --------------------------------------------------
071 YP_003987465.1   --------------------------------------------------
072 NP_861410.1      --------------------------------------------------
073 YP_009345071.1   --------------------------------------------------
074 YP_009508582.1   --------------------------------------------------
075 NP_569141.1      --------------------------------------------------
076 YP_002916057.1   --------------------------------------------------
077 NP_777384.2      --------------------------------------------------
078 YP_233110.1      --------------------------------------------------
079 NP_542258.1      --------------------------------------------------
080 YP_009130664.1   --------------------------------------------------
081 YP_009508566.1   --------------------------------------------------
082 YP_006607892.1   --------------------------------------------------
083 YP_009506270.1   --------------------------------------------------
084 NP_042513.1      --------------------------------------------------
085 YP_006907834.1   --------------------------------------------------
086 YP_004442827.1   --------------------------------------------------
087 YP_009408594.1   --------------------------------------------------
088 YP_009508561.1   --------------------------------------------------
089 NP_056907.1      --------------------------------------------------
090 YP_002308474.1   --------------------------------------------------
091 YP_007761644.1   --------------------------------------------------
092 NP_040550.1      --------------------------------------------------
093 NP_758808.1      --------------------------------------------------
094 YP_009345075.1   --------------------------------------------------
095 NP_068729.1      --------------------------------------------------
096 NP_047255.1      LIDLLTEEPPPYPGGHGPPPSGPRTPTASP-------------IASRLRE
097 YP_006273075.1   --------------------------------------------------
098 YP_009508888.1   --------------------------------------------------
099 NP_395469.1      --------------------------------------------------
100 YP_009508546.1   --------------------------------------------------
101 NP_056762.1      --------------------------------------------------
102 YP_009508556.1   --------------------------------------------------
```

```
103 NP_569153.1        --------------------------------------------------
104 NP_043924.1        --------------------------------------------------
105 YP_009243641.1     --------------------------------------------------
106 YP_004300274.1     --------------------------------------------------
107 NP_056848.1        --------------------------------------------------
108 YP_006495799.1     --------------------------------------------------
109 YP_009508409.1     --------------------------------------------------
110 NP_687035.1        --------------------------------------------------
111 NP_045937.2        --------------------------------------------------
112 NP_612577.1        --------------------------------------------------
113 NP_619548.1        --------------------------------------------------
114 NP_056880.1        DWNDDDLSPEDWDDLEEQAAHYHDDDELILPVKRKVVKKKPQALRRKPLP
115 YP_009121747.1     --------------------------------------------------
116 YP_009113237.2     --------------------------------------------------
117 YP_009508408.1     --------------------------------------------------
118 NP_955579.1        --------------------------------------------------
119 YP_009229919.1     --------------------------------------------------
120 YP_004442833.1     --------------------------------------------------
121 NP_862833.2        --------------------------------------------------
122 NP_040563.1        --------------------------------------------------
123 YP_443922.1        FLDQPFSSPPPSGSLSYRPLPQSLPSPLSRAVTTPHSYTGEHQQLRQNDI
124 NP_057933.2        LIDLLTEDPPPYRDPRPPPSDRDGNGGEATPAGEAPDPSPMASRLRGRRE
125 YP_009116631.1     --------------------------------------------------
126 YP_009513249.1     --------------------------------------------------
127 YP_223871.1        --------------------------------------------------
128 NP_127504.1        --------------------------------------------------
129 NP_044929.1        --------------------------------------------------
130 NP_057860.1        --------------------------------------------------
131 NP_040973.1        --------------------------------------------------
132 NP_777317.1        --------------------------------------------------
133 NP_054716.1        --------------------------------------------------
134 YP_001856242.1     --------------------------------------------------
135 NP_954565.2        PPPYNKATPSAPTVMAVVNPKEELKEKIAQLEEQIKLEELHQALISKLQK
136 NP_569150.1        --------------------------------------------------
137 NP_659397.1        --------------------------------------------------
138 YP_009513211.1     --------------------------------------------------
139 YP_233107.1        --------------------------------------------------
140 YP_009109692.1     --------------------------------------------------
141 NP_057849.4        --------------------------------------------------
142 NP_049560.1        --------------------------------------------------
143 YP_009506267.1     --------------------------------------------------
144 NP_039820.1        --------------------------------------------------
145 NP_056790.1        --------------------------------------------------
146 YP_002321513.1     --------------------------------------------------
147 NP_955591.1        --------------------------------------------------
148 YP_001956722.2     --------------------------------------------------
149 NP_056902.1        --------------------------------------------------
150 NP_056886.1        KTYCGKTSPSKKEIGAMLSLLQKEGLLMSPSDLYSPGSWDPITAALSQRA
151 NP_041261.1        --------------------------------------------------
```

```
001 YP_004442836.1     --------------------------------------------------
002 NP_056728.1        --------------------------------------------------
003 YP_009508537.1     --------------------------------------------------
004 YP_004732983.2     --------------------------------------------------
005 YP_009508407.1     --------------------------------------------------
006 NP_040939.1        --------------------------------------------------
007 YP_003864102.1     --------------------------------------------------
008 YP_009002585.1     --------------------------------------------------
009 YP_003284237.1     --------------------------------------------------
010 YP_009268869.1     --------------------------------------------------
011 YP_009506251.1     --------------------------------------------------
012 NP_955611.1        --------------------------------------------------
013 YP_008567619.1     --------------------------------------------------
014 NP_149413.1        --------------------------------------------------
015 YP_004442824.1     --------------------------------------------------
016 YP_001931967.1     --------------------------------------------------
017 YP_002455786.1     --------------------------------------------------
018 YP_009109694.1     --------------------------------------------------
019 NP_789739.1        --------------------------------------------------
020 NP_955619.1        --------------------------------------------------
021 YP_224289.1        --------------------------------------------------
```

```
022 YP_009182100.1    --------------------------------------------------
023 NP_705927.1       --------------------------------------------------
024 YP_009508577.1    --------------------------------------------------
025 YP_004442830.1    --------------------------------------------------
026 NP_758887.1       --------------------------------------------------
027 YP_009508406.1    --------------------------------------------------
028 YP_610965.1       --------------------------------------------------
029 YP_001931961.1    --------------------------------------------------
030 YP_241114.1       --------------------------------------------------
031 YP_006732334.1    --------------------------------------------------
032 YP_009041481.1    --------------------------------------------------
033 YP_001036293.1    --------------------------------------------------
034 YP_605811.1       --------------------------------------------------
035 NP_955577.1       --------------------------------------------------
036 YP_003208050.1    --------------------------------------------------
037 NP_789740.1       --------------------------------------------------
038 YP_004442839.1    --------------------------------------------------
039 YP_009507791.1    --------------------------------------------------
040 YP_004581513.1    --------------------------------------------------
041 NP_043933.1       --------------------------------------------------
042 YP_009507248.1    --------------------------------------------------
043 YP_002117531.1    --------------------------------------------------
044 YP_004347415.1    --------------------------------------------------
045 NP_056891.1       LKTGNETVTHPDTAGGLSRTPHWPGQHIPKGKCCASREKEEQIPKDIFPV
046 YP_009513242.1    --------------------------------------------------
047 YP_009352866.1    --------------------------------------------------
048 YP_009165750.1    --------------------------------------------------
049 NP_040840.1       --------------------------------------------------
050 YP_004222728.1    --------------------------------------------------
051 YP_009506264.1    --------------------------------------------------
052 YP_567050.1       --------------------------------------------------
053 YP_009508571.1    --------------------------------------------------
054 NP_056803.1       --------------------------------------------------
055 NP_955564.1       --------------------------------------------------
056 YP_009508411.1    --------------------------------------------------
057 YP_009140788.1    --------------------------------------------------
058 YP_009508410.1    --------------------------------------------------
059 NP_040333.1       --------------------------------------------------
060 YP_008992013.1    --------------------------------------------------
061 NP_041734.1       --------------------------------------------------
062 NP_056895.1       --------------------------------------------------
063 YP_009109689.1    GQDGPSTWQSSLFPLRTVNRTIQYWPFSASDLYNWKTHNPSFSQDPQALT
064 YP_002519387.1    --------------------------------------------------
065 YP_595725.1       --------------------------------------------------
066 YP_009508551.1    --------------------------------------------------
067 YP_001497148.1    DPNGPPTWQSSLFPLRTVNRTVQYWPFSASDLYNWKTHNPSFSQEPQALT
068 NP_041186.1       --------------------------------------------------
069 NP_663784.1       -----------MGARNSVLRGKKADELEKVRLRPGGKKKYRLKHIVWAAN
070 YP_009508443.1    --------------------------------------------------
071 YP_003987465.1    --------------------------------------------------
072 NP_861410.1       --------------------------------------------------
073 YP_009345071.1    --------------------------------------------------
074 YP_009508582.1    --------------------------------------------------
075 NP_569141.1       --------------------------------------------------
076 YP_002916057.1    --------------------------------------------------
077 NP_777384.2       --------------------------------------------------
078 YP_233110.1       --------------------------------------------------
079 NP_542258.1       --------------------------------------------------
080 YP_009130664.1    --------------------------------------------------
081 YP_009508566.1    --------------------------------------------------
082 YP_006607892.1    --------------------------------------------------
083 YP_009506270.1    --------------------------------------------------
084 NP_042513.1       --------------------------------------------------
085 YP_006907834.1    --------------------------------------------------
086 YP_004442827.1    --------------------------------------------------
087 YP_009408594.1    --------------------------------------------------
088 YP_009508561.1    --------------------------------------------------
089 NP_056907.1       --------------------------------------------------
090 YP_002308474.1    --------------------------------------------------
091 YP_007761644.1    --------------------------------------------------
092 NP_040550.1       --------------------------------------------------
093 NP_758808.1       --------------------------------------------------
094 YP_009345075.1    --------------------------------------------------
```

```
095 NP_068729.1       --------------------------------------------------
096 NP_047255.1       RRENPAEESQALPLREGPNNRPQYWPFSASDLYNWKSHNPPFSQDPVALT
097 YP_006273075.1    --------------------------------------------------
098 YP_009508888.1    --------------------------------------------------
099 NP_395469.1       --------------------------------------------------
100 YP_009508546.1    --------------------------------------------------
101 NP_056762.1       --------------------------------------------------
102 YP_009508556.1    --------------------------------------------------
103 NP_569153.1       --------------------------------------------------
104 NP_043924.1       --------------------------------------------------
105 YP_009243641.1    --------------------------------------------------
106 YP_004300274.1    --------------------------------------------------
107 NP_056848.1       --------------------------------------------------
108 YP_006495799.1    --------------------------------------------------
109 YP_009508409.1    --------------------------------------------------
110 NP_687035.1       -----------MGGGHSALSGRSLDTFEKIRLRPNGKKKYQIKHLIWAGK
111 NP_045937.2       --------------------------------------------------
112 NP_612577.1       --------------------------------------------------
113 NP_619548.1       --------------------------------------------------
114 NP_056880.1       PVGFAGAMAEAREKGDLTFT---------------------------FPV
115 YP_009121747.1    --------------------------------------------------
116 YP_009113237.2    --------------------------------------------------
117 YP_009508408.1    --------------------------------------------------
118 NP_955579.1       --------------------------------------------------
119 YP_009229919.1    --------------------------------------------------
120 YP_004442833.1    --------------------------------------------------
121 NP_862833.2       --------------------------------------------------
122 NP_040563.1       --------------------------------------------------
123 YP_443922.1       DREYLTSYNMPQEEDENDPAPTQEQIWTAAEQLNREQFPSGGGGPYQLRG
124 NP_057933.2       PPVADSTTSQAFPLRAGGNGQLQYWPFSSSDLYNWKNNNPSFSEDPGKLT
125 YP_009116631.1    --------------------------------------------------
126 YP_009513249.1    --------------------------------------------------
127 YP_223871.1       --------------------------------------------------
128 NP_127504.1       --------------------------------------------------
129 NP_044929.1       --------------------------------------------------
130 NP_057860.1       --------------------------------------------------
131 NP_040973.1       --------------------------------------------------
132 NP_777317.1       --------------------------------------------------
133 NP_054716.1       --------------------------------------------------
134 YP_001856242.1    --------------------------------------------------
135 NP_954565.2       LKTGNETVTHPDTAGGLSRTPHWPGQHIPKGKCCASREKEEQIPKDIFPV
136 NP_569150.1       --------------------------------------------------
137 NP_659397.1       --------------------------------------------------
138 YP_009513211.1    --------------------------------------------------
139 YP_233107.1       --------------------------------------------------
140 YP_009109692.1    --------------------------------------------------
141 NP_057849.4       -----------MGARASVLSGGELDRWEKIRLRPGGKKKYKLKHIVWASR
142 NP_049560.1       --------------------------------------------------
143 YP_009506267.1    --------------------------------------------------
144 NP_039820.1       --------------------------------------------------
145 NP_056790.1       --------------------------------------------------
146 YP_002321513.1    --------------------------------------------------
147 NP_955591.1       --------------------------------------------------
148 YP_001956722.2    --------------------------------------------------
149 NP_056902.1       --------------------------------------------------
150 NP_056886.1       MILGKSGELKTWGLVLGALKAAREEQVTSEQAKFWLGLGGGRVSPPGPEC
151 NP_041261.1       --------------------------------------------------


001 YP_004442836.1    --------------------------------------------------
002 NP_056728.1       --------------------------------------------------
003 YP_009508537.1    --------------------------------------------------
004 YP_004732983.2    --------------------------------------------------
005 YP_009508407.1    --------------------------------------------------
006 NP_040939.1       --------------------------------------------------
007 YP_003864102.1    --------------------------------------------------
008 YP_009002585.1    --------------------------------------------------
009 YP_003284237.1    --------------------------------------------------
010 YP_009268869.1    --------------------------------------------------
011 YP_009506251.1    --------------------------------------------------
012 NP_955611.1       --------------------------------------------------
013 YP_008567619.1    --------------------------------------------------
```

```
014 NP_149413.1        --------------------------------------------------
015 YP_004442824.1     --------------------------------------------------
016 YP_001931967.1     --------------------------------------------------
017 YP_002455786.1     --------------------------------------------------
018 YP_009109694.1     --------------------------------------------------
019 NP_789739.1        --------------------------------------------------
020 NP_955619.1        --------------------------------------------------
021 YP_224289.1        --------------------------------------------------
022 YP_009182100.1     --------------------------------------------------
023 NP_705927.1        --------------------------------------------------
024 YP_009508577.1     --------------------------------------------------
025 YP_004442830.1     --------------------------------------------------
026 NP_758887.1        --------------------------------------------------
027 YP_009508406.1     --------------------------------------------------
028 YP_610965.1        --------------------------------------------------
029 YP_001931961.1     --------------------------------------------------
030 YP_241114.1        --------------------------------------------------
031 YP_006732334.1     --------------------------------------------------
032 YP_009041481.1     --------------------------------------------------
033 YP_001036293.1     --------------------------------------------------
034 YP_605811.1        --------------------------------------------------
035 NP_955577.1        --------------------------------------------------
036 YP_003208050.1     --------------------------------------------------
037 NP_789740.1        --------------------------------------------------
038 YP_004442839.1     --------------------------------------------------
039 YP_009507791.1     --------------------------------------------------
040 YP_004581513.1     --------------------------------------------------
041 NP_043933.1        --------------------------------------------------
042 YP_009507248.1     --------------------------------------------------
043 YP_002117531.1     --------------------------------------------------
044 YP_004347415.1     --------------------------------------------------
045 NP_056891.1        TETVDGQGQAWRHHNGFDFAVIKELKTAASQYGATAPYTLAIVESVADNW
046 YP_009513242.1     --------------------------------------------------
047 YP_009352866.1     --------------------------------------------------
048 YP_009165750.1     --------------------------------------------------
049 NP_040840.1        --------------------------------------------------
050 YP_004222728.1     --------------------------------------------------
051 YP_009506264.1     --------------------------------------------------
052 YP_567050.1        --------------------------------------------------
053 YP_009508571.1     --------------------------------------------------
054 NP_056803.1        --------------------------------------------------
055 NP_955564.1        --------------------------------------------------
056 YP_009508411.1     --------------------------------------------------
057 YP_009140788.1     --------------------------------------------------
058 YP_009508410.1     --------------------------------------------------
059 NP_040333.1        --------------------------------------------------
060 YP_008992013.1     --------------------------------------------------
061 NP_041734.1        --------------------------------------------------
062 NP_056895.1        -----------------MGNSPSYNPPAGISPSDWLNLRQSAQRLNPRPS
063 YP_009109689.1     SLIESILLTHQPTWDDCQQLLQVLLTTEERQRVLLEARKNVPGPGGLPTQ
064 YP_002519387.1     --------------------------------------------------
065 YP_595725.1        --------------------------------------------------
066 YP_009508551.1     --------------------------------------------------
067 YP_001497148.1     SLIESILLTHQPTWDDCQQLLQVLLTTEERQRVLLEARKNVPGPGGFPTQ
068 NP_041186.1        --------------------------------------------------
069 NP_663784.1        ELDKFGLAESLLESKEGCQKILRVLDPLVPTGSENLKSLFNTVCVIWCLH
070 YP_009508443.1     --------------------------------------------------
071 YP_003987465.1     --------------------------------------------------
072 NP_861410.1        --------------------------------------------------
073 YP_009345071.1     --------------------------------------------------
074 YP_009508582.1     --------------------------------------------------
075 NP_569141.1        --------------------------------------------------
076 YP_002916057.1     --------------------------------------------------
077 NP_777384.2        -----------------MGNSPSYNPPAGISPSDWLNLRQSAQRLNPRPS
078 YP_233110.1        --------------------------------------------------
079 NP_542258.1        --------------------------------------------------
080 YP_009130664.1     --------------------------------------------------
081 YP_009508566.1     --------------------------------------------------
082 YP_006607892.1     --------------------------------------------------
083 YP_009506270.1     --------------------------------------------------
084 NP_042513.1        --------------------------------------------------
085 YP_006907834.1     --------------------------------------------------
086 YP_004442827.1     --------------------------------------------------
```

```
087 YP_009408594.1    --------------------------------------------------
088 YP_009508561.1    --------------------------------------------------
089 NP_056907.1       -----------MGQTYGLSSSPIPKAPRGLSTHHWLNFLQAAYRLQPGPS
090 YP_002308474.1    --------------------------------------------------
091 YP_007761644.1    --------------------------------------------------
092 NP_040550.1       --------------------------------------------------
093 NP_758808.1       --------------------------------------------------
094 YP_009345075.1    --------------------------------------------------
095 NP_068729.1       --------------------------------------------------
096 NP_047255.1       NLIESILVTHQPTWDDCQQLLQALLTGEERQRVLLEARKQVPGEDGRPTQ
097 YP_006273075.1    --------------------------------------------------
098 YP_009508888.1    --------------------------------------------------
099 NP_395469.1       --------------------------------------------------
100 YP_009508546.1    --------------------------------------------------
101 NP_056762.1       --------------------------------------------------
102 YP_009508556.1    --------------------------------------------------
103 NP_569153.1       --------------------------------------------------
104 NP_043924.1       --------------------------------------------------
105 YP_009243641.1    --------------------------------------------------
106 YP_004300274.1    --------------------------------------------------
107 NP_056848.1       --------------------------------------------------
108 YP_006495799.1    --------------------------------------------------
109 YP_009508409.1    --------------------------------------------------
110 NP_687035.1       EMERFGLHEKLLETKEGCQKIIEVLTPLEPTGSEGLKALFNLCCVIWCIH
111 NP_045937.2       --------------------------------------------------
112 NP_612577.1       --------------------------------------------------
113 NP_619548.1       --------------------------------------------------
114 NP_056880.1       VFMGESDEDDTPVWEPLPLKTLKELQSAVRTMGPSAPYTLQVVDMVASQW
115 YP_009121747.1    --------------------------------------------------
116 YP_009113237.2    --------------------------------------------------
117 YP_009508408.1    --------------------------------------------------
118 NP_955579.1       --------------------------------------------------
119 YP_009229919.1    --------------------------------------------------
120 YP_004442833.1    --------------------------------------------------
121 NP_862833.2       --------------------------------------------------
122 NP_040563.1       --------------------------------------------------
123 YP_443922.1       TFLGEVQDLQTELSQPGPRRSQRLIEQRNRTHLQAPLRLCATGRGEQYSP
124 NP_057933.2       ALIESVLITHQPTWDDCQQLLGTLLTGEEKQRVLLEARKAVRGDDGRPTQ
125 YP_009116631.1    --------------------------------------------------
126 YP_009513249.1    --------------------------------------------------
127 YP_223871.1       --------------------------------------------------
128 NP_127504.1       --------------------------------------------------
129 NP_044929.1       --------------------------------------------------
130 NP_057860.1       -----------MGQIFSRSASPIPRPPRGLAAHHWLNFLQAAYRLEPGPS
131 NP_040973.1       --------------------------------------------------
132 NP_777317.1       --------------------------------------------------
133 NP_054716.1       --------------------------------------------------
134 YP_001856242.1    --------------------------------------------------
135 NP_954565.2       TETVDGQGQAWRHHNGFDFAVIKELKTAASQYGATAPYTLAIVESVADNW
136 NP_569150.1       --------------------------------------------------
137 NP_659397.1       --------------------------------------------------
138 YP_009513211.1    --------------------------------------------------
139 YP_233107.1       --------------------------------------------------
140 YP_009109692.1    --------------------------------------------------
141 NP_057849.4       ELERFAVNPGLLETSEGCRQILGQLQPSLQTGSEELRSLYNTVATLYCVH
142 NP_049560.1       --------------------------------------------------
143 YP_009506267.1    --------------------------------------------------
144 NP_039820.1       --------------------------------------------------
145 NP_056790.1       --------------------------------------------------
146 YP_002321513.1    --------------------------------------------------
147 NP_955591.1       --------------------------------------------------
148 YP_001956722.2    --------------------------------------------------
149 NP_056902.1       --------------------------------------------------
150 NP_056886.1       IEKPATERRIDKGEEVGETTVQRDAKMAPEETATPKTVGTSCYHCGTAIG
151 NP_041261.1       --------------------------------------------------
```

```
001 YP_004442836.1    --------------------------------------------------
002 NP_056728.1       --------------------------------------------------
003 YP_009508537.1    --------------------------------------------------
004 YP_004732983.2    --------------------------------------------------
005 YP_009508407.1    --------------------------------------------------
```

```
006 NP_040939.1          --------------------------------------------------
007 YP_003864102.1       --------------------------------------------------
008 YP_009002585.1       --------------------------------------------------
009 YP_003284237.1       --------------------------------------------------
010 YP_009268869.1       --------------------------------------------------
011 YP_009506251.1       --------------------------------------------------
012 NP_955611.1          --------------------------------------------------
013 YP_008567619.1       --------------------------------------------------
014 NP_149413.1          --------------------------------------------------
015 YP_004442824.1       --------------------------------------------------
016 YP_001931967.1       --------------------------------------------------
017 YP_002455786.1       --------------------------------------------------
018 YP_009109694.1       --------------------------------------------------
019 NP_789739.1          --------------------------------------------------
020 NP_955619.1          --------------------------------------------------
021 YP_224289.1          --------------------------------------------------
022 YP_009182100.1       --------------------------------------------------
023 NP_705927.1          --------------------------------------------------
024 YP_009508577.1       --------------------------------------------------
025 YP_004442830.1       --------------------------------------------------
026 NP_758887.1          --------------------------------------------------
027 YP_009508406.1       --------------------------------------------------
028 YP_610965.1          --------------------------------------------------
029 YP_001931961.1       --------------------------------------------------
030 YP_241114.1          --------------------------------------------------
031 YP_006732334.1       --------------------------------------------------
032 YP_009041481.1       --------------------------------------------------
033 YP_001036293.1       --------------------------------------------------
034 YP_605811.1          --------------------------------------------------
035 NP_955577.1          --------------------------------------------------
036 YP_003208050.1       --------------------------------------------------
037 NP_789740.1          --------------------------------------------------
038 YP_004442839.1       --------------------------------------------------
039 YP_009507791.1       --------------------------------------------------
040 YP_004581513.1       --------------------------------------------------
041 NP_043933.1          --------------------------------------------------
042 YP_009507248.1       --------------------------------------------------
043 YP_002117531.1       --------------------------------------------------
044 YP_004347415.1       --------------------------------------------------
045 NP_056891.1          LTPTDWNTLVRAVLSGGDHLLWKSEFFENCRDTAKRNQQAGNGWD-FDML
046 YP_009513242.1       --------------------------------------------------
047 YP_009352866.1       --------------------------------------------------
048 YP_009165750.1       --------------------------------------------------
049 NP_040840.1          --------------------------------------------------
050 YP_004222728.1       --------------------------------------------------
051 YP_009506264.1       --------------------------------------------------
052 YP_567050.1          --------------------------------------------------
053 YP_009508571.1       --------------------------------------------------
054 NP_056803.1          --------------------------------------------------
055 NP_955564.1          --------------------------------------------------
056 YP_009508411.1       --------------------------------------------------
057 YP_009140788.1       --------------------------------------------------
058 YP_009508410.1       --------------------------------------------------
059 NP_040333.1          --------------------------------------------------
060 YP_008992013.1       --------------------------------------------------
061 NP_041734.1          --------------------------------------------------
062 NP_056895.1          PSDFTDLKNYIHWFHKTQKKPWTFTSGGPASCPPGKFG------------
063 YP_009109689.1       LPNEIDEGFPLTRPDWD---------------------------------
064 YP_002519387.1       --------------------------------------------------
065 YP_595725.1          --------------------------------------------------
066 YP_009508551.1       --------------------------------------------------
067 YP_001497148.1       LPNEIDEGFPLTRPDWD---------------------------------
068 NP_041186.1          --------------------------------------------------
069 NP_663784.1          AEEKVKDTEEAKKLAQRHLVAETGTAEKMPNTSRPTAPPSGKRGNYPVQQ
070 YP_009508443.1       --------------------------------------------------
071 YP_003987465.1       --------------------------------------------------
072 NP_861410.1          --------------------------------------------------
073 YP_009345071.1       --------------------------------------------------
074 YP_009508582.1       --------------------------------------------------
075 NP_569141.1          --------------------------------------------------
076 YP_002916057.1       --------------------------------------------------
077 NP_777384.2          PSDFTDLKNYIHWFHKTQKKPWTFTSGGPASCPPGKFG------------
078 YP_233110.1          --------------------------------------------------
```

```
079 NP_542258.1       --------------------------------------------------
080 YP_009130664.1    --------------------------------------------------
081 YP_009508566.1    --------------------------------------------------
082 YP_006607892.1    --------------------------------------------------
083 YP_009506270.1    --------------------------------------------------
084 NP_042513.1       --------------------------------------------------
085 YP_006907834.1    --------------------------------------------------
086 YP_004442827.1    --------------------------------------------------
087 YP_009408594.1    --------------------------------------------------
088 YP_009508561.1    --------------------------------------------------
089 NP_056907.1       DFDFQQLRRFLKLALKTPIWLNPIDYSLLASLIPKGYPGRTIEIINVLIK
090 YP_002308474.1    --------------------------------------------------
091 YP_007761644.1    --------------------------------------------------
092 NP_040550.1       --------------------------------------------------
093 NP_758808.1       --------------------------------------------------
094 YP_009345075.1    --------------------------------------------------
095 NP_068729.1       --------------------------------------------------
096 NP_047255.1       LPNVIDETFPLTRPNWD---------------------------------
097 YP_006273075.1    --------------------------------------------------
098 YP_009508888.1    --------------------------------------------------
099 NP_395469.1       --------------------------------------------------
100 YP_009508546.1    --------------------------------------------------
101 NP_056762.1       --------------------------------------------------
102 YP_009508556.1    --------------------------------------------------
103 NP_569153.1       --------------------------------------------------
104 NP_043924.1       --------------------------------------------------
105 YP_009243641.1    --------------------------------------------------
106 YP_004300274.1    --------------------------------------------------
107 NP_056848.1       --------------------------------------------------
108 YP_006495799.1    --------------------------------------------------
109 YP_009508409.1    --------------------------------------------------
110 NP_687035.1       AEQKVKDTEEAVVTVKQHYHLVDKNEKAAKKKNETTAPPGGESRNYPVVN
111 NP_045937.2       --------------------------------------------------
112 NP_612577.1       --------------------------------------------------
113 NP_619548.1       --------------------------------------------------
114 NP_056880.1       LTPSDWHQTARATLSPGDYVLWRTEYEEKSKEMVQKAAGKRKGKVSLDML
115 YP_009121747.1    --------------------------------------------------
116 YP_009113237.2    --------------------------------------------------
117 YP_009508408.1    --------------------------------------------------
118 NP_955579.1       --------------------------------------------------
119 YP_009229919.1    --------------------------------------------------
120 YP_004442833.1    --------------------------------------------------
121 NP_862833.2       --------------------------------------------------
122 NP_040563.1       --------------------------------------------------
123 YP_443922.1       WKLTDLTSMMEQLPSLHGGASAWLLQLQTLTSGFRLGLGDMRALLARATD
124 NP_057933.2       LPNEVDAAFPLERPDWD---------------------------------
125 YP_009116631.1    --------------------------------------------------
126 YP_009513249.1    --------------------------------------------------
127 YP_223871.1       --------------------------------------------------
128 NP_127504.1       --------------------------------------------------
129 NP_044929.1       --------------------------------------------------
130 NP_057860.1       SYDFHQLKKFLKIALETPVWICPINYSLLASLLPKGYPGRVNEILHILIQ
131 NP_040973.1       --------------------------------------------------
132 NP_777317.1       --------------------------------------------------
133 NP_054716.1       --------------------------------------------------
134 YP_001856242.1    --------------------------------------------------
135 NP_954565.2       LTPTDWNTLVRAVLSGGDHLLWKSEFFENCRDTAKRNQQAGNGWD-FDML
136 NP_569150.1       --------------------------------------------------
137 NP_659397.1       --------------------------------------------------
138 YP_009513211.1    --------------------------------------------------
139 YP_233107.1       --------------------------------------------------
140 YP_009109692.1    --------------------------------------------------
141 NP_057849.4       QRIEIKDTKEALDKIEEEQNKSKKKAQQAAADTGHSNQVSQN--YPIVQN
142 NP_049560.1       --------------------------------------------------
143 YP_009506267.1    --------------------------------------------------
144 NP_039820.1       --------------------------------------------------
145 NP_056790.1       --------------------------------------------------
146 YP_002321513.1    --------------------------------------------------
147 NP_955591.1       --------------------------------------------------
148 YP_001956722.2    --------------------------------------------------
149 NP_056902.1       --------------------------------------------------
150 NP_056886.1       CNCATASAPPPPYVGSGLYPSLAGVGEQQGQGGDTPPGAEQSRAEPGHAG
151 NP_041261.1       --------------------------------------------------
```

```
001 YP_004442836.1 --------------------------------------------------
002 NP_056728.1    --------------------------------------------------
003 YP_009508537.1 --------------------------------------------------
004 YP_004732983.2 --------------------------------------------------
005 YP_009508407.1 --------------------------------------------------
006 NP_040939.1    --------------------------------------------------
007 YP_003864102.1 --------------------------------------------------
008 YP_009002585.1 --------------------------------------------------
009 YP_003284237.1 --------------------------------------------------
010 YP_009268869.1 --------------------------------------------------
011 YP_009506251.1 --------------------------------------------------
012 NP_955611.1    --------------------------------------------------
013 YP_008567619.1 --------------------------------------------------
014 NP_149413.1    --------------------------------------------------
015 YP_004442824.1 --------------------------------------------------
016 YP_001931967.1 --------------------------------------------------
017 YP_002455786.1 --------------------------------------------------
018 YP_009109694.1 --------------------------------------------------
019 NP_789739.1    --------------------------------------------------
020 NP_955619.1    --------------------------------------------------
021 YP_224289.1    --------------------------------------------------
022 YP_009182100.1 --------------------------------------------------
023 NP_705927.1    --------------------------------------------------
024 YP_009508577.1 --------------------------------------------------
025 YP_004442830.1 --------------------------------------------------
026 NP_758887.1    --------------------------------------------------
027 YP_009508406.1 --------------------------------------------------
028 YP_610965.1    --------------------------------------------------
029 YP_001931961.1 --------------------------------------------------
030 YP_241114.1    --------------------------------------------------
031 YP_006732334.1 --------------------------------------------------
032 YP_009041481.1 --------------------------------------------------
033 YP_001036293.1 --------------------------------------------------
034 YP_605811.1    --------------------------------------------------
035 NP_955577.1    --------------------------------------------------
036 YP_003208050.1 --------------------------------------------------
037 NP_789740.1    --------------------------------------------------
038 YP_004442839.1 --------------------------------------------------
039 YP_009507791.1 --------------------------------------------------
040 YP_004581513.1 --------------------------------------------------
041 NP_043933.1    --------------------------------------------------
042 YP_009507248.1 --------------------------------------------------
043 YP_002117531.1 --------------------------------------------------
044 YP_004347415.1 --------------------------------------------------
045 NP_056891.1    TGSGNYSSTDAQMQYDPGLFAQIQAAATKAWRKLPVKGDPGASLTGVKQG
046 YP_009513242.1 --------------------------------------------------
047 YP_009352866.1 --------------------------------------------------
048 YP_009165750.1 --------------------------------------------------
049 NP_040840.1    --------------------------------------------------
050 YP_004222728.1 --------------------------------------------------
051 YP_009506264.1 --------------------------------------------------
052 YP_567050.1    --------------------------------------------------
053 YP_009508571.1 --------------------------------------------------
054 NP_056803.1    --------------------------------------------------
055 NP_955564.1    --------------------------------------------------
056 YP_009508411.1 --------------------------------------------------
057 YP_009140788.1 --------------------------------------------------
058 YP_009508410.1 --------------------------------------------------
059 NP_040333.1    --------------------------------------------------
060 YP_008992013.1 --------------------------------------------------
061 NP_041734.1    --------------------------------------------------
062 NP_056895.1    ---------RVPLVLATLNEVLSNDEGAPGASAPEEQPPPYDPPAILPII
063 YP_009109689.1 ------------YETAPGRESLRIYRQALLAGLKGAGKRPTNLAKVRTIT
064 YP_002519387.1 --------------------------------------------------
065 YP_595725.1    --------------------------------------------------
066 YP_009508551.1 --------------------------------------------------
067 YP_001497148.1 ------------YETAPGRESLRIYRQALLAGLKGAGKRPTNLAKVRTII
068 NP_041186.1    --------------------------------------------------
069 NP_663784.1    AGGNYVHVPLSPRTLNAWVKLVEEKKFGAEVVPGFQALSEGCTPYDINQM
070 YP_009508443.1 --------------------------------------------------
```

071 YP_003987465.1 --------------------------------------------------
072 NP_861410.1 --------------------------------------------------
073 YP_009345071.1 --------------------------------------------------
074 YP_009508582.1 --------------------------------------------------
075 NP_569141.1 --------------------------------------------------
076 YP_002916057.1 --------------------------------------------------
077 NP_777384.2 ---------RVPLVLATLNEVLSNDEGAPGASAPEEQPPPYDPPAILPII
078 YP_233110.1 --------------------------------------------------
079 NP_542258.1 --------------------------------------------------
080 YP_009130664.1 --------------------------------------------------
081 YP_009508566.1 --------------------------------------------------
082 YP_006607892.1 --------------------------------------------------
083 YP_009506270.1 --------------------------------------------------
084 NP_042513.1 --------------------------------------------------
085 YP_006907834.1 --------------------------------------------------
086 YP_004442827.1 --------------------------------------------------
087 YP_009408594.1 --------------------------------------------------
088 YP_009508561.1 --------------------------------------------------
089 NP_056907.1 NQTSPTPPPAPSLPEPANPPPLQQPSAPPEPHTPPPYIKPPATHCLPILH
090 YP_002308474.1 --------------------------------------------------
091 YP_007761644.1 --------------------------------------------------
092 NP_040550.1 --------------------------------------------------
093 NP_758808.1 --------------------------------------------------
094 YP_009345075.1 --------------------------------------------------
095 NP_068729.1 --------------------------------------------------
096 NP_047255.1 ------------FATPAGREHLRLYRQLLLAGLRGAARRPTNLAQVKQVV
097 YP_006273075.1 --------------------------------------------------
098 YP_009508888.1 --------------------------------------------------
099 NP_395469.1 --------------------------------------------------
100 YP_009508546.1 --------------------------------------------------
101 NP_056762.1 --------------------------------------------------
102 YP_009508556.1 --------------------------------------------------
103 NP_569153.1 --------------------------------------------------
104 NP_043924.1 -----MASNKWFVYSDEPTKVILKRDKSKEKDETKKKKIKTEQNSDAAYE
105 YP_009243641.1 --------------------------------------------------
106 YP_004300274.1 --------------------------------------------------
107 NP_056848.1 --------------------------------------------------
108 YP_006495799.1 --------------------------------------------------
109 YP_009508409.1 --------------------------------------------------
110 NP_687035.1 QNNAWVHQPLSPRTLNAWVKCVEEKRWGAEVVPMFQALSEGCLSYDVNQM
111 NP_045937.2 --------------------------------------------------
112 NP_612577.1 --------------------------------------------------
113 NP_619548.1 --------------------------------------------------
114 NP_056880.1 LGTGQFLSPSSQIKLSKDVLKDVTTNAVLAWRAIPPPGVKKTVLAGLKQG
115 YP_009121747.1 --------------------------------------------------
116 YP_009113237.2 --------------------------------------------------
117 YP_009508408.1 --------------------------------------------------
118 NP_955579.1 --------------------------------------------------
119 YP_009229919.1 --------------------------------------------------
120 YP_004442833.1 --------------------------------------------------
121 NP_862833.2 --------------------------------------------------
122 NP_040563.1 --------------------------------------------------
123 YP_443922.1 HVTMNVIMKTAGLDHLPGPTPPEVYTGPLWAELRRNYPTERDYAALSSFL
124 NP_057933.2 ------------YTTQAGRNHLVHYRQLLLAGLQNAGRSPTNLAKVKGIT
125 YP_009116631.1 --------------------------------------------------
126 YP_009513249.1 --------------------------------------------------
127 YP_223871.1 --------------------------------------------------
128 NP_127504.1 --------------------------------------------------
129 NP_044929.1 --------------------------------------------------
130 NP_057860.1 -----TQAQIPSRPAPPPPSSSTHDPPDSDPQIPPPYVEPTAPQVLPVMH
131 NP_040973.1 --------------------------------------------------
132 NP_777317.1 --------------------------------------------------
133 NP_054716.1 --------------------------------------------------
134 YP_001856242.1 --------------------------------------------------
135 NP_954565.2 TGSGNYSSTDAQMQYDPGLFAQIQAAATKAWRKLPVKGDPGASLTGVKQG
136 NP_569150.1 --------------------------------------------------
137 NP_659397.1 --------------------------------------------------
138 YP_009513211.1 --------------------------------------------------
139 YP_233107.1 --------------------------------------------------
140 YP_009109692.1 --------------------------------------------------
141 NP_057849.4 IQGQMVHQAISPRTLNAWVKVVEEKAFSPEVIPMFSALSEGATPQDLNTM
142 NP_049560.1 --------------------------------------------------
143 YP_009506267.1 --------------------------------------------------

```
144 NP_039820.1       --------------------------------------------------
145 NP_056790.1       --------------------------------------------------
146 YP_002321513.1    --------------------------------------------------
147 NP_955591.1       --------------------------------------------------
148 YP_001956722.2    --------------------------------------------------
149 NP_056902.1       --------------------------------------------------
150 NP_056886.1       QAPGPALTDWARVREELASTGPPVVAMPVVIKTEGPAWTPLEPKLITRLA
151 NP_041261.1       --------------------------------------------------
```

```
001 YP_004442836.1    --------------------------------------------------
002 NP_056728.1       --------------------------------------------------
003 YP_009508537.1    --------------------------------------------------
004 YP_004732983.2    --------------------------------------------------
005 YP_009508407.1    --------------------------------------------------
006 NP_040939.1       --------------------------------------------------
007 YP_003864102.1    --------------------------------------------------
008 YP_009002585.1    --------------------------------------------------
009 YP_003284237.1    --------------------------------------------------
010 YP_009268869.1    --------------------------------------------------
011 YP_009506251.1    --------------------------------------------------
012 NP_955611.1       --------------------------------------------------
013 YP_008567619.1    --------------------------------------------------
014 NP_149413.1       --------------------------------------------------
015 YP_004442824.1    --------------------------------------------------
016 YP_001931967.1    --------------------------------------------------
017 YP_002455786.1    --------------------------------------------------
018 YP_009109694.1    --------------------------------------------------
019 NP_789739.1       --------------------------------------------------
020 NP_955619.1       --------------------------------------------------
021 YP_224289.1       --------------------------------------------------
022 YP_009182100.1    --------------------------------------------------
023 NP_705927.1       --------------------------------------------------
024 YP_009508577.1    --------------------------------------------------
025 YP_004442830.1    --------------------------------------------------
026 NP_758887.1       --------------------------------------------------
027 YP_009508406.1    --------------------------------------------------
028 YP_610965.1       --------------------------------------------------
029 YP_001931961.1    --------------------------------------------------
030 YP_241114.1       --------------------------------------------------
031 YP_006732334.1    --------------------------------------------------
032 YP_009041481.1    --------------------------------------------------
033 YP_001036293.1    --------------------------------------------------
034 YP_605811.1       --------------------------------------------------
035 NP_955577.1       --------------------------------------------------
036 YP_003208050.1    --------------------------------------------MPEAAA
037 NP_789740.1       --------------------------------------------------
038 YP_004442839.1    --------------------------------------------------
039 YP_009507791.1    --------------------------------------------------
040 YP_004581513.1    --------------------------------------------------
041 NP_043933.1       --------------------------------------------------
042 YP_009507248.1    --------------------------------------------------
043 YP_002117531.1    --------------------------------------------------
044 YP_004347415.1    --------------------------------------------------
045 NP_056891.1       PDEPFADFVHRLITTAGRIFGSAEAGVDYVKQLAYENANPACQAAIRPYR
046 YP_009513242.1    --------------------------------------------------
047 YP_009352866.1    --------------------------------------------------
048 YP_009165750.1    --------------------------------------------------
049 NP_040840.1       --------------------------------------------------
050 YP_004222728.1    --------------------------------------------------
051 YP_009506264.1    --------------------------------------------------
052 YP_567050.1       --------------------------------------------------
053 YP_009508571.1    --------------------------------------------------
054 NP_056803.1       --------------------------------------------------
055 NP_955564.1       --------------------------------------------------
056 YP_009508411.1    --------------------------------------------------
057 YP_009140788.1    --------------------------------------------------
058 YP_009508410.1    --------------------------------------------------
059 NP_040333.1       --------------------------------------------------
060 YP_008992013.1    --------------------------------------------------
061 NP_041734.1       --------------------------------------------------
062 NP_056895.1       SEGNRNRHRAWALRELQDIKKEIENKAPGSQVWIQTLRLAILQADPTPAD
```

```
063 YP_009109689.1    QGKDESPAAFMERLLEGFRMYT----------------------------
064 YP_002519387.1    --------------------------------------------------
065 YP_595725.1       --------------------------------------------------
066 YP_009508551.1    --------------------------------------------------
067 YP_001497148.1    QGKEESPAAFMERLLEGFRMYT----------------------------
068 NP_041186.1       --------------------------------------------------
069 NP_663784.1       LNCVGDHQAAMQIIREIINEEAADWDSQHPIPG-PLPAGQLRDPRGSDIA
070 YP_009508443.1    --------------------------------------------------
071 YP_003987465.1    --------------------------------------------------
072 NP_861410.1       --------------------------------------------------
073 YP_009345071.1    --------------------------------------------------
074 YP_009508582.1    --------------------------------------------------
075 NP_569141.1       --------------------------------------------------
076 YP_002916057.1    --------------------------------------------------
077 NP_777384.2       SEGNRNRHRAWALRELQDIKKEIENKAPGSQVWIQTLRLAILQADPTPAD
078 YP_233110.1       --------------------------------------------------
079 NP_542258.1       --------------------------------------------------
080 YP_009130664.1    --------------------------------------------------
081 YP_009508566.1    --------------------------------------------------
082 YP_006607892.1    --------------------------------------------------
083 YP_009506270.1    --------------------------------------------------
084 NP_042513.1       --------------------------------------------------
085 YP_006907834.1    --------------------------------------------------
086 YP_004442827.1    --------------------------------------------------
087 YP_009408594.1    --------------------------------------------------
088 YP_009508561.1    --------------------------------------------------
089 NP_056907.1       PHGAPSAHSPWQMKDLQAIKQEVSTSAPGSPQFMQTVRLAIQQFDPTAKD
090 YP_002308474.1    --------------------------------------------------
091 YP_007761644.1    --------------------------------------------------
092 NP_040550.1       --------------------------------------------------
093 NP_758808.1       --------------------------------------------------
094 YP_009345075.1    --------------------------------------------------
095 NP_068729.1       --------------------------------------------------
096 NP_047255.1       QGKEETPAAFLERLKEAYRMYT----------------------------
097 YP_006273075.1    --------------------------------------------------
098 YP_009508888.1    --------------------------------------------------
099 NP_395469.1       --------------------------------------------------
100 YP_009508546.1    --------------------------------------------------
101 NP_056762.1       --------------------------------------------------
102 YP_009508556.1    --------------------------------------------------
103 NP_569153.1       --------------------------------------------------
104 NP_043924.1       TPGTAPVQKPLLETTPEAELEKVLKGLEEWGYKALEKKRDPELWNPNQEG
105 YP_009243641.1    --------------------------------------------------
106 YP_004300274.1    --------------------------------------------------
107 NP_056848.1       --------------------------------------------------
108 YP_006495799.1    --------------------------------------------------
109 YP_009508409.1    --------------------------------------------------
110 NP_687035.1       LNVIGDHQGALQILKEVINEEAAEWDRTHRPPAGPLPAGQLRDPTGSDIA
111 NP_045937.2       --------------------------------------------------
112 NP_612577.1       --------------------------------------------------
113 NP_619548.1       --------------------------------------------------
114 NP_056880.1       NEESYETFISRLEEAVYRMMPRGEGSDILIKQLAWENANSLCQDLIRPIR
115 YP_009121747.1    --------------------------------------------------
116 YP_009113237.2    --------------------------------------------------
117 YP_009508408.1    --------------------------------------------------
118 NP_955579.1       --------------------------------------------------
119 YP_009229919.1    --------------------------------------------------
120 YP_004442833.1    --------------------------------------------------
121 NP_862833.2       --------------------------------------------------
122 NP_040563.1       --------------------------------------------------
123 YP_443922.1       MNPGEGPAEYLDRAKTTWRTVHGEPHDASETSLNMWKEVVINGTPAPVKA
124 NP_057933.2       QGPNESPSAFLERLKEAYRRYT----------------------------
125 YP_009116631.1    --------------------------------------------------
126 YP_009513249.1    --------------------------------------------------
127 YP_223871.1       --------------------------------------------------
128 NP_127504.1       --------------------------------------------------
129 NP_044929.1       --------------------------------------------------
130 NP_057860.1       PHGAPPNHRPWQMKDLQAIKQEVSQAAPGSPQFMQTIRLAVQQFDPTAKD
131 NP_040973.1       --------------------------------------------------
132 NP_777317.1       --------------------------------------------------
133 NP_054716.1       --------------------------------------------------
134 YP_001856242.1    --------------------------------------------------
135 NP_954565.2       PDEPFADFVHRLITTAGRIFGSAEAGVDYVKQLAYENANPACQAAIRPYR
```

```
136 NP_569150.1        --------------------------------------------------
137 NP_659397.1        --------------------------------------------------
138 YP_009513211.1     --------------------------------------------------
139 YP_233107.1        --------------------------------------------------
140 YP_009109692.1     --------------------------------------------------
141 NP_057849.4        LNTVGGHQAAMQMLKETINEEAAEWDRVHPVHAGPIAPGQMREPRGSDIA
142 NP_049560.1        --------------------------------------------------
143 YP_009506267.1     --------------------------------------------------
144 NP_039820.1        --------------------------------------------------
145 NP_056790.1        --------------------------------------------------
146 YP_002321513.1     --------------------------------------------------
147 NP_955591.1        --------------------------------------------------
148 YP_001956722.2     --------------------------------------------------
149 NP_056902.1        --------------------------------------------------
150 NP_056886.1        DTVRTKGLRSPITMAEVEALMSSPLLPHDVTNLMRVILGPAPYALWMDAW
151 NP_041261.1        --------------------------------------------------
```

```
001 YP_004442836.1     --------------------------------------------------
002 NP_056728.1        --------------------------------------------------
003 YP_009508537.1     --------------------------------------------------
004 YP_004732983.2     --------------------------------------------------
005 YP_009508407.1     --------------------------------------------------
006 NP_040939.1        --------------------------------------------------
007 YP_003864102.1     --------------------------------------------------
008 YP_009002585.1     --------------------------------------------------
009 YP_003284237.1     --------------------------------------------------
010 YP_009268869.1     --------------------------------------------------
011 YP_009506251.1     --------------------------------------------------
012 NP_955611.1        --------------------------------------------------
013 YP_008567619.1     --------------------------------------------------
014 NP_149413.1        --------------------------------------------------
015 YP_004442824.1     --------------------------------------------------
016 YP_001931967.1     --------------------------------------------------
017 YP_002455786.1     --------------------------------------------------
018 YP_009109694.1     --------------------------------------------------
019 NP_789739.1        --------------------------------------------------
020 NP_955619.1        --------------------------------------------------
021 YP_224289.1        --------------------------------------------------
022 YP_009182100.1     --------------------------------------------------
023 NP_705927.1        --------------------------------------------------
024 YP_009508577.1     --------------------------------------------------
025 YP_004442830.1     --------------------------------------------------
026 NP_758887.1        --------------------------------------------------
027 YP_009508406.1     --------------------------------------------------
028 YP_610965.1        --------------------------------------------------
029 YP_001931961.1     --------------------------------------------------
030 YP_241114.1        --------------------------------------------------
031 YP_006732334.1     --------------------------------------------------
032 YP_009041481.1     --------------------------------------------------
033 YP_001036293.1     --------------------------------------------------
034 YP_605811.1        --------------------------------------------------
035 NP_955577.1        --------------------------------------------------
036 YP_003208050.1     PAAYAANWPKGAVLYNGVNFLAADGKTVILSDAAKRIKAGVAYLRRKQRR
037 NP_789740.1        --------------------------------------------------
038 YP_004442839.1     --------------------------------------------------
039 YP_009507791.1     --------------------------------------------------
040 YP_004581513.1     --------------------------------------------------
041 NP_043933.1        --------------------------------------------------
042 YP_009507248.1     --------------------------------------------------
043 YP_002117531.1     --------------------------------------------------
044 YP_004347415.1     --------------------------------------------------
045 NP_056891.1        KKTDLTGYIRLCSDIGPSYQQGLAMAAAFSGQTVKDFL-------NNKNK
046 YP_009513242.1     --------------------------------------------------
047 YP_009352866.1     --------------------------------------------------
048 YP_009165750.1     --------------------------------------------------
049 NP_040840.1        --------------------------------------------------
050 YP_004222728.1     --------------------------------------------------
051 YP_009506264.1     --------------------------------------------------
052 YP_567050.1        --------------------------------------------------
053 YP_009508571.1     --------------------------------------------------
054 NP_056803.1        --------------------------------------------------
```

```
055 NP_955564.1        --------------------------------------------------
056 YP_009508411.1     --------------------------------------------------
057 YP_009140788.1     --------------------------------------------------
058 YP_009508410.1     --------------------------------------------------
059 NP_040333.1        --------------------------------------------------
060 YP_008992013.1     --------------------------------------------------
061 NP_041734.1        --------------------------------------------------
062 NP_056895.1        LEQLCQYIASPVDQTAHMTSLTAAIAAAEAANTLQGFNPQNGTLTQQSAQ
063 YP_009109689.1     ------PFDPEAPEHKATVAMSFIDQAALDIKGKLQRLDGIQTHGLQELV
064 YP_002519387.1     --------------------------------------------------
065 YP_595725.1        --------------------------------------------------
066 YP_009508551.1     --------------------------------------------------
067 YP_001497148.1     ------PFTPEAPEHKATVAMSFIDQAASDIKGKLQRLDGIQTYGLQELV
068 NP_041186.1        --------------------------------------------------
069 NP_663784.1        GTTSTVDEQIQWMYRPQNPVPVGNIYRRWIQIGLQKCVRKYNPTNILDIK
070 YP_009508443.1     --------------------------------------------------
071 YP_003987465.1     --------------------------------------------------
072 NP_861410.1        --------------------------------------------------
073 YP_009345071.1     --------------------------------------------------
074 YP_009508582.1     --------------------------------------------------
075 NP_569141.1        --------------------------------------------------
076 YP_002916057.1     --------------------------------------------------
077 NP_777384.2        LEQLCQYIASPVDQTAHMTSLTAAIAAAEAANTLQGFNPQNGTLTQQSAQ
078 YP_233110.1        --------------------------------------------------
079 NP_542258.1        --------------------------------------------------
080 YP_009130664.1     --------------------------------------------------
081 YP_009508566.1     --------------------------------------------------
082 YP_006607892.1     --------------------------------------------------
083 YP_009506270.1     --------------------------------------------------
084 NP_042513.1        --------------------------------------------------
085 YP_006907834.1     --------------------------------------------------
086 YP_004442827.1     --------------------------------------------------
087 YP_009408594.1     --------------------------------------------------
088 YP_009508561.1     --------------------------------------------------
089 NP_056907.1        LQDLLQYLCSSLVVSLHHQQFHTLITEAETR-GMTGYNPMAGPLRMQANN
090 YP_002308474.1     --------------------------------------------------
091 YP_007761644.1     --------------------------------------------------
092 NP_040550.1        --------------------------------------------------
093 NP_758808.1        --------------------------------------------------
094 YP_009345075.1     --------------------------------------------------
095 NP_068729.1        --------------------------------------------------
096 NP_047255.1        ------PYDPEDPGQAASVILSFIYQSSPDIRNKLQRLEGLQGFTLSDLL
097 YP_006273075.1     --------------------------------------------------
098 YP_009508888.1     --------------------------------------------------
099 NP_395469.1        --------------------------------------------------
100 YP_009508546.1     --------------------------------------------------
101 NP_056762.1        --------------------------------------------------
102 YP_009508556.1     --------------------------------------------------
103 NP_569153.1        --------------------------------------------------
104 NP_043924.1        LDEYLFRGWVQGGLADSKKALEKNMEKFVPLFVVTMSQAVPYWRQTMQAR
105 YP_009243641.1     --------------------------------------------------
106 YP_004300274.1     --------------------------------------------------
107 NP_056848.1        --------------------------------------------------
108 YP_006495799.1     --------------------------------------------------
109 YP_009508409.1     --------------------------------------------------
110 NP_687035.1        GTTSSIQEQIEWTFNANPRIDVGAQYRKWVILGLQKVVQMYNPQKVLDIR
111 NP_045937.2        ------------MGNSSSTPPPSALKNSDLFKTMLRTQYSGSVKTRRINQ
112 NP_612577.1        --------------------------------------------------
113 NP_619548.1        --------------------------------------------------
114 NP_056880.1        KTGTIQDYIRACLDASPAVVQGMAYAAAMRGQKYSTFVKQTYGGGKGGQG
115 YP_009121747.1     --------------------------------------------------
116 YP_009113237.2     ----------------------MSLRFEESIQSLYNTERLGLIQLLDLSQ
117 YP_009508408.1     --------------------------------------------------
118 NP_955579.1        --------------------------------------------------
119 YP_009229919.1     --------------------------------------------------
120 YP_004442833.1     --------------------------------------------------
121 NP_862833.2        --------------------------------------------------
122 NP_040563.1        --------------------------------------------------
123 YP_443922.1        KLRQTVGLCSKSKAEFCSHVHHHISQHHKDAGGAESEIKALQVQLLKLQL
124 NP_057933.2        ------PYDPEDPGQETNVSMSFIWQSAPDIGRKLERLEDLKNKTLGDLV
125 YP_009116631.1     --------------------------------------------------
126 YP_009513249.1     --------------------------------------------------
127 YP_223871.1        --------------------------------------------------
```

```
128 NP_127504.1          --------------------------------------------------
129 NP_044929.1          --------------------------------------------------
130 NP_057860.1          LQDLLQYLCSSLVASLHHQQLDSLISEAETR-GITGYNPLAGPLRVQANN
131 NP_040973.1          --------------------------------------------------
132 NP_777317.1          --------------------------------------------------
133 NP_054716.1          --------------------------------------------------
134 YP_001856242.1       --------------------------------------------------
135 NP_954565.2          KKTDLTGYIRLCSDIGPSYQQGLAMAAAFSGQTVKDFL-------NNKNK
136 NP_569150.1          --------------------------------------------------
137 NP_659397.1          --------------------------------------------------
138 YP_009513211.1       --------------------------------------------------
139 YP_233107.1          --------------------------------------------------
140 YP_009109692.1       --------------------------------------------------
141 NP_057849.4          GTTSTLQEQIGWMT-NNPPIPVGEIYKRWIILGLNKIVRMYSPTSILDIR
142 NP_049560.1          --------------------------------------------------
143 YP_009506267.1       --------------------------------------------------
144 NP_039820.1          --------------------------------------------------
145 NP_056790.1          --------------------------------------------------
146 YP_002321513.1       --------------------------------------------------
147 NP_955591.1          --------------------------------------------------
148 YP_001956722.2       --------------------------------------------------
149 NP_056902.1          --------------------------------------------------
150 NP_056886.1          GVQLQTVIAAATRDPRHPANGQGRGERTNLNRLKGLADGMVGNPQGQAAL
151 NP_041261.1          --------------------------------------------------
```

```
001 YP_004442836.1       ---------------------------------------------MASRP
002 NP_056728.1          --------------------------------------------------
003 YP_009508537.1       --------------------------------------------------
004 YP_004732983.2       ---------------------------------------------MSRSR
005 YP_009508407.1       --------------------------------------------------
006 NP_040939.1          --------------------------------------------------
007 YP_003864102.1       --------------------------------------------------
008 YP_009002585.1       -------------------------------------------MSTTTTQ
009 YP_003284237.1       ---------------------------------------------MSTRG
010 YP_009268869.1       --------------------------------------------------
011 YP_009506251.1       -------------------------------------------MSQTR--
012 NP_955611.1          --------------------------------------------------
013 YP_008567619.1       --------------------------------------------------
014 NP_149413.1          -------------------------------------------------M
015 YP_004442824.1       ---------------------------------------------MASRP
016 YP_001931967.1       --------------------------------------------------
017 YP_002455786.1       --------------------------------------------------
018 YP_009109694.1       --------------------------------------------------
019 NP_789739.1          --------------------------------------------------
020 NP_955619.1          --------------------------------------------------
021 YP_224289.1          -----------------------------------------MTTRRSSLP
022 YP_009182100.1       --------------------------------------------------
023 NP_705927.1          --------------------------------------------------
024 YP_009508577.1       --------------------------------------------------
025 YP_004442830.1       ---------------------------------------------MAFTG
026 NP_758887.1          --------------------------------------------------
027 YP_009508406.1       --------------------------------------------------
028 YP_610965.1          -------------------------------------------MASYRRP
029 YP_001931961.1       --------------------------------------------------
030 YP_241114.1          --------------------------------------------------
031 YP_006732334.1       --------------------------------------------------
032 YP_009041481.1       ---------------------------------------------MSQT-
033 YP_001036293.1       -----------------------------------------------MSR
034 YP_605811.1          ---------------------------------------------MASRP
035 NP_955577.1          --------------------------------------------------
036 YP_003208050.1       WAKAQMRRAVCAANKARKAYYRLTASTPIIQVGDYGFAEPISKGFRYKGP
037 NP_789740.1          --------------------------------------------------
038 YP_004442839.1       ---------------------------------------------MASRP
039 YP_009507791.1       --------------------------------------------------
040 YP_004581513.1       --------------------------------------------------
041 NP_043933.1          --------------------------------------------------
042 YP_009507248.1       --------------------------------------------------
043 YP_002117531.1       ---------------------------------------------MSR--
044 YP_004347415.1       --------------------------------------------------
045 NP_056891.1          EKGGCCFKCGKKGHFAKNCHEHAHNNAEPKVPGLCPRCKRGKHWANECKS
046 YP_009513242.1       --------------------------------------------------
```

```
047 YP_009352866.1   ---------------------------------------------MSRT-
048 YP_009165750.1   --------------------------------------------------
049 NP_040840.1      --------------------------------------------------
050 YP_004222728.1   --------------------------------------------------
051 YP_009506264.1   ---------------------------------------------MANRG
052 YP_567050.1      --------------------------------------------------
053 YP_009508571.1   --------------------------------------------------
054 NP_056803.1      --------------------------------------------------
055 NP_955564.1      --------------------------------------------------
056 YP_009508411.1   -----------------------------------------------MSL
057 YP_009140788.1   -------------------------------------------MSSSRSR
058 YP_009508410.1   --------------------------------------------------
059 NP_040333.1      --------------------------------------------------
060 YP_008992013.1   --------------------------------------------------
061 NP_041734.1      -------------------------------------------MSRARP-
062 NP_056895.1      PNAGDLRSQYQNLWLQAWKNLPTRPSVQPWSTIVQGPAESYVEFVNRLQI
063 YP_009109689.1   REAEKVYNKRETPEEREARLIKEQEERE-----------DRRDRKRDKHL
064 YP_002519387.1   --------------------------------------------------
065 YP_595725.1      -------------------------------------------------M
066 YP_009508551.1   --------------------------------------------------
067 YP_001497148.1   REAEKVYNKRETPEEKEARLAKEQEARE-----------ERRDRKRDKHL
068 NP_041186.1      --------------------------------------------------
069 NP_663784.1      QGPKEPFQSYVDRFYKSLRAEQTDPAVKNWMTQTLLIQNANPDCKLVLKG
070 YP_009508443.1   -----------------------------------------------MSR
071 YP_003987465.1   ---------------------------------------------MMN--
072 NP_861410.1      --------------------------------------------------
073 YP_009345071.1   ----------------------------------------------MTSR
074 YP_009508582.1   --------------------------------------------------
075 NP_569141.1      --------------------------------------------------
076 YP_002916057.1   --------------------------------------------------
077 NP_777384.2      PNAGDLRSQYQNLWLQAWKNLPTRPSVQPWSTIVQGPAESYVEFVNRLQI
078 YP_233110.1      ---------------------------------------------MASRP
079 NP_542258.1      --------------------------------------------------
080 YP_009130664.1   -----------------------------------------MEMTTSTTA
081 YP_009508566.1   --------------------------------------------------
082 YP_006607892.1   --------------------------------------------------
083 YP_009506270.1   ---------------------------------------------MAQRP
084 NP_042513.1      --------------------------------------------------
085 YP_006907834.1   --------------------------------------------------
086 YP_004442827.1   ---------------------------------------------MAFQG
087 YP_009408594.1   ---------------------------------------------MVVCH
088 YP_009508561.1   --------------------------------------------------
089 NP_056907.1      PAQQGLRREYQNLWLAAFSALPGNTRDPSWAAILQGLEEPYCAFVERLNV
090 YP_002308474.1   --------------------------------------------------
091 YP_007761644.1   --------------------------------------------------
092 NP_040550.1      --------------------------------------------------
093 NP_758808.1      ----------------------------------------MSL----AVR
094 YP_009345075.1   ---------------------------------------MSSQTRTTTQA
095 NP_068729.1      --------------------------------------------------
096 NP_047255.1      KEAEKIYNKRETPEEREERLWQRQEERDKKRH-----------KEMTKVL
097 YP_006273075.1   ---------------------------------------------MNRSR
098 YP_009508888.1   --------------------------------------------------
099 NP_395469.1      --------------------------------------------------
100 YP_009508546.1   --------------------------------------------------
101 NP_056762.1      --------------------------------------------------
102 YP_009508556.1   --------------------------------------------------
103 NP_569153.1      -------------------------------------------MSTSRAR
104 NP_043924.1      NQNGKKQKNRIAELEKEVADLTSAGRGADQVIAGMDKELKKTAEKYQAKL
105 YP_009243641.1   --------------------------------------------------
106 YP_004300274.1   --------------------------------------------------
107 NP_056848.1      --------------------------------------------------
108 YP_006495799.1   ---------------------------------------------MTTR-
109 YP_009508409.1   --------------------------------------------------
110 NP_687035.1      QGPKEPFQDYVDRFYKALRAEQAPQDVKNWMTQTLLIQNANPDCKLILKG
111 NP_045937.2      DIKKQYPLWPDQGTCATKHWEQAVLIPLDSVSEETAKVLNFLRVKIQARK
112 NP_612577.1      --------------------------------------------------
113 NP_619548.1      --------------------------------------------------
114 NP_056880.1      AEGPVCFSCGKTGHIRKDCKDEKGSKRAP--PGLCPRCKKGYHWKSECKS
115 YP_009121747.1   ----------------------------------------MSFGRMPSVL
116 YP_009113237.2   EPKVLNKDLSNNIDFAIKHIVLQSRVLLKHFHSIATRQEALEKMSRYGSR
117 YP_009508408.1   -----------------------------------------------MSL
118 NP_955579.1      --------------------------------------------------
119 YP_009229919.1   ----------------------------------------MSRTRTETRA
```

120 YP_004442833.1 ---------------------------------------------MALRG
121 NP_862833.2 --------------------------------------------------
122 NP_040563.1 --------------------------------------------------
123 YP_443922.1 KDAQKTAQPKKQMVEGAVPLPQLVHETVKQMLATPSPNPAPYPQPPYQPP
124 NP_057933.2 REAEKIFNKRETPEEREERIRRETEEKEERRRTEDEQKEKERDRRRHREM
125 YP_009116631.1 ---------------------------------------------MSRS-
126 YP_009513249.1 --------------------------------------------------
127 YP_223871.1 --------------------------------------------------
128 NP_127504.1 --------------------------------------------------
129 NP_044929.1 --------------------------------------------------
130 NP_057860.1 PQQQGLRREYQQLWLAAFAALPGSAKDPSWASILQGLEEPYHAFVERLNI
131 NP_040973.1 --------------------------------------------------
132 NP_777317.1 ----------------------------------------------MTRP
133 NP_054716.1 --------------------------------------------------
134 YP_001856242.1 --------------------------------------------------
135 NP_954565.2 EKGGCCFKCGKKGHFAKNCHEHAHNNAEPKVPGLCPRCKRGKHWANECKS
136 NP_569150.1 ---------------------------------------------MAQRP
137 NP_659397.1 --------------------------------------------------
138 YP_009513211.1 --------------------------------------------------
139 YP_233107.1 -----------------------------------------MTTQRAHLP
140 YP_009109692.1 --------------------------------------------------
141 NP_057849.4 QGPKEPFRDYVDRFYKTLRAEQASQEVKNWMTETLLVQNANPDCKTILKA
142 NP_049560.1 --------------------------------------------------
143 YP_009506267.1 -----------------------------------------------MER
144 NP_039820.1 --------------------------------------------MATRRL
145 NP_056790.1 --------------------------------------------------
146 YP_002321513.1 ----------------------------------------MSSSRTQTLE
147 NP_955591.1 --------------------------------------------------
148 YP_001956722.2 --------------------------------------------------
149 NP_056902.1 --------------------------------------------------
150 NP_056886.1 LRPGELVAITASALQAFREVARLAEPAGPWADIMQGPSESFVDFANRLIK
151 NP_041261.1 --------------------------------------------------

001 YP_004442836.1 RVTGSGSRTQTVAEAGTP-----LIDDQIREY------------------
002 NP_056728.1 --------------------------------------------------
003 YP_009508537.1 --------------------------------------------------
004 YP_004732983.2 TQTTELPRATRRSTSP-----VERLDDQIRGY------------------
005 YP_009508407.1 --------------------------------------------------
006 NP_040939.1 --------------------------------------------------
007 YP_003864102.1 --------------------------------------------------
008 YP_009002585.1 TVPPVTRNATRRQQGERE---TPLFEDQIRDY------------------
009 YP_003284237.1 RVTGTS--SQTVTEPGQP-----LVEDQIRDY------------------
010 YP_009268869.1 --------------------------------------------------
011 YP_009506251.1 --THQMPSVTTTVNEQG---EGPLFEDQVRNY------------------
012 NP_955611.1 --------------------------------------------------
013 YP_008567619.1 -----MAQAQAK----------TLVEDQITQY------------------
014 NP_149413.1 AQVVRGTGSSTVLEDGA------VLDDQIRDY------------------
015 YP_004442824.1 RVSGTGSRTATIAEPGVP-----LIEDQIREY------------------
016 YP_001931967.1 --------------------------------------------------
017 YP_002455786.1 --------------------------------------------------
018 YP_009109694.1 --------------------------------------------------
019 NP_789739.1 --------------------------------------------------
020 NP_955619.1 --------------------------------------------------
021 YP_224289.1 TVTETMG-PSTSGRDGTP-----LIEDQIRDY------------------
022 YP_009182100.1 --------------------------------------------------
023 NP_705927.1 --------------------------------------------------
024 YP_009508577.1 --------------------------------------------------
025 YP_004442830.1 QGRRLGTGSTTVLEDNT------ILSDQIRDY------------------
026 NP_758887.1 --------------------------------------------------
027 YP_009508406.1 --------------------------------------------------
028 YP_610965.1 AVHNNEASTSTVP----------NQEDQIRDY------------------
029 YP_001931961.1 --------------------------------------------------
030 YP_241114.1 --------------------------------------------------
031 YP_006732334.1 --------------------------------------------------
032 YP_009041481.1 -QTVTTP----------------VVEDQIRDY------------------
033 YP_001036293.1 TATQ--EVSTSQ----------RQFEDQIRDY------------------
034 YP_605811.1 RVSGSTTRT-MVAEPGVP-----LVDDQIREY------------------
035 NP_955577.1 --------------------------------------------------
036 YP_003208050.1 GAPKKEPRTTSPAERRRRVGPKAQKEEKAQDWSVLDDGNFLASLWEPEDW
037 NP_789740.1 --------------------------------------------------
038 YP_004442839.1 RVLGSTTRT-MVAEPGTP-----LVDDQIRDY------------------

```
039 YP_009507791.1   --------------------------------------------------
040 YP_004581513.1   --------------------------------------------------
041 NP_043933.1      --------------------------------------------------
042 YP_009507248.1   --------------------------------------------------
043 YP_002117531.1   --SVGGA-STSAQASGEP-----LVEDQVRDY------------------
044 YP_004347415.1   --------------------------------------------------
045 NP_056891.1      KTDNQGNPIPPHQGNRVEG---------PAPGPETSLWGSQLCSSQQKQP
046 YP_009513242.1   --------------------------------------------------
047 YP_009352866.1   -ETVTIP----------------VVEDQIRDY------------------
048 YP_009165750.1   --------------------------------------------------
049 NP_040840.1      --------------------------------------------------
050 YP_004222728.1   --------------------------------------------------
051 YP_009506264.1   RVTGRSG-STTTTEPGQP-----LVEDQIRDY------------------
052 YP_567050.1      --------------------------------------------------
053 YP_009508571.1   --------------------------------------------------
054 NP_056803.1      --------------------------------------------------
055 NP_955564.1      --------------------------------------------------
056 YP_009508411.1   ARTTEGVASTSR----------PTHEDQIRDY------------------
057 YP_009140788.1   TTTQQLPAVSSRRPESIHNRDQPLFEDQVRDY------------------
058 YP_009508410.1   --------------------------------------------------
059 NP_040333.1      --------------------------------------------------
060 YP_008992013.1   --------------------------------------------------
061 NP_041734.1      --QHPVPSVTTTTSEQNR--EGPLYEDQIRDY------------------
062 NP_056895.1      SLADNLPDGVPKEPIIDSLSYANANKECQQILQGRGLVAAPVGQKLQACA
063 YP_009109689.1   TKILAAVVTEKRAGKSGETRRRPKV---------DKDQCAYCKERGHWIK
064 YP_002519387.1   --------------------------------------------------
065 YP_595725.1      TQRVRGTGSSTITEDGA------LLDHQIRDY------------------
066 YP_009508551.1   --------------------------------------------------
067 YP_001497148.1   TKILAAVVTENRTGKSGETKRRPKI---------EKDQCAYCKERGHWIK
068 NP_041186.1      --------------------------------------------------
069 NP_663784.1      LGMNPTLEEMLTACQGVGGPGQKARLMAEALKEAMGPS---PIPFAAAQQ
070 YP_009508443.1   TQLS--APSTSG----------NTMEDQIRDY------------------
071 YP_003987465.1   ---------TTPREVEQP-----LVEDQIWGY------------------
072 NP_861410.1      --------------------------------------------------
073 YP_009345071.1   TETVTQPQTGIN------------FEDQIRGY------------------
074 YP_009508582.1   --------------------------------------------------
075 NP_569141.1      --------------------------------------------------
076 YP_002916057.1   --------------------------------------------------
077 NP_777384.2      SLADNLPDGVPKEPIIDSLSYANANKECQQILQGRGLVAAPVGQKLQACA
078 YP_233110.1      RVSGSTTRT-MIAEPGVP-----LVDDQIREY------------------
079 NP_542258.1      --------------------------------------------------
080 YP_009130664.1   RTQVQPSTSRTT----------PLFEDQIRNY------------------
081 YP_009508566.1   --------------------------------------------------
082 YP_006607892.1   --------------------------------------------------
083 YP_009506270.1   RVTGNGSRTEAIAEAGTP-----LIDDQIREY------------------
084 NP_042513.1      --------------------------------------------------
085 YP_006907834.1   --------------------------------------------------
086 YP_004442827.1   TRRRLGTGSTTVLEDNS------VLADQIRDY------------------
087 YP_009408594.1   SHSDGYTSTADRP-------------------------------------
088 YP_009508561.1   --------------------------------------------------
089 NP_056907.1      ALDNGLPEGTPKEPILRSLAYSNANKDCQKLLQARGHTNSPLGDMLRACQ
090 YP_002308474.1   --------------------------------------------------
091 YP_007761644.1   --------------------------------------------------
092 NP_040550.1      --------------------------------------------------
093 NP_758808.1      DRGSNPSTSSTVP----------SQQDQIRDY------------------
094 YP_009345075.1   PVRDAPATSVANA------------EDQIRDY------------------
095 NP_068729.1      --------------------------------------------------
096 NP_047255.1      ATVVAQNRDKDREESKLGDQRKIPL---------GKDQCAYCKEKGHWVR
097 YP_006273075.1   TVTQQLPAVSTRRREEIN-RDQPLFEDQIRDY------------------
098 YP_009508888.1   --------------------------------------------------
099 NP_395469.1      --------------------------------------------------
100 YP_009508546.1   --------------------------------------------------
101 NP_056762.1      --------------------------------------------------
102 YP_009508556.1   --------------------------------------------------
103 NP_569153.1      TVIEQLPPATTARVEERD--NTPLYDDQIRDY------------------
104 NP_043924.1      EELEEQLAAMTVEKEELESQVEGLKESLVEAETKKVSLMEVLTMPTRSKG
105 YP_009243641.1   --------------------------------------------------
106 YP_004300274.1   --------------------------------------------------
107 NP_056848.1      --------------------------------------------------
108 YP_006495799.1   TRTEQVPRATSTRPSSRG-EASSTLDDQIRGY------------------
109 YP_009508409.1   --------------------------------------------------
110 NP_687035.1      LGMNPTLEEMLIACQGVGGPQHKAKLMVEMMSNGQNMVQVGPQKKG---P
111 NP_045937.2      GETARQMTAHTIKKLIVGTIDKNKQQTEILQKTDESDEEMDTTNTMLFIA
```

```
112 NP_612577.1        --------------------------------------------------
113 NP_619548.1        --------------------------------------------------
114 NP_056880.1        KFDKDGNPLPPLETNAENSKNLVKGQSPSPAQKGDGVKGSGLNPEAPPFT
115 YP_009121747.1     GRSATTQTQTDLP----------TQEDQIRGY------------------
116 YP_009113237.2     SRTMVQQPAVTERVTEGSDGRIPLAEDQIRSY------------------
117 YP_009508408.1     VTRTREVGSTSA----------PTHEDQIRDY------------------
118 NP_955579.1        --------------------------------------------------
119 YP_009229919.1     PIRDGEGTSGNTP----------TFEDQIRSY------------------
120 YP_004442833.1     --RRLGTGSTTVLEDGT------VLSDQIRNY------------------
121 NP_862833.2        --------------------------------------------------
122 NP_040563.1        --------------------------------------------------
123 YP_443922.1        PPPLSPQYPQAYPGYGYPPRGPPQAGGHWGSRYMNNGACYNCGQPGHQAR
124 NP_057933.2        SKLLATVVSGQKQDRQGGERRRSQL---------DRDQCAYCKEKGHWAK
125 YP_009116631.1     HTRLQAPPATERATSSSD-SGTPTLEDQIRGY------------------
126 YP_009513249.1     --------------------------------------------------
127 YP_223871.1        --------------------------------------------------
128 NP_127504.1        --------------------------------------------------
129 NP_044929.1        --------------------------------------------------
130 NP_057860.1        ALDNGLPEGTPKDPILRSLAYSNANKECQKLLQARGHTNSPLGDMLRACQ
131 NP_040973.1        --------------------------------------------------
132 NP_777317.1        RIEGPVSRT-ETAEAGAP-----LVDDQIREY------------------
133 NP_054716.1        --------------------------------------------------
134 YP_001856242.1     --------------------------------------------------
135 NP_954565.2        KTDNQGNPIPPHQGNRVEG---------PAPGPETSLWGSQLCSSQQKQP
136 NP_569150.1        RITGSRT---TTAEEGTP-----LIDDQIREY------------------
137 NP_659397.1        --------------------------------------------------
138 YP_009513211.1     --------------------------------------------------
139 YP_233107.1        SINGTEVAS-TSRQPGTP-----LVEDQIRDY------------------
140 YP_009109692.1     --------------------------------------------------
141 NP_057849.4        LGPAATLEEMMTACQGVGGPGHKARVLAEAMSQVTNSATIMMQRGNFRNQ
142 NP_049560.1        --------------------------------------------------
143 YP_009506267.1     PQTS-TTLATTP----------SVFEDQVRDY------------------
144 NP_039820.1        PAVTQTDGSRTATESGVP-----EYEDQIRSY------------------
145 NP_056790.1        --------------------------------------------------
146 YP_002321513.1     ERTRPSLFYQPSAPPEEGEERPISYNDQIRDY------------------
147 NP_955591.1        --------------------------------------------------
148 YP_001956722.2     --------------------------------------------------
149 NP_056902.1        --------------------------------------------------
150 NP_056886.1        AVEGSDLPPSARAPVIIDCFRQKSQPDIQQLIRTAPSTLTTPGEIIKYVL
151 NP_041261.1        --------------------------------------------------


001 YP_004442836.1     --------------------------------------------------
002 NP_056728.1        --------------------------------------------------
003 YP_009508537.1     --------------------------------------------------
004 YP_004732983.2     --------------------------------------------------
005 YP_009508407.1     --------------------------------------------------
006 NP_040939.1        --------------------------------------------------
007 YP_003864102.1     --------------------------------------------------
008 YP_009002585.1     --------------------------------------------------
009 YP_003284237.1     --------------------------------------------------
010 YP_009268869.1     --------------------------------------------------
011 YP_009506251.1     --------------------------------------------------
012 NP_955611.1        --------------------------------------------------
013 YP_008567619.1     --------------------------------------------------
014 NP_149413.1        --------------------------------------------------
015 YP_004442824.1     --------------------------------------------------
016 YP_001931967.1     --------------------------------------------------
017 YP_002455786.1     --------------------------------------------------
018 YP_009109694.1     --------------------------------------------------
019 NP_789739.1        --------------------------------------------------
020 NP_955619.1        ------------VQPKKPPPNQPCFRCGKAGHWSRDCTQPRPPPGPCPLC
021 YP_224289.1        --------------------------------------------------
022 YP_009182100.1     --------------------------------------------------
023 NP_705927.1        --------------------------------------------------
024 YP_009508577.1     --------------------------------------------------
025 YP_004442830.1     --------------------------------------------------
026 NP_758887.1        ----------------------------------------------FFRE
027 YP_009508406.1     --------------------------------------------------
028 YP_610965.1        --------------------------------------------------
029 YP_001931961.1     --------------------------------------------------
030 YP_241114.1        --------------------------------------------------
```

```
031 YP_006732334.1    --------------------------------------------------
032 YP_009041481.1    --------------------------------------------------
033 YP_001036293.1    --------------------------------------------------
034 YP_605811.1       --------------------------------------------------
035 NP_955577.1       -------------------NLGDQESQGQDPPP-----------------
036 YP_003208050.1    ACPCSPIAESPAQMGPISARTIQGPPKRPTTRESAQEPIEECCFMMRQGS
037 NP_789740.1       --------------------------------------------------
038 YP_004442839.1    --------------------------------------------------
039 YP_009507791.1    --------------------------------------------------
040 YP_004581513.1    --------------------------------------------------
041 NP_043933.1       --------------------------------------------------
042 YP_009507248.1    --------------------------------------------------
043 YP_002117531.1    --------------------------------------------------
044 YP_004347415.1    --------------------------------------------------
045 NP_056891.1       ISKLTRATPGSAGLDLCSTSHTVLTPEMGPQALSTGIYGPLPPNTFGLIL
046 YP_009513242.1    --------------------------------------------------
047 YP_009352866.1    --------------------------------------------------
048 YP_009165750.1    --------------------------------------------------
049 NP_040840.1       --------------------------------------------------
050 YP_004222728.1    --------------------------------------------------
051 YP_009506264.1    --------------------------------------------------
052 YP_567050.1       --------------------------------------------------
053 YP_009508571.1    --------------------------------------------------
054 NP_056803.1       --------------------------------------------------
055 NP_955564.1       --------------------------------------------------
056 YP_009508411.1    --------------------------------------------------
057 YP_009140788.1    --------------------------------------------------
058 YP_009508410.1    --------------------------------------------------
059 NP_040333.1       -------------------TLDDQGGQGQEPPP-----------------
060 YP_008992013.1    -------------------------------------------------M
061 NP_041734.1       --------------------------------------------------
062 NP_056895.1       HWAPKVKQPAILVHTPGPKMPGP-----------RQPAPKRPPPGPCYRC
063 YP_009109689.1    DCPKRPRDQKKPAPVLTLGEDSEQGCQGSGAPP-----------------
064 YP_002519387.1    --------------------------------------------------
065 YP_595725.1       --------------------------------------------------
066 YP_009508551.1    --------------------------------------------------
067 YP_001497148.1    DCPKRPRDSKKSTPVLTLGEESEEGRQGSGAPP-----------------
068 NP_041186.1       --------------------------------------------------
069 NP_663784.1       RKAIRYWNCGKEGHSARQCRAPRRQGCWKCGKPGHIMANCPERQAGFFRV
070 YP_009508443.1    --------------------------------------------------
071 YP_003987465.1    --------------------------------------------------
072 NP_861410.1       --------------------------------------------------
073 YP_009345071.1    --------------------------------------------------
074 YP_009508582.1    --------------------------------------------------
075 NP_569141.1       --------------------------------------------------
076 YP_002916057.1    --------------------------------------------------
077 NP_777384.2       HWAPKVKQPAILVHTPGPKMPGP-----------RQPAPKRPPPGPCYRC
078 YP_233110.1       --------------------------------------------------
079 NP_542258.1       --------------------------------------------------
080 YP_009130664.1    --------------------------------------------------
081 YP_009508566.1    --------------------------------------------------
082 YP_006607892.1    --------------------------------------------------
083 YP_009506270.1    --------------------------------------------------
084 NP_042513.1       --------------------------------------------------
085 YP_006907834.1    --------------------------------------------------
086 YP_004442827.1    --------------------------------------------------
087 YP_009408594.1    --------------------------------------------------
088 YP_009508561.1    --------------------------------------------------
089 NP_056907.1       AWTPKDKARVLVVQPRKPPPTQPCFRCGKTGHWSRDCTLPRPPPGPCPLC
090 YP_002308474.1    --------------------------------------------------
091 YP_007761644.1    --------------------------------------------------
092 NP_040550.1       --------------------------------------------------
093 NP_758808.1       --------------------------------------------------
094 YP_009345075.1    --------------------------------------------------
095 NP_068729.1       --------------------------------------------------
096 NP_047255.1       DCPKRPRKKPAN---STLLNLGDQESQGQDPPP-----------------
097 YP_006273075.1    --------------------------------------------------
098 YP_009508888.1    --------------------------------------------------
099 NP_395469.1       --------------------------------------------------
100 YP_009508546.1    --------------------------------------------------
101 NP_056762.1       --------------------------------------------------
102 YP_009508556.1    --------------------------------------------------
103 NP_569153.1       --------------------------------------------------
```

```
104 NP_043924.1        PKKRGPDLKQIRSLHVMADSLGMDSDGIDWDWLARQAWDYEGDEDPHVKE
105 YP_009243641.1     --------------------------------------------------
106 YP_004300274.1     --------------------------------------------------
107 NP_056848.1        --------------------------------------------------
108 YP_006495799.1     --------------------------------------------------
109 YP_009508409.1     --------------------------------------------------
110 NP_687035.1        RGPLKCFNCGKFGHMQRECKAPRQIKCFKCGKIGHMAKDCKNGQANFFRV
111 NP_045937.2        RNKRERIAQQQQADLAAQQQVLLLQREQQREQREKDIKKRDEKKKKLLPD
112 NP_612577.1        --------------------------------------------------
113 NP_619548.1        --------------------------------------------------
114 NP_056880.1        IHDLPRGTPGSAGLDLSSQKDLILSLEDGVSLVPTLVKGTLPEGTTGLII
115 YP_009121747.1     --------------------------------------------------
116 YP_009113237.2     --------------------------------------------------
117 YP_009508408.1     --------------------------------------------------
118 NP_955579.1        --------------------------------------------------
119 YP_009229919.1     --------------------------------------------------
120 YP_004442833.1     --------------------------------------------------
121 NP_862833.2        --------------------------------------------------
122 NP_040563.1        --------------------------------------------------
123 YP_443922.1        NCPLPQTDSQRRFLLNRSQGQAGRGRGQPRGPLRAPQPIPAPYSPAGPSL
124 NP_057933.2        DCPKKPRGPRGPRPQTSLLTLDDQG-QGQEPPP-----------------
125 YP_009116631.1     --------------------------------------------------
126 YP_009513249.1     --------------------------------------------------
127 YP_223871.1        ------------------------GRQGSSALR-----------------
128 NP_127504.1        --------------------------------------------------
129 NP_044929.1        --------------------------------------------------
130 NP_057860.1        AWTPKDKTKVLVVQPKKPPPNQPCFRCGKAGHWSRDCTQPRPPPGPCPLC
131 NP_040973.1        --------------------------------------------------
132 NP_777317.1        --------------------------------------------------
133 NP_054716.1        --------------------------------------------------
134 YP_001856242.1     --------------------------------------------------
135 NP_954565.2        ISKLTRATPGSAGLDLCSTSHTVLTPEMGPQALSTGIYGPLPPNTFGLIL
136 NP_569150.1        --------------------------------------------------
137 NP_659397.1        --------------------------------------------------
138 YP_009513211.1     --------------------------------------------------
139 YP_233107.1        --------------------------------------------------
140 YP_009109692.1     ----------------TLGEDSEQGCQGSGAPP-----------------
141 NP_057849.4        RKIVKCFNCGKEGHTARNCRAPRKKGCWKCGKEGHQMKDC------TERQ
142 NP_049560.1        --------------------------------------------------
143 YP_009506267.1     --------------------------------------------------
144 NP_039820.1        --------------------------------------------------
145 NP_056790.1        ------------------------GSQGSDPLP-----------------
146 YP_002321513.1     --------------------------------------------------
147 NP_955591.1        --------------------------------------------------
148 YP_001956722.2     --------------------------------------------------
149 NP_056902.1        --------------------------------------------------
150 NP_056886.1        DRQKTAPLTDQGIAAAMSSAIQPLIMAVVNRERDGQTGSGGRARGLCYTC
151 NP_041261.1        --------------------------------------------------
```

```
001 YP_004442836.1     ------------RSSRRAAYEAQRVARQAGNIIGRIV--GRQPREHTLSM
002 NP_056728.1        --------------------------------------------------
003 YP_009508537.1     ----------------------------MDLLKPLTVERKGVKIKGYWDS
004 YP_004732983.2     ------------RRMARARYLAEQRIRRSFSRNYR----------ETLER
005 YP_009508407.1     --------------------------------------------------
006 NP_040939.1        ----------------------------------MSQLWKERTYAKRMQR
007 YP_003864102.1     --------------------------------------------------
008 YP_009002585.1     ------------RNSQRRAYTTQRAVRRLRRRITG-----RQPYQETLEQ
009 YP_003284237.1     ------------RRAQRARHEAARVARNIGNIGRTIV--GRQPREHTLAL
010 YP_009268869.1     --------------------------------------------------
011 YP_009506251.1     ------------RRNQRRLFNLRKKARRAQKALTGS----RSRHQETLEQ
012 NP_955611.1        --------------------------------------------------
013 YP_008567619.1     ------------RQNQRRMHNLRQAVRRLTN--SQA----STSGQEIIEQ
014 NP_149413.1        ------------RRSQHLKHELSRKTAKALSWVK-TT--DDNPREQTLEM
015 YP_004442824.1     ------------RNSHRAAYEAQRLTRQAGNIVGRIV--GRQPREHTLSM
016 YP_001931967.1     --------------------------------------------------
017 YP_002455786.1     --------------------------------------------------
018 YP_009109694.1     --------------------------------------------------
019 NP_789739.1        --------------------------------------------------
020 NP_955619.1        QDPTHWKRDCPRLKPTIPEPEPEEDALLLDLPADIPHPKNLHRGGGLTSP
021 YP_224289.1        ------------RASARRRYEAQRVARNIGNVGRRLI--GRQTREDTLAL
022 YP_009182100.1     --------------------------------------------------
```

```
023 NP_705927.1          --------------------------------------------------
024 YP_009508577.1       --------------------------MTTPPLLQLPVEVKKTELNGFWDT
025 YP_004442830.1       ------------NRRQRAKHEMQNIVAKPSAIFKRII--GGSQSEKTLEM
026 NP_758887.1          YPLGQWQTQELPG----DAI-----DPNGPSNARDGRPSRENAVRLHEER
027 YP_009508406.1       --------------------------------------------------
028 YP_610965.1          ------------RANARLRHDAQR---QLQRVRRTLTGRQ--GYRRTLEQ
029 YP_001931961.1       --------------------------------------------------
030 YP_241114.1          --------------------------------------------------
031 YP_006732334.1       --------------------------------------------------
032 YP_009041481.1       ------------RRMQRARHEAQRILPPILGGRRRR----QIQGPRRIEE
033 YP_001036293.1       ------------RLGARRRYNLRQGLNKFKNVLTLG----T-YYAPTLEQ
034 YP_605811.1          ------------RTAARAAYEAQRIARRTGNIIGRIV--GRQPREHTLAM
035 NP_955577.1          -------------------------------EPRITLKIGGQPVTFLVDT
036 YP_003208050.1       TSSTLPPVRGSNTAAAPIQEDQVRDYRRWQRLRWNMRNRWNTTFRRTLES
037 NP_789740.1          ----------LQG----KAR-----EFSSEQTRANSPTRRELQVWGRDNN
038 YP_004442839.1       ------------RNAARAAYEAQRVARQTGNIIGRIV--GRQPREHTLAM
039 YP_009507791.1       --------------------------------------------------
040 YP_004581513.1       --------------------------------------------------
041 NP_043933.1          --------------------------------------------------
042 YP_009507248.1       --------------------------------------------------
043 YP_002117531.1       ------------RRGQRRRYEAQRVARNLQNIGRTIV--GARPREHTLAL
044 YP_004347415.1       --------------------------------------------------
045 NP_056891.1          GRSSITMKGLQVYPGVIDNDYTGEIKIMAKAVNNIVTVSQGNRIAQLILL
046 YP_009513242.1       -------------------------MDPLQLLQPLEAEIKGTKLKAHWDS
047 YP_009352866.1       ------------RRMQRARHEAQRRLPPILGGRRR-----DRQGPQRIEE
048 YP_009165750.1       --------------------------------------------------
049 NP_040840.1          ----------------------------------MPSLWKKRTYAKGLP-
050 YP_004222728.1       --------------------------------------------------
051 YP_009506264.1       ------------RRAQRARYEAQRVARSLGNIGRTIV--GRQPREHTLAL
052 YP_567050.1          --------------------------------------------------
053 YP_009508571.1       -------------------------MNPLQLLQPLQAEVKGNKLIAHWDS
054 NP_056803.1          -------------------------MNPLQLLQPLPAEVKGTKLLAHWDS
055 NP_955564.1          --------------------------------------------------
056 YP_009508411.1       ------------RLGQRRRHNLQQGMKKLSAKLFGG----P-YSR-TLEQ
057 YP_009140788.1       ------------RRNQRRRFIAEQALRRTIGKIRG------RRYNQTLEQ
058 YP_009508410.1       --------------------------------------------------
059 NP_040333.1          -------------------------------EPRITLRVGGQPVTFLVDT
060 YP_008992013.1       ASTSSSNLSTPLTLSSTLSLPKSCTSHTANKIENLVEYSYIPESAQISES
061 NP_041734.1          ------------RRGQRRIFNLRRRARRLRRSMMG------SRYQETLEQ
062 NP_056895.1          LKEGHWARDCPTKTTGPPPGPCPICKRSFPLETRLSNPQIKKLIEGGLSA
063 YP_009109689.1       -------------------------------EPRLTLSVGGHPTTFLVDT
064 YP_002519387.1       --------------------------------------------------
065 YP_595725.1          ------------RRAQHAKHEAQRIAGQALAFLR-VT--SDDPREKTLEM
066 YP_009508551.1       -------------------------MNPLQLLQPLPAEIKGTKLLAHWDS
067 YP_001497148.1       -------------------------------EPRLTLSVGGHPTTFLVDT
068 NP_041186.1          --------------------------------------------------
069 NP_663784.1          GPTGKEASQLPRDPSPSGADTNSTSGRSSSGTVGEIYAAREKAEGAEGET
070 YP_009508443.1       ------------RLGARRRYNMGLRAQKLIKTLSLG----T-FYNPTLEQ
071 YP_003987465.1       ------------RWTQRACYEAQEAGRRLANIGRTIL--GMQPREHTLAM
072 NP_861410.1          --------------------------------------------------
073 YP_009345071.1       ------------RTGQRRRYNWARQARRLRNRIRR-----SPRQPETLET
074 YP_009508582.1       --------------------------MTAPPLLQLPVEVKKTELNGFWDT
075 NP_569141.1          --------------------------------------------------
076 YP_002916057.1       --------------------------------------------------
077 NP_777384.2          LKEGHWARDCPTKTTGPPPGPCPICKRSFPLETRLSNPQIKKLIEGGLSA
078 YP_233110.1          ------------RSAARVAYEAQRIARRTGNILGRIV--GRQPREHTLAM
079 NP_542258.1          --------------------------------------------------
080 YP_009130664.1       ------------RQRERRLHNARQVVGRLHRRITGR----SQLHNQTLEQ
081 YP_009508566.1       --------------------------MYQP-QHQLQVEIHDQKLIGYWDT
082 YP_006607892.1       --------------------------------------------------
083 YP_009506270.1       ------------RNTRRAAYEAQRMARQAGNIVGRIV--GRQPREHTLSM
084 NP_042513.1          --------------------------------------------------
085 YP_006907834.1       --------------------------------------------------
086 YP_004442827.1       ------------SRGQRIKHEMQNIVARPTALFKRIT--GGSASEKTLEM
087 YP_009408594.1       ------------------------------------------------LP
088 YP_009508561.1       --------------------------MAQT---NIPVEIKGTQLNGFWDT
089 NP_056907.1          KDPSHWKRDCPQFKPPPTEEEPLLLDLPSDAIATEEKKL-----------
090 YP_002308474.1       --------------------------------------------------
091 YP_007761644.1       --------------------------------------------------
092 NP_040550.1          --------------------------------------------------
093 NP_758808.1          ------------RNMQRVRHTAERAARRIFPGR----------FNRTLES
094 YP_009345075.1       ------------RRARRTMWNASR---QLTGRRN---------NLRTLEQ
095 NP_068729.1          --------------------------------------------------
```

```
096 NP_047255.1         -------------------------------EPRITLKIGGQPVTFLVDT
097 YP_006273075.1      ------------RRNQRRRFVAEQAVRRTMGRMRG------RRYNQTLEQ
098 YP_009508888.1      -------------------------MSLQHLLQPLEAEVKGTKLKAHWDS
099 NP_395469.1         --------------------------------------------------
100 YP_009508546.1      -------------------------MNPLQLLQPLPAEIKGTKLLAHWDS
101 NP_056762.1         -------------MSLRPFTGTSRTITQDSTSESNIKKGKNSTKRELIEE
102 YP_009508556.1      -------------------------MDPLQLLQPLEAEIKGTKLKAHWDS
103 NP_569153.1         ------------RQWQRRRHNMG----RRWNQLIG-----R-PYNQTLEQ
104 NP_043924.1         AEEEAWRERQTSQPSQPSQLRPFVAAGNGHREDQWRPLTVTELPAAVTAV
105 YP_009243641.1      --------------------------------------------------
106 YP_004300274.1      --------------------------------------------------
107 NP_056848.1         --------------------------------------------------
108 YP_006495799.1      ------------RQTARLRHNVRRRLRSLSPGGFR----------NTLEQ
109 YP_009508409.1      --------------------------------------------------
110 NP_687035.1         WPLGRSETKKFCA----IQRRHSWSGTNSPPNGNSLRSSKEAPPAVCREG
111 NP_045937.2         TTQKVEQTDIGEASSSDASAQKPISTDNNPDLKVDGVLTRSQHTTVPSNI
112 NP_612577.1         --------------------------------------------------
113 NP_619548.1         --------------------------------------------------
114 NP_056880.1         GRSSNYKKGLEVLPGVIDSDFQGEIKVMVKAAKNAVIIHKGERIAQLLLL
115 YP_009121747.1      ------------RTNARRMYNLNQAVRSRLGRR----------YRRNLET
116 YP_009113237.2      ------------RRNARRRYNVQNAARRITARLSN------RPYTRTLEQ
117 YP_009508408.1      ------------RLGARRRYNMGLKAQKLVKTLSRG----K-FYNPTLEQ
118 NP_955579.1         --------------------------------------------------
119 YP_009229919.1      ------------RNIQRVKHQAGR---KLKQLRG--------QHSRSLES
120 YP_004442833.1      ------------RNGQRALYEAQNIVARPNAIFKRIF--GGSETEKTLEM
121 NP_862833.2         --------------------------------------------------
122 NP_040563.1         ------------------------TAKMLPLWQTWPPSKKLQVKKREVLL
123 YP_443922.1         PFQGMYDDNTPWEXRCPPPPEAEVQRYSVYTEPTVSCQIEGVNHRFLVDT
124 NP_057933.2         -------------------------------EPRITLKVGGQPVTFLVDT
125 YP_009116631.1      ------------RRSARMRHQAQQRLRRTFGRDFR----------NTIER
126 YP_009513249.1      ----------------------------MDLLKPLTVERKGVKIKGYWDS
127 YP_223871.1         -------------------------------EPRLKVKVGGQIIDFLVDT
128 NP_127504.1         -MTSPSDYQSNSSLATTYSNAPKLSKALSNKYDYLYEVDILKENQKISDT
129 NP_044929.1         ----------------------------MPALRPLQVEIKGNHLKGYWDS
130 NP_057860.1         QDPTHWKRDCPRLKPTIPEPEPEEDALLLDLPADIPHPKNLHRGGGLTSP
131 NP_040973.1         --------------------------------------------------
132 NP_777317.1         ------------RSIRRTAYEARRRAGMAGRTMGRML--GQQPSNRAAPS
133 NP_054716.1         ----------------------------MQALQPLQVQIKGNSLKGFYDT
134 YP_001856242.1      --------------------------------------------------
135 NP_954565.2         GRSSITMKGLQVYPGVIDNDYTGEIKIMAKAVNNIVTVSQGNRIAQLILL
136 NP_569150.1         ------------RSSRRAAYEAQRIARQTGNIIGRVV--GRQPREHTLSM
137 NP_659397.1         --------------------------------------------------
138 YP_009513211.1      --------------------------------------------------
139 YP_233107.1         ------------RRSARARYEAQRMGRNLVNIGRTII--GRRPREHTLAL
140 YP_009109692.1      -------------------------------EPRLTLSVGGHPTTFLVDT
141 NP_057849.4         ANFLREDLAFLQG----KAR-----EFSSEQTRANSPTRRELQVWGRDNN
142 NP_049560.1         --------------------------------------------------
143 YP_009506267.1      ------------RLGARRRYNMSQAAQKLAKTLTFG----K-YSQPTLEQ
144 NP_039820.1         ------------RNDQRRRHIWAGRGRRLLSIMP-----GVSSSERTLEM
145 NP_056790.1         -------------------------------EPRVTLTVEGTPIEFLVDT
146 YP_002321513.1      ------------RRMRRMGYETIRAIRQPLNRIFAR-------RREALLS
147 NP_955591.1         --------------------------------------------------
148 YP_001956722.2      -------------------------MDPLQLLQPLEAEIKGTKLKAHWDS
149 NP_056902.1         -----MQKCSKKREARGSREAPETNFPDTTEESAQQICCTRDSSDSKSVP
150 NP_056886.1         GSPGHYQAQCPKKRKSGNSRERCQLCNGMGHNAKQCRKRDGNQGQRPGKG
151 NP_041261.1         --------------------------------------------------
```

```
001 YP_004442836.1      VVNPNSELSRSLAHRARTVPGEVLYMTQRDSP-VNRVYRNRTEERM-LVT
002 NP_056728.1         --------------------------------------------------
003 YP_009508537.1      QADITCVPKDLLQEEEPVRQQNVTTIHGTQEEDVYYVNLKIDGRRINTEV
004 YP_004732983.2      RLDPEAELQLSRRRRANLVPAEVLYSLNYNEP-QNRVYQHYEEVRS--HV
005 YP_009508407.1      --------------------------------------------------
006 NP_040939.1         KERHKGKTAGKREEGDTCGAVRSSYG----ITSAPPMVQVRIGSQQRNLL
007 YP_003864102.1      --------------------------------------------------
008 YP_009002585.1      EVDPQADLQMSMQERARLVPAEVLYRSRRDTI-HHRVYSHRSEEAI-LCT
009 YP_003284237.1      LMNPEVELQRSMQERARTVPAEVLYMTRRDDI-HHRVYHHRSEERM-LVL
010 YP_009268869.1      ----SNITAGKQQEGATCGAVRAPYV----VTEAPPKIDIKVGTNWKKVL
011 YP_009506251.1      EIDPQVTLRLSMQERARLVPAEVLYRSRRDTV-HHRVYTHRSEESI-LCV
012 NP_955611.1         --------------------------------------------------
013 YP_008567619.1      SLDPQKQLMRSMTQRASIVPAEVLYHSRKDDV-NHKVYIHRSEEAL-SVT
014 NP_149413.1         VMSPEAELERSLKKRARAFPAEVIYSPRRDDI-HHRVFVGSSSQDV-MIV
```

```
015 YP_004442824.1    VVNPDSELSRSLAHRARTVPAEVLYMTQRDSP-VNRVYRNRTEERM-LVT
016 YP_001931967.1    --------------------------------------------------
017 YP_002455786.1    --------------------------------------------------
018 YP_009109694.1    --------------------------------------------------
019 NP_789739.1       --------------------------------------------------
020 NP_955619.1       PTLQQVLPNQDPTSILPVIPLDPARRPVIKAQIDTQTSHPKTIEA----L
021 YP_224289.1       LMDPEVELQRSMRERARTVPAEVLYMSRRDDV-HHRVYHFRSEERM-MVS
022 YP_009182100.1    --------------------------------------------------
023 NP_705927.1       --------------------------------------------------
024 YP_009508577.1    GAQITCIPEAFLKEEIPIGEAQIKTLHGTKLQSVYYLKFKVLGRKVEAEV
025 YP_004442830.1    VLDPEAELEKSMKRRAKAFPAEVVYAPRSDNK-LHKVFQGRTSQDI-MVV
026 NP_758887.1       ATAESGKQTGKEREGSIRGS----LQLPQFSLWNRPTTVVEIEGQKVEAL
027 YP_009508406.1    --------------------------------------------------
028 YP_610965.1       HIDPNEQLQLSQKKKAAIVPAEVLYDDGWHST-VHKVYQHYDEQRVLVLE
029 YP_001931961.1    --------------------------------------------------
030 YP_241114.1       --------------------------------------------------
031 YP_006732334.1    --------------------------------------------------
032 YP_009041481.1    ELDPNMELSASSRRRATMVPAEVLYSTGDTRV-RNRVYEHYSEQRG--LI
033 YP_001036293.1    QVDPERTLQLSAIKRASMIPAEVLYHARPDTV-NHKVYMHWSEEAD-LIV
034 YP_605811.1       VVDPNSELERSLAHRARTIPAEVLYMTQRGEP-TNRVYRNRTEERM-LVT
035 NP_955577.1       GAQHSVLTRPDGPLSDRTALVQGATG-SKNYRWTTDRRVQLATGKVTHSF
036 YP_003208050.1    ELDPEAEIQLSQRRRANMVPAEVLYEEGWSSTRLHRVYQHYSEERILST-
037 NP_789740.1       SPSEAG----ADRQGTVS------FNFPQVTLWQRPLVTIKIGGQLKEAL
038 YP_004442839.1    IVDPNSELERSLAHRARTIPAEVLYMTQRGEP-TNRVYRNRTEERM-LVT
039 YP_009507791.1    --------------------------------------------------
040 YP_004581513.1    --------------------------------------------------
041 NP_043933.1       --------------------------------------------------
042 YP_009507248.1    -MIPARSSKLVRIPINATDGEVLVEEQMFCNCIVHECVTMVKDGRGYVEL
043 YP_002117531.1    LMDPEAELTRSMQERARTVPAEVLYMTRRDDI-HHRVYYHRSEERL-LII
044 YP_004347415.1    --------------------------------------------------
045 NP_056891.1       PLIETDNKVQQPYRGQGSF-GSSDIYWVQPITCQKPSLTLWLDDKMFTGL
046 YP_009513242.1    GATITCIPEAFLEDEQPIQTMLIKTIHGEKQQNVYYLTFKIQGRKVEAEV
047 YP_009352866.1    EINPNRELSASSRRRASMVPAEVLYNSGDTNV-RNRVYEHYSEQRG--LI
048 YP_009165750.1    --------------------------------------------------
049 NP_040840.1       ----AEETAGKQQEGATCGAVRAPYV----VTEAPPKIEIKVGTRWKKLL
050 YP_004222728.1    --------------------------------------------------
051 YP_009506264.1    LMDPDVELQRSMSERARTVPAEVLYMTRRDDV-HHRVYHHRSEERM-LVL
052 YP_567050.1       --------------------------------------------------
053 YP_009508571.1    GASITCIPESFLEEETPIKKTIIKTIHGQKEQKVYYLTFKVNGRKVEAEV
054 NP_056803.1       GATITCIPESFLEDEQPIKQTLIKTIHGEKQQNVYYLTFKVKGRKVEAEV
055 NP_955564.1       ------------------------HWVQEISDSRPMLHIYLNGRRFLGL
056 YP_009508411.1    QVDPEKTLQVSMYQRASLVPAEVLYHARPDTI-HHKVYLHWSEEAV-LVI
057 YP_009140788.1    TIDPEEELSQSMQERANLVPAEVLYRTRRDNI-NHQVYNHRSEEAM-LCV
058 YP_009508410.1    --------------------------------------------------
059 NP_040333.1       GAQHSVLTQNPGPLSDKSAWVQGATG-GKRYRWTTDRRVHLATGKVTHSF
060 YP_008992013.1    TYPLISPYHLYKRPNSFTRS-IRTLISTKRPHPKEYIQSSRLDQCALKAT
061 NP_041734.1       EIDPQTTLRLSMQERARLVPAEVLYRSRRDTV-HHRVYTHRSEESV-LCV
062 NP_056895.1       PQTVTPITDPLSEAELECLLSIPLARSRPSVAVYLSGPWLQPSQNQALML
063 YP_009109689.1    GAQHSVLTKANGPLSSRTSWVQGATG-RKMHKWTNRRTVNLGQGMVTHSF
064 YP_002519387.1    --------------------------------------------------
065 YP_595725.1       LMQPDVELTRSMKKRARAFPAEVLYGPRSDDI-HHKVFQGSSSQDI-LLI
066 YP_009508551.1    GATITCIPESFLEDEQPIKKTLIKTIHGEKQQNVYYVTFKVKGRKVEAEV
067 YP_001497148.1    GAQHSVLTKANGPLSSRTSWVQGATG-RKIHKWTNRRTVDLGQGTVTHSF
068 NP_041186.1       --------------------------------------------------
069 NP_663784.1       IQRGDGGLAAPRAERDTSQRGDRGLAAPQFSLWKRPVVTAYIEDQPVEVL
070 YP_009508443.1    QVDPEKTLQLSSVQRASLVPAEILYHARTDTV-NHKVYMHWSEESD-LII
071 YP_003987465.1    VIDPEAELQRSMSERARTVPAEVLYMTRRDNI-HHRVYYFRCEERM-RVL
072 NP_861410.1       --------------------------------------------------
073 YP_009345071.1    QIDPQAELSMSLQRRASIVPAEVLYHSRQDDI-NHRVYQHWSEEAV-NCI
074 YP_009508582.1    GAQITCIPEAFLKEEIPIGEAQIKTLHGTKLQPVYYLKFKILGRKVEAEV
075 NP_569141.1       --------------------------------------------------
076 YP_002916057.1    --------------------------------------------------
077 NP_777384.2       PQTVTPITDPLSEAELECLLSIPLARSRPSVAVYLSGPWLQPSQNQALML
078 YP_233110.1       VVDPNSELERSLAHRARTIPAEVLYMTQRGEP-TNRVYRNRTEERM-LVT
079 NP_542258.1       --------------------------------------------------
080 YP_009130664.1    QIDPQVHLRNSMQERAAIVPAEVLYHSRCDDE-HHRVYVHRSEEAL-LCT
081 YP_009508566.1    GAQITCIPQVYLEQEKPIGKHVIETVNGKTQRDAYYIKLKINGKKIETEV
082 YP_006607892.1    --------------------------------------------------
083 YP_009506270.1    VVDPNSELSRSLAHRARTVPGEVLYMTQRDSP-VNRVYRNRTGERM-LVT
084 NP_042513.1       --------------------------------------------------
085 YP_006907834.1    --------------------------------------------------
086 YP_004442827.1    ILDPEAELEKSMKRRAKAFPAEVVYAPRYDNK-LHKVFQGRSSQDI-MVV
087 YP_009408594.1    PVDSHDLPEIPDDQDSEQFLAILTFTGDPQYLLPDPVYDTSSD-------
```

088 YP_009508561.1 GAQITCIPQSFLLDEQPIGTTDIETIHGKQKQKLYYLKFKVLGRKVEAEV
089 NP_056907.1 -PGGGDVISPQQISMLPVIPLEQQHQPLLDVQVSIAGAPPRPTQA----L
090 YP_002308474.1 --------------------------------------------------
091 YP_007761644.1 --------------------------------------------------
092 NP_040550.1 --------------------------------------------------
093 NP_758808.1 QINPEAEIRLSQQRRAAMVPAEVLYNTSPSTR-NQKVYQHYSEERILCTG
094 YP_009345075.1 EIDPEAQLRLSTQQRAGIVPAEVLYRNGWSER-AHRVYRHYSEERIMLTN
095 NP_068729.1 --------------------------------------------------
096 NP_047255.1 GAQHSVLTRPDGPLSDRTALVQGATG-SKNYRWTTDRRVQLATGKVTHSF
097 YP_006273075.1 IVDPEVELNQSMQERANLVPAEVLYRSRRDNI-NHQIYNHRSEEAM-LCV
098 YP_009508888.1 GTTITCIPTVFLTDEIPIKDVLIKTIHGERRQPAYYLTFKINGRKVQAEV
099 NP_395469.1 --------------------------------------------------
100 YP_009508546.1 GATITCVPESFLEDEQPIKQTLIKTIHGEKQQKVYYLTFKIKGRKVEAEV
101 NP_056762.1 VDVNQEVENFDWKKLSGIKPNKLYEKNWQEKVKLKQQSIVSAYKEEAISV
102 YP_009508556.1 GATITCIPEAFLEDEQPIQTMLIKTIHGEKRQNVYYLTFKIQGRKVEAEV
103 NP_569153.1 VVDPEVALQLSMQERARLVPAEVLYRSRTDDR-HHQVYIHKSEEAI-LCV
104 NP_043924.1 GGAWDPTRETGSARWKKIVKAAEAIGWGTGDVCQVVTAMSPSWADVPPEI
105 YP_009243641.1 --------------------------------------------------
106 YP_004300274.1 --------------------------------------------------
107 NP_056848.1 --------------------------------------------------
108 YP_006495799.1 QLSPNQELALSRRRRANLVPAEVLYTHNNNEV-VNRVYQHYEERSA--HV
109 YP_009508409.1 --------------------------------------------------
110 NP_687035.1 TAPERGERTDKETEGERSG-----GCFLELPLWRRPMKRVIIEGTPVQAL
111 NP_045937.2 TIKKDGTSVQYQHPIRNYPTGEGNLTAQVRNPFRPLELQQLRKDCPALPE
112 NP_612577.1 --------------------------------------------------
113 NP_619548.1 --------------------------------------------------
114 NP_056880.1 PYLKLPNPVIKEERGSEGFGSTSHVHWVQEISDSRPMLHIYLNGRRFLGL
115 YP_009121747.1 ELDPDQQLEVSRQRRAELVPAEVLYGSNSQTV-RHRVYQHYSEQRILCTE
116 YP_009113237.2 EVDPEATLRLSMQERARLVPAEVLYRSRSDDV-NHRVYIHRDEEAV-LCT
117 YP_009508408.1 QVEPEKTLQLSSVQRASLVPAEVLYHSRPDTV-NHKVYMHWSEESD-LII
118 NP_955579.1 --------------------------------------------------
119 YP_009229919.1 RIDPDAQLQISRRRRSDMVPAEVLYEAQRDNP-IHRVYQHYSERRIRVTE
120 YP_004442833.1 VMDVEAELEKSMKRRARAFPAEVVYSPRRDNK-LHKVFQGRASQDL-MVV
121 NP_862833.2 --------------------------------------------------
122 NP_040563.1 CPLWAEEPTTEQFSPEQHEFCDPICTPSYIRLDKQPFIKVFIGGRWVKGL
123 YP_443922.1 GCTYSAIKT-MQPLSMDSIQVVGVSGRPETQLKTQPLLFRWGPSTVRHQF
124 NP_057933.2 GAQHSVLTQNPGPLSDKSAWVQGATG-GKRYRWTTDRKVHLATGKVTHSF
125 YP_009116631.1 QLDPDAELSLSRRRRANLVPAEVLYAHNGQEP-VNRVYEHYSELSA--HV
126 YP_009513249.1 QADITCVPKDLLQGEEPVRQQNVTTIHGTQEGDVYYVNLKIDGRRINTEV
127 YP_223871.1 GATHSVVQKPVGPMSKESVAIIGATGNIRNYPKSEGRLVDLGRGLVTHSF
128 NP_127504.1 YLPLLNPYSAFAKRSVTPWSQIRSLVQSKPRHVKEYVAASKLDQHPVFAT
129 NP_044929.1 GAEITCVPAIYIIEEQPVGKKLITTIHNEKEHDVYYVEMKIEKRKVQCEV
130 NP_057860.1 PTLQQVLPNQDPTSILPVIPLDPARRPVIKAQIDTQTSHPKTIEA----L
131 NP_040973.1 --------------------------------------------------
132 NP_777317.1 IVNPDEELERSLTRRARTIPAEVLYLAQRREV-DNRVYRNMTEERM-LIT
133 NP_054716.1 GAEITCVPAIFLIEEEPIGERTIQTIHGITKEKVYYLTFKIQGRKLAAEV
134 YP_001856242.1 --------------------------------------------------
135 NP_954565.2 PLIETDNKVQQPYRGQGSF-GSSDIYWVQPITCQKPSLTLWLDDKMFTGL
136 NP_569150.1 VVDPNSELSRSLAHRARTVPGEVLYMTQRDSP-VNRIYRNRTEERM-LVT
137 NP_659397.1 --------------------------------------------------
138 YP_009513211.1 ----------MGKMGSKRTVVAGATG-SKVYPWTTKRLLKIGQKQVTHSF
139 YP_233107.1 LMDPEVELRRSMQERARTVPAEVLYMTRRDDI-HHRVYHYRSEEGM-LIT
140 YP_009109692.1 GAQHSVLTKANGPLSSRTSWVQGATG-RKMHKWTNRRTVNLGQGMVTHSF
141 NP_057849.4 SPSEAG----ADRQGTVS------FNFPQVTLWQRPLVTIKIGGQLKEAL
142 NP_049560.1 --------------------------------------------------
143 YP_009506267.1 QVDPEKTLQLSSMQRASLVPAEVLYHSRPDTV-NHKVYMHWSEESD-LVV
144 NP_039820.1 QMNPEVQLQRSMNHRAEAVPAEVLYRTFHGSV-NHRVYSHRSEERM-MVV
145 NP_056790.1 GAEHSVLTQPMGKVGSRRTVVEGATG-SKVYPWTTKRLLKIGHKQVTHSF
146 YP_002321513.1 PEQVSRRLQRSGSRRPEAVPAEVLYTAGQDSA-QTRVYESVAEESI-LVI
147 NP_955591.1 --------------------------------------------------
148 YP_001956722.2 GATITCVPQAFLEEEVPIKNIWIKTIHGEKEQPVYYLTFKIQGRKVEAEV
149 NP_056902.1 RSERNKKGIQCQGEGSSRGSQPGQFVGVTYNLEKRPTTIVLINDTPLNVL
150 NP_056886.1 LSSGPWPGPEPPAVSLAMTMEHKDRPLVRVILTNTGSHPVKQRSVYITAL
151 NP_041261.1 --------------------------------------------------

001 YP_004442836.1 NG-QQDRSFIYPESFEELVNAGFEYIHLGVLQVRLQIMHRTYAGTMALVV
002 NP_056728.1 --------------------------------------------------
003 YP_009508537.1 IGTALDYAIITPGDAPWILKKPLELT------------------------
004 YP_004732983.2 IDRQQDFRFIEEQSYRTLVQEGMQHIHPGMLMVRIQVLHRVDAGISAMIV
005 YP_009508407.1 --------------------------------------------------
006 NP_040939.1 FDTGADRTIVRWHEGSGNPAGRIKLQGIGGIV------------------

```
007 YP_003864102.1    --------------------------------------------------
008 YP_009002585.1    DGNQVDRTFIQPESMEALQRTGISFIHIGIMQVRLQILHRQNEGTMALVV
009 YP_003284237.1    RNDQQDRTFIQQESYEALAQAGLEYIHLGVLQVRFQILHRRYAGTVAFLV
010 YP_009268869.1    VDTGADRTIVRYHDNSGIPTGRIKLQGIGGII------------------
011 YP_009506251.1    GGNQVDRTFIQPSSLERLQRTGMSFIHIGILQVRIQILHRQEEGTMALVV
012 NP_955611.1       --------------------------------------------------
013 YP_008567619.1    EGNQVDRTFIQEESHKKLVKSGIKFIHLGVLQVRLQILHRKEEGTLALVV
014 NP_149413.1       DDNQIDMTYIKEESFEKLEQAGFRYIHLGAMAIRIQPLHAAWSGKLAFIV
015 YP_004442824.1    NG-QQDRTFIYPESFEELVNAGFEYIHLGVLQVRLQIMHRTYAGTIALVV
016 YP_001931967.1    --------------------------------------------------
017 YP_002455786.1    --------------------------------------------------
018 YP_009109694.1    --------------------------------------------------
019 NP_789739.1       --------------------------------------------------
020 NP_955619.1       LDTGADMTVLPIALFSSNTPLKNTSVLGAGGQ------------------
021 YP_224289.1       DADQQDRTFISEESFERLQQAGLTYIHLGVLQVRFQILHRTFAGTMALLV
022 YP_009182100.1    --------------------------------------------------
023 NP_705927.1       --------------------------------------------------
024 YP_009508577.1    TTSPFDYVIISPSDIPWYKPQPLELT------------------------
025 YP_004442830.1    DDNQLDMTFIREETFEQLERAGLRYIHIGTLQVRIQPLHQRWSGTMAFIA
026 NP_758887.1       LDTGADDTVIKDLDLKG----NWKPQIIGGIG------------------
027 YP_009508406.1    --------------------------------------------------
028 YP_610965.1       -GQQVELPFINEQSYRQLLSSGFQHIHVGMVLVRVHALHSRHAGTMALLC
029 YP_001931961.1    --------------------------------------------------
030 YP_241114.1       --------------------------------------------------
031 YP_006732334.1    --------------------------------------------------
032 YP_009041481.1    VNNQEDFRFIQESSYNTLRNSGYEHMHVGMMMMRVLTLHRTYTGVTALVT
033 YP_001036293.1    NN-QHDSTFIRPESYLELRKANMQFIHIGLMQVRIQVLHRHEAGTMALIV
034 YP_605811.1       HG-QQDRTFILPESYEELREAGFEYIHLGVLQVRIQIMHRSYAGTMALVV
035 NP_955577.1       LYVPECPYPLLGRDLLTKLKAQIHFTGEGANV------------------
036 YP_003208050.1    -GEQVELPFVTRRSYERLQRTGYQQIHLGLVLIRVHTLHRRDAGVKALVV
037 NP_789740.1       LDTGADDTVLEEMSLPG----RWKPKMIGGIG------------------
038 YP_004442839.1    QG-QQDRSFILQESYEALLEAGFEYIHLGVLQVRIQIMHRSYAGTMALVV
039 YP_009507791.1    ------------------------MQGVGGTN------------------
040 YP_004581513.1    --------------------------------------------------
041 NP_043933.1       --------------------------------------------------
042 YP_009507248.1    ENPTPNDVIFYLDQPASAELFNIKCTQVEQSQ------------------
043 YP_002117531.1    GNTQVDRTFIPQQSYEALSKEHMEYIHLGVIQVRLQILHRKFAGTLALIS
044 YP_004347415.1    --------------------------------------------------
045 NP_056891.1       IDTGADVTIIKLEDWPPNWPITDTLTNLRGIG------------------
046 YP_009513242.1    LASPYDYILLNPSDVPWLMKKPLQLT------------------------
047 YP_009352866.1    VNSQEDFRYIQESSYNTLRNSGYEHIHIGMMMVRVLTLHRTHAGVKALVT
048 YP_009165750.1    --------------------------------------------------
049 NP_040840.1       VDTGADKTIVTSHDMSGIPKGRIILQGIGGII------------------
050 YP_004222728.1    --------------------------------------------------
051 YP_009506264.1    ANDQQDRTFIVRESYEALERAGFEYVHLEVMQVRFQILHRRFAGTLAFIV
052 YP_567050.1       --------------------------------------------------
053 YP_009508571.1    IASPYDYILLSPMDVPWLVQKPLQLT------------------------
054 NP_056803.1       IASPYEYILLSPTDVPWLTQQPLQLT------------------------
055 NP_955564.1       LDTGADKTCIAGRDWPANWPIHQTESSLQGLG------------------
056 YP_009508411.1    QG-QYDSTFIKTESYEALQRAGMQFIHIGLMQVRIQVLHRLEAGTMALVA
057 YP_009140788.1    G-EQQDRSFIQPDSFQELQRSGMQFIHLGVLQVRIQILHRADEGTMALVV
058 YP_009508410.1    --------------------------------------------------
059 NP_040333.1       LHVPDCPYPLLGRDLLTKLKAQIHFEGSGAQV------------------
060 YP_008992013.1    SAEQYVTLEIPSELISNWKREGYTHLHLGGIRLILTLHGRKGLPVTARVA
061 NP_041734.1       GGSQVDRAFIQPESLEQLQRTGMSFIHIGILQVRIQILHRQEEGTMALVV
062 NP_056895.1       VDTGAENTVLPQNWLVRDYPRIPAAVLGAGGV------------------
063 YP_009109689.1    LVVPECPYPLLGRDLLTKLGAQIHFSEAGAQV------------------
064 YP_002519387.1    --------------------------------------------------
065 YP_595725.1       DDNQLDMTFIKEETFEQLEQAGLRYIHPGILAVRIQPLHPDWSGKLVFIV
066 YP_009508551.1    IASPYEYILLSPTDVPWLTQQPLQLT------------------------
067 YP_001497148.1    LVVPECPYPLLGRDLLTKLGAQIHFSETGAQV------------------
068 NP_041186.1       --------------------------------------------------
069 NP_663784.1       LDTGADDSIVAGIELGDN----YTPKIVGGIG------------------
070 YP_009508443.1    NG-QHDSTFIRPESFQVLQRSGMQFIHIGLMQVRIQVLHRLEAGTMAMVV
071 YP_003987465.1    DSDQQDRTFITPEAFEHLKEAGFEYIHLGILQVRFQILHRRYAGTMAFIA
072 NP_861410.1       --------------------------------------------------
073 YP_009345071.1    DGQQIDRTLVTMESYSTLNRAGFRYIHIGVIQVRLQILHRREMGTLAYVV
074 YP_009508582.1    TTSPFDYVIISPSDIPWYKPQPLELT------------------------
075 NP_569141.1       --------------------------------------------------
076 YP_002916057.1    --------------------------------------------------
077 NP_777384.2       VDTGAENTVLPQNWLVRDYPRIPAAVLGAGGV------------------
078 YP_233110.1       HG-QQDRTFILPESYEELREAGFEYIHLGVLQVRIQIMHRTYDGTMALVV
079 NP_542258.1       --------------------------------------------------
```

```
080 YP_009130664.1   DNQQQDRIFIQEESFKQLQRSRMQFIHIGVIQLRIQALHRHDEGTLALVV
081 YP_009508566.1   IPSPFSYALITPNDIPWFKPGGIELT------------------------
082 YP_006607892.1   --------------------------------------------------
083 YP_009506270.1   NG-QQDRSFIYPESFEELVAAGFEYIHLGVLQVRLQIMHRTYAGTMALVV
084 NP_042513.1      --------------------------------------------------
085 YP_006907834.1   --------------------------------------------------
086 YP_004442827.1   DDNQLDMTFIKEETFEQLERAGLRYIHIGALQVRIQPLHQKWYGTMAFLA
087 YP_009408594.1   --SDSDSDTESTTSDGPLPDLGPPDHVWATDPRPSYAVWEENRSGNCPKA
088 YP_009508561.1   TSSTLAYVILAPIDIPWYKPTPLELT------------------------
089 NP_056907.1      LDTGADLTVLPQALAPESVSVSDTTVLGAGGQ------------------
090 YP_002308474.1   --------------------------------------------------
091 YP_007761644.1   --------------------------------------------------
092 NP_040550.1      --------------------------------------------------
093 NP_758808.1      QNQQLNLPFINESSYRALRESGQQHLHIGLIMIRVHPLHRRNAGTTALIV
094 YP_009345075.1   PNQQRELRVITDQSYQKLREEGKQHVHCGLVMIRVHALHRKQAGVMVLVV
095 NP_068729.1      --------------------------------------------------
096 NP_047255.1      LYVPECPYPLLGRDLLTKLKAQIHFTGEGANV------------------
097 YP_006273075.1   G-EQQDRMFIQPNSFQELQRSGMQFIHLGVLQVRLQILHRADGGTMALVV
098 YP_009508888.1   IASPYDYILLCPADVPWLQQQPLQLT------------------------
099 NP_395469.1      --------------------------------------------------
100 YP_009508546.1   IASPYEYILLSPTDVPWLTQQPLQLT------------------------
101 NP_056762.1      THNAYTTTLFPQEVIKNVKNQGKLYYHIGMMAIGVKGLHRRKIGTKVMIM
102 YP_009508556.1   LASPYDYILLNPSDVPWLMKKPLQLT------------------------
103 NP_569153.1      DGDQVDRLLIQPESAEQLSRSGMSFIHMGIVQVRIQILHRQHEGTTALVV
104 NP_043924.1      RNRVATEKEIKAWLMKQGPGGGQGLLEFTKLRQGPTENPSNYLEKALELY
105 YP_009243641.1   --------------------------------------------------
106 YP_004300274.1   --------------------------------------------------
107 NP_056848.1      --------------------------------------------------
108 YP_006495799.1   VDRQMDLRFIEEESYRRLVDEGLQFIHLGMGMVRIHMLHRNSAGIEAMIV
109 YP_009508409.1   --------------------------------------------------
110 NP_687035.1      LDTGADDTIIQEKDLHFPPHKPWRSKVVGGIG------------------
111 NP_045937.2      GIPQLAEWLTQTMAIYNCDEADVEQLARVIFPTPVRQIAGVINGHAAANT
112 NP_612577.1      --------------------------------------------------
113 NP_619548.1      --------------------------------------------------
114 NP_056880.1      LDTGADKTCIAGRDWPANWPIHQTESSLQGLG------------------
115 YP_009121747.1   ENQ-TDLRLCNRQSYQSLRESGMQHIHLGMFMIRILTMHRRGAGTTALIV
116 YP_009113237.2   GDAQVDRALIQPQSAERLLQSGYSFIHIGIIQARIQILHRSQEGTMALVL
117 YP_009508408.1   NG-QHDSTFIRPESYQLLQRSGMQFIHIGLMQVRVQVLHRLEAGTMAMIV
118 NP_955579.1      --------------------------------------------------
119 YP_009229919.1   EGEQEDLRFITEPSYGTLRREGFQHIHLGLMMVRVHGLHARHAGTQVLVV
120 YP_004442833.1   DDNQMDMTFIKEETFEQLERAGLRYIHIGALQVRIQPLHQRWSGKMAFLA
121 NP_862833.2      --------------------------------------------------
122 NP_040563.1      VDTGADEVVLKNIHWDRIKGYPGTPIKQIGVN------------------
123 YP_443922.1      LYCPNCPINLLGRDLLCRLGCFINLTEDGVEVSINSPKSGRVLLLPSLPV
124 NP_057933.2      LHVPDCPYPLLGRDLLTKLKAQIHFEGSGAQV------------------
125 YP_009116631.1   VDRQQDFRFIEEASYQRLTREGMQFIHVGMAMVRIQMLHRTDAGISALVV
126 YP_009513249.1   IGTTLDYAIITPGDVPWILKKPLELT------------------------
127 YP_223871.1      LVIPECPDPLLGRDLLQKLRATISFTGGGPPE------------------
128 NP_127504.1      GEEQFVTLHIPEEFASHWKSHQFTHIHFGAVKIALTYHGRKGQPVVARLA
129 NP_044929.1      IATALDYVLVAPVDIPWYKPGPLELT------------------------
130 NP_057860.1      LDTGADMTVLPIALFSSNTPLKNTSVLGAGGQ------------------
131 NP_040973.1      --------------------------------------------------
132 NP_777317.1      DG-QQNRTFIRPESYSTLVASGYEYIHLGVLQVRIQTMHRAFAGTMALVV
133 NP_054716.1      IGTQLDYVIIAPSDIPWYKKYELELT------------------------
134 YP_001856242.1   --------------------------------------------------
135 NP_954565.2      IDTGADVTIIKLEDWPPNWPITDTLTNLRGIG------------------
136 NP_569150.1      NG-QQDRSFIYQESFEELASAGFEYIHLGVLQVRIQIMHRTYAGTMALIV
137 NP_659397.1      --------------------------------------------------
138 YP_009513211.1   LVIPECPAPLLGRDLLTKLKAQIQFSTEGPQV------------------
139 YP_233107.1      GSDQQDRTFITEESYEHLAQAEIEYIHLGILQVRFQILHRRYAGTMALLV
140 YP_009109692.1   LVVPECPYPLLGRDLLTKLGAQIHFSEAGAQV------------------
141 NP_057849.4      LDTGADDTVLEEMSLPG----RWKPKMIGGIG------------------
142 NP_049560.1      --------------------------------------------------
143 YP_009506267.1   NR-QYDSTFIRPESYTTLMKSGMQFIHIGLMQVRIQILHRAEAGTLAMIV
144 NP_039820.1      NGSQVDRSFIQESSFEVLSRTGIEFIHIGVMLVRIQILHRKFAGTMALIV
145 NP_056790.1      LVIPECPAPLLGRDLLTKLKAQIQFSAEGPQV------------------
146 YP_002321513.1   GETQRDLLFMEPQSLQEIRRQNFRHIHIGMVIIKIRPLHTDARGTSALLV
147 NP_955591.1      --------------------------------------------------
148 YP_001956722.2   ISSPYDYILVSPSDIPWLMKKPLQLT------------------------
149 NP_056902.1      LDTGADTSVLTTAHYNRLKYRGRKYQGTGIIG------------------
150 NP_056886.1      LDSGADITIISEEDWPTDWPVMEAANPQIHGI------------------
151 NP_041261.1      --------------------------------------------------
```

```
001 YP_004442836.1    FRDTRWTQEGD-EDRSIIAAMEADLSQG-HQLIYVIPDIMMTIKDFYQHV
002 NP_056728.1       --------------------------------------------------
003 YP_009508537.1    -------------------------------------IKLDLEEQQRTLL
004 YP_004732983.2    FRDTRWNDER-----QIISAMTVDMARG-AQLVYAIPDLMMSIHDFYHHL
005 YP_009508407.1    --------------------------------------------------
006 NP_040939.1       ------------------------------EGEKWNNVELEYKGE-----
007 YP_003864102.1    --------------------------------------------------
008 YP_009002585.1    FRDTRWNGDQ-----SIFATMEVDLTEG-RQIIYVVPDTMMTIGDFARNI
009 YP_003284237.1    FRDTRWNE-DD---RSIIAAMEVDLAEG-NQLVYVIPDIMMTIKDFYRHI
010 YP_009268869.1    ------------------------------EGEKWDKVVIQYKEK-----
011 YP_009506251.1    FRDNRWAGDQ-----AIFAQMEVDLTKG-SQLVFVVPDTMMTIGDFARNV
012 NP_955611.1       --------------------------------------------------
013 YP_008567619.1    FRDNRWKGDK-----GIIATMEVDLTKG-CQMVYIIPDIMMTVNDFYHNI
014 NP_149413.1       LRDVRNSP------PTTLGAMEVDLSKG-AQLIYVLPNFMCTIKDFYHGI
015 YP_004442824.1    FRDTRWTQEGE-EDRSIIAAMEADLSQG-HQLIYVIPDIMMTIRDFYQHV
016 YP_001931967.1    --------------------------------------------------
017 YP_002455786.1    --------------------------------------------------
018 YP_009109694.1    -----------------------------------------TVSLQDEHR
019 NP_789739.1       --------------------------------------------------
020 NP_955619.1       ------------------------------TQDHFKLTSLPVLIRLPFRT
021 YP_224289.1       FRDTRWTA-DD---RSIISAMEVDLAEG-NQLIYVIPNIMMTIGDFYRHI
022 YP_009182100.1    --------------------------------------------------
023 NP_705927.1       --------------------------------------------------
024 YP_009508577.1    -------------------------------------VKLPVQDFKKELI
025 YP_004442830.1    FQDTRSQP------ATVIAAMEVDLSKG-AQMIYIIPDFMTTIGDFYQGI
026 NP_758887.1       ------------------------------GSINVKQFFNCKVTIAG---
027 YP_009508406.1    --------------------------------------------------
028 YP_610965.1       LRDTRWKSSR-----GIIGSMEVDLTAG-SQLVYVVPDILLSVQDFYNHI
029 YP_001931961.1    --------------------------------------------------
030 YP_241114.1       --------------------------------------------------
031 YP_006732334.1    --------------------------------------------------
032 YP_009041481.1    FRDTRWHGEL-----SILGQMELDLSEG-SQLVFTTPNLLCSIHDFYHHI
033 YP_001036293.1    FRDCSWKGRR-----STIARMEVDLTKG-SQLVYIAPNITKKLTDFYNNI
034 YP_605811.1       FRDTRWTQENH-QGRSIIATMEADLSQG-HQLIYVIPDIMMTIRDFYQHI
035 NP_955577.1       ------------------------------VGPKGLPLQVLTLQLEEEYR
036 YP_003208050.1    FRDTRWRDDR-----SIIGSMEIDLSIG-TQLAYIAPDMMMSIHDFYNHV
037 NP_789740.1       ------------------------------GFIKVRQYDQILIEICG---
038 YP_004442839.1    FRDTRWTQEDA-EDRSIIAAMEADLSQG-HQLIYVIPDIMMTIRDFYQHI
039 YP_009507791.1    ------------------------------RGEKYINVHFEAYFDKEVRC
040 YP_004581513.1    --------------------------------------------------
041 NP_043933.1       --------------------------------------------------
042 YP_009507248.1    -----------------------------------------RVDDVLSRL
043 YP_002117531.1    FRDTRWAGDDD---RQLISVMEVDLSEG-CQLIYVIPDMMLTLKDFYRHI
044 YP_004347415.1    --------------------------------------------------
045 NP_056891.1       ------------------------------QSNNPKQSSKYLTWRDKENN
046 YP_009513242.1    -------------------------------------VLVPLQDYQERLL
047 YP_009352866.1    FRDTRWQGEL-----SILGQMELDLSEG-SQPVFTAPNLMCSIHDFFHHI
048 YP_009165750.1    --------------------------------------------------
049 NP_040840.1       ------------------------------EGEKWEQVHLQYKDK-----
050 YP_004222728.1    --------------------------------------------------
051 YP_009506264.1    FRDTRWHE-DD---TSIIAAMEVDLAEG-NQLVYVIPDIMMTIKDFYRHI
052 YP_567050.1       --------------------------------------------------
053 YP_009508571.1    -------------------------------------ILVPLQDYQSRIL
054 NP_056803.1       -------------------------------------ILVPLQEYQDRIL
055 NP_955564.1       ------------------------------MACGVARSSQPLRWQHEDK-
056 YP_009508411.1    FRDCSWKGRR-----SIIAKMEIDLSKG-SQLVYVAPDITKRVADFYRNI
057 YP_009140788.1    FRDNRWQGDQ-----SIFAIAEVDLTRG-SQIVYVIPDTMMTIGDFYRNV
058 YP_009508410.1    --------------------------------------------------
059 NP_040333.1       ------------------------------VGPMGQPLQVLTLNIEDEYR
060 YP_008992013.1    LLDTRFKQFQD----AVIGTVLTTLHAG-SVLLTFYPNFNLSLQDPNLPT
061 NP_041734.1       FRDNRWSGDQ-----SIFAQMEIDLTKG-SQLVFVIPDTMMTIGDFARNV
062 NP_056895.1       ------------------------------SRNRYNWLQGPLTLALKPEG
063 YP_009109689.1    ------------------------------LDRDGQPIQILTVSLQDEHR
064 YP_002519387.1    --------------------------------------------------
065 YP_595725.1       FRDIRDNP------PRVLGAMEIDLSKG-PQMVYVINSFMTTIKDFFHGI
066 YP_009508551.1    -------------------------------------ILVPLQEYQEKIL
067 YP_001497148.1    ------------------------------LNRDGQPIQVLTVSLHDEHR
068 NP_041186.1       --------------------------------------------------
069 NP_663784.1       ------------------------------GFINTKEYKNVEIKVLN---
070 YP_009508443.1    FRDCSWKGKR-----SIIARMEVDLSKG-SQLVYIAPNIIKKLADFYNNI
```

| | | |
|---|---|---|
| 071 | YP_003987465.1 | YRDTRWD--NR---KSIIATMEVDLSEG-NQIVYIIPDMMTTIRDCYEHI |
| 072 | NP_861410.1 | -------------------------------------------------- |
| 073 | YP_009345071.1 | FRDNRLLGDQ-----AILAQMEVSLADGGHQMIYVVPDIMMTIGDFYRNI |
| 074 | YP_009508582.1 | -------------------------------------VKLPVQDFKKELI |
| 075 | NP_569141.1 | -------------------------------------------------- |
| 076 | YP_002916057.1 | -------------------------------------------------- |
| 077 | NP_777384.2 | ------------------------------SRNRYNWLQGPLTLALKPEG |
| 078 | YP_233110.1 | FRDTRWTQENH-QDRSIIATMEADLSQG-HQLIYVIPDIMMTIRDFYQHI |
| 079 | NP_542258.1 | -------------------------------------------------- |
| 080 | YP_009130664.1 | FRDNRWPDDR-----SLFATMEVDLSQG-SQLVYVIPNTMMTIGDFYNNV |
| 081 | YP_009508566.1 | -------------------------------------EKLPIQDYKDNIV |
| 082 | YP_006607892.1 | -------------------------------------------------- |
| 083 | YP_009506270.1 | FRDTRWTQEGE-EDRSIIAAMEADLSQG-HQLIYVIPDIMMTIRDFYQHV |
| 084 | NP_042513.1 | -------------------------------------------------- |
| 085 | YP_006907834.1 | -------------------------------------------------- |
| 086 | YP_004442827.1 | FQDTRSQP------AKVIAAMEIDLSKG-AQMVYITPDFMTTIGDFYQGI |
| 087 | YP_009408594.1 | FWRS-YQPPA-----SYLDPVPDPSTAS-WGPDFDEP----LVPPEGGGE |
| 088 | YP_009508561.1 | -------------------------------------IKLPVQDLKNTLV |
| 089 | NP_056907.1 | ------------------------------TSSQFKLLQSPLCVYLPFRR |
| 090 | YP_002308474.1 | -------------------------------------------------- |
| 091 | YP_007761644.1 | -------------------------------------------------- |
| 092 | NP_040550.1 | -------------------------------------------------- |
| 093 | NP_758808.1 | PRDIRWNDDR-----SIIGTMEIDLSAG-SQIVYIAPNIMLSVEDFYRNI |
| 094 | YP_009345075.1 | IRDSRWGSDPR----GILGQMEIDLSTG-TELVFFVPDQMMTIKDFYDHF |
| 095 | NP_068729.1 | -------------------------------------------------- |
| 096 | NP_047255.1 | ------------------------------VGPKGLPLQVLTLQLEEEYR |
| 097 | YP_006273075.1 | FRDNRWQGDQ-----SIFAISEVDLTRG-SQIVYVIPDTMMIIGDFYRNV |
| 098 | YP_009508888.1 | -------------------------------------VLVPLEQYKERIL |
| 099 | NP_395469.1 | -------------------------------------------------- |
| 100 | YP_009508546.1 | -------------------------------------ILVPLQEYQDRIL |
| 101 | NP_056762.1 | FYDDSFGKAR------EASIGSIEMDMNAGCGVFYSCPDFAKYIKDLSHL |
| 102 | YP_009508556.1 | -------------------------------------VLVPLQDYQDRLL |
| 103 | NP_569153.1 | FRDNRWQGDQ-----SIFATMELDLTKG-MQMVYIIPDTMMTVRDFCRNV |
| 104 | NP_043924.1 | LDSQPGDRDGNKDDPAFLQQATQGLLPWLKKAVILGGKNTSWQEMTSFCQ |
| 105 | YP_009243641.1 | -------------------------------------------------- |
| 106 | YP_004300274.1 | -------------------------------------------------- |
| 107 | NP_056848.1 | -------------------------------------------------- |
| 108 | YP_006495799.1 | FRDTRWTDDR-----QVIASMSVDMTHG-SQLVYYVIPDAMLSIHDFYNHI |
| 109 | YP_009508409.1 | -------------------------------------------------- |
| 110 | NP_687035.1 | ------------------------------GGIHVKEYQGVQVQLED--- |
| 111 | NP_045937.2 | AAKIQNYVTACRQHYPAVCDWGTIQAFTYKPPQTAHEYVKHAEIIFKNNS |
| 112 | NP_612577.1 | -------------------------------------------------- |
| 113 | NP_619548.1 | -------------------------------------------------- |
| 114 | NP_056880.1 | ------------------------------MACGVARSSQPLRWQHEDK- |
| 115 | YP_009121747.1 | LRDTRWGDDR-----QIIGTMEVDLSAG-TQLVYMVPDMVLSIDDFHNHI |
| 116 | YP_009113237.2 | FRDTRWTGDR-----SIFATMEVDLTEG-SQLTYAIPNTMMTIRDFCNHV |
| 117 | YP_009508408.1 | FRDCSWKGKR-----STIARMEVDLSKG-SQLVYVAPNIIKRLADFYNNI |
| 118 | NP_955579.1 | -----------------------------------------TLQLEEEYR |
| 119 | YP_009229919.1 | LRDTRWPDER-----QVIGTMEVDMSTG-TQLVYVVPDIMMSIEDFYEHI |
| 120 | YP_004442833.1 | FQDTRSQP------ATIIAAMEVDLSKG-AQMIYITPDFMTTIGDFYQGI |
| 121 | NP_862833.2 | -------------------------------------------------- |
| 122 | NP_040563.1 | ------------------------------GVNVAKRKTHVEWRFKD--- |
| 123 | YP_443922.1 | PPLSLTQEVYWLKCTQTGPSTPAIQFQFNTWRRLIYALHPYKTPLPEVHC |
| 124 | NP_057933.2 | ------------------------------MGPMGQPLQVLTLNIEDEHR |
| 125 | YP_009116631.1 | FRDTRWSDDR-----QVIGSMSVDMTRG-AQLVYIIPNAMMSIHDFYNRI |
| 126 | YP_009513249.1 | -------------------------------------IKLDLEEQQGTLL |
| 127 | YP_223871.1 | ------------------------------IRTEGKLLV--TAPLEEEYR |
| 128 | NP_127504.1 | LLDTRYLEYQH----ANLGTAEITLNAG-TVFITLFPNFTMSLSDANLST |
| 129 | NP_044929.1 | -------------------------------------IKIDVESQKHTLI |
| 130 | NP_057860.1 | ------------------------------TQDHFKLTSLPVLIRLPFRT |
| 131 | NP_040973.1 | -------------------------------------------------- |
| 132 | NP_777317.1 | FRDTRWTREVSGEDRSIIAAMEVDLSRG-HQLIYVIPNIIMTIRDFYQHI |
| 133 | NP_054716.1 | -------------------------------------IKIDIQKQQEQLL |
| 134 | YP_001856242.1 | -------------------------------------------------- |
| 135 | NP_954565.2 | ------------------------------QSNNPKQSSKYLTWRDKENN |
| 136 | NP_569150.1 | FRDTRWTQEGE-EGRSIIGAMEADLSQG-HQLIYVIPDIMMTIRDFYQHV |
| 137 | NP_659397.1 | -------------------------------------------------- |
| 138 | YP_009513211.1 | ------------------------------TWED-RPAMCLVLNLEEEYR |
| 139 | YP_233107.1 | FRDTRWNSDDR----SIIAAMEVDLSEG-NQLIYIMPDMMMTIKDFYRHI |
| 140 | YP_009109692.1 | ------------------------------LDRDGQPIQILTVSLQDEHR |
| 141 | NP_057849.4 | ------------------------------GFIKVRQYDQILIEICG--- |
| 142 | NP_049560.1 | -------------------------------------------------- |
| 143 | YP_009506267.1 | FRDCSWRGRR-----SMIAKMEVDLTKG-SQMVYIAPNLIKRLTDFYNNI |

```
144 NP_039820.1         FRDTRWSD-----DRAVLAAMEIDLSEG-NQIVYVLPDIMMTIKSFYRHI
145 NP_056790.1         ------------------------------TWGE-RPTMCLVLNLEEEYR
146 YP_002321513.1      LRDTRFNDSR-----EILGTMEVDFTNG-AQLVYVAPNYMSTISDFYNHF
147 NP_955591.1         ------------------------------------------LNIEDEHR
148 YP_001956722.2      -------------------------------------TLVPLQEYEERLL
149 NP_056902.1         ------------------------------VGGNVETFSTPVTIKK----
150 NP_056886.1         ------------------------------GGGIPMRKSRDMIELGVINR
151 NP_041261.1         --------------------------------------------------


001 YP_004442836.1      QISILTKGYQG--FQ-GEANLLITRSCRCRLTNVPNVGFAYNIQKVVEYL
002 NP_056728.1         --------------------------------------------------
003 YP_009508537.1      NNSILS--------------------------------------------
004 YP_004732983.2      QVSITTRGYGTG-WVGGESNLIVTRSLTGRITNTSQANFNYQIEGVADYL
005 YP_009508407.1      --------------------------------------------------
006 NP_040939.1         TRKGTIVVLPQS--------------------------------------
007 YP_003864102.1      --------------------------------------------------
008 YP_009002585.1      QISILTRGYSN--WQNGEANLLVTRGIVGRLSNTPNVAFAYEIGNVTDYL
009 YP_003284237.1      QISIKTKGYDT--WEGGEANLLITRSITSRLSNTPNVGFAYKIERVAEYL
010 YP_009268869.1      RIEGTIVVLPSS--------------------------------------
011 YP_009506251.1      QLSILTRGYDN--WQNGEANLLVTRSMTGRLSNTPNVAFACQIASATDYL
012 NP_955611.1         --------------------------------------------------
013 YP_008567619.1      HISVLTRGYEG--WQHGEANLLVTRGMIGRLSNTPNVGFAYDIQEVVDYF
014 NP_149413.1         QLSIKTKGYEG--WQ-GEANLHLERLITGRLSNTSNVAFKYKIDGVTAYL
015 YP_004442824.1      QISVLTKGYNG--FQ-GEANLLITRSCRCRLTNVPNVGFAYNIQKVVEYL
016 YP_001931967.1      --------------------------------------------------
017 YP_002455786.1      --------------------------------------------------
018 YP_009109694.1      LFDIPV--------------------------------------------
019 NP_789739.1         --------------------------------------------------
020 NP_955619.1         TPIVLTSCLVDTKNNWA-----------------------------IIG-
021 YP_224289.1         QISIRTRGYDS--WEGGEANLLITRSVTARLSNTSNVGFAYRIDRVAEYL
022 YP_009182100.1      --------------------------------------------------
023 NP_705927.1         --------------------------------------------------
024 YP_009508577.1      NKANIN--------------------------------------------
025 YP_004442830.1      QLSIKTKGYNN--WR-GEANLHLERMVTARLSSTSNTGFEIKIDKVVQFI
026 NP_758887.1         --KTTHASVLVGP-------------------------------------
027 YP_009508406.1      --------------------------------------------------
028 YP_610965.1         EVVVMTHGYQD--WVDGESNLLITKALVGRLTNTSYAHFKYNVEGVADYL
029 YP_001931961.1      --------------------------------------------------
030 YP_241114.1         --------------------------------------------------
031 YP_006732334.1      --------------------------------------------------
032 YP_009041481.1      QVAIITRGYDT--WVAGESNLLITRGLVARVTNSSVSGFRFGIEGVLNYF
033 YP_001036293.1      QVSILTKGYDD--YQNSEANLLITKGMIGRLSNTSNVGFAYTVDGVAEYF
034 YP_605811.1         QISILTKGYEG--FQ-GEANLLITRSCRCRLSNVPNVGFQYNIQNVVEFL
035 NP_955577.1         LFEPES--------------------------------------------
036 YP_003208050.1      EVVVATRGYEE--WQGGESNLLITRGLVGRLTNTSHASFNYNVEQVAEHL
037 NP_789740.1         --HKAIGTVLVGP-------------------------------------
038 YP_004442839.1      QISILTKGYNG--FQ-GESNLLVTRSCRCRLSNVPNVGFQYNIQNVVEFL
039 YP_009507791.1      ICNGKICVIPQNN-------------------------------------
040 YP_004581513.1      --------------------------------------------------
041 NP_043933.1         --------------------------------------------------
042 YP_009507248.1      RTDHLN--------------------------------------------
043 YP_002117531.1      QVSIQTKGYDT--WTNGEANLLVTRSCTARLSNTPNVGFNYKTERVAEYL
044 YP_004347415.1      --------------------------------------------------
045 NP_056891.1         SGLIKPFVIPNLPVNLWGRDLLSQMKIMMCSPNDIVTAQMLAQGYSPGKG
046 YP_009513242.1      KQTTLP--------------------------------------------
047 YP_009352866.1      QVAVNTMEYDT--WIAGESNLLITRGLVARITNSSVSGFRFGIEGVLNYF
048 YP_009165750.1      --------------------------------------------------
049 NP_040840.1         IIRGTIVVLATS--------------------------------------
050 YP_004222728.1      --------------------------------------------------
051 YP_009506264.1      QISIRTKGYDS--WQGGEANILITRSITARLSNTPNVGFAYRIEQVAEYL
052 YP_567050.1         --------------------------------------------------
053 YP_009508571.1      EKTALS--------------------------------------------
054 NP_056803.1         NKTALP--------------------------------------------
055 NP_955564.1         SGIIHPFVIPTLPFTLWGRDIMKDIKVR----------------------
056 YP_009508411.1      QISVMTKGYED--YTNSEANLLITKSMVGRLSNTSNVGFQYSVSGIADYF
057 YP_009140788.1      QISIQTTGYES--WQNGEANLLITRGMTGRLSNTPNVGFAYRISHVTDYL
058 YP_009508410.1      --------------------------------------------------
059 NP_040333.1         LHETSK--------------------------------------------
060 YP_008992013.1      TLKVQVQ-IQGAEQISTAKIATLHHQLVYRLQNHALDLPTPEHHSDTLMV
061 NP_041734.1         QLSILTRGYEN--WQNGEANLLITRGMTGRLSNTPNVAFAYQIASATDYL
062 NP_056895.1         PFITIPKILVDTFDKWQ-----------------------------ILG-
```

```
063 YP_009109689.1    LFDIPV--------------------------------------------
064 YP_002519387.1    --------------------------------------------------
065 YP_595725.1       QLTVKVKGYEG--WQ-GEANLHIERLITARLSNTTNVYFKYKVEGVASFI
066 YP_009508551.1    SKTALP--------------------------------------------
067 YP_001497148.1    LFETPV--------------------------------------------
068 NP_041186.1       --------------------------------------------------
069 NP_663784.1       --KRVRATIMTGD-------------------------------------
070 YP_009508443.1    QISILTKGYDD--YHNSEANLLITKGLVGRLSNTSNVGFVYSISGIADYF
071 YP_003987465.1    QISIRTIGYDD--DWNGESNLLITRGMTARLSNTPNVGFAYNISRVADYL
072 NP_861410.1       --------------------------------------------------
073 YP_009345071.1    QISIQTRGYGN--WRNAEANLLVTRGLVGRLSNTSNVGFEYSISNVTDYL
074 YP_009508582.1    NKANIN--------------------------------------------
075 NP_569141.1       --------------------------------------------------
076 YP_002916057.1    --------------------------------------------------
077 NP_777384.2       PFITIPKILVDTFDKWQ-----------------------------ILG-
078 YP_233110.1       QISILTKGYEG--FQ-GEANLLITRSCRCRLSNVPNVGFQYNIQNVVEFL
079 NP_542258.1       --------------------------------------------------
080 YP_009130664.1    QVSILTRGYEA--WR-GEANLLITRGLVGRLSNTPNVGFAYEVQSVVDYL
081 YP_009508566.1    KRADIT--------------------------------------------
082 YP_006607892.1    --------------------------------------------------
083 YP_009506270.1    QISVLTKGYQG--FQ-GEANLLITRSCRCRLTNVPNVGFAYNIQKVVEYL
084 NP_042513.1       --------------------------------------------------
085 YP_006907834.1    --------------------------------------------------
086 YP_004442827.1    QLSIKTKGYIN--WR-GEANLHLERMITARLSSTSNTGFEIKIDKVVQYM
087 YP_009408594.1    HCVAATRAENE--ILDDNDALFSEHDLFG-------HDLSYDPD--QDLF
088 YP_009508561.1    SQANIG--------------------------------------------
089 NP_056907.1       APVTLPSCLVDTNSKWA-----------------------------IIGR
090 YP_002308474.1    --------------------------------------------------
091 YP_007761644.1    --------------------------------------------------
092 NP_040550.1       --------------------------------------------------
093 NP_758808.1       QLAIQTQGYEN--WNSAESNLLISRALIGRLTNDSFTGFQYNISNVAEYL
094 YP_009345075.1    QLVVKVKGYDS--WIGGEANLLITKVFVGRLSNTSNTAFRYAVDDLGEYL
095 NP_068729.1       --------------------------------------------------
096 NP_047255.1       LFEPES--------------------------------------------
097 YP_006273075.1    QISIQTSGYEN--WQNGEANLLITRGMTGRLSNTPNVGFAYRISHVTDYL
098 YP_009508888.1    KETALE--------------------------------------------
099 NP_395469.1       --------------------------------------------------
100 YP_009508546.1    NKTALP--------------------------------------------
101 NP_056762.1       KIGIQTLGYEN----YEGKNLSVAIKTIGRLTTNIQSKYKINVKDIVEQI
102 YP_009508556.1    KQTALP--------------------------------------------
103 NP_569153.1       QISILTKGYGN--WQNGEANLLVTRGIVGRLSNTPNVAFAYQIQNVTDYL
104 NP_043924.1       RLWLVRDQFADKTGVSKARPIVRNEGPRPQQGHSKIVFGGNCRNCGKAGH
105 YP_009243641.1    --------------------------------------------------
106 YP_004300274.1    --------------------------------------------------
107 NP_056848.1       --------------------------------------------------
108 YP_006495799.1    QVSIQTRGYNGG-WTGGDSNMIVTRSLIGRLTNQSTTNFGYQIQGVTDYL
109 YP_009508409.1    --------------------------------------------------
110 NP_687035.1       --KIITGSILIGS-------------------------------------
111 NP_045937.2       GLEWQHATVPFINMVVQGLPPKVTRSLMSGNPDWSTKTIPQIIPLMQHYL
112 NP_612577.1       --------------------------------------------------
113 NP_619548.1       --------------------------------------------------
114 NP_056880.1       SGIIHPFVIPTLPFTLWGRDIMKDIKVR----------------------
115 YP_009121747.1    QIAIQTHGYSD--WQGGESNLLITVGLVGRISNTSYTGFQYNVENVVDHL
116 YP_009113237.2    EISVLTRGYEE--WN-GEANLLITRGLVGRLTNTSNAGFAYSIQNVTDYL
117 YP_009508408.1    QISILTKGYED--YQNSEANLLITKGLIGRLSNTSNIGFAYSIDGIADYF
118 NP_955579.1       LFEPES--------------------------------------------
119 YP_009229919.1    QLAIQTRGYEG--WQ-GESNLIVTTALVGRLTNTSYASFRYNVQNVAEHL
120 YP_004442833.1    QLSIKTKGYNN--WR-GEANLHLERMVTARLSSTSNTGFEIKIDKVIQYM
121 NP_862833.2       --------------------------------------------------
122 NP_040563.1       --KTGIIDVLFSD-------------------------------------
123 YP_443922.1       TMNVTD--------------------------------------------
124 NP_057933.2       LHETSK--------------------------------------------
125 YP_009116631.1    QVSIQTRGYGTG-WEGGDSNMIITRSLVGRLTNTSITSFEYRIDNVTDYL
126 YP_009513249.1    NNSILS--------------------------------------------
127 YP_223871.1       LFLEAP--------------------------------------------
128 NP_127504.1       ALKIQVQ-IQGAPLTKDSIQATLHYQIAWRVQNHAMDLTLPGGEEALFLK
129 NP_044929.1       TESTLS--------------------------------------------
130 NP_057860.1       TPIVLTSCLVDTKNNWA-----------------------------IIG-
131 NP_040973.1       -------VLEDNS-------------------------------------
132 NP_777317.1       QMSILTKGYTG--FQ-GEANLLVTRSCRCRLTSVPNAGFAFNIQRVVEYL
133 NP_054716.1       HTTNLS--------------------------------------------
134 YP_001856242.1    --------------------------------------------------
135 NP_954565.2       SGLIKPFVIPNLPVNLWGRDLLSQMKIMMCSPNDIVTAQMLAQGYSPGKG
```

| | | |
|---|---|---|
| 136 | NP_569150.1 | QISILTKGYQG--FQ-GEANLLITRSCRCRLTNVPNVGFAYNIQRVVEYL |
| 137 | NP_659397.1 | -------------------------------------------------- |
| 138 | YP_009513211.1 | LHEKPV-------------------------------------------- |
| 139 | YP_233107.1 | QMSIQTRGYDS--WTGAEANLLITRSITSRLSNTPNVGFAFQVNKVAEYL |
| 140 | YP_009109692.1 | LFDIPV-------------------------------------------- |
| 141 | NP_057849.4 | --HKAIGTVLVGP------------------------------------- |
| 142 | NP_049560.1 | -------------------------------------------------- |
| 143 | YP_009506267.1 | QISILTKGYDD--YQNSEANLLITKGLVGRLSNTSNVGFAYSISGITDYF |
| 144 | NP_039820.1 | QICVMTKGYDG--WQ-GEDNLLITRGLTGRLSNTSNVGFAYDVKAMVEHL |
| 145 | NP_056790.1 | LHEKPV-------------------------------------------- |
| 146 | YP_002321513.1 | QLAIQTRGYESWVGQNAEANIGLSRQVIGRLSNDNVDRYSYNTTAVTDLL |
| 147 | NP_955591.1 | LHETSK-------------------------------------------- |
| 148 | YP_001956722.2 | KQTMLT-------------------------------------------- |
| 149 | NP_056902.1 | KGRHIKTRMLVAD------------------------------------- |
| 150 | NP_056886.1 | DGSLERPLLLFPAVAMVRGSILG--------------------------- |
| 151 | NP_041261.1 | -------------------------------------------------- |

| | | |
|---|---|---|
| 001 | YP_004442836.1 | NSKGVKAIQAQKLSTKKFQGTEWNIKPS-NVVVP-MQPASMITRVNYDSS |
| 002 | NP_056728.1 | -------------------------------------------------- |
| 003 | YP_009508537.1 | -----KKGKEELKRLFEKYSALWQSWENQ---VGHRKIRPHKIATGTVK- |
| 004 | YP_004732983.2 | ASHGVQSIPGQPWRDINQEG-SWNLRPSSIQAPT-QVPTGLVSRQSATGN |
| 005 | YP_009508407.1 | -------------------------------------------------- |
| 006 | NP_040939.1 | -----------PVEVLGRDNMARFGIKIIMANLEEKRIPITKVKLKEGCT |
| 007 | YP_003864102.1 | --------------------KGIWKFLTTVTDMLAPQKYADPIIWKSDT- |
| 008 | YP_009002585.1 | TSQGVQAIAGRRYDSRELQNQQWILRPPQVERLP-QAPKNVDTRNLIDGS |
| 009 | YP_003284237.1 | RSKGVKAINATKHSAKQFQGGEWTLKPS-RVVVP-MQPSSMNTSTKYDGS |
| 010 | YP_009268869.1 | -----------PVEVLGRDNMAKLDIGIIMANLEEKKIPITQVKLKEGCK |
| 011 | YP_009506251.1 | ASHGVKAIAGKKVNLQQLRNQQWVLRPPQADVLP-MQPRSVETRNLIDGS |
| 012 | NP_955611.1 | ---------------------------------TVALHLAIPLKWKPDHT |
| 013 | YP_008567619.1 | ITHGVNALPGAKYDTTRLQGQQWILKKP-ELIVP-TRPGHINSKDLVDGT |
| 014 | NP_149413.1 | KTNGIKAIDAKKESTKKIRGGEWNIQPS-TVEVV-MEPRDMMQSQNFDGT |
| 015 | YP_004442824.1 | NSKGVKAIQAPKLDTKKFRGTDWNIKPS-EVVVP-MEPTRMITRVNYDSS |
| 016 | YP_001931967.1 | -------------------------------------------------- |
| 017 | YP_002455786.1 | -----------MPGGPLLTGGPPGPDPVIPG-----DHPCSHR-LRTSSR |
| 018 | YP_009109694.1 | -----TTS--LPDVWLQDFPQAWAETGGL----GRAKCQAPIIIDLKPT- |
| 019 | NP_789739.1 | ---------------------------------PISPIETVPVKLKPGMD |
| 020 | NP_955619.1 | -----------RDALQQCQGVLYLPEAKRPPVILPIQAPAVLG-LEHLPR |
| 021 | YP_224289.1 | RTKGVRAIDATKHDARRFQHGEWNLRPS-TAVVP-VQPTSITSAVNYDRS |
| 022 | YP_009182100.1 | -------------------------------------------------- |
| 023 | NP_705927.1 | ---------------------------------PISPIETVPVKLKPGMD |
| 024 | YP_009508577.1 | -----NEEKKQLAKLLDKYDVLWQQWENQ---VGHRKIPPHNIATGTVA- |
| 025 | YP_004442830.1 | RTNGIKAIEAEKKSVKQFQGKEWNIQPI-KVEVA-MKPNKMTITDEYDNS |
| 026 | NP_758887.1 | ----------TPVNIVGRNVLKKLGC-TLNF--PVSKVETVKVTLKPGTD |
| 027 | YP_009508406.1 | -------------------------------------------------- |
| 028 | YP_610965.1 | ASNGIVALAGQRHTTTELQGLDWNLKPK-QSTAP-VQPQEVTYNNRLDGS |
| 029 | YP_001931961.1 | -------------------------------------------------- |
| 030 | YP_241114.1 | -----------------SRSKRLGVFSLGTSD--SPVTHADPIDWKSEE- |
| 031 | YP_006732334.1 | -------------------------------------------------- |
| 032 | YP_009041481.1 | NSRGVMAIKAQRRSTVPQRGIAWELRPSSIPRPI-QAPTNAVIRETTRGT |
| 033 | YP_001036293.1 | ITHGVQALKGMPMTTDDLQGTLWNIKPS-RIIVP-SQPTQAEDRELADGS |
| 034 | YP_605811.1 | KSKGVKALNATKLSTRRFQGSEWNIRPS-EVVVP-MQPTTMIVRVNYDSS |
| 035 | NP_955577.1 | -----TQKQEMD-IWLKNFPQAWAETGGM----GTAHCQAPVLIQLKAT- |
| 036 | YP_003208050.1 | SSHGIVALPGRRHSADAHNGEMWSIRPP-RSNSV-RNPREVQISNRLDGS |
| 037 | NP_789740.1 | ----------TPVNIIGRNLLTQIGC-TLNF--PISPIETVPVKLKPGMD |
| 038 | YP_004442839.1 | KSKGVKALNATKISTKKFQGGEWNIKPS-EVVVP-MQPTRMITRVNYDSS |
| 039 | YP_009507791.1 | ----------LLEPLFGRDNMKLLDIKLVMANIS-TKIPIVKAKLVDPNK |
| 040 | YP_004581513.1 | -------------------------------------------------- |
| 041 | NP_043933.1 | -------------------------------------------------- |
| 042 | YP_009507248.1 | --------EEEKANLLRLCSRYSDVFYIDGEALTFTNKIKHRIRTTDEVP |
| 043 | YP_002117531.1 | HSRGVQAINATQHNALKLRNTEWHLQPS-QATAP-MQPQTVQTVGRYDGS |
| 044 | YP_004347415.1 | -------------------------------------------------- |
| 045 | NP_056891.1 | LGKKENGILHPIPNQGQSNKKGFGNFLTAAIDILAPQQCAEPITWKSDE- |
| 046 | YP_009513242.1 | -----KEQKDQLEKLFLKYDALWQHWENQ---VGHRRIKPHNIATGTLA- |
| 047 | YP_009352866.1 | NSKGVMAIKARPRSTLAQRGIAWELRPTSIPRPI-QAPTSAVVRETSRGN |
| 048 | YP_009165750.1 | -------------------------------------------------- |
| 049 | NP_040840.1 | -----------PVEVLGRDNMRELGIGLIMANLEEKKIPSTRVRLKEGCK |
| 050 | YP_004222728.1 | ---------------------------------TVALHLAIPLKWKPDHT |
| 051 | YP_009506264.1 | RSKGVKAINATKKSARQFQGGEWLLRPS-QVVVP-MQPSSLTTSTRYDGT |
| 052 | YP_567050.1 | ---------------------------------------------EHFPP |
| 053 | YP_009508571.1 | -----EEFKKQLQTLFLKYDNLWQHWENQ---VCHRKIRPHNIATGDYP- |
| 054 | NP_056803.1 | -----EEQKQQLKALFTKYDNLWQHWENQ---VGHRKIRPHNIATGDYP- |

```
055 NP_955564.1       -----------------LMTDSPDDSQDLMIGAIESNLFADQISWKSDQ-
056 YP_009508411.1    ATRGVQALKGTPFTAEEYQGELWNLKPS-RLTIA-MQPQESEERELPDHS
057 YP_009140788.1    ASRGVQALPGRRYNSDMLRGRNWIIRQPQLQVT--MMPRSVQTTNLLDGS
058 YP_009508410.1    --------------------------------------------------
059 NP_040333.1       -----GPDVPLGSTWLSDFPQAWAETGGM----GLAVRQAPLIIPLKAT-
060 YP_008992013.1    LAESDQIPTIIQIPRQIPRHELLKLMPLEWISNYEQFHNNTAPVQTSESM
061 NP_041734.1       ASHGVKAIAGKKMNLQHLRNQQWILRPPQTDITP-MQPRSVETRNLVDGS
062 NP_056895.1       -----------RDVLSRLQASISIPEEVRPPMVGVLDAPPSHIGLEHLPV
063 YP_009109689.1    -----TTS--LPDVWLQDFPQAWAETGGL----GRAKCQAPIIIDLKPT-
064 YP_002519387.1    --------------------------------------------------
065 YP_595725.1       KTKGIKAIEATKKSVKGIRGGEWNILPS-KLEVV-MQPTKVQTTENYDGT
066 YP_009508551.1    -----EDQKQQLKTLFVKYDNLWQHWENQ---VGHRKIRPHNIATGDYP-
067 YP_001497148.1    -----TTN--LLEAWLQDFPQAWAETGGL----GRAKCQAPIIIDLKPT-
068 NP_041186.1       --------------------------------------------------
069 NP_663784.1       ----------TPINIFGRNILTALGM-SLNL--PVAKIEPIKVTLKPGKD
070 YP_009508443.1    ITHGVQAIIGAPVSTEDIQGTLWNVKPT-TLVVP-VQPTAVDDRELADGS
071 YP_003987465.1    KSKGVKAIDASKEKTTQFKGREWVLKPSQASVTP-IMPTIADAHERYDGS
072 NP_861410.1       --------------------------------------------------
073 YP_009345071.1    TSHGVRAIPGKKVSESEHLGRSWILREP-EVSII-QQPTIADTRNLPNGS
074 YP_009508582.1    -----NEEKKQLAKLLDKYDILWQQWENQ---VGHRKIPPHNIATGTVA-
075 NP_569141.1       --------------------------------------------------
076 YP_002916057.1    --------------------------------------------------
077 NP_777384.2       -----------RDVLSRLQASISIPEEVRPPMVGVLDAPPSHIGLEHLPV
078 YP_233110.1       KSRGVKALNATKLSTRRFQGGEWNIRPS-EVVVP-MQPTTMIVRVNYDSS
079 NP_542258.1       ---------------PTLKCSP-------------CPNTQAYW-VRTSSP
080 YP_009130664.1    VSHGVQAIPGRRRTVQELQGYNWVVRPT-QASIP-RQPTEVDTRTNIDGT
081 YP_009508566.1    -----KEEKGMLYKLLDKYDPLWQQWENQ---VGNRQITPHIIATGTIN-
082 YP_006607892.1    --------------------------------------------------
083 YP_009506270.1    NSKGVKAIQAQKLSTKKFQGTDWNIKPS-NVVVP-MQPTTMITRVNYDSS
084 NP_042513.1       --------------------------------------------------
085 YP_006907834.1    --------------------------------------------------
086 YP_004442827.1    KTNGIQAIEAQKESIKKYQGAEWNIKPI-KVEVT-MKPKSMIVQDEYDNS
087 YP_009408594.1    LEHDLFTPP-----PDWDAWLDTDLSET-AESQG-PNPMMIQVTSINDHE
088 YP_009508561.1    -----KEDKIKLAKLLDKYDDLWQQWDNQ---VGNRKITPHNIATGTYP-
089 NP_056907.1       DILQKCQSVLYLPEDNLCKGTPRLPDGITPPRLLPVTTPNVIG-LEHFPP
090 YP_002308474.1    ----------------------------------------CHW-AGTSPP
091 YP_007761644.1    --------------------------------------------------
092 NP_040550.1       ---------------------------------TVALHLAIPLKWKPDHT
093 NP_758808.1       HSHGVQAIEG--QAHPRTLGNRWILQAP-APPRS-LVPQNVETTTLLDGN
094 YP_009345075.1    ATKGIRAVPGQRYDASQYEAERWQLEPP-AGATV-RVPSRAHVWTDNQGS
095 NP_068729.1       --------------------------------------------------
096 NP_047255.1       -----TQKQEMD-IWLKNFPQAWAETGGM----GTAHCQAPVLIQLKAT-
097 YP_006273075.1    ASRGVQALPGRRYRSEMLRGRNWIIRQPQIQAA--MMPRNVETRNLIDGT
098 YP_009508888.1    -----GQFKQQLQNILSTFDTLWQHWENQ---VGHRKIPPHNIATGTHP-
099 NP_395469.1       --------------------------------------------------
100 YP_009508546.1    -----EEQKQRLKTLFAKYDTLWQHWENQ---VGHRRIRPHNIATGDYP-
101 NP_056762.1       SSQGIIMVAPMEIDSSHLDGNEWDLSKFLNHENTSRVPTKALIYQNLQGG
102 YP_009508556.1    -----KEQKDQLEKLFLKYDALWQHWENQ---VGHRRIKPHNIATGTLA-
103 NP_569153.1       VSHGIQALPGRRYSTADIQGQQWFLRPSNIPAVP-MAPTNVDTRNMIDGS
104 NP_043924.1       MARDCWAKGGGQEGKGPRQNTTWKPKSGAIASAPPAESPYADCAKQLADI
105 YP_009243641.1    -------------------------------------MEPIALSWKSNE-
106 YP_004300274.1    --------------------------------------------------
107 NP_056848.1       --------------------------------------------------
108 YP_006495799.1    TSHGVSCIPGEQWSVTNRSG-EWRLQQSSIAPPE-QVPTEARLRPGPRGE
109 YP_009508409.1    --------------------------------------------------
110 NP_687035.1       ----------TPINIIGRNILAQAGM-KLVMGVLSSQIEETKVQLKEGKD
111 NP_045937.2       NLQSRQDAKIKQTPLVLQLAMPAQTMNGNKGYVGSYPTNEPYYSFQQQQR
112 NP_612577.1       --------------------------------------------------
113 NP_619548.1       --------------------------------------------------
114 NP_056880.1       -----------------LMTDSPDDSQDLMIGAIESNLFADQISWKSDQ-
115 YP_009121747.1    ATNGITAIPGERRSIEDLEGMSWNLRPA-PLTMI-RIPARVEVNQRLDRS
116 YP_009113237.2    TSHGVKALPGRPRSTEDIRGLNWIVRPPAPRIQP-MQPQHLETHDLLGGG
117 YP_009508408.1    ITNGVQAIKGAPMSTEDIQGTLWNIKPT-TLVVP-VQPTAADDRELADGS
118 NP_955579.1       -----TQKQEMD-IWLKNFPQAWAETGGM----GTAHCQAPVLIQLKAT-
119 YP_009229919.1    ATNGIQAIPGQPRTIDQLKGERWELKTT-KVARS-QAPREVRLQQRSNGN
120 YP_004442833.1    KTNGIQAIEATKQSVKKYQGQDWNIRPA-KVEVA-MKPTAMEVKNEYDNS
121 NP_862833.2       -----------------SRTKRFGVFSLGTSD--SPVTHADPIDWKSGE-
122 NP_040563.1       ----------TPVNLFGRSLLRSIVT-CFTLLVHTEKIEPLPVKVRGP--
123 YP_443922.1       -----DEDNPYTEDWDENMSHQTPSIRCLSIVCGPEGVAAPVILPSHLKP
124 NP_057933.2       -----EPDVSLGSTWLSDFPQAWAETGGM----GLAVRQAPLIIPLKAT-
125 YP_009116631.1    ASNGVACIPGQKWSVANRSG-EWELQPSRIAAPL-AVPTDARLRQNPNGN
126 YP_009513249.1    -----KKGKEELKQLFEKYSALWQSWENQ---VGHRRIRPHKIATGTVK-
127 YP_223871.1       -----IQNVTLLEQWKRGIPKVWAEINPP----GLASTQAPIHVQLLST-
```

| | | |
|---|---|---|
| 128 | NP_127504.1 | IDAGSGATQCTQVPRQLSKEDLIKILPDSWVTNYEKLREPEEPLRSTEVS |
| 129 | NP_044929.1 | -----PQGQMRLKKLLDQYQALWQCWENQ---VGHRRIEPHKIATGALK- |
| 130 | NP_057860.1 | -----------RDALQQCQGVLYLPEAKRPPVILPIQAPAVLG-LEHLPR |
| 131 | NP_040973.1 | ----------LIQPLLGRDNMIKFNIRLVMAQIS-DKIPVVKVKMKDPNK |
| 132 | NP_777317.1 | NSTGVRALEAQRVGNQQYQGTDWHIEPS-DIIVP-AQPSQLRTRINYDGT |
| 133 | NP_054716.1 | -----SEGKKYLKDLFIKYDNLWQKWENQ---VGHRRITPHKIATGTLN- |
| 134 | YP_001856242.1 | ---------------------------------PISPIETVPVKLKPGMD |
| 135 | NP_954565.2 | LGKKENGILHPIPNQGQSNKKGFGNFLTAAIDILAPQQCAEPITWKSDE- |
| 136 | NP_569150.1 | NSKGVKAIQAQKLSTTKYQGTEWNIKPS-NVVVP-MQPTNLITRVNYDNS |
| 137 | NP_659397.1 | -------------------------------------------------- |
| 138 | YP_009513211.1 | -----PPS--IDPSWLQLFPMVWAEKAGM----GLANQVPPVVVELKSD- |
| 139 | YP_233107.1 | RSKGIKAIDAMKYSTGQFQHSRWNLRPS-TVVIP-VLPSTLVTSTNYDGS |
| 140 | YP_009109692.1 | -----TTS--LPDVWLQDFPQAWAETGGL----GRAKCQAPIIIDLKPT- |
| 141 | NP_057849.4 | ----------TPVNIIGRNLLTQIGC-TLNF--PISPIETVPVKLKPGMD |
| 142 | NP_049560.1 | -------------------------------------------------- |
| 143 | YP_009506267.1 | VTHGVQAIKGQPVSTEEFQETLWNIKPS-NLVVP-TQPTFAEERELADGS |
| 144 | NP_039820.1 | QSNGVKAIKGEKWDAKRFHNGQWNIEPS-KVVVP-MQPTEMKAVSNYDGT |
| 145 | NP_056790.1 | -----PSS--IDPSWLQLFPTVWAERAGM----GLANQVPPVVVELRSG- |
| 146 | YP_002321513.1 | ASRGIRAIEGARSMPDRSNS--WNLRPSSVQNQQSMIPTAYTARRHINGD |
| 147 | NP_955591.1 | -----EPDVSLGSTWLSDFPQAWAETGGM----GLAVRQAPLIIPLKAT- |
| 148 | YP_001956722.2 | -----GSYKEKLQSLFLKYDALWQHWENQ---VGHRRIKPHHIATGTVN- |
| 149 | NP_056902.1 | ----------IPVTILGRDILQDLGAKLVLAQLS-KEIKFRKIELKEGTM |
| 150 | NP_056886.1 | ------------RDCLQGLGLRLTNLIGRATVLTVALHLAIPLKWKPDHT |
| 151 | NP_041261.1 | -------------------------------------------------- |

| | | |
|---|---|---|
| 001 | YP_004442836.1 | RSIRFGDYQASTST--APPRYNNDGDSDDEI------------------- |
| 002 | NP_056728.1 | -------------------------------------------------- |
| 003 | YP_009508537.1 | -------------------------------------------------- |
| 004 | YP_004732983.2 | ISLRFTGFQDQVQTVETEEESGMTDTGERVTHYALVG------------- |
| 005 | YP_009508407.1 | -------------------------------------------------- |
| 006 | NP_040939.1 | -------------------------------------------------- |
| 007 | YP_003864102.1 | -------------------------------------------------- |
| 008 | YP_009002585.1 | ISLRFRDYATTSTP--ELPSYNEQDEEVDPDE------------------ |
| 009 | YP_003284237.1 | ISIRFGDYEASTSS--RPPKYNTADDE--ID------------------- |
| 010 | YP_009268869.1 | -------------------------------------------------- |
| 011 | YP_009506251.1 | ISIRFHDYEVAPST--SGPHYNENDEEVESES------------------ |
| 012 | NP_955611.1 | -------------------------------------------------- |
| 013 | YP_008567619.1 | ISLRFSNYTTQAS--TSKLKYNANDEEIQSDEEILAY------------- |
| 014 | NP_149413.1 | TSFRFTNFQASASR--PAPAYNDEDEV--EM------------------- |
| 015 | YP_004442824.1 | RTLKFTDYQASTST--APPKYNEDGDTDD--------------------- |
| 016 | YP_001931967.1 | -------------------------------------------------- |
| 017 | YP_002455786.1 | -------------------------------------------------- |
| 018 | YP_009109694.1 | -------------------------------------------------- |
| 019 | NP_789739.1 | -------------------------------------------------- |
| 020 | NP_955619.1 | -------------------------------------------------- |
| 021 | YP_224289.1 | ISLRFGDYQAASSS--RPPVYNKNDDE--VD------------------- |
| 022 | YP_009182100.1 | -------------------------------------------------- |
| 023 | NP_705927.1 | -------------------------------------------------- |
| 024 | YP_009508577.1 | -------------------------------------------------- |
| 025 | YP_004442830.1 | KSFRFGHYQASSNQ--PAPEYNSEDEI-IQE------------------- |
| 026 | NP_758887.1 | -------------------------------------------------- |
| 027 | YP_009508406.1 | -------------------------------------------------- |
| 028 | YP_610965.1 | VTLRFTGYAATRRPTLVEVNEDDLEEFAGMVNF----------------- |
| 029 | YP_001931961.1 | -------------------------------------------------- |
| 030 | YP_241114.1 | -------------------------------------------------- |
| 031 | YP_006732334.1 | -------------------------------------------------- |
| 032 | YP_009041481.1 | LSTRFSGYDDIPARDSYQSPFDKTEYAEDGDTSLPKR------------- |
| 033 | YP_001036293.1 | VSMRFGAYKAAPAL--MSS------EEFDSDENQVIN------------- |
| 034 | YP_605811.1 | RSIRFGDYEASTSS--SAPKYEQDGDEDEAL------------------- |
| 035 | NP_955577.1 | -------------------------------------------------- |
| 036 | YP_003208050.1 | VSLRFTGYEIA--PAVTEALQS---------------------------- |
| 037 | NP_789740.1 | -------------------------------------------------- |
| 038 | YP_004442839.1 | RSIRFGDYEASTSS--APPTYSTNDDEDEVL------------------- |
| 039 | YP_009507791.1 | -------------------------------------------------- |
| 040 | YP_004581513.1 | -------------------------------------------------- |
| 041 | NP_043933.1 | -------------------------------------------------- |
| 042 | YP_009507248.1 | -------------------------------------------------- |
| 043 | YP_002117531.1 | LSLQFGNYTHAESS--RPPVYTAHDEERDID------------------- |
| 044 | YP_004347415.1 | -------------------------------------------------- |
| 045 | NP_056891.1 | -------------------------------------------------- |
| 046 | YP_009513242.1 | -------------------------------------------------- |

```
047 YP_009352866.1   LSTRFFGYEDVPAKEPFKSPFDKTEYEEDGETSQPRR--------------
048 YP_009165750.1   ---------------------------------------------------
049 NP_040840.1      ---------------------------------------------------
050 YP_004222728.1   ---------------------------------------------------
051 YP_009506264.1   LSIRFGDYEASSSS--KPPVYNEHDDE--ID--------------------
052 YP_567050.1      ---------------------------------------------------
053 YP_009508571.1   ---------------------------------------------------
054 NP_056803.1      ---------------------------------------------------
055 NP_955564.1      ---------------------------------------------------
056 YP_009508411.1   ISMRFTRYVAAREP--ARIAFNSQDEEVSEDEQHVLA--------------
057 YP_009140788.1   ISARFANYTQAPEP--QQPRYNGHDEEEASDEA------------------
058 YP_009508410.1   ---------------------------------------------------
059 NP_040333.1      ---------------------------------------------------
060 YP_008992013.1   FERRPDGTVRMTFKPPPSAPQEPPRLSFTYSS----MITAVQT--------
061 NP_041734.1      ISIRFHDYEAATSA--SRPHYNEEDEEVESET-------------------
062 NP_056895.1      ---------------------------------------------------
063 YP_009109689.1   ---------------------------------------------------
064 YP_002519387.1   ---------------------------------------------------
065 YP_595725.1      TSFRFTNYEGASSS--KPVEHNSDD--------------------------
066 YP_009508551.1   ---------------------------------------------------
067 YP_001497148.1   ---------------------------------------------------
068 NP_041186.1      ---------------------------------------------------
069 NP_663784.1      ---------------------------------------------------
070 YP_009508443.1   ISVRFSNYKAARQP--AQITYNQDDEQIDEDEAHIIA--------------
071 YP_003987465.1   LGIKFGNYQEASTS--GPPQYNEHDDE--IV--------------------
072 NP_861410.1      ---------------------------------------------------
073 YP_009345071.1   VSVRFGGYEAAPET--SRVRFDENDNEIMDDDE------------------
074 YP_009508582.1   ---------------------------------------------------
075 NP_569141.1      ---------------------------------------------------
076 YP_002916057.1   ---------------------------------------------------
077 NP_777384.2      ---------------------------------------------------
078 YP_233110.1      RSIRFGDYEASTSS--SAPRYEQDGDEDEAL--------------------
079 NP_542258.1      ---------------------------------------------------
080 YP_009130664.1   ISLRFGNYTSARTSTTRRIAYNLRDEEIQSDEEQIIA--------------
081 YP_009508566.1   ---------------------------------------------------
082 YP_006607892.1   ---------------------------------------------------
083 YP_009506270.1   RSIRFGDYQASTSS--APPRYNDDGDSGDEI--------------------
084 NP_042513.1      ---------------------------------------------------
085 YP_006907834.1   ---------------------------------------------------
086 YP_004442827.1   KSFKFGNYQATSNT--PAPEYNSEDEI-IQE--------------------
087 YP_009408594.1   IEEAYHLPTIP----------------------------------------
088 YP_009508561.1   ---------------------------------------------------
089 NP_056907.1      ---------------------------------------------------
090 YP_002308474.1   ---------------------------------------------------
091 YP_007761644.1   ---------------------------------------------------
092 NP_040550.1      ---------------------------------------------------
093 NP_758808.1      VSIRFSNYHQAPVN-----DTQD-NSHPDIQ--------------------
094 YP_009345075.1   TSIRFTGYNTQAAPAPEYNSRDEEILEE-----------------------
095 NP_068729.1      ---------------------------------------------------
096 NP_047255.1      ---------------------------------------------------
097 YP_006273075.1   ISARSADYTQAPEP--RQPHYNEQDEEVASDEE------------------
098 YP_009508888.1   ---------------------------------------------------
099 NP_395469.1      ---------------------------------------------------
100 YP_009508546.1   ---------------------------------------------------
101 NP_056762.1      ESLRFSNYKQTRMHDPTENNSDEDEDLKILGEQLN----------------
102 YP_009508556.1   ---------------------------------------------------
103 NP_569153.1      ISLRFNSYQPAPDP--TPVAYNQHDEEVPPDED------------------
104 NP_043924.1      EKRLKDLTTAGGGPKGPNPFHKPX---------------------------
105 YP_009243641.1   ---------------------------------------------------
106 YP_004300274.1   ---------------------------------------------------
107 NP_056848.1      ---------------------------------------------------
108 YP_006495799.1   ISMRFTNFRDQAVRERALDE--DTDSGRPETSGRDET--------------
109 YP_009508409.1   ---------------------------------------------------
110 NP_687035.1      ---------------------------------------------------
111 NP_045937.2      PAPRAPPGNVPSNTCFFCKQPGHWKADCPNKTRN-----------------
112 NP_612577.1      ---------------------------------------------------
113 NP_619548.1      ---------------------------------------------------
114 NP_056880.1      ---------------------------------------------------
115 YP_009121747.1   TSLRFRSYRNNPQPPRFSVDSQDREVQANLS--------------------
116 YP_009113237.2   ISMRFTNYEAAPRE--PERRFNSQDEELASDE-------------------
117 YP_009508408.1   ISVRFSKYKAIQEP--TTPVFNDQDEELDPNEQHILA--------------
118 NP_955579.1      ---------------------------------------------------
119 YP_009229919.1   ISFRFTNYAGVPRQQPISVNQHDEEQFP-----------------------
```

```
120 YP_004442833.1     MTFSFHKYQATSST--PAPQYNSEDEIQLEE-------------------
121 NP_862833.2        --------------------------------------------------
122 NP_040563.1        --------------------------------------------------
123 YP_443922.1        WYQLAPDSSPHVTLAVGNGFEARSLGPMIRRASKLDWVPTATPGIMKAST
124 NP_057933.2        --------------------------------------------------
125 YP_009116631.1     ISLRFTDFRDQRIVEEGETS--EPE-GRPETK-EDES-------------
126 YP_009513249.1     --------------------------------------------------
127 YP_223871.1        --------------------------------------------------
128 NP_127504.1        MSKRHDKSVAISFDHSHYKKLRNTHHFMGMISDDVIVLDDPET-------
129 NP_044929.1        --------------------------------------------------
130 NP_057860.1        --------------------------------------------------
131 NP_040973.1        --------------------------------------------------
132 NP_777317.1        RSIRFADYQASAST--TLPRGSSDTDDHDAT-------------------
133 NP_054716.1        --------------------------------------------------
134 YP_001856242.1     --------------------------------------------------
135 NP_954565.2        --------------------------------------------------
136 NP_569150.1        RSIRFGNYQASTSS--APPKYNEDGDSDDDI-------------------
137 NP_659397.1        --------------------------------------------------
138 YP_009513211.1     --------------------------------------------------
139 YP_233107.1        TTLQFGDYQAASSS--RPPIYN---NEDDEI-------------------
140 YP_009109692.1     --------------------------------------------------
141 NP_057849.4        --------------------------------------------------
142 NP_049560.1        --------------------------------------------------
143 YP_009506267.1     ISIRFSRYKPVPQL--PEVSYNANDEELGDDERHIIA-------------
144 NP_039820.1        TSLRFSNYAAASTS--KPPQYNEKDEEINED-------------------
145 NP_056790.1        --------------------------------------------------
146 YP_002321513.1     TSIRFHDHVANLAQRPQNSEELDEEEAIFSRLPNDQHGSSHFTHTVMMAI
147 NP_955591.1        --------------------------------------------------
148 YP_001956722.2     --------------------------------------------------
149 NP_056902.1        --------------------------------------------------
150 NP_056886.1        --------------------------------------------------
151 NP_041261.1        --------------------------------------------------
```

```
001 YP_004442836.1     ------------------------------------------------QE
002 NP_056728.1        --------------------------------------------------
003 YP_009508537.1     --------------------------------------------------
004 YP_004732983.2     ---------------------------------TFEWLEECPSYQQRRNQ
005 YP_009508407.1     --------------------------------------------------
006 NP_040939.1        --------------------------------------------------
007 YP_003864102.1     --------------------------------------------------
008 YP_009002585.1     ------------------------------------------------EQ
009 YP_003284237.1     ------------------------------------------------SD
010 YP_009268869.1     --------------------------------------------------
011 YP_009506251.1     -----------------------------------------------EAE
012 NP_955611.1        --------------------------------------------------
013 YP_008567619.1     ---------------------------------IHQERQEVTTPEVLGVI
014 NP_149413.1        ------------------------------------------------EA
015 YP_004442824.1     ------------------------------------------------EA
016 YP_001931967.1     --------------------------------------------------
017 YP_002455786.1     --------------------------------------------------
018 YP_009109694.1     --------------------------------------------------
019 NP_789739.1        --------------------------------------------------
020 NP_955619.1        --------------------------------------------------
021 YP_224289.1        ------------------------------------------------SD
022 YP_009182100.1     --------------------------------------------------
023 NP_705927.1        --------------------------------------------------
024 YP_009508577.1     --------------------------------------------------
025 YP_004442830.1     ------------------------------------------------EV
026 NP_758887.1        --------------------------------------------------
027 YP_009508406.1     --------------------------------------------------
028 YP_610965.1        -----------------------------------------------YDD
029 YP_001931961.1     --------------------------------------------------
030 YP_241114.1        --------------------------------------------------
031 YP_006732334.1     --------------------------------------------------
032 YP_009041481.1     -------------------------------------------FNNVTEG
033 YP_001036293.1     ---------------------------------VLT----TTVSS-LLIK
034 YP_605811.1        ------------------------------------------------GD
035 NP_955577.1        --------------------------------------------------
036 YP_003208050.1     ------------------------------------------------DD
037 NP_789740.1        --------------------------------------------------
038 YP_004442839.1     ------------------------------------------------GE
```

```
039 YP_009507791.1   --------------------------------------------------
040 YP_004581513.1   --------------------------------------------------
041 NP_043933.1      --------------------------------------------------
042 YP_009507248.1   --------------------------------------------------
043 YP_002117531.1   ------------------------------------------------EE
044 YP_004347415.1   --------------------------------------------------
045 NP_056891.1      --------------------------------------------------
046 YP_009513242.1   --------------------------------------------------
047 YP_009352866.1   -------------------------------------------FTRVNEE
048 YP_009165750.1   --------------------------------------------------
049 NP_040840.1      --------------------------------------------------
050 YP_004222728.1   --------------------------------------------------
051 YP_009506264.1   ------------------------------------------------DE
052 YP_567050.1      --------------------------------------------------
053 YP_009508571.1   --------------------------------------------------
054 NP_056803.1      --------------------------------------------------
055 NP_955564.1      --------------------------------------------------
056 YP_009508411.1   ---------------------------------VLH----TSEQF-LKVQ
057 YP_009140788.1   ----------------------------------------------ELAA
058 YP_009508410.1   --------------------------------------------------
059 NP_040333.1      --------------------------------------------------
060 YP_008992013.1   --------------------------------------------------
061 NP_041734.1      -----------------------------------------------ESE
062 NP_056895.1      --------------------------------------------------
063 YP_009109689.1   --------------------------------------------------
064 YP_002519387.1   --------------------------------------------------
065 YP_595725.1      ------------------------------------------------EA
066 YP_009508551.1   --------------------------------------------------
067 YP_001497148.1   --------------------------------------------------
068 NP_041186.1      --------------------------------------------------
069 NP_663784.1      --------------------------------------------------
070 YP_009508443.1   ---------------------------------YLH----MEEPC-LKVR
071 YP_003987465.1   ------------------------------------------------ED
072 NP_861410.1      --------------------------------------------------
073 YP_009345071.1   ------------------------------------------------SQ
074 YP_009508582.1   --------------------------------------------------
075 NP_569141.1      --------------------------------------------------
076 YP_002916057.1   --------------------------------------------------
077 NP_777384.2      --------------------------------------------------
078 YP_233110.1      ------------------------------------------------GD
079 NP_542258.1      --------------------------------------------------
080 YP_009130664.1   ---------------------------------VFKLDCYEAQPAIWDTL
081 YP_009508566.1   --------------------------------------------------
082 YP_006607892.1   --------------------------------------------------
083 YP_009506270.1   ------------------------------------------------QT
084 NP_042513.1      --------------------------------------------------
085 YP_006907834.1   --------------------------------------------------
086 YP_004442827.1   ------------------------------------------------EV
087 YP_009408594.1   ------------------------------------------------DD
088 YP_009508561.1   --------------------------------------------------
089 NP_056907.1      --------------------------------------------------
090 YP_002308474.1   --------------------------------------------------
091 YP_007761644.1   --------------------------------------------------
092 NP_040550.1      --------------------------------------------------
093 NP_758808.1      ------------------------------------------------ED
094 YP_009345075.1   ------------------------------------------------EE
095 NP_068729.1      --------------------------------------------------
096 NP_047255.1      --------------------------------------------------
097 YP_006273075.1   ----------------------------------------------ELAE
098 YP_009508888.1   --------------------------------------------------
099 NP_395469.1      --------------------------------------------------
100 YP_009508546.1   --------------------------------------------------
101 NP_056762.1      --------------------------------------------------
102 YP_009508556.1   --------------------------------------------------
103 NP_569153.1      -----------------------------------------------EEQ
104 NP_043924.1      --------------------------------------------------
105 YP_009243641.1   --------------------------------------------------
106 YP_004300274.1   --------------------------------------------------
107 NP_056848.1      --------------------------------------------------
108 YP_006495799.1   ---------------------------------TPYVGALRFDFKGRSKP
109 YP_009508409.1   --------------------------------------------------
110 NP_687035.1      --------------------------------------------------
111 NP_045937.2      --------------------------------------------------
```

```
112 NP_612577.1       --------------------------------------------------
113 NP_619548.1       --------------------------------------------------
114 NP_056880.1       --------------------------------------------------
115 YP_009121747.1    ------------------------------------------------ED
116 YP_009113237.2    -------------------------------------------------E
117 YP_009508408.1    ---------------------------------YLS----EASDMPLKIR
118 NP_955579.1       --------------------------------------------------
119 YP_009229919.1    ------------------------------------------------ED
120 YP_004442833.1    ------------------------------------------------EV
121 NP_862833.2       --------------------------------------------------
122 NP_040563.1       --------------------------------------------------
123 YP_443922.1       EQMWRIVLV-----------------------------------------
124 NP_057933.2       --------------------------------------------------
125 YP_009116631.1    ---------------------------------THYV----LMFKHSSPR
126 YP_009513249.1    --------------------------------------------------
127 YP_223871.1       --------------------------------------------------
128 NP_127504.1       --------------------------------------------------
129 NP_044929.1       --------------------------------------------------
130 NP_057860.1       --------------------------------------------------
131 NP_040973.1       --------------------------------------------------
132 NP_777317.1       ------------------------------------------------AV
133 NP_054716.1       --------------------------------------------------
134 YP_001856242.1    --------------------------------------------------
135 NP_954565.2       --------------------------------------------------
136 NP_569150.1       ------------------------------------------------QA
137 NP_659397.1       --------------------------------------------------
138 YP_009513211.1    --------------------------------------------------
139 YP_233107.1       ------------------------------------------------DE
140 YP_009109692.1    --------------------------------------------------
141 NP_057849.4       --------------------------------------------------
142 NP_049560.1       --------------------------------------------------
143 YP_009506267.1    ---------------------------------YLS----TPNLV---VR
144 NP_039820.1       ------------------------------------------------EQ
145 NP_056790.1       --------------------------------------------------
146 YP_002321513.1    QPSWEICQGIVPEDTFYTNGVFHGDMRNIDPYFNEYQVDTMPWIEETNEQ
147 NP_955591.1       --------------------------------------------------
148 YP_001956722.2    --------------------------------------------------
149 NP_056902.1       --------------------------------------------------
150 NP_056886.1       --------------------------------------------------
151 NP_041261.1       --------------------------------------------------
```

```
001 YP_004442836.1    TIN---MLYIEDDLED----------------------------------
002 NP_056728.1       --------------------------------------------------
003 YP_009508537.1    --------------------------------------------------
004 YP_004732983.2    ETEENGWVKHVEGDKGFNFKVRMTPPAWSHDPQPI---------------
005 YP_009508407.1    --------------------------------------------------
006 NP_040939.1       --------------------------------------------------
007 YP_003864102.1    --------------------------------------------------
008 YP_009002585.1    LMEHVVAAFVEEEPKEIWDT------------------------------
009 YP_003284237.1    EEHTIAMIT-EEIDE-----------------------------------
010 YP_009268869.1    --------------------------------------------------
011 YP_009506251.1    IREHTVAVWIIGEEEYPDDNG-----------------------------
012 NP_955611.1       --------------------------------------------------
013 YP_008567619.1    RLTPEARLPVRTSPSAAGYNLFGTERLVIQPGERQ---------------
014 NP_149413.1       HMALCE--ELDEEL------------------------------------
015 YP_004442824.1    EVHHVNMLFFE--AED----------------------------------
016 YP_001931967.1    --------------------------------------------------
017 YP_002455786.1    --------------------------------------------------
018 YP_009109694.1    --------------------------------------------------
019 NP_789739.1       --------------------------------------------------
020 NP_955619.1       --------------------------------------------------
021 YP_224289.1       EETFQINMVLLGDDMD----------------------------------
022 YP_009182100.1    -----------------MSGFKLSSPSDEDLNRWLTDELDRVQNKIIGKV
023 NP_705927.1       --------------------------------------------------
024 YP_009508577.1    --------------------------------------------------
025 YP_004442830.1    RMTLCESEDEDEGI------------------------------------
026 NP_758887.1       --------------------------------------------------
027 YP_009508406.1    --------------------------------------------------
028 YP_610965.1       SSPLLCTEILEEE-------------------------------------
029 YP_001931961.1    --------------------------------------------------
030 YP_241114.1       --------------------------------------------------
```

| | | |
|---|---|---|
| 031 | YP_006732334.1 | -------------------------------------------------- |
| 032 | YP_009041481.1 | TSTNFALVILASCNPDFPEDDQLADGNFYT-------------------- |
| 033 | YP_001036293.1 | LMHRDAIIPQRITDGSVGYDLAACEDVVIQPRRPT--------------- |
| 034 | YP_605811.1 | IHQVNMITIIEDDAED---------------------------------- |
| 035 | NP_955577.1 | -------------------------------------------------- |
| 036 | YP_003208050.1 | ESELAAMAIQESG------------------------------------- |
| 037 | NP_789740.1 | -------------------------------------------------- |
| 038 | YP_004442839.1 | IHQVNMITFLEDDDEF---------------------------------- |
| 039 | YP_009507791.1 | -------------------------------------------------- |
| 040 | YP_004581513.1 | -------------------------------------------------- |
| 041 | NP_043933.1 | -------------------------------------------------- |
| 042 | YP_009507248.1 | -------------------------------------------------- |
| 043 | YP_002117531.1 | SIAYLYSSDGEADDTDLEFLT----------------------------- |
| 044 | YP_004347415.1 | -------------------------------------------------- |
| 045 | NP_056891.1 | -------------------------------------------------- |
| 046 | YP_009513242.1 | -------------------------------------------------- |
| 047 | YP_009352866.1 | TKDNYALVIIATESPDFPEEG-LAEGDYYT-------------------- |
| 048 | YP_009165750.1 | -------------------------------------------------- |
| 049 | NP_040840.1 | -------------------------------------------------- |
| 050 | YP_004222728.1 | -------------------------------------------------- |
| 051 | YP_009506264.1 | HLVAFLDVEEEEEDP----------------------------------- |
| 052 | YP_567050.1 | -------------------------------------------------- |
| 053 | YP_009508571.1 | -------------------------------------------------- |
| 054 | NP_056803.1 | -------------------------------------------------- |
| 055 | NP_955564.1 | -------------------------------------------------- |
| 056 | YP_009508411.1 | VLHPDARIPQRMTAEAAGYDLTVIESAQLQPGEQK--------------- |
| 057 | YP_009140788.1 | TQHHVVAMFRLSGYTEVEEFKPKFKEQYERYQREQ--------------- |
| 058 | YP_009508410.1 | -------------------------------------------------- |
| 059 | NP_040333.1 | -------------------------------------------------- |
| 060 | YP_008992013.1 | -------------------------------------------------- |
| 061 | NP_041734.1 | IREHTIAVWIG-EEEIPDQTG----------------------------- |
| 062 | NP_056895.1 | -------------------------------------------------- |
| 063 | YP_009109689.1 | -------------------------------------------------- |
| 064 | YP_002519387.1 | -------------------------------------------------- |
| 065 | YP_595725.1 | YMALFE--EEEEE------------------------------------- |
| 066 | YP_009508551.1 | -------------------------------------------------- |
| 067 | YP_001497148.1 | -------------------------------------------------- |
| 068 | NP_041186.1 | -------------------------------------------------- |
| 069 | NP_663784.1 | -------------------------------------------------- |
| 070 | YP_009508443.1 | LLNEDAIIPQRLSHQAVGYDIAVVQAAEVPALGQT--------------- |
| 071 | YP_003987465.1 | VVLAIIDDLPEDDEEDYFQQILNDYQPDSQVALDYPVLIPTED----PFA |
| 072 | NP_861410.1 | -------------------------------------------------- |
| 073 | YP_009345071.1 | IAEEVHVMIINEDDEEWEELN----------------------------- |
| 074 | YP_009508582.1 | -------------------------------------------------- |
| 075 | NP_569141.1 | -------------------------------------------------- |
| 076 | YP_002916057.1 | -------------------------------------------------- |
| 077 | NP_777384.2 | -------------------------------------------------- |
| 078 | YP_233110.1 | IHQVNMITIIEDDAED---------------------------------- |
| 079 | NP_542258.1 | -------------------------------------------------- |
| 080 | YP_009130664.1 | GE-PSGKFGYYVRYDTHDDNLHIPLEHIIATG------------------ |
| 081 | YP_009508566.1 | -------------------------------------------------- |
| 082 | YP_006607892.1 | -------------------------------------------------- |
| 083 | YP_009506270.1 | TVEHVNMLYILDEAED---------------------------------- |
| 084 | NP_042513.1 | -------------------------------------------------- |
| 085 | YP_006907834.1 | -------------------------------------------------- |
| 086 | YP_004442827.1 | RMALCDSDSDSETDD----------------------------------- |
| 087 | YP_009408594.1 | ES--TGTEVLDIN------------------------------------- |
| 088 | YP_009508561.1 | -------------------------------------------------- |
| 089 | NP_056907.1 | -------------------------------------------------- |
| 090 | YP_002308474.1 | -------------------------------------------------- |
| 091 | YP_007761644.1 | -------------------------------------------------- |
| 092 | NP_040550.1 | -------------------------------------------------- |
| 093 | NP_758808.1 | ENQFIGFLS---DLGEE--------------------------------- |
| 094 | YP_009345075.1 | SEIETARMVLFVN------------------------------------- |
| 095 | NP_068729.1 | -------------------------------------------------- |
| 096 | NP_047255.1 | -------------------------------------------------- |
| 097 | YP_006273075.1 | TQQHIVAMLRFPDYVEIQEFKPKYKEQYDKYKREQ--------------- |
| 098 | YP_009508888.1 | -------------------------------------------------- |
| 099 | NP_395469.1 | -------------------------------------------------- |
| 100 | YP_009508546.1 | -------------------------------------------------- |
| 101 | NP_056762.1 | -------------------------------------------------- |
| 102 | YP_009508556.1 | -------------------------------------------------- |
| 103 | NP_569153.1 | IRNHTIALWREDD--EVWDT------------------------------ |

```
104 NP_043924.1          --------------------------------------------------
105 YP_009243641.1       --------------------------------------------------
106 YP_004300274.1       --------------------------------------------------
107 NP_056848.1          --------------------------------------------------
108 YP_006495799.1       REEECFEVAAPDVLQSWMQQLSRSPPRQNR--------------------
109 YP_009508409.1       --------------------------------------------------
110 NP_687035.1          --------------------------------------------------
111 NP_045937.2          --------------------------------------------------
112 NP_612577.1          --------------------------------------------------
113 NP_619548.1          --------------------------------------------------
114 NP_056880.1          --------------------------------------------------
115 YP_009121747.1       EEQFVGICIQEREIHNP---------------------------------
116 YP_009113237.2       EVETILVLIAEEPEKEVWDT------------------------------
117 YP_009508408.1       LLNADAVLPQRMSYGAVGYDIAITTSEDIPPYRQV---------------
118 NP_955579.1          --------------------------------------------------
119 YP_009229919.1       EREFVGAILEEVEQPQENKPWWHDYDPDLPKHWGKQCDGCLMN-------
120 YP_004442833.1       RMAICSDDESEEPS------------------------------------
121 NP_862833.2          --------------------------------------------------
122 NP_040563.1          --------------------------------------------------
123 YP_443922.1          --------------------------------------------------
124 NP_057933.2          --------------------------------------------------
125 YP_009116631.1       WDTLGQPSGKYDYMVRYDAPEPTTWPTTNR--------------------
126 YP_009513249.1       --------------------------------------------------
127 YP_223871.1          --------------------------------------------------
128 NP_127504.1          --------------------------------------------------
129 NP_044929.1          --------------------------------------------------
130 NP_057860.1          --------------------------------------------------
131 NP_040973.1          --------------------------------------------------
132 NP_777317.1          QTEYINMLYTIE-GDD----------------------------------
133 NP_054716.1          --------------------------------------------------
134 YP_001856242.1       --------------------------------------------------
135 NP_954565.2          --------------------------------------------------
136 NP_569150.1          TIEHVNMLYIEDTSDT----------------------------------
137 NP_659397.1          --------------------------------------------------
138 YP_009513211.1       --------------------------------------------------
139 YP_233107.1          EQHIVAVITIIDDQED----------------------------------
140 YP_009109692.1       --------------------------------------------------
141 NP_057849.4          --------------------------------------------------
142 NP_049560.1          --------------------------------------------------
143 YP_009506267.1       LVNENAILPQRMSHGAVGYDIAITEAAEILPGQQK---------------
144 NP_039820.1          EINHSLNLILNDEESTDEDEEYYQYQRYAWSQVGDSTFYYDTDGVWEEID
145 NP_056790.1          --------------------------------------------------
146 YP_002321513.1       LADQAEVIIEEMDRDQHFIVITDGSNYPPGYFDFPDSDTEEPDDY-----
147 NP_955591.1          --------------------------------------------------
148 YP_001956722.2       --------------------------------------------------
149 NP_056902.1          --------------------------------------------------
150 NP_056886.1          --------------------------------------------------
151 NP_041261.1          --------------------------------------------------
```

```
001 YP_004442836.1       ----------------------------------------------DYP-
002 NP_056728.1          --------------------------------------------------
003 YP_009508537.1       --------------------------------------------------
004 YP_004732983.2       ------------------------IATGWGDDFNNPPPPPPPKTEEEEIL
005 YP_009508407.1       --------------------------------------------------
006 NP_040939.1          --------------------------------------------------
007 YP_003864102.1       --------------------------------------------------
008 YP_009002585.1       ---------------------LGQPSGKFDFLVKYSAPASSKIRIEDIVP
009 YP_003284237.1       -------------------------WLILAEELG----------YDLTEE
010 YP_009268869.1       --------------------------------------------------
011 YP_009506251.1       -----------------RIKVWEESSNGNGRFFRYYTPPSVSN--EPIIA
012 NP_955611.1          --------------------------------------------------
013 YP_008567619.1       -----------------LQRTGIAIRVPEGHYGRIAARSSAAWQLGIVVG
014 NP_149413.1          ------------------------------------------------TD
015 YP_004442824.1       ----------------------------------------------DYPS
016 YP_001931967.1       --------------------------------------------------
017 YP_002455786.1       --------------------------------------------------
018 YP_009109694.1       --------------------------------------------------
019 NP_789739.1          --------------------------------------------------
020 NP_955619.1          --------------------------------------------------
021 YP_224289.1          -------------------------YPMFRELEK---------LYITSES
022 YP_009182100.1       KDVDPDHVAMAEPGENHLRNGKVLIHQMNRMFATINTMFEKQLWFEQRQK
```

```
023 NP_705927.1        --------------------------------------------------
024 YP_009508577.1     --------------------------------------------------
025 YP_004442830.1     ------------------------------------------------SA
026 NP_758887.1        --------------------------------------------------
027 YP_009508406.1     --------------------------------------------------
028 YP_610965.1        ----------------------------DDEDLWVPVPPTCRASPYSDAE
029 YP_001931961.1     --------------------------------------------------
030 YP_241114.1        --------------------------------------------------
031 YP_006732334.1     --------------------------------------------------
032 YP_009041481.1     -----------------------------GQVLTD-WEDDY--LQDWLTD
033 YP_001036293.1     -----------------LVKTGIAIEVPAGTYAQLYLKSSFALKTGLQIT
034 YP_605811.1        ----------------------------------------------DYPR
035 NP_955577.1        --------------------------------------------------
036 YP_003208050.1     ----------------------------EDED-----------------D
037 NP_789740.1        --------------------------------------------------
038 YP_004442839.1     ----------------------------------------------DYPQ
039 YP_009507791.1     --------------------------------------------------
040 YP_004581513.1     --------------------------------------------------
041 NP_043933.1        --------------------------------------------------
042 YP_009507248.1     --------------------------------------------------
043 YP_002117531.1     --------------------QLDQVIPPVENVEGEEVSLFSLLDELDAEA
044 YP_004347415.1     --------------------------------------------------
045 NP_056891.1        --------------------------------------------------
046 YP_009513242.1     --------------------------------------------------
047 YP_009352866.1     -----------------------------GQVLSDNWEDDY--LEDWLEE
048 YP_009165750.1     --------------------------------------------------
049 NP_040840.1        --------------------------------------------------
050 YP_004222728.1     --------------------------------------------------
051 YP_009506264.1     -------------------------WLALERELG----------YDADYE
052 YP_567050.1        --------------------------------------------------
053 YP_009508571.1     --------------------------------------------------
054 NP_056803.1        --------------------------------------------------
055 NP_955564.1        --------------------------------------------------
056 YP_009508411.1     -----------------LFRTGVAIEVPTGTYAHLFPRSGIALKKGINIG
057 YP_009140788.1     -----------------QWDTLGQPSGKYDYYVQYTAPPAATIPLTEIQP
058 YP_009508410.1     --------------------------------------------------
059 NP_040333.1        --------------------------------------------------
060 YP_008992013.1     ---------------------------------AQENLPITGFTSNGFPV
061 NP_041734.1        -----------------RKKVWEESSNGNGRFFRYYTPPPTFE--GQIIA
062 NP_056895.1        --------------------------------------------------
063 YP_009109689.1     --------------------------------------------------
064 YP_002519387.1     --------------------------------------------------
065 YP_595725.1        ------------------------------------------------DD
066 YP_009508551.1     --------------------------------------------------
067 YP_001497148.1     --------------------------------------------------
068 NP_041186.1        --------------------------------------------------
069 NP_663784.1        --------------------------------------------------
070 YP_009508443.1     -----------------MFSTGIAVEVPNGTYAHLFIKSGVAKRKGLILN
071 YP_003987465.1     SSYAPLTPPALTPLPASGDNTPDGIRPVLQRIVVGEKDFTDFNTLDFTLR
072 NP_861410.1        --------------------------------------------------
073 YP_009345071.1     ----------------------RQMMADFAEEEQKHTEPIIAEEDIYLLP
074 YP_009508582.1     --------------------------------------------------
075 NP_569141.1        --------------------------------------------------
076 YP_002916057.1     --------------------------------------------------
077 NP_777384.2        --------------------------------------------------
078 YP_233110.1        ----------------------------------------------DYPR
079 NP_542258.1        --------------------------------------------------
080 YP_009130664.1     --------------------------WDDDDDDCTTSSSDEASYLQHLS-
081 YP_009508566.1     --------------------------------------------------
082 YP_006607892.1     --------------------------------------------------
083 YP_009506270.1     ----------------------------------------------EYPT
084 NP_042513.1        --------------------------------------------------
085 YP_006907834.1     --------------------------------------------------
086 YP_004442827.1     ----------------------------------------------PQSA
087 YP_009408594.1     --------------------------------------------------
088 YP_009508561.1     --------------------------------------------------
089 NP_056907.1        --------------------------------------------------
090 YP_002308474.1     --------------------------------------------------
091 YP_007761644.1     --------------------------------------------------
092 NP_040550.1        --------------------------------------------------
093 NP_758808.1        ----------------------------YELEYPSFTPVHADEFIFIIIN
094 YP_009345075.1     ----------------------------ATNKGTEIRRIERPTVPAVYPA
095 NP_068729.1        --------------------------------------------------
```

```
096 NP_047255.1         --------------------------------------------------
097 YP_006273075.1      -----------------GWDTLGEPSGKYDYYVRYTAPP-ATTPIEDIQP
098 YP_009508888.1      --------------------------------------------------
099 NP_395469.1         --------------------------------------------------
100 YP_009508546.1      --------------------------------------------------
101 NP_056762.1         ----------------------------IKMARFYTMQTPEEELREVIQQ
102 YP_009508556.1      --------------------------------------------------
103 NP_569153.1         ---------------------LGEPSGKFDFYVRYTRPAHALQDPAHIVA
104 NP_043924.1         ---------------------------GVAALAAPLCSLKGRPHVSVEIE
105 YP_009243641.1      --------------------------------------------------
106 YP_004300274.1      --------------------------------------------------
107 NP_056848.1         --------------------------------------------------
108 YP_006495799.1      ---------------------------YQGPEWDTLGEPSG-KYDYKVRY
109 YP_009508409.1      --------------------------------------------------
110 NP_687035.1         --------------------------------------------------
111 NP_045937.2         --------------------------------------------LRNMGN
112 NP_612577.1         --------------------------------------------------
113 NP_619548.1         --------------------------------------------------
114 NP_056880.1         --------------------------------------------------
115 YP_009121747.1      ----------------------------YDICFCSVCLSEAKRVEEEPLN
116 YP_009113237.2      ---------------------LGEPSGKFDYFVKYSAPEWAKNP-VKIEP
117 YP_009508408.1      -----------------KLSTGLAIEVPYGTYAQLYIKSGVAHRTGLMLN
118 NP_955579.1         --------------------------------------------------
119 YP_009229919.1      -----------------PIIGTRWKYTEDDYDLCASCMIKYKAQQGSYAA
120 YP_004442833.1      -----------------------------------------------TSA
121 NP_862833.2         --------------------------------------------------
122 NP_040563.1         --------------------------------------------------
123 YP_443922.1         --------------------------------------------------
124 NP_057933.2         --------------------------------------------------
125 YP_009116631.1      ---------------------------GWDDDPPKPPSPKG-SYEVSLRG
126 YP_009513249.1      --------------------------------------------------
127 YP_223871.1         --------------------------------------------------
128 NP_127504.1         ---------------------------------FSKTLPSLMQTHDWIHH
129 NP_044929.1         --------------------------------------------------
130 NP_057860.1         --------------------------------------------------
131 NP_040973.1         --------------------------------------------------
132 NP_777317.1         ----------------------------------------------TESA
133 NP_054716.1         --------------------------------------------------
134 YP_001856242.1      --------------------------------------------------
135 NP_954565.2         --------------------------------------------------
136 NP_569150.1         ----------------------------------------------DYPV
137 NP_659397.1         --------------------------------------------------
138 YP_009513211.1      --------------------------------------------------
139 YP_233107.1         ----------------------------------------------DYPA
140 YP_009109692.1      --------------------------------------------------
141 NP_057849.4         --------------------------------------------------
142 NP_049560.1         --------------------------------------------------
143 YP_009506267.1      -----------------LLPTGISLQVPNGFYAHLFIKSGVALRKGLILN
144 NP_039820.1         RCNDLPEYVPSETSTPTIDESEAIIDEFLEHAYEQRCDSDESLQSGDPRK
145 NP_056790.1         --------------------------------------------------
146 YP_002321513.1      -----------------AEFLQDLAEAPSQDRSSLYGSEDRSPEVSTKLD
147 NP_955591.1         --------------------------------------------------
148 YP_001956722.2      --------------------------------------------------
149 NP_056902.1         --------------------------------------------------
150 NP_056886.1         --------------------------------------------------
151 NP_041261.1         --------------------------------------------------


001 YP_004442836.1      -AAEEEQVLLEDMVGEED----------------IISKFLEELELSD---
002 NP_056728.1         --------------------------------------------------
003 YP_009508537.1      --------------------------------------------------
004 YP_004732983.2      ELYPVRRRPDPVQIARKEKAAVFSQAVNTIFEHEGKDVSRMKPSGEAPDS
005 YP_009508407.1      --------------------------------------------------
006 NP_040939.1         --------------------------------------------------
007 YP_003864102.1      --------------------------------------------------
008 YP_009002585.1      TGWGDEWEQPEQQQSPQQQQLLSCSQE------ESDDEDDWVSRFQFLAQ
009 YP_003284237.1      VEDVPPEQSSRSMVGEEK----------------IISQFLEKL-------
010 YP_009268869.1      --------------------------------------------------
011 YP_009506251.1      TGWDDDEDDDYDYDPRWDTNLESEGEEPDEPLRSEEDEEEEVQDEPVWHQ
012 NP_955611.1         --------------------------------------------------
013 YP_008567619.1      AGVIDADFQGEIKVLIFNLSNSPVEIEAGDAVAQLVLERISTPSLSVWRV
014 NP_149413.1         QQWIDRYLSEYSAQQKVVG---------------EIDEE-----------
```

```
015 YP_004442824.1   LAALEKIFSPEGMVGEEE----------------TINTFLQNLNLDT---
016 YP_001931967.1   --------------------------------------------------
017 YP_002455786.1   --------------------------------------------------
018 YP_009109694.1   --------------------------------------------------
019 NP_789739.1      --------------------------------------------------
020 NP_955619.1      --------------------------------------------------
021 YP_224289.1      MVGEDDEEEKSEDLDDEE----------------AHADFQKFLNYSDEVN
022 YP_009182100.1   VLFQNLWDFLNSYDENQKAQQKALKDSQDNIALATRNIARMLEQVKGIEE
023 NP_705927.1      --------------------------------------------------
024 YP_009508577.1   --------------------------------------------------
025 YP_004442830.1   LQEIENFLSALPP-EGGGS---------------NFSAARCSEN------
026 NP_758887.1      --------------------------------------------------
027 YP_009508406.1   --------------------------------------------------
028 YP_610965.1      EEFLQNLFSAGFTPATEEQDMEYPVSIPIPLSPIYDTYPEDVVHTDHGVL
029 YP_001931961.1   --------------------------------------------------
030 YP_241114.1      --------------------------------------------------
031 YP_006732334.1   --------------------------------------------------
032 YP_009041481.1   EPAPQPTAAPLPTLSLDEAEDIMNDFLDRYFSDEKDSTTVSSFSVEREVE
033 YP_001036293.1   GGVVDPDYRGELQIILINNTFTCVPILKGTFVAQIVLHQVATPAVQETTT
034 YP_605811.1      LSALERIIAPESMVGEED----------------TIAEFLGNLSLDSSTD
035 NP_955577.1      --------------------------------------------------
036 YP_003208050.1   DPFQDFLYMASMPRTAYES------------QPIWD--------------
037 NP_789740.1      --------------------------------------------------
038 YP_004442839.1   LAAFEQIFALESGVGEEDD---------------TIIEFLDNLRLSD---
039 YP_009507791.1   --------------------------------------------------
040 YP_004581513.1   --------------------------------------------------
041 NP_043933.1      --------------------------------------------------
042 YP_009507248.1   --------------------------------------------------
043 YP_002117531.1   AAAAAAAAVTEEPAVADP----------------IIEEFLQALDSAEQEL
044 YP_004347415.1   --------------------------------------------------
045 NP_056891.1      --------------------------------------------------
046 YP_009513242.1   --------------------------------------------------
047 YP_009352866.1   S-------QPQINISFSEEEDIISTFLG---NDEKESCYQSSFFGDRVEE
048 YP_009165750.1   --------------------------------------------------
049 NP_040840.1      --------------------------------------------------
050 YP_004222728.1   --------------------------------------------------
051 YP_009506264.1   VADT----TSEALVGDEE----------------VIQTFLSQVQD-----
052 YP_567050.1      --------------------------------------------------
053 YP_009508571.1   --------------------------------------------------
054 NP_056803.1      --------------------------------------------------
055 NP_955564.1      --------------------------------------------------
056 YP_009508411.1   AGVIDPDYTGEVKVLIQNISTQPCDIYAGEACAQIILEQHLTPEVQEVRA
057 YP_009140788.1   SGWDDQ--------PQWD-----------------DEPQ---ATTIRFEE
058 YP_009508410.1   --------------------------------------------------
059 NP_040333.1      --------------------------------------------------
060 YP_008992013.1   YPAKQNGHFLWDVPGSGHC--DPDCPCW------DDWEEDNDYVT-----
061 NP_041734.1      TGWGSD-DDNEKTPPKWD-----------------ESPDEEGPTEPIWDQ
062 NP_056895.1      --------------------------------------------------
063 YP_009109689.1   --------------------------------------------------
064 YP_002519387.1   --------------------------------------------------
065 YP_595725.1      ITFLNRILSKYSTQQKVVG---------------EEEFS-----------
066 YP_009508551.1   --------------------------------------------------
067 YP_001497148.1   --------------------------------------------------
068 NP_041186.1      --------------------------------------------------
069 NP_663784.1      --------------------------------------------------
070 YP_009508443.1   AGVIDPDYTGEVMLLVCNPTATPQFIQKGEFIAQIVLQQVLTPPVQDVAI
071 YP_003987465.1   GLDVELQTTDDFTAEEEE----------------LIGSFLNQLEEEENLY
072 NP_861410.1      --------------------------------------------------
073 YP_009345071.1   GMEDLLNISTDADLSNGNNLTSTVNGFIIVSVYHNGENMKILKVRNDTKE
074 YP_009508582.1   --------------------------------------------------
075 NP_569141.1      --------------------------------------------------
076 YP_002916057.1   --------------------------------------------------
077 NP_777384.2      --------------------------------------------------
078 YP_233110.1      LSALERIIAPESMVGEED----------------TIAEFLSNLSLDSSTD
079 NP_542258.1      --------------------------------------------------
080 YP_009130664.1   -----------------NLGGTPVPIEQDLPYPVPISDMINP--------
081 YP_009508566.1   --------------------------------------------------
082 YP_006607892.1   --------------------------------------------------
083 YP_009506270.1   LAAEEESFPLENMVGEEA----------------IISQFLESLNLSN---
084 NP_042513.1      --------------------------------------------------
085 YP_006907834.1   --------------------------------------------------
086 YP_004442827.1   IYQIEQYLATLSPQEEGGD---------------CVSAARSRNTECA---
087 YP_009408594.1   --------------------------------------------------
```

```
088 YP_009508561.1   --------------------------------------------------
089 NP_056907.1      --------------------------------------------------
090 YP_002308474.1   --------------------------------------------------
091 YP_007761644.1   --------------------------------------------------
092 NP_040550.1      --------------------------------------------------
093 NP_758808.1      GEEIPD-----------------------DFVSSFCSNFSPPPIPEPEPT
094 YP_009345075.1   NPXVHKQFEENMEERQSQGERWGTLGDDDSFLVKFSAPPTIPIEDIVPSG
095 NP_068729.1      --------------------------------------------------
096 NP_047255.1      --------------------------------------------------
097 YP_006273075.1   TGWEEE--------PKWD-----------------DSPDEKSLATIRFEE
098 YP_009508888.1   --------------------------------------------------
099 NP_395469.1      --------------------------------------------------
100 YP_009508546.1   --------------------------------------------------
101 NP_056762.1      LEREKQAMIAKLEAKMKESSKMAIVEDNFNPNNEYLEDTYSEYEDLEFEK
102 YP_009508556.1   --------------------------------------------------
103 NP_569153.1      TGWDDLDNDP-STSSPSNNILTYLTPS------SSSDEDDDMSYLQYLAQ
104 NP_043924.1      GHKIECLVDTGAEVSLTSLQLQAQRFEQVVGLGGKPVRVGIADHVDTTVG
105 YP_009243641.1   --------------------------------------------------
106 YP_004300274.1   --------------------------------------------------
107 NP_056848.1      --------------------------------------------------
108 YP_006495799.1   SAPPPTPWPNKPTGWGDEEEDLPPPRFDSLSESFS--GGGEIEPLEYHES
109 YP_009508409.1   --------------------------------------------------
110 NP_687035.1      --------------------------------------------------
111 NP_045937.2      MGRGGRMGGPPYRSQPYPAFIQPPQNHQNQYNGRMDRSQLQASAQEWLPG
112 NP_612577.1      --------------------------------------------------
113 NP_619548.1      --------------------------------------------------
114 NP_056880.1      --------------------------------------------------
115 YP_009121747.1   KAKRGNKPNKSRRRSYEKWSTLGEHSGKWDYYVRYDSPKDTTPIDQIIAT
116 YP_009113237.2   TGWESSDEGDEDDEDFRDINSFTAGVS-------LQEQEEQQQMPEVIPE
117 YP_009508408.1   AGVIDPDYRGEISLLLHNPTDSTVSVRKGDFLTQLVLHQILTPPVQEENA
118 NP_955579.1      --------------------------------------------------
119 YP_009229919.1   TYFIPIHGGNTLGQPSGKYDYQVKYTVPEDWKPAHKSQIIATGWDTESSL
120 YP_004442833.1   IQEIEEFLAALSPREEGSE---------------SVSTARCKN-------
121 NP_862833.2      --------------------------------------------------
122 NP_040563.1      --------------------------------------------------
123 YP_443922.1      --------------------------------------------------
124 NP_057933.2      --------------------------------------------------
125 YP_009116631.1   EKK----LKEKELAEFTPETDLVSQWLNQLSNSAHNSGASSSDDEPKFDE
126 YP_009513249.1   --------------------------------------------------
127 YP_223871.1      --------------------------------------------------
128 NP_127504.1      FQLDGRAVSWYKDPFDGHCPWDIDCQCYSCLYSEDEEDFEDGFPT-----
129 NP_044929.1      --------------------------------------------------
130 NP_057860.1      --------------------------------------------------
131 NP_040973.1      --------------------------------------------------
132 NP_777317.1      LAALERLLAPESMVGEHRQPIFDLDDPNPDIIEEDFLNFLDSLTLND---
133 NP_054716.1      --------------------------------------------------
134 YP_001856242.1   --------------------------------------------------
135 NP_954565.2      --------------------------------------------------
136 NP_569150.1      MAAEEEIFPLENMVGEDD----------------IISQFLENLDIT----
137 NP_659397.1      --------------------------------------------------
138 YP_009513211.1   --------------------------------------------------
139 YP_233107.1      LAAVEKQIFPENMVGEEE---------------AIISSFLQKLELSE---
140 YP_009109692.1   --------------------------------------------------
141 NP_057849.4      --------------------------------------------------
142 NP_049560.1      --------------------------------------------------
143 YP_009506267.1   AGVIDPDYTGELKLLLLNPTGAPVHVQAGEFIAQAVLQAIITPPVQEAQA
144 NP_039820.1      YEYPTPQSSPEHLDNESRSRSSSASSTSMQDDVEEIVRLMKEMRMKKQKK
145 NP_056790.1      --------------------------------------------------
146 YP_002321513.1   PNDPNVDFKIHEYCCLSPTHQSCGFFQASTSTAHQSAEQSVVGTSNNQNN
147 NP_955591.1      --------------------------------------------------
148 YP_001956722.2   --------------------------------------------------
149 NP_056902.1      --------------------------------------------------
150 NP_056886.1      --------------------------------------------------
151 NP_041261.1      --------------------------------------------------


001 YP_004442836.1   --------------------------------------------------
002 NP_056728.1      --------------------------------------------------
003 YP_009508537.1   --------------------------------------------------
004 YP_004732983.2   DPDSPVWKVKKSPYPHKPMKLKD---------------------------
005 YP_009508407.1   --------------------------------------------------
006 NP_040939.1      --------------------------------------------------
```

007 YP_003864102.1 --------------------------------------------------
008 YP_009002585.1 FDEEP---------------------------------------------
009 YP_003284237.1 --------------------------------------------------
010 YP_009268869.1 --------------------------------------------------
011 YP_009506251.1 EEEEEE--------------------------------------------
012 NP_955611.1 --------------------------------------------------
013 YP_008567619.1 FTEETVRGTASPLDEDLPYPQLR-------------------------QQ
014 NP_149413.1 --------------------------------------------------
015 YP_004442824.1 --------------------------------------------------
016 YP_001931967.1 --------------------------------------------------
017 YP_002455786.1 --------------------------------------------------
018 YP_009109694.1 --------------------------------------------------
019 NP_789739.1 --------------------------------------------------
020 NP_955619.1 --------------------------------------------------
021 YP_224289.1 --------------------------------------------------
022 YP_009182100.1 VINTVFQTIEGILNVNKQFMTAEKQREQILQSQLKQIKDQLAEPVVLSKP
023 NP_705927.1 --------------------------------------------------
024 YP_009508577.1 --------------------------------------------------
025 YP_004442830.1 --------------------------------------------------
026 NP_758887.1 --------------------------------------------------
027 YP_009508406.1 --------------------------------------------------
028 YP_610965.1 LSDHEVWVPPSHPAEAGGEPCPSTFCE-----------------------
029 YP_001931961.1 --------------------------------------------------
030 YP_241114.1 --------------------------------------------------
031 YP_006732334.1 --------------------------------------------------
032 YP_009041481.1 VPVE-------APEIRKTLEPEP---------------------------
033 YP_001036293.1 LTPTVRGTRGFGTLTAEASNMITKVRVEERPVNPVPTPTDHMEEDDEDAE
034 YP_605811.1 GEFYDADNSLF---------------------------------------
035 NP_955577.1 --------------------------------------------------
036 YP_003208050.1 --------------------------------------------------
037 NP_789740.1 --------------------------------------------------
038 YP_004442839.1 --------------------------------------------------
039 YP_009507791.1 --------------------------------------------------
040 YP_004581513.1 --------------------------------------------------
041 NP_043933.1 --------------------------------------------------
042 YP_009507248.1 --------------------------------------------------
043 YP_002117531.1 Q-------------------------------------------------
044 YP_004347415.1 --------------------------------------------------
045 NP_056891.1 --------------------------------------------------
046 YP_009513242.1 --------------------------------------------------
047 YP_009352866.1 E---------------DSLNPKL---------------------------
048 YP_009165750.1 --------------------------------------------------
049 NP_040840.1 --------------------------------------------------
050 YP_004222728.1 --------------------------------------------------
051 YP_009506264.1 --------------------------------------------------
052 YP_567050.1 --------------------------------------------------
053 YP_009508571.1 --------------------------------------------------
054 NP_056803.1 --------------------------------------------------
055 NP_955564.1 --------------------------------------------------
056 YP_009508411.1 MRPTLRGDQGFGSTSAAVFSIFE-----------------------DDDD
057 YP_009140788.1 FS------------------------------------------------
058 YP_009508410.1 --------------------------------------------------
059 NP_040333.1 --------------------------------------------------
060 YP_008992013.1 --------------------------------------------------
061 NP_041734.1 EEEE----------------------------------------------
062 NP_056895.1 --------------------------------------------------
063 YP_009109689.1 --------------------------------------------------
064 YP_002519387.1 --------------------------------------------------
065 YP_595725.1 --------------------------------------------------
066 YP_009508551.1 --------------------------------------------------
067 YP_001497148.1 --------------------------------------------------
068 NP_041186.1 --------------------------------------------------
069 NP_663784.1 --------------------------------------------------
070 YP_009508443.1 LHQTLRGDRAFGEVTSNYLAEA-----------STSYLADMCP--TFEED
071 YP_003987465.1 C-------------------------------------------------
072 NP_861410.1 --------------------------------------------------
073 YP_009345071.1 IKLIS---------------------------------------------
074 YP_009508582.1 --------------------------------------------------
075 NP_569141.1 --------------------------------------------------
076 YP_002916057.1 --------------------------------------------------
077 NP_777384.2 --------------------------------------------------
078 YP_233110.1 EEFYDADNSLF---------------------------------------
079 NP_542258.1 --------------------------------------------------

080 YP_009130664.1 FASTEGGGDKAPINN-----------------------------------
081 YP_009508566.1 --------------------------------------------------
082 YP_006607892.1 --------------------------------------------------
083 YP_009506270.1 --------------------------------------------------
084 NP_042513.1 --------------------------------------------------
085 YP_006907834.1 --------------------------------------------------
086 YP_004442827.1 --------------------------------------------------
087 YP_009408594.1 --------------------------------------------------
088 YP_009508561.1 --------------------------------------------------
089 NP_056907.1 --------------------------------------------------
090 YP_002308474.1 --------------------------------------------------
091 YP_007761644.1 --------------------------------------------------
092 NP_040550.1 --------------------------------------------------
093 NP_758808.1 --------------------------------------------------
094 YP_009345075.1 WDDK----------------------------------------------
095 NP_068729.1 --------------------------------------------------
096 NP_047255.1 --------------------------------------------------
097 YP_006273075.1 YKEEES--------------------------------------------
098 YP_009508888.1 --------------------------------------------------
099 NP_395469.1 --------------------------------------------------
100 YP_009508546.1 --------------------------------------------------
101 NP_056762.1 LGLTG---------------------------------------------
102 YP_009508556.1 --------------------------------------------------
103 NP_569153.1 QSPVPSPTQDFTNPFSEGGGESTYPYPSFQPPFDLQSDDSYGTLATWSEY
104 NP_043924.1 --------------------------------------------------
105 YP_009243641.1 --------------------------------------------------
106 YP_004300274.1 --------------------------------------------------
107 NP_056848.1 --------------------------------------------------
108 YP_006495799.1 SEDS------YEEVRRRDMENHN---------------------------
109 YP_009508409.1 --------------------------------------------------
110 NP_687035.1 --------------------------------------------------
111 NP_045937.2 TYPAX---------------------------------------------
112 NP_612577.1 --------------------------------------------------
113 NP_619548.1 --------------------------------------------------
114 NP_056880.1 --------------------------------------------------
115 YP_009121747.1 GWDE----------------------------------------------
116 YP_009113237.2 EEPETP--------------------------------------------
117 YP_009508408.1 LTPTLRGDRGFGALTAAAFIPT-----------SVPTEHNEADDEANEAD
118 NP_955579.1 --------------------------------------------------
119 YP_009229919.1 DDDD-----HPDDSQPRWDTSPEEDNE-----------------------
120 YP_004442833.1 --------------------------------------------------
121 NP_862833.2 --------------------------------------------------
122 NP_040563.1 --------------------------------------------------
123 YP_443922.1 --------------------------------------------------
124 NP_057933.2 --------------------------------------------------
125 YP_009116631.1 ADDE------DDVYNQKTWEKED---------------------------
126 YP_009513249.1 --------------------------------------------------
127 YP_223871.1 --------------------------------------------------
128 NP_127504.1 --------------------------------------------------
129 NP_044929.1 --------------------------------------------------
130 NP_057860.1 --------------------------------------------------
131 NP_040973.1 --------------------------------------------------
132 NP_777317.1 --------------------------------------------------
133 NP_054716.1 --------------------------------------------------
134 YP_001856242.1 --------------------------------------------------
135 NP_954565.2 --------------------------------------------------
136 NP_569150.1 --------------------------------------------------
137 NP_659397.1 --------------------------------------------------
138 YP_009513211.1 --------------------------------------------------
139 YP_233107.1 --------------------------------------------------
140 YP_009109692.1 --------------------------------------------------
141 NP_057849.4 --------------------------------------------------
142 NP_049560.1 --------------------------------------------------
143 YP_009506267.1 LQMTLRGDRGFGEMSRTRLPPVESDDG------SPSSPLSEEVRILFEDD
144 NP_039820.1 KKAQQALSSQAQEEP-----------------------------------
145 NP_056790.1 --------------------------------------------------
146 YP_002321513.1 FPNTTEWLDDTPLEFPEERLSYSQWFSEDPQVLAQPPWSQHSAKVCHVLD
147 NP_955591.1 --------------------------------------------------
148 YP_001956722.2 --------------------------------------------------
149 NP_056902.1 --------------------------------------------------
150 NP_056886.1 --------------------------------------------------
151 NP_041261.1 --------------------------------------------------

```
001 YP_004442836.1   --------------------------------------------------
002 NP_056728.1      --------------------------------------------------
003 YP_009508537.1   --------------------------------------------------
004 YP_004732983.2   --------------------------------------------------
005 YP_009508407.1   --------------------------------------------------
006 NP_040939.1      --------------------------------------------------
007 YP_003864102.1   --------------------------------------------------
008 YP_009002585.1   --ATQPQSPQSP-----------------------------------TWY
009 YP_003284237.1   --------------------------------------------------
010 YP_009268869.1   --------------------------------------------------
011 YP_009506251.1   --------------------------------------------------
012 NP_955611.1      --------------------------------------------------
013 YP_008567619.1   QDDTSPLLAETGGHLPN---------------------------------
014 NP_149413.1      --------------------------------------------------
015 YP_004442824.1   --------------------------------------------------
016 YP_001931967.1   --------------------------------------------------
017 YP_002455786.1   --------------------------------------------------
018 YP_009109694.1   --------------------------------------------------
019 NP_789739.1      --------------------------------------------------
020 NP_955619.1      --------------------------------------------------
021 YP_224289.1      --------------------------------------------------
022 YP_009182100.1   VSDPALEGHLKSINRGVKNLVERAELLIDYEELAAKIGKNISTQGIAVSE
023 NP_705927.1      --------------------------------------------------
024 YP_009508577.1   --------------------------------------------------
025 YP_004442830.1   --------------------------------------------------
026 NP_758887.1      --------------------------------------------------
027 YP_009508406.1   --------------------------------------------------
028 YP_610965.1      --------------------------------------------------
029 YP_001931961.1   --------------------------------------------------
030 YP_241114.1      --------------------------------------------------
031 YP_006732334.1   --------------------------------------------------
032 YP_009041481.1   --------------------------------------------------
033 YP_001036293.1   AYGDYLSYLSSRSFSREG------------------------------GE
034 YP_605811.1      --------------------------------------------------
035 NP_955577.1      --------------------------------------------------
036 YP_003208050.1   --------------------------------------------------
037 NP_789740.1      --------------------------------------------------
038 YP_004442839.1   --------------------------------------------------
039 YP_009507791.1   --------------------------------------------------
040 YP_004581513.1   --------------------------------------------------
041 NP_043933.1      --------------------------------------------------
042 YP_009507248.1   --------------------------------------------------
043 YP_002117531.1   --------------------------------------------------
044 YP_004347415.1   --------------------------------------------------
045 NP_056891.1      --------------------------------------------------
046 YP_009513242.1   --------------------------------------------------
047 YP_009352866.1   --------------------------------------------------
048 YP_009165750.1   --------------------------------------------------
049 NP_040840.1      --------------------------------------------------
050 YP_004222728.1   --------------------------------------------------
051 YP_009506264.1   --------------------------------------------------
052 YP_567050.1      --------------------------------------------------
053 YP_009508571.1   --------------------------------------------------
054 NP_056803.1      --------------------------------------------------
055 NP_955564.1      --------------------------------------------------
056 YP_009508411.1   DDDDFPSFAE----------------------------------------
057 YP_009140788.1   --------------------------------------------------
058 YP_009508410.1   --------------------------------------------------
059 NP_040333.1      --------------------------------------------------
060 YP_008992013.1   --------------------------------------------------
061 NP_041734.1      --------------------------------------------------
062 NP_056895.1      --------------------------------------------------
063 YP_009109689.1   --------------------------------------------------
064 YP_002519387.1   --------------------------------------------------
065 YP_595725.1      --------------------------------------------------
066 YP_009508551.1   --------------------------------------------------
067 YP_001497148.1   --------------------------------------------------
068 NP_041186.1      --------------------------------------------------
069 NP_663784.1      --------------------------------------------------
070 YP_009508443.1   PFYEDPTILTLQQVE-----------------------------------
```

```
071 YP_003987465.1    --------------------------------------------------
072 NP_861410.1       --------------------------------------------------
073 YP_009345071.1    --------------------------------------------------
074 YP_009508582.1    --------------------------------------------------
075 NP_569141.1       --------------------------------------------------
076 YP_002916057.1    --------------------------------------------------
077 NP_777384.2       --------------------------------------------------
078 YP_233110.1       --------------------------------------------------
079 NP_542258.1       --------------------------------------------------
080 YP_009130664.1    ----FPIFDKS---------------------------------------
081 YP_009508566.1    --------------------------------------------------
082 YP_006607892.1    --------------------------------------------------
083 YP_009506270.1    --------------------------------------------------
084 NP_042513.1       --------------------------------------------------
085 YP_006907834.1    --------------------------------------------------
086 YP_004442827.1    --------------------------------------------------
087 YP_009408594.1    --------------------------------------------------
088 YP_009508561.1    --------------------------------------------------
089 NP_056907.1       --------------------------------------------------
090 YP_002308474.1    --------------------------------------------------
091 YP_007761644.1    --------------------------------------------------
092 NP_040550.1       --------------------------------------------------
093 NP_758808.1       --------------------------------------------------
094 YP_009345075.1    --------------------------------------------------
095 NP_068729.1       --------------------------------------------------
096 NP_047255.1       --------------------------------------------------
097 YP_006273075.1    --------------------------------------------------
098 YP_009508888.1    --------------------------------------------------
099 NP_395469.1       --------------------------------------------------
100 YP_009508546.1    --------------------------------------------------
101 NP_056762.1       --------------------------------------------------
102 YP_009508556.1    --------------------------------------------------
103 NP_569153.1       DAMSQSNSPSSHSDAIQHLSFQHPSADTVLDFDRYSFTTSEDDVVQSAWI
104 NP_043924.1       --------------------------------------------------
105 YP_009243641.1    --------------------------------------------------
106 YP_004300274.1    --------------------------------------------------
107 NP_056848.1       --------------------------------------------------
108 YP_006495799.1    --------------------------------------------------
109 YP_009508409.1    --------------------------------------------------
110 NP_687035.1       --------------------------------------------------
111 NP_045937.2       --------------------------------------------------
112 NP_612577.1       --------------------------------------------------
113 NP_619548.1       --------------------------------------------------
114 NP_056880.1       --------------------------------------------------
115 YP_009121747.1    --------------------------------------------------
116 YP_009113237.2    --------------------------------------------------
117 YP_009508408.1    DEMEYLSYLSTAPFSG----------------------------------
118 NP_955579.1       --------------------------------------------------
119 YP_009229919.1    --------------------------------------------------
120 YP_004442833.1    --------------------------------------------------
121 NP_862833.2       --------------------------------------------------
122 NP_040563.1       --------------------------------------------------
123 YP_443922.1       --------------------------------------------------
124 NP_057933.2       --------------------------------------------------
125 YP_009116631.1    --------------------------------------------------
126 YP_009513249.1    --------------------------------------------------
127 YP_223871.1       --------------------------------------------------
128 NP_127504.1       --------------------------------------------------
129 NP_044929.1       --------------------------------------------------
130 NP_057860.1       --------------------------------------------------
131 NP_040973.1       --------------------------------------------------
132 NP_777317.1       --------------------------------------------------
133 NP_054716.1       --------------------------------------------------
134 YP_001856242.1    --------------------------------------------------
135 NP_954565.2       --------------------------------------------------
136 NP_569150.1       --------------------------------------------------
137 NP_659397.1       --------------------------------------------------
138 YP_009513211.1    --------------------------------------------------
139 YP_233107.1       --------------------------------------------------
140 YP_009109692.1    --------------------------------------------------
141 NP_057849.4       --------------------------------------------------
142 NP_049560.1       --------------------------------------------------
143 YP_009506267.1    PFLEDPTIYTIDSFNHS---------------------------------
```

```
144 NP_039820.1       --------------------------------------------------
145 NP_056790.1       --------------------------------------------------
146 YP_002321513.1    LNTGTSCKPVKPLKTEEQLRYERFERGKPTQADRDAVKYGIEKYWGLLGE
147 NP_955591.1       --------------------------------------------------
148 YP_001956722.2    --------------------------------------------------
149 NP_056902.1       --------------------------------------------------
150 NP_056886.1       --------------------------------------------------
151 NP_041261.1       --------------------------------------------------


001 YP_004442836.1    ------------------DDTKSQ-------VSQ---QVMNLEE-EDFPQ
002 NP_056728.1       --------------------------------------------------
003 YP_009508537.1    --------------------------------------------------
004 YP_004732983.2    ---------EKGKSPFEDLELKQDLVQSWIAQLGSGSGSRTEKPIFDTTS
005 YP_009508407.1    --------------------------------------------------
006 NP_040939.1       --------------------------------------------------
007 YP_003864102.1    --------------------------------------------------
008 YP_009002585.1    NP--FAEGGGSDEND-------ENPDPEESEEMGNEMGY-------ETEV
009 YP_003284237.1    -------------------ELEEE---------TTSMASLNIAMMSEYPE
010 YP_009268869.1    --------------------------------------------------
011 YP_009506251.1    --------------------EEEEEEEEEEEDEYDPNTY-------MAYL
012 NP_955611.1       --------------------------------------------------
013 YP_008567619.1    -------TNYEEEEAISPAVTTPEDIQDDLYQHDLAAIN------TEGPV
014 NP_149413.1       ------------------EEED-IITTFLSAIPDEEEAYPAEEIEEEYPA
015 YP_004442824.1    ------------------EDTRSH-------ISE---KVHHIED---YPS
016 YP_001931967.1    --------------------------------------------------
017 YP_002455786.1    --------------------------------------------------
018 YP_009109694.1    --------------------------------------------------
019 NP_789739.1       --------------------------------------------------
020 NP_955619.1       --------------------------------------------------
021 YP_224289.1       -----------TIQDFLNQYDADI---------SEDEANYDVGVLEEYPE
022 YP_009182100.1    RSKSFMSFDHPKPVESVLPKARMFSMFAGGKPKSSIVRSLELEVEEQPKV
023 NP_705927.1       --------------------------------------------------
024 YP_009508577.1    --------------------------------------------------
025 YP_004442830.1    ----------------CEDDEESIINDFLKLS--HEEVYMAEE-DSDYPA
026 NP_758887.1       --------------------------------------------------
027 YP_009508406.1    --------------------------------------------------
028 YP_610965.1       ----YDSNIEEQFLYDLECLYNAEQSDLASEEEPATVAVGDDACVLSPSQ
029 YP_001931961.1    --------------------------------------------------
030 YP_241114.1       --------------------------------------------------
031 YP_006732334.1    --------------------------------------------------
032 YP_009041481.1    ---------VSDEEVQWEVETIGTEVSDVTQQALP---------IYEEAA
033 YP_001036293.1    KSREVEAEADDDEDLYFRPSPARVTFSDDPFSEDPTIATIRESGDASQPI
034 YP_605811.1       ----------EEEEYDGDDDSEVS-------TPRSKYNIFALED--EYPK
035 NP_955577.1       --------------------------------------------------
036 YP_003208050.1    --------------------------DSDSEEEPDQGFINPFSEDGGGSK
037 NP_789740.1       --------------------------------------------------
038 YP_004442839.1    ----------EEEEYNGDNETETS-------TPRSKYNIFTFED--DYPQ
039 YP_009507791.1    --------------------------------------------------
040 YP_004581513.1    --------------------------------------------------
041 NP_043933.1       --------------------------------------------------
042 YP_009507248.1    --------------------------------------------------
043 YP_002117531.1    ----------VLIEDDRQRQAPARRAAHAEKRRRVEAAEQNRQNIEDFPD
044 YP_004347415.1    --------------------------------------------------
045 NP_056891.1       --------------------------------------------------
046 YP_009513242.1    --------------------------------------------------
047 YP_009352866.1    ---------ETDE--EWEVETIRTETTEVSQTVLP---------IYQEAQ
048 YP_009165750.1    --------------------------------------------------
049 NP_040840.1       --------------------------------------------------
050 YP_004222728.1    --------------------------------------------------
051 YP_009506264.1    ----------LTSPSNSDSETESL---------AESVRSLNIAAL-EYPD
052 YP_567050.1       --------------------------------------------------
053 YP_009508571.1    --------------------------------------------------
054 NP_056803.1       --------------------------------------------------
055 NP_955564.1       --------------------------------------------------
056 YP_009508411.1    ---------GGGEMDYFPFQIEESDEDPID---------------LAQLV
057 YP_009140788.1    ------------------------------SEAELELAY-------QQEV
058 YP_009508410.1    --------------------------------------------------
059 NP_040333.1       --------------------------------------------------
060 YP_008992013.1    -----------------KKKKKPK------KKSHASCHHSTSRPPQDPPP
061 NP_041734.1       -------------------------------DEYDPNVY-------RAYL
062 NP_056895.1       --------------------------------------------------
```

```
063 YP_009109689.1   --------------------------------------------------
064 YP_002519387.1   --------------------------------------------------
065 YP_595725.1      ------------------PEEDQIISDFLGKT---EEAYP-AEIEEEYPA
066 YP_009508551.1   --------------------------------------------------
067 YP_001497148.1   --------------------------------------------------
068 NP_041186.1      --------------------------------------------------
069 NP_663784.1      --------------------------------------------------
070 YP_009508443.1   --------EEDDDISYLTYLYDHHLFFGGEKEEEEAAAAKIFDEVAAAPI
071 YP_003987465.1   ----------KESPSCLFTDDNNG---------DDEGEHHVVAASLEYPE
072 NP_861410.1      --------------------------------------------------
073 YP_009345071.1   ----------------------------------------------KGDI
074 YP_009508582.1   --------------------------------------------------
075 NP_569141.1      --------------------------------------------------
076 YP_002916057.1   --------------------------------------------------
077 NP_777384.2      --------------------------------------------------
078 YP_233110.1      ----------EEEEYDG--DSEVS-------TPRSKYNIFALED--EYPK
079 NP_542258.1      --------------------------------------------------
080 YP_009130664.1   ----------DEEEEILPSSDS--------CRKEIASYI------AAAFP
081 YP_009508566.1   --------------------------------------------------
082 YP_006607892.1   --------------------------------------------------
083 YP_009506270.1   ------------------DDDEDS-------RSQ---HVMNLDE-EEFPE
084 NP_042513.1      --------------------------------------------------
085 YP_006907834.1   --------------------------------------------------
086 YP_004442827.1   ----------------EQDEEEAIINAFLKLTAEEEVLRIEDEEDCEYPA
087 YP_009408594.1   --------------------------------TDQTIAMLHDDDDWSNYP
088 YP_009508561.1   --------------------------------------------------
089 NP_056907.1      --------------------------------------------------
090 YP_002308474.1   --------------------------------------------------
091 YP_007761644.1   --------------------------------------------------
092 NP_040550.1      --------------------------------------------------
093 NP_758808.1      ------------------------------AIEETAFTLEEQFN------
094 YP_009345075.1   ------EEESQEQYDSEKEELAMATRETEIAIEEFSLEQVSTSSEEEEEE
095 NP_068729.1      --------------------------------------------------
096 NP_047255.1      --------------------------------------------------
097 YP_006273075.1   ----------------------QSDEEHYQPDAEAEFDY-------PLKR
098 YP_009508888.1   --------------------------------------------------
099 NP_395469.1      --------------------------------------------------
100 YP_009508546.1   --------------------------------------------------
101 NP_056762.1      ----------------WEDLDQDSIETEEITEWENPNQVLHREIRAYKSV
102 YP_009508556.1   --------------------------------------------------
103 NP_569153.1      SENLFRENTGNGEVHNLVPPRPDTPRGDEVKGTQESMAH-------TVAV
104 NP_043924.1      --------------------------------------------------
105 YP_009243641.1   --------------------------------------------------
106 YP_004300274.1   --------------------------------------------------
107 NP_056848.1      --------------------------------------------------
108 YP_006495799.1   ---------NLIAEGFTQEPRFPGLYAPARIEVIEEDHYEGDASEQSRSE
109 YP_009508409.1   --------------------------------------------------
110 NP_687035.1      --------------------------------------------------
111 NP_045937.2      -------------DPIDCPYEKSGTKTTQDVITTKNAEIMVTVNHTKIPM
112 NP_612577.1      --------------------------------------------------
113 NP_619548.1      --------------------------------------------------
114 NP_056880.1      --------------------------------------------------
115 YP_009121747.1   ----------------------YEDDPWRDDLEEGTVQIKEEDTEQSDKE
116 YP_009113237.2   -----------VEERLRRESYWNEEIDSDEEDPWGLFLN-------RNQY
117 YP_009508408.1   -------VEEGEDEGYFNLAAAEDYPTTIIDAAETSTIG-----FLSAPI
118 NP_955579.1      --------------------------------------------------
119 YP_009229919.1   ----EAESSDSFTEGGGSDNKSDNTPQKSQQEEDERVEVETASESESEKE
120 YP_004442833.1   ----------------KLEEEEEIINDFLKIS--EESVRMAEEEETDYPA
121 NP_862833.2      --------------------------------------------------
122 NP_040563.1      --------------------------------------------------
123 YP_443922.1      --------------------------------------------------
124 NP_057933.2      --------------------------------------------------
125 YP_009116631.1   ---------QEKRE--LELQGWKPTGRPGLYEMIPEQEEE--VYLRYEAE
126 YP_009513249.1   --------------------------------------------------
127 YP_223871.1      --------------------------------------------------
128 NP_127504.1      -----------------KYKGIPRPGSIAERKMQEEANLKKLYEEKDPFV
129 NP_044929.1      --------------------------------------------------
130 NP_057860.1      --------------------------------------------------
131 NP_040973.1      --------------------------------------------------
132 NP_777317.1      ------------DQNSNGQPDDDE-------VSTWSAQINHINDQSEPPA
133 NP_054716.1      --------------------------------------------------
134 YP_001856242.1   --------------------------------------------------
135 NP_954565.2      --------------------------------------------------
```

136 NP_569150.1 ------------------DDEEDS-------RSQ---YVMNLEDNEEFPQ
137 NP_659397.1 --------------------------------------------------
138 YP_009513211.1 --------------------------------------------------
139 YP_233107.1 -------------DEESYYGDLES-------EPRSHYEINHLDD--DYPE
140 YP_009109692.1 --------------------------------------------------
141 NP_057849.4 --------------------------------------------------
142 NP_049560.1 --------------------------------------------------
143 YP_009506267.1 -------FNDEEEMDYISYLYHLDDIPAN-----------------DAPI
144 NP_039820.1 -----IIEENIEENKQAQEEPTQEEIPTHKENQPEEIQNEEIHVFEEEPA
145 NP_056790.1 --------------------------------------------------
146 YP_002321513.1 ESGKFDYFVKYRTPPWATHSVCTINEDLPPAFNPDEWDDLPFEYTTDLHS
147 NP_955591.1 --------------------------------------------------
148 YP_001956722.2 --------------------------------------------------
149 NP_056902.1 --------------------------------------------------
150 NP_056886.1 --------------------------------------------------
151 NP_041261.1 --------------------------------------------------

001 YP_004442836.1 LQEIERLLST----------------------------------------
002 NP_056728.1 --------------------------------------------------
003 YP_009508537.1 --------------------------------------------------
004 YP_004732983.2 SDSDSLSDVSSKVLAYAGVE-------------------------EAVME
005 YP_009508407.1 --------------------------------------------------
006 NP_040939.1 --------------------------------------------------
007 YP_003864102.1 --------------------------------------------------
008 YP_009002585.1 EEQEEKVAMI----------------------------------------
009 YP_003284237.1 LKKISELVNA----------------------------------------
010 YP_009268869.1 --------------------------------------------------
011 YP_009506251.1 QKEEDDWTEI----------------------------------------
012 NP_955611.1 --------------------------------------------------
013 YP_008567619.1 WDNSEEEEE-----------------------------------------
014 NP_149413.1 IKRLEELMRTRVIVKEVPEKSE----------------------------
015 YP_004442824.1 LAKVEQVLS-----------------------------------------
016 YP_001931967.1 --------------------------------------------------
017 YP_002455786.1 --------------------------------------------------
018 YP_009109694.1 --------------------------------------------------
019 NP_789739.1 --------------------------------------------------
020 NP_955619.1 --------------------------------------------------
021 YP_224289.1 LLKLENNLAE----------------------------------------
022 YP_009182100.1 LKDYKVPKLSVKQLYEQGSMMQRNAYRIKQIERKIPYDPNTKVIHLVDKG
023 NP_705927.1 --------------------------------------------------
024 YP_009508577.1 --------------------------------------------------
025 YP_004442830.1 IRKLEELIKQ----------------------------------------
026 NP_758887.1 --------------------------------------------------
027 YP_009508406.1 --------------------------------------------------
028 YP_610965.1 SESKSGRELV----------------------------------------
029 YP_001931961.1 --------------------------------------------------
030 YP_241114.1 --------------------------------------------------
031 YP_006732334.1 --------------------------------------------------
032 YP_009041481.1 EQETIATQ-LADELSYKS--------------------------------
033 YP_001036293.1 FDESDDNED----------------------------------------H
034 YP_605811.1 LQQLESLVLS----------------------------------------
035 NP_955577.1 --------------------------------------------------
036 YP_003208050.1 DFEAKTQTAL----------------------------------------
037 NP_789740.1 --------------------------------------------------
038 YP_004442839.1 LQKLDNLVLS----------------------------------------
039 YP_009507791.1 --------------------------------------------------
040 YP_004581513.1 --------------------------------------------------
041 NP_043933.1 --------------------------------------------------
042 YP_009507248.1 --------------------------------------------------
043 YP_002117531.1 EEALMDLLRQ----------------------------------------
044 YP_004347415.1 --------------------------------------------------
045 NP_056891.1 --------------------------------------------------
046 YP_009513242.1 --------------------------------------------------
047 YP_009352866.1 TLKNEAVQGLAEEMSYKN--------------------------------
048 YP_009165750.1 --------------------------------------------------
049 NP_040840.1 --------------------------------------------------
050 YP_004222728.1 --------------------------------------------------
051 YP_009506264.1 AKAINEVIGM----------------------------------------
052 YP_567050.1 --------------------------------------------------
053 YP_009508571.1 --------------------------------------------------
054 NP_056803.1 --------------------------------------------------

```
055 NP_955564.1        --------------------------------------------------
056 YP_009508411.1     LEAEEEMLK-----------------------------------------
057 YP_009140788.1     FAAVEEEAE-----------------------------------------
058 YP_009508410.1     --------------------------------------------------
059 NP_040333.1        --------------------------------------------------
060 YP_008992013.1     -----PSAPLH---------------------------------------
061 NP_041734.1        QKEEDEWQEI----------------------------------------
062 NP_056895.1        --------------------------------------------------
063 YP_009109689.1     --------------------------------------------------
064 YP_002519387.1     --------------------------------------------------
065 YP_595725.1        LRRLEQLMKTKVVVQEIEEPS-----------------------------
066 YP_009508551.1     --------------------------------------------------
067 YP_001497148.1     --------------------------------------------------
068 NP_041186.1        --------------------------------------------------
069 NP_663784.1        --------------------------------------------------
070 YP_009508443.1     YDDYDDADDE-----------------------------------EEVQR
071 YP_003987465.1     LRKISEIINS----------------------------------------
072 NP_861410.1        --------------------------------------------------
073 YP_009345071.1     IGQGEVQQEE----------------------------------------
074 YP_009508582.1     --------------------------------------------------
075 NP_569141.1        --------------------------------------------------
076 YP_002916057.1     --------------------------------------------------
077 NP_777384.2        --------------------------------------------------
078 YP_233110.1        LQQLESLVLS----------------------------------------
079 NP_542258.1        --------------------------------------------------
080 YP_009130664.1     ENNAD---------------------------------------------
081 YP_009508566.1     --------------------------------------------------
082 YP_006607892.1     --------------------------------------------------
083 YP_009506270.1     MKEIERVLAS----------------------------------------
084 NP_042513.1        --------------------------------------------------
085 YP_006907834.1     --------------------------------------------------
086 YP_004442827.1     IKRLEELLKQ----------------------------------------
087 YP_009408594.1     KFRPIELIAA----------------------------------------
088 YP_009508561.1     --------------------------------------------------
089 NP_056907.1        --------------------------------------------------
090 YP_002308474.1     --------------------------------------------------
091 YP_007761644.1     --------------------------------------------------
092 NP_040550.1        --------------------------------------------------
093 NP_758808.1        --------------------------------------------------
094 YP_009345075.1     PDWENPFAVL----------------------------------------
095 NP_068729.1        --------------------------------------------------
096 NP_047255.1        --------------------------------------------------
097 YP_006273075.1     IHYLNPFSEG----------------------------------------
098 YP_009508888.1     --------------------------------------------------
099 NP_395469.1        --------------------------------------------------
100 YP_009508546.1     --------------------------------------------------
101 NP_056762.1        SEQIEDIFGELLKEHG----------------------------------
102 YP_009508556.1     --------------------------------------------------
103 NP_569153.1        TTEESKHEAE----------------------------------------
104 NP_043924.1        --------------------------------------------------
105 YP_009243641.1     --------------------------------------------------
106 YP_004300274.1     --------------------------------------------------
107 NP_056848.1        --------------------------------------------------
108 YP_006495799.1     EEPEFDYPQIAAVVNQNN---------------------------EPELE
109 YP_009508409.1     --------------------------------------------------
110 NP_687035.1        --------------------------------------------------
111 NP_045937.2        LVDTGACLTAIGGAATVVPDLKLTN-------------------------
112 NP_612577.1        --------------------------------------------------
113 NP_619548.1        --------------------------------------------------
114 NP_056880.1        --------------------------------------------------
115 YP_009121747.1     FTEKIEMIQMIVSPTQEEEEHY------------------AFPTIESIGE
116 YP_009113237.2     WDPEPVHEEP----------------------------------------
117 YP_009508408.1     YDTDGEEGD------------------------------------EDNDD
118 NP_955579.1        --------------------------------------------------
119 YP_009229919.1     GESQAGYMAN----------------------------------------
120 YP_004442833.1     IKRLEELLQQ----------------------------------------
121 NP_862833.2        --------------------------------------------------
122 NP_040563.1        --------------------------------------------------
123 YP_443922.1        --------------------------------------------------
124 NP_057933.2        --------------------------------------------------
125 YP_009116631.1     DEEEDQELQVIGAATMD----------------------------EPEME
126 YP_009513249.1     --------------------------------------------------
127 YP_223871.1        --------------------------------------------------
```

```
128 NP_127504.1         GSLSRPGKYEY---------------------------------------
129 NP_044929.1         --------------------------------------------------
130 NP_057860.1         --------------------------------------------------
131 NP_040973.1         --------------------------------------------------
132 NP_777317.1         LLQLEQVLS-----------------------------------------
133 NP_054716.1         --------------------------------------------------
134 YP_001856242.1      --------------------------------------------------
135 NP_954565.2         --------------------------------------------------
136 NP_569150.1         LREIEKVLSS----------------------------------------
137 NP_659397.1         --------------------------------------------------
138 YP_009513211.1      --------------------------------------------------
139 YP_233107.1         LLNVEKILS-----------------------------------------
140 YP_009109692.1      --------------------------------------------------
141 NP_057849.4         --------------------------------------------------
142 NP_049560.1         --------------------------------------------------
143 YP_009506267.1      YDDYPEEGE---------------------------------------VD
144 NP_039820.1         FKHLAAQLSELVN-------------------------------------
145 NP_056790.1         --------------------------------------------------
146 YP_002321513.1      SDEEKPVPDYLSMQTPKPKYYYGASSSSSSSESVASRYRPKKNEKVYKPK
147 NP_955591.1         --------------------------------------------------
148 YP_001956722.2      --------------------------------------------------
149 NP_056902.1         --------------------------------------------------
150 NP_056886.1         --------------------------------------------------
151 NP_041261.1         --------------------------------------------------


001 YP_004442836.1      -------TAESAISS-----------------------------------
002 NP_056728.1         --------------------------------------------------
003 YP_009508537.1      --------------------------------------------------
004 YP_004732983.2      YPRR-VKTATAKLADMEKAFAG-----------------ETTAAVGGDSE
005 YP_009508407.1      --------------------------------------------------
006 NP_040939.1         --------------------------------------------------
007 YP_003864102.1      --------------------------------------------------
008 YP_009002585.1      QEDPDLLPYPHLRELL-------------------------KNEEKVFST
009 YP_003284237.1      -------SSEVTSS------------------------------------
010 YP_009268869.1      --------------------------------------------------
011 YP_009506251.1      IAS-----LNPEMEYP-------------------------QRRTEVAET
012 NP_955611.1         --------------------------------------------------
013 YP_008567619.1      GYAAEEEEEIDDMEYP---------------------HVRALRLKMQ-EV
014 NP_149413.1         ----QPQEAKMASSEG----------------------------------
015 YP_004442824.1      -------TGESAISS-----------------------------------
016 YP_001931967.1      --------------------------------------------------
017 YP_002455786.1      --------------------------------------------------
018 YP_009109694.1      --------------------------------------------------
019 NP_789739.1         --------------------------------------------------
020 NP_955619.1         --------------------------------------------------
021 YP_224289.1         -------IPILSEVT-----------------------------------
022 YP_009182100.1      EMDRMKAKGYEYVHFGAVQVVFRLLARRDLNCSVLSICRDNRLKSLQKGL
023 NP_705927.1         --------------------------------------------------
024 YP_009508577.1      --------------------------------------------------
025 YP_004442830.1      ------EEVRMSDTG-----------------------------------
026 NP_758887.1         --------------------------------------------------
027 YP_009508406.1      --------------------------------------------------
028 YP_610965.1         ------FMELEALPYPTL---------------------ARMEK-----E
029 YP_001931961.1      --------------------------------------------------
030 YP_241114.1         --------------------------------------------------
031 YP_006732334.1      --------------------------------------------------
032 YP_009041481.1      -----LHKMQKKLVEN-------------------------MALSGAES-
033 YP_001036293.1      ELIQELVSLEADFDYP---------------------QLRQLED----CV
034 YP_605811.1         -------TTESAISR-----------------------------------
035 NP_955577.1         --------------------------------------------------
036 YP_003208050.1      ------MTSMEELPYPKL---------------------KKLEA-----A
037 NP_789740.1         --------------------------------------------------
038 YP_004442839.1      -------TNESAISS-----------------------------------
039 YP_009507791.1      --------------------------------------------------
040 YP_004581513.1      --------------------------------------------------
041 NP_043933.1         --------------------------------------------------
042 YP_009507248.1      --------------------------------------------------
043 YP_002117531.1      -------PCTETLEHPRENDDVSLNLAVLEEDNKDELYDYPTAIRNLENL
044 YP_004347415.1      --------------------------------------------------
045 NP_056891.1         --------------------------------------------------
046 YP_009513242.1      --------------------------------------------------
```

```
047 YP_009352866.1     -----LHKMQEKLAKN-------------------------MALSGVES-
048 YP_009165750.1     --------------------------------------------------
049 NP_040840.1        --------------------------------------------------
050 YP_004222728.1     --------------------------------------------------
051 YP_009506264.1     -------SSVTSDYRPP---------------------------------
052 YP_567050.1        --------------------------------------------------
053 YP_009508571.1     --------------------------------------------------
054 NP_056803.1        --------------------------------------------------
055 NP_955564.1        --------------------------------------------------
056 YP_009508411.1     --EVKLTTATLDLDYP---------------------ELEELQR-----I
057 YP_009140788.1     ----------LELAYP-------------------------QRFKAIEKL
058 YP_009508410.1     --------------------------------------------------
059 NP_040333.1        --------------------------------------------------
060 YP_008992013.1     --------------------------------------------------
061 NP_041734.1        TAS-----LREEMEYP-------------------------KRRPQ-TEM
062 NP_056895.1        --------------------------------------------------
063 YP_009109689.1     --------------------------------------------------
064 YP_002519387.1     --------------------------------------------------
065 YP_595725.1        ----QPVEAKMSTSTG----------------------------------
066 YP_009508551.1     --------------------------------------------------
067 YP_001497148.1     --------------------------------------------------
068 NP_041186.1        --------------------------------------------------
069 NP_663784.1        --------------------------------------------------
070 YP_009508443.1     LINPKLASISTEFEYP---------------------QLRELEQ-----V
071 YP_003987465.1     -------TVVDSNP------------------------------------
072 NP_861410.1        --------------------------------------------------
073 YP_009345071.1     ---VHFTEEEARAELR-----------------------KLIAGEEVHME
074 YP_009508582.1     --------------------------------------------------
075 NP_569141.1        --------------------------------------------------
076 YP_002916057.1     --------------------------------------------------
077 NP_777384.2        --------------------------------------------------
078 YP_233110.1        -------TTESAISR-----------------------------------
079 NP_542258.1        --------------------------------------------------
080 YP_009130664.1     ----------DEIDYP---------------------QLKNL---QQ-HL
081 YP_009508566.1     --------------------------------------------------
082 YP_006607892.1     --------------------------------------------------
083 YP_009506270.1     -------VSESAISS-----------------------------------
084 NP_042513.1        --------------------------------------------------
085 YP_006907834.1     --------------------------------------------------
086 YP_004442827.1     ------ETVQMSEAS-----------------------------------
087 YP_009408594.1     ------ETALASS-------------------------------------
088 YP_009508561.1     --------------------------------------------------
089 NP_056907.1        --------------------------------------------------
090 YP_002308474.1     --------------------------------------------------
091 YP_007761644.1     --------------------------------------------------
092 NP_040550.1        --------------------------------------------------
093 NP_758808.1        -----------DLDYP-----------------------TLISMEKQLVQ
094 YP_009345075.1     ------AADDNDMDYPEL---------------------RALQEIVRSTG
095 NP_068729.1        --------------------------------------------------
096 NP_047255.1        --------------------------------------------------
097 YP_006273075.1     GGKDDHL-NSEEMDYP-------------------------QRMKKIEKL
098 YP_009508888.1     --------------------------------------------------
099 NP_395469.1        --------------------------------------------------
100 YP_009508546.1     --------------------------------------------------
101 NP_056762.1        --------------------------------------------------
102 YP_009508556.1     --------------------------------------------------
103 NP_569153.1        FDYPAFARLQAHEESG-------------------------RPKPKTEKV
104 NP_043924.1        --------------------------------------------------
105 YP_009243641.1     --------------------------------------------------
106 YP_004300274.1     --------------------------------------------------
107 NP_056848.1        --------------------------------------------------
108 YP_006495799.1     YPTKKFQELMAKIKGFQNQK---------------------VEIAAMNAE
109 YP_009508409.1     --------------------------------------------------
110 NP_687035.1        --------------------------------------------------
111 NP_045937.2        ----------------------------------------TEIIAVGISA
112 NP_612577.1        --------------------------------------------------
113 NP_619548.1        --------------------------------------------------
114 NP_056880.1        --------------------------------------------------
115 YP_009121747.1     EAIKQDEVDEEDLDYPRVREIKRMIKKAKKEG-----YTPPIHQEQHIFP
116 YP_009113237.2     EAQEIVTVIQEELDYPHLKRLEE--------------ALKGKTVPVEKKE
117 YP_009508408.1     ILPKRIAVLSFEHDYP---------------------QLRGLEE---HVV
118 NP_955579.1        --------------------------------------------------
119 YP_009229919.1     ------EEEELELEYPTL---------------------QKLKEKV--EH
```

```
120 YP_004442833.1    ------EQVKMSEVSS----------------------------------
121 NP_862833.2       --------------------------------------------------
122 NP_040563.1       --------------------------------------------------
123 YP_443922.1       --------------------------------------------------
124 NP_057933.2       --------------------------------------------------
125 YP_009116631.1    YPTR-LEKVMGKLKNVSMEKL--------------------FPVSGMDSE
126 YP_009513249.1    --------------------------------------------------
127 YP_223871.1       --------------------------------------------------
128 NP_127504.1       --------------------------------------------------
129 NP_044929.1       --------------------------------------------------
130 NP_057860.1       --------------------------------------------------
131 NP_040973.1       --------------------------------------------------
132 NP_777317.1       -------TNTSAISR-----------------------------------
133 NP_054716.1       --------------------------------------------------
134 YP_001856242.1    --------------------------------------------------
135 NP_954565.2       --------------------------------------------------
136 NP_569150.1       -------VAETAISS-----------------------------------
137 NP_659397.1       --------------------------------------------------
138 YP_009513211.1    --------------------------------------------------
139 YP_233107.1       -------TNESAISN-----------------------------------
140 YP_009109692.1    --------------------------------------------------
141 NP_057849.4       --------------------------------------------------
142 NP_049560.1       --------------------------------------------------
143 YP_009506267.1    VFNLATLDSGFDLDYP---------------------TLQKLEA------
144 NP_039820.1       -------MAESSGQSGVG--------------------------------
145 NP_056790.1       --------------------------------------------------
146 YP_002321513.1    PKPIQTQKAQYQPKKPKPQKEQKVTKEWKPKF-----QVHKLSQDISTEC
147 NP_955591.1       --------------------------------------------------
148 YP_001956722.2    --------------------------------------------------
149 NP_056902.1       --------------------------------------------------
150 NP_056886.1       --------------------------------------------------
151 NP_041261.1       --------------------------------------------------
```

```
001 YP_004442836.1    --------YRPPDVDMSGIAPGYAPATSTT-------------------G
002 NP_056728.1       --------------------------------------------------
003 YP_009508537.1    --------------------------------------------------
004 YP_004732983.2    MTTGQSSRSTLIPPNEGGGPIRYPPAERPSTSAS------------TYNT
005 YP_009508407.1    --------------------------------------------------
006 NP_040939.1       --------------------------------------------------
007 YP_003864102.1    --------------------------------------------------
008 YP_009002585.1    QQSEVSS--YRPPQDTAMTPAGYAPARGEAS-----------------SS
009 YP_003284237.1    --------YRPPEIDMTGPA-GYAPATSQQ-------------------G
010 YP_009268869.1    --------------------------------------------------
011 YP_009506251.1    VFSEIVD--YTPPGDTLMTPVGYPPASSSRS--------------TVTTP
012 NP_955611.1       --------------------------------------------------
013 YP_008567619.1    FSAQSEISQYRPPQDTAMGPSSYPPAVN-----------------ITNEA
014 NP_149413.1       ----SSAIWQPGTMDLDGNFPGYAPAQGAT-------------------G
015 YP_004442824.1    --------YRPPEVEMGGVAPGYAPASGTS-------------------G
016 YP_001931967.1    --------------------------------------------------
017 YP_002455786.1    --------------------------------------------------
018 YP_009109694.1    --------------------------------------------------
019 NP_789739.1       --------------------------------------------------
020 NP_955619.1       --------------------------------------------------
021 YP_224289.1       ------SQYRPADIDMAGPT-GYAPATSQQ-------------------G
022 YP_009182100.1    LGALQASLNNQVAYFVCIPNFSCTIEDAHEALVLCVQTHGTDFKEGYTDL
023 NP_705927.1       --------------------------------------------------
024 YP_009508577.1    --------------------------------------------------
025 YP_004442830.1    ----SSAMIP-ADIDMEGDLPRYAPAQGQT-------------------G
026 NP_758887.1       --------------------------------------------------
027 YP_009508406.1    --------------------------------------------------
028 YP_610965.1       SARSSVTSPYTPPTDSGMAPPGYPPASAGG----------AASSSQMPIL
029 YP_001931961.1    --------------------------------------------------
030 YP_241114.1       --------------------------------------------------
031 YP_006732334.1    --------------------------------------------------
032 YP_009041481.1    ---SGSGAGVYNPYSMREDNSNRPPGYAAAQGLP------------QNIP
033 YP_001036293.1    IAASSAVSNYTPPEDTTMNPPTYPPARID--------------QAGPSHS
034 YP_605811.1       --------FRPTDTDMTGIGPAYAPATGTA-------------------G
035 NP_955577.1       --------------------------------------------------
036 YP_003208050.1    FSSGSVVSNYAPPQDSNMGPPTYPPAPTNG----------ASSSRPMPTL
037 NP_789740.1       --------------------------------------------------
038 YP_004442839.1    --------YNPGDIDMEGVGPGYAPATGGT-------------------S
```

```
039 YP_009507791.1   --------------------------------------------------
040 YP_004581513.1   --------------------------------------------------
041 NP_043933.1      --------------------------------------------------
042 YP_009507248.1   --------------------------------------------------
043 YP_002117531.1   FSSEVTGSYSPPDTDMVGQT-GYAPATSTQ-------------------G
044 YP_004347415.1   --------------------------------------------------
045 NP_056891.1      --------------------------------------------------
046 YP_009513242.1   --------------------------------------------------
047 YP_009352866.1   ---SGSGAGVYNPYDMREDTSNRPPGFAAAQGLR------------QNIP
048 YP_009165750.1   --------------------------------------------------
049 NP_040840.1      --------------------------------------------------
050 YP_004222728.1   --------------------------------------------------
051 YP_009506264.1   -----ALDYRPADVDMTGPS-GYAPSTSAS-------------------G
052 YP_567050.1      --------------------------------------------------
053 YP_009508571.1   --------------------------------------------------
054 NP_056803.1      --------------------------------------------------
055 NP_955564.1      --------------------------------------------------
056 YP_009508411.1   VGASTAMSEYSPPQDTSMNPPTYPPARRE-----------SAASAPSTST
057 YP_009140788.1   LSTSEVTGPYRPPDDAAMGRPSYPPARSISEE-------------GVSSS
058 YP_009508410.1   --------------------------------------------------
059 NP_040333.1      --------------------------------------------------
060 YP_008992013.1   -----LYKKELQWLAKRCTSTISSPVPDPSPPLS-----------CMMFS
061 NP_041734.1      AFSETVD--YTPPGDTMMTPVGYPPASSSRS--------------TVTTP
062 NP_056895.1      --------------------------------------------------
063 YP_009109689.1   --------------------------------------------------
064 YP_002519387.1   --------------------------------------------------
065 YP_595725.1      ----SSAMIP-ANMDMDGNMPGYAPAQEAR-------------------G
066 YP_009508551.1   --------------------------------------------------
067 YP_001497148.1   --------------------------------------------------
068 NP_041186.1      --------------------------------------------------
069 NP_663784.1      --------------------------------------------------
070 YP_009508443.1   IAASSAVSAYNPPEDTTMNPPTYPPAQTP----SMPSTSYGGSTSYGGTF
071 YP_003987465.1   -----GNQYMIGDTDMDGPP-GYAPAQAQA-------------------A
072 NP_861410.1      --------------------------------------------------
073 YP_009345071.1   AESSINSGSYRLPTDATMGPPIYPPARQRFD----------------EPS
074 YP_009508582.1   --------------------------------------------------
075 NP_569141.1      --------------------------------------------------
076 YP_002916057.1   --------------------------------------------------
077 NP_777384.2      --------------------------------------------------
078 YP_233110.1      --------FRPADTDMTGVGPGYAPATGTA-------------------G
079 NP_542258.1      --------------------------------------------------
080 YP_009130664.1   YASPSAITGYKPPQDVAMGPPNYGPTPLR---------------VIRPEA
081 YP_009508566.1   --------------------------------------------------
082 YP_006607892.1   --------------------------------------------------
083 YP_009506270.1   --------FRPPEVEMGGIAPAYAPATSTT-------------------G
084 NP_042513.1      --------------------------------------------------
085 YP_006907834.1   --------------------------------------------------
086 YP_004442827.1   -----SAIIP-AEVDMEGNLPGYAPATTAT-------------------G
087 YP_009408594.1   -STTIAMSPYNPPEDTMMGPPQYPPATARPN---------MPNPWAVPDY
088 YP_009508561.1   --------------------------------------------------
089 NP_056907.1      --------------------------------------------------
090 YP_002308474.1   --------------------------------------------------
091 YP_007761644.1   --------------------------------------------------
092 NP_040550.1      --------------------------------------------------
093 NP_758808.1      SS---VTSAYNPPTEPLMGQVVYPPASAPRP---------QAETSS---T
094 YP_009345075.1   DSSSSIVTNLQPPMDSAMGPPVFPPAREGSR---------PTYLPGNLGF
095 NP_068729.1      --------------------------------------------------
096 NP_047255.1      --------------------------------------------------
097 YP_006273075.1   LSTSEVTSPYRPPEDAAMGRPSYPPARTIPGG-------------GASSS
098 YP_009508888.1   --------------------------------------------------
099 NP_395469.1      --------------------------------------------------
100 YP_009508546.1   --------------------------------------------------
101 NP_056762.1      -NYDMALKNLEEKYDLDKIEKAKSIEEIAKSSTSSEIRPTKRPKEEQTAY
102 YP_009508556.1   --------------------------------------------------
103 NP_569153.1      LSSAISS--YTPPTDTAMTPVAYPPAQNIASP-------------SYNPS
104 NP_043924.1      -QVKGKGCWRISQELAENILGNDLLRSLGLIVDQCNGVLWQASEGLGPDN
105 YP_009243641.1   --------------------------------------------------
106 YP_004300274.1   --------------------------------------------------
107 NP_056848.1      --------------------------------------------------
108 YP_006495799.1   SSGSGVTSGNFVPPIATTGPSVYPPAT--GTHGT------------NIGP
109 YP_009508409.1   --------------------------------------------------
110 NP_687035.1      --------------------------------------------------
111 NP_045937.2      EPVPHVLAKPTKIQIENTNIDISPWYNPDQTFHILGRDTLSKMRAIVSFE
```

```
112 NP_612577.1       --------------------------------------------------
113 NP_619548.1       --------------------------------------------------
114 NP_056880.1       --------------------------------------------------
115 YP_009121747.1    STSEGASGVYNPPPDQVMGPPVYPPARGNVP---------QNSFNLGYPK
116 YP_009113237.2    TVASAVSSAYNPPADTAMMPPAYPPGSADPG------------------P
117 YP_009508408.1    AATSTAVSAYRPPEDTTMNPPTYPPASNLPSAGSVPSTSYP-STSYPTTQ
118 NP_955579.1       --------------------------------------------------
119 YP_009229919.1    MAKSSASSPYRPPQEPYMGTYIYPPASGIT----------EASTSTASTM
120 YP_004442833.1    ----SSALIP-ADVDMEGNLPGYAPAQTKA-------------------G
121 NP_862833.2       --------------------------------------------------
122 NP_040563.1       --------------------------------------------------
123 YP_443922.1       -------------------------------------------------D
124 NP_057933.2       --------------------------------------------------
125 YP_009116631.1    SSITGGGGG-FIPPSPVPGAQGYPPAT--TSTMS------------TIGP
126 YP_009513249.1    --------------------------------------------------
127 YP_223871.1       --------------------------------------------------
128 NP_127504.1       -----LVRYDAPSWAKDPHLTVEPTGWDSDEPIPPKQPFTTRNTLPKIYM
129 NP_044929.1       --------------------------------------------------
130 NP_057860.1       --------------------------------------------------
131 NP_040973.1       --------------------------------------------------
132 NP_777317.1       --------YRPADEDMTGIGPAYTPATSTR-------------------G
133 NP_054716.1       --------------------------------------------------
134 YP_001856242.1    --------------------------------------------------
135 NP_954565.2       --------------------------------------------------
136 NP_569150.1       --------YRPPDAEMGEEAPAYAPATSAT-------------------G
137 NP_659397.1       --------------------------------------------------
138 YP_009513211.1    --------------------------------------------------
139 YP_233107.1       --------YRVP-QDEEMTVPGYAPAGSSR-------------------G
140 YP_009109692.1    --------------------------------------------------
141 NP_057849.4       --------------------------------------------------
142 NP_049560.1       --------------------------------------------------
143 YP_009506267.1    TIASTAVSEYRPPEDTTMNPPIYPPARIS---GQASTSTGPTSTSYTSGP
144 NP_039820.1       ---FQPPVNAQPDVNMEGPA-GYAPATSQA-------------------T
145 NP_056790.1       --------------------------------------------------
146 YP_002321513.1    PNLDKFLKTFEPKKKEPTPPVEKIFSASSSYHVPEDSSMGDTAYTPAPNM
147 NP_955591.1       --------------------------------------------------
148 YP_001956722.2    --------------------------------------------------
149 NP_056902.1       --------------------------------------------------
150 NP_056886.1       --------------------------------------------------
151 NP_041261.1       --------------------------------------------------


001 YP_004442836.1    WAG-NNQFPFTTRR-PRRWDSNNEFYMLPPAQSKQGAMFIMPLDFDV--K
002 NP_056728.1       --------------------------------------------------
003 YP_009508537.1    --------------------------------------------------
004 YP_004732983.2    TAPPLFEGTVR----PGRYGRPLAPWSLPSAQHSQGALLILPPEVASHAD
005 YP_009508407.1    --------------------------------------------------
006 NP_040939.1       --------------------------------------------------
007 YP_003864102.1    --------------------------------------------------
008 YP_009002585.1    THQPMFEGYMPRRP-QFKQDRSTEYWQFPTAQGQTGAMFVIPRQIGLFHD
009 YP_003284237.1    WAS---SHPPTSGRN-FRYKDPSGYFALPSAQQQNGSILLLSPTYDT--K
010 YP_009268869.1    --------------------------------------------------
011 YP_009506251.1    SRPPLYEGRIPSASRVLRRDDYTEWWQLPSSQGTTGALFVMPKQIGLFHD
012 NP_955611.1       --------------------------------------------------
013 YP_008567619.1    GPSRPMFEGYNKSGVRFKHKDLADNWTLPSAQQQQGAVFVIPTQLGLFDS
014 NP_149413.1       WDNSASSSKKNYGGYSRKFKNESEFFNLPSAMATSGAILVLTMNNYA--K
015 YP_004442824.1    WAG-TSSQLFPFPRRPKKWDDSSEWFALPPAQSRQGAMFVMPYDFDV--K
016 YP_001931967.1    --------------------------------------------------
017 YP_002455786.1    --------------------------------------------------
018 YP_009109694.1    --------------------------------------------------
019 NP_789739.1       --------------------------------------------------
020 NP_955619.1       --------------------------------------------------
021 YP_224289.1       LLG--STAPDRLGKGSFKWKSPTEYFNLPSAQQQAGAMFVMPANFDP--K
022 YP_009182100.1    AIEYIVTYKFMTSDMEAMSKLKATEGFNSMVISDRNNHTISHTKKIDWKE
023 NP_705927.1       --------------------------------------------------
024 YP_009508577.1    --------------------------------------------------
025 YP_004442830.1    WS-DYNPQQKHFGGTSKKWSNPVNHWNLPSANAANGSMLVLSIGKNN--D
026 NP_758887.1       --------------------------------------------------
027 YP_009508406.1    --------------------------------------------------
028 YP_610965.1       GAPFTAADVRANLKNLRGSRSNAGMFQLPSAQQLRGALLVLPEDIGLYQD
029 YP_001931961.1    --------------------------------------------------
030 YP_241114.1       --------------------------------------------------
```

```
031 YP_006732334.1    --------------------------------------------------
032 YP_009041481.1    DVSHFTASARPPPKSKQPYGAP-AMWNLPSAQQTQGVMLVLPYDIGKYAE
033 YP_001036293.1    GPSYS-----VPSGTRFKPMNYSANLQLPSAQQTTGAMFYMPLELDKFDE
034 YP_605811.1       YTG-ASSSDFPYPRRPRKWDNNSEWFNLPTANARQASIFVMPQDFDT--K
035 NP_955577.1       --------------------------------------------------
036 YP_003208050.1    G----LEGARPKVSDFR-RRETSNQWNLPSAQQVNGAMLVLPEDIGLYEE
037 NP_789740.1       --------------------------------------------------
038 YP_004442839.1    YTG-AGTSDFPYPRRPRKWENSSEWFNLPMANARQASVFIMPQDFDT--K
039 YP_009507791.1    --------------------------------------------------
040 YP_004581513.1    --------------------------------------------------
041 NP_043933.1       --------------------------------------------------
042 YP_009507248.1    --------------------------------------------------
043 YP_002117531.1    WVG-SHKFEGSTSKARFKWKDPSEMFTLPSAYQVEGAIFVMPPDYDP--K
044 YP_004347415.1    --------------------------------------------------
045 NP_056891.1       --------------------------------------------------
046 YP_009513242.1    --------------------------------------------------
047 YP_009352866.1    EVSNFVGGARPPPKSKVPYGAP-ANWNLPSAQQSQGVMLVLPYDIGMYAE
048 YP_009165750.1    --------------------------------------------------
049 NP_040840.1       --------------------------------------------------
050 YP_004222728.1    --------------------------------------------------
051 YP_009506264.1    YKE---GERSFGFRGGFRWKNPSENFQLPSAQQQTGAIFIMPPNFDP--K
052 YP_567050.1       --------------------------------------------------
053 YP_009508571.1    --------------------------------------------------
054 NP_056803.1       --------------------------------------------------
055 NP_955564.1       --------------------------------------------------
056 YP_009508411.1    GISSG--APTVSAGTRFRGKDYSGAWTLPSAQQTTGAIFNMPLELGKFDE
057 YP_009140788.1    NYPSKFN----AARTRFRGGYNDEMWTLPSAQQKNGAMFVLPEQLGLFND
058 YP_009508410.1    --------------------------------------------------
059 NP_040333.1       --------------------------------------------------
060 YP_008992013.1    SASSDYSSYFPPLDTHTDSQRNVVSKPFIPSPITSAGHLEPPKPFESVLN
061 NP_041734.1       SRPPLFEGRTTHVPRFLKRDEYTEWWQLPSSQGTTGALFVMPKQMGLFHE
062 NP_056895.1       --------------------------------------------------
063 YP_009109689.1    --------------------------------------------------
064 YP_002519387.1    --------------------------------------------------
065 YP_595725.1       WD-SGETSRRNYGGHSRKWKDESQFFNLPSAMATSGAMLVLTMGNYA--K
066 YP_009508551.1    --------------------------------------------------
067 YP_001497148.1    --------------------------------------------------
068 NP_041186.1       --------------------------------------------------
069 NP_663784.1       --------------------------------------------------
070 YP_009508443.1    GGTSG-----VGSGTRFRSKDYSGNWNLPSAQQQTGAMLYIPLELGKFDE
071 YP_003987465.1    AVPPTYGGTSPSVRGPFRWKNPHEHFELPSAHQLSGAIFVMPQNFDP--K
072 NP_861410.1       --------------------------------------------------
073 YP_009345071.1    TSTVPMYPPQYRAPLRAKVPDFQNWWNLPTAQINGGAMFIIPSDFSKFDD
074 YP_009508582.1    --------------------------------------------------
075 NP_569141.1       --------------------------------------------------
076 YP_002916057.1    --------------------------------------------------
077 NP_777384.2       --------------------------------------------------
078 YP_233110.1       YTG-ASSSDFPYPRRPRKWDNNSEWFNLPTANARQASIFVMPQDFDT--K
079 NP_542258.1       --------------------------------------------------
080 YP_009130664.1    GPLRPAFEGYKPGEVRYKAKDYSEWWNLPSAQHGTGAIFIIPTQLGMFND
081 YP_009508566.1    --------------------------------------------------
082 YP_006607892.1    --------------------------------------------------
083 YP_009506270.1    WAG-TTEFPFTTARRPRVWDSNNEFYMLPPAQSRQGAIFIMPMDFDV--K
084 NP_042513.1       --------------------------------------------------
085 YP_006907834.1    --------------------------------------------------
086 YP_004442827.1    WS-DYNPKINRIGGTSSKWNNQTNYWNLPNANAVNGSMLILCYGKNS--E
087 YP_009408594.1    AT-------RNAGRARFQGGIPGEQWQLPSAQQIAGAMLVLPDDIGMYND
088 YP_009508561.1    --------------------------------------------------
089 NP_056907.1       --------------------------------------------------
090 YP_002308474.1    --------------------------------------------------
091 YP_007761644.1    --------------------------------------------------
092 NP_040550.1       --------------------------------------------------
093 NP_758808.1       SERF-------KNFRAKP--YSTPTIFLPPAYNQQGAILVLPDDIGLYED
094 YP_009345075.1    GDG-----LAXPKRYPTARLPTQNFWELPSAQINRGALLILPNDIGKYDD
095 NP_068729.1       --------------------------------------------------
096 NP_047255.1       --------------------------------------------------
097 YP_006273075.1    AEPPRFD----AARTRFKGGYNDEMWTLPSAQQKNGAMFVIPEQLGLFND
098 YP_009508888.1    --------------------------------------------------
099 NP_395469.1       --------------------------------------------------
100 YP_009508546.1    --------------------------------------------------
101 NP_056762.1       EDDMRDDWKRKELTVNPIEASKDRNFERIGSSYKKNFYPSRSEILNLDNV
102 YP_009508556.1    --------------------------------------------------
103 NP_569153.1       PQMPMFEGYYPKRP-NFKRD-NHAFISLPSAQQNTGALFIMPQQIGLFHE
```

```
104 NP_043924.1       WMAAETLRIYSIKSPGHYNLPELLATKDEQLADILWNNVEAFATHRNDCG
105 YP_009243641.1    --------------------------------------------------
106 YP_004300274.1    --------------------------------------------------
107 NP_056848.1       --------------------------------------------------
108 YP_006495799.1    QDQGGWGGRMPRSRMPGGYGRPQRPWTLPSAQTENGVMLIIPEDLTLAAD
109 YP_009508409.1    --------------------------------------------------
110 NP_687035.1       --------------------------------------------------
111 NP_045937.2       KNGEMTVLLPPTYHKQLSCQTKNTLNIDEYLLQFPDQLWASLPTDIGRML
112 NP_612577.1       --------------------------------------------------
113 NP_619548.1       --------------------------------------------------
114 NP_056880.1       --------------------------------------------------
115 YP_009121747.1    YARE-------SDFNGDYGRYHSSQWTLPPALTDSGAMLVLPADPGLWSD
116 YP_009113237.2    STTPIFKG-YDGAKVKIKREEGSEYWQLPSAQNQTGALFVIPRQLGMFDE
117 YP_009508408.1    GATSG-----VGSGTRFRAKDYSGNWNLPSAQQTTGALFYIPLELGKFDE
118 NP_955579.1       --------------------------------------------------
119 YP_009229919.1    R--------LPRLRHQG-RDPTVNWWSLPPAQNQQGAMLILPQDIGLYTD
120 YP_004442833.1    YS-DYNPGETRFGGYSRKWSNPINNWSLPVANAATGSMLVLSIGKNS--E
121 NP_862833.2       --------------------------------------------------
122 NP_040563.1       --------------------------------------------------
123 YP_443922.1       TTEICKPERLSLPRHHGKPYSDHPSATALLDSIDEAIWTTSPFDVGRLAV
124 NP_057933.2       --------------------------------------------------
125 YP_009116631.1    ADMQGWGGRVPRSRSPLGYGRPQQPWSLPSAQSDNGCMLVLPQDFTLIPD
126 YP_009513249.1    --------------------------------------------------
127 YP_223871.1       --------------------------------------------------
128 NP_127504.1       FNPLNYENNFPPLSSFSKDGADHTPKIPKRNVVLPSGAKDPTGDLEATVN
129 NP_044929.1       --------------------------------------------------
130 NP_057860.1       --------------------------------------------------
131 NP_040973.1       --------------------------------------------------
132 NP_777317.1       WTG-ENS-NFPPVTAPRRWDNSSERFQLPSAQGTHGAIFVMPYDFDT--K
133 NP_054716.1       --------------------------------------------------
134 YP_001856242.1    --------------------------------------------------
135 NP_954565.2       --------------------------------------------------
136 NP_569150.1       WAG-SRPFPFMPKGGPRRWDSNNEFYSLPPAQSRQGAMFVMPMDFDI--K
137 NP_659397.1       --------------------------------------------------
138 YP_009513211.1    --------------------------------------------------
139 YP_233107.1       WAT-NIDDAMLHQRKPKRWDNSSEWFQLPSANARQGSIFVMPYDFDV--K
140 YP_009109692.1    --------------------------------------------------
141 NP_057849.4       --------------------------------------------------
142 NP_049560.1       --------------------------------------------------
143 YP_009506267.1    GPSSSGYSQRTPSGVHFRARDYSNNWSLPSAQQNTGAMFYLPLELVKFDE
144 NP_039820.1       WSN---GVNIPVKSANFRWKGPVGNFQLPSAQGKDGAMLVFGMNYSP--E
145 NP_056790.1       --------------------------------------------------
146 YP_002321513.1    ITPYNPPGVDARRGRARGRPYVPPTFNLPLAGNDTGLMLELS-DPSMYNE
147 NP_955591.1       --------------------------------------------------
148 YP_001956722.2    --------------------------------------------------
149 NP_056902.1       --------------------------------------------------
150 NP_056886.1       --------------------------------------------------
151 NP_041261.1       --------------------------------------------------
```

```
001 YP_004442836.1    VFERWESITLLHMTE--RSFDNADDKMRYMENLLGEDEKKHFIEWRM--K
002 NP_056728.1       --------------------------------------------------
003 YP_009508537.1    -------------PTPQKQYHINPKAKPDIQIVINDLLKQGVLIQK----
004 YP_004732983.2    AITTWETITLNHLMN--ISFDSLQDRVDYIENLLGPREREAWVTWRM--A
005 YP_009508407.1    --------------------------------------------------
006 NP_040939.1       -------------GPHVPQWPLTEEKLKGLTEIIDKLVEEGKLGKAP--P
007 YP_003864102.1    -------------PVWVDQWPLTKEKLTAAQQLVQEQLQAGHIVES----
008 YP_009002585.1    AFNRWESITKNYVAS--QGFTDSKDKAEFMENLLGETEKLTWVQWRM--N
009 YP_003284237.1    IFERWESTTLNHMAD--KSFATAEDKLIYLENLLGEMEKKTFQTWRM--A
010 YP_009268869.1    -------------GPHIAQWPLTQEKLEGLKEIVDKLEKEGKVGRAP--P
011 YP_009506251.1    VFSRWESITKNYVSA--QGITDPTEKMEFIENLLGETEKLTWIQWRM--R
012 NP_955611.1       -------------PVWIDQWPLPEGKLVALTQLVEKELQLGHIEPS----
013 YP_008567619.1    VFARWESITKNLVAA--QGFTDPADKVDFIENLLGEAEKLTWIQWRS--T
014 NP_149413.1       EFERWQSINTNLLAT--QSFETADDKITRIENLLGETEKLMFQTWRM--A
015 YP_004442824.1    VFERWESITLLYLSE--RSFDTADDKLRTVENLLGENEKKMFIAWRM--K
016 YP_001931967.1    --------------------------------------------------
017 YP_002455786.1    -------------APRGQPVSFKPERLQALIDLVSKALEAGHIEPY----
018 YP_009109694.1    -----------AVPVSIKQYPMSLEAHMGIRQHIIKFLELGVLRPC----
019 NP_789739.1       -------------GPKVKQWPLTEEKIKALVEICTEMEKEGKISKIG--P
020 NP_955619.1       -------------PPEISQFPLNPERLQALQHLVRKALEAGHIEPY----
021 YP_224289.1       VFERWESITLNHMAD--KVFSTAEDKLIYLENLLGEAEKIMFQSWRM--T
022 YP_009182100.1    LNQSEFPKEWSINSEEPPEAKSSSVITGIYQHENGNISMRFHQAGEG--S
```

```
023 NP_705927.1      -------------GPKVKQWPLTEEKIKALVEICTEMEKEGKISKIG--P
024 YP_009508577.1   -------------PRPQRQYHINTKAKPSIQQVIDDLLKQGVLIKQ----
025 YP_004442830.1   IFERWESSTLNYMAS--QNISGSEEKIARIENLLGETEKKTFIGWRM--A
026 NP_758887.1      -------------GPKIKQWPLSKEKILALQEICSQMEKEGQISKIG--P
027 YP_009508406.1   --------------------------------------------------
028 YP_610965.1      VISRWETITLNVLAD--KVFSDNRSKLMYVENLLGEDEKKAWIQWRM--A
029 YP_001931961.1   --------------------------------------------------
030 YP_241114.1      -------------PVWVDQWPLTQEKLSAAQQLVQEQLRLGHIEPS----
031 YP_006732334.1   --------------------------------------------------
032 YP_009041481.1   VINRWESINVNLVNS--MNWDSNQDKVDFMENLLGETEKQIFVSWRM--Q
033 YP_001036293.1   VFSRWESITKNITSQ--HSFISGREKADFIENLLGETEKLTWIQWRT--T
034 YP_605811.1      VFERWESSVLLHMSD--KVFDDPQDKLTYVENLLGESEKKMFITWRM--M
035 NP_955577.1      -----------ATPISIRQYPMPHEAYQGIKPHIRRMLDQGILKPC----
036 YP_003208050.1   VISRWESITINLLND--RTWDNNKARVIYVENLLGESEKKIWQQWRT--T
037 NP_789740.1      -------------GPKVKQWPLTEEKIKALVEICTEMEKEGKISKIG--P
038 YP_004442839.1   VFERWESSVLLHMTD--KIFDDPQDKLTYIENLLGESEKKMFITWRM--K
039 YP_009507791.1   -------------GPKIKQWPLTNEKIEALTEIVERLETEGKVKRAD--P
040 YP_004581513.1   --------------------------------------------------
041 NP_043933.1      --------------------------------------------------
042 YP_009507248.1   --------------VYTKSYRYPFIHRQEVRDQITKMLDQGIIRPSDSAW
043 YP_002117531.1   VFSRWESITLNHMAP--KAFANATDKITYIENLLGENEKITFQTWRM--S
044 YP_004347415.1   --------------------------------------------------
045 NP_056891.1      -------------PVWVDQWPLTNDKLAAAQQLVQEQLEAGHITES----
046 YP_009513242.1   -------------PRPQKQYPINPKAKPSIQIVIDDLLKQGVLIQQ----
047 YP_009352866.1   VINRWESINVNVVNS--MNWDSNQDKVDFIENLLGETEKQIVVSWRM--Q
048 YP_009165750.1   --------------------------------------------------
049 NP_040840.1      -------------GPHIAQWPLTQEKLEGLKEIVDRLEKEGKVGRAP--P
050 YP_004222728.1   -------------PVWIDQWPLPEGKLVALTQLVEKELQLGHIEPS----
051 YP_009506264.1   VFERWESVVLNFLAD--KNFPTAEDKLIYIENMLGEMEKITFQTWRM--A
052 YP_567050.1      -------------PPQIDQFPFKPERLQALTDLVSKALEASYIEPY----
053 YP_009508571.1   -------------PRPQKQYPINPKARSSIQVVIDDLLKQGVLVQQ----
054 NP_056803.1      -------------PRPQKQYPINPKAKPSIQIVIDDLLKQGVLTPQ----
055 NP_955564.1      -------------PVWLNQWPLKQEKLQALQQLVTEQLQLGHLEES----
056 YP_009508411.1   VFMRWESVTKNLTAQ--HVFTSGRDKAEFIENLLGEYEKLTWLQWRT--A
057 YP_009140788.1   AFSRWESVTKNHVAT--QSFTDVRDKMEYMENLLGEIEKLIWIQWRM--T
058 YP_009508410.1   --------------------------------------------------
059 NP_040333.1      -----------STPVSIKQYPMSQEARLGIKPHIQRLLDQGILVPC----
060 YP_008992013.1   WQTQNARAQNDTLLNINSKVEKISLRTEHLET---KVDSISAQLQQI--H
061 NP_041734.1      VFSRWESITKNYVAA--QGFTDPTEKMEFMENLLGETEKLTWIQWRM--N
062 NP_056895.1      -------------PPEVPQFPLNLERLQALQDLVHRSLEAGYISPW----
063 YP_009109689.1   -----------AVPVSIKQYPMSLEAHMGIRQHIIKFLELGVLRPC----
064 YP_002519387.1   --------------------------------------------------
065 YP_595725.1      EFDRWQSINTNLLAS--QTFENAEDKITRIENLLGETEKLMFQTWRM--A
066 YP_009508551.1   -------------PRPQKQYPINPKAKPSIQIVIDDLLKQGVLTPQ----
067 YP_001497148.1   -----------AMPVSIRQYPMSKEAHMGIQPHITRFLELGVLRPC----
068 NP_041186.1      --------------------------------------------------
069 NP_663784.1      -------------GPRLKQWPLTKEKIEALKEICEKMEKEGQLEEAP--P
070 YP_009508443.1   VFMRWESITKNIISQ--HTFISGKEKADYIENLLGEYEKLTWIQWRT--A
071 YP_003987465.1   VFGRWESIVLNHLAD--RNFPTALDKIIYVENLLGEMEKITFQTWRM--R
072 NP_861410.1      --------------------------------------------------
073 YP_009345071.1   VFMRWESITKNVVAL--QNFSDNKDKAEFIENLLGETEKLAWMQWRS--I
074 YP_009508582.1   -------------PRPQRQYHINTKAKPSIQQVIDDLLKQGVLVKQ----
075 NP_569141.1      --------------------------------------------------
076 YP_002916057.1   --------------------------------------------------
077 NP_777384.2      -------------PPEVPQFPLNLERLQALQDLVHRSLEAGYISPW----
078 YP_233110.1      VFERWESSVLLHMSD--KVFDDPQDKLTYVENLLGESEKKMFITWRM--M
079 NP_542258.1      -------------APGGLAVPVKPERLQALTDLVSRALEAKHIEPY----
080 YP_009130664.1   TFMRWESITKNLVSQ--QGFTDPNDKVEFIENLLGEAEKIAWIQWRM--T
081 YP_009508566.1   -------------PKPQKQYHINPKAKPSIQIVINDLLKQGVLKQQ----
082 YP_006607892.1   --------------------------------------------------
083 YP_009506270.1   VFERWESITLLHMAE--RSFDNADDKMRYIENLLGEDEKKHFVEWRM--K
084 NP_042513.1      --------------------------------------------------
085 YP_006907834.1   --------------------------------------------------
086 YP_004442827.1   IFERWESSTLNYMAA--QNISGGEEKISRIENLLGESEKRTFIGWRT--A
087 YP_009408594.1   VISRWESITTNLVDS--KVWGDVTTKINFIENLLGEIEKKLFVQWRM--M
088 YP_009508561.1   -------------PKPQKQYHINPKAKPSIQIVINDLLKQGVLRQS----
089 NP_056907.1      -------------PPQIDQFPFKPERLQALTDLVSKALEASYIEPY----
090 YP_002308474.1   -------------APRSSTISVKPERLQALTDLVSKALEAKYIEPY----
091 YP_007761644.1   --------------------------------------------------
092 NP_040550.1      -------------PVWIDQWPLPEGKLVALTQLVEKELQLGHIEPS----
093 NP_758808.1      TISRWESITLNMMNE--KVWPSNEAKAKYMENLLGEMEKKTWIQWRT--T
094 YP_009345075.1   VITRWSTINSNLLSQ--TDFSDNATKVQFVENLLGEDEKRLFQQWRS--N
095 NP_068729.1      --------------------------------------------------
```

```
096 NP_047255.1         -----------ATPISIRQYPMPHEAYQGIKPHIRRMLDQGILKPC----
097 YP_006273075.1      AFSRWESVTKNHVAT--QGFTDTRDKIKYMENLLGEIEKLIWIQWRM--T
098 YP_009508888.1      -------------PRPQKQYPINPKAKESIQIVINDLLKQGVLIQQ----
099 NP_395469.1         --------------------------------------------------
100 YP_009508546.1      -------------PRPQKQYPINPKAKPSIQIVIDDLLKQGVLTPQ----
101 NP_056762.1         PPQFYYDQLVTWEGIVKNEWEARKKDGMDMWSWMDGRITGLVLYLVQ--D
102 YP_009508556.1      -------------PRPQKQYPINPKAKPSIQIVIDDLLKQGVLIQQ----
103 NP_569153.1         VFTSWEAITKAYVAQ--QGITDPRDKAEFIENMLGPTEKIIWTQWRM--G
104 NP_043924.1         NLQGMTASFTADHPKMIKQYPVPDASHASIKETVEALLEQGVLRKCN---
105 YP_009243641.1      -------------PIWIEQWPLTREKLQAAEELVEEQLQKGHIETS----
106 YP_004300274.1      --------------------------------------------------
107 NP_056848.1         --------------------------------------------------
108 YP_006495799.1      AINRWESITINVVSK--LAFDNMQDKVDYVENLLGEREKEVWTTWRM--K
109 YP_009508409.1      --------------------------------------------------
110 NP_687035.1         -------------GPKLKQWPLSREKIEALTEICKQMEEEGKLSRIG--G
111 NP_045937.2         VPPITIKIKDNASLPSIRQYPLPKDKTEGLRPLISSLENQGILIKC----
112 NP_612577.1         --------------------------------------------------
113 NP_619548.1         --------------------------------------------------
114 NP_056880.1         -------------PVWLNQWPLKQEKLQALQQLVTEQLQLGHLEES----
115 YP_009121747.1      VLARWESITINRLND--TIWANNKAKLMFVENLLGENEKKMWQQWRT--A
116 YP_009113237.2      VFTRWESITKNYVSL--QMFSSGQEKMDFIENLLGEKEKLVWISWRM--A
117 YP_009508408.1      TFMRWESITKNLVSQ--HTFTSGRDKAEFIENLLGEYEKLAWIQWRT--S
118 NP_955579.1         -----------ATPISIRQYPMPHEAYQGIKPHIRRMLDQGILKPC----
119 YP_009229919.1      VVSRWESVTLNVIDDPMKSWSSNRQKISFIENLLGEDEKKIWQQWKA--A
120 YP_004442833.1      IFERWESTTLNYMAS--QNIIGAEEKISRIENLLGETEKKIFIGWRT--Q
121 NP_862833.2         -------------PVWVDQWPLTQEKLSAAQQLVQEQLRLGHIEPS----
122 NP_040563.1         -------------GPKVPQWPLTKEKYQALKEIVKDLLAEGKISEAA--W
123 YP_443922.1         PPVRVQLTHQGQLPVYRSQYRLKYEQIEGIRPTVEGLLGADCIYAT----
124 NP_057933.2         -----------STPVSIKQYPMSQEARLGIKPHIQRLLDQGILVPC----
125 YP_009116631.1      VINRWESITVNLINK--MMFDSLQDKADYVENLLGEREKETWMTWRM--Q
126 YP_009513249.1      -------------PTPQKQYHINPKAKPDIQIVINDLLKQGVLIQK----
127 YP_223871.1         -----------ALPVRVRQYPITLEAKRSLRETIHKFRAAGILRPV----
128 NP_127504.1         WQTENALAQNRMLTTIDRTLKETVTKVDRVTDQSSKNQGLIKVLEQQ--L
129 NP_044929.1         -------------PRPQKQYHINPRAKADIQIVIDDLLRQGVLRQQ----
130 NP_057860.1         -------------PPEISQFPLNPERLQALQHLVRKALEAGHIEPY----
131 NP_040973.1         -------------GPQIKQWPLTNEKIEALTEIVERLEREGKVKRAD--P
132 NP_777317.1         VFDRWESTTLLYLSE--RNFDTAEDKIRVIENLLGESEKKMFIAWRM--N
133 NP_054716.1         -------------PKPQKQYRINPKAKADIQIVIDDLLKQGVLKQQ----
134 YP_001856242.1      -------------GPKVKQWPLTEEKIKALVEICTEMEKEGKISKIG--P
135 NP_954565.2         -------------PVWVDQWPLTNDKLAAAQQLVQEQLEAGHITES----
136 NP_569150.1         VFERWESITLLHMTE--KIFDNADDKMRYMENLLGEDEKKHFIEWRM--K
137 NP_659397.1         --------------------------------------------------
138 YP_009513211.1      -----------ASPVAVRQYPMSKEAREGIRPHIQRFLDLGILVPC----
139 YP_233107.1         VFERWESSVLVHLAD--KNFDTPEDKVIYIENLLGESEKKAFMTWRM--K
140 YP_009109692.1      -----------AVPVSIKQYPMSLEAHMGIRQHIIKFLELGVLRPC----
141 NP_057849.4         -------------GPKVKQWPLTEEKIKALVEICTEMEKEGKISKIG--P
142 NP_049560.1         --------------------------------------------------
143 YP_009506267.1      VFSRWESITKNLVSQ--QSFITGKEKAEFIENLLGETEKLTWIQWRT--A
144 NP_039820.1         VFDRWASITRNYISS--FNFNDGGDKIAWMEDLLGETERKIFVSWRM--R
145 NP_056790.1         -----------ASPVAVRQYPMSKEAREGIRPHIQKFLDLGVLVPC----
146 YP_002321513.1      ILDQWESSTKNLLNG--QQFDSNRSKVEYVENLLGSTAKKTFLQWKV--A
147 NP_955591.1         -----------STPVSIKQYPMSQEARLGIKPHIQRLLDQGILVPC----
148 YP_001956722.2      -------------PRPQKQYPINPKAKASIQTVINDLLKQGVLIQQ----
149 NP_056902.1         -------------GPKIPQWPLTKEKLEGAKEIVQRLLSEGKISEAS--D
150 NP_056886.1         -------------PVWIDQWPLPEGKLVALTQLVEKELQLGHIEPS----
151 NP_041261.1         --------------------------------------------------
```

```
001 YP_004442836.1      YTTEYETMKAQALGDQGTQNIINQMRLIFFLENPQVGTTTSQDAAYKTLK
002 NP_056728.1         --------------------------------------------------
003 YP_009508537.1      ESTMNTPVYPVPKP-NGRWRMVLDYRAVNKVTPLIAVQNQHSYGILGSLF
004 YP_004732983.2      YDTEYRQLVEISGE---PRNVTSTIKRVLGINDPYTGTTHIQNQAYADLE
005 YP_009508407.1      --------------------------------------------------
006 NP_040939.1         HWTCNTPIFCIKKK-SGKWRMLIDFRELNKQTEDLTEAQLGLP-HPGGLQ
007 YP_003864102.1      NSPWNTPIFVIKKK-SGKWRLLQDLRAVNATMILMGALQPGLP-SPVAIP
008 YP_009002585.1      FPTEYQEMVNAADGREGTQNILSQMRRVFTLEDPTTGSTAVQDEAYRDLE
009 YP_003284237.1      FPTEFELMKTQALGANGTQNSISQIRRIFYLEDPKTGSTTSQDAAYKAIK
010 YP_009268869.1      HWTCNTPIFCIKKK-SGKWRMLIDFRELNKQTEDLAEAQLGLP-HPGGLQ
011 YP_009506251.1      YEAEYQQLLTIAEGRQGTQNILSQIKRVFSLEDPASGSTRIQDSAYRDLE
012 NP_955611.1         LSCWNTPVFVIRKA-SGSYRLLHDLRAVNAKLVPFGAVQQGAP-VLSALP
013 YP_008567619.1      YQAEYQQMMNSADGRAGTENILSQVRRIFALEDPSTGSTKIQDEAYRDLE
014 NP_149413.1         FPNAFDTLRGQALGTNGTANVFAQIKRVLLGEVPEQGTTGVQDAAYKKIK
```

```
015 YP_004442824.1    YTNEYETMKAQALGDNGSQNILNQLRLIFFLENPQVGTTDSQDAAYKTLK
016 YP_001931967.1    --------------------------------------------------
017 YP_002455786.1    SGPGNNPVFPVKKP-NGKWRFIHDLRATNAITTTLASPSPGPP-DLTSLP
018 YP_009109694.1    RSPWNTPLLPVKKPGTQDYRPVQDLREINKRTVDIHPTVPNPYNLLSTLK
019 NP_789739.1       ENPYNTPVFAIKKKDSTKWRKLVDFRELNKRTQDFWEVQLGIP-HPAGLK
020 NP_955619.1       TGPGNNPVFPVKKA-NGTWRFIHDLRATNSLTIDLSSSSPGPP-DLSSLP
021 YP_224289.1       YANEYEEMKGQALGNNGTQNLLSQIRRIFYLEDPKTGTTVSQDAAYKAIK
022 YP_009182100.1    SHSKAQTHPVHIFDDEPIQLDVSKSGRPKARMIRASTRKEVVVELHDINS
023 NP_705927.1       ENPYNTPVFAIKKKDSTKWRKLVDFRELNKRTQDFWEVQLGIP-HPAGLK
024 YP_009508577.1    TSVMNTPIYPVPKP-DGKWRMVLDYRAVNKTVPLIGAQNQHSLGILTNLV
025 YP_004442830.1    YTTEFELLKAQAIGVNGTQNVISQIRRILLGDSPKQGTTLTQDAAYKRIK
026 NP_758887.1       ENPYNTPVFCIKKKDGTSWRKLVDFRQLNKVTQDFFEVQLGIP-HPGGLK
027 YP_009508406.1    --------------------------------------------------
028 YP_610965.1       FEQEYESLIAVADD---SQAIISQVRRIITLEDPSQGSTEEQDRAYMDLG
029 YP_001931961.1    --------------------------------------------------
030 YP_241114.1       TSAWNSPIFVIKKK-SGKWRLLQDLRKVNETMMHMGALQPGLP-TPSAIP
031 YP_006732334.1    --------------------------------------------------
032 YP_009041481.1    FEDDYRKLVSVAGD---IRNVTSAIRVIFLTEDPAQGSTMEQDRAYADLE
033 YP_001036293.1    FNAEFDSLIAAADGREGTQNIISQIRTIFSLEDPFRGSTKIQEDAYRDLE
034 YP_605811.1       FTAEYEEMKNNALGSNGTQNILNQIRMIFFLENPQVGTTNTQDAAYKTLK
035 NP_955577.1       QSPWNTPLLPVKKPGTEDYRPVQDLREVNKRVEDIHPTVPNPYNLLSTLP
036 YP_003208050.1    YTKEYEELVAIADD---PQDITSQVRMVITLEDPYRGSTEEQSRAYLDLE
037 NP_789740.1       ENPYNTPVFAIKKKDSTKWRKLVDFRELNKRTQDFWEVQLGIP-HPAGLK
038 YP_004442839.1    FTTEYEEMRANALGSNGTQNILNQIRMIFFLENPQVGTTNSQDAAYKTLK
039 YP_009507791.1    NNPWNTPIFCIKKK-SGKWRMLIDFRELNKLTLKGAEVQLGLP-HPAGLS
040 YP_004581513.1    --------------------------------------------------
041 NP_043933.1       --------------------------------------------------
042 YP_009507248.1    SSPIWVVPKKIDASGKQKWRLVVDFRKLNEKTIDDKYPIPNISDVLDKLG
043 YP_002117531.1    YEQEYEALKTQAVGENGTQNVLSHIRKIFFLEDPKQGTTSSQDAAYRSLK
044 YP_004347415.1    --------------------------------------------------
045 NP_056891.1       SSPWNTPIFVIKKK-SGKWRLLQDLRAVNATMVLMGALQPGLP-SPVAIP
046 YP_009513242.1    NSTMNTPVYPVPKP-DGKWRMVLDYREVNKTIPLIAAQNQHSAGILSSIY
047 YP_009352866.1    FEKDYKRLVENAGD---IRNVTSAIRVIFLTEDPAQGSTVEQDRAYADLE
048 YP_009165750.1    --------------------------------------------------
049 NP_040840.1       HWTCNTPIFCIKKK-SGKWRMLIDFRELNKQTEDLAEAQLGLP-HPGGLQ
050 YP_004222728.1    LSCWNTPVFVIRKA-SGSYRLLHDLRAVNAKLVPFGAVQQGAP-VLSALP
051 YP_009506264.1    YEAEYTAMKGQALGNNGTQNILSQIRRIFYLEDPKSGTTLTQDAAYKAIK
052 YP_567050.1       SGPGNNPVFPVKKP-NGKWRFIHDLRATNAITTTLASPSPGPP-DLTSLS
053 YP_009508571.1    NSTMNTPVYPIPKP-DGRWGMVLDYREVNKTIPLIAAQNQHSAGILATIV
054 NP_056803.1       NSTMNTPVYPVPKP-DGRWRMVLDYREVNKTIPLTAAQNQHSAGILATIV
055 NP_955564.1       NSPWNTPVFVIKKK-SGKWRLLQDLRAVNATMHDMGALQPGLP-SPVAVP
056 YP_009508411.1    YPAEYDTLLASADGREGTQNILSQIRTIFTLEDPFRGSTKVQEDAYRDLE
057 YP_009140788.1    YVAEYESLIAIGDGRDGTQNILSQMRRVFSLEDPAQGSTVIQDEAYRELE
058 YP_009508410.1    --------------------------------------------------
059 NP_040333.1       QSPWNTPLLPVKKPGTNDYRPVQDLREVNKRVEDIHPTVPNPYNLLSGLP
060 YP_008992013.1    QNLQSRIAQLISELRTMLAQRYHGPEFDQKEREIRHLKAELAQIESDKQK
061 NP_041734.1       YEAEYQQLLTQADGRQGTQNILSQIKRIFSLEDPASGSTRIQDAAYRDLE
062 NP_056895.1       DGPGNNPVFPVRKP-NGTWRFVHDLRATNALTKPIPALSPGPP-DLTAIP
063 YP_009109689.1    RSPWNTPLLPVKKPGTQDYRPVQDLREINKRTVDIHPTVPNPYNLLSTLK
064 YP_002519387.1    --------------------------------------------------
065 YP_595725.1       FPTAFEAMKTQATGTNGTQNVFSQMKRILLGEVPEQGTTNTQDAAYKRIK
066 YP_009508551.1    NSTMNTPVYPVPKP-DGRWRMVLDYREVNKTIPLTAAQNQHSAGILATIV
067 YP_001497148.1    RSPWNTPLLPVKKPGTRDYRPVQDLREVNKRTMDIHPTVPNPYNLLSTLS
068 NP_041186.1       -------------------------------MMHMGALQPGLP-TPSAIP
069 NP_663784.1       TNPYNTPTFAIKKKDKNKWRMLIDFRELNKVTQDFTEIQLGIP-HPAGLA
070 YP_009508443.1    FPEEYEKLLASADGREGTQNIISQLRTIFTLEDPFRGSTKVQEEAYRDLE
071 YP_003987465.1    YEEEFVSLQTFVLGNNGTQNLLAQIRKVFFLEDPKGGTTVSQDAAYKAIK
072 NP_861410.1       --------------------------------------------------
073 YP_009345071.1    YETEYQALIGTMEGRSGTQNMLSQIRRIFTLEDPYQGSTMSQDQAYKDLE
074 YP_009508582.1    TSVMNTPVYPVPKP-DGKWRMVLDYRAVNKTIPLIGAQNQHSLGILTNLV
075 NP_569141.1       --------------------------------------------------
076 YP_002916057.1    --------------------------------------------------
077 NP_777384.2       DGPGNNPVFPVRKP-NGTWRFVHDLRATNALTKPIPALSPGPP-DLTAIP
078 YP_233110.1       FTAEYEEMKNNALGSNGTQNILNQIRMIFFLENPQVGTTNTQDAAYKTLK
079 NP_542258.1       QGPGNNPIFPVKKP-NGKWRFIHDLRATNSLTRDLASPSPGPP-DLTSLP
080 YP_009130664.1    YPEAFQQMLESADGRQGTQNILSQIYRVFTLEDPYQGSTTMQEEALRDLE
081 YP_009508566.1    NSIMNTPIYPVPKT-EGKWRMVLDYRAVNKTIPLIAAQNQHSAGILTNLV
082 YP_006607892.1    --------------------------------------------------
083 YP_009506270.1    YTVEYETMKAQALGDQGTQNIINQIRLIFFLENPQVGTTTSQDAAYKTLK
084 NP_042513.1       --------------------------------------------------
085 YP_006907834.1    --------------------------------------------------
086 YP_004442827.1    FPTDFEALKNQAIGTNGTQNVISQVRRILLGDSPKQGTTATQDAAYKKIK
087 YP_009408594.1    YPDQYQRLIAIGDD---TQNILSQIRRIITLEDPATGSTAERDQAYLDLE
```

| | | |
|---|---|---|
| 088 | YP_009508561.1 | TSPMNTPVYPVPKP-DGKWRMVLDYRAVNKTIPLIAAQNQHSLGILTNLI |
| 089 | NP_056907.1 | SGPGNNPVFPVKKP-NGKWRFIHDLRATNAITTTLASPSPGPP-DLTSLS |
| 090 | YP_002308474.1 | QGPGNNPIFPVKKP-NGKWRFIHDLRATNCLTKTLTSPSPGPP-DLTSLP |
| 091 | YP_007761644.1 | -------------------------------------------------- |
| 092 | NP_040550.1 | LSCWNTPVFVIRKA-SGSYRLLHDLRAVNAKLVPFGAVQQGAP-VLSALP |
| 093 | NP_758808.1 | YVSEYDALVQQSDE---TQNLLSQVRRIFLLQDPYQGSTAEQDQAYNDLE |
| 094 | YP_009345075.1 | FPEEYDTLVEHADD---NINMFSQIRRMFTLEDPYSGSTMAQERAYYDLE |
| 095 | NP_068729.1 | -------------------------------------------------- |
| 096 | NP_047255.1 | QSPWNTPLLPVKKPGTEDYRPVQDLREVNKRVEDIHPTVPNPYNLLSTLP |
| 097 | YP_006273075.1 | YATEYEALVAIGEGRDGTQNILSQMRRVFSLEDPAQGSTVIQDEAYRELE |
| 098 | YP_009508888.1 | NSIMNTPVYPVPKP-DGRWRMVLDYREVNKTIPLIAAQNQHSAGILASIY |
| 099 | NP_395469.1 | -------------------------------------------------- |
| 100 | YP_009508546.1 | NSTMNTPVYPVPKP-DGRWRMVLDYREVNKTIPLTAAQNQHSAGILATIV |
| 101 | NP_056762.1 | WISKNQAAYNDIKSRGDRPENFVKMVKDRFLIEDPTDERRTALQRLAQRE |
| 102 | YP_009508556.1 | NSTMNTPVYPVPKP-DGKWRMVLDYREVNKTIPLIAAQNQHSARILSSFF |
| 103 | NP_569153.1 | YADEYENLVTTADGREGTQNILSQMRRVFSLEDPTTGSTAVQDEAYRDLE |
| 104 | NP_043924.1 | -STVNSAIWPVGKP-DGSWRLTIDYRPLNSAVSCPYPTVASTPELFAKLE |
| 105 | YP_009243641.1 | TSPWNSPIFVIRKK-SGKWRLLTDLRAVNASMFPMGALQPGLP-APVMLP |
| 106 | YP_004300274.1 | -------------------------------------------------- |
| 107 | NP_056848.1 | -------------------------------------------------- |
| 108 | YP_006495799.1 | WEDEYRRMVAISDD---TRNLTAAIKRVFGVHDPFTGSTFLQNQAYADLE |
| 109 | YP_009508409.1 | -------------------------------------------------- |
| 110 | NP_687035.1 | ENPYNTPVFAIKKKDKTQWRMLVDFRELNKATQDFFEVQLGIP-HPAGLQ |
| 111 | NP_045937.2 | HSPCNTPIFPIKKAGRDEYRMIHDLRAINNIVAPLTAVVASPTTVLSNLA |
| 112 | NP_612577.1 | -------------------------------------------------- |
| 113 | NP_619548.1 | -------------------------------------------------- |
| 114 | NP_056880.1 | NSPWNTPVFVIKKK-SGKWRLLQDLRAVNATMHDMGALQPGLP-SPVAVP |
| 115 | YP_009121747.1 | FPEAYEALETMADD---PQNITSQIRQITIMEDPYRGSTEEQDRAYRSLE |
| 116 | YP_009113237.2 | YQDEYNQLIAKSDGYEGVQNILSQIRRVFTLQDPATGSTKAQDQAYRDIE |
| 117 | YP_009508408.1 | YPDEYEKILESSDGREGTQNILSQIRTVFTLEDPFRGSTKVQEDAYRDLE |
| 118 | NP_955579.1 | QSPWNTPLLPVKKPGTEDYRPVQDLREVNKRVEDIHPTVPNPYNLLSTLP |
| 119 | YP_009229919.1 | YPEDYNSLEAIAEE---TQNVLSQIRKVFLLEVPYQGDLAEQNQAYIDLE |
| 120 | YP_004442833.1 | YTTEFEILKTQAIGTNGTQNVISQIRRILLGDSPKQGTTLTQDAAYKRIK |
| 121 | NP_862833.2 | TSAWNSPIFVIKKK-SGKWRLLQDLRKVNETMMHMGALQPGLP-TPSAIP |
| 122 | NP_040563.1 | DNPYNTPVFVIKKKGTGRWRMLMDFRELNKITVKGQEFSTGLP-YPPGIK |
| 123 | YP_443922.1 | SSPWNTPILPVPKADGKTYRMVQDFRAVNDVTTGVDLPVPDPHVSLSNLS |
| 124 | NP_057933.2 | QSPWNTPLLPVKKPGTNDYRPVQDLREVNKRVEDIHPTVPNPYNLLSGLP |
| 125 | YP_009116631.1 | YEEEYKQLLTMSGD---VRNITAAVKRVFGVHDPHTGSVHIQNQAYAELE |
| 126 | YP_009513249.1 | ESTMNTPVYPVPKP-NGRWRMVLDYRAVNKVTPLIAVQNQHSYGILGSLF |
| 127 | YP_223871.1 | HSPWNTPLLPVRKSGTSEYRMVQDLREVNKRVETIHPTVPNPYTLLSLLP |
| 128 | NP_127504.1 | QDLNKRICPPGTSLFHFFDQ--QKSEMASLKEQIRLLKEQPQKNETDT-- |
| 129 | NP_044929.1 | NSEMNTPVYPVPKA-DGRWRMVLDYREVNKVTPLVATQNCHSASILNTLY |
| 130 | NP_057860.1 | TGPGNNPVFPVKKA-NGTWRFIHDLRATNSLTIDLSSSSPGPP-DLSSLP |
| 131 | NP_040973.1 | NNPWNTPVFAIKKK-SGKWRMLIDFRELNKLTEKGAEVQLGLP-HPAGLQ |
| 132 | NP_777317.1 | FTEAYEELKAQAMGYNGSLNILNQIRLIFFLESPQAGTTDSQDAAFKTLK |
| 133 | NP_054716.1 | TSPMNTPVYPVPKP-DGRWRMVLDYRAVNKVTPAIATQNCHSASLLNTLY |
| 134 | YP_001856242.1 | ENPYNTPVFAIKKKDSTKWRKLVDFRELNKRTQDFWEVQLGIP-HPAGLK |
| 135 | NP_954565.2 | SSPWNTPIFVIKKK-SGKWRLLQDLRAVNATMVLMGALQPGLP-SPVAIP |
| 136 | NP_569150.1 | YTAEYETMKAQALGDQGTQNIINQIRLIFFLENPQVGTTTSQDAAYKTLK |
| 137 | NP_659397.1 | -------------------------------------------------- |
| 138 | YP_009513211.1 | QSPWNTPLLPVKKPGTNDYRPVQDLREVNKRVQDIHPTVPNPYNLLSSLP |
| 139 | YP_233107.1 | YLPEFEALKAAALGNNGTQNILNQIRMIFFLENPKVGTTDEQDAAYKTIK |
| 140 | YP_009109692.1 | RSPWNTPLLPVKKPGTQDYRPVQDLREINKRTVDIHPTVPNPYNLLSTLK |
| 141 | NP_057849.4 | ENPYNTPVFAIKKKDSTKWRKLVDFRELNKRTQDFWEVQLGIP-HPAGLK |
| 142 | NP_049560.1 | -------------------------------------------------- |
| 143 | YP_009506267.1 | FPTEYETLLASADGREGTQNILSHIRTIFTLEDPYRGSTKVQEQAYLDLE |
| 144 | NP_039820.1 | FKDEYQNLAKIANQDGGTQAILSQIRRIFLGEDPVLGQNTVQNIAFRKLK |
| 145 | NP_056790.1 | RSPWNTPLLPVKKPGTNDYRPVQDLREINKRVQDIHPTVPNPYNLLSSLP |
| 146 | YP_002321513.1 | YPTDYQQLITVADD---PQNITSMIRRIFTMEDPYTGSTLLQAVAYKDVE |
| 147 | NP_955591.1 | QSPWNTPLLPVKKPGTNDYRPVQDLREVNKRVEDIHPTVPNPYNLLSGLP |
| 148 | YP_001956722.2 | NSIMNTPVYPVPKP-DGKWRMVLDYREVNKTIPLIAAQNQHSAGILSSIF |
| 149 | NP_056902.1 | NNPYNSPIFVIKKR-SGKWRLLQDLRELNKTVQVGTEISRGLP-HPGGLI |
| 150 | NP_056886.1 | LSCWNTPVFVIRKA-SGSYRLLHDLRAVNAKLVPFGAVQQGAP-VLSALP |
| 151 | NP_041261.1 | -------------------------------MVPMGALQPGLP-SPVAIP |

| | | |
|---|---|---|
| 001 | YP_004442836.1 | SLVCTEMTDTAIYRYMNDYFHLSAKTGRAWATEE---LSKEFFTKLPRGL |
| 002 | NP_056728.1 | -------------------------------------------------- |
| 003 | YP_009508537.1 | K-GRYKTTIDLSNGFWAHPIVPEDYWITAFTWQGK-------QYCWTVLP |
| 004 | YP_004732983.2 | RLQCKNLE--SVMPFLNSYFQLAAKSGKMWSSPE---LSEKLFRKLPPEI |
| 005 | YP_009508407.1 | -------------------------------------------------- |
| 006 | NP_040939.1 | KKK-HVTILDIGDAYFTIPLYEPYREYTCFTLLSPNNLGPCKRYYWKVLP |

```
007 YP_003864102.1 QNY-LKIIIIDLKDCFFTIPLHPNDQKRFAFSLPSTNFKEPMKRYQWKVLP
008 YP_009002585.1 RLSCTNIK--DIVSFLNDYARLAAKSGRMFISPE---LSEKLWLKMPYDL
009 YP_003284237.1 SLVCTDMSGIAIKRYMNSYMDLAATSGRMWVSAE---LSDEFFTKLPNGL
010 YP_009268869.1 KKK-HVTILDIGDAYFTIPLYEPYRPYTCFTMLSPNNLGPCTRYYWKVLP
011 YP_009506251.1 RLTCHNIK--DIVQFLNDYGRLAAKSGRLFIGTE---LSEKLWMKMPPEL
012 NP_955611.1 RGW-PLMVLDLKDCFFSIPLAEQDREAFAFTLPSVNNQAPARRFQWKVLP
013 YP_008567619.1 QLTCSNIK--DLVPFLNDYLRLAAKTGRMFLGTE---LSEKLWMKLPGNL
014 NP_149413.1 SLVCTEMTYPALMRYMVGYRNLAARSGRAWANNE---LTNEFFTKLPGKL
015 YP_004442824.1 SLVCTEMTDTAIYRYMNDYFHLAAKSGRAWANEE---LSQEFFTKLPRGL
016 YP_001931967.1 --------------------------------------------------
017 YP_002455786.1 QALPHLQTIDLTDAFFQIPLPKRFQPYFAFTIPQPLNHGPGSRYAWTVLP
018 YP_009109694.1 PDYSWYTVLDLKDAFFCLPLAPQSQELFAFEWKDPE-RGISGQLTWTRLP
019 NP_789739.1 KKK-SVTVLDVGDAYFSVPLDEDFRKYTAFTIPSINNETPGIRYQYNVLP
020 NP_955619.1 TTLAHLQTIDLKDAFFQIPLPKQFQPYFAFTVPQQCNYGPGTRYAWRVLP
021 YP_224289.1 SLVCHEMTGTAIKRYMADYWTLAAKTGRIWQGPE---LSDEFFTKLPSGL
022 YP_009182100.1 IREISDEKLDEIYATTSDPQYFNYTVLPEMQRRINQGISVRAISQSVNRP
023 NP_705927.1 KKK-SVTVLDVGDAYFSVPLDEDFRKYTAFTIPSINNETPGIRYQYNVLP
024 YP_009508577.1 R-QKYKSTIDLSNGFWAHPITKDSQWITAFTWEGK-------QHVWTRLP
025 YP_004442830.1 SLVCTEISYPAIMRYLVGYFHLSARTGRMWSSKE---LSDEFFTKLPDEI
026 NP_758887.1 QCE-QITVLDIGDAYFSCPLDEDFRKYTAFTIPSVNNQGPGIRYQYNVLP
027 YP_009508406.1 --------------------------------------------------
028 YP_610965.1 RLSCTNMK--DLLDYMNDYKTLAAKSGRMYVNHE---LSDKFFDKMPQII
029 YP_001931961.1 --------------------------------------------------
030 YP_241114.1 DKS-YIIVIDLKDCFYTIPLAPQDCKRFAFSVPSVNFKEPMKRYQWKVLP
031 YP_006732334.1 --------------------------------------------------
032 YP_009041481.1 RLPTPEMK--NIFQFLNQYKKLAAASGRMWITPE---LSEKLFRKLPPVI
033 YP_001036293.1 RISCKDVQ--DIIPFMNEYMRIAAKTGRLFISSE---LSEKFWMKLPGDL
034 YP_605811.1 QLVCTEMSGPAIYRYLNDYFHLAAKSGRAWASDE---LSKEFFTKLPKGL
035 NP_955577.1 PSHPWYTVLDLKDAFFCLRLHSESQLLFAFEWRDPE-IGLSGQLTWTRLP
036 YP_003208050.1 RLVCDDIK--DILPYLNDFKVLAAKSGRMFTSPE---LSEKLFRKLPPLI
037 NP_789740.1 KKK-SVTVLDVGDAYFSVPLDEDFRKYTAFTIPSINNETPGIRYQYNVLP
038 YP_004442839.1 QLVCTEMSGPAIYRYLNDYFHLSAKSGRAWASEE---LSKEFFTKLPKGL
039 YP_009507791.1 MRK-QVTVLDIGDAYFTIPLDPDYQPYTAFTLPNKNNQGPGRRYVWCSLP
040 YP_004581513.1 --------------------------------------------------
041 NP_043933.1 --------------------------------------------------
042 YP_009507248.1 KCQ-YFTTLDLASGFYQVEMDPQDISKTAFNVEHG-------HFEFLRMP
043 YP_002117531.1 SLTCMEMTGTAIRRYMASYFDLAAKSGRMWSSEE---LSKEFFQKLPGRL
044 YP_004347415.1 --------------------------------------------------
045 NP_056891.1 QGY-LKIIIDLKDCFFSIPLHPSDQKRFAFSLPSTNFKEPMQRFQWKVLP
046 YP_009513242.1 R-GKYKTTLDLTNGFWAHPITPESYWLTAFTWQGK-------QYCWTRLP
047 YP_009352866.1 RLPTPEMK--NIFQFLNQYKKLAAASGKMWITSE---LSEKLFRKLPPII
048 YP_009165750.1 --------------------------------------------------
049 NP_040840.1 RKK-HVTILDIGDAYFTIPLYEPYRQYTCFTMLSPNNLGPCVRYYWKVLP
050 YP_004222728.1 RGW-PLMVLDLKDCFFSIPLAEQDREAFAFTLPSVNNQAPARRFQWKVLP
051 YP_009506264.1 SLVCNDENGEAVKRYMLSYFDLAARSGRMWTSQE---LSEEFFTKLPDGL
052 YP_567050.1 TALPYLQTIDLTDAFFQIPLPKQFQPYFAFTIPQPCNYGPGARYAWTVLP
053 YP_009508571.1 R-KKYKTTLVLANGFWAHPITPESYWLTAFIWQGK-------QYCWTRLP
054 NP_056803.1 R-QKYKTTLDLANGFWAHPITPDSYWLTAFTWQGK-------QYCWTRLP
055 NP_955564.1 KGW-EIIIIDLQDCFFNIKLHPEDCKRFAFSVPSPNFKRPYQRFQWKVLP
056 YP_009508411.1 RISCSDMK--NIIAFMNEYMRLASKSGRLFVSPE---LSEKFWLKLPGDL
057 YP_009140788.1 RLSCTDVK--YIIPFLNEYLRLAAKSGRIFLGGE---LSEKIWMKMPGDL
058 YP_009508410.1 --------------------------------------------------
059 NP_040333.1 PSHQWYTVLDLKDAFFCLRLHPTSQSLFAFEWRDPE-MGISGQLTWTRLP
060 YP_008992013.1 PSLFTSSPPIPSIGPTYHPFASMLSLIRQYEPSKLFGMTHTLFRDNPLPP
061 NP_041734.1 RLTCHNIK--DIVQFLNDYGRLAAKSGRLFLGTE---LSEKLWMKMPPEL
062 NP_056895.1 THLPHIICLDLKDAFFQIPVEDRFRSYFAFTLPTPGGLQPHRRFAWRVLP
063 YP_009109689.1 PDYSWYTVLDLKDAFFCLPLAPQSQELFAFEWKDPE-RGISGQLTWTRLP
064 YP_002519387.1 --------------------------------------------------
065 YP_595725.1 SLVCQEMTYPAIMRYLVGYRNLAARSGRAWVNNE---LTDEFFTKLPGKL
066 YP_009508551.1 R-QKYKTTLDLANGFWAHPITPESYWLTAFTWQGK-------QYCWTRLP
067 YP_001497148.1 PDRTWYTVLDLKDAFFCLPLAPQSQELFAFEWRDPE-RGISGQLTWTRLP
068 NP_041186.1 DKS-YIIVIDLKDCFYTIPLAPQDCKRFAFSLPSVNFKEPMQRYQWRVLP
069 NP_663784.1 KKK-RISILDVGDAYFSIPLHEDFRQYTAFTLPAVNNMEPGKRYIYKVLP
070 YP_009508443.1 RISCRDVK--DLIPFMNEYMRIASKTGRLFISSE---LSEKFWFKLPGDL
071 YP_003987465.1 SLTCQEMSGQAIRKYMSSYFDLAARSGRMWVNEE---LSDDFFTKLPMGL
072 NP_861410.1 --------------------------------------------------
073 YP_009345071.1 RLTCTDVK--HILEYLRQFMYLAAKTGRLFTSSE---LSSKLFVKMPGDL
074 YP_009508582.1 R-QKYKSTIDLSNGFWAHPITKDSQWITAFTWEGK-------QHVWTRLP
075 NP_569141.1 --------------------------------------------------
076 YP_002916057.1 --------------------------------------------------
077 NP_777384.2 THLPHIICLDLKDAFFQIPVEDRFRSYFAFTLPTPGGLQPHRRFAWRVLP
078 YP_233110.1 QLVCTEMSGPAIYRYLNDYFHLAAKSGRAWASDE---LSKEFFTKLPRGL
079 NP_542258.1 QDLPHLRTIDLTDAFFQIPLPAVFQPYFAFTLPQPNNHGPGTRYSWRVLP
```

| No. | ID | Sequence |
|---|---|---|
| 080 | YP_009130664.1 | KLSCHNIK--DIVAYMNEYMRLACKSRRLFINAD---LSEKFWFKMPGEL |
| 081 | YP_009508566.1 | R-QKYKSTIDLSNGFWAHPIDQDSQWITAFTWEGK-------QYVWTRLP |
| 082 | YP_006607892.1 | -------------------------------------------------- |
| 083 | YP_009506270.1 | SLVCTEMTDTAIYRYMNDYFHLSAKTGRAWASEE---LSKEFFTKLPRGL |
| 084 | NP_042513.1 | -------------------------------------------------- |
| 085 | YP_006907834.1 | -------------------------------------------------- |
| 086 | YP_004442827.1 | SLVCTELSYPAIMRYMVGYFHLASKSGRMWTSNE---LSDEFFTKLPDEI |
| 087 | YP_009408594.1 | RLSCEHMK--DLIPYMNRYKVLSAKTGRLFINSE---LSDKFFRKMPTLI |
| 088 | YP_009508561.1 | R-HKYKSTIDLSNGFWAHPITEDSQWITAFTWEGK-------QHVWTRLP |
| 089 | NP_056907.1 | TALPYLQTIDLTDAFFQIPLPKQFQPYFAFTIPQPCNYGPGARYAWTVLP |
| 090 | YP_002308474.1 | QGLPHLRTIDLTDAFFQIPLPVAFQPYFAFTLPQPNNHGPGARYSWKVLP |
| 091 | YP_007761644.1 | -------------------------------------------------- |
| 092 | NP_040550.1 | RGW-PLMVLDLKDCFFSIPLAEQDREAFAFTLPSVNNQAPARRFQWKVLP |
| 093 | NP_758808.1 | RISCDNIK--DLIPYLIQFRNLAAKSGRLFLGPE---LSEKLFRKMPPLI |
| 094 | YP_009345075.1 | RLTCSNIN--NVIQFMNEYKKLAAASGRAFISTE---LSEKFFRKLPPLY |
| 095 | NP_068729.1 | -------------------------------------------------- |
| 096 | NP_047255.1 | PSHPWYTVLDLKDAFFCLRLHSESQLLFAFEWRDPE-IGLSGQLTWTRLP |
| 097 | YP_006273075.1 | RMSCTDVK--YIIPFLNEYMRLAAKTGRLFLGGE---LSEKIWMKMPGDL |
| 098 | YP_009508888.1 | R-GTYKTTLDLANGFWAHPITPNSYWLTAFTWQGK-------QHCWTRLP |
| 099 | NP_395469.1 | -------------------------------------------------- |
| 100 | YP_009508546.1 | R-QKYKTTLDLANGFWAHPITPESYWLTAFTWQGK-------QYCWTRLP |
| 101 | NP_056762.1 | LEALNCEDPTKIQPFMAEYLKKASEAKKGFDVVY----VERLFDRLPEAV |
| 102 | YP_009508556.1 | S-GKYKTTLDLTNGFWAHPITPESYWLTAFTWRGK-------QYCRTKLP |
| 103 | NP_569153.1 | RLTCDSVK--HIVQYLNDFMRIAAKTGRMFIGPE---LSEKLWLKMPGDL |
| 104 | NP_043924.1 | KKYQVYSSLDISNGFWSIRLEEECQYLFAFTFDT-------QQYTWTRLP |
| 105 | YP_009243641.1 | RDW-PLIIIDLLDCFFTIPIHPQDKQRFAFSIPKINNAGPHTRYQWRVLP |
| 106 | YP_004300274.1 | -------------------------------------------------- |
| 107 | NP_056848.1 | -------------------------------------------------- |
| 108 | YP_006495799.1 | RLSCKRME--DVMPFLFTYYQLAAKSGRMWTNEE---LSDKLFRKLPEAV |
| 109 | YP_009508409.1 | -------------------------------------------------- |
| 110 | NP_687035.1 | KKK-QITVIDIGDAYYSIPLCKEFRKYTAFTIPSVNNTGPGIRYQFNCLP |
| 111 | NP_045937.2 | PSLHWFTVIDLSNAFFSVPIHKDSQYLFAFTFEG-------HQYTWTVLP |
| 112 | NP_612577.1 | -------------------------------------------------- |
| 113 | NP_619548.1 | -------------------------------------------------- |
| 114 | NP_056880.1 | KGW-EIIIIDLQDCFFNIKLHPEDCKRFAFSVPSPNFKRPYQRFQWKVLP |
| 115 | YP_009121747.1 | RITCEDTR--DLWKFLNEFRILASKSGRLFF-PA---TTDKFFTKLPMTL |
| 116 | YP_009113237.2 | KLNCHDVK--DIVSFLNDYLRLAAKTGRMYIGME---LSEKLWMKMPGEL |
| 117 | YP_009508408.1 | RISCKDMK--DIIPFMNEYMRLASKTGRLFISSE---LSEKFWMKLPGDL |
| 118 | NP_955579.1 | PSHPWYTVLDLKDAFFCLRLHSESQLLFAFEWRDPE-IGLSGQLTWTRLP |
| 119 | YP_009229919.1 | RLTCESTR--DIWRYLNDFKTLAAKSGRIYF-PE---TSEKLFRKMPPLI |
| 120 | YP_004442833.1 | SLVCDKETYPDIMRYMVGYLHLAARTGRMWISKE---LSDEFFLKLPSEI |
| 121 | NP_862833.2 | DRS-YIIVIDLKDCFYTIPLAPQDCKRFAFSLPSVNFKEPMQRYQWKVLP |
| 122 | NP_040563.1 | ECE-HLTAIDIKDAYFTIPLHEDFRPFTAFSVVPVNREGPIERFQWNVLP |
| 123 | YP_443922.1 | PKHQYFTVVDLANAFFSIPLHVDSQPLFAFTYDNQ-------QYTYSVLP |
| 124 | NP_057933.2 | PSHQWYTVLDLKDAFFCLRLHPTSQPLFAFEWRDPE-MGISGQLTWTRLP |
| 125 | YP_009116631.1 | RLYCKRTD--DVIPFLYDYYQLAAKSGRMWLGPE---LSEKLFRKLPPEI |
| 126 | YP_009513249.1 | K-GRYKTTIDLSNGFWAHPIVPEDYWITAFTWQGK-------QYCWTVLP |
| 127 | YP_223871.1 | PDRIWYSVLDLKDAFFCIPLAPESQLIFAFEWADAE-EGESGQLTWTRLP |
| 128 | NP_127504.1 | PSYQSSYQPFHSFSSPYMPSNPPNSPFTTFANTPQP--QPSLFSQYPIQP |
| 129 | NP_044929.1 | R-GPYKSTLDLANGFWAHPIKPEDYWITAFTWGGK-------TYCWTVLP |
| 130 | NP_057860.1 | TTLAHLQTIDLKDAFFQIPLPKQFQPYFAFTVPQQCNYGPGTRYAWRVLP |
| 131 | NP_040973.1 | IKK-QVTVLDIGDAYFTIPLDPDYAPYTAFTLPRKNNAGPGRRFVWCSLP |
| 132 | NP_777317.1 | SLVCTEMTDTSLYRYMNDYFHLAAKSGRAWANED---LSKEFFTKLPRGL |
| 133 | NP_054716.1 | R-GQYKTTLDLANGFWAHPIQESDQWITSFTWNGK-------SYVWTTLP |
| 134 | YP_001856242.1 | KKK-SVTVLDVGDAYFSVPLDEDFRKYTAFTIPSINNETPGIRYQYNVLP |
| 135 | NP_954565.2 | QGY-LKIIIDLKDCFFSIPLHPSDQKRFAFSLPSTNFKEPMQRFQWKVLP |
| 136 | NP_569150.1 | SLVCTEMTDTAIYRYMNDYFHLSAKTGRAWASEE---LSKEFFTKLPRGL |
| 137 | NP_659397.1 | -------------------------------------------------- |
| 138 | YP_009513211.1 | PSHTWYSVLDLKDAFFCLKLHPNSQPLFAFEWRDPE-KGNTGQLTWTRLP |
| 139 | YP_233107.1 | SLVCNEMTDKAVYRYMNDYFHLASKSGRMWANEE---LSTEFFTKLPRHL |
| 140 | YP_009109692.1 | PDYSWYTVLDLKDAFFCLPLAPQSQELFAFEWKDPE-RGISGQLTWTRLP |
| 141 | NP_057849.4 | KKK-SVTVLDVGDAYFSVPLDEDFRKYTAFTIPSINNETPGIRYQYNVLP |
| 142 | NP_049560.1 | -------------------------------------------------- |
| 143 | YP_009506267.1 | RISCRDVK--DLIPFLNEYMRLAAKTGRLFISTE---LSDKIWMKLPGDL |
| 144 | NP_039820.1 | QLVCPNY--QSIRRYLMDYLTLAAETGLMWSETEGPAISEELFTKMPAAI |
| 145 | NP_056790.1 | PSYTWYSVLDLKDAFFCLRLHPNSQPLFAFEWKDPE-KGNTGQLTWTRLP |
| 146 | YP_002321513.1 | RLQCDRIE--HIVPFMMEYFKISAETGRLYVDNE---MSDKFFRKLPPLY |
| 147 | NP_955591.1 | PSHQWYTVLDLKDAFFCLRLHPTSQPLFAFEWRDPE-MGISGQLTWTRLP |
| 148 | YP_001956722.2 | R-GKYKTTLDLSNGFWAHSITPESYWLTAFTWLGQ-------QYCWTRLP |
| 149 | NP_056902.1 | KCK-HMTVLDIGDAYFTIPLDPEFRPYTAFTIPSINHQEPDKRYVWKCLP |
| 150 | NP_056886.1 | RGW-PLMVLDLKDCFFSIPLAEQDREAFAFTLPSVNNQAPARRFQWKVLP |
| 151 | NP_041261.1 | LNY-HKIVIDLKDCFFTIPLHPEDRPYFAFSVPQINFQSPMPRYQWKVLP |

```
001 YP_004442836.1   GDEVEKAFKEKHPSNTIGVTARITFTK--RYLKELCEKVALQKSIGKMDF
002 NP_056728.1      --------------------------------------------------
003 YP_009508537.1   QGFLNSPGLFTGDVVDLLQGI--------PNVEVYVDDVYISHDSEKEHL
004 YP_004732983.2   GPTIAKEYAERYPGMLIGVNARIQFVS--EYLQDLCKQADLQRKLKNLNF
005 YP_009508407.1   --------------------------------------------------
006 NP_040939.1      QGWKLSPSVYQFTMQEILEDWIQQHPE--IQFGIYMDDIYIGSDLEIKKH
007 YP_003864102.1   QGMANSPTLCQKFVAMAIQTVRDTWKQ--IYIIHYMDDILLAGADGQQVL
008 YP_009002585.1   GNKIRTAYEAKHEGNTIGVFPRILFTY--KYLEQECKEAAFRRSLKNLSF
009 YP_003284237.1   GDRVAKAFKEKHPGNTVGVPARVTFTQ--NYLEEVCREAAYQRSLKNLDF
010 YP_009268869.1   QGWKLSPSVYQFTMQEILRDWIAKHPM--IQFGIYMDDIYIGSDLDIMKH
011 YP_009506251.1   GHRMREAFQKEYEGNEVGVFPRILFAY--RYLEQECKDAAFKRSLKSLSF
012 NP_955611.1      QGMTCSPTICQLVVGQVLEPLRLKHPS--LCMLHYMDDLLLAASSHDGLE
013 YP_008567619.1   GRRIKEAFDNKYKGNEAGVIPRILFTH--KYLEDQCKEAAFQRSLKGMTF
014 NP_149413.1      GDKVKEAFVAKYPSVERHVPAAIRFTY--DYLEEICTENNFQKQLRSLNF
015 YP_004442824.1   GDRIEKAFKEKHPSNTIGVAARITFTK--NYMKEICQEAVFQNQLKKLGF
016 YP_001931967.1   --------------------------------------------------
017 YP_002455786.1   QGFKNSPTLFEQQLASVLGPARKAFPT--SVIVQYMDDILLACPSQHELD
018 YP_009109694.1   QGFKNSPTLFDEALHRDLTDFRTQHPE--VTLLQYVDDLLLAAPTKKACT
019 NP_789739.1      QGWKGSPAIFQSSMTKILEPFRKQNPD--IVIYQYMDDLYVGSDLEIGQH
020 NP_955619.1      QGFKNSPTLFEMQLAHILQPIRQAFPQ--CTILQYMDDILLASPSHADLQ
021 YP_224289.1      GDRVAKAFKEKYPGNTVGVPARITFTQ--LYLEELCREAAYQRSLKNLDF
022 YP_009182100.1   STSGDSKEVPITSEVPSSSTLEGKGKT--KKFTPAGSSIKLPLITGVPTG
023 NP_705927.1      QGWKGSPAIFQSSMTKILEPFRKQNPD--IVIYQYMDDLYVGSDLEIGQH
024 YP_009508577.1   QGFLNSPALFTADVVDLLKNI--------PGISVYVDDIYFSTETVSEHL
025 YP_004442830.1   SDSAKEAFDKKFPGTNIHVPARIEFTY--NYLEEICTENNKQRRLGKLNF
026 NP_758887.1      QGWKGSPAIFQATADKILQPFRERHPD--VVIYQYMDDLFVGSDRVAPEH
027 YP_009508406.1   --------------------------------------------------
028 YP_610965.1      GPDMKKAFSEKYPGAQMGVLPRITFAY--KYLSDICKQAAVQKGVKDLAF
029 YP_001931961.1   --------------------------------------------------
030 YP_241114.1      QGMTNSPTLCQKLVATAIAPVRQRFPQ--IYLVHYMDDILLAHTDEHLLY
031 YP_006732334.1   --------------------------------------------------
032 YP_009041481.1   GPAIEKAYKQRYPGMDIGVPTRINFIY--QYLMEVCKQAALQRSLKDLSF
033 YP_001036293.1   GKTIKDSFDKTYVGNTVGVHPRILFTY--KYLQEECKKAALSRSLKNLQF
034 YP_605811.1      GDRVEKEFKKRYPNNTIGVAPRITFTR--NYMKEICQEAVFQSQLKKLDF
035 NP_955577.1      QGFKNSPTLFDEALHSDLADFRVRYPA--LVLLQYVDDLLLAAATRTECL
036 YP_003208050.1   GGEIEAAFKEKYPGNTVGVLPRIVFIH--HYLTDLCKKAAIQRGIKDLSF
037 NP_789740.1      QGWKGSPAIFQSSMTKILEPFRKQNPD--IVIYQYMDDLYVGSDLEIGQH
038 YP_004442839.1   GDRVEKSFKEKYPNNTIGVAARITFTR--AYMKEICQEAVFQNQLKRLDF
039 YP_009507791.1   QGWVLSPLIYQSTLDNILQPWRKKYPN--IDVYQYMDDIYIGSDFSRLEH
040 YP_004581513.1   --------------------------------------------------
041 NP_043933.1      --------------------------------------------------
042 YP_009507248.1   MGLKNSPSTFQRVMDNVLRGLQN------NICLVYIDDIIVYSTSLQEHL
043 YP_002117531.1   GERVEQAFNAKFQGNTIGVVPRITFTQ--RYLTEICEEASYQRSLKNLNF
044 YP_004347415.1   --------------------------------------------------
045 NP_056891.1      QGMANSPTLCQKYVATAIHKVRHAWKQ--MYIIHYMDDILIAGKDGQQVL
046 YP_009513242.1   QGFLNSPALFTADVVDLLKEV--------PNVQAYVDDIYMSHDDPQEHL
047 YP_009352866.1   GPAVETAFKLRYPGLDIGVPTRINFIY--HYLMEVCKQAALQRSMEDLSF
048 YP_009165750.1   --------------------------------------------------
049 NP_040840.1      QGWKLSPAVYQFTMQKILRGWIEEHPM--IQFGIYMDDIYIGSDLGLEEH
050 YP_004222728.1   QGMTCSPTICQLIVGQVLEPLRLKHPS--LCMLHYMDDLLLAASSHDRLE
051 YP_009506264.1   RDRAAEAFKKKFPGNTIGVPARITFLQ--NYLEEACKEDAYQRSIKNLNF
052 YP_567050.1      QGFKNSPTLFEQQLAAVLSPIRKTFPM--STIIQYMDDILLASPTQEELQ
053 YP_009508571.1   QGFLNSPALFTADVVDLLKEI--------SNVQAYVDDIYLSHDDPQEHL
054 NP_056803.1      QGFLNSPALFTADAVDLLKEV--------PNVQVYVDDIYLSHDNPHEHI
055 NP_955564.1      QGMKNSPTLCQKFVDKAILTVRDKYQD--SYIVHYMDDILLAHPSRSIVD
056 YP_009508411.1   GKRIKEKFDQAYAGNNVGVHPRILFTY--KFLEEECKAAAFSRSLKGLQF
057 YP_009140788.1   GKRIKEEFDRKYPGVISGVVPRVLFAY--KFLENECKEAAFKRSLKNLSF
058 YP_009508410.1   --------------------------------------------------
059 NP_040333.1      QGFKNSPTLFDEALHRDLADFRIQHPD--LILLQYVDDLLLAATSELDCQ
060 YP_008992013.1   --PPKSKPKPKPQPRPVTINPSSTIIP--YQQSPGYTPASPSPPEPSKSQ
061 NP_041734.1      GHRMKEAFQKEYSGNEVGVFPRILFAY--RYLEQECKDAAFKRSLKSLSF
062 NP_056895.1      QGFINSPALFERALQEPLRQVSAAFSQ--SLLVSYMDDILIASPTEEQRS
063 YP_009109689.1   QGFKNSPTLFDEALHRDLTDFRTQHPE--VTLLQYVDDLLLAAPTKKACT
064 YP_002519387.1   --------------------------------------------------
065 YP_595725.1      GDRVKEAFKKKYPGVERHVPAATRFTY--DYLEEICTENNFQKQLRSLNF
066 YP_009508551.1   QGFLNSPALFTADVVDLLKEI--------PNVQVYVDDIYLSHDDPKEHV
067 YP_001497148.1   QGFKNSPTLFDEALHRDLTDFRTQHPE--VTLLQYVDDLLLAAPTEEACT
068 NP_041186.1      QGMTNSPTLCQKFVATAIAPVRQRFPQ--LYLVHYMDDILLAHTDEHLLY
069 NP_663784.1      QGWKGSPAIFQYTMRQVLEPFRKANPD--VILIQYMDDILIASDRTGLEH
070 YP_009508443.1   GKRVKEAFDQTYSGNTVGVHPRILFTY--KFLEEECKKAAFARSLKGLQF
```

| | | |
|---|---|---|
| 071 | YP_003987465.1 | GDRVRQAFLSRYPGNTIGVPARITFTQ--QYLEEVCREAAYQRSLKTLDF |
| 072 | NP_861410.1 | -------------------------------------------------- |
| 073 | YP_009345071.1 | GKRIEDSFKAKYPGNTVGVFPRIVFSF--KYLENQCKEASFQRSLKSLEF |
| 074 | YP_009508582.1 | QGFLNSPALFTADVVDLLKDI--------PGISVYVDDIYFSTETVSEHL |
| 075 | NP_569141.1 | -------------------------------------------------- |
| 076 | YP_002916057.1 | -------------------------------------------------- |
| 077 | NP_777384.2 | QGFINSPALFERALQEPLRQVSAAFSQ--SLLVSYMDDILIASPTEEQRS |
| 078 | YP_233110.1 | GDRVEKKFKEKYPNNTIGVAPRITFTR--NYIKEICQEAVFQSQLKRLDF |
| 079 | NP_542258.1 | QGFKNSPTLFEQQLSHILAPVRKAFPN--SLIIQYMDDILLASPALRELT |
| 080 | YP_009130664.1 | GPRIKEAYEARYPGNTIGVFPRVLFAY--KFLENECRDAAYKRSLKNLSF |
| 081 | YP_009508566.1 | QGFLNSPALFTADVVDLLKEI--------PNVNVYVDDIYVSTETINQHF |
| 082 | YP_006607892.1 | -------------------------------------------------- |
| 083 | YP_009506270.1 | GDAVEKAFKEKHPSNTVGIAARITFTK--RYLKELCEKVALQKSIGKMDF |
| 084 | NP_042513.1 | -------------------------------------------------- |
| 085 | YP_006907834.1 | -------------------------------------------------- |
| 086 | YP_004442827.1 | ADSAKTAFNKKFPGTAIHIPARIEFTY--NYLEEICTENNKQRKLGKLNF |
| 087 | YP_009408594.1 | GKHLEQKYNAAFPGNTIGVFPRIHFAY--QELAEMCKQAQLQRGLKDLAF |
| 088 | YP_009508561.1 | QGFLNSPALFTADVVDILKEV--------PGVSVYVDDIYISSPTMEEHF |
| 089 | NP_056907.1 | QGFKNSPTLFEQQLAAVLSPIRKTFPM--STIIQYMDDILLASPTQEELQ |
| 090 | YP_002308474.1 | QGFKNSPTLFEQQLSHILTPVRQAFPK--SIVIQYMDDILLASPTLEESI |
| 091 | YP_007761644.1 | -------------------------------------------------- |
| 092 | NP_040550.1 | QGMTCSPTICQLVVGQVLEPLRLKHPS--LRMLHYMDDLLLAASSHDGLE |
| 093 | NP_758808.1 | GKEIETAFIAKHGNANITVMPRIHFAY--HYLAELCKKAALQRSLKDLSF |
| 094 | YP_009345075.1 | GKKIEAAFNEKHPGNNTGVMPRIHFTY--HWLAELCKSAAMQRGIKSLDF |
| 095 | NP_068729.1 | -------------------------------------------------- |
| 096 | NP_047255.1 | QGFKNSPTLFDEALHSDLADFRVRYPA--LVLLQYVDDLLLAAATRTECL |
| 097 | YP_006273075.1 | RKTIKEDFDNKYPGVVSGVVPRVLFAY--KFLEKECRDAAFKRSLKNLSF |
| 098 | YP_009508888.1 | QGFLNSPALFTADVVDLMKHI--------PNVQVYVDDLYLSHDDPQEHL |
| 099 | NP_395469.1 | -------------------------------------------------- |
| 100 | YP_009508546.1 | QGFLNSPALFTADVVDLLKEI--------PNVQVYVDDIYLSHDDPQEHI |
| 101 | NP_056762.1 | GKVVKADFVKDGNSYEAGIGIAVSYIS--TWMRAKCIKETEAKTQKKASL |
| 102 | YP_009508556.1 | QGFLNSPALFTAHVVDLLKEI--------PHVPAYVDDIYISHDDPHEHL |
| 103 | NP_569153.1 | GQRMKKAYEEKHPGNIVGVCPRILFAY--KYLEGECKDAAFRRSLKNLSF |
| 104 | NP_043924.1 | QGFHASPGIFHQALYNGLASCKTAIESQGCKLLQYVDDILLMSEDRDHHL |
| 105 | YP_009243641.1 | QGMMNSPTICQLYVDAALKPIRKRYPF--LHIYHYMDDILFAGPKKEFLQ |
| 106 | YP_004300274.1 | -------------------------------------------------- |
| 107 | NP_056848.1 | -------------------------------------------------- |
| 108 | YP_006495799.1 | GPTIEKAYKERYPGLVIGVMARINFII--EYLQNVCKQAALQRSLKNLNF |
| 109 | YP_009508409.1 | -------------------------------------------------- |
| 110 | NP_687035.1 | QGWKGSPTIFQNTAANILEEIKRHTPG--LEIVQYMDDLWLASDHDETRH |
| 111 | NP_045937.2 | QGFIHSPTLFSQALYQSLHKIKFKIS---SEICIYMDDVLIASKDRDTNL |
| 112 | NP_612577.1 | -------------------------------------------------- |
| 113 | NP_619548.1 | -------------------------------------------------- |
| 114 | NP_056880.1 | QGMKNSPTLCQKFVDKAILTVRDKYQD--SYIVHYMDDILLAHPSRSIVD |
| 115 | YP_009121747.1 | SKKIEEAFRNKYPGLTAGVLPAIKFTH--TYITEMCTEAALQKELRDLSL |
| 116 | YP_009113237.2 | GVRIKEAFDREY-GEIIGVAPRVMFAY--GFLERQCREAAFARSLKGLDF |
| 117 | YP_009508408.1 | GKRVKELYEASYSGNTIGVHPRILFTY--KFLEEECKKAAFSRSLKGLQF |
| 118 | NP_955579.1 | QGFKNSPTLFDEALHSDLADFRVRYPA--LVLLQYVDDLLLAAATRTECL |
| 119 | YP_009229919.1 | GEEVEKAWNEKYPGAAAAVIPRIYFTY--QYLSKMCKQAALQREIKDLSF |
| 120 | YP_004442833.1 | GDQARKAFDAKFPGTHIHVPARIDFTY--NYLEEVCTENNRQRKLAKLDF |
| 121 | NP_862833.2 | QGMTNSPTLCQKFVAAAIAPVRQCFPQ--LYLVHYMDDILLAHINEHLLY |
| 122 | NP_040563.1 | QGWVCSPAIYQTTTQKIIENIKKSHPD--VMLYQYMDDLLIGSNRD--DH |
| 123 | YP_443922.1 | QGYRCSPGIFNRVLKDHLKEIVIPD-G--VVLIQYVDDLLLAACSSDQCL |
| 124 | NP_057933.2 | QGFKNSPTLFDEALHRDLADFRIQHPD--LILLQYVDDLLLAATSELDCQ |
| 125 | YP_009116631.1 | GPTIEQAYKDRYPGLTIGVLARANFIL--EYLQNVCKQAALQRSLKSLSF |
| 126 | YP_009513249.1 | QGFLNSPGLFTGDVVDLLQGI--------PNVEVYVDDVYISHDSEKEHL |
| 127 | YP_223871.1 | QGFKNSPTLFDVALNRDLQGFRLDHPS--VSLLQYVDDLLIAADTQAACL |
| 128 | NP_127504.1 | KSPNTFDLAKLVWEKKDAIAAEKRAKK--KLQKDEVKQKTSLPPESKRPD |
| 129 | NP_044929.1 | QGFLNSPALFTADVVDILKDI--------PNVQVYVDDVYVSSATEQEHL |
| 130 | NP_057860.1 | QGFKNSPTLFEMQLAHILQPIRQAFPQ--CTILQYMDDILLASPSHADLQ |
| 131 | NP_040973.1 | QGWILSPLIYQSTLDNIIQPFIRQNPQ--LDIYQYMDDIYIGSNLSKKEH |
| 132 | NP_777317.1 | GDRIEKAFTEKHPTNTIGVAARIAFTR--NYLREICQEAVFQSSLKRLGF |
| 133 | NP_054716.1 | QGFLNSPALFTADVVDLLKDI--------PNVEVYVDDVYFSNDTEEEHL |
| 134 | YP_001856242.1 | QGWKGSPAIFQSSMTKILEPFRKQNPD--IVIYQYMDDLYVGSDLEIGQH |
| 135 | NP_954565.2 | QGMANSPTLCQKYVATAIHKVRHAWKQ--MYIIHYMDDILIAGKDGQQVL |
| 136 | NP_569150.1 | GDEVEKAFMEKHPSNTVGITARITFTK--RYLKELCEKVALQKSIGKMDF |
| 137 | NP_659397.1 | -------------------------------------------------- |
| 138 | YP_009513211.1 | QGFKNSPTLFDEALHRDLASFRALNPQ--VVMLQYVDDLLVAAPTYRDCK |
| 139 | YP_233107.1 | GDKVEKAFKERHPTNSIGVTARIAFTR--NYLKDMCQEALFQSQLKKMNF |
| 140 | YP_009109692.1 | QGFKNSPTLFDEALHRDLTDFRTQHPE--VTLLQYVDDLLLAAPTKKACT |
| 141 | NP_057849.4 | QGWKGSPAIFQSSMTKILEPFRKQNPD--IVIYQYMDDLYVGSDLEIGQH |
| 142 | NP_049560.1 | -----------MQLAHILQPIRQAFSQ--CTILQYMDDILLASPSHKDLQ |
| 143 | YP_009506267.1 | GKRIKTKFEEQYAGNTIGVHPRIMFIY--KFLEEECKNAAFMRSMKNLQF |

```
144 NP_039820.1      GERVAQAYKIMDPTSAVNLPSRVYFTI--NYLTEQCKIASYMRSLKALDF
145 NP_056790.1      QGFKNSPTLFDEALHRDLAPFRALNPQ--VVLLQYVDDLLVAAPTYEDCK
146 YP_002321513.1   GEDIEKNYKQVH-GDLPGVAPRIYFTY--KYLQEMCKNAEKQRGLKSLNF
147 NP_955591.1      QGFKNSPTLFDEALHRDLADFRIQHPD--LILLQYVDDLLLAATSELDCQ
148 YP_001956722.2   QGFLNSPALFTADVVDLLKEV--------PNVQVYVDDIYISHDDPREHL
149 NP_056902.1      QGFVLSPYIYQKTLQEILQPFRERYPE--VQLYQYMDDLFMGSNGSKKQH
150 NP_056886.1      QGMTCSPTICQLVVGQVLEPLRLKHPS--LCMLHYMDDLLLAASSHDGLE
151 NP_041261.1      QGMANSPTLCQKFVAAAIAPVRSQWPE--AYILHYMDDILLACDSAEAAK

001 YP_004442836.1   CRN---TPVHGLYGRERSH--KRYGARKSTSYKGKPHK-SHVRIGKKKHL
002 NP_056728.1      --------------------------------------------------
003 YP_009508537.1   EYL---EILFNRLNEAGYIVSLKKSNIANSIVDFLGFQITNEGRGLTDTF
004 YP_004732983.2   CKA---IPIPGYYDQGV---KKKYGLRKSKTYKGKPHD-SHVKVIKNKYK
005 YP_009508407.1   --------------------------------------------------
006 NP_040939.1      REI---VKDLANYIAQYGFTLPEEKRQKGYPAKWLGFELHPQTWKFQKHT
007 YP_003864102.1   QCF---AQLKEKLVIAGLHIAPEKIQLHD-PYTYLGFQLNGPKITTQKA-
008 YP_009002585.1   CSE---MPLPGYYGEK-----KRYGVRRSTTYKGKPHP-THARIEKQKHL
009 YP_003284237.1   CKE---FPVPGYY-KKPGY---KYGRRKTTTYKGKPHK-THVRIDKRKHL
010 YP_009268869.1   REI---VEELASYIAQYGFMLPEEKRQEGYPAKWLGFELHPEKWRFQKHT
011 YP_009506251.1   CKD---MPLTGYYDKS-----SRYGMRRSKTYKGKPHA-SHARIEKRKHL
012 NP_955611.1      AAG---EEVISTLERAGFTISPDKVQREP-GVQYLGYKLG-STYVAPVG-
013 YP_008567619.1   CNS---IPIPGYYS-KE----KKMGVRRSTTYKGKPHS-SHVRIEKRKHL
014 NP_149413.1      CKG---FPVVNPVGTKKYG--KRYGTRKARSYRGKPHS-SHVRIERTRYL
015 YP_004442824.1   CRN---TPVHGLYGKDKPY--KKYGARKSQNYHGKPHK-SHVRIAKRKHL
016 YP_001931967.1   --------------------------------------------------
017 YP_002455786.1   QLA---TLTAQLLSSHGLPVSQEKTQRTPGKIHFLGQIIHPDHITYETTP
018 YP_009109694.1   QGT---RHLLQELGEKGYRASAKKAQICQTKVTYLGYILSEGKRWLTPGR
019 NP_789739.1      RTK---IEELRQHLLRWGLTTPDKKHQKEPPFLWMGYELHPDKWTVQPIV
020 NP_955619.1      LLS---EATMASLISHGLPVSENKTQQTPGTIKFLGQIISPNHLTYDAVP
021 YP_224289.1      CRE---FPIPGYY-KKPGR---KFGVRKSTSYKGKPHK-THIKIDKRKYL
022 YP_009182100.1   KPN---EFNTERYKPRIEPNPAVSPNGVWLDIDNNPDPRGNIDEWVNTMN
023 NP_705927.1      RTK---IEELRQHLLRWGLTTPDKKHQKEPPFLWMGYELHPDKWTVQPIV
024 YP_009508577.1   KIL---EKVFKILLEAGYIVSLKKSALLRYEVTFLGFSITQTGRGLTSEF
025 YP_004442830.1   CRG---FPVINPL-SKKY---KSLGTRRTTTYKGKPHH-THARIDRRKYL
026 NP_758887.1      SRM---IQELRDHLLFWGLETPDKKFQKEPPFEWMGYILHPKKWTVQKVQ
027 YP_009508406.1   --------------------------------------------------
028 YP_610965.1      CRR---IPLPGYYKDGP---KKKLGLRKAKNYRGKPHD-THVRLVKNK--
029 YP_001931961.1   --------------------------------------------------
030 YP_241114.1      QAF---SILKQHLSLNGLVIADEKIQTHF-PYNYLGFSLYPRTYNTQLV-
031 YP_006732334.1   --------------------------------------------------
032 YP_009041481.1   CAQ---VPIPGDYNRE----GKKYGLRRASTYKGKPHA-THVRTFKRKDA
033 YP_001036293.1   CSK---IPIPGYYKGRE----KKYGVRKSTTYKGKPHD-SHVRIERRKHL
034 YP_605811.1      CKG---TPVHGLYGKDKAYG-RKYGVRKSTSYKGKPHK-SHVRIDKKKHL
035 NP_955577.1      EGT---KALLETLGNKGYRASAKKAQICLQEVTYLGYSLKDGQRWLTKAR
036 YP_003208050.1   CRK---FPIPGYYQK-S---QKKYGLRKSQSYKGKPHD-THVRVVKNK--
037 NP_789740.1      RTK---IEELRQHLLRWGLTTPDKKHQKEPPFLWMGYELHPDKWTVQPIV
038 YP_004442839.1   CKG---TPVHGLYGKDKSYG-RRYGVRKSTSYKGKPHK-SHVRIDKKKHL
039 YP_009507791.1   EKI---IQELRDLLIFWGFETPEDKLQQEPPYKWMGYTLYPNKWTIQKTK
040 YP_004581513.1   --------------------------------------------------
041 NP_043933.1      --------------------------------------------------
042 YP_009507248.1   ENL---ERVFQRLRESNFKIQMDKSEFLKLETAYLGHIISRDGIKPNPDK
043 YP_002117531.1   CSS---FPIPNYQ-KGAGR---KFGLRKSKSYKGKPHK-THVRIEKSKYL
044 YP_004347415.1   --------------------------------------------------
045 NP_056891.1      QCF---DQLKQELTAAGLHIAPEKVQLQD-PYTYLGFELNGPKITNQKA-
046 YP_009513242.1   EQL---EKVFSILLNAGYVVSLKKSEIAQREVEFLGFNITKEGRGLTETF
047 YP_009352866.1   CSQ---IPIPGDYNRE----GKKYGLRKAGTYKGKPHN-THVRAFKKKDQ
048 YP_009165750.1   --------------------------------------------------
049 NP_040840.1      RGI---VNELASYIAQYGFMLPEDKRQEGYPAKWLGFELHPEKWKFQKHT
050 YP_004222728.1   AAG---EEVISTLERAGFTISPDKVQREP-GVQYLGYKLG-STYVAPVG-
051 YP_009506264.1   CKE---FPIPGYYHKKPTK---KYGLRKTTTYRGKPHK-THVRIDKSKHL
052 YP_567050.1      QLS---KMTLQALVTHGLPVSQEKTQQTPGQIRFLGQVISPNHITYETTP
053 YP_009508571.1   DQL---EKVFQILLQAGYVVSLKKSEVAQKTVEFLGFNITKEGRGLTEAF
054 NP_056803.1      QQL---EKVFQILLQAGYVVSLKKSEIGQRTVEFLGFNITKEGRGLTDTF
055 NP_955564.1      EIL---TSMIQALNKHGLVVSTEKIQKYD-NLKYLGTHIQGDSVSYQKL-
056 YP_009508411.1   CSQ---IPIPGYYKNSG----KKYGVRRSTTYKGKPHE-THVRIEKKKYL
057 YP_009140788.1   CSS---IPIPGYYKNTG----KKYGVRRSKTYKGKPHE-SHARIEKRKHL
058 YP_009508410.1   --------------------------------------------------
059 NP_040333.1      QGT---RALLQTLGDLGYRASAKKAQICQKQVKYLGYLLKEGQRWLTEAR
060 YP_008992013.1   PPT---ASKDKEPMHQFS-----AHTIDHPSSTDDQTSDSNLAVSDSHTE
061 NP_041734.1      CKD---MPLTGYYDKT-----SKYGMRKSRTYKGKPHA-SHARVEKRKHL
062 NP_056895.1      QCY---QALAARLRDLGFQVASEKTRQTPSPVPFLGQMVHNQIVTYQSLP
```

| | | |
|---|---|---|
| 063 | YP_009109689.1 | Q G T - - - R H L L Q E L G E K G Y R A S A K K A Q I Q Q T K V T Y L G Y I L S E G K R W L T P G R |
| 064 | YP_002519387.1 | - - - - - - - - - - - - - - - - - - - - - - - - - - - - - - - - - - - - - - - - - - - - - - - - - - |
| 065 | YP_595725.1 | C K G - - - F P V V N P V G T R K Y G - - K K Y G T R K A R S Y R G K P H K - S H V R I E K K K Y L |
| 066 | YP_009508551.1 | Q Q L - - - E K V F Q I L L Q A G Y V V S L K K S E I G Q K T V E F L G F N I T K E G R G L T D T F |
| 067 | YP_001497148.1 | R G T - - - K H L L R E L G D K G Y R A S A K K A Q I Q Q T K V T Y L G Y I L S E G K R W L T P G R |
| 068 | NP_041186.1 | Q A F - - - S I L K Q H L S L N G L V I A D E K I Q T H F - P Y N Y L G F S L Y P R V Y N T Q L V - |
| 069 | NP_663784.1 | D K V - - - V L Q L K E L L N G L G F S T P D E K F Q K D P P F Q W M G C E L W P T K W K L Q K L Q |
| 070 | YP_009508443.1 | C N Q - - - I P I P G Y Y K G K E - - - - K R Y G A R R S T T Y K G K P H D - S H V R I E K K K H L |
| 071 | YP_003987465.1 | C K E - - - F P I P N F Y - K G P Q K - - - K Y G V R K S T T Y R G K P H K - S H I R I D K T K H L |
| 072 | NP_861410.1 | - - - - - - - - - - - - - - - - - - - - - - - - - - - - - - - - - - - - - - - - - - - - - - - - - - |
| 073 | YP_009345071.1 | C K Q - - - M P I P G Y Y G T D R - - - - K R L G I R K A T H Y K G K P H S - T H V K I E K H K H L |
| 074 | YP_009508582.1 | K I L - - - E K V F K I L L E A G Y I V S L K K S A L L R H E V T F L G F S I T Q T G R G L T S E F |
| 075 | NP_569141.1 | - - - - - - - - - - - - - - - - - - - - - - - - - - - - - - - - - - - - - - - - - - - - - - - - - - |
| 076 | YP_002916057.1 | - - - - - - - - - - - - - - - - - - - - - - - - - - - - - - - - - - - - - - - - - - - - - - - - - - |
| 077 | NP_777384.2 | Q C Y - - - Q A L A A R L R D L G F Q V A S E K T R Q T P S P V P F L G Q M V H N Q I V T Y Q S L P |
| 078 | YP_233110.1 | C K G - - - T P V H G L Y G K E K A Y G - R K Y G V R K S T S Y K G K P H K - S H V R I D K K K H L |
| 079 | NP_542258.1 | A L T - - - D K V T N A L T K E G L P M S L E K T Q A T P G S I H F L G Q V I S P D C I T Y E T L P |
| 080 | YP_009130664.1 | C S S - - - I P I P G Y Y K - I E - - - - K K Y G V R R S R T Y K G K P H A - S H A R I E R R K H L |
| 081 | YP_009508566.1 | Q V L - - - D K I F Q K L L Q A G Y V V S L K K S N L Q R Y E V T F L G F T I S K Y G R G L T E E F |
| 082 | YP_006607892.1 | - - - - - - - - - - - - - - - - - - - - - - - - - - - - - - - - - - - - - - - - - - - - - - - - - - |
| 083 | YP_009506270.1 | C R N - - - T P V H G L Y G K E K S Y - - R K Y G A R K S T S Y K G K P H K - S H V R I G K K K H L |
| 084 | NP_042513.1 | - - - - - - - - - - - - - - - - - - - - - - - - - - - - - - - - - - - - - - - - - - - - - - - - - - |
| 085 | YP_006907834.1 | - - - - - - - - - - - - - - - - - - - - - - - - - - - - - - - - - - - - - - - - - - - - - - - - - - |
| 086 | YP_004442827.1 | C K G - - - F P V V N P M - T R K Y - - - K S L G T R R T T T Y K G K P H K - T H A R I D K K K Y L |
| 087 | YP_009408594.1 | C S Q - - - I P L P G Y Y G P - - - - - R K K Y G L R K A K N Y S G K P H K - S H V R V I K K K Y Q |
| 088 | YP_009508561.1 | Q V L - - - D S I F R K L L E T G Y I V S L K K S A L A R Y E V N F L G F V I S E T G R G L T S E F |
| 089 | NP_056907.1 | Q L S - - - K M T L Q A L V T H G L P V S Q E K T Q Q T P G Q I R F L G Q V I S P N H I T Y E T T P |
| 090 | YP_002308474.1 | V L A - - - Q E I T N A L A Q E G L P M S T E K T Q S T P G P I H F L G Q T I S K K Y I T Y E T L P |
| 091 | YP_007761644.1 | - - - - - - - - - - - - - - - - - - - - - - - - - - - - - - - - - - - - - - - - - - - - - - - - - - |
| 092 | NP_040550.1 | A A G - - - E E V I S T L E R A G F T I S P D K I Q R E P - G V Q Y L G Y K L G - S T Y V A P V G - |
| 093 | NP_758808.1 | C N Q - - - I P L P G I Y T K - G - - - N K K F G L R K A R T Y K G K P H P - T H V R V F K - - - K |
| 094 | YP_009345075.1 | C K D - - - I P I P G V Y P R - - - - - E K K Y S L R K A K N Y K G K P H D - T H V R V F K R K - - |
| 095 | NP_068729.1 | - - - - - - - - - - - - - - - - - - - - - - - - - - - - - - - - - - - - - - - - - - - - - - - - - - |
| 096 | NP_047255.1 | E G T - - - K A L L E T L G N K G Y R A S A K K A Q I Q L Q E V T Y L G Y S L K D G Q R W L T K A R |
| 097 | YP_006273075.1 | C S S - - - I P I P G Y Y K N T G - - - - K K Y G V R R S K T Y K G K P H E - S H A R I E K R K H L |
| 098 | YP_009508888.1 | Q V L - - - Q Q V L H I L H D A G Y V V S L K K S A I A Q K V V E F L G F N I T K T G R G L T D A F |
| 099 | NP_395469.1 | - - - - - - - - - - - - - - - - - - - - - - - - - - - - - - - - - - - - - - - - - - - - - - - - - - |
| 100 | YP_009508546.1 | Q Q L - - - E K V F Q I L L Q A G Y V V S L K K S E I G Q K T V E F L G F N I T K E G R G L T D T F |
| 101 | NP_056762.1 | A F C R S I Y T I G D Y K K R K I L K R V T N Y N K N R R K N Y V R R P S I K K K C R C Y I C Q D E |
| 102 | YP_009508556.1 | E Q L - - - E K V F S I L L N A G Y V V S L K K S E I A Q R E V E F L G F N I T K D G R G L T E T F |
| 103 | NP_569153.1 | C S S - - - I P I P G Y Y G G K S G E - - K R Y G V R R T T T Y K G K P H S - T H A R I E K T K H L |
| 104 | NP_043924.1 | R S L - - - A I L L Q G L K D L G V K I N P K K S H F Q K D Q V Q Y L G V N V G A D T R S L I D A R |
| 105 | YP_009243641.1 | D L L - - - E Q L P L Y F K P Y G L I I A P E K I Q L E D - I S H Y L G F V V S R T T V H P Q K M - |
| 106 | YP_004300274.1 | - - - - - - - - - - - - - - - - - - - - - - - - - - - - - - - - - - - - - - - - - - - - - - - - - - |
| 107 | NP_056848.1 | - - - - - - - - - - - - - - - - - - - - - - - - - - - - - - - - - - - - - - - - - - - - - - - - - - |
| 108 | YP_006495799.1 | C R T - - - M P V P G Y Y E K - - - - - - K K Y G V R K S T T Y K G K P H D - T H V K V I K N K D K |
| 109 | YP_009508409.1 | - - - - - - - - - - - - - - - - - - - - - - - - - - - - - - - - - - - - - - - - - - - - - - - - - - |
| 110 | NP_687035.1 | N Q Q - - - V D I V R K M L L E K G L E T P D K K V Q R E P P W E W M G Y K L H P N K W T I N K I E |
| 111 | NP_045937.2 | K D T - - - A V M L Q H L A S E G H K V S K K K I Q L Q Q Q E V V Y L G Q L L T P E G R K I L P D R |
| 112 | NP_612577.1 | - - - - - - - - - - - - - - - - - - - - - - - - - - - - - - - - - - - - - - - - - - - - - - - - - - |
| 113 | NP_619548.1 | - - - - - - - - - - - - - - - - - - - - - - - - - - - - - - - - - - - - - - - - - - - - - - - - - - |
| 114 | NP_056880.1 | E I L - - - T S M I Q A L N K H G L V V S T E K I Q K Y D - N L K Y L G T H I Q G D S V S Y Q K L - |
| 115 | YP_009121747.1 | C S S - - - I P I P G Y Y D R - Q - - - R K K Y G I R K S R T Y K G K P H D K T H V K V F K R K Y K |
| 116 | YP_009113237.2 | C K D - - - I P I P G Y Y Q S R R N T - G K R N S L R R S Q T Y K G K P H S - T H I R I D K A K Y L |
| 117 | YP_009508408.1 | C N Q - - - I P I P G Y Y K G K E - - - - K R Y G V R R S T T Y K G K P H D - S H V R I E K K K H L |
| 118 | NP_955579.1 | E G T - - - K A L L E T L G N K G Y R A S A K K A Q I Q L Q E V T Y L G Y S L K D G Q R W L T K A R |
| 119 | YP_009229919.1 | C K E - - - V H I P G Y Y K T - - - - - G K R Y G L R K A K T Y K G K P H D - T H V R V F K K K - - |
| 120 | YP_004442833.1 | C R G - - - F P V I N P L - Q R K Y - - - R S L G T R R T R T Y K G K P H S - S H A R I D K R K Y L |
| 121 | NP_862833.2 | Q A F - - - S L L K Q H L S L N G L V X A D E K I Q T H F - P Y N Y L G F H L Y P R A Y T T Q L V - |
| 122 | NP_040563.1 | K Q I - - - V Q E I R D K L G S Y G F K T P D E K V Q E E - R V K W I G F E L T P K K W R F Q P R Q |
| 123 | YP_443922.1 | A A T - - - K I L L D F L A T K G Y K V K R S K V Q T Q R R T V L F L G R E V S A E G A G L S T A H |
| 124 | NP_057933.2 | Q G T - - - R A L L Q T L G N L G Y R A S A K K A Q I Q Q K Q V K Y L G Y L L K E G Q R W L T E A R |
| 125 | YP_009116631.1 | C R N - - - M P V P G Y Y E K - - - - - - K Q Y G I R K A K T Y K G K P H P - T H V K V I K N K Y K |
| 126 | YP_009513249.1 | E Y L - - - D I L F N R L K E A G Y I I S L K K S N I A N S I V D F L G F Q I T N E G R G L T D T F |
| 127 | YP_223871.1 | S A T - - - R D L L M T L A E L G Y R V S G K K A Q L Q Q E E V T Y L G F K I H K G S R S L S N S R |
| 128 | NP_127504.1 | P Q S - - - S S H L G D Q F M I S D P A L P K V Y A L N E P S V P S E D T S - S Q S Y I S T E E S V |
| 129 | NP_044929.1 | D I L - - - E T I F N R L S T A G Y I V S L K K S K L A K E T V E F L G F S I S Q N G R G L T D S Y |
| 130 | NP_057860.1 | L L S - - - E A T M A S L I S H G L P V S E N K T Q Q T P G T I K F L G Q I I S P N H L T Y D A V P |
| 131 | NP_040973.1 | K E K - - - V E E L R K L L L W W G F E T P E D K L Q E E P P Y T W M G Y E L H P L T W T I Q Q K Q |
| 132 | NP_777317.1 | C K S - - - T P V H G V Y G E R K S Y K - G R F G A R K S T T Y N G K P H K - S H V R I A R K K H L |
| 133 | NP_054716.1 | K T M - - - D L L F Q K L Q T A G Y I V S L K K S K L G Q H T V D F L G F Q I T Q T G R G L T D S Y |
| 134 | YP_001856242.1 | R T K - - - I E E L R Q H L L R W G L T T P D K K H Q K E P P F L W M G Y E L H P D K W T V Q P I V |
| 135 | NP_954565.2 | Q C F - - - D Q L K Q E L T A A G L H I A P E K V Q L Q D - P Y T Y L G F E L N G P K I T N Q K A - |

```
136 NP_569150.1       CRS---TPVHGLY-RDKSY--RKYGARKSTSYKGKPHK-SHVRIGKKKYL
137 NP_659397.1       --------------------------------------------------
138 YP_009513211.1    EGT---RRLLQELSKLGYRVSAKKAQLQREEVTYLGYLLKGGKRWLTPAR
139 YP_233107.1       CGN---TPVHGIYGKNKEGGFRKYGVRKNTSYKGKPHK-SHIRIGKQKHL
140 YP_009109692.1    QGT---RHLLQELGEKGYRASAKKAQIQQTKVTYLGYILSEGKRWLTPGR
141 NP_057849.4       RTK---IEELRQHLLRWGLTTPDKKHQKEPEFLWMGYELHPDKWTVQPIV
142 NP_049560.1       LLS---EVTMASLISHGLPVSENKTQQTPGTIKFLGQIISPNHLTYDAVP
143 YP_009506267.1    CSK---IPIPGYYGGAE----KKYGVRKSKTYKGKPHQ-SHVRIEKRKHL
144 NP_039820.1       CRD---FPIEGYYGRSGEK--KKYTARKATKYTGKAHD-NHIRVTKAKY-
145 NP_056790.1       KGT---QKLLQELSKLGYRVSAKKAQLQQREVTYLGYLLKEGKRWLTPAR
146 YP_002321513.1    CKD---IRLPGQYARDS---SKRYGMRKANTYKGKPHR-THARVIKNKEK
147 NP_955591.1       QGT---RALLQTLGNLGYRASAKKAQIQQKQVKYLGYLLKEGQRWLTEAR
148 YP_001956722.2    EQL---EKVFSLLLNAGYVVSLKKSEIAQHEVEFLGFNITKEGRGLTETF
149 NP_056902.1       KEL---IIELRAILLEKGFETPDDKLQEVPEYSWLGYQLCPENWKVQKMQ
150 NP_056886.1       AAG---EEVISTLERAGFTISPDKVQREP-GVQYLGYKLG-STYVAPVG-
151 NP_041261.1       ACY---AHIISCLTSYGLKIAPDKVQVSE-EFSYLGFELHHQQVFTPRV-

001 YP_004442836.1    AAR-----KRNCRCYACGEEGHFASECT-NQKKIIDRVRVLDSLDLEDGL
002 NP_056728.1       --------------------------------------------------
003 YP_009508537.1    KEKLENVTAPTTLKQLQSILGLLNFARNFIPDFTELIAPLYALIPKSTK-
004 YP_004732983.2    GAQG-----RKCKCYLCGIEGHYARECPKKHVR-PERAAYFEGMGLDVNW
005 YP_009508407.1    --------------------------------------------------
006 NP_040939.1       LPELTKG--TITLNKLQKLVGELVWR---QSIIGKSIPNILKLMEGDREL
007 YP_003864102.1    VIRKDK---LQTLNDYQKLLGDINWLRPYLKLTTGDLRPLFDILKGDPNP
008 YP_009002585.1    VR------NKRCKCYLCGEMGHFARECP-NDKKNTRRVAMYEQLNIPEGF
009 YP_003284237.1    KNR-----K--CKCYACGEEGHYASECR-NKKQYTDRVNIIENIALKEDE
010 YP_009268869.1    LPEIKEG--TITLNKLQKLVGDLVWR---QSLIGKSIPNILKLMEGDRAL
011 YP_009506251.1    VR------NKRCKCYLRGDEGHFARECP-NNKRDVKRVAIFEGINLPDDY
012 NP_955611.1       LVAEPR---IATLWDVQKLVGSLQWLRPALGIPPRLMGPFYEQLRG-SDP
013 YP_008567619.1    LR------NKNCKCYLCGEEGHFAKDCN-RNKRDIRRVAIFEGLELPEDH
014 NP_149413.1       KQRKDGKGEKKCRCFVCGSTEHLMKDCK-SPMKRQDRVNLAREIDIPDNY
015 YP_004442824.1    AMR-----KKDCKCFACGETGHFASECK-NPRKLVERVAIMDSLELPEDV
016 YP_001931967.1    --------------------------------------------------
017 YP_002455786.1    TIPIKA---HWTLTELQTLLGELQWVSKGTPVLREHLHCLYSALRGLKDP
018 YP_009109694.1    IETVARIPPPRNPREVREFLGTAGFCRLWIPGFAELAAPLYALTKES---
019 NP_789739.1       LPEKD----SWTVNDIQKLVGKLNWAS--QIYPGIKVRQLCKLLRGTKAL
020 NP_955619.1       KVPIRS---RWALPELQALLGEIQWVSKGTPTLRQPLHSLYCALQRHTDP
021 YP_224289.1       RNK-----K--CKCYACGEEGHYASECR-NPRKLMERVSVLQDLELGKDM
022 YP_009182100.1    MLIVSETGKAWSPDEARTYYACTFKGIARKWFDKYKETDEYVLWLSTIER
023 NP_705927.1       LPEKD----SWTVNDIQKLVGKLNWAS--QIYPGIKVRQLCKLLRGTKAL
024 YP_009508577.1    KDKIQNITSPRTLKELQSILGLFNFARNFVPNFSEIIKPLYSLISTAEG-
025 YP_004442830.1    QAR-----KKNCRCYVCGEEGHFAKECK-SQKKIQERVNIVNELDIPEGY
026 NP_758887.1       LPEKE----EWTVNDIQKLVGKLNWAS--QIYSGIKTKELCRLIRGAKPL
027 YP_009508406.1    --------------------------------------------------
028 YP_610965.1       -D---KGRTTKCRCYICGQEGHFARECR-NGRGNIARAAIIKDLDLPSDF
029 YP_001931961.1    --------------------------------------------------
030 YP_241114.1       KLQTDH---LKTLNDFQKLLGDINWIRPYLKLPTYTLQPLFDILKGDSDP
031 YP_006732334.1    --------------------------------------------------
032 YP_009041481.1    GKQ------RKCACFICGEPDHFAKECPRNGKGNINRVHYYDNLNIPEDW
033 YP_001036293.1    IR------SKKCKCFLCGAEGHFARECP-NDKRNVKRIAIFEGMEIPEDC
034 YP_605811.1       LMK-----RKDCKCFACGDVGHFASECP-NPKKLMHRVQILQSLELDDGI
035 NP_955577.1       KEAILSIPVPKNSRQVREFLGTAGYCRLWIPGFAELAAPLYPLTRPG---
036 YP_003208050.1    -D---KARSVKCKCFICGEEGHFARECR-SKKGNIARAAVVDNLDLPPDF
037 NP_789740.1       LPEKD----SWTVNDIQKLVGKLNWAS--QIYPGIKVRQLCKLLRGTKAL
038 YP_004442839.1    TMK-----RKDCKCFACGDIGHFASECP-NPKKLMHRVQILQSLELEDGI
039 YP_009507791.1    L-DIPE---VPTLNQLQKLAGVINWAT--QNVGGIKIKALTELMKGNQQL
040 YP_004581513.1    --------------------------------------------------
041 NP_043933.1       --------------------------------------------------
042 YP_009507248.1    ISAIQKYLIPKTPKEIKQFLGLLGYYRKFIPDFARLTKPLTQCLKKGSKV
043 YP_002117531.1    RSK-----K--CKCYACGEEGHFAKDCK-NPRKLTERVAILEDLDLGDDY
044 YP_004347415.1    --------------------------------------------------
045 NP_056891.1       VIRKDK---LQTLNDFQKLLGDINWLRPYLKLTTGDLKPLFDTLKGDSDP
046 YP_009513242.1    KQKLLNVIPPKDLKQLQSILGLLNFARNFIPNYSELVKPLYTIVANANG-
047 YP_009352866.1    QKQ------RKCACFICGEPDHFARECPRSGKGNINRVHYYDNLNIPQDW
048 YP_009165750.1    --------------------------------------------------
049 NP_040840.1       LPEITEG--PITLNKLQKLVGDLVWR---QSLIGKSIPNILKLMEGDRAL
050 YP_004222728.1    LVAEPR---IATLWDVQKLVGSLQWLRPALGIPPRLMGPFYEQLRG-SDP
051 YP_009506264.1    KNR-----K--CKCYACGEEGHFARDCR-NPRKIMDRVNVLEDLELKDGM
052 YP_567050.1       TIPIKS---QWTLTELQTVLGEIQWVSKGTPILRKHLQCLYSALRGYQDP
053 YP_009508571.1    KAKLLDITPPKDLKQLQSILGLLNFARNFILNFAELVKPLYSLISSAKG-
054 NP_056803.1       KTKLLNVTPPKDLKQLQSILGLLNFARNFIPNFAELVQTLYNLIASSKG-
```

```
055 NP_955564.1     QIRTDK---LRTLNDFQKLLGNINWIRPFLKLTTGELKPLFEILNGDSNP
056 YP_009508411.1  ERGS-ALKNKKCKCFLCGEEGHFARECS-SDRRNIKRVAIFEGIELPEDC
057 YP_009140788.1  VR------NKRCKCFLCGQEGHFARECP-NDRKSSKRMAMFENLELPDDC
058 YP_009508410.1  --------------------------------------------------
059 NP_040333.1     KETVMGQPTPKTPRQLREFLGTAGFCRLWIPGFAEMAAPLYPLTKTG---
060 YP_008992013.1  IDTESSASTSDSEKSYADITRILMAQPD------PQTSRTEPYVNIPSDV
061 NP_041734.1     IR------NKKCKCYLCGDEGHFARECP-NQRRDVKRVAIFEGIDLPEGF
062 NP_056895.1     TLQISS---PISLHQLQAVLGDLQWVSRGTPTTRRPLQLLYSSLKGIDDP
063 YP_009109689.1  IETVARIPPPRNPREVREFLGTAGFCRLWIPGFAELAAPLYALTKES---
064 YP_002519387.1  --------------------------------------------------
065 YP_595725.1     QQR-----EKKCRCYVCGSPDHLMKDCK-SPMKRQERVNLANELDIPDGY
066 YP_009508551.1  KTKLLNITPPKDLKQLQSILGLLNFARNFIPNFAELVQPLYNLIASAKG-
067 YP_001497148.1  IETVARIPPPQSPREVREFLGTAGFCRLWIPGFAELAAPLYALTKES---
068 NP_041186.1     KLQTDH---LKTLNDFQKLLGDINWIRPYLKLPTYTLQPLFDILKGDSDP
069 NP_663784.1     LPQKD----IWTVNDIQKLVGVLNWAA--QIYSGIKTKHLCRLIRGKMTL
070 YP_009508443.1  VR------ERRCKCFLCGAEGHFARECP-NDKRNIKRVAVFEGLEIPEDC
071 YP_003987465.1  KNR-----K--CKCYICGEEGHFARDCK-NPKKIVDRVHVLEELDIQDGM
072 NP_861410.1     --------------------------------------------------
073 YP_009345071.1  RT-------KKCKCYLCGEEGHFARECK-NDNRNIHRQTMYQSLELPKEF
074 YP_009508582.1  KDKIQNITPPKTLKELQSILGLFNFARNFVPNFSEIIKPLYSLISTAEG-
075 NP_569141.1     --------------------------------------------------
076 YP_002916057.1  --------------------------------------------------
077 NP_777384.2     TLQISS---PISLHQLQAVLGDLQWVSRGTPTTRRPLQLLYSSLKGIDDP
078 YP_233110.1     LMK-----RKDCKCFACGDIGHFASECP-NPKKLMHRVQILQSLELDDGI
079 NP_542258.1     SIHVKS---IWSLAELQSMLGELQWVSKGTPVLRSSLHQLYLALRGHRDP
080 YP_009130664.1  QR------NKRCKCYLCGQEGHFARECP-NDRRSIKRVAIFENLDLPEDC
081 YP_009508566.1  QEKLRNISPPNSLKQLQSILGLLNFARNFIPNFSELIKPLYELISTAQG-
082 YP_006607892.1  --------------------------------------------------
083 YP_009506270.1  ALR-----KKNCRCYACGEEGHFASECR-NPRKIMDRVKVLDSLELEDGL
084 NP_042513.1     --------------------------------------------------
085 YP_006907834.1  --------------------------------------------------
086 YP_004442827.1  QGR-----KKDCRCYVCGEEGHFAKECR-SKKRIQERVNIVQELDIPDGY
087 YP_009408594.1  HD---RGRVKKCKCFICGKEGHFARECR-NNVGNKERVAMLNELDLPEDY
088 YP_009508561.1  RERLQEITPPTTLKQLQSILGFLNFARNFVPNFSELVQPLYQLISTASG-
089 NP_056907.1     TIPIKS---QWTLTELQTVLGEIQWVSKGTPILRKHLQCLYSALRGYQDP
090 YP_002308474.1  TIHVKP---NWTLTELQSTLGELQWVSKGTPTLRSSLHQLYTALRGHHDP
091 YP_007761644.1  --------------------------------------------------
092 NP_040550.1     LVAEPR---IATLWDVQKLVGSLQWLRPALGIPPRLMGPFYEQLRG-SDP
093 NP_758808.1     AK---YQRTKKCKCFICGEPGHFARECT-KQRGNIVRATVHQELAIPDNF
094 YP_009345075.1  ------KPPVKCKCYICGREGHFARDCN-NNNGNIARAAVIENLELPPDW
095 NP_068729.1     --------------------------------------------------
096 NP_047255.1     KEAILSIPVPKNSRQVREFLGTAGYCRLWIPGFAELAAPLYPLTRPG---
097 YP_006273075.1  VR------NKRCKCFLCGQEGHFARKCP-NDKRSSKRMAMFENLEIPEDC
098 YP_009508888.1  KEKLLNISPPQNLKQLQSILGLMNFARNFIPNYAERVKPFYSLISTAKS-
099 NP_395469.1     --------------------------------------------------
100 YP_009508546.1  KTKLLNVTPPKDLKQLQSILGLLNFARNFIPNFAELVQPLYNLIPSARG-
101 NP_056762.1     NHLANRCPRRYTNQARASLIDGLDEDIVSIASDDEDIENFLEIIELDEFI
102 YP_009508556.1  KLKLLDITPPKDLKQLQSILGLLNFARNFIPNYSELVKPLYTIVANANG-
103 NP_569153.1     R-------NKKCKCYLCGEEGHFARECP-NDRRNVKRVAMFEGLDLPDDC
104 NP_043924.1     SQLIRTLDIPLTVQGLRSALGLFNFCRAWIPEFSRKTQSLYDMLKGDCKS
105 YP_009243641.1  TIRRDN---LKTLNDFQKLLGDINWLRPSLGIPTYQLHNLFDTLKGDTDL
106 YP_004300274.1  --------------------------------------------------
107 NP_056848.1     --------------------------------------------------
108 YP_006495799.1  NAPG-----RKCKCYLCGIEGHYARECPKKTVQ-PQRMAYFEGLALADNW
109 YP_009508409.1  --------------------------------------------------
110 NP_687035.1     LPPLEG---EWTVNKIQKVVGVLNWAS--QIYPGIKTKHTCAMLRGKKNL
111 NP_045937.2     KVTVSQFQQPTTIRQIRAFLGLVGYCRHWIPEFSIHSKFLEKQLKKDTAE
112 NP_612577.1     --------------------------------------------------
113 NP_619548.1     --------------------------------------------------
114 NP_056880.1     QIRTDK---LRTLNDFQKLLGNINWIRPFLKLTTGELKPLFEILNGDSNP
115 YP_009121747.1  ED---RGRVSKCKCYICGREGHFARDCK-SKTGNMARRAVIDNLDLKEDW
116 YP_009113237.2  Q-------NRKCKCYACGEEGHFSKDCP-NDKKSSKRVALLQNLEIPEDY
117 YP_009508408.1  MR------EKRCKCFLCGNEGHFARECP-NDKRNIKRVAVFEGLEVPDDC
118 NP_955579.1     KEAILSIPVPKNSRQVREFLGTAGYCRLWIPGFAELAAPLYPLTRPG---
119 YP_009229919.1  -H---ADKVRKCKCFICGEEGHFARECK-RGKGNIARAAVLKDLDIPDDY
120 YP_004442833.1  QGR-----KKECRCYVCGEEGHFAKECK-SKKKIQERVNIVNELDIPEGY
121 NP_862833.2     KLQTDH---LKTLNDFQKLLGDINWIRPYLKLPTYILQPLFDILKGNSDP
122 NP_040563.1     LKIKN----PLTVNELQQLVGNCVWVQ--PEVK-IPLYPLTDLLRDKTNL
123 YP_443922.1     RNSILHHPRPTTVSGLLSFLGLTGYSRTHIPDYTCRTEPLREIVRQAGAR
124 NP_057933.2     KETVMGQPTPKTPRQLREFLGTAGFCRLWIPGFAEMAAPLYPLTKTG---
125 YP_009116631.1  HTQG-----KKCKCYLCGIEGHYARECPKKVVK-PQRAAYFNGMGLDDNW
126 YP_009513249.1  KEKLENITAPTTLKQLQSILGLLNFARNFIPDFTELIAPLYALIPKSTK-
127 YP_223871.1     TQAILQIPVPKTKRQVREFLGTIGYCRLWIPGFAELAQPLYAATRGGN--
```

```
128 NP_127504.1        EDTDSFSVVSEESTQLSQLSSSSNDSPENNENTLPQTFMVRPTEPEISEV
129 NP_044929.1        KQKLMDLQPPTTLRQLQSILGLINFARNFLPNFAELVAPLYQLIPKAKG-
130 NP_057860.1        KVPIRS---RWALPELQALLGEIQWVSKGTPTLRQPLHSLYCALQRHTDP
131 NP_040973.1        L-DIPE---QPTLNELQKLAGKINWAS--QAIPDLSIKALTNMMRGNQNL
132 NP_777317.1        ALR-----KSECKCYACGELGHFASDCN-NPRKLTQRVAILDSLELEKGI
133 NP_054716.1        KSKLLDITPPNTLKQLQSILGLLNFARNFIPNYSELITPLYQLIPLAKG-
134 YP_001856242.1     LPEKD----SWTVNDIQKLVGKLNWAS--QIYPGIKVRQLCKLLRGTKAL
135 NP_954565.2        VIRKDK---LQTLNDFQKLLGDINWLRPYLKLTTGDLKPLFDTLKGDSDP
136 NP_569150.1        SLR-----KKNCRCYACGEEGHFASECK-NPRKIMDRVKVLDSLDLEDGL
137 NP_659397.1        --------------------------------------------------
138 YP_009513211.1     KATVMKIPTPTTPRQVREFLGTAGFCRLWIPGFASLAAPLYPLTREK---
139 YP_233107.1        NLR-----KKDCKCFACGETGHYASECT-NPKKFTHRVAILESLDLQEGL
140 YP_009109692.1     IETVARIPPPRNPREVREFLGTAGFCRLWIPGFAELAAPLYALTKES---
141 NP_057849.4        LPEKD----SWTVNDIQKLVGKLNWAS--QIYPGIKVRQLCKLLRGTKAL
142 NP_049560.1        TVPIRS---RWALPELQALLGEIQWVSKGTPTLRQPLHSLYCALQRHTDP
143 YP_009506267.1     TR------EKRCKCFLCGKEGHYARECP-NDRRDIKRVAIFEGLSIPDDF
144 NP_039820.1        --------QRKCKCYICGQEGHYANQCR-NKHKDQQRVAILQSLDLKENE
145 NP_056790.1        KATVMKIPVPTTPRQVREFLGTAGFCRLWIPGFASLAAPLYPLTKES---
146 YP_002321513.1     SKK---GVIRKCKCYICGEDGHFARECR-SKMVNVQRAAIFNELELDDSL
147 NP_955591.1        KETVMGQPTPKTPRQLREFLGTAGFCRLWIPGFAEMAAPLYPLTKTG---
148 YP_001956722.2     KQKLLNITPPRDLKQLQSILGLLNFARNFIPNFSELVKPLYNIIATANG-
149 NP_056902.1        LDMVKN----PTLNDVQKLMGNITWMS--SGIPGLTVKHIAATTKGCLEL
150 NP_056886.1        LVAEPR---IATLWDVQKLVGSLQWLRPALGIPPRLMGPFYEQLRG-SDP
151 NP_041261.1        CLKTDH---LKTLNDFQKLLGDIQWLRPYLKLPTSALVPLNNILKGDPNP
```

```
001 YP_004442836.1     DVVSVGFDEDDVSDIYSIDEEAENNKFTNED-------------------
002 NP_056728.1        --------------------------------------------------
003 YP_009508537.1     ---NYVPWQTEHST------------------------------------
004 YP_004732983.2     DVISVDPGDQDGSDICSISEGEAQHGMEDLAAFKAQLPYPV---------
005 YP_009508407.1     --------------------------------------------------
006 NP_040939.1        QS--ERKIEEVHVK------------------------------------
007 YP_003864102.1     NS--PRSLSDEALI------------------------------------
008 YP_009002585.1     EVVSVEEGEEVSDAIFSISENEDEDALHTGGAFAETIF------------
009 YP_003284237.1     EIVSVGEDEDEVSDIWSLSEGEDGH-----G-------------------
010 YP_009268869.1     QS--ERRIELRHVK------------------------------------
011 YP_009506251.1     EIVSVEEGEEDSDAIYSLSENEE---ELQDEVLHEKVF------------
012 NP_955611.1        NE--AREWNLDMKM------------------------------------
013 YP_008567619.1     DVVSVSEGEDDSDAIYSVSEGE-------DGSIATMT-------------
014 NP_149413.1        DIVSVGFDEEDTEDIYSVSENEENQAHLGVSYEDPQLPTIP---------
015 YP_004442824.1     DVISVGMDEDDISEIYSVGEEIDEYQFQQEE-------------------
016 YP_001931967.1     --------------------------------------------------
017 YP_002455786.1     RD--TITLRHPHLH------------------------------------
018 YP_009109694.1     ---TPFTWQTEHQL------------------------------------
019 NP_789739.1        TE--VIPLTEEAEL------------------------------------
020 NP_955619.1        RD--QIYLNPSQVQ------------------------------------
021 YP_224289.1        EIVSVEENETELSDIYSISEGEESGPPKSEI-------------------
022 YP_009182100.1     SKSPSDFATCLYKQFCREQDPSKKEKDLEILKGKLVNLTISDLKYFLSYV
023 NP_705927.1        TE--VIPLTEEAEL------------------------------------
024 YP_009508577.1     ---NNIKWTSEHTR------------------------------------
025 YP_004442830.1     DIVSVGYDEKSVEDIFSISEGEDQQAHILKE-EDEQLPTTP---------
026 NP_758887.1        DE--KVEWTREAEL------------------------------------
027 YP_009508406.1     --------------------------------------------------
028 YP_610965.1        DVLSVDNNDPMSDAICSFSEGEAG-GSQTIKTALSDLP------------
029 YP_001931961.1     --------------------------------------------------
030 YP_241114.1        AS--PRTLSLEGQT------------------------------------
031 YP_006732334.1     --------------------------------------------------
032 YP_009041481.1     DIMSVDPEEPMSDAICSMSEGE-AGFTATAYNFEDDLPYG----------
033 YP_001036293.1     EIVSVQEGDELSDSIFSVSEGE---EVLQSLNPFIQQ-------------
034 YP_605811.1        DVISVGFDESDVSDIYSVSEGEDNYQFNNED-------------------
035 NP_955577.1        ---TLFQWGTEQQL------------------------------------
036 YP_003208050.1     DVLSLDLNESDSDAICSFSEGEIANGARYVQEELHKLP------------
037 NP_789740.1        TE--VIPLTEEAEL------------------------------------
038 YP_004442839.1     DVISVGFDESDVSDIYSVSEGEDNYQFKHED-------------------
039 YP_009507791.1     DS--YRQWTPEALGHLQKIKAGEPKAVTLRQGPKESYKDFIDR-------
040 YP_004581513.1     --------------------------------------------------
041 NP_043933.1        --------------------------------------------------
042 YP_009507248.1     TLSPEYVN------------------------------------------
043 YP_002117531.1     DVVSVGLDEGDQSDIYSVSEGEEGHS----D-------------------
044 YP_004347415.1     --------------------------------------------------
045 NP_056891.1        NS--HRSLSKEALA------------------------------------
046 YP_009513242.1     ---KFISWTEENSN------------------------------------
```

```
047 YP_009352866.1   DIMSVDPDEPMSDAICSMSEGE-AGFSATAFQFDDDLPYG----------
048 YP_009165750.1   --------------------------------------------------
049 NP_040840.1      QS--ERYIESIHVR------------------------------------
050 YP_004222728.1   NE--AREWNLDMKM------------------------------------
051 YP_009506264.1   EVVSVGENEDDLSDVYSLSEGETG------G-------------------
052 YP_567050.1      RA--HLLLQKQQLH------------------------------------
053 YP_009508571.1   ---KYIEWSNENTV------------------------------------
054 NP_056803.1      ---KYIEWTEDNTK------------------------------------
055 NP_955564.1      IS--TRKLTPEACK------------------------------------
056 YP_009508411.1   EIVSVKEGEAQSDAIYSISEGEDDNEVHN-LNSITHS-------------
057 YP_009140788.1   DIVSVQEGEELSDAIYSISEGEDN---------DEELQ------------
058 YP_009508410.1   --------------------------------------------------
059 NP_040333.1      ---TLFEWGPDQQK------------------------------------
060 YP_008992013.1   EEEMPESSATNQPPP-AQTTSSSQKSSNGPWFTYDDIPSYKWR-------
061 NP_041734.1      DIVSVEEGEEESDAIYSISENEDG--ELDTEVVHEKVF------------
062 NP_056895.1      RA--IIQLSPEQLQ------------------------------------
063 YP_009109689.1   ---TPFTWQTEHQL------------------------------------
064 YP_002519387.1   --------------------------------------------------
065 YP_595725.1      DLVSVGYDESDIDEIYSVSENEECQAHLGLN-EDEQLPKVP---------
066 YP_009508551.1   ---KYIEWSEENTK------------------------------------
067 YP_001497148.1   ---APFTWQEKHQS------------------------------------
068 NP_041186.1      AS--PRTLSLEGRT------------------------------------
069 NP_663784.1      TE--EVQWTELAEA------------------------------------
070 YP_009508443.1   EIVSVHEGDDLSDAIYSVSEGENNEEIDQGVNVFINR-------------
071 YP_003987465.1   DVVSVGENEDELSDIYSVSEGEES--------------------------
072 NP_861410.1      --------------------------------------------------
073 YP_009345071.1   ELVSAEENDS-GDEIYSMSEGEDPTTMFMNLTPLAQIEDCLTLDWEYKDV
074 YP_009508582.1   ---NNIKWTSEHTR------------------------------------
075 NP_569141.1      --------------------------------------------------
076 YP_002916057.1   --------------------------------------------------
077 NP_777384.2      RA--IIQLSPEQLQ------------------------------------
078 YP_233110.1      DVISVGFDESDVSDIYSVSEGEDSYQFNNED-------------------
079 NP_542258.1      RD--TIELTSTQVQ------------------------------------
080 YP_009130664.1   DIVSVQEGDNESDAIYNISEGEEMEDLTKSLNVMSVS-------------
081 YP_009508566.1   ---QSISWEPKHSQ------------------------------------
082 YP_006607892.1   --------------------------------------------------
083 YP_009506270.1   DVVSVGFDEDDVSDIYSIDEEADNYKFTNEE-------------------
084 NP_042513.1      --------------------------------------------------
085 YP_006907834.1   --------------------------------------------------
086 YP_004442827.1   DIISVGYDEKSIEDIYSISEGEDNQAHLGRD-EEQ-LPEVP---------
087 YP_009408594.1   DIVSVDLNEPDSDAICSVSEGETS-TEQQCQAATLEFP------------
088 YP_009508561.1   ---NFIQWTAEHTL------------------------------------
089 NP_056907.1      RA--HLLLQKQQLH------------------------------------
090 YP_002308474.1   RD--TIQLTPPQLQ------------------------------------
091 YP_007761644.1   --------------------------------------------------
092 NP_040550.1      NE--AREWNLDMKM------------------------------------
093 NP_758808.1      DVVSVDADESDSSGIYSYSENEAPLQEVNSFIHDENIF------------
094 YP_009345075.1   DVVSVDINDPDSDAICSVSDGEPEGQEEVLNGISPDP-------------
095 NP_068729.1      --------------------------------------------------
096 NP_047255.1      ---TLFQWGTEQQL------------------------------------
097 YP_006273075.1   DIVSVEEGEDMSDAIYSISEGEDG--------ADELQ-------------
098 YP_009508888.1   ---NNILWNDELTS------------------------------------
099 NP_395469.1      --------------------------------------------------
100 YP_009508546.1   ---KYIEWSKENTK------------------------------------
101 NP_056762.1      AHSSQEHEHTWEIGG-----------------------------------
102 YP_009508556.1   ---KFISWTEENSN------------------------------------
103 NP_569153.1      EIVSIDEGDPDSDAIFSISEGEE------AGTLEEQCF------------
104 NP_043924.1      TDKLKWTEDNLNKFKLLKDEVASACVLGLPDPTLPFRHLIG---------
105 YP_009243641.1   NS--ARCLTPAAEE------------------------------------
106 YP_004300274.1   --------------------------------------------------
107 NP_056848.1      --------------------------------------------------
108 YP_006495799.1   DVLSVDVGESDSEGVCSISEGETAGRMDELAAFKTQLPYPV---------
109 YP_009508409.1   --------------------------------------------------
110 NP_687035.1      LE--EIVWTEEAEA------------------------------------
111 NP_045937.2      PFQLDDQQVEAFNK------------------------------------
112 NP_612577.1      --------------------------------------------------
113 NP_619548.1      --------------------------------------------------
114 NP_056880.1      IS--TRKLTPEACK------------------------------------
115 YP_009121747.1   DIVSADMSDHDT--VYSISEGEGGATEMNIGAMVEESP------------
116 YP_009113237.2   VAVSVDEGDPQSDAIYSISDNEDE--YQNELLQEESVF------------
117 YP_009508408.1   EIVSVQEGDNQSDAIYSVSEGEVADDVQQGVNVFIQK-------------
118 NP_955579.1      ---TLFQWGTEQQL------------------------------------
119 YP_009229919.1   DVVSVDVGDPDSDGICSFSEGEIK--NYATKAVLEDDF------------
```

```
120 YP_004442833.1    DIVSVGYDEKSLSDIFSISEGEDNQAHL-RD-EEG-LPEVP---------
121 NP_862833.2       AS--PRILSPEGRS------------------------------------
122 NP_040563.1       QE--KIQLTPEAIK------------------------------------
123 YP_443922.1       NLHASLIWTPEATC------------------------------------
124 NP_057933.2       ---TLFNWGPDQQK------------------------------------
125 YP_009116631.1    DVVSVEPGEEDDDEICSISEGENTGGMHELMAFKTQLPYPV---------
126 YP_009513249.1    ---NYVPWQIEHST------------------------------------
127 YP_223871.1       ---DPLVWGEKEEE------------------------------------
128 NP_127504.1       EDEVDGMTEEPIPERRPEITPPKMVGTGFHTFSLDDISITKWP-------
129 NP_044929.1       ---QCIPWTMDHTT------------------------------------
130 NP_057860.1       RD--QIYLNPSQVQ------------------------------------
131 NP_040973.1       NS--TRQWTKEAR-------------------------------------
132 NP_777317.1       DVVSVGFDEDDVSDIYSVGEEIDEYQFTHEN-------------------
133 NP_054716.1       ---IYIPWETKHTA------------------------------------
134 YP_001856242.1    TE--VIPLTEEAEL------------------------------------
135 NP_954565.2       NS--HRSLSKEALA------------------------------------
136 NP_569150.1       DVISVGFDEDDVSDIYSIDEEADNYRFTNEE-------------------
137 NP_659397.1       --------------------------------------------------
138 YP_009513211.1    ---VPFTWTEAHQE------------------------------------
139 YP_233107.1       EVVSVGMDESDVSDIYSISENEDNHQITEE--------------------
140 YP_009109692.1    ---TPFTWQTEHQL------------------------------------
141 NP_057849.4       TE--VIPLTEEAEL------------------------------------
142 NP_049560.1       RD--QICLNPSQVQ------------------------------------
143 YP_009506267.1    DIVSVHEGEEQSDAIYSVSEGE-NESIDQEVNTFISYGEGG---------
144 NP_039820.1       EVVSADDKEEEDDEIFSVLGEEDYQEETIMVLEED---------------
145 NP_056790.1       ---IPFIWTEEHQQ------------------------------------
146 YP_002321513.1    DVVSVDMGEPDSDHIVSMSDGVGPEATRKYLDSQFTLP------------
147 NP_955591.1       ---TLFNWGPDQQK------------------------------------
148 YP_001956722.2    ---KYITWTTDNSQ------------------------------------
149 NP_056902.1       NQ--KVIWTEEAQK------------------------------------
150 NP_056886.1       NE--AREWNLDMKM------------------------------------
151 NP_041261.1       LS--VRALTPEAKQ------------------------------------


001 YP_004442836.1    --------------------------------------------------
002 NP_056728.1       --------------------------------------------------
003 YP_009508537.1    --------------------------------------------------
004 YP_004732983.2    --------------------------------------------------
005 YP_009508407.1    --------------------------------------------------
006 NP_040939.1       --------------------------------------------------
007 YP_003864102.1    --------------------------------------------------
008 YP_009002585.1    --------------------------------------------------
009 YP_003284237.1    --------------------------------------------------
010 YP_009268869.1    --------------------------------------------------
011 YP_009506251.1    --------------------------------------------------
012 NP_955611.1       --------------------------------------------------
013 YP_008567619.1    --------------------------------------------------
014 NP_149413.1       --------------------------------------------------
015 YP_004442824.1    --------------------------------------------------
016 YP_001931967.1    --------------------------------------------------
017 YP_002455786.1    --------------------------------------------------
018 YP_009109694.1    --------------------------------------------------
019 NP_789739.1       --------------------------------------------------
020 NP_955619.1       --------------------------------------------------
021 YP_224289.1       --------------------------------------------------
022 YP_009182100.1    NEYNLAYYNLGIFNNNTYRDMFIQKLPDPWKEYLTNERAKEAGNEEWTLG
023 NP_705927.1       --------------------------------------------------
024 YP_009508577.1    --------------------------------------------------
025 YP_004442830.1    --------------------------------------------------
026 NP_758887.1       --------------------------------------------------
027 YP_009508406.1    --------------------------------------------------
028 YP_610965.1       --------------------------------------------------
029 YP_001931961.1    --------------------------------------------------
030 YP_241114.1       --------------------------------------------------
031 YP_006732334.1    --------------------------------------------------
032 YP_009041481.1    --------------------------------------------------
033 YP_001036293.1    --------------------------------------------------
034 YP_605811.1       --------------------------------------------------
035 NP_955577.1       --------------------------------------------------
036 YP_003208050.1    --------------------------------------------------
037 NP_789740.1       --------------------------------------------------
038 YP_004442839.1    --------------------------------------------------
```

```
039 YP_009507791.1    --------------------------------------------------
040 YP_004581513.1    --------------------------------------------------
041 NP_043933.1       --------------------------------------------------
042 YP_009507248.1    --------------------------------------------------
043 YP_002117531.1    --------------------------------------------------
044 YP_004347415.1    --------------------------------------------------
045 NP_056891.1       --------------------------------------------------
046 YP_009513242.1    --------------------------------------------------
047 YP_009352866.1    --------------------------------------------------
048 YP_009165750.1    --------------------------------------------------
049 NP_040840.1       --------------------------------------------------
050 YP_004222728.1    --------------------------------------------------
051 YP_009506264.1    --------------------------------------------------
052 YP_567050.1       --------------------------------------------------
053 YP_009508571.1    --------------------------------------------------
054 NP_056803.1       --------------------------------------------------
055 NP_955564.1       --------------------------------------------------
056 YP_009508411.1    --------------------------------------------------
057 YP_009140788.1    --------------------------------------------------
058 YP_009508410.1    --------------------------------------------------
059 NP_040333.1       --------------------------------------------------
060 YP_008992013.1    --------------------------------------------------
061 NP_041734.1       --------------------------------------------------
062 NP_056895.1       --------------------------------------------------
063 YP_009109689.1    --------------------------------------------------
064 YP_002519387.1    --------------------------------------------------
065 YP_595725.1       --------------------------------------------------
066 YP_009508551.1    --------------------------------------------------
067 YP_001497148.1    --------------------------------------------------
068 NP_041186.1       --------------------------------------------------
069 NP_663784.1       --------------------------------------------------
070 YP_009508443.1    --------------------------------------------------
071 YP_003987465.1    --------------------------------------------------
072 NP_861410.1       --------------------------------------------------
073 YP_009345071.1    YEYRHFGTCTSDIQQPQPWKQQTKVYEEGGNGHDRAFYRYYGPPKGRENW
074 YP_009508582.1    --------------------------------------------------
075 NP_569141.1       --------------------------------------------------
076 YP_002916057.1    --------------------------------------------------
077 NP_777384.2       --------------------------------------------------
078 YP_233110.1       --------------------------------------------------
079 NP_542258.1       --------------------------------------------------
080 YP_009130664.1    --------------------------------------------------
081 YP_009508566.1    --------------------------------------------------
082 YP_006607892.1    --------------------------------------------------
083 YP_009506270.1    --------------------------------------------------
084 NP_042513.1       --------------------------------------------------
085 YP_006907834.1    --------------------------------------------------
086 YP_004442827.1    --------------------------------------------------
087 YP_009408594.1    --------------------------------------------------
088 YP_009508561.1    --------------------------------------------------
089 NP_056907.1       --------------------------------------------------
090 YP_002308474.1    --------------------------------------------------
091 YP_007761644.1    --------------------------------------------------
092 NP_040550.1       --------------------------------------------------
093 NP_758808.1       --------------------------------------------------
094 YP_009345075.1    --------------------------------------------------
095 NP_068729.1       --------------------------------------------------
096 NP_047255.1       --------------------------------------------------
097 YP_006273075.1    --------------------------------------------------
098 YP_009508888.1    --------------------------------------------------
099 NP_395469.1       --------------------------------------------------
100 YP_009508546.1    --------------------------------------------------
101 NP_056762.1       --------------------------------------------------
102 YP_009508556.1    --------------------------------------------------
103 NP_569153.1       --------------------------------------------------
104 NP_043924.1       --------------------------------------------------
105 YP_009243641.1    --------------------------------------------------
106 YP_004300274.1    --------------------------------------------------
107 NP_056848.1       --------------------------------------------------
108 YP_006495799.1    --------------------------------------------------
109 YP_009508409.1    --------------------------------------------------
110 NP_687035.1       --------------------------------------------------
111 NP_045937.2       --------------------------------------------------
```

```
112 NP_612577.1        --------------------------------------------------
113 NP_619548.1        --------------------------------------------------
114 NP_056880.1        --------------------------------------------------
115 YP_009121747.1     --------------------------------------------------
116 YP_009113237.2     --------------------------------------------------
117 YP_009508408.1     --------------------------------------------------
118 NP_955579.1        --------------------------------------------------
119 YP_009229919.1     --------------------------------------------------
120 YP_004442833.1     --------------------------------------------------
121 NP_862833.2        --------------------------------------------------
122 NP_040563.1        --------------------------------------------------
123 YP_443922.1        --------------------------------------------------
124 NP_057933.2        --------------------------------------------------
125 YP_009116631.1     --------------------------------------------------
126 YP_009513249.1     --------------------------------------------------
127 YP_223871.1        --------------------------------------------------
128 NP_127504.1        --------------------------------------------------
129 NP_044929.1        --------------------------------------------------
130 NP_057860.1        --------------------------------------------------
131 NP_040973.1        --------------------------------------------------
132 NP_777317.1        --------------------------------------------------
133 NP_054716.1        --------------------------------------------------
134 YP_001856242.1     --------------------------------------------------
135 NP_954565.2        --------------------------------------------------
136 NP_569150.1        --------------------------------------------------
137 NP_659397.1        --------------------------------------------------
138 YP_009513211.1     --------------------------------------------------
139 YP_233107.1        --------------------------------------------------
140 YP_009109692.1     --------------------------------------------------
141 NP_057849.4        --------------------------------------------------
142 NP_049560.1        --------------------------------------------------
143 YP_009506267.1     --------------------------------------------------
144 NP_039820.1        --------------------------------------------------
145 NP_056790.1        --------------------------------------------------
146 YP_002321513.1     --------------------------------------------------
147 NP_955591.1        --------------------------------------------------
148 YP_001956722.2     --------------------------------------------------
149 NP_056902.1        --------------------------------------------------
150 NP_056886.1        --------------------------------------------------
151 NP_041261.1        --------------------------------------------------
```

```
001 YP_004442836.1     -------------------------------MEDFKNYEVYMMEAEELDE
002 NP_056728.1        --------------------------------------------------
003 YP_009508537.1     --------------------------------------------------
004 YP_004732983.2     --------------------------EAQYEQHQALVVIQTTFKKEDKPQ
005 YP_009508407.1     --------------------------------------------------
006 NP_040939.1        --------------------------------------------------
007 YP_003864102.1     --------------------------------------------------
008 YP_009002585.1     -------------------------------MFHEGTDNVYWAGKKDG--
009 YP_003284237.1     -------------------------------EEEVAAMVFALEDEEED--
010 YP_009268869.1     --------------------------------------------------
011 YP_009506251.1     -------------------------------MIREG-DQSYWMGKANH--
012 NP_955611.1        --------------------------------------------------
013 YP_008567619.1     --------------------------HYINVFFEED--RTYWVGKAGS--
014 NP_149413.1        --------------------------TSFEEWEEYYKEEFTLMAEVDDS-
015 YP_004442824.1     -------------------------------FEDFTNYEVYMIHVEEI--
016 YP_001931967.1     --------------------------------------------------
017 YP_002455786.1     --------------------------------------------------
018 YP_009109694.1     --------------------------------------------------
019 NP_789739.1        --------------------------------------------------
020 NP_955619.1        --------------------------------------------------
021 YP_224289.1        -------------------------------MSLELFMLHEIEDEPEA--
022 YP_009182100.1     NLIEFVKTKLADLAIQAWRAKRMQKLYSKSGLSFCQHLIDIPTQWGGRNK
023 NP_705927.1        --------------------------------------------------
024 YP_009508577.1     --------------------------------------------------
025 YP_004442830.1     --------------------------LSWEDWENYYKKEFLYMAEIEEVS
026 NP_758887.1        --------------------------------------------------
027 YP_009508406.1     --------------------------------------------------
028 YP_610965.1        -------------------------FLEGSFMLKETV-------------
029 YP_001931961.1     --------------------------------------------------
030 YP_241114.1        --------------------------------------------------
```

```
031 YP_006732334.1    --------------------------------------------------
032 YP_009041481.1    --------------------------EPTNSFNFGLVLIQGPVPDVNAS-
033 YP_001036293.1    ---------------------------NLLYIREED--GKHFLGGGG---
034 YP_605811.1       -------------------------------FDVIG-HDVFMFTIEEQ--
035 NP_955577.1       --------------------------------------------------
036 YP_003208050.1    -------------------------YET-ALMLQSQN-------------
037 NP_789740.1       --------------------------------------------------
038 YP_004442839.1    -------------------------------FGVVD-HDVFMITIEE---
039 YP_009507791.1    --------------------------------------------------
040 YP_004581513.1    --------------------------------------------------
041 NP_043933.1       --------------------------------------------------
042 YP_009507248.1    --------------------------------------------------
043 YP_002117531.1    -------------------------------LHDETIFSLTEEDDEAA--
044 YP_004347415.1    --------------------------------------------------
045 NP_056891.1       --------------------------------------------------
046 YP_009513242.1    --------------------------------------------------
047 YP_009352866.1    --------------------------EGAG-INLGFVMIQGPIPNANEN-
048 YP_009165750.1    --------------------------------------------------
049 NP_040840.1       --------------------------------------------------
050 YP_004222728.1    --------------------------------------------------
051 YP_009506264.1    -------------------------------IEEIYALEDEMEPEQ----
052 YP_567050.1       --------------------------------------------------
053 YP_009508571.1    --------------------------------------------------
054 NP_056803.1       --------------------------------------------------
055 NP_955564.1       --------------------------------------------------
056 YP_009508411.1    ---------------------------SLFYLRAED--NTYWLGGGG---
057 YP_009140788.1    -------------------------------RSLQTMEIVLMLGEVDGG-
058 YP_009508410.1    --------------------------------------------------
059 NP_040333.1       --------------------------------------------------
060 YP_008992013.1    --------------------------DRLNEMSAWIDLQMLRSGATTES-
061 NP_041734.1       -------------------------------MMREE-DQSYWLGKTNH--
062 NP_056895.1       --------------------------------------------------
063 YP_009109689.1    --------------------------------------------------
064 YP_002519387.1    --------------------------------------------------
065 YP_595725.1       --------------------------QTFEEWEEYYKDEFIMMADIEESE
066 YP_009508551.1    --------------------------------------------------
067 YP_001497148.1    --------------------------------------------------
068 NP_041186.1       --------------------------------------------------
069 NP_663784.1       --------------------------------------------------
070 YP_009508443.1    --------------------------ETLYYLREED--HTYFLGGGG---
071 YP_003987465.1    --------------------------------THEAYHVFSLVEDC----
072 NP_861410.1       --------------------------------------------------
073 YP_009345071.1    PETPIPTGWGSDFDHDPDKEIDDIPDDEEIYMMQDDQFDEHWIGKRGG--
074 YP_009508582.1    --------------------------------------------------
075 NP_569141.1       --------------------------------------------------
076 YP_002916057.1    --------------------------------------------------
077 NP_777384.2       --------------------------------------------------
078 YP_233110.1       -------------------------------FDVIG-HDVFMFTIEEQ--
079 NP_542258.1       --------------------------------------------------
080 YP_009130664.1    --------------------------EHLYYFREED--QTYWIGGEG---
081 YP_009508566.1    --------------------------------------------------
082 YP_006607892.1    --------------------------------------------------
083 YP_009506270.1    -------------------------------MEDFKNYEVYMMNLEEIEE
084 NP_042513.1       --------------------------------------------------
085 YP_006907834.1    --------------------------------------------------
086 YP_004442827.1    --------------------------TTWEDWDSYYKKEFLMMAEVLEVS
087 YP_009408594.1    -------------------------GIEGLFMLGAPE-------------
088 YP_009508561.1    --------------------------------------------------
089 NP_056907.1       --------------------------------------------------
090 YP_002308474.1    --------------------------------------------------
091 YP_007761644.1    --------------------------------------------------
092 NP_040550.1       --------------------------------------------------
093 NP_758808.1       -------------------------FLSDADEFESPQQHLHETVNMLQSR
094 YP_009345075.1    ---------------------------ESIFMLREELEHKPP--------
095 NP_068729.1       --------------------------------------------------
096 NP_047255.1       --------------------------------------------------
097 YP_006273075.1    -------------------------------KSLQTLEIVLMLGEIDGG-
098 YP_009508888.1    --------------------------------------------------
099 NP_395469.1       --------------------------------------------------
100 YP_009508546.1    --------------------------------------------------
101 NP_056762.1       --------------------------------------------------
102 YP_009508556.1    --------------------------------------------------
103 NP_569153.1       -------------------------------VFQEECNGTYWLGKRGG--
```

104 NP_043924.1 --------------------------------------------------
105 YP_009243641.1 --------------------------------------------------
106 YP_004300274.1 --------------------------------------------------
107 NP_056848.1 --------------------------------------------------
108 YP_006495799.1 --------------------------EFEQG-MFVLTIEHVTAPAVASG-
109 YP_009508409.1 --------------------------------------------------
110 NP_687035.1 --------------------------------------------------
111 NP_045937.2 --------------------------------------------------
112 NP_612577.1 --------------------------------------------------
113 NP_619548.1 --------------------------------------------------
114 NP_056880.1 --------------------------------------------------
115 YP_009121747.1 -------------------------LEEYCFMFHQSHEEIHKRVQEAVS-
116 YP_009113237.2 -------------------------------VIVEKTESRKYLLGKEGS-
117 YP_009508408.1 --------------------------ETLYYLRQED--NAYFLGGGG---
118 NP_955579.1 --------------------------------------------------
119 YP_009229919.1 -------------------------LDELAFVVTEKKEIDY---------
120 YP_004442833.1 --------------------------TTWDQWDEYYKKEFIYMADTKEVL
121 NP_862833.2 --------------------------------------------------
122 NP_040563.1 --------------------------------------------------
123 YP_443922.1 --------------------------------------------------
124 NP_057933.2 --------------------------------------------------
125 YP_009116631.1 --------------------------EYEAS-TPQFLMPWTQVPVEKSDK
126 YP_009513249.1 --------------------------------------------------
127 YP_223871.1 --------------------------------------------------
128 NP_127504.1 --------------------------ERIQDFHTWMLTKQLVE-REPFL-
129 NP_044929.1 --------------------------------------------------
130 NP_057860.1 --------------------------------------------------
131 NP_040973.1 --------------------------------------------------
132 NP_777317.1 -------------------------------LEDFKNYEVNFLEDHEEDE
133 NP_054716.1 --------------------------------------------------
134 YP_001856242.1 --------------------------------------------------
135 NP_954565.2 --------------------------------------------------
136 NP_569150.1 -------------------------------MEGFKNYEVYMLRMEEMDE
137 NP_659397.1 --------------------------------------------------
138 YP_009513211.1 --------------------------------------------------
139 YP_233107.1 -------------------------------------WDVLMLQEEEIPE
140 YP_009109692.1 --------------------------------------------------
141 NP_057849.4 --------------------------------------------------
142 NP_049560.1 --------------------------------------------------
143 YP_009506267.1 -----------------TGSTTHFSNQTLYYLKEEGTVTHYYLGGGG---
144 NP_039820.1 --------------------------DIQQIIKEFSKFGDLSRRNVGPNF
145 NP_056790.1 --------------------------------------------------
146 YP_002321513.1 ------------------------WATEFIGVFHVTTDTLSMNENLG---
147 NP_955591.1 --------------------------------------------------
148 YP_001956722.2 --------------------------------------------------
149 NP_056902.1 --------------------------------------------------
150 NP_056886.1 --------------------------------------------------
151 NP_041261.1 --------------------------------------------------

001 YP_004442836.1 --------------------------------------------------
002 NP_056728.1 --------------------------------------------------
003 YP_009508537.1 --------------------------------------------------
004 YP_004732983.2 --------------------------------------------------
005 YP_009508407.1 --------------------------------------------------
006 NP_040939.1 --------------------------------------------------
007 YP_003864102.1 --------------------------------------------------
008 YP_009002585.1 --------------------------------------------------
009 YP_003284237.1 --------------------------------------------------
010 YP_009268869.1 --------------------------------------------------
011 YP_009506251.1 --------------------------------------------------
012 NP_955611.1 --------------------------------------------------
013 YP_008567619.1 --------------------------------------------------
014 NP_149413.1 -SDEEK--------------------------------------------
015 YP_004442824.1 --------------------------------------------------
016 YP_001931967.1 --------------------------------------------------
017 YP_002455786.1 --------------------------------------------------
018 YP_009109694.1 --------------------------------------------------
019 NP_789739.1 --------------------------------------------------
020 NP_955619.1 --------------------------------------------------
021 YP_224289.1 --------------------------------------------------
022 YP_009182100.1 ELKHKKKGKKIKSFKRYKFRKSKKKGFEKSKINPQKNFKRRFKQKSGKEI

```
023 NP_705927.1       --------------------------------------------------
024 YP_009508577.1    --------------------------------------------------
025 YP_004442830.1    SEEEKED-------------------------------------------
026 NP_758887.1       --------------------------------------------------
027 YP_009508406.1    --------------------------------------------------
028 YP_610965.1       --------------------------------------------------
029 YP_001931961.1    --------------------------------------------------
030 YP_241114.1       --------------------------------------------------
031 YP_006732334.1    --------------------------------------------------
032 YP_009041481.1    --------------------------------------------------
033 YP_001036293.1    --------------------------------------------------
034 YP_605811.1       --------------------------------------------------
035 NP_955577.1       --------------------------------------------------
036 YP_003208050.1    --------------------------------------------------
037 NP_789740.1       --------------------------------------------------
038 YP_004442839.1    --------------------------------------------------
039 YP_009507791.1    --------------------------------------------------
040 YP_004581513.1    --------------------------------------------------
041 NP_043933.1       --------------------------------------------------
042 YP_009507248.1    --------------------------------------------------
043 YP_002117531.1    --------------------------------------------------
044 YP_004347415.1    --------------------------------------------------
045 NP_056891.1       --------------------------------------------------
046 YP_009513242.1    --------------------------------------------------
047 YP_009352866.1    --------------------------------------------------
048 YP_009165750.1    --------------------------------------------------
049 NP_040840.1       --------------------------------------------------
050 YP_004222728.1    --------------------------------------------------
051 YP_009506264.1    --------------------------------------------------
052 YP_567050.1       --------------------------------------------------
053 YP_009508571.1    --------------------------------------------------
054 NP_056803.1       --------------------------------------------------
055 NP_955564.1       --------------------------------------------------
056 YP_009508411.1    --------------------------------------------------
057 YP_009140788.1    --------------------------------------------------
058 YP_009508410.1    --------------------------------------------------
059 NP_040333.1       --------------------------------------------------
060 YP_008992013.1    --------------------------------------------------
061 NP_041734.1       --------------------------------------------------
062 NP_056895.1       --------------------------------------------------
063 YP_009109689.1    --------------------------------------------------
064 YP_002519387.1    --------------------------------------------------
065 YP_595725.1       NSDEEK--------------------------------------------
066 YP_009508551.1    --------------------------------------------------
067 YP_001497148.1    --------------------------------------------------
068 NP_041186.1       --------------------------------------------------
069 NP_663784.1       --------------------------------------------------
070 YP_009508443.1    --------------------------------------------------
071 YP_003987465.1    --------------------------------------------------
072 NP_861410.1       --------------------------------------------------
073 YP_009345071.1    --------------------------------------------------
074 YP_009508582.1    --------------------------------------------------
075 NP_569141.1       --------------------------------------------------
076 YP_002916057.1    --------------------------------------------------
077 NP_777384.2       --------------------------------------------------
078 YP_233110.1       --------------------------------------------------
079 NP_542258.1       --------------------------------------------------
080 YP_009130664.1    --------------------------------------------------
081 YP_009508566.1    --------------------------------------------------
082 YP_006607892.1    --------------------------------------------------
083 YP_009506270.1    --------------------------------------------------
084 NP_042513.1       --------------------------------------------------
085 YP_006907834.1    --------------------------------------------------
086 YP_004442827.1    SDEEKE--------------------------------------------
087 YP_009408594.1    --------------------------------------------------
088 YP_009508561.1    --------------------------------------------------
089 NP_056907.1       --------------------------------------------------
090 YP_002308474.1    --------------------------------------------------
091 YP_007761644.1    --------------------------------------------------
092 NP_040550.1       --------------------------------------------------
093 NP_758808.1       --------------------------------------------------
094 YP_009345075.1    --------------------------------------------------
095 NP_068729.1       --------------------------------------------------
```

096 NP_047255.1 --------------------------------------------------
097 YP_006273075.1 --------------------------------------------------
098 YP_009508888.1 --------------------------------------------------
099 NP_395469.1 --------------------------------------------------
100 YP_009508546.1 --------------------------------------------------
101 NP_056762.1 --------------------------------------------------
102 YP_009508556.1 --------------------------------------------------
103 NP_569153.1 --------------------------------------------------
104 NP_043924.1 --------------------------------------------------
105 YP_009243641.1 --------------------------------------------------
106 YP_004300274.1 --------------------------------------------------
107 NP_056848.1 --------------------------------------------------
108 YP_006495799.1 --------------------------------------------------
109 YP_009508409.1 --------------------------------------------------
110 NP_687035.1 --------------------------------------------------
111 NP_045937.2 --------------------------------------------------
112 NP_612577.1 --------------------------------------------------
113 NP_619548.1 --------------------------------------------------
114 NP_056880.1 --------------------------------------------------
115 YP_009121747.1 --------------------------------------------------
116 YP_009113237.2 --------------------------------------------------
117 YP_009508408.1 --------------------------------------------------
118 NP_955579.1 --------------------------------------------------
119 YP_009229919.1 --------------------------------------------------
120 YP_004442833.1 SEEENE--------------------------------------------
121 NP_862833.2 --------------------------------------------------
122 NP_040563.1 --------------------------------------------------
123 YP_443922.1 --------------------------------------------------
124 NP_057933.2 --------------------------------------------------
125 YP_009116631.1 --------------------------------------------------
126 YP_009513249.1 --------------------------------------------------
127 YP_223871.1 --------------------------------------------------
128 NP_127504.1 --------------------------------------------------
129 NP_044929.1 --------------------------------------------------
130 NP_057860.1 --------------------------------------------------
131 NP_040973.1 --------------------------------------------------
132 NP_777317.1 RWDTLGEPSGKFDYYVKYSAPKHHQLMEEIIPSGWDTEEEEEEYPESVEE
133 NP_054716.1 --------------------------------------------------
134 YP_001856242.1 --------------------------------------------------
135 NP_954565.2 --------------------------------------------------
136 NP_569150.1 --------------------------------------------------
137 NP_659397.1 --------------------------------------------------
138 YP_009513211.1 --------------------------------------------------
139 YP_233107.1 --------------------------------------------------
140 YP_009109692.1 --------------------------------------------------
141 NP_057849.4 --------------------------------------------------
142 NP_049560.1 --------------------------------------------------
143 YP_009506267.1 --------------------------------------------------
144 NP_039820.1 PG------------------------------------------------
145 NP_056790.1 --------------------------------------------------
146 YP_002321513.1 --------------------------------------------------
147 NP_955591.1 --------------------------------------------------
148 YP_001956722.2 --------------------------------------------------
149 NP_056902.1 --------------------------------------------------
150 NP_056886.1 --------------------------------------------------
151 NP_041261.1 --------------------------------------------------

001 YP_004442836.1 ----------PKEYLVGEPSEWRSKMRVSRRQYYCK-HDWKFEE----SH
002 NP_056728.1 --------------------------------------------------
003 YP_009508537.1 -------------------------------------------------T
004 YP_004732983.2 -------------------GSWRMSKPIPETQQQCQ-HTWDDMYALAEGQ
005 YP_009508407.1 --------------------------------------------------
006 NP_040939.1 -------------------------------------------------E
007 YP_003864102.1 -------------------------------------------------A
008 YP_009002585.1 ---------------------YLPYKRITREQFQCE-HEWQMNQEVATVE
009 YP_003284237.1 -------------TLIGKPGTWRSQVRVTRKEYECI-HLWDFNS----EG
010 YP_009268869.1 -------------------------------------------------E
011 YP_009506251.1 ---------------------WTAMVRVSYQQYHCP-HQWEHNKEIPIMA
012 NP_955611.1 -------------------------------------------------A
013 YP_008567619.1 ---------------------WQPMIKVTQKVHDCQ-HCWMENTDVPNDG
014 NP_149413.1 -----------GPFLVGPKDGFRHQMIVSQKQYKCT-HDWEYQK----DR

```
015 YP_004442824.1 ----------PREFLVGEPSDWRSKMKVSINQYYCI-HNWKFDQ----ES
016 YP_001931967.1 --------------------------------------------------
017 YP_002455786.1 -------------------------------------------------A
018 YP_009109694.1 -------------------------------------------------A
019 NP_789739.1    -------------------------------------------------E
020 NP_955619.1    -------------------------------------------------S
021 YP_224289.1    -------------YLIGATDTWRNQMKVSRKEYYCQ-HEWDHTT----KV
022 YP_009182100.1 EDRKKQSKSKPTSSKGVANVCWNCNQQGHFSRECPRKNARLIELQEAADT
023 NP_705927.1    -------------------------------------------------E
024 YP_009508577.1 -------------------------------------------------Y
025 YP_004442830.1 ----------NGPWLVGKEGGFMYQMKVSRKQYHCQ-HEWFYNQ----PK
026 NP_758887.1    -------------------------------------------------E
027 YP_009508406.1 --------------------------------------------------
028 YP_610965.1    --------------------SWRPTVSLPANQANCQ-HLWEDNS-QSTGH
029 YP_001931961.1 --------------------------------------------------
030 YP_241114.1    -------------------------------------------------A
031 YP_006732334.1 --------------------------------------------------
032 YP_009041481.1 --------------------SWMPRHPLEGLQLTCI-HVWEENAPIQATE
033 YP_001036293.1 ---------------------WRAAVQVTPQEHDCH-HQWEYYI----LN
034 YP_605811.1    -----------RNCLVETTSAWRSAMKVTSEEKNCL-HTWSFEE----KT
035 NP_955577.1    -------------------------------------------------A
036 YP_003208050.1 --------------------SWRRTVKLPAYQDCCG-HQPTLGPNLTFEQ
037 NP_789740.1    -------------------------------------------------E
038 YP_004442839.1 -----------RNYLVPTTSEWRAAMKVTKEEYLCL-HQWEFSS----ER
039 YP_009507791.1 -------------------------------------LFQQIDQEQALEE
040 YP_004581513.1 --------------------------------------------------
041 NP_043933.1    --------------------------------------------------
042 YP_009507248.1 -------------------------------------------------A
043 YP_002117531.1 -------------LLVGKAGTWRPQMKVSKKEFSCE-HEWDYEV----SG
044 YP_004347415.1 --------------------------------------------------
045 NP_056891.1    -------------------------------------------------S
046 YP_009513242.1 -------------------------------------------------Q
047 YP_009352866.1 --------------------SWMVRYPLEGSQQSCD-HQWEEGAAIPQGE
048 YP_009165750.1 --------------------------------------------------
049 NP_040840.1    -------------------------------------------------E
050 YP_004222728.1 -------------------------------------------------A
051 YP_009506264.1 -------------LLIGKPNTWRTQIRVSRKEFHCL-HEWDFTK----TE
052 YP_567050.1    -------------------------------------------------A
053 YP_009508571.1 -------------------------------------------------Q
054 NP_056803.1    -------------------------------------------------Q
055 NP_955564.1    -------------------------------------------------A
056 YP_009508411.1 ---------------------WRFAVKVTPQEHNCA-HSWSYYE----EQ
057 YP_009140788.1 ---------------------YRPQIKLPDEQFNCA-HNWVHNGEILVQK
058 YP_009508410.1 --------------------------------------------------
059 NP_040333.1    -------------------------------------------------A
060 YP_008992013.1 ------------------------VLREFATRFTGALRDWFDSLGPYRQL
061 NP_041734.1    ---------------------WTAMVRVSSQQYHCM-HQWEHNKEILVVA
062 NP_056895.1    -------------------------------------------------G
063 YP_009109689.1 -------------------------------------------------A
064 YP_002519387.1 --------------------------------------------------
065 YP_595725.1    -----------GPFLVGPKGGFRHQMEVSYKQYKCE-HDWDFTR----TR
066 YP_009508551.1 -------------------------------------------------Q
067 YP_001497148.1 -------------------------------------------------A
068 NP_041186.1    -------------------------------------------------A
069 NP_663784.1    -------------------------------------------------E
070 YP_009508443.1 ---------------------WRCSVKVSQQEHDCK-HRWDYYT----NT
071 YP_003987465.1 --------------LIGKANTWRYQVRVSPKEYYCK-HEWGFNS----KI
072 NP_861410.1    --------------------------------------------------
073 YP_009345071.1 ---------------------WQPLIKLSDQEFNCV-HHWQHNIQVP-DC
074 YP_009508582.1 -------------------------------------------------H
075 NP_569141.1    --------------------------------------------------
076 YP_002916057.1 --------------------------------------------------
077 NP_777384.2    -------------------------------------------------G
078 YP_233110.1    -----------RNCLVETTSAWRSAMKVTPEEKNCL-HTWSFEE----KT
079 NP_542258.1    -------------------------------------------------A
080 YP_009130664.1 ---------------------YRSMIKATKTQHDCN-HVWEYN----GAG
081 YP_009508566.1 -------------------------------------------------A
082 YP_006607892.1 --------------------------------------------------
083 YP_009506270.1 ----------PNDYLVGEPSDWRSKMKVSRRQYFCK-HEWSFEE----AH
084 NP_042513.1    --------------------------------------------------
085 YP_006907834.1 --------------------------------------------------
086 YP_004442827.1 ----------EGPWLVGKEGGYMQQMYVTKKQFHCT-HDWYYDQ----PR
087 YP_009408594.1 --------------------GSRVQIKVSDEILNCV-HQWQMNEDVDVLY
```

```
088 YP_009508561.1   -------------------------------------------------R
089 NP_056907.1      -------------------------------------------------A
090 YP_002308474.1   -------------------------------------------------A
091 YP_007761644.1   --------------------------------------------------
092 NP_040550.1      -------------------------------------------------A
093 NP_758808.1      -------------------SAYLPQVAVGEEKLNCS-HIWLQDVDIPSDK
094 YP_009345075.1   --------------------SWFYKMPIPKEYQECL-HAWEANAPIVQGR
095 NP_068729.1      --------------------------------------------------
096 NP_047255.1      -------------------------------------------------A
097 YP_006273075.1   ---------------------YRPQIKLPEEQMNCV-HNWIHNGDIFVQR
098 YP_009508888.1   -------------------------------------------------Q
099 NP_395469.1      --------------------------------------------------
100 YP_009508546.1   -------------------------------------------------Q
101 NP_056762.1      ------------------------------------------------KK
102 YP_009508556.1   -------------------------------------------------Q
103 NP_569153.1      ---------------------YQDLVQISKEIYYCQ-HEWEENQPINDPA
104 NP_043924.1      ------------------------IRQGHFLCSLVQKDDNGMWHVLGFYS
105 YP_009243641.1   -------------------------------------------------E
106 YP_004300274.1   --------------------------------------------------
107 NP_056848.1      --------------------------------------------------
108 YP_006495799.1   ---------------------WRRRTELRGDQKVCP-HQWSDVAEVSLED
109 YP_009508409.1   --------------------------------------------------
110 NP_687035.1      -------------------------------------------------E
111 NP_045937.2      --------------------------------------------------
112 NP_612577.1      --------------------------------------------------
113 NP_619548.1      --------------------------------------------------
114 NP_056880.1      -------------------------------------------------A
115 YP_009121747.1   ---------------------WKPRMELPKGSKECE-HKWQENQ--VTCH
116 YP_009113237.2   ---------------------YLPWVAVKRKQYKCD-HEWTQGRDACLTE
117 YP_009508408.1   ---------------------WRCAVRVTQQEHDCR-HQWEYYS----PT
118 NP_955579.1      -------------------------------------------------A
119 YP_009229919.1   --------------------GWRYKVVLTKDQERCR-HTLEANCEVEAGK
120 YP_004442833.1   ----------NGPWLVGQPGGFEKQMYVTKKQYHCEGHEWFHNQ----PK
121 NP_862833.2      -------------------------------------------------A
122 NP_040563.1      -------------------------------------------------C
123 YP_443922.1      -------------------------------------------------A
124 NP_057933.2      -------------------------------------------------A
125 YP_009116631.1   -------------------PSWRRRKDISQVQKDCT-HTWSDTQEVPISD
126 YP_009513249.1   -------------------------------------------------T
127 YP_223871.1      -------------------------------------------------A
128 NP_127504.1      ------------------------ILSEFTARLSGTLREWWNSVGPDDKN
129 NP_044929.1      -------------------------------------------------Q
130 NP_057860.1      -------------------------------------------------S
131 NP_040973.1      ------------------------------------------------LE
132 NP_777317.1      HDVFMLREDEPDDYLVGNPPSWRSKLRVSRQQYYCQ-HNWQYDH----KE
133 NP_054716.1      -------------------------------------------------I
134 YP_001856242.1   -------------------------------------------------E
135 NP_954565.2      -------------------------------------------------S
136 NP_569150.1      ----------PREYLVGEPSEWRSKMKVSRRQYFCK-HEWKFEE----TH
137 NP_659397.1      --------------------------------------------------
138 YP_009513211.1   -------------------------------------------------A
139 YP_233107.1      ----------PSEYYIGEPG-WRSQMKVSKTEFHCQ-HEWKFDK----ED
140 YP_009109692.1   -------------------------------------------------A
141 NP_057849.4      -------------------------------------------------E
142 NP_049560.1      -------------------------------------------------S
143 YP_009506267.1   ---------------------TKVSVKVTQKEHDCT-HDWVLYG---EN
144 NP_039820.1      -------PAEVQMGVLKPKSSWRRPIQATLEEINCH-HNWTAIS----TG
145 NP_056790.1      -------------------------------------------------A
146 YP_002321513.1   ---------------------WRSKPKLSDDQLLCD-HEGFKICAKPAEA
147 NP_955591.1      -------------------------------------------------A
148 YP_001956722.2   -------------------------------------------------Q
149 NP_056902.1      -------------------------------------------------E
150 NP_056886.1      -------------------------------------------------A
151 NP_041261.1      -------------------------------------------------S
```

```
001 YP_004442836.1   RTICRACGSEAVHKHRIDCLKCDMTICLMCQPWFYKDINTEEVKK-----
002 NP_056728.1      --------------------------------------------------
003 YP_009508537.1   LETLITKLNEAEYLQGRRGDKTLIMKVNASY--TTGYIRYYNEGEK----
004 YP_004732983.2   -QACSTCQTITVLGRRATCTLCLLNLCSLCAGLDFGLKIVPKTA------
005 YP_009508407.1   --------------------------------------------------
006 NP_040939.1      WEACRKKLEEMEGN-YYNKDKDVYGQLAWG-DKAIEYIVYQEKGKP----
```

```
007 YP_003864102.1  LDKVETAIAE-QFVTHINYSQPLILLVFNTALTPTGLFWQSN--------
008 YP_009002585.1  LEKCSFCKQETRRRMRLHCPKCRITSCVLCSSNYCGIAIKPE-PV-----
009 YP_003284237.1  YEKCKACRLEARANERMKCKECKMVICCLCSNYCFDITIPRE-------R
010 YP_009268869.1  WEECRRKLAEMEGN-YYDEEKDVYGQIDWG-DKAIEYIVFQERGKP----
011 YP_009506251.1  LINCHFCKQPTNLRGRVHCPTCQLTSCYMCAPIYCDVQVQQQ-PK-----
012 NP_955611.1     WREIVRLSTT-AALERWDPALPLEGAVARCEQGAIGVLGQGLSTHP----
013 YP_008567619.1  PIQCLCCKRNSQKNFRIHCPICKVTSCGLCSTFCFDKSIQVKKEY-----
014 NP_149413.1     TRPCKRCLKTVSKGEYAYCKICKVGVCNDCALWCYQIKAPTCSQE-----
015 YP_004442824.1  MARCNACHHEALKGSRMDCLKCDLTICILCRPHYYPKIEIP-VEK-----
016 YP_001931967.1  --------------------------------------------------
017 YP_002455786.1  LHNIQQALHH-NCRGRLDSTLPLLGLIFLSPSGTTSVLFQTNHK------
018 YP_009109694.1  FEALKKALLSAPALGLPDTSKPFTLFLDERQGIAKGVLTQKLGPWK----
019 NP_789739.1     LAENREILKEPVHGVYYDPSKDLIAEIQKQGQGQWTYQIYQEPFKN----
020 NP_955619.1     LVQLRQALSQ-NCRSRLVQTLPLLGAIMLTLTGTTTVVFQSKQQ------
021 YP_224289.1     VERCRACTYYITSQPRIKCTKCKIPICNLCTNYCYGIKMADEK------E
022 YP_009182100.1  WEIPLMRIEEDDDYSDTSSIYSLETESDSESSDSDQESEDEDTVRMIKTQ
023 NP_705927.1     LAENREILKEPVHGVYYDPSKDLIAEIQKQGQGQWTYQIYQEPFKN----
024 YP_009508577.1  LEEIVSALNHAGNLEQRDNESPLVVKLNASP--KTGYIRYYNKGGQ----
025 YP_004442830.1  ARVCQRCFCKASKGQYIICGKCKVKVCNLCSDYCYGISLVKKEED-----
026 NP_758887.1     YEENKLIVQEEVHGVYYQPEKPLMAKVQKLTQGQWSYQIEQEDNKP----
027 YP_009508406.1  --------------------------------------------------
028 YP_610965.1     SW-CTYCKAPTSVLYRTKCTLCLLLCCPYCALHFH--QHRVRKER-----
029 YP_001931961.1  --------------------------------------------------
030 YP_241114.1     LQLIEEAIRR-QQITYCDYQRSWGLYILPTPRAPTGVLYQDK--------
031 YP_006732334.1  --------------------------------------------------
032 YP_009041481.1  NKRCGFCRNETRVTSRILCPACKLVACLLCASRRLGIEVRKEEE------
033 YP_001036293.1  TGLCKFCKRDITQRWHAKCMHCNLVTCGFCSQHYLDLEIPCQREE-----
034 YP_605811.1     SDYCRACKNRALQGSRADCTQCKMIICSLCKPYYFQDGSPIPAQG-----
035 NP_955577.1     FEDIKKALLSSPALGLPDITKPFELFIDENSGFAKGVLVQKLGPWK----
036 YP_003208050.1  AK-CAFCGVQTNPEMRMYCDICRLTACPMCSKYYL--DIPVRMNK-----
037 NP_789740.1     LAENREILKEPVHGVYYDPSKDLIAEIQKQGQGQWTYQIYQEPFKN----
038 YP_004442839.1  AGYCRACKHQALKDSRADCSTCKITICSLCKPHYYKDAAPIPVQK-----
039 YP_009507791.1  VRIAKEAIENLTDMSYFDNKKPVYCKFSIVGVHQVAYIVYQDQGQ-----
040 YP_004581513.1  --------------------------------------------------
041 NP_043933.1     --------------------------------------------------
042 YP_009507248.1  FEHCKTLLTNDPILQYPDFTREFNLTTDASNFAIGAVLSQGPIGSD----
043 YP_002117531.1  ELKCRCCFLPTRSKDRIHCKKCKITVCNMCSEFCYNIKIVRELPS----S
044 YP_004347415.1  --------------------------------------------------
045 NP_056891.1     LEKVETAIAE-QFVTHINYSLPLIFLIFNTALTPTGLFWQDN--------
046 YP_009513242.1  LQYIISVLNQADNLEERNPETRLILKVNSSP--SAGYIRYYNEGSK----
047 YP_009352866.1  PKNCGFCRLETRITSRISCPQCQLKACLLCAKRRLGIEVKKEEE------
048 YP_009165750.1  --------------------------------------------------
049 NP_040840.1     WEACRQKLKEMEGN-YYDEEKDIYGQLDWG-NKAIEYIVFQEKGKP----
050 YP_004222728.1  WREIVQLSTT-AALERWDPALPLEGAVARCEQGAIGVLGQGLSTHP----
051 YP_009506264.1  QADCRACGLKARSGGRMDCGKCGIIICCLCSQYCYDVTIPRE-------Q
052 YP_567050.1     LHAIQQALQH-NCRSRLNPALPILGLISLSSSGTTSVLFQARQR------
053 YP_009508571.1  LQTIIKALNNADNLEERIPEKRLIIKVNTSP--SAGYVRYYNETGK----
054 NP_056803.1     LNKVIEALNTASNLEERLPDQRLVIKVNTSP--SAGYVRYYNESGK----
055 NP_955564.1     LQLMNERLST-ARVKRLDLSQPWSLCILKTEYTPTACLWQDG--------
056 YP_009508411.1  SRTCRFCKREVMKRFNAHCSKCSITTCGMCSNHYLGFQIPCQQEE-----
057 YP_009140788.1  AIKCSFCRRESLKRARIHCPLCLLTACNLCGPYYLKIEVPVD-PA-----
058 YP_009508410.1  --------------------------------------------------
059 NP_040333.1     YQEIKQALLTAPALGLPDLTKPFELFVDEKQGYAKGVLTQKLGPWR----
060 YP_008992013.1  QFVQLPEVSSALAILHDQFLGDPAAAFEAARRDYLSMKCCSLDAKDLDFH
061 NP_041734.1     HINCHFCKQPTQLRSRIHCPTCQLTSCFMCAPIYCNMIVQQQ-PK-----
062 NP_056895.1     IAELRQALSH-NARSRYNEQEPLLAYVHLTRAGSTLVLFQKGAQ------
063 YP_009109689.1  FEALKKALLSAPALGLPDTSKPFTLFLDERQGIAKGVLTQKLGPWK----
064 YP_002519387.1  --------------------------------------------------
065 YP_595725.1     VKPCKRCLKTVTKGQYIYCRTCKITVCHECSEFCYNIKIEGAEAV-----
066 YP_009508551.1  LNMVIEALNTASNLEERLPEQRLVIKVNTSP--SAGYVRYYNETGK----
067 YP_001497148.1  FEALKEALLSAPALGLPDTSKPFTLFIDEKQGIAKGVLTQKLGPWK----
068 NP_041186.1     LQSIEEAIRQ-QQITYCDYQRSWGLYILPTPRAPTGVLYQDK--------
069 NP_663784.1     LEENKIILSQEQEGYYYQEEKELEATIQKSQGHQWTYKIHQEE-KI----
070 YP_009508443.1  ADKCRFCKREVMKRWNARCQHCNILTCGMCSKHYVDLELPCQQEE-----
071 YP_003987465.1  LKNCRACGATAQTGERMDCPLCNMTICSSCSNYCFDITIPKNEERVQAIV
072 NP_861410.1     --------------------------------------------------
073 YP_009345071.1  PEHCRCCNKETSDRCRIQCGYCKITACSLCAFSYFDIKVTQRKEF-----
074 YP_009508582.1  LEEIVSALNHAGNLEQRDDESPLVVKLNASP--KTGYIRYYNKGGQ----
075 NP_569141.1     --------------------------------------------------
076 YP_002916057.1  --------------------------------------------------
077 NP_777384.2     IAELRQALSH-NARSRYNEQEPLLAYVHLTRAGSTLVLFQKGAQ------
078 YP_233110.1     TDHCRACKNLALRGSRADCTQCKIIICSLCKPYYFQDGSPIPAQS-----
079 NP_542258.1     LKIIQKALAL-NCRSRLVSQLPILALIILRPTGTTAVLFQTKQK------
```

```
080 YP_009130664.1   SFKCFCCKEETSVNYRMHCPQCLITTCNLCALYYFGQKVPASRPN-----
081 YP_009508566.1   LNNLIIALNHADNLEQRNGEVPLVIKINASN--TTGYIRFYNKNGK----
082 YP_006607892.1   --------------------------------------------------
083 YP_009506270.1   VTTCKACGSEAVPKHRVDCKKCGMTVCLMCQPWYYKDINTEEVKK-----
084 NP_042513.1      --------------------------------------------------
085 YP_006907834.1   --------------------------------------------------
086 YP_004442827.1   VRICQRCFQKSPKGQYIICGICKVKVCHQCTEYCYGITMKGKEVQ-----
087 YP_009408594.1   IR-CLFCKYNTKKRMRIHCTQCKVTACPMCSDYYL--QVHITPG------
088 YP_009508561.1   LNELISALNHAGNLEQRRGDSPLVVKVNASD--KTGYIRYYNDNSL----
089 NP_056907.1      LHAIQQALQH-NCRSRLNPALPILGLISLSSSGTTSVLFQARQR------
090 YP_002308474.1   LNTLQKALTH-NCRSRIVSNLPILALIMLRPTGTTAVLFQTKQK------
091 YP_007761644.1   --------------------------------------------------
092 NP_040550.1      WREIVQLSTT-AALERWDPALPLEGAVARCEQGAIGVLGQGLSTHP----
093 NP_758808.1      HK-CHTCRRDTQKHYRLECQKCKFLVCSLCTIPYLGITMQFRQKQ-----
094 YP_009345075.1   NFNCLYCKNLTKQTRRWHCPKCQFTTCAYCAYLQLGKKMPPAPTK-----
095 NP_068729.1      --------------------------------------------------
096 NP_047255.1      FEDIKKALLSSPALGLPDITKPFELFIDENSGFAKGVLVQKLGPWK----
097 YP_006273075.1   AVHCSFCRRESIKRARIHCPLCLLTACNLCGPYYLKIEVPVE-PA-----
098 YP_009508888.1   LQELITLLNQADNLEERKPTTRLIIKVNSSS--HAGYIRYYNEGSK----
099 NP_395469.1      --------------------------------------------------
100 YP_009508546.1   LNTVIEALNTASNLEERLPNQRLVIKVNTSP--SAGYVRYYNESGK----
101 NP_056762.1      DKVCEICSYFTDYNKTVSCKTCETQYCKTCSDQLALEVTEVKKPTKEETM
102 YP_009508556.1   LQYIISVLNQADNLEERNPETRLILKVNSSP--SAGYIRYYNEGSR----
103 NP_569153.1      HVRCYPCKRETTQRARLHCKLCHITSCLMCGPTYFNKKITVQ-PM-----
104 NP_043924.1      RKMTPVESNLGICEQYAECAAWAISACNLVSGFGRKIIVTSHSPVKFILQ
105 YP_009243641.1   LQFFEHQLKS-KFLMRIDPLLPLTLYLINTFMYPTALLGQNE--------
106 YP_004300274.1   --------------------------------------------------
107 NP_056848.1      --------------------------------------------------
108 YP_006495799.1   -RFCSLCGDNTPVGRRVHCTTCKANLCPVCAWLEFGIRIVAAKT------
109 YP_009508409.1   --------------------------------------------------
110 NP_687035.1      YKNNQGIVQETQEGTYYDPLKELIATVQKQGEGQWTYQFTQEG-AV----
111 NP_045937.2      ---LKHAITTAPVLVVPDPAKPFQLYTSHSEHASIAVLTQKHAGRT----
112 NP_612577.1      --------------------------------------------------
113 NP_619548.1      --------------------------------------------------
114 NP_056880.1      LQLMNERLST-ARVKRLDLSQPWSLCILKTEYTPTACLWQDG--------
115 YP_009121747.1   SI-CHFCGIATTEISRLHCPECRLTACALCALHYLKIKIEVKRSK-----
116 YP_009113237.2   AK-CSFCKGERLQRQYVYCHKCDLLACGMCSWTYVNQKIEVK-PA-----
117 YP_009508408.1   VANCRFCKREVMQRWNAKCSKCSILTCGMCSEHYLGLNIPCQREE-----
118 NP_955579.1      FEDIKKALLSSPALGLPDITKPFELFIDENSGFAKGVLVQKLGPWK----
119 YP_009229919.1   EK-CSYCQYPTFKRMRCHCKGCGITACPMCSKFYL--EYLVTPEP-----
120 YP_004442833.1   VRSCQRCFGKFTKGQYVICGKCKVRVCHLCIQYCYGFSITKEEIP-----
121 NP_862833.2      LQFVEEAIRQ-QQITYCDYQRSWGLYILSTPRAPTGVLYQDK--------
122 NP_040563.1      VEEFNLKLKDPEWKDRIREGAELVIKIQMVPRGIVFDLLQDGNPIW----
123 YP_443922.1      FGLLKTDLSVAAALAAPDYSKPFHLDVSEKEGFASAILFQKQEGER----
124 NP_057933.2      YQEIKQALLTAPALGLPDLTKPFELFVDEKQGYAKGVLTQKLGPWR----
125 YP_009116631.1   -RVCSICSDETPHGRRVTCTTCNINLCPICARMDYGIMLIAAKDTK----
126 YP_009513249.1   LETLITKLNGAEYLQGRKGDKTLIMKVNASY--TTGYIRYYNEGEK----
127 YP_223871.1      FQSLKLALTQPPALALPSLDKPFQLFVEETSGAAKGVLTQALGPWK----
128 NP_127504.1      RFLTSQDFTWNIRILYSYFCGDQSQNKEELRRQIFEMKCLSYDRKKIDRH
129 NP_044929.1      LKTIIQALNSTENLEERRPDVDLIMKVHISN--TAGYIRFYNHGGQ----
130 NP_057860.1      LVQLRQALSQ-NCRSRLVQTLPLLGAIMLTLTGTTTVVFQSKQQ------
131 NP_040973.1      VQKAKKAIEEQVQLGYYDPSKELYAKLSLVGPHQISYQVYQKDPEKI---
132 NP_777317.1      PTTCKGCNIYAAPKNRMDCPQCKLTICVLCEPHYYKGAMALSKPE-----
133 NP_054716.1      LQKIIKELNASENLEQRKPDVELIVKVHVSP--TAGYIKFANKGSI----
134 YP_001856242.1   LAENREILKEPVHGVYYDPSKDLIAEIQKQGQGQWTYQIYQEPFKN----
135 NP_954565.2      LEKVETAIAE-QFVTHINYSLPLIFLIFNTALTPTGLFWQDN--------
136 NP_569150.1      VTICKACGSEAAPKHRIDCLKCEMTVCLMCQPWFYKNVNTEEVKK-----
137 NP_659397.1      --------------------------------------------------
138 YP_009513211.1   FGRIKEALLSAPALALPDLTKPFALYVDEKEGVARGVLTQTLGPWR----
139 YP_233107.1      SKHCRDCKFEARRDNRMDCSKCQLTICALCTYHCFK----ILIPR-----
140 YP_009109692.1   FEALKKALLSAPALGLPDTSKPFTLFLDERQGIAKGVLTQKLGPWK----
141 NP_057849.4      LAENREILKEPVHGVYYDPSKDLIAEIQKQGQGQWTYQIYQEPFKN----
142 NP_049560.1      LVQLRQALSQ-NCRSRLVQTVPLLGLVMLTLTGTTTVVFQSKQQ------
143 YP_009506267.1   NAHCRFCRREVMQRWNAKCNKCDITTCGMCSDHYVGLRVPCKRPE-----
144 NP_039820.1      QLACRSCKQFLAG---VQCHHCHAVYCFMCAEAYHDVQAEKILSKDYSFS
145 NP_056790.1      FDHIKKALLSAPALALPDLTKPFTLYIDERAGVARGVLTQTLGPWR----
146 YP_002321513.1   RIKCQECSIFTTPQARFLCDSCNLLLCAMCAQGYYGINIVEPVVPE----
147 NP_955591.1      YQEIKQALLTAPALGLPDLTKPFELFVDEKQGYAKGVLTQKLGPWR----
148 YP_001956722.2   LQNIISMLNSAENLEERNPEVRLIMKVNTSP--SAGYIRFYNEFAK----
149 NP_056902.1      LEENNEKIKNAQGLQYYNPEEEMLCEVEITKNYEATYVIKQSQGI-----
150 NP_056886.1      WREIVRLSTT-AALERWDPALPLEGAVARCEQGAIGVLGQGLSTHP----
151 NP_041261.1      LALINKAIQN-QSVQQISYNLPLVLLLLPTPHTPTAVFWQPNGTDPTK--
```

```
001 YP_004442836.1   TRVRIERVIDWKEIALRQ--HEVLKTSIANEK-----------QLSEENE
002 NP_056728.1      --------------------------------------------------
003 YP_009508537.1   --------------------------------------------------
004 YP_004732983.2   -TRADWKFQDRDSLIASL--YEHNAFLLRQVEGLKQELQAAKEQLQLLHS
005 YP_009508407.1   --------------------------------------------------
006 NP_040939.1      --------------------------------------------------
007 YP_003864102.1   --------------------------------------------------
008 YP_009002585.1   --QTSFFNEKS--LLTQQ--QEYIAYQTAEIK-----------KLQEQLK
009 YP_003284237.1   SFSSQHHEPKWKEIATAL--GEINAKLEQEKK-----------ALIEELN
010 YP_009268869.1   --------------------------------------------------
011 YP_009506251.1   --TPIPFSPNA--LLQQQ--AAYIRWLESENK-----------RLNELVE
012 NP_955611.1      --------------------------------------------------
013 YP_008567619.1   --V-APYNPRN--LLEEQ--QRYIEWCELEMK-----------RLKESII
014 NP_149413.1      --KKPSSSINYEKLARQL--LLENSKLKEEKK-----------ILIEELN
015 YP_004442824.1   IKTRETRSVDWKKIVLEQ--HEALLSISAREK-----------SLRKELE
016 YP_001931967.1   --------------------------------------------------
017 YP_002455786.1   --------------------------------------------------
018 YP_009109694.1   --------------------------------------------------
019 NP_789739.1      --------------------------------------------------
020 NP_955619.1      --------------------------------------------------
021 YP_224289.1      KEKEKEKESDLLGLVNNC--REIMAKYRTEKE-----------VLLEELN
022 YP_009182100.1   FGIKTIRLNNWPHDTQEIGDVVSFEAILEYEKFECDFCKTSADQYTPVIQ
023 NP_705927.1      --------------------------------------------------
024 YP_009508577.1   --------------------------------------------------
025 YP_004442830.1   --PEVD----YKSLAASL--ISENERLKLEKK-----------VLLEELN
026 NP_758887.1      --------------------------------------------------
027 YP_009508406.1   --------------------------------------------------
028 YP_610965.1      --PPVEAP---EQLIQNL--IGYIDSLLKENE-----------RLKEER-
029 YP_001931961.1   --------------------------------------------------
030 YP_241114.1      --------------------------------------------------
031 YP_006732334.1   --------------------------------------------------
032 YP_009041481.1   ---VTWKFQNKDELINSL--YHHNAFLIKENS-----LLKEQLQLAQRRI
033 YP_001036293.1   --V-QVYNPQR--LLQEQ--QNYIQVCEVEIE-----------KLKESVK
034 YP_605811.1      STPSGYSHDDWMGSANRW--KAHYEFSQARRK-----------SLKADLE
035 NP_955577.1      --------------------------------------------------
036 YP_003208050.1   --PPPPPPENKDELIHEL--VVYTETLLKRVK-----------ELEEQL-
037 NP_789740.1      --------------------------------------------------
038 YP_004442839.1   KVVK----DDWMNMAKRF--MDSWKKAEDEKM-----------ELKEELN
039 YP_009507791.1   --------------------------------------------------
040 YP_004581513.1   --------------------------------------------------
041 NP_043933.1      --------------------------------------------------
042 YP_009507248.1   --------------------------------------------------
043 YP_002117531.1   SLLKDRKKVDYIGVCIEV--LEENKKLRTEKE-----------VLIQDLN
044 YP_004347415.1   --------------------------------------------------
045 NP_056891.1      --------------------------------------------------
046 YP_009513242.1   --------------------------------------------------
047 YP_009352866.1   ---VTWKFQNKDELINNL--YLHNAFLIKENS-----LLKEQLQLAQRRV
048 YP_009165750.1   --------------------------------------------------
049 NP_040840.1      --------------------------------------------------
050 YP_004222728.1   --------------------------------------------------
051 YP_009506264.1   NQ-VAYASPNWREIALRQ--QEIIRDLEAEKR-----------ALIEEVN
052 YP_567050.1      --------------------------------------------------
053 YP_009508571.1   --------------------------------------------------
054 NP_056803.1      --------------------------------------------------
055 NP_955564.1      --------------------------------------------------
056 YP_009508411.1   --VRPVYEPRQ--LMNEQ--MTYIKHCEEEIA-----------RLKQEVA
057 YP_009140788.1   --PSTPLNPRR--IIQEQ--HNYIGWCEAEIE-----------RLNKEVK
058 YP_009508410.1   --------------------------------------------------
059 NP_040333.1      --------------------------------------------------
060 YP_008992013.1   YKRMSLLFYKLNGFNEPTLKHVFLASLPEELQPDIQRQLTASNLVIDNIS
061 NP_041734.1      --PPVPFNTHT--LLQQQ--AAYIQWLEKENQ-----------RLTEAVE
062 NP_056895.1      --------------------------------------------------
063 YP_009109689.1   --------------------------------------------------
064 YP_002519387.1   --------------------------------------------------
065 YP_595725.1      --KPPEKKSNYELLAKQL--LIENSKLKMEKE-----------ILIEELN
066 YP_009508551.1   --------------------------------------------------
067 YP_001497148.1   --------------------------------------------------
068 NP_041186.1      --------------------------------------------------
069 NP_663784.1      --------------------------------------------------
070 YP_009508443.1   --IRQVYNPQQ--LMMEQ--QNYIQSCEMEIQ-----------RLKDELA
```

```
071 YP_003987465.1    SSSTASREESWKTIALEQ--RKLIKKLENEKE-----------ALIVELN
072 NP_861410.1       --------------------------------------------------
073 YP_009345071.1    --VSPYYATPS--LLKEN--QEYISWCQAELN-----------KTGEAMT
074 YP_009508582.1    --------------------------------------------------
075 NP_569141.1       --------------------------------------------------
076 YP_002916057.1    --------------------------------------------------
077 NP_777384.2       --------------------------------------------------
078 YP_233110.1       STPSGYSYDDWMGSANRW--KAHYEFSQARRK-----------SLKADLE
079 NP_542258.1       --------------------------------------------------
080 YP_009130664.1    --P-IQYNPLS--LMHEQ--QGYITWCEAEIT-----------RLKQEVE
081 YP_009508566.1    --------------------------------------------------
082 YP_006607892.1    --------------------------------------------------
083 YP_009506270.1    TRVRIERVIDWKDIALKQ--HEVLKTSIAREK-----------QLSEEVE
084 NP_042513.1       --------------------------------------------------
085 YP_006907834.1    --------------------------------------------------
086 YP_004442827.1    --PEED----YKELAKSL--LAENKKLKLERK-----------LLLEELN
087 YP_009408594.1    --PPVQKEFDEKPVLREL--LQYVTWLQTEND-----------RLKAELR
088 YP_009508561.1    --------------------------------------------------
089 NP_056907.1       --------------------------------------------------
090 YP_002308474.1    --------------------------------------------------
091 YP_007761644.1    --------------------------------------------------
092 NP_040550.1       --------------------------------------------------
093 NP_758808.1       --KSQPENPNLVRELLEH--AIFLEEKCKNQE-----------LLSETQI
094 YP_009345075.1    --EPAVRSKVQSSAVHAE--LSRLRNERDHLE-----------RIVDSFL
095 NP_068729.1       --------------------------------------------------
096 NP_047255.1       --------------------------------------------------
097 YP_006273075.1    --PSTPINPRR--IIQEQ--HNYIGWCEVEIE-----------RLNKEVK
098 YP_009508888.1    --------------------------------------------------
099 NP_395469.1       --------------------------------------------------
100 YP_009508546.1    --------------------------------------------------
101 NP_056762.1       IDDLKLNVKNLEFRVTILEHKVEMQNLQDKFETMQIR-------------
102 YP_009508556.1    --------------------------------------------------
103 NP_569153.1       --PQAPFNQKG--LLQQQ--QEYIAWCNNEIA-----------RLKEEVA
104 NP_043924.1       TTPNVSNQRLARWHRILTQEDITIETDASIQG------------------
105 YP_009243641.1    --------------------------------------------------
106 YP_004300274.1    --------------------------------------------------
107 NP_056848.1       --------------------------------------------------
108 YP_006495799.1    -DATKWNYLNKDELIAQL--YEHNAFLTHQNK----ALQGRIEELTKGRN
109 YP_009508409.1    --------------------------------------------------
110 NP_687035.1       --------------------------------------------------
111 NP_045937.2       --------------------------------------------------
112 NP_612577.1       --------------------------------------------------
113 NP_619548.1       --------------------------------------------------
114 NP_056880.1       --------------------------------------------------
115 YP_009121747.1    --GSSKGS-----EVIVP--DEETQQPNSIWK-----------LLAQKDN
116 YP_009113237.2    --AKPVPKPLSTDLVRLQ--SEHLLWQEKQID-----------KLKEEVL
117 YP_009508408.1    --IRQVYNPRQ--LMMEQ--QSYIQSCELEIQ-----------RLKEEIQ
118 NP_955579.1       --------------------------------------------------
119 YP_009229919.1    --PAPKQYSSKDRLIQEL--MSYNVYLLSENE-----------RLQEIIN
120 YP_004442833.1    --KEEEN---WKELASTL--MIENRKLKLEKK-----------LLLKELN
121 NP_862833.2       --------------------------------------------------
122 NP_040563.1       --------------------------------------------------
123 YP_443922.1       --------------------------------------------------
124 NP_057933.2       --------------------------------------------------
125 YP_009116631.1    -SAAHWQYQNKDELIQHL--YEHNAFLTRKVA----ELTSQLQEFHNRRP
126 YP_009513249.1    --------------------------------------------------
127 YP_223871.1       --------------------------------------------------
128 NP_127504.1       FQRMIKLFYHIG--GDISLKQAFISSLPPILSERISALIKERGTSVTQMH
129 NP_044929.1       --------------------------------------------------
130 NP_057860.1       --------------------------------------------------
131 NP_040973.1       --------------------------------------------------
132 NP_777317.1       PAPEEPKHLYWMRYAKEQ--LDKYKEAQKEKL-----------ELRKRVE
133 NP_054716.1       --------------------------------------------------
134 YP_001856242.1    --------------------------------------------------
135 NP_954565.2       --------------------------------------------------
136 NP_569150.1       TRVRIERVIDWKDIALKQ--HEVLKTSIANEK-----------QLSEEVE
137 NP_659397.1       --------------------------------------------------
138 YP_009513211.1    --------------------------------------------------
139 YP_233107.1       KQTTPSMKHDWRELAERQ--SELLRKFCQREK-----------ELEQEVQ
140 YP_009109692.1    --------------------------------------------------
141 NP_057849.4       --------------------------------------------------
142 NP_049560.1       --------------------------------------------------
143 YP_009506267.1    --VQQVYQPTN--LLVEQ--RNYIMSCEEEIA-----------RLKNEIE
```

```
144 NP_039820.1         ARGKKGKAVIIEEDEIEG--EFLISQLQQENQ-----------RLQKQVE
145 NP_056790.1         --------------------------------------------------
146 YP_002321513.1      -LQHHIERPKIDKGKKKVSWQEEIQDLEGMSEIPYARARQSLRMLSSDEE
147 NP_955591.1         --------------------------------------------------
148 YP_001956722.2      --------------------------------------------------
149 NP_056902.1         --------------------------------------------------
150 NP_056886.1         --------------------------------------------------
151 NP_041261.1         --------------------------------------------------


001 YP_004442836.1      ELKEKI-------------KELKGKELITVEEISEEEG------------
002 NP_056728.1         --------------------------------------------------
003 YP_009508537.1      --------------------------------------------------
004 YP_004732983.2      VDMINLSD-----------DGLENFSLEEKSFLRGGGGTSSSS-------
005 YP_009508407.1      --------------------------------------------------
006 NP_040939.1         --------------------------------------------------
007 YP_003864102.1      --------------------------------------------------
008 YP_009002585.1      QTE----------EKLAEAERKIRGYENMEVLEKERKEMAL---------
009 YP_003284237.1      RALEQI-------------KDYKQKGLPTAIIQEVDE-------------
010 YP_009268869.1      --------------------------------------------------
011 YP_009506251.1      FYK----------RLLDGGSRLQQ------ELEADQKELEE---------
012 NP_955611.1         --------------------------------------------------
013 YP_008567619.1      QNDELRMENDRLQLENRDLKRQIEKLQDKQKMKALMEDLGEEF-------
014 NP_149413.1         KEIKAR-------------QEAEKNKAVFVEETSSEVN------------
015 YP_004442824.1      QALEQL-------------KHYKRESEIIIEEE----P------------
016 YP_001931967.1      --------------------------------------------------
017 YP_002455786.1      --------------------------------------------------
018 YP_009109694.1      --------------------------------------------------
019 NP_789739.1         --------------------------------------------------
020 NP_955619.1         --------------------------------------------------
021 YP_224289.1         NALKQA-------------KEYKEK--FEHLEGICEEQ------------
022 YP_009182100.1      RSDLSKEGTISYHTWCFTSLVMRETKNAEVPQLVQREVDLAKLRDAKKKF
023 NP_705927.1         --------------------------------------------------
024 YP_009508577.1      --------------------------------------------------
025 YP_004442830.1      KEILEK-------------NQLRN-ELPVIEETNSEAA------------
026 NP_758887.1         --------------------------------------------------
027 YP_009508406.1      --------------------------------------------------
028 YP_610965.1         DDLQQK-------LQRTEERLQSLLGDYLHEASEDMAA------------
029 YP_001931961.1      --------------------------------------------------
030 YP_241114.1         --------------------------------------------------
031 YP_006732334.1      --------------------------------------------------
032 YP_009041481.1      ANNAQLTG----ADLQAFMEDYRNLTLETAQKIATETPKPVGK-------
033 YP_001036293.1      AANKIA--------EDKIMAASIRHAVELEELRKQLQAIK----------
034 YP_605811.1         KAEEEL-------------KFYKQKEKERARLKNQIP-------------
035 NP_955577.1         --------------------------------------------------
036 YP_003208050.1      KEKELE-------IARLELMIN-------DEGKQETAE------------
037 NP_789740.1         --------------------------------------------------
038 YP_004442839.1      DIKEQI-------------KFYKGKEKE--VCTEEIP-------------
039 YP_009507791.1      --------------------------------------------------
040 YP_004581513.1      --------------------------------------------------
041 NP_043933.1         --------------------------------------------------
042 YP_009507248.1      --------------------------------------------------
043 YP_002117531.1      SALEQL-------------KNTKGK--EIIVEEESPE-------------
044 YP_004347415.1      --------------------------------------------------
045 NP_056891.1         --------------------------------------------------
046 YP_009513242.1      --------------------------------------------------
047 YP_009352866.1      ANNAMLTG----ADLQAFMEDYRNLTVETSQRIAEGTRGPLGQ-------
048 YP_009165750.1      --------------------------------------------------
049 NP_040840.1         --------------------------------------------------
050 YP_004222728.1      --------------------------------------------------
051 YP_009506264.1      MSRDEI-------------KQYKAKGLTEIIEETPEET------------
052 YP_567050.1         --------------------------------------------------
053 YP_009508571.1      --------------------------------------------------
054 NP_056803.1         --------------------------------------------------
055 NP_955564.1         --------------------------------------------------
056 YP_009508411.1      AAKILS--------ENEIVSMKIREAVETDTLRKENETLR----------
057 YP_009140788.1      HYK----------GLYEALLTEKS-------LKKDYEDLRE---------
058 YP_009508410.1      --------------------------------------------------
059 NP_040333.1         --------------------------------------------------
060 YP_008992013.1      LGKIFQLAKTCLDKLCEQKQFFKDLLKDKEPFRSACKKPYLQIKCKQKKD
061 NP_041734.1         FYK----------KEAEE-LRLER------DLEQDRRSLEP---------
062 NP_056895.1         --------------------------------------------------
```

```
063 YP_009109689.1    --------------------------------------------------
064 YP_002519387.1    --------------------------------------------------
065 YP_595725.1       KEIKAH-------------QETKKGKELYIEEASTEVE------------
066 YP_009508551.1    --------------------------------------------------
067 YP_001497148.1    --------------------------------------------------
068 NP_041186.1       --------------------------------------------------
069 NP_663784.1       --------------------------------------------------
070 YP_009508443.1    REKKVA--------EEMVVAARISEAVSQQESKKQLEEMK----------
071 YP_003987465.1    QALEQL-------------KQTKGKGIPEHQDDALEKELLKELLREEKEK
072 NP_861410.1       --------------------------------------------------
073 YP_009345071.1    HIKKEN------AAQSEVITAQEHVINKLKEEKLDRQKIVEELKQELQ--
074 YP_009508582.1    --------------------------------------------------
075 NP_569141.1       --------------------------------------------------
076 YP_002916057.1    --------------------------------------------------
077 NP_777384.2       --------------------------------------------------
078 YP_233110.1       RAEEEL-------------KFYKQKEKEKAKLKDQIP-------------
079 NP_542258.1       --------------------------------------------------
080 YP_009130664.1    TS-----------------KKQVEYW--KQKYLSTQEEV-----------
081 YP_009508566.1    --------------------------------------------------
082 YP_006607892.1    --------------------------------------------------
083 YP_009506270.1    FLRKQN-------------KELKSKEPIVFEEDTEET-------------
084 NP_042513.1       --------------------------------------------------
085 YP_006907834.1    --------------------------------------------------
086 YP_004442827.1    NQILEN-------------TKIKR-ELPVIEETSSEAA------------
087 YP_009408594.1    AATERR-------PREDVGDVEID-----AELLKDLED------------
088 YP_009508561.1    --------------------------------------------------
089 NP_056907.1       --------------------------------------------------
090 YP_002308474.1    --------------------------------------------------
091 YP_007761644.1    --------------------------------------------------
092 NP_040550.1       --------------------------------------------------
093 NP_758808.1       ERIVSS-------EKQVKFYGILPTKKSNKSAGYDLQSNIDIEIPPG---
094 YP_009345075.1    DQLPSE-------LQRSVVEPSQNQKLQQQEEPSSQKG------------
095 NP_068729.1       --------------------------------------------------
096 NP_047255.1       --------------------------------------------------
097 YP_006273075.1    HYK----------GLYEALLMEKS-------LRKDYEELKE---------
098 YP_009508888.1    --------------------------------------------------
099 NP_395469.1       --------------------------------------------------
100 YP_009508546.1    --------------------------------------------------
101 NP_056762.1       --------------------------------------------------
102 YP_009508556.1    --------------------------------------------------
103 NP_569153.1       FYK----------QLAQERELQLQ-------LEQSRKELAG---------
104 NP_043924.1       --------------------------------------------------
105 YP_009243641.1    --------------------------------------------------
106 YP_004300274.1    --------------------------------------------------
107 NP_056848.1       --------------------------------------------------
108 YP_006495799.1    NDLIDLCEGDPDLMAEICDELFRKPASGETSILRGKERVKLEI-------
109 YP_009508409.1    --------------------------------------------------
110 NP_687035.1       --------------------------------------------------
111 NP_045937.2       --------------------------------------------------
112 NP_612577.1       --------------------------------------------------
113 NP_619548.1       --------------------------------------------------
114 NP_056880.1       --------------------------------------------------
115 YP_009121747.1    DWKQAI-------RDQAKEEALWQIKEKELRAQINEKD------------
116 YP_009113237.2    YWKNKYLAS-IERDLEQEFQTLPVREEDREKKRKEDDEMEEDRVSSNQKP
117 YP_009508408.1    NVKKVA--------EEQVVAAKINEAVAVKEMQEQIQELK----------
118 NP_955579.1       --------------------------------------------------
119 YP_009229919.1    KKLIED-------LEKGRMEAER-----FDQAKERRKG------------
120 YP_004442833.1    DQIEAN-------------KKIKE-EFPVIEETNSDAA------------
121 NP_862833.2       --------------------------------------------------
122 NP_040563.1       --------------------------------------------------
123 YP_443922.1       --------------------------------------------------
124 NP_057933.2       --------------------------------------------------
125 YP_009116631.1    EDLISLAD---DLEDVSILDNASNRGKEEKELFQFGTTIPIDH-------
126 YP_009513249.1    --------------------------------------------------
127 YP_223871.1       --------------------------------------------------
128 NP_127504.1       VGDIRQTAFYVLDDLCSKRKFFNQMKKMSRDLEKACTKSDLIIKGDKGCS
129 NP_044929.1       --------------------------------------------------
130 NP_057860.1       --------------------------------------------------
131 NP_040973.1       --------------------------------------------------
132 NP_777317.1       ILETMI-------------PQHKELEEE----------------------
133 NP_054716.1       --------------------------------------------------
134 YP_001856242.1    --------------------------------------------------
135 NP_954565.2       --------------------------------------------------
```

```
136 NP_569150.1       ILRKQS-------------KELKEKEPIIFEEDTEET-------------
137 NP_659397.1       --------------------------------------------------
138 YP_009513211.1    --------------------------------------------------
139 YP_233107.1       ELRREI-------------STLKTDLANLSVIQETPE-------------
140 YP_009109692.1    --------------------------------------------------
141 NP_057849.4       --------------------------------------------------
142 NP_049560.1       --------------------------------------------------
143 YP_009506267.1    AMKISS--------ENQLVAEKIARAVEEEHLKKEVQALK----------
144 NP_039820.1       RLQEEL-------------MKLHREKDEALKHSEKASR------------
145 NP_056790.1       --------------------------------------------------
146 YP_002321513.1    QPNRPQRQAEKESDTIIKMASIMAEQTKEISYWRDKAVELKQALGDLQRD
147 NP_955591.1       --------------------------------------------------
148 YP_001956722.2    --------------------------------------------------
149 NP_056902.1       --------------------------------------------------
150 NP_056886.1       --------------------------------------------------
151 NP_041261.1       --------------------------------------------------
```

```
001 YP_004442836.1    ---------AAMIEKIENLEKENELLNAIIT-------------------
002 NP_056728.1       --------------------------------------------------
003 YP_009508537.1    --------------------------------------------------
004 YP_004732983.2    ---------IKISSTTTP--PGFPTTPNRFQP------------------
005 YP_009508407.1    --------------------------------------------------
006 NP_040939.1       --------------------------------------------------
007 YP_003864102.1    --------------------------------------------------
008 YP_009002585.1    ---------KDMQRKIDEEIKKAAMQELPTEK------------------
009 YP_003284237.1    -------------SKTEELERENELLNAVLE-------------------
010 YP_009268869.1    --------------------------------------------------
011 YP_009506251.1    ---------LEADRKELEAHDKGKRVQFTEEEK-----------------
012 NP_955611.1       --------------------------------------------------
013 YP_008567619.1    --------DYEEEVNVLVEDWDNKRVTKELD-------------------
014 NP_149413.1       ---------DEIEMWKSKAELYEALYHEALNK------------------
015 YP_004442824.1    ---------VELMEELENTKRENELLNALLER------------------
016 YP_001931967.1    --------------------------------------------------
017 YP_002455786.1    --------------------------------------------------
018 YP_009109694.1    --------------------------------------------------
019 NP_789739.1       --------------------------------------------------
020 NP_955619.1       --------------------------------------------------
021 YP_224289.1       ------------EAENKALAQENQDLKEKVE-------------------
022 YP_009182100.1    EQAQLKEEQERLRATRRPVILEVYDDAEELEEMSKIGGHLEELSIKLRRP
023 NP_705927.1       --------------------------------------------------
024 YP_009508577.1    --------------------------------------------------
025 YP_004442830.1    ---------LEIDMLQEVVTSLQRQLEQKDQI------------------
026 NP_758887.1       --------------------------------------------------
027 YP_009508406.1    --------------------------------------------------
028 YP_610965.1       ---------EAERRRKGKALATSVPRATTGGPVI----------------
029 YP_001931961.1    --------------------------------------------------
030 YP_241114.1       --------------------------------------------------
031 YP_006732334.1    --------------------------------------------------
032 YP_009041481.1    ---------AALTWREEAEAAYKEELATRKNW------------------
033 YP_001036293.1    ---------LDKERLELENPDLKKILE------------------------
034 YP_605811.1       ---------EAIQAKLDDLEREKELNNILRIE------------------
035 NP_955577.1       --------------------------------------------------
036 YP_003208050.1    ---------EGEE-------------------------------------
037 NP_789740.1       --------------------------------------------------
038 YP_004442839.1    ---------QIYTQKLELLENE----NSLLKD------------------
039 YP_009507791.1    --------------------------------------------------
040 YP_004581513.1    --------------------------------------------------
041 NP_043933.1       ----------------------MENNLQLLEELN----HLEEG-------
042 YP_009507248.1    --------------------------------------------------
043 YP_002117531.1    -----------LLARISYLEQRELFMQELYE-------------------
044 YP_004347415.1    --------------------------------------------------
045 NP_056891.1       --------------------------------------------------
046 YP_009513242.1    --------------------------------------------------
047 YP_009352866.1    ---------AARNWRKEADVEWTEEEED----------------------
048 YP_009165750.1    --------------------------------------------------
049 NP_040840.1       --------------------------------------------------
050 YP_004222728.1    --------------------------------------------------
051 YP_009506264.1    ------------AKEIERLEAEKALLAELLE-------------------
052 YP_567050.1       --------------------------------------------------
053 YP_009508571.1    --------------------------------------------------
054 NP_056803.1       --------------------------------------------------
```

```
055 NP_955564.1         --------------------------------------------------
056 YP_009508411.1      ---------LENERLQLEVADLNKIIKGKQPIIPEDEELE----------
057 YP_009140788.1      ---------GDIKRGKGVMIQEPEDINYLGEN------------------
058 YP_009508410.1      --------------------------------------------------
059 NP_040333.1         --------------------------------------------------
060 YP_008992013.1      CDCSPKKKRHFRKFKSPEFSSRP-RRSRKPYRF-----------------
061 NP_041734.1         ---------TLLD--------KGKKVQILDPD------------------
062 NP_056895.1         --------------------------------------------------
063 YP_009109689.1      --------------------------------------------------
064 YP_002519387.1      --------------------------------------------------
065 YP_595725.1         ---------NEIETWKSRAELFEALYNEEVKKN-----------------
066 YP_009508551.1      --------------------------------------------------
067 YP_001497148.1      --------------------------------------------------
068 NP_041186.1         --------------------------------------------------
069 NP_663784.1         --------------------------------------------------
070 YP_009508443.1      ---------LENDRLTLENEDLKKEIRS----------------------
071 YP_003987465.1      AKKAEESLQELLKEEKEKAKKAEESLQELLR-------------------
072 NP_861410.1         --------------------------------------------------
073 YP_009345071.1      ---------QEKEKFLKEIEEQKELFMRQIAEEK----------------
074 YP_009508582.1      --------------------------------------------------
075 NP_569141.1         --------------------------------------------------
076 YP_002916057.1      --------------------------------------------------
077 NP_777384.2         --------------------------------------------------
078 YP_233110.1         ---------EAVQAKLDDLEKEKELNNILRIE------------------
079 NP_542258.1         --------------------------------------------------
080 YP_009130664.1      ---------FEKEFDELYKEKGKGVVIQEAT-------------------
081 YP_009508566.1      --------------------------------------------------
082 YP_006607892.1      --------------------------------------------------
083 YP_009506270.1      ---------AQLVQKLEDMEKENELMKILIS-------------------
084 NP_042513.1         --------------------------------------------------
085 YP_006907834.1      --------------------------------------------------
086 YP_004442827.1      ---------IEIDTLQEALERANELNKEKDAM------------------
087 YP_009408594.1      ---------LEIH-------------------------------------
088 YP_009508561.1      --------------------------------------------------
089 NP_056907.1         --------------------------------------------------
090 YP_002308474.1      --------------------------------------------------
091 YP_007761644.1      --------------------------------------------------
092 NP_040550.1         --------------------------------------------------
093 NP_758808.1         -------KCTVISTGTFLQMPDNMYGRLVERTSLAIQG------------
094 YP_009345075.1      ---------VCIQEPPPQKPAITFTRNEGSTVTLPTKKTRRRAGWDFYSP
095 NP_068729.1         --------------------------------------------------
096 NP_047255.1         --------------------------------------------------
097 YP_006273075.1      ---------GDKERRKGIMIQEPEEVNYLGED------------------
098 YP_009508888.1      --------------------------------------------------
099 NP_395469.1         --------------------------------------------------
100 YP_009508546.1      --------------------------------------------------
101 NP_056762.1         --------------------------------------------------
102 YP_009508556.1      --------------------------------------------------
103 NP_569153.1         ---------VDSRRRKDKGIVIDEGSCYFNPE------------------
104 NP_043924.1         --------------------------------------------------
105 YP_009243641.1      --------------------------------------------------
106 YP_004300274.1      --------------------------------------------------
107 NP_056848.1         --------------------------------------------------
108 YP_006495799.1      ---------LEEEVRGYKEEQELRTPLQKLSEGASTS-------------
109 YP_009508409.1      --------------------------------------------------
110 NP_687035.1         --------------------------------------------------
111 NP_045937.2         --------------------------------------------------
112 NP_612577.1         --------------------------------------------------
113 NP_619548.1         --------------------------------------------------
114 NP_056880.1         --------------------------------------------------
115 YP_009121747.1      -------KKAQEESQKWAEERQSLYKDIDDLLKIQQQ-------------
116 YP_009113237.2      YKS----STPAKERDRRTVIIQEPHQETVQKTV-----------------
117 YP_009508408.1      ---------LENDRLTLENHDLKELLK-----------------------
118 NP_955579.1         --------------------------------------------------
119 YP_009229919.1      ---------VVIRTPSGS--------------------------------
120 YP_004442833.1      ---------IEIEMLQELLNEANKSNNQ----------------------
121 NP_862833.2         --------------------------------------------------
122 NP_040563.1         --------------------------------------------------
123 YP_443922.1         --------------------------------------------------
124 NP_057933.2         --------------------------------------------------
125 YP_009116631.1      ---------IQNLENVAKIIEKWKDTPRVVIK------------------
126 YP_009513249.1      --------------------------------------------------
127 YP_223871.1         --------------------------------------------------
```

128 NP_127504.1 GYCNPSRRRKYKRFKLPSFKERDGRQYRKRRRF-----------------
129 NP_044929.1 --------------------------------------------------
130 NP_057860.1 --------------------------------------------------
131 NP_040973.1 --------------------------------------------------
132 NP_777317.1 ---------NELKIRLEDEKRENSLLQRLYE-------------------
133 NP_054716.1 --------------------------------------------------
134 YP_001856242.1 --------------------------------------------------
135 NP_954565.2 --------------------------------------------------
136 NP_569150.1 ---------AQLIQKLEDVERENELLNILIK-------------------
137 NP_659397.1 --------------------------------------------------
138 YP_009513211.1 --------------------------------------------------
139 YP_233107.1 ----------EDEEELQRLRQENTLLELLNQ-------------------
140 YP_009109692.1 --------------------------------------------------
141 NP_057849.4 --------------------------------------------------
142 NP_049560.1 --------------------------------------------------
143 YP_009506267.1 ---------LENDRLTLENEDQRKELEKK---------------------
144 NP_039820.1 ----------VFSTIQESDEAELNLIKEELR-------------------
145 NP_056790.1 --------------------------------------------------
146 YP_002321513.1 YAQ----LQKEVELLRWHNVVKNSQPKEADESLETNEDSIN---------
147 NP_955591.1 --------------------------------------------------
148 YP_001956722.2 --------------------------------------------------
149 NP_056902.1 --------------------------------------------------
150 NP_056886.1 --------------------------------------------------
151 NP_041261.1 --------------------------------------------------

001 YP_004442836.1 -----------------------------------QKEEEEVGYLNKIQE
002 NP_056728.1 --------------------------------------------------
003 YP_009508537.1 --------------------------------------------------
004 YP_004732983.2 -----------------------------------------LAQEKLKGI
005 YP_009508407.1 --------------------------------------------------
006 NP_040939.1 --------------------------------------------------
007 YP_003864102.1 --------------------------------------------------
008 YP_009002585.1 ---------------------------------------------ELWEQ
009 YP_003284237.1 -----------------------------------KKDKE----------
010 YP_009268869.1 --------------------------------------------------
011 YP_009506251.1 --------------------------------------VQLTEELEEIHM
012 NP_955611.1 --------------------------------------------------
013 YP_008567619.1 -----------------------------------------------DLE
014 NP_149413.1 ---------------------------------PSSSMIKEGLYQTQITH
015 YP_004442824.1 --------------------------------EKTEREELAIRASEEKEA
016 YP_001931967.1 --------------------------------------------------
017 YP_002455786.1 --------------------------------------------------
018 YP_009109694.1 --------------------------------------------------
019 NP_789739.1 --------------------------------------------------
020 NP_955619.1 --------------------------------------------------
021 YP_224289.1 -----------------------------------ELKEENEILKDQNTT
022 YP_009182100.1 VAASPGATSSNPLEENVDTPKDLKGQILLCPIFKGPIPKRTEADWKREEE
023 NP_705927.1 --------------------------------------------------
024 YP_009508577.1 --------------------------------------------------
025 YP_004442830.1 -----------------------------------------------IIQ
026 NP_758887.1 --------------------------------------------------
027 YP_009508406.1 --------------------------------------------------
028 YP_610965.1 --------------------------------------------REPLEG
029 YP_001931961.1 --------------------------------------------MSTSQR
030 YP_241114.1 --------------------------------------------------
031 YP_006732334.1 --------------------------------------MLSQQMMNHQEQ
032 YP_009041481.1 ------------------------------------------RQKKKIGE
033 YP_001036293.1 --------------------------------------------------
034 YP_605811.1 --------------------------------AEAELKALKESFKEREEA
035 NP_955577.1 --------------------------------------------------
036 YP_003208050.1 ------------------------------------------------AN
037 NP_789740.1 --------------------------------------------------
038 YP_004442839.1 --------------------------------LEEERKIQDIQVFKEIQA
039 YP_009507791.1 --------------------------------------------------
040 YP_004581513.1 --------------------------------------------------
041 NP_043933.1 ------------------------------------WEMLTDEDFKLQQN
042 YP_009507248.1 --------------------------------------------------
043 YP_002117531.1 -----------------------------------KEKFEKDLLQAQMRS
044 YP_004347415.1 --------------------------------------------------
045 NP_056891.1 --------------------------------------------------
046 YP_009513242.1 --------------------------------------------------

```
047 YP_009352866.1   -------------------------------------------------E
048 YP_009165750.1   --------------------------------------------------
049 NP_040840.1      --------------------------------------------------
050 YP_004222728.1   --------------------------------------------------
051 YP_009506264.1   -----------------------------------KEQAKNKMKIGELEA
052 YP_567050.1      --------------------------------------------------
053 YP_009508571.1   --------------------------------------------------
054 NP_056803.1      --------------------------------------------------
055 NP_955564.1      --------------------------------------------------
056 YP_009508411.1   --------------------------------------ELKQQVQQLKKS
057 YP_009140788.1   --------------------------------------------------
058 YP_009508410.1   --------------------------------------------------
059 NP_040333.1      --------------------------------------------------
060 YP_008992013.1   ---------------------------------------FRKRSSSSKDS
061 NP_041734.1      ---------------------------------------------EDQHT
062 NP_056895.1      --------------------------------------------------
063 YP_009109689.1   --------------------------------------------------
064 YP_002519387.1   --------------------------------------------MTKSKI
065 YP_595725.1      --------------------------------KASTSSVTEGMYQVQIDH
066 YP_009508551.1   --------------------------------------------------
067 YP_001497148.1   --------------------------------------------------
068 NP_041186.1      --------------------------------------------------
069 NP_663784.1      --------------------------------------------------
070 YP_009508443.1   ----------------------------------------------LKIQ
071 YP_003987465.1   -----------------------------------KEKEKVRKAEESQKE
072 NP_861410.1      --------------------------------------------------
073 YP_009345071.1   ------------------------------------------AKFLSLKQ
074 YP_009508582.1   --------------------------------------------------
075 NP_569141.1      --------------------------------------------------
076 YP_002916057.1   --------------------------------------------------
077 NP_777384.2      --------------------------------------------------
078 YP_233110.1      --------------------------------AETELKALKESFKEKEEA
079 NP_542258.1      --------------------------------------------------
080 YP_009130664.1   -----------------------------------------------NVF
081 YP_009508566.1   --------------------------------------------------
082 YP_006607892.1   --------------------------------------------MNMNTM
083 YP_009506270.1   -----------------------------------QKEAEEIQYLNKITE
084 NP_042513.1      --------------------------------------------------
085 YP_006907834.1   --------------------------------------------------
086 YP_004442827.1   -----------------------------------------------ILK
087 YP_009408594.1   -------------------------------------------------D
088 YP_009508561.1   --------------------------------------------------
089 NP_056907.1      --------------------------------------------------
090 YP_002308474.1   --------------------------------------------------
091 YP_007761644.1   --------------MNQTRFEDDPIEHDLNAQSWIKWFHLMRRQDTGYQW
092 NP_040550.1      --------------------------------------------------
093 NP_758808.1      ---------------------------------------ITVQGGVIDPD
094 YP_009345075.1   GDIALPIGAILTIQTQLKVQLPSGTYMQLAAKSSLAKLGVQVLGGVIDED
095 NP_068729.1      --------------------------------------------------
096 NP_047255.1      --------------------------------------------------
097 YP_006273075.1   --------------------------------------------------
098 YP_009508888.1   --------------------------------------------------
099 NP_395469.1      --------------------------------------------------
100 YP_009508546.1   --------------------------------------------------
101 NP_056762.1      --------------------------------------------------
102 YP_009508556.1   --------------------------------------------------
103 NP_569153.1      ---------------------------------------------ETTR-
104 NP_043924.1      --------------------------------------------------
105 YP_009243641.1   --------------------------------------------------
106 YP_004300274.1   --------------------------------------------------
107 NP_056848.1      --------------------------------------------------
108 YP_006495799.1   ---------------------------------------YKAPEESDEER
109 YP_009508409.1   --------------------------------------------------
110 NP_687035.1      --------------------------------------------------
111 NP_045937.2      --------------------------------------------------
112 NP_612577.1      --------------------------------------------MSLRNR
113 NP_619548.1      --------------------------------------MTSSSLFREGEL
114 NP_056880.1      --------------------------------------------------
115 YP_009121747.1   ----------------------------------------------IKDQ
116 YP_009113237.2   --------------------------------------EVEGKGKEKAEE
117 YP_009508408.1   -------------------------------------------------N
118 NP_955579.1      --------------------------------------------------
119 YP_009229919.1   ---------------------------------------------VHEAS
```

```
120 YP_004442833.1    --------------------------------------------------
121 NP_862833.2       --------------------------------------------------
122 NP_040563.1       --------------------------------------------------
123 YP_443922.1       --------------------------------------------------
124 NP_057933.2       --------------------------------------------------
125 YP_009116631.1    ----------------------------------------ETPESSTSNT
126 YP_009513249.1    --------------------------------------------------
127 YP_223871.1       --------------------------------------------------
128 NP_127504.1       ---------------------------------------FRR--SKTSKA
129 NP_044929.1       --------------------------------------------------
130 NP_057860.1       --------------------------------------------------
131 NP_040973.1       --------------------------------------------------
132 NP_777317.1       -----------------------------------DERTKNTELETKIKE
133 NP_054716.1       --------------------------------------------------
134 YP_001856242.1    --------------------------------------------------
135 NP_954565.2       --------------------------------------------------
136 NP_569150.1       -----------------------------------QKEKDEIQYLNEIIE
137 NP_659397.1       --------------------------------------MKRNTLFKWGEF
138 YP_009513211.1    --------------------------------------------------
139 YP_233107.1       ------------------------------------KQQKDIEALQQQNE
140 YP_009109692.1    --------------------------------------------------
141 NP_057849.4       --------------------------------------------------
142 NP_049560.1       --------------------------------------------------
143 YP_009506267.1    ----------------------------------------NKEVLDLQSY
144 NP_039820.1       --------------------------------------QFKEETRMAIAQ
145 NP_056790.1       --------------------------------------------------
146 YP_002321513.1    ----------------------------------------DGQEDASGAE
147 NP_955591.1       --------------------------------------------------
148 YP_001956722.2    --------------------------------------------------
149 NP_056902.1       --------------------------------------------------
150 NP_056886.1       --------------------------------------------------
151 NP_041261.1       --------------------------------------------------
```

```
001 YP_004442836.1    LEERIRKLEQQMKYRDEEMQVILK--------------------------
002 NP_056728.1       --------------------------------------------------
003 YP_009508537.1    --------------------------------------------------
004 YP_004732983.2    QEDLSLAVQFDDVR------------------------------------
005 YP_009508407.1    --------------------------------------------------
006 NP_040939.1       --------------------------------------------------
007 YP_003864102.1    --------------------------------------------------
008 YP_009002585.1    REGKAHIAGAVIAA------------------------------------
009 YP_003284237.1    ----IKDLKER---------------LQWLSQ------------------
010 YP_009268869.1    --------------------------------------------------
011 YP_009506251.1    AYLEEDTITRVVGH------------------------------------
012 NP_955611.1       --------------------------------------------------
013 YP_008567619.1    AELNKVGVIIPEEK------------------------------------
014 NP_149413.1       LTKELRQLEAGMEVNQ----------------------------------
015 YP_004442824.1    LKTEIRELRQENEALKTG--------------------------------
016 YP_001931967.1    --------------------------------------------------
017 YP_002455786.1    --------------------------------------------------
018 YP_009109694.1    --------------------------------------------------
019 NP_789739.1       --------------------------------------------------
020 NP_955619.1       --------------------------------------------------
021 YP_224289.1       MGQEMVKIQEKQQEAEKFLRREIKRREQANTE------------------
022 YP_009182100.1    ERNRKLVESYRKLKLFERYKEVRNEVLQQIPQHQIPINTIITVKSSPTEM
023 NP_705927.1       --------------------------------------------------
024 YP_009508577.1    --------------------------------------------------
025 YP_004442830.1    LTNKLKGYEQASSSSE----------------------------------
026 NP_758887.1       --------------------------------------------------
027 YP_009508406.1    --------------------------------------------------
028 YP_610965.1       CVEECSDEDTEEAA------------------------------------
029 YP_001931961.1    SLSSTQIYPIRKALMN----------------------------------
030 YP_241114.1       --------------------------------------------------
031 YP_006732334.1    NKKEILFKAIMAKIRQR---------------------------------
032 YP_009041481.1    EEDEEEEEEEIIVG------------------------------------
033 YP_001036293.1    -EFEQVNVLICDG-------------------------------------
034 YP_605811.1       LMGEITALEEGVKVHKEEAEELQEENQKLKEKILAFEKEA----------
035 NP_955577.1       --------------------------------------------------
036 YP_003208050.1    FIEESSTIHCEEDE------------------------------------
037 NP_789740.1       --------------------------------------------------
038 YP_004442839.1    MRKEIESLKAAK---EEENKAHEEEKKALKEEILALKRSI----------
```

```
039 YP_009507791.1   --------------------------------------------------
040 YP_004581513.1   ----------MIQG------------------------------------
041 NP_043933.1      E-------------------------------------------------
042 YP_009507248.1   --------------------------------------------------
043 YP_002117531.1   MAAEMERLQQQVS---------QLEEQIAQKA------------------
044 YP_004347415.1   --------------------------------------------------
045 NP_056891.1      --------------------------------------------------
046 YP_009513242.1   --------------------------------------------------
047 YP_009352866.1   DEGEQVGGVVFPEG------------------------------------
048 YP_009165750.1   -----------MDLL-----------------------------------
049 NP_040840.1      --------------------------------------------------
050 YP_004222728.1   --------------------------------------------------
051 YP_009506264.1   EKALLAELLEREQ-----AKCE-ALEEAQQTT------------------
052 YP_567050.1      --------------------------------------------------
053 YP_009508571.1   --------------------------------------------------
054 NP_056803.1      --------------------------------------------------
055 NP_955564.1      --------------------------------------------------
056 YP_009508411.1   LEEKRLGVIICEG-------------------------------------
057 YP_009140788.1   -----------EKV------------------------------------
058 YP_009508410.1   --------------------------------------------------
059 NP_040333.1      --------------------------------------------------
060 YP_008992013.1   KRRKSSRCFICRKKGHFAK-------------------------------
061 NP_041734.1      AYLEEDTISRVIGH------------------------------------
062 NP_056895.1      --------------------------------------------------
063 YP_009109689.1   --------------------------------------------------
064 YP_002519387.1   T-------------------------------------------------
065 YP_595725.1      LRKELREVEATLEVNK----------------------------------
066 YP_009508551.1   --------------------------------------------------
067 YP_001497148.1   --------------------------------------------------
068 NP_041186.1      --------------------------------------------------
069 NP_663784.1      --------------------------------------------------
070 YP_009508443.1   IEDMAVNVLICDG-------------------------------------
071 YP_003987465.1   TALLVEKLQEQLQ-----QALSNKTTCNEEVR------------------
072 NP_861410.1      --------------------------------------------------
073 YP_009345071.1   KQEKEEECLFIKEE------------------------------------
074 YP_009508582.1   --------------------------------------------------
075 NP_569141.1      --------------------------------------------------
076 YP_002916057.1   --------------------------------------------------
077 NP_777384.2      --------------------------------------------------
078 YP_233110.1      LKEEITALEEEVRIHKEEAEELQEENQKLKEKIIAFEKDV----------
079 NP_542258.1      --------------------------------------------------
080 YP_009130664.1   RQIETQRILS----------------------------------------
081 YP_009508566.1   --------------------------------------------------
082 YP_006607892.1   FLSSLKNPQIQMEQLSI---------------------------------
083 YP_009506270.1   LSERIKRLEQQQKY------------------------------------
084 NP_042513.1      --------------------------------------------------
085 YP_006907834.1   -----------MKPQPL---------------------------------
086 YP_004442827.1   LTEKLKGLEEASTS------------------------------------
087 YP_009408594.1   KGRESIQYLGEYVV------------------------------------
088 YP_009508561.1   --------------------------------------------------
089 NP_056907.1      --------------------------------------------------
090 YP_002308474.1   --------------------------------------------------
091 YP_007761644.1   HKSLSEQEIINEARIRAFITVIPRPIVTEPINNGQTSNQPQSPTETIEEI
092 NP_040550.1      --------------------------------------------------
093 NP_758808.1      FTGEIQIVLFNHNTAPYPVKKTYRLAQIIFEKFYTPIFIQEPFTSTQQGS
094 YP_009345075.1   YKEEIFITLYNASKEDLVFPAQSQIAQG----------------------
095 NP_068729.1      --------------------------------------------------
096 NP_047255.1      --------------------------------------------------
097 YP_006273075.1   -----------EKI------------------------------------
098 YP_009508888.1   --------------------------------------------------
099 NP_395469.1      --------------------------------------------------
100 YP_009508546.1   --------------------------------------------------
101 NP_056762.1      --------------------------------------------------
102 YP_009508556.1   --------------------------------------------------
103 NP_569153.1      -----------IIA------------------------------------
104 NP_043924.1      --------------------------------------------------
105 YP_009243641.1   --------------------------------------------------
106 YP_004300274.1   --------------------------------------------------
107 NP_056848.1      --------------------------------------------------
108 YP_006495799.1   VGALLAEGELEKLA------------------------------------
109 YP_009508409.1   --------------------------------------------------
110 NP_687035.1      --------------------------------------------------
111 NP_045937.2      --------------------------------------------------
```

```
112 NP_612577.1         --------------------------------------------------
113 NP_619548.1         GHFCLN--------------------------------------------
114 NP_056880.1         --------------------------------------------------
115 YP_009121747.1      WNEEKAKLEAQN---EYLQQENKRLREEGFQPYPP---------------
116 YP_009113237.2      ISEESSEEDAEIIA------------------------------------
117 YP_009508408.1      KEPEQLEVLICEG-------------------------------------
118 NP_955579.1         --------------------------------------------------
119 YP_009229919.1      SDEETAMAITEETS------------------------------------
120 YP_004442833.1      LRKRIAELEGASTS------------------------------------
121 NP_862833.2         --------------------------------------------------
122 NP_040563.1         --------------------------------------------------
123 YP_443922.1         --------------------------------------------------
124 NP_057933.2         --------------------------------------------------
125 YP_009116631.1      IGALLAEEGIEELA------------------------------------
126 YP_009513249.1      --------------------------------------------------
127 YP_223871.1         --------------------------------------------------
128 NP_127504.1         MRQKPRSCFTCGKIGHFSR-------------------------------
129 NP_044929.1         --------------------------------------------------
130 NP_057860.1         --------------------------------------------------
131 NP_040973.1         --------------------------------------------------
132 NP_777317.1         LQKELEQAKHQNTVLTAQAKGKTEE-------------------------
133 NP_054716.1         --------------------------------------------------
134 YP_001856242.1      --------------------------------------------------
135 NP_954565.2         --------------------------------------------------
136 NP_569150.1         LKERIKDLEQQQKD------------------------------------
137 NP_659397.1         SP------------------------------------------------
138 YP_009513211.1      --------------------------------------------------
139 YP_233107.1         LLLQLAKIDKKEIESLRS--------------------------------
140 YP_009109692.1      --------------------------------------------------
141 NP_057849.4         --------------------------------------------------
142 NP_049560.1         --------------------------------------------------
143 YP_009506267.1      IEEHDVNVLICEG-------------------------------------
144 NP_039820.1         LKEAIIVQEEDTIEER----------------------------------
145 NP_056790.1         --------------------------------------------------
146 YP_002321513.1      ATNDMDEILSVHSQHS----------------------------------
147 NP_955591.1         --------------------------------------------------
148 YP_001956722.2      --------------------------------------------------
149 NP_056902.1         --------------------------------------------------
150 NP_056886.1         --------------------------------------------------
151 NP_041261.1         --------------------------------------------------
```

```
001 YP_004442836.1      ------------QQREEKVNALEEVSVNALR-PRNNHLNIRCEVEIKEKK
002 NP_056728.1         ----MDHLLLKTQTQTEQVMNVTNP----------NSIYIKGRLYFKGYK
003 YP_009508537.1      ---------------------KPISYVSIVFSKTELKFTELEKLLTTVHK
004 YP_004732983.2      -------------------QQE-QAYTEMPRGAHNKLYHVVVTFRIPDAK
005 YP_009508407.1      --------------------------------------------------
006 NP_040939.1         ------------------------LWVNVVHNIKNLSI--PQQVIKAAQK
007 YP_003864102.1      ----------------------PLMWIHSHASPKRVLLPYYDAIADLIIL
008 YP_009002585.1      -------------------STETGREERRPVKG-NMLYNMDVEIDIPSVP
009 YP_003284237.1      --------EKVQMLTEEDEKVFSAS-GRSGP-RYNGLYNVKVGIEVEKEI
010 YP_009268869.1      ------------------------LWVNVVHNIKNLSQ--SQQIIKAAQK
011 YP_009506251.1      -------------------TTEGPSEAKKVVKKSNMLYNLEVILIIPEVG
012 NP_955611.1         ---------------------RPCLWLFS-TQPTKAFTAWLEVLTLLITK
013 YP_008567619.1      -------------------ICISKEGEKIKPISKSMLIKFQVKLEIPDVP
014 NP_149413.1         -----------AVESEEEDEVEEVTLMASAV--KDQMFRFPVVIEVPEVG
015 YP_004442824.1      ------------RPIIEDVNALEECIVSTLG-PRNNLLNMKVSFEVEERR
016 YP_001931967.1      ----------------MSKRNLTNP----------NSIYILGTFQFPGYK
017 YP_002455786.1      ---------------------WPLVWLHAPHPPT-SLCPWGHILACTVLT
018 YP_009109694.1      ---------------------RPVAYLSKKLDPVAAGWPPCLRIMAATAM
019 NP_789739.1         ------------------------LKTGKYARMRGAHTNDVKQLTEAVQK
020 NP_955619.1         ---------------------WPLVWLHAPLPHT-SQCPWGQLLASAVLL
021 YP_224289.1         --------KLALMMNEVCTEREYEQINATT-VQNNLYNIKIGLEVDGEK
022 YP_009182100.1      LADIEKLMEEVKKLQQKESTPEDKPDATPEFRHKDNSIYISAAFRFKGYE
023 NP_705927.1         ------------------------LKTGKYARMRGAHTNDVKQLTEAVQK
024 YP_009508577.1      ---------------------KPIAYASHVFTNTELKFTPLEKLLVTMHK
025 YP_004442830.1      -----------KILMAKEKL--QETVLISAR--KDPMYRFDVHVEVEGK-
026 NP_758887.1         ------------------------LKVGKYARTKNAHTNELRVLAGLVQK
027 YP_009508406.1      --------------------------------------------------
028 YP_610965.1         ----------------ALLAEAAKIGQQHQKKLINRLYNTVVKFTIPGAE
029 YP_001931961.1      ----RVKNLINNQIYSGDVTNITNP----------HSIYIKGRLKFPGYK
030 YP_241114.1         ----------------------PLRWIYLSATPTKHLLPYYELVAKIVAK
```

```
031 YP_006732334.1 --KHQCLSLQTVDNGLCLIANRTSP----------NSIYIKAKLSFKGYK
032 YP_009041481.1 -------------------AMHESILSSTVKGATRRLYNIRVTFEIPDVP
033 YP_001036293.1 -------------------REK-SERIASIKERRNGLYNLMVQLEIPGAA
034 YP_605811.1    --------VQESEEVIELVNNVEEHLVLTGQ-QKNNLLNIKITLEIKEKK
035 NP_955577.1    ---------------------RPVAYLSKKLDTVASGWPPCLRMVAAIAI
036 YP_003208050.1 ----------------FGASAAVTG-----RKVINRLYNVKILFTIPNVK
037 NP_789740.1    ------------------------LKTGKYARMRGAHTNDVKQLTEAVQK
038 YP_004442839.1 --------T-EIQKPVELVNNVEEHLVLTGK-QRNNLLNIKITIEIKEKK
039 YP_009507791.1 ------------------------LWMGRMNRQKKKAENTCDIVLRAINK
040 YP_004581513.1 -------------------AVSPQIVSSASGELNNRLYNMKVCIRIRGCP
041 NP_043933.1    --DEEQLLESVDSEERFSTLTKTNP----------NSIYIRGNFYFKGYK
042 YP_009507248.1 ---------------------KPVCYASRTLNESELNYSTIEKELLAIVW
043 YP_002117531.1 --------ESVGMLMEENEQVFSASQALQPKGQYNGLYNLQIEICCDSWT
044 YP_004347415.1 ---------------------------------MNRMTYMTVKITIPTYM
045 NP_056891.1    ----------------------PIMWIHLPASPKKVLLPYYDAIADLIIL
046 YP_009513242.1 ---------------------RPIMYVNYVFSKAEVKFTQTEKMLTTMHK
047 YP_009352866.1 -------------------INSEECYSTSVKKASKKLYNVRVTFEIPGVE
048 YP_009165750.1 ------QKVQIQNNHQQLSLNATNT----------SSIYISGKLKFQGYK
049 NP_040840.1    ------------------------LWVNVVHSIKNLSQ--AQQIIKAAQK
050 YP_004222728.1 ---------------------KPCLWLFS-TQPTKAFTAWLEVLTLLITK
051 YP_009506264.1 --------ENALFLEEEESKVFSYERLRAGP-RYNGLYNLKIALEINGVI
052 YP_567050.1    ---------------------WPLVWLHTPHPPT-SLCPWGHLLACTILT
053 YP_009508571.1 ---------------------KPIMYLNYVFSKAELKFTLLEKLLTTMHK
054 NP_056803.1    ---------------------KPIMYLNYVFSKAELKFSMLEKLLTTMHK
055 NP_955564.1    ----------------------VVEWIHLPHISPKVITPYDIFCTQLIIK
056 YP_009508411.1 -------------------KGEQVLATTSS---RNGLYNLVVELSIPGIL
057 YP_009140788.1 -------------------LIAGAEERKRLVK--NMLYNFDIEFEIPDVP
058 YP_009508410.1 --------------------------------------------------
059 NP_040333.1    ---------------------RPVAYLSKKLDPVAAGWPPCLRMVAAIAV
060 YP_008992013.1 -----------DCPNKRAKSIRLVEHLQATTDYSPQQDELEFYFSEQDEP
061 NP_041734.1    -------------------TVE-QQEVRKPVKKGNMLYNLDVVLHIPEVG
062 NP_056895.1    ---------------------FPLAYFQTPLTDN-QASPWGLLLLLGCQY
063 YP_009109689.1 ---------------------RPVAYLSKKLDPVAAGWPPCLRIMAATAM
064 YP_002519387.1 -------LLMNLNLKTSGKSTKTNP----------YSTYVTVGFYFNGYK
065 YP_595725.1    -----------VEESEEE---AEEVMMASAV--KDEMYRFPVIIEVPEVG
066 YP_009508551.1 ---------------------KPIMYLNYVFSKAELKFSMLEKLLTTMHK
067 YP_001497148.1 ---------------------RPVAYLSKKLDPVAAGWPPCLRIMAATAM
068 NP_041186.1    ----------------------PLRWIYLSATPTKHLLPYYELVAKIIAK
069 NP_663784.1    ------------------------LKVGKYAKIKNTHTNGVRLLAQVVQK
070 YP_009508443.1 -------------------KDKHVAATLPLKERRNGLYNLIVELDIPGST
071 YP_003987465.1 --------EEVAVITEEEN--LSYSAIKSPT-RYNGLYNLKAKIEVEGET
072 NP_861410.1    ------------------MKSKGNP----------NATFITVKINDV---
073 YP_009345071.1 -------------------PEEEEQRVEETLAISGKLIIIEGRIEIEGVK
074 YP_009508582.1 ---------------------KPIAYASHVFTNTESKFTPLEKLLVTMHK
075 NP_569141.1    -----------------------------------MKIYILAKIIVEGYY
076 YP_002916057.1 ---------------------------------------MRVSIRIRGCP
077 NP_777384.2    ---------------------FPLAYFQTPLTDN-QASPWGLLLLLGCQY
078 YP_233110.1    --------TQGPEEVIELVNNVEEHLVLTGQ-QKNNLLNIKITLEVKEKR
079 NP_542258.1    ---------------------WPLVWLHTPHPAT-SLRPWGQLLANAIIT
080 YP_009130664.1 ---------------------IEKNELSPPKKTKNMLYNLILEINIPDVA
081 YP_009508566.1 ---------------------RPIAYASHVFNHTEQKFTPVEKLLTTMHK
082 YP_006607892.1 --HLQNHQVQIQNNSEMNYLNSTNP----------NSIYVKGILKFSGYK
083 YP_009506270.1 -------------REEEQVNALEEVSVNALR-PRNNHLNIKCEVEVKERK
084 NP_042513.1    ---------------------MSSK----------NSSFIKVKLFNK---
085 YP_006907834.1 --KTMEHLLQKTQIQTEQVMNITNP----------NSIYIRGRLYFSGYK
086 YP_004442827.1 -----------VLMAQEGQ----ETVMLTAR--KDPMYRFDVHVEVDGI-
087 YP_009408594.1 ----------------CRASTDQKE--YHPKRMINRLFNMKVTFHIPTMD
088 YP_009508561.1 ---------------------IPIAYASHVFSTAELKFTPLEKLLVTMHR
089 NP_056907.1    ---------------------WPLVWLHTPHPPT-SLCPWGHLLACTILT
090 YP_002308474.1 ---------------------WPLVWLHTPHPAT-SLRLWGQLLANAIIT
091 YP_007761644.1 DLQIQLLELKLKKVKLQKKLKSIPP----------PPLPNPDRIETVEES
092 NP_040550.1    ---------------------RPCLWLFS-TQPTKAFTAWLEVLTLLITK
093 NP_758808.1    SNFGSTAKPLQITENIEVMSETVANQVAKSSVLP-RLYSIQAHIHIAPDI
094 YP_009345075.1 -----------IVIPLAAQDTAMVIKEESARAVNNQLYNFEVEIMIPGVK
095 NP_068729.1    --------MNTEIVQKHRVLTKGNP----------NVTFIKVSIGKR---
096 NP_047255.1    ---------------------RPVAYLSKKLDTVASGWPPCLRMVAAIAI
097 YP_006273075.1 -------------------LAAGTEEKKKFVK--NMLYNFDIEFEIPDVS
098 YP_009508888.1 ---------------------KPILYINYVFSKAEEKFSMLEKLLTTLHK
099 NP_395469.1    ----------------------MNQ----------NNTFIKITISQK---
100 YP_009508546.1 ---------------------KPIMYLNYVFSKAELKFSILEKLLTTMHK
101 NP_056762.1    ----------NKSEITEIPTTSLAMRANESNYIKTSINKTAGCYVETKIS
102 YP_009508556.1 ---------------------RPIMYVNYVFSKAEVKFTQTEKILTTMHK
103 NP_569153.1    -------------------HGDTQVTKTRPVK--NMLYNMDVRMEIPGIP
```

104 NP_043924.1 -----------------WFVPEPYEGEQHQCRPHDDTITWRVSTRAMPTG
105 YP_009243641.1 ---------------------EPCEWLYTRHRFHRSIMSYEQQIAILIQQ
106 YP_004300274.1 -----------------------------------MKIHILVKVICEGYK
107 NP_056848.1 ---------------------------------MNKITYMTIKISIPKYM
108 YP_006495799.1 -------------------AVAGESYSRMPKGGLNKLYHLNVQFVIPKAE
109 YP_009508409.1 --------------------------------------------------
110 NP_687035.1 ------------------------LKVGRYAKQRETHTNDLRTLAHLVQK
111 NP_045937.2 ---------------------RPIAFLSSKFDAIESGLPPCLKACASIHR
112 NP_612577.1 ----------------------TNP----------NSIYVKGILKFPGYQ
113 NP_619548.1 -------------KQEMLHLNVTNP----------NSIYIEGKLSFEGYK
114 NP_056880.1 ----------------------VVEWIHLPHISPKVITPYDIFCTQLIIK
115 YP_009121747.1 ------LGPHQINTLIEVAMRADGSEQPKVRKINNQLYNVEVEFDIPGIK
116 YP_009113237.2 -------------------VQKEVHHAEGSSKRRNNLYNIDVILEVIDEA
117 YP_009508408.1 -------------------KERQYIAAASLKERRNGLYNLMVEIDIPGSL
118 NP_955579.1 ---------------------RPVAYLSKKLDTVASGWPPCLRMVAAIAI
119 YP_009229919.1 ----------------LSSTTEAPS---HGRKVINRLYNMVVNINIPGVK
120 YP_004442833.1 -----------IILTAQEQY--NETVLVTQR--KDCMYRFDVFVEVEGK-
121 NP_862833.2 ----------------------PLRWIYLSATPAKHLLPYYELIAKIVAK
122 NP_040563.1 ---------------------------GGVKGLNYDHSNKIKKILRTMNE
123 YP_443922.1 ---------------------RVLMYHSSKLDHIETGQTTCSRYVAAVAK
124 NP_057933.2 ---------------------RPVAYLSKKLDPVAAGWPPCLRMVAAIAV
125 YP_009116631.1 -------------------AAVDTAYTEMPKGGLNKLYNTIVEFVIPQEK
126 YP_009513249.1 ---------------------KPISYVSIVFSKTELKFTELEKLLTTVHK
127 YP_223871.1 ---------------------RPVAYLSKRLDPVAAGWPRCLRAIAAAAL
128 NP_127504.1 -----------NCP-QNKKSIKLISEIQKYTGID-IEDDLESVFSIEDEP
129 NP_044929.1 ---------------------KPIAYNNALFTSTELKFTPTEKIMATIHK
130 NP_057860.1 ---------------------WPLVWLHAPLPHT-SQCPWGQLLASAVLL
131 NP_040973.1 ------------------------LWYGKMSRQKKKAENTCDIALRACYK
132 NP_777317.1 ------------GEALMHYADEEDCVVNLLR-PRNNLLNIKVAVEVGSTK
133 NP_054716.1 ---------------------KPIAYHNVVFSKTELKFTITEKVMTTIHK
134 YP_001856242.1 ------------------------LKTGKYARMRGAHTNDVKQLTEAVQK
135 NP_954565.2 ----------------------PIMWIHLPASPKKVLLPYYDAIADLIIL
136 NP_569150.1 -------------KEE-QVNVLEEVSINALR-PRNNHLNIKCEIEVKNKK
137 NP_659397.1 -------------NKFCNLANKTNP----------NSIYIEAKVKFKGYQ
138 YP_009513211.1 ---------------------RPVAYLSKKLDPVASGWPTCLKAIAAVAL
139 YP_233107.1 -------------QHPLPVLHLDDFAVTVVQ-PSNNLLNIKVIVEIDGKQ
140 YP_009109692.1 ---------------------RPVAYLSKKLDPVAAGWPPCLRIMAATAM
141 NP_057849.4 ------------------------LKTGKYARMRGAHTNDVKQLTEAVQK
142 NP_049560.1 ---------------------WPLVWLHAPLPHT-SQCPWGQLLASAVLL
143 YP_009506267.1 -------------------NDKQVAASIPVQDRRNGLYNLIVELEIPGCA
144 NP_039820.1 ------------CAMILEEKHTENIYSATAKAEYNGLYNVKVGIKPDNME
145 NP_056790.1 ---------------------RPVAYLSKKLDPVASGWPTCLKAVAAVAL
146 YP_002321513.1 ----------------MKHFEFDALVNIERPGQLNKLYNFGCEIQCNGGS
147 NP_955591.1 ---------------------RPVAYLSKKLDPVAAGWPPCLRMVAAIAV
148 YP_001956722.2 ---------------------RPIMYLNYVYTKAEVKFTNTEKLLTTIHK
149 NP_056902.1 ------------------------LWAGKKIMKANKGWSTVKNLMLLLQH
150 NP_056886.1 ---------------------RPCLWLFS-TQPTKAFTAWLEVLTLLITK
151 NP_041261.1 -------------------NGSPLLWLHLPASPSKVLLTYPSLLAMLIIK

001 YP_004442836.1 V----IIN-AILDTGATVCVADSRMIPAEMR--EQAKNRIIIRGVNGVTE
002 NP_056728.1 K----IILHCFVDTGASLCIASKFVIPEEHWVNAER--PIMVKIADGSSI
003 YP_009508537.1 G-----ILKALDLSMGQNIHVYSPIVSMQNIQKTP-----QTAKKALASR
004 YP_004732983.2 GQLLEFIINAIIDTGCTCCCINLTKVPDGAI--ENASIIQEVSGINSKTV
005 YP_009508407.1 --------------------------------------------------
006 NP_040939.1 L-----TQEVIIRTGKIPWILLP------------------GKEEDWRLE
007 YP_003864102.1 G-----KGHSRTYFGIEPSTIIQ------------------PYSKSQIDW
008 YP_009002585.1 R----FTVKAILDTGATSCCISMDKIPKAAI--EPNTFEVTFRGINSVQK
009 YP_003284237.1 K----YIN-AIVDTGATTCVVREARLTEKML--EESPVNVTLRGMNSISR
010 YP_009268869.1 L-----TQEVIIRIGKIPWILLP------------------GKEEDWILE
011 YP_009506251.1 KP---LIVKAILDTGATTCCININSVPKMAI--EQNTFLVQFRGINSTQS
012 NP_955611.1 L-----KASAVRTFGKEVDILLL------------------PACFRED--
013 YP_008567619.1 E----FTVNAILDTGATTCCIDERVIPQEAL--EANPYLVHFRGINSKTT
014 NP_149413.1 K----VQLTALLDTGATRSCINKPFIEDKML--QDTAFKVKISGVNSVTY
015 YP_004442824.1 I----MIN-AILDTGATVSVCSEEIVPQEFR--TNAAVKALIKGVNGTTA
016 YP_001931967.1 I----IIIHCLVDTGASMCIASRYIIPPELWKPTKK--VHKVKTANG-QV
017 YP_002455786.1 L-----IKYALQHYGQLCKSFHH------------------NMSTQALHD
018 YP_009109694.1 L-----VKDSAKLTLGQPLTVITPHTLEAIVRQPPDRWITNARLTHYQAL
019 NP_789739.1 I-----TTESIVIWGKTPKFKLP------------------IQKETWETW
020 NP_955619.1 L-----IKYTLQSYGLLCQTIHH------------------NISTQTFNQ
021 YP_224289.1 K----FIN-AILDTGAATCVVSQAKVSERFL--EPSAMKVELRGVNGTSK
022 YP_009182100.1 E----YILHCLVDTGASITLATQWAIPEEKWITIPHGNEIKITVANGETC

| | | |
|---|---|---|
| 096 | NP_047255.1 | L-----VKDAGKLTLGQPLTILTSHPVEALVRQPPNKWLSNARMTHYQAM |
| 097 | YP_006273075.1 | R----FIVRAILDTGASTCCINEGAVPKEAL--EESPYEVQLNGVNSVQK |
| 098 | YP_009508888.1 | A-----IIKAVDLAMGTEIMVYSPIVSMTKIQKTP-----LPERKALPVR |
| 099 | NP_395469.1 | ------TILAYIDTGASLCLLPEYNLPKQLWKELKK--PITIRVADKREL |
| 100 | YP_009508546.1 | A-----IIKAMDLAMGQEILVYSPIVSMTKIQKTP-----LPERKALPIR |
| 101 | NP_056762.1 | FNNENRIITALIDSGSTHNIICPTLIPASWINNTHREIIMFAVDNSKYNL |
| 102 | YP_009508556.1 | G-----IIKAMDLAMGQEILVYSPIVSMTKIQKTP-----LPERKALPVR |
| 103 | NP_569153.1 | A----FTVKAILDTGATTCCIDSRSVPKDAL--EENSFVVNFSGINSKQQ |
| 104 | NP_043924.1 | EKWWIDGSRYWDHDKGGYVTGWAALREDKKNQLGGALEGHVSAQVAELVA |
| 105 | YP_009243641.1 | G-----KQRCISLSGYDVSTIVL------------------PLTKSQFFF |
| 106 | YP_004300274.1 | S----RYYTPMIDTGAEISICKMNCLPDEYWKETRND--LIVRGFNNEGS |
| 107 | NP_056848.1 | S----RIYHGLFDTGANICICKKKVLPDELWHKTEN---LVLRGFNDEKH |
| 108 | YP_006495799.1 | GGHVSFTVAAIIDTGCTCCCINGAKVPEEAQ--EEASYAMTFAGVNSRGT |
| 109 | YP_009508409.1 | -------------------------------------------------- |
| 110 | NP_687035.1 | I-----GKEALTIWGRLPRVQLP------------------VDKKTWDMW |
| 111 | NP_045937.2 | S-----ITQADSFILGAPLIIYTTHAICTLLQRDRSQLVTASRFSKWEAD |
| 112 | NP_612577.1 | TN---LILHCYVDTGSSLCMASKYVIPEEYWQTAEK--PLNIKIANGKII |
| 113 | NP_619548.1 | S----FNIHCYVDTGASLCIASRYIIPEELWENSPK--DIQVKIANQELI |
| 114 | NP_056880.1 | G-----KHRSKELFSKDPDYIVV------------------PYTKVQFDL |
| 115 | YP_009121747.1 | L----FIVKAIIDTGATSCCINKVIVPREAL--EEMNKTVFFNGLNSRQP |
| 116 | YP_009113237.2 | P----VTVKAIIDTGATVCCIDERTVPAQIL--EENTYRVEFSGVNSKSI |
| 117 | YP_009508408.1 | P----INVNAILDTGATTCCINEKGLPKEIL--EDNTYEVKFVGANSEMT |
| 118 | NP_955579.1 | L-----VKDAGKLTLGQPLTILTSHPVEALVRQPPNKWLSNARMTHYQAM |
| 119 | YP_009229919.1 | P----FSVRAILDTGATTCCVDKGAVPAEAL--EKSPYAVHFSGINSRQL |
| 120 | YP_004442833.1 | K----EKLKALLDTGATKSCISERYIPKEFL--TDSKFTVNIAGVNSMTR |
| 121 | NP_862833.2 | G-----KYEAIQYFGLEPPFICV------------------PYALEQQDW |
| 122 | NP_040563.1 | L-----NRTVVIMTGREASFLLP------------------GSSEDWEAA |
| 123 | YP_443922.1 | A-----IEKTAHLVMCHKMEIHTHHGVAAYLTAKEFTFSADR-----KNK |
| 124 | NP_057933.2 | L-----TKDAGKLTMGQPLVILAPHAVEALVKQPPDRWLSNARMTHYQAL |
| 125 | YP_009116631.1 | GAPTKFIVRAVIDTGCTCTCINSKKVPKEAL--EEAKYQMNFAGVNSTGE |
| 126 | YP_009513249.1 | G-----ILKALDLSMGQNIHVYSPIVSMQNIQKTP-----QTAKKALASR |
| 127 | YP_223871.1 | L-----TREASKLTFGQDIEITSSHNLESLLRSPPDKWLTNARITQYQVL |
| 128 | NP_127504.1 | SEDTLFSLEFYEEYAGEQYQITSYEAPKTENPPLPKIHTIVEIPQTEVKI |
| 129 | NP_044929.1 | G-----ILKALDLSLGKEIHVYSAIASMTKLQKTP-----LSERKALSIR |
| 130 | NP_057860.1 | L-----IKYTLQSYGLLCQTIHH------------------NISTQTFNQ |
| 131 | NP_040973.1 | I-----KEESIIRIGKEPRYEIP------------------TSREAWESN |
| 132 | NP_777317.1 | F----NIN-AILDTGATVCVCSEEMIPKEIR--EPSKTNTVIRGVNGANK |
| 133 | NP_054716.1 | A-----ILKAFDLAMGQPIWVYSPIHSMTRIQKTP-----LTERKALSIR |
| 134 | YP_001856242.1 | I-----TTESIVIWGKTPKFKLP------------------IQKETWETW |
| 135 | NP_954565.2 | G-----KDHSKKYFGIEPSTIIQ------------------PYSKSQIDW |
| 136 | NP_569150.1 | V----VIN-AILDTGATVCVADERMIPSGMK--EQAKNKIIIRGVNGVTE |
| 137 | NP_659397.1 | T----LSLHCYVDTGASLCLASKHVIPDDFWENAPR--SIPVKIANQSSI |
| 138 | YP_009513211.1 | L-----IKDADKLTLGQNVLVIAPHNLESIVRQPPDRWMTNARMTHYQSL |
| 139 | YP_233107.1 | L----PIN-AILDTGAAICVCNGEVIPQEYR--RSSLTDSLIQGVNGATR |
| 140 | YP_009109692.1 | L-----VKDSAKLTLGQPLTVITPHTLEAIVRQPPDRWITNARLTHYQAL |
| 141 | NP_057849.4 | I-----TTESIVIWGKTPKFKLP------------------IQKETWETW |
| 142 | NP_049560.1 | L-----IKYTLQSYGLLCQTIHH------------------NISTQTFDQ |
| 143 | YP_009506267.1 | P----LKVNAILDTGATTCCINSEGLPKEMM--EDNTFTVKFTGANSVMK |
| 144 | NP_039820.1 | P----YYINAIVDTGATACLIQISAIPENYY--EDAKVTVNFRSVLGIGT |
| 145 | NP_056790.1 | L-----IKDADKLTLGQNVTVIASHSLESIVRQPPDRWMTNARMTHYQSL |
| 146 | YP_002321513.1 | P----FTARAILDTGASICCICTSITPRNCL--VET-KEASFNGITSSSK |
| 147 | NP_955591.1 | L-----TKDAGKLTMGQPLVILAPHAVEALVKQPPDRWLSNARMTHYQAL |
| 148 | YP_001956722.2 | G-----IIKALDLGMGQEILVYSPIVSMTKIQKTP-----LPERKALPIR |
| 149 | NP_056902.1 | V-----ATESITRVGKCPTFKVP------------------FTKEQVMWE |
| 150 | NP_056886.1 | L-----KASAVRTFGKEVDILLL------------------PACFRED-- |
| 151 | NP_041261.1 | G-----KYTGRQLFGRDPHSIII------------------PYTQDQLTW |

| | | |
|---|---|---|
| 001 | YP_004442836.1 | VNEVTAI-GRLWVGKQ-----WFTLPQTFIMP---TLGDGVHMIIGMNFI |
| 002 | NP_056728.1 | TISKVCKDIDLIIAGE-----IFRIPTVYQQ------ESGIDFIIGNNFC |
| 003 | YP_009508537.1 | WLSWLSYLEDPRIRFFYDPQMPALKDL--PAVDIGENN-KKHP-SNFQHV |
| 004 | YP_004732983.2 | VTKKLRQ-GKMILAGN-----DFYIPYVSAFE---MNMPGIDMLIGCNFI |
| 005 | YP_009508407.1 | -------------------------------------------------- |
| 006 | NP_040939.1 | LQLG-NITWMP--------------------------KFWSCYRGHTRWR |
| 007 | YP_003864102.1 | LMQNTETWPIACASYAGKIDNHYPPDK----------LLQFCRLHTFIFP |
| 008 | YP_009002585.1 | SNKKLKY-GQMRIGGH-----MFRIPYTYCFES-LTLGDDVQYVLGCNFI |
| 009 | YP_003284237.1 | ASKVVKQ-GKIWIGEQ-----YFRTPRTFALDI--SLSGGIDMILGCNFI |
| 010 | YP_009268869.1 | LQIG-NITWMP--------------------------SFWSCYRGSIRWK |
| 011 | YP_009506251.1 | VDKKLRY-GRMTISGH-----QFRFPYCYAFP--LSLGDGIEMILGCNFI |
| 012 | NP_955611.1 | -LPLPEGILLALKGFAGKIRSSDTPS-----------IFDIARPLHVSLK |
| 013 | YP_008567619.1 | ANKKLKY-GRMIIGDQ-----QFKVPYTYAFP--MTISDDIQMILGCNFI |
| 014 | NP_149413.1 | LDKQVKDGAKLWAGSS-----WFRLPLTYVGNM--HMGEKTQMLIGCNFL |

```
015 YP_004442824.1 VTEVLKE-GKLWVGNQ-----YFRMPRTFVMP---KLSEGLHLIIGLNFI
016 YP_001931967.1 TIDKVCLSQVITVAGK-----EFLIPTIYQL------NTGIDMVLGNNFL
017 YP_002455786.1 FVKNSSHPSVAILIHHMHRFCDLGRQPPGPWRTLLQLPALLREPQLLRPA
018 YP_009109694.1 LLDTDRVQFGPPVTLNPATLLPVPENQPSPHDCRQVLAETHGTREDLKDQ
019 NP_789739.1    WTEYWQATWIP-------------------------EWEFVNTPPLVKLW
020 NP_955619.1    FIQTSDHPSVPILLHHSHRFKNLGAQTGELWNTFLKTTAPLAPVKALMPV
021 YP_224289.1    VNKVLKE-GKIWIGNQ-----YFRLPRTYALEI--ELSGGIDMIIGTNFL
022 YP_009182100.1 KIDKYATNVPIYISGI-----KFTIEKVLQFN-----KQGCDFLIGNDFF
023 NP_705927.1    WTEYWQATWIP-------------------------EWEFVNTPPLVKLW
024 YP_009508577.1 WITWLSYLEDPRITFYYDKTLPDLKNV--PETVTDKKP-KMXPIIEYAAV
025 YP_004442830.1 VTKQLKE-AKLWVGGT-----FFTLPITYVGKL--DLGERTQMVIGCNFI
026 NP_758887.1    WPEYWQATWIP-------------------------EWEFVSTPHLIGLW
027 YP_009508406.1 --------------------------------------------------
028 YP_610965.1    SRQRLKE-GRMTIGDH-----DFPIPLVYCFP-LDSSRDGVQMLIGCNFI
029 YP_001931961.1 EINKVCENLNMYIAGE-----CFHIPRVFMQ------ESGIDFILGNNFC
030 YP_241114.1    LFQFSDNWAIAFANYPGRIIHHYPPDK----------LLQFASSHAFIFP
031 YP_006732334.1 QINKVCKNLILQIVGE-----EFVIPTIYQQ------ETGIDILLGNNFC
032 YP_009041481.1 AKKRIKA-GKMFLGDN-----YFKIPYIYCFQ---MQVKNIDMLIGCNFI
033 YP_001036293.1 ARKKLKG-GIMKIGDN-----QFRIPYTFAFP--LKLGGGEQLIIGCNFI
034 YP_605811.1    VKEILEE-GKIWIGNQ-----WFRIPRTYIMP---QLSEGLHFIIGMNFI
035 NP_955577.1    LLDAERVHFGPTVSLNPATLLPLPSGG-NHHDCLQILAETHGTRPDLTDQ
036 YP_003208050.1 ARHKIKG-GQISMGES-----NFRLPFIYAFK-MDS-KDEIQMLLGCNFL
037 NP_789740.1    WTEYWQATWIP-------------------------EWEFVNTPPLVKLW
038 YP_004442839.1 VQEILEE-GRIWIGNQ-----WFRIPRTYVMP---QLSEGLHFIIGMNFI
039 YP_009507791.1 LITS-IYLKNPPP----------------------EVQFIHASLMMARML
040 YP_004581513.1 TRKKLKE-GYMLIGEH-----MFPLPFVYALN-PMRIGRGIQFIIGCNFI
041 NP_043933.1    TFNKVAELMQVQIADE-----TFIIPTLYQAT-----TKG-DITLGNNFC
042 YP_009507248.1 WRLRLSEYDFSVVYKK---------------------GKSNTNADALSRV
043 YP_002117531.1 GTKSLRP-GLMKIQGG-----TYRIPKTYVIDI--QLSEGIDMILGCNFI
044 YP_004347415.1 VIKYRADNIKIMIAKK-----EFIIPYIYAMD-----NINADIIIGATFY
045 NP_056891.1    LMQNTEMWPIACASFVGILDNHYPPNK----------LIQFCKLHTFVFP
046 YP_009513242.1 WITWMTYLEDPRIQFHYDKTLPELQQT--PSVTEDVIA-KTKHPSEFAMV
047 YP_009352866.1 AKKRLKA-GKIFLGDN-----YFKIPYIYCFQ---MQVKGIDMLIGCNFI
048 YP_009165750.1 TINKVCKNIDIMIADT-----IFHIPTIYQQ------EIGIDLLLGNNFL
049 NP_040840.1    LQMG-NINWMP--------------------------SFWSCYKGSVRWK
050 YP_004222728.1 -LPLPEGILLALKGFAGKIRSSDTPS-----------IFDIARPLHVSLK
051 YP_009506264.1 TNQVLKG-GKIWIGEQ-----YYRIPRTMALEV--ALSEGIDMIIGCNFI
052 YP_567050.1    FVKNSPHPSVGILIHHMGRFHNLGSQPSGPWRTLLHLPALLQEPRLLRPL
053 YP_009508571.1 WITWMTYLEDPRIQFHYDKTLPELKNI--PDVLTENSSKIMIHPSQYNSV
054 NP_056803.1    WITWMTYLEDPRIQFHYDKTLPELKHI--PDVYTSSIP-PLKHPSQYEGV
055 NP_955564.1    LLQEKEDWPISLLGFLGEVHFHLPKDP----------LLTFTLQTAIIFP
056 YP_009508411.1 ANKKLKG-GMMKIGDH-----QFRIPYTFAFP--LRLGGGEQLILGCNFI
057 YP_009140788.1 TKKKLKY-GRMMIGMN-----SFRIPFTYSLP--LVIGDNIQMIVGCNFI
058 YP_009508410.1 --------------------------------------------------
059 NP_040333.1    LLDTDRVQFGPIVALNPATLLPLPEEG-LQHDCLDILAEAHGTRPDLTDQ
060 YP_008992013.1 LPSKFQRPIPAIGLIDTGAQRSMINPHILPPDSWTQSEEHFKAVNGKLFT
061 NP_041734.1    VDKKLKY-GRMTISNH-----QFRIPYCYAFP--LSLGDGIEMILGCNFI
062 NP_056895.1    WIQSSEDPRVQELLQLWPQISSQGIQPPGPWKTLITRAEVFLTP------
063 YP_009109689.1 LLDTDRVQFGPPVTLNPATLLPVPENQPSPHDCRQVLAETHGTREDLKDQ
064 YP_002519387.1 EIRYVTRNLKVSMTDTNGNKHTFIIPTVYQQ------NTGMDFIFGQNFL
065 YP_595725.1    LDRQVKDGAKLWAGEN-----WFRLPITYVGPM--YMGEKTQMLIGCNFM
066 YP_009508551.1 WITWMTYLEDPRIQFHYDKTLPELKHI--PDVYTSSQS-PVKHPSQYEGV
067 YP_001497148.1 LLDTDRVQFGPPVTLNPATLLPAPKDQQSAHDCRQVLAETHGTREDLKDQ
068 NP_041186.1    LFQFSDNWSIAFANYPGQITHHYPSDK----------LLQFASSHAFIFP
069 NP_663784.1    WDNYWQVTWIP-------------------------EWDFVSTPPLVRLT
070 YP_009508443.1 AIKKLKN-GTMRIGDH-----QFRIPYTYAFP--LKLGGGEQLILGCNFI
071 YP_003987465.1 GRKVLKG-GKLWLGEQ-----FFRLPRTLVADM--VLTEGVEMIVGCNFL
072 NP_861410.1    EIWHRAEMVEIWIRNY-----KFVAATVCQK------SSGMDFVIGNNFL
073 YP_009345071.1 VNQKLKQ-GKLIIGSQ-----EFRLPYTYVLPMNKNLVTDMQMIIGCNFI
074 YP_009508582.1 WITWLSYLEDPRIIFHYDKTLPDLKNV--PETITEKQP-KILPIIEYAAV
075 NP_569141.1    MINYKARNVKIQIWDK-----ILTIEEIYNFE-----FTTKDMLLGMPFL
076 YP_002916057.1 TRKKLKE-GYMIIADH-----TFPLPFVYALN-PMRIGRGIQFIIGCNFI
077 NP_777384.2    WIQSSEDPRVQELLQLWPQISSQGIQPPGPWKTLITRAEVFLTP------
078 YP_233110.1    VKEILEE-GKIWIGNQ-----WFRIPRTYIMP---QLSEGLHFIIGMNFI
079 NP_542258.1    YLHTSDQSSVAILLQHSHRFHNLGAQPSGPWRSLLQVPQIFQNIDVLRPP
080 YP_009130664.1 TNKKLRY-GKMIIGDN-----AFRIPYTYVFP--MQLGDDIQMILGCNFI
081 YP_009508566.1 WLSWLSYIEDPRFLFIYDKTLPDLKEM--PPTQTDDYN-PMLPLHQYLAV
082 YP_006607892.1 TLTKVCKNVKIMLAGE-----LFHIPTIYQQ------EVEIDLILGNNIL
083 YP_009506270.1 VNEVTSA-GKLWVGKQ-----WFYLPQTYIMP---TLADGVHMIIGMNFI
084 NP_042513.1    HIWYKAVDLVLLLEGK-----RFPLPSVYQQ------DAGLPLILGNNFL
085 YP_006907834.1 TLDKICRNLDIIIAGE-----RFHIDVVFQQ------ESGIDFIIGNNFC
086 YP_004442827.1 VTKQLKE-AKLWVKDT-----FFTLPITYVGNL--DLGRNTQMIIGCNFI
087 YP_009408594.1 ANWKLKG-GQMQIGEN-----KFKIPFTYSFN-MNL-GDGIQMIIGCNFV
```

```
088 YP_009508561.1     WVTWISYLEDPRITFYYDKTLPDLKHV--PASTDNNII-TLLPITEYEAV
089 NP_056907.1        FVKNSPHPSVGILIHHMGRFHNLGSQPSGPWRTLLHLPALLQEPRLLRPL
090 YP_002308474.1     YLHTSDQSSVAILLQHSHRFHNLGAQPSGPWKGLLQVPQIFQNVATLSPP
091 YP_007761644.1     AVEIIQSLSTLRYIDLEFQLEGYSEYRIRSFY-----DTGASLMLARYHI
092 NP_040550.1        -LPLEEGILLALRGFAGKIRSSDTPS-----------IFDIARPLHVSLK
093 NP_758808.1        IQHRIKR-GTLEIASN-----KYALPLCYIIE-LND-KDDFSMILGCNFF
094 YP_009345075.1     VNEKIKD-GQMRIAGN-----SFRIPFTYCFP-LGA-HDNVDMLIGCNFI
095 NP_068729.1        YIREAISNVFLKIENK-----EFLIPIIYLH------DSGLDLIIGNNFL
096 NP_047255.1        LLDAERVHFGPTVSLNPATLLPLPSGG-NHHDCLQILAETHGTRPDLTDQ
097 YP_006273075.1     TKKKIKY-GRMMIGMN-----TFRIPFTYSLP--LVIGDNIQMIVGCNFI
098 YP_009508888.1     WITWMTYLEDPRITFHYDKTLPELKDV--PSVYQNDIP-IVPHPSQYSMV
099 NP_395469.1        QINKVALMITILIEKR-----KFLVPTIYQF------DSGVPMIIGNNFL
100 YP_009508546.1     WITWMTYLEDPRIQFHYDKTLPELKHI--PDVYTSEQP-PSKHPSQYEGV
101 NP_056762.1        NQELIDDIKLQFQEVDETFGIKYKLGQTYVAP-----KPTKTFIIGHRFL
102 YP_009508556.1     WITWMTYLEDPRIQFYYDKTLPELQQI--PSVTEDVVA-KTKHPSEFAMV
103 NP_569153.1        VKQKIKT-GKMFINEH-----YFRIPYCYSFE--MQIGDGIQLILGCNFI
104 NP_043924.1        LREAIRLQRPLTLYTDSTYVLGICTKYLAVWKRRGMVNADGSQISNQNIL
105 YP_009243641.1     LLQRSEALQIALAQFVGNIEYTLPKGK----------LWDFFKRHNFVIN
106 YP_004300274.1     VIDRKASNISIQIWDK-----IVKIDHIYQFE-----IQGKDIILGMNFI
107 NP_056848.1        VAEYRADNITIMIAKE-----KFIIPYIYAMD-----EMSPDIIIGATFY
108 YP_006495799.1     TRKKMRA-GKMVISGN-----DFYTPYISVFD---MDLPDIDMLIGCNFI
109 YP_009508409.1     --------------------------------------------------
110 NP_687035.1        WQDYWQVSWIP-------------------------EWEFVSTPLLVKLW
111 NP_045937.2        LLRPELTFVACSAVSPAHLYMQSCENNIPPHDCVLLTHTISRPRPDLSDL
112 NP_612577.1        QLTKVCSKLPIRLGGE-----RFLIPTLFQQ------ESGIDLLLGNNFC
113 NP_619548.1        KITKVCKNLKVKFAGK-----SFEIPTVYQQ------ETGIDFLIGNNFC
114 NP_056880.1        LLQEKEDWPISLLGFLGEVHFHLPKDP----------LLTFTLQTAIIFP
115 YP_009121747.1     SSQKIKA-GNFRIGGN-----KFRIPLIYAMD-MMT-GDGIDMLIGTNFT
116 YP_009113237.2     ASKKIKY-GRMKIGEN-----TFRIPYTYAFQ--MNVGKEIQFIIGCNFI
117 YP_009508408.1     ATKKIKA-GNMKIGDH-----TFRIPYTFAFP--LELGGGEQLILGCNFI
118 NP_955579.1        LLDAERVHFGPTVSLNPATLLPLPSGG-NHHDCLQILAETHGTRPDLTDQ
119 YP_009229919.1     AEYKEKQ-GQMEIGEN-----RFRIPFTYSFP-MGM-KDGIEMLLGCNFI
120 YP_004442833.1     VTKQIKE-AKLWIKET-----FFSLPITYVGNL--DLGTNTQMIIGCNFI
121 NP_862833.2        LFRYSDNWAIAFANYPGRIIHHYPSDK----------LLQFASSHAFIFP
122 NP_040563.1        LQKEESLTQIFP----------------------------VKFYRHSCRW
123 YP_443922.1        IQNKCTQSHITFVNTDKNMADALNADEGSPHVCAERAAQELKLRPDLGND
124 NP_057933.2        LLDTDRVQFGPVVALNPATLLPLPEEG-LQHNCLDILAEAHGTRPDLTDQ
125 YP_009116631.1     TKLKMKN-GKMIVSGS-----DFYTPYIAAFP---MELPDVDMLIGCNFL
126 YP_009513249.1     WLSWISYLEDPRIRFFYDPQMPALKDL--PAVDTGKDN-KKHP-SNFQHI
127 YP_223871.1        LLDPERVRFKQTAALNPATLLPETDDTLPIHHCLDTLDSLTSTRPDLTDQ
128 NP_127504.1        YTSKWDKPISVIAFYDTGAAYSIMDPAILPSEYWIPHFRHFGTADDGILT
129 NP_044929.1        WLKWQTYFEDPRIKFHHDATLPDLQNLPVPQQDTGKEM-TILPLLHYEAI
130 NP_057860.1        FIQTSDHPSVPILLHHSHRFKNLGAQTGELWNTFLKTTAPLAPVKALMPV
131 NP_040973.1        LINS-PYLKAPPP----------------------EVEYIHAALNIKRAL
132 NP_777317.1        VTEVIKD-GRLWVGDQ-----YFRLPRTFVMP---KLSDGLHIILGMNFI
133 NP_054716.1        WLKWQTYFEDPRLIFHYDDTLPDLQNL--PQTTLGNEV-DILPLSEYEVV
134 YP_001856242.1     WTEYWQATWIP-------------------------EWEFVNTPPLVKLW
135 NP_954565.2        LMQNTEMWPIACASFVGILDNHYPPNK----------LIQFCKLHTFVFP
136 NP_569150.1        VNEVTSA-GKLWVGKQ-----WFYLPQTFIMP---SLADGVHMIIGMNFI
137 NP_659397.1        TLNKVCRGLKINIAGN-----EFFIPTIYQQ------ETGIDILLGNNFC
138 YP_009513211.1     LLN-ERVSFAPPAILNPATLLPVESDDTPIHICSEILAEETGTRPDLRDQ
139 YP_233107.1        VNEIIKD-GKLWVGDQ-----YFRIPRTYIMP---TMHKGLEFIIGMNFI
140 YP_009109692.1     LLDTDRVQFGPPVTLNPATLLPVPENQPSPHDCRQVLAETHGTREDLKDQ
141 NP_057849.4        WTEYWQATWIP-------------------------EWEFVNTPPLVKLW
142 NP_049560.1        FIQTSDHPSVPILYHHSHRFKNLGAQTGELWNTFLKTAAPLAPVKALMPM
143 YP_009506267.1     ADKKIKY-GMMKIEGH-----VFRIPYTFAFP--LRLGGGEQLVIGCNFI
144 NP_039820.1        STQMIKA-GRILIGEQ-----YFRMPVTYVMNM--GLSPGIQMIIGCSFI
145 NP_056790.1        LLN-ERVSFAPPAVLNPATLLPVESEATPVHRCSEILAEETGTRRDLEDQ
146 YP_002321513.1     CNQRIAR-GTFKIGDH-----MFQIPLVYAFQ--MNPRDDVKFIIGCNFI
147 NP_955591.1        LLDTDRVQFGPVVALNPATLLPLPEEG-LQHNCLDILAEAHGTRPDLTDQ
148 YP_001956722.2     WITWMSYLEDPRIQFHYDKTLPELQQV--PTVTDDIIA-KIKHPSEFSMV
149 NP_056902.1        MQKGWYYSWLP--------------------------EIVYTHQVVHDDW
150 NP_056886.1        -LPLEEGILLALKGFAGKIRSSDTPS-----------IFDIARPLHVSLK
151 NP_041261.1        LLQTSDEWAIALSSFTGDIDNHYPSDP----------VIQFAKLHQFIFP
```

```
001 YP_004442836.1     RTVG--LRIENGEVTIYKIMTTVPAPPIAHDL------------------
002 NP_056728.1        QLYEPFIQFTDRVIFTK---NKSYPVHIAKLT------------------
003 YP_009508537.1     FYTDGSAITSPTKEGHLNAGMGIVYFMNKDG-------------------
004 YP_004732983.2     RAMKGGIRLEGTEVTFYKTITRIQTTLEPQ--------------------
005 YP_009508407.1     --------------------------------------------------
006 NP_040939.1        KRNIIEEVVEGPTYYTDGGK-KNKVGSLG-FI------------------
```

```
007 YP_003864102.1   KIISKSPLDNALLVFTDGSSTG-IAAYTFAD-------------------
008 YP_009002585.1   RNMHGGVRLEGTEVTFYKNITTIQTRLMAPIVE-----------------
009 YP_003284237.1   RSLEGGVRIEGNDITFYKLVTSVRTQREAHQV------------------
010 YP_009268869.1   KRNVITEVVEGPTYYTDGGK-KNGKGSLG-FI------------------
011 YP_009506251.1   RGMYGGLRIEGSTITFYKNVTTIQTKLAAGMVGGTTSFYPGEGGGSHFEF
012 NP_955611.1      VRVTDHPVPGPTVFTDASSSTHKGVVVWREGP------------------
013 YP_008567619.1   RAMHGGVRIEGNEVTFYKYVTKIMAQQGN-AMVRT---------------
014 NP_149413.1      QSLAGGVRLEGRTVTFYKYISSIQAN------------------------
015 YP_004442824.1   HAMEGGVRIEKGEVTFYKIVTTTPTSPIVHDI------------------
016 YP_001931967.1   NLYGPFIQFTNCIILHLRD-NPKTSVRVTKVT------------------
017 YP_002455786.1   FSLSPVVIDQAPCLFSDGSPQKAAYVIWDKVI------------------
018 YP_009109694.1   ELPDADHTWYTDGSSYLDSGTRRAGAAVVD--------------------
019 NP_789739.1      YQLEKEPIVGAETF----------------
020 NP_955619.1      FTLSPVIINTAPCLFSDGSTSQAAYILWDKHI------------------
021 YP_224289.1      RAMSGGVRIEGETVTFYKIVTTTDAPKVPHQI------------------
022 YP_009182100.1   MKYAPYTQYPDRISIAK----NGISYITNVVERPVSVAY-----------
023 NP_705927.1      YQLEKEPIVGAETFYVDGAANRETKLGKAGYV------------------
024 YP_009508577.1   FYTDGSAIRSPDKNKSHSSGMGIVHAVFKP--------------------
025 YP_004442830.1   QSLRGGLRLEGRTVTFYKLTTHVQAD------------------------
026 NP_758887.1      YNLLREPVPGEDVYYVDGAANRNSKEGKAGYV------------------
027 YP_009508406.1   --------------------------------------------------
028 YP_610965.1      RSKQGGVRFEGTTVTFYKQLTTVQTEIEAIKS------------------
029 YP_001931961.1   QNYEPLKQYTDRIILTL---ENR-EIIIGKIR------------------
030 YP_241114.1      KIVRRQPIPNATLIFTDGSSNG-TAALIINH-------------------
031 YP_006732334.1   QLYNPFIQWVDRIAFHK---NEE-IILVPKVR------------------
032 YP_009041481.1   KSMGGGLRIEGPTITFYKNVTSIQTQAEELGVISMEKGE-----------
033 YP_001036293.1   RAMNGGVRIEGDTVTFYKNVTTIQTQQQVPKLLAL---------------
034 YP_605811.1      RAMEGGIRIEQGMVTFYKMVTQAQAPPMVHDI------------------
035 NP_955577.1      PLPDADLTWYTDGSSFIRNGEREAGAAVTT--------------------
036 YP_003208050.1   RSMAGGVRLEGTTITFYKQITTINTTLAVESAKKA---------------
037 NP_789740.1      YQLEKEPIVGAETFYVDGAANRETKLGKAGYV------------------
038 YP_004442839.1   RAMEGGIRIEQGIVTFYKMITQAQAPPITHDI------------------
039 YP_009507791.1   TMLTSEPIEGIETWFIDGGR-RLGKKAISAYW------------------
040 YP_004581513.1   RRMKGGLRIEGPTVTFYRNVSTIETQEKSTVAAT----------------
041 NP_043933.1      RLYEPFVQYKDMITFHK----DGRAVSTKKVT------------------
042 YP_009507248.1   EIHTTEIDEIDSVIENIKELSSMINNPSETSR------------------
043 YP_002117531.1   RAMEGGLRIEGDTLSFYKLLTTVQTSRHAQQI------------------
044 YP_004347415.1   NKYSPITIDMEKGIIKFTNKGEVYPNYLVKYPRKKV--------------
045 NP_056891.1      QIISKTPLNNALLVFTDGSSTG-MAAYTLTD-------------------
046 YP_009513242.1   FYTDGSAIKHPDINKSHSAGMGIAQVQFQP--------------------
047 YP_009352866.1   KSMGGGLRIEGTTVTFYKNVTTVQTQAEDLGAPNAGDQD-----------
048 YP_009165750.1   LLYGPFTQYTDRIILFK---ENQ-PVIIGKVS------------------
049 NP_040840.1      KRNVIAEVVPGPTYYTDGGK-KNGRGSLG-YI------------------
050 YP_004222728.1   VRVTDHPVPGPTVFTDASSSTHKGVVVWREGP------------------
051 YP_009506264.1   RSLEGGVRIEGDKITFYKLTTHIDTSKEAFTV------------------
052 YP_567050.1      FTLSPVVLTTAPCLFSDGSSQKAAYVLWDQTI------------------
053 YP_009508571.1   FYTDGSAIRSPDPTKSHNAGMGIVQVKFSP--------------------
054 NP_056803.1      FCTDGSAIKSPDPTKSNNAGMGIVHAIYNP--------------------
055 NP_955564.1      HMTSTTPLEKGIVIFTDGSANGRSVTYIQGR-------------------
056 YP_009508411.1   RAMQGGLRIEGDEVTFYKNVTTIKTQQMVSKVLAL---------------
057 YP_009140788.1   RAMYGGVRIEGNEVTFYKNLTKISTSPEVSVNV-----------------
058 YP_009508410.1   --------------------------------------------------
059 NP_040333.1      PLPDADHTWYTDGSSFLQEGQRKAGAAVTT--------------------
060 YP_008992013.1   TSLITKKPIGIQIFPNCVIWTKVIGSTLPNKDILLG--------------
061 NP_041734.1      RGMYGGLRIEGHTITFYKNVTTIQTRLAAVMVGGTTASELGGGEESKSDS
062 NP_056895.1      -QFSPEPIPAALCLFSDGATGRGAYCLWKDHL------------------
063 YP_009109689.1   ELPDADHTWYTDGSSYLDSGTRRAGAAVVD--------------------
064 YP_002519387.1   KLYRPFTQDLYEIYLTP-------KLLKFKIQQ-----------------
065 YP_595725.1      QSLAGGVRLEGRTVTFYKYIASIKAN------------------------
066 YP_009508551.1   FYTDGSAIKSPDPTKSNNAGMGIVHATYKP--------------------
067 YP_001497148.1   ELPDADHSWYTDGSSYIDSGTRRAGAAVVD--------------------
068 NP_041186.1      KIVRRQPIPEATLIFTDGSSNG-TAALIINH-------------------
069 NP_663784.1      FNLVGDPIPGAETFYTDGSCNRQSKEGKAGYV------------------
070 YP_009508443.1   RSMNGGVRIEGNEVTFYKNVTTIQTQQEVPKVLAL---------------
071 YP_003987465.1   RSLEGGLRIEGEVVTFYKLVTNVQSCRSTHAFG-----------------
072 NP_861410.1      RLYQPFIQGLNYIKLRAP------LDKDINQPS-----------------
073 YP_009345071.1   RSLHGGVRIEGNEITFYKYLTKLESINLIEEDN-----------------
074 YP_009508582.1   FYTDGSAIRSPDKNKSHSSGMGIVQAIFKP--------------------
075 NP_569141.1      EKLYPHIIT--KTHWWFTTPCKNKVGAKRVNNKQRK--------------
076 YP_002916057.1   RRMKGGLRIEGPTVTFYRNVSTIETQEKSTVAAX----------------
077 NP_777384.2      -QFSPEPIPAALCLFSDGATGRGAYCLWKDHL------------------
078 YP_233110.1      RAMEGGIRIEQGTVTFYKMVTQAQAPPMVHDI------------------
079 NP_542258.1      FIISPVVIDHAPCLFSDGATSKAAFILWDKQV------------------
```

```
080 YP_009130664.1     RSMQGGVRIEGDTVTFYKNVTQIQTQQTVPIIAAI---------------
081 YP_009508566.1     FYTDGSSIKSPDPTKTHSSGMGIVQAIYEP--------------------
082 YP_006607892.1     QLHEPFVQFTDRIIIHK---DGE-PVTIGKVF------------------
083 YP_009506270.1     RTVG--LRIENGEVTIYKIMTTVQAPPIVHEL------------------
084 NP_042513.1        KLYNPFIQTLETISLRCP------QLE--KQPS-----------------
085 YP_006907834.1     QEYSPFIQFTGHIVFTM---KDQYQVPITKLR------------------
086 YP_004442827.1     QSLRGAVRLEGRSVTFYKLISTVEAD------------------------
087 YP_009408594.1     RAMQGGIRIEGNQVTFYKQLTTIETTMSAGPCIQE---------------
088 YP_009508561.1     FYTDGSAIKSPKTEQTHSAGMGIVMVVYTP--------------------
089 NP_056907.1        FTLSPVVLTTAPCLFSDGSSQKAAYVLWDQTI------------------
090 YP_002308474.1     FTISPVVINHAPCLFSDGSNSQAAFTIWDKKI------------------
091 YP_007761644.1     FPPELWIKQTPKLVSYANQSTGSIMYKAVQVPFKLG--------------
092 NP_040550.1        VRVTDHPVPGPTAFTDASSSTHKGVVVWREGP------------------
093 NP_758808.1        KHMGGGMRFEGPHVTFYKGITTLST---------S---------------
094 YP_009345075.1     KANYGGMRIEGSKITFYKNVGSIETSPVVSSAHFA---------------
095 NP_068729.1        KLYQPFIQRLETIELRWK------NLNNPKES------------------
096 NP_047255.1        PLPDADLTWYTDGSSFIRNGEREAGAAVTT--------------------
097 YP_006273075.1     RAMYGGVRIEGNEVTFYKNLTRISTSPEVSVNS-----------------
098 YP_009508888.1     FYTDGSAIKNPNPTKTHSAGMGVVQGKFNP--------------------
099 NP_395469.1        RLYYPFCQYLSYITLRCP------KMINQKQ-------------------
100 YP_009508546.1     FYTDGSAIKSPDPTKSNNAGMGIVHVTYKP--------------------
101 NP_056762.1        TNENGSVTIHKDYITIQKTTGIYPTARHELKSEFAR--------------
102 YP_009508556.1     FYTDGSAIKHPDINKSHSAGMGIAQVQFQP--------------------
103 NP_569153.1        RSMYGGVRLEGNTITFYKQITSINTRLAAPLLK-----------------
104 NP_043924.1        QEIWQLIEHDSTQTLGIVKVKAHTQRKCSTHEQQLNNDVDQPAKQYAKEE
105 YP_009243641.1     QIISDIPLE-GPNVFTDGSRQR--SAYWAEK-------------------
106 YP_004300274.1     RMYLPHKIS--KDFWYLTTPCGKMIGAKIVENKERF--------------
107 NP_056848.1        NKYSPIELDIGKGIIKFTKNNEKYPNYLVKYPKKRK--------------
108 YP_006495799.1     KAMQGGIRFEGTEVTIYKKVTTIQTTLEPI--------------------
109 YP_009508409.1     --------------------------------------------------
110 NP_687035.1        YSLVKEPIKGEDVYYVDGAASKVTKLGKAGYL------------------
111 NP_045937.2        PIPDPDMTLFSDGSYTTGRGGAAVVMHRPVTD------------------
112 NP_612577.1        QLYSPFIQYTDRIYFHL---NKQ-SVIIGKIT------------------
113 NP_619548.1        RLYNPFIQWEDRIAFHL----KNEMVLIKKVT------------------
114 NP_056880.1        HMTSTTPLEKGIVIFTDGSANGRSVTYIQGR-------------------
115 YP_009121747.1     RSMNGGIRIEGDEVTLYKKVTKIKTNPTVEITA-A---------------
116 YP_009113237.2     RAMNGGLRIEGPTLTFYKLLTSIETKTVAALIK-----------------
117 YP_009508408.1     RSMSGGVRIEGDNVTFYKNVTTIKTQQEVSKVLSL---------------
118 NP_955579.1        PLPDADLTWYTDGSSFIRNGEREAGAAVTT--------------------
119 YP_009229919.1     RSMYGGLRIEGNLLTFYKNVTTVTTVQNTEASKLA---------------
120 YP_004442833.1     QSLKGAVRLEGRSVTFYKLVSTIEAD------------------------
121 NP_862833.2        KIVRQQPIPEATLIFTDGSSNG-TAALIINQ-------------------
122 NP_040563.1        TSICGPVRENLTTYYTDGGK--KGKTAAAVYW------------------
123 YP_443922.1        PLPQADLWLYTDGCCYRGKDGNIAAYAVVQQDSNG---------------
124 NP_057933.2        PLPDADHTWYTDGSSLLQEGQRKAGAAVTT--------------------
125 YP_009116631.1     RAMKGGVRLEGTEVTIYKKVTTIQTTLEPQ--------------------
126 YP_009513249.1     FYTDGSAITSPTKEGHLNAGMGIVYFINKDG-------------------
127 YP_223871.1        PLAQAEATLFTDGSSYVRDGKRYAGAAVVT--------------------
128 NP_127504.1        TTVKTKHPITIEFFPGFKYTTKLLGSDIPGKDLLIG--------------
129 NP_044929.1        FYTDGSAIRSPKPNKTHSAGMGIIQAKFEP--------------------
130 NP_057860.1        FTLSPVIINTAPCLFSDGSTSQAAYILWDKHI------------------
131 NP_040973.1        SMIKDAPIPGAETWYIDGGR-KLGKAAKAAYW------------------
132 NP_777317.1        SSMEGGIRIEQGMVTFYKLVTQAETSPIVHEL------------------
133 NP_054716.1        FYTDGSSIKSPKKDKQHSAGMGIIAVRYQP--------------------
134 YP_001856242.1     YQLEKEPIVGAETFYVDGAANRETKLGKAGYV------------------
135 NP_954565.2        QIISKTPLNNALLVFTDGSSTG-MAAYTLTD-------------------
136 NP_569150.1        RTVG--LRIENGEVTIYKIMTTVQAPPIVHEL------------------
137 NP_659397.1        QTYQPFIQWVDHIAFHIKENNEEKRVLIPKVR------------------
138 YP_009513211.1     PLPGVP-AWYTDGSSFIMDGRRQAGAAIVD--------------------
139 YP_233107.1        KAMEGGLRIEKGEVTFYKLVTTVNTSPKPHEV------------------
140 YP_009109692.1     ELPDADHTWYTDGSSYLDSGTRRAGAAVVD--------------------
141 NP_057849.4        YQLEKEPIVGAETFYVDGAANRETKLGKAGYV------------------
142 NP_049560.1        FTLSPVIINAAPCLFSDGSTSRAAYILWDKHI------------------
143 YP_009506267.1     RSMNGGVRIEGNDVTFYKNITTVKTQTEVPKILAL---------------
144 NP_039820.1        RSLEGGLRIEKDIITFYKLVTSIETSRTTQVA------------------
145 NP_056790.1        PLPGVP-TWYTDGSSFITEGKRRAGAPIVD--------------------
146 YP_002321513.1     RSMKGGIKIEGNTITFYERSYSIQTCEESGEIAAK---------------
147 NP_955591.1        PLPDADHTWYTDGSSLLQEGQRKAGAAVTT--------------------
148 YP_001956722.2     FYTDGSAIKHPNVNKSHNAGMGIAQVQFKP--------------------
149 NP_056902.1        RMKLVEEPTSGITIYTDGGK-QNGE-GIAAYV------------------
150 NP_056886.1        VRVTDHPVPGPTVFTDASSSTHKGVVVWREGP------------------
151 NP_041261.1        KITKCAPIPQATLVFTDGSSNG-IAAYVIDN-------------------
```

```
001 YP_004442836.1    --NYIEELELELYEYYDICATQ----------------------------
002 NP_056728.1       ---RAVRVG--TEGFLESMKKRS---------------------------
003 YP_009508537.1    --------------------------------------------------
004 YP_004732983.2    -----KIAYLEELVEA-EDLHYE---------------------------
005 YP_009508407.1    --------------------------------------------------
006 NP_040939.1       --------------------------------------------------
007 YP_003864102.1    --------------------------------------------------
008 YP_009002585.1    EDDEDDEVEQ---ETNWIRTMAS---------------------------
009 YP_003284237.1    --AAIEELDLNEDEYYDIAISD----------------------------
010 YP_009268869.1    --------------------------------------------------
011 YP_009506251.1    KASEESDDDAEIDFEIHQQIISH---------------------------
012 NP_955611.1       --------------------------------------------------
013 YP_008567619.1    -----EVISEEEEEYSAVEEICT---------------------------
014 NP_149413.1       --------EFLQAEVEEINIAT----------------------------
015 YP_004442824.1    --NYIEELELEIPEYYDICSTVG---------------------------
016 YP_001931967.1    ---KAYAYG--KPGFIQSMKKGS---------------------------
017 YP_002455786.1    --------------------------------------------------
018 YP_009109694.1    --------------------------------------------------
019 NP_789739.1       --------------------------------------------------
020 NP_955619.1       --------------------------------------------------
021 YP_224289.1       --NLIEELEMSEEEFIEVQIST----------------------------
022 YP_009182100.1    --RKEFLERRKIPDHQQSHKVKCGENYKVQKQAQENDFTIEDYGTTDSEA
023 NP_705927.1       --------------------------------------------------
024 YP_009508577.1    --------------------------------------------------
025 YP_004442830.1    --------EYLKMAEEQLMIAQ----------------------------
026 NP_758887.1       --------------------------------------------------
027 YP_009508406.1    --------------------------------------------------
028 YP_610965.1       -----AQFDEPEELMEIICPQWH---------------------------
029 YP_001931961.1    ---RAHRVG--VPGFLESLKKKS---------------------------
030 YP_241114.1       --------------------------------------------------
031 YP_006732334.1    ---KAMRKG--YEGFLKTMERDS---------------------------
032 YP_009041481.1    GAHQQQIAELTTTESVLDVQEAY---------------------------
033 YP_001036293.1    -----EELEMDEDEYIELQQSIS---------------------------
034 YP_605811.1       --SYLEELELELPIYYDICATN----------------------------
035 NP_955577.1       --------------------------------------------------
036 YP_003208050.1    ----IPELDLDAEMLAEIQELVY---------------------------
037 NP_789740.1       --------------------------------------------------
038 YP_004442839.1    --SYIEELELELPLYYDICAAN----------------------------
039 YP_009507791.1    --------------------------------------------------
040 YP_004581513.1    ----------------------I---------------------------
041 NP_043933.1       ---KAYFHG--LPGFLESKKVGS---------------------------
042 YP_009507248.1    --------------------------------------------------
043 YP_002117531.1    --AAIQELELDEIEYQNLGALD----------------------------
044 YP_004347415.1    --LIPWEKGN----------------------------------------
045 NP_056891.1       --------------------------------------------------
046 YP_009513242.1    --------------------------------------------------
047 YP_009352866.1    GLPQQQVAELLTAEAVLDHQEAY---------------------------
048 YP_009165750.1    ---KAYLHG--MPGYLDSMKKNS---------------------------
049 NP_040840.1       --------------------------------------------------
050 YP_004222728.1    --------------------------------------------------
051 YP_009506264.1    --ASIEELDLNEDEYYDIALMD----------------------------
052 YP_567050.1       --------------------------------------------------
053 YP_009508571.1    --------------------------------------------------
054 NP_056803.1       --------------------------------------------------
055 NP_955564.1       --------------------------------------------------
056 YP_009508411.1    -----EELEMTEDEFIEVNETLS---------------------------
057 YP_009140788.1    -----LEEEVCEEEYLQIQDMVV---------------------------
058 YP_009508410.1    --------------------------------------------------
059 NP_040333.1       --------------------------------------------------
060 YP_008992013.1    -FDILHQIK-HLQIIPHGIRVKS---------------------------
061 NP_041734.1       ESMFDLSETEEFDSETHQQIVSH---------------------------
062 NP_056895.1       --------------------------------------------------
063 YP_009109689.1    --------------------------------------------------
064 YP_002519387.1    ---RAFKQA--SPGFLDSIRKQKRGE------------------------
065 YP_595725.1       --------EYLQAEAEEILVAT----------------------------
066 YP_009508551.1    --------------------------------------------------
067 YP_001497148.1    --------------------------------------------------
068 NP_041186.1       --------------------------------------------------
069 NP_663784.1       --------------------------------------------------
070 YP_009508443.1    -----EELEMGEEEYIAIQESCS---------------------------
```

```
071 YP_003987465.1   --AIIEELEMAEDEYIDLNLML----------------------------
072 NP_861410.1      ---KMIYIPVTTPSKILQFAILE---------------------------
073 YP_009345071.1   -------FSKNFSEYMLLNDTGT---------------------------
074 YP_009508582.1   --------------------------------------------------
075 NP_569141.1      --TTEWIRGS----------------------------------------
076 YP_002916057.1   ----------------------I---------------------------
077 NP_777384.2      --------------------------------------------------
078 YP_233110.1      --SYLEELELELPIYYDICATN----------------------------
079 NP_542258.1      --------------------------------------------------
080 YP_009130664.1   -----EELELEEEEYISIATICA---------------------------
081 YP_009508566.1   --------------------------------------------------
082 YP_006607892.1   ---KAYYHG--KPGYLESMKKNS---------------------------
083 YP_009506270.1   --NYIEELELELQEYYNICSAE----------------------------
084 NP_042513.1      ---SLITTKIYNTFSLFGGVIVN---------------------------
085 YP_006907834.1   ---RAFKRG--IPGFLESMKKRS---------------------------
086 YP_004442827.1   --------EYIQKIEEQVLIVQ----------------------------
087 YP_009408594.1   ----EEDLD-----AAVMTELSL---------------------------
088 YP_009508561.1   --------------------------------------------------
089 NP_056907.1      --------------------------------------------------
090 YP_002308474.1   --------------------------------------------------
091 YP_007761644.1   --YKTYSFDFWCHDAMSHDIIVG---------------------------
092 NP_040550.1      --------------------------------------------------
093 NP_758808.1      ----YANTGIDTEHEQITSTT-----------------------------
094 YP_009345075.1   ----IEELEMDELDYLKIHESVF---------------------------
095 NP_068729.1      ---QMISTKILTKNEVLKLSFEK---------------------------
096 NP_047255.1      --------------------------------------------------
097 YP_006273075.1   -----LDEEVCEEEYLQVQEMVI---------------------------
098 YP_009508888.1   --------------------------------------------------
099 NP_395469.1      ---EVIKIPIHHSSQLIKAKLLN---------------------------
100 YP_009508546.1   --------------------------------------------------
101 NP_056762.1      ------KHGGRPPLFSNIPETYN---------------------------
102 YP_009508556.1   --------------------------------------------------
103 NP_569153.1      QEEEEKEEELNLEEHRLIQEMVA---------------------------
104 NP_043924.1      PNMSVIAPLQVYPLWIGLVPCKE---------------------------
105 YP_009243641.1   --------------------------------------------------
106 YP_004300274.1   --KCEWEKGD----------------------------------------
107 NP_056848.1      --LVPWTKGN----------------------------------------
108 YP_006495799.1   -----KLGYAEFDPDIQVELERA---------------------------
109 YP_009508409.1   --------------------------------------------------
110 NP_687035.1      --------------------------------------------------
111 NP_045937.2      --------------------------------------------------
112 NP_612577.1      ---KAYQYG--VKGFLESMKKKS---------------------------
113 NP_619548.1      ---KAFSVS--NPSFLENMKKDS---------------------------
114 NP_056880.1      --------------------------------------------------
115 YP_009121747.1   ----IQELEIDEEFYQELQGTXM---------------------------
116 YP_009113237.2   ------EEEVNYEELDQIKEIVA---------------------------
117 YP_009508408.1   -----EELEMNEDEYISIQQSVS---------------------------
118 NP_955579.1      --------------------------------------------------
119 YP_009229919.1   ----VEELGLEAEEYYSIREMVY---------------------------
120 YP_004442833.1   --------EYIRRAEEQICIAH----------------------------
121 NP_862833.2      --------------------------------------------------
122 NP_040563.1      --------------------------------------------------
123 YP_443922.1      --------------------------------------------------
124 NP_057933.2      --------------------------------------------------
125 YP_009116631.1   -----KISLLRAEAEVGEELERM---------------------------
126 YP_009513249.1   --------------------------------------------------
127 YP_223871.1      --------------------------------------------------
128 NP_127504.1      -FDIYRQLNNKLRIGADGIRWKN---------------------------
129 NP_044929.1      --------------------------------------------------
130 NP_057860.1      --------------------------------------------------
131 NP_040973.1      --------------------------------------------------
132 NP_777317.1      --NYIEELELELPEYYNICAAA----------------------------
133 NP_054716.1      --------------------------------------------------
134 YP_001856242.1   --------------------------------------------------
135 NP_954565.2      --------------------------------------------------
136 NP_569150.1      --NYIDELELELHEYYNICAAE----------------------------
137 NP_659397.1      ---IAMRKG--HPGFLEAMRKGS---------------------------
138 YP_009513211.1   --------------------------------------------------
139 YP_233107.1      --CLLDELDLELPEYYDICAAI----------------------------
140 YP_009109692.1   --------------------------------------------------
141 NP_057849.4      --------------------------------------------------
142 NP_049560.1      --------------------------------------------------
143 YP_009506267.1   -----EELEMEEEEYLALQSKYS---------------------------
```

```
144 NP_039820.1         --NSIEELELSEDEYLNIAASV----------------------------
145 NP_056790.1         --------------------------------------------------
146 YP_002321513.1      --YAIPELELNENEYQCLSESIY---------------------------
147 NP_955591.1         --------------------------------------------------
148 YP_001956722.2      --------------------------------------------------
149 NP_056902.1         --------------------------------------------------
150 NP_056886.1         --------------------------------------------------
151 NP_041261.1         --------------------------------------------------


001 YP_004442836.1      --------------------------------------------------
002 NP_056728.1         --------------------------------------------------
003 YP_009508537.1      --------------------------------------------------
004 YP_004732983.2      --------------------------------------------------
005 YP_009508407.1      --------------------------------------------------
006 NP_040939.1         --------------------------------------------------
007 YP_003864102.1      --------------------------------------------------
008 YP_009002585.1      --------------------------------------------------
009 YP_003284237.1      --------------------------------------------------
010 YP_009268869.1      --------------------------------------------------
011 YP_009506251.1      --------------------------------------------------
012 NP_955611.1         --------------------------------------------------
013 YP_008567619.1      --------------------------------------------------
014 NP_149413.1         --------------------------------------------------
015 YP_004442824.1      --------------------------------------------------
016 YP_001931967.1      --------------------------------------------------
017 YP_002455786.1      --------------------------------------------------
018 YP_009109694.1      --------------------------------------------------
019 NP_789739.1         --------------------------------------------------
020 NP_955619.1         --------------------------------------------------
021 YP_224289.1         --------------------------------------------------
022 YP_009182100.1      EEPIYDAIYDKGFQPIESDDEPNDEIQPINDEPNEEVRPGFQPIESNDEP
023 NP_705927.1         --------------------------------------------------
024 YP_009508577.1      --------------------------------------------------
025 YP_004442830.1      --------------------------------------------------
026 NP_758887.1         --------------------------------------------------
027 YP_009508406.1      --------------------------------------------------
028 YP_610965.1         --------------------------------------------------
029 YP_001931961.1      --------------------------------------------------
030 YP_241114.1         --------------------------------------------------
031 YP_006732334.1      --------------------------------------------------
032 YP_009041481.1      --------------------------------------------------
033 YP_001036293.1      --------------------------------------------------
034 YP_605811.1         --------------------------------------------------
035 NP_955577.1         --------------------------------------------------
036 YP_003208050.1      --------------------------------------------------
037 NP_789740.1         --------------------------------------------------
038 YP_004442839.1      --------------------------------------------------
039 YP_009507791.1      --------------------------------------------------
040 YP_004581513.1      --------------------------------------------------
041 NP_043933.1         --------------------------------------------------
042 YP_009507248.1      --------------------------------------------------
043 YP_002117531.1      --------------------------------------------------
044 YP_004347415.1      --------------------------------------------------
045 NP_056891.1         --------------------------------------------------
046 YP_009513242.1      --------------------------------------------------
047 YP_009352866.1      --------------------------------------------------
048 YP_009165750.1      --------------------------------------------------
049 NP_040840.1         --------------------------------------------------
050 YP_004222728.1      --------------------------------------------------
051 YP_009506264.1      --------------------------------------------------
052 YP_567050.1         --------------------------------------------------
053 YP_009508571.1      --------------------------------------------------
054 NP_056803.1         --------------------------------------------------
055 NP_955564.1         --------------------------------------------------
056 YP_009508411.1      --------------------------------------------------
057 YP_009140788.1      --------------------------------------------------
058 YP_009508410.1      --------------------------------------------------
059 NP_040333.1         --------------------------------------------------
060 YP_008992013.1      --------------------------------------------------
061 NP_041734.1         --------------------------------------------------
062 NP_056895.1         --------------------------------------------------
```

```
063 YP_009109689.1    --------------------------------------------------
064 YP_002519387.1    --------------------------------------------------
065 YP_595725.1       --------------------------------------------------
066 YP_009508551.1    --------------------------------------------------
067 YP_001497148.1    --------------------------------------------------
068 NP_041186.1       --------------------------------------------------
069 NP_663784.1       --------------------------------------------------
070 YP_009508443.1    --------------------------------------------------
071 YP_003987465.1    --------------------------------------------------
072 NP_861410.1       --------------------------------------------------
073 YP_009345071.1    --------------------------------------------------
074 YP_009508582.1    --------------------------------------------------
075 NP_569141.1       --------------------------------------------------
076 YP_002916057.1    --------------------------------------------------
077 NP_777384.2       --------------------------------------------------
078 YP_233110.1       --------------------------------------------------
079 NP_542258.1       --------------------------------------------------
080 YP_009130664.1    --------------------------------------------------
081 YP_009508566.1    --------------------------------------------------
082 YP_006607892.1    --------------------------------------------------
083 YP_009506270.1    --------------------------------------------------
084 NP_042513.1       --------------------------------------------------
085 YP_006907834.1    --------------------------------------------------
086 YP_004442827.1    --------------------------------------------------
087 YP_009408594.1    --------------------------------------------------
088 YP_009508561.1    --------------------------------------------------
089 NP_056907.1       --------------------------------------------------
090 YP_002308474.1    --------------------------------------------------
091 YP_007761644.1    --------------------------------------------------
092 NP_040550.1       --------------------------------------------------
093 NP_758808.1       --------------------------------------------------
094 YP_009345075.1    --------------------------------------------------
095 NP_068729.1       --------------------------------------------------
096 NP_047255.1       --------------------------------------------------
097 YP_006273075.1    --------------------------------------------------
098 YP_009508888.1    --------------------------------------------------
099 NP_395469.1       --------------------------------------------------
100 YP_009508546.1    --------------------------------------------------
101 NP_056762.1       --------------------------------------------------
102 YP_009508556.1    --------------------------------------------------
103 NP_569153.1       --------------------------------------------------
104 NP_043924.1       --------------------------------------------------
105 YP_009243641.1    --------------------------------------------------
106 YP_004300274.1    --------------------------------------------------
107 NP_056848.1       --------------------------------------------------
108 YP_006495799.1    --------------------------------------------------
109 YP_009508409.1    --------------------------------------------------
110 NP_687035.1       --------------------------------------------------
111 NP_045937.2       --------------------------------------------------
112 NP_612577.1       --------------------------------------------------
113 NP_619548.1       --------------------------------------------------
114 NP_056880.1       --------------------------------------------------
115 YP_009121747.1    --------------------------------------------------
116 YP_009113237.2    --------------------------------------------------
117 YP_009508408.1    --------------------------------------------------
118 NP_955579.1       --------------------------------------------------
119 YP_009229919.1    --------------------------------------------------
120 YP_004442833.1    --------------------------------------------------
121 NP_862833.2       --------------------------------------------------
122 NP_040563.1       --------------------------------------------------
123 YP_443922.1       --------------------------------------------------
124 NP_057933.2       --------------------------------------------------
125 YP_009116631.1    --------------------------------------------------
126 YP_009513249.1    --------------------------------------------------
127 YP_223871.1       --------------------------------------------------
128 NP_127504.1       --------------------------------------------------
129 NP_044929.1       --------------------------------------------------
130 NP_057860.1       --------------------------------------------------
131 NP_040973.1       --------------------------------------------------
132 NP_777317.1       --------------------------------------------------
133 NP_054716.1       --------------------------------------------------
134 YP_001856242.1    --------------------------------------------------
135 NP_954565.2       --------------------------------------------------
```

```
136 NP_569150.1       --------------------------------------------------
137 NP_659397.1       --------------------------------------------------
138 YP_009513211.1    --------------------------------------------------
139 YP_233107.1       --------------------------------------------------
140 YP_009109692.1    --------------------------------------------------
141 NP_057849.4       --------------------------------------------------
142 NP_049560.1       --------------------------------------------------
143 YP_009506267.1    --------------------------------------------------
144 NP_039820.1       --------------------------------------------------
145 NP_056790.1       --------------------------------------------------
146 YP_002321513.1    --------------------------------------------------
147 NP_955591.1       --------------------------------------------------
148 YP_001956722.2    --------------------------------------------------
149 NP_056902.1       --------------------------------------------------
150 NP_056886.1       --------------------------------------------------
151 NP_041261.1       --------------------------------------------------

001 YP_004442836.1    --------------------NQRGEIAEEFIEP-----------------
002 NP_056728.1       ------KTQQPE-------PVNIS--TNKIENPL----EEIAILSEGRRL
003 YP_009508537.1    --------------------NLQKQQEWSISLGN----------------
004 YP_004732983.2    --------------------LAAASMPEPTAEGLR---------------
005 YP_009508407.1    --------------------------------------------------
006 NP_040939.1       ---------------------VSTGEKFRKHEE-----------------
007 YP_003864102.1    -----------------------SIIKFQTK--F----------------
008 YP_009002585.1    --------------------STMDAPYEKLRSS-----------------
009 YP_003284237.1    --------------------QEKAYINKEIVDS-----------------
010 YP_009268869.1    ---------------------ASTGVKFRKHEE-----------------
011 YP_009506251.1    --------------------AVAQATTPKLSPK-----------------
012 NP_955611.1       ---------------------RWEIKEIADL--G----------------
013 YP_008567619.1    ----------------------GTVIHPEFGNK-----------------
014 NP_149413.1       --------------------SEGGFIDKPFMSKN----------------
015 YP_004442824.1    --------------------AIQGGISEEFISP-----------------
016 YP_001931967.1    ------KVKPPT-------PENISRLTDKIVSFSGGESSEDTTKTNFEND
017 YP_002455786.1    ---------------------LSQRSVPLPPHAN----------------
018 YP_009109694.1    --------------------GHNTIWAQSLPPG-----------------
019 NP_789739.1       --------------------------------------------------
020 NP_955619.1       ---------------------LSQRSFPLPP-PH----------------
021 YP_224289.1       --------------------STEGYINKEIMDS-----------------
022 YP_009182100.1    NNEIQPSNDEPNKEVRPKPTKDIR--PKPITEEDFQAELRLVEAHYEKQA
023 NP_705927.1       ---------------------TNRGRQKVVTLTD----------------
024 YP_009508577.1    --------------------ELTIEHQWSIPLGD----------------
025 YP_004442830.1    --------------------QPSPFIEKSFMTRN----------------
026 NP_758887.1       ---------------------TARNKSRVIALEN----------------
027 YP_009508406.1    --------------------------------------------------
028 YP_610965.1       --------------------IEHPTDAIGVGLR-----------------
029 YP_001931961.1    ------KLTMQK-------GTNIA--PSKT--------SFDKRGELGYTL
030 YP_241114.1       -----------------------QTYYTQTS--F----------------
031 YP_006732334.1    ------KSQPTP-------GTNIT--QEVIDE------------------
032 YP_009041481.1    --------------------FARPEQPVDLKFKHV---------------
033 YP_001036293.1    --------------------SIEP--NNYFSNK-----------------
034 YP_605811.1       --------------------PSGGEINSDLIPP-----------------
035 NP_955577.1       --------------------ESEVIWAAPLPPG-----------------
036 YP_003208050.1    --------------------YQSPARNPRFESR-----------------
037 NP_789740.1       ---------------------TNRGRQKVVTLTD----------------
038 YP_004442839.1    --------------------PIGGMFNNDLISP-----------------
039 YP_009507791.1    ---------------------TSEGKEKYEYIE-----------------
040 YP_004581513.1    --------------------GSINEGRTLIFPR-----------------
041 NP_043933.1       ------STSTPN-------PENIT--PVTINQDNISKIFKGEEIGESEQ-
042 YP_009507248.1    --------------------PQRQTENTDEIRQ-----------------
043 YP_002117531.1    ---------------------DQAFFNKKFMEQN----------------
044 YP_004347415.1    --------------------PSIIQPLQNEQIN-----------------
045 NP_056891.1       -----------------------TTIKFQTN--L----------------
046 YP_009513242.1    --------------------EYKVIHQWSIPLGD----------------
047 YP_009352866.1    --------------------FAQPEKPIDQKFKHV---------------
048 YP_009165750.1    ------RNPVPP-------PINIT--SNKIEDSKRFSVLKLQLLERGREI
049 NP_040840.1       ---------------------TSTGEKFRIHEE-----------------
050 YP_004222728.1    ---------------------RWEIKEIADL--G----------------
051 YP_009506264.1    --------------------EEKGYINREIVDS-----------------
052 YP_567050.1       ---------------------LHHDSVTLPPHGS----------------
053 YP_009508571.1    --------------------ELQVINQWSIPLGN----------------
054 NP_056803.1       --------------------EYKILNQWSIPLGH----------------
```

```
055 NP_955564.1       -----------------------EPIIKENT--Q----------------
056 YP_009508411.1    --------------------VVGR----QPQPK-----------------
057 YP_009140788.1    --------------------CNIGESRESFLKK-----------------
058 YP_009508410.1    --------------------------------------------------
059 NP_040333.1       --------------------ETEVVWAKALPAG-----------------
060 YP_008992013.1    ------------------MFKPFTDVLKLYNLSETP--------------
061 NP_041734.1       --------------------VAAQAQQQKLDPK-----------------
062 NP_056895.1       ---------------------LDFQAVPAP--------------------
063 YP_009109689.1    --------------------GHNTIWAQSLPPG-----------------
064 YP_002519387.1    ----IISKHATN-------PINITKLEDNLRIVETILEDSNSKTLSEVSD
065 YP_595725.1       --------------------SEQEFINRSFMSKN----------------
066 YP_009508551.1    --------------------EYQVLNQWSIPLGN----------------
067 YP_001497148.1    --------------------GHHIIWAQSLPPG-----------------
068 NP_041186.1       -----------------------QTYYAQTS--F----------------
069 NP_663784.1       ---------------------TDRGKDKVKVLEQ----------------
070 YP_009508443.1    --------------------TICPVVNINFAQR-----------------
071 YP_003987465.1    --------------------ATEPAVGEEFAKL-----------------
072 NP_861410.1       ------------------KLQDILFELHVQEN------------------
073 YP_009345071.1    --------------------DELQNEQENLRIKKA---------------
074 YP_009508582.1    --------------------ELTIEHQWTIPLGD----------------
075 NP_569141.1       --------------------EKITQKLENINKNTTT--------------
076 YP_002916057.1    --------------------GNINEERTMVFPR-----------------
077 NP_777384.2       ---------------------LDFQAVPAP--------------------
078 YP_233110.1       --------------------PSGGEINSDLISP-----------------
079 NP_542258.1       ---------------------IHQQVLPLP--ST----------------
080 YP_009130664.1    --------------------YGGMEISSPFQFS-----------------
081 YP_009508566.1    --------------------NFQIKHQWSIPLGD----------------
082 YP_006607892.1    ------KKPVPS-------PLNIT--PVKIEEQE---TLRLQLLRRGREI
083 YP_009506270.1    --------------------TSKGEIAEEFISP-----------------
084 NP_042513.1       ------------------ILKQQIYIAIEDE-------------------
085 YP_006907834.1    ------TAQQPE-------PLNIS--TNKTVS-----------LSRGRRF
086 YP_004442827.1    --------------------EPAPYIDRPFMTKN----------------
087 YP_009408594.1    --------------------FSAVPSNQKFLQQ-----------------
088 YP_009508561.1    --------------------EPNITQQWSIPLGD----------------
089 NP_056907.1       ---------------------LHHDSVTLPPHGS----------------
090 YP_002308474.1    ---------------------IHQQVLPLP--TA----------------
091 YP_007761644.1    ----NPFWIYLQQSFMSITGNTVHTKDNSFVFYEHKNPARETPWQSEFTH
092 NP_040550.1       ---------------------RWEIKEIADL--G----------------
093 NP_758808.1       --------------------------SQSFKER-----------------
094 YP_009345075.1    --------------------YNVGQKLGNHGPE-----------------
095 NP_068729.1       ------------------IHICLEKYLFFKT-------------------
096 NP_047255.1       --------------------ESEVIWAAPLPPG-----------------
097 YP_006273075.1    --------------------CNIGESRKSFLKK-----------------
098 YP_009508888.1    --------------------EFQVVNQWSIPLGN----------------
099 NP_395469.1       ------------------LVTNIEEQLLMEQ-------------------
100 YP_009508546.1    --------------------EYQILNQWSIPLGN----------------
101 NP_056762.1       ------------------KIPHLHSYQPQPILGYKN--------------
102 YP_009508556.1    --------------------EYKVIHQWSIPLGD----------------
103 NP_569153.1       --------------------YSTERPFVQFQQK-----------------
104 NP_043924.1       -------------------PKLWENIQHHITKVDLP--------------
105 YP_009243641.1    -----------------------KYWSQKQN--T----------------
106 YP_004300274.1    --------------------KTLNQKLRNVYD------------------
107 NP_056848.1       --------------------PSVTETMENIGIN-----------------
108 YP_006495799.1    --------------------YYAAPVSEAELIKLR---------------
109 YP_009508409.1    --------------------------------------------------
110 NP_687035.1       ---------------------SERGKSRIRELEN----------------
111 NP_045937.2       -----------------------DFIIIHQQPGG----------------
112 NP_612577.1       ------KVNRPE-------PINIT--SNQH--------LFLEEG--GNHV
113 NP_619548.1       ------KTEQIP-------GTNIS--KNIINP------------------
114 NP_056880.1       -----------------------EPIIKENT--Q----------------
115 YP_009121747.1    --------------------FSKE-DAESLKER-----------------
116 YP_009113237.2    --------------------MNVHEPYPGFAQR-----------------
117 YP_009508408.1    --------------------AISPGIHSSFTTR-----------------
118 NP_955579.1       --------------------ESEVIWAAPLPPG-----------------
119 YP_009229919.1    --------------------YSLGHTSPKFQER-----------------
120 YP_004442833.1    --------------------DPASFVQETFMKKN----------------
121 NP_862833.2       -----------------------QTYYAHTN--F----------------
122 NP_040563.1       ---------------------CEG--RTKSKVFP----------------
123 YP_443922.1       --------------------NHSTLESGVIPQP-----------------
124 NP_057933.2       --------------------ETEVIWAKALPAG-----------------
125 YP_009116631.1    --------------------YYANDYSEEGISRLK---------------
126 YP_009513249.1    --------------------NLQKQQEWSISLGN----------------
127 YP_223871.1       --------------------LDSVIWAEPLPIG-----------------
```

128 NP_127504.1 ------------------QFKRYTEIPRLFQLTTSN--------------
129 NP_044929.1 --------------------DFRIVHLWSFPLGD----------------
130 NP_057860.1 ---------------------LSQRSFPLPP-PH----------------
131 NP_040973.1 ---------------------TDTGKWQVMELE-----------------
132 NP_777317.1 --------------------PESGVISEDFLDK-----------------
133 NP_054716.1 --------------------QMNIIQEWSIPLGD----------------
134 YP_001856242.1 ---------------------TNRGRQKVVTLTD----------------
135 NP_954565.2 -----------------------TTIKFQTN--L----------------
136 NP_569150.1 --------------------SSRGEISEEFISP-----------------
137 NP_659397.1 ------GKKPVP-------GTNIT--QEVIDD------------------
138 YP_009513211.1 --------------------NKRTVWASNLPEG-----------------
139 YP_233107.1 --------------------PRQGSINEEFISP-----------------
140 YP_009109692.1 --------------------GHNTIWAQSLPPG-----------------
141 NP_057849.4 ---------------------TNRGRQKVVTLTD----------------
142 NP_049560.1 ---------------------LSQKSFPLPP-PH----------------
143 YP_009506267.1 --------------------SICPVIQENFKAR-----------------
144 NP_039820.1 --------------------ETPSFLDQEFARKN----------------
145 NP_056790.1 --------------------GKRTVWASSLPEG-----------------
146 YP_002321513.1 --------------------LNIGTTSDGFKSR-----------------
147 NP_955591.1 --------------------ETEVIWAKALPAG-----------------
148 YP_001956722.2 --------------------EFTVINTWSIPLGD----------------
149 NP_056902.1 ---------------------TSNGRTKQKRLGP----------------
150 NP_056886.1 ---------------------RWEIKEIADL--G----------------
151 NP_041261.1 -----------------------QPISIKSP--Y----------------

001 YP_004442836.1 -------------------AMMTRMKKLGYIGEEPLKHWTKNQV------
002 NP_056728.1 SE-------EKLFITQQR-MQKIEELLEKVCSENPLDPNKTKQW------
003 YP_009508537.1 --------------------HTAQFAEIAAFEFALKKCLPLGGN------
004 YP_004732983.2 -----------------NTKLLAELKDQGYIGEEPLKHWSKNRV------
005 YP_009508407.1 ------------------------------MN-HTEEYKRALQA------
006 NP_040939.1 --------------------GTNQQLELRAIEEALKQGPQT---------
007 YP_003864102.1 --------------------SSAQLVELQAFIAVLSAFPNQP--------
008 YP_009002585.1 -----------------FGPLMKELKDQGYIGENPMKHWARNKV------
009 YP_003284237.1 -------------------SIFRRLKELGYIGEEPLKHWRKNQV------
010 YP_009268869.1 --------------------GTNQQLELRAIEEACKQGPEK---------
011 YP_009506251.1 -----------------LKHLMGQLKDQGFIGENPMKHWASNKI------
012 NP_955611.1 --------------------ASVQQLEARAVAMALLLWPTTP--------
013 YP_008567619.1 -----------------YQKLIEDMKNQGYIGEDPMKYWQKNQV------
014 NP_149413.1 ------------------KQLIEEMKEQGYMGEDTLKHWEKNKV------
015 YP_004442824.1 -------------------EFIQKMKKLGYIGEEPLKHWEKNQV------
016 YP_001931967.1 FL-------KTENIFEIQSTSEIEELLELACSENPLDPEKSKGL------
017 YP_002455786.1 --------------------NSAQKGELVGLLLGLQAAQPWPS-------
018 YP_009109694.1 --------------------TSAQKAELIALTKALELSKGKK--------
019 NP_789739.1 --------------------------------------------------
020 NP_955619.1 --------------------KSAQRAELLGLLHGLSSARSWRC-------
021 YP_224289.1 -------------------KLFKELKDQGYIGEEPLKHWQKNMV------
022 YP_009182100.1 RPKQRPNLNVLEEQISRCSIEELTKQLEPNISDNPIH-KWDVTQ------
023 NP_705927.1 --------------------TTNQKTELQAIYLALQDSGLE---------
024 YP_009508577.1 --------------------HTAQYAEISAVEFACKKANNISGP------
025 YP_004442830.1 ------------------KEIINEMKELGYIGDEPLRYWANNKV------
026 NP_758887.1 --------------------TTNQKAELEAIKMALQDSGPK---------
027 YP_009508406.1 --------------------------------------------------
028 YP_610965.1 -----------------AQKIVQRLLQQN-ISDDPLKFWAKNKV------
029 YP_001931961.1 GL-------KKLFEEDLKKKSIIEKLLDQVCSENPLDPLKTKKW------
030 YP_241114.1 --------------------SSAQVVELFAVHQALLTVPTS---------
031 YP_006732334.1 ---------ENNFIIDISRYQEIQELLKRVCSENPIDPQKSKGW------
032 YP_009041481.1 -----------------LGPILKDLMAQGYIGEDPVRHWVKNQV------
033 YP_001036293.1 -----------------LSQVFQELKAAGYIGDDPLKFWSKNQV------
034 YP_605811.1 -------------------SEIQKLKDLGYIGEEPLKHWAKNQV------
035 NP_955577.1 --------------------TSAQRAELIALTQALKMAEGKK--------
036 YP_003208050.1 -----------------FEGLIGRLRNLGFIGENPVKHWARNQV------
037 NP_789740.1 --------------------TTNQKTELQAIYLALQDSGLE---------
038 YP_004442839.1 -------------------VEIQKLKDLGYIGEEPLKHWAKNQV------
039 YP_009507791.1 --------------------GSNQQAEVNALRMALVDGPSE---------
040 YP_004581513.1 -----------------FRKEVAALIKEGFIGNNPLLHWTKNRV------
041 NP_043933.1 ----------LFSTIS--AYTEVEKLLDSICSEHPLDSRINKGK------
042 YP_009507248.1 --------------------NSTTDTVHTSDEHPILEVPITNEP------
043 YP_002117531.1 ------------------RALMERLKKIGIIGEDPLKFWERNKV------
044 YP_004347415.1 ----------NIN----EVEELNNILGEDIYGDNPLKHWEKHKT------
045 NP_056891.1 --------------------NSAQLVELQALIAVLSAFPNQP--------
046 YP_009513242.1 --------------------HTAQLAEIAAVEFACKKALKISGP------

047 YP_009352866.1 -----------------LGPILADLKAQGFIGEDPVKHWKKNQV------
048 YP_009165750.1 IN-------RRLFLTQQDQLSTIEKLLDEVCSENPIDPHKSKKW------
049 NP_040840.1 --------------------GTNQQLELRAIEEACKQGPEK---------
050 YP_004222728.1 --------------------ASVQQLEARAVAMALLLWPTTP---------
051 YP_009506264.1 -------------------QLFRNLKEAGYIGEEPLKHWSQNQI------
052 YP_567050.1 --------------------NSAQKGELLALLSGIRAAKSWPS-------
053 YP_009508571.1 --------------------HTAQMAEIAAVEFACKKALKITGP------
054 NP_056803.1 --------------------HTAQMAEIAAVEFACKKALKVPGP------
055 NP_955564.1 --------------------NTAQQAEIVAVITAFEEVSQP---------
056 YP_009508411.1 -----------------YEAILSRMKQQGYIGDKPMRHWSKNKV------
057 YP_009140788.1 -----------------FKPLIEEFQEAGYIGENPLQHWEKNGV------
058 YP_009508410.1 ------------------------------MNSESLEYKRALQA------
059 NP_040333.1 --------------------TSAQRAELIALTQALKMAEGKK--------
060 YP_008992013.1 ---------PSYQDVSTKLLSFCPESHSEFT--HPNPLWKNQSF------
061 NP_041734.1 -----------------LQQLMVQLQDQGFIGENPMQHWAKNKI------
062 NP_056895.1 --------------------ESAQKGELAGLLAGLAAAPPEP--------
063 YP_009109689.1 --------------------TSAQKAELIALTKALELSKGKK--------
064 YP_002519387.1 SKTLSEDSDDYFVYSIQRHNTIVNDLLTQACSENPLDENKNHNG------
065 YP_595725.1 ------------------KRLLEEMKEQGYMGEDTLAHWNKNQI------
066 YP_009508551.1 --------------------HTAQMAEIAAVEFACKKALKIPGP------
067 YP_001497148.1 --------------------TSAQKAELIALTKALELSEGKK--------
068 NP_041186.1 --------------------SSAQVVELFAVHQALLTVPTS---------
069 NP_663784.1 --------------------TTNQQAELEVFRMALADSGPK---------
070 YP_009508443.1 -----------------FEDLILELKSAGYIGEEPMKFWSHNKV------
071 YP_003987465.1 -------------------PLYQQLKAEGFIGENPLLHWQRNQV------
072 NP_861410.1 --------------SKTPLELKVSSTLEEVCDENPLDV-KNTNT------
073 YP_009345071.1 -----------------VAHKLLQLSDMGYIGEEPLKHWKKNQV------
074 YP_009508582.1 --------------------HTAQYAEISAVEFACKKANNISGP------
075 NP_569141.1 --------QLEIIIFTIDKVKIIQKKLEKLYNDNPLQGWEKHKT------
076 YP_002916057.1 -----------------FRKEVARLIQEGYIGENPLRHWSKNKV------
077 NP_777384.2 --------------------ESAQKGELAGLLAGLAAAPPEP--------
078 YP_233110.1 -------------------SEIRKLKDLGYIGEEPLKHWAKNQV------
079 NP_542258.1 --------------------CSAQAGELFGLLAGLQKSKPWPA-------
080 YP_009130664.1 -----------------YQKLIQELKDMGYIGDDPMKYWSSNKI------
081 YP_009508566.1 --------------------HTAQYAEIAAVEFACKKALQVTGP------
082 YP_006607892.1 VT-------QKLFSIQQDHLSKIHNLLDRVCSENPIDPEKSKRW------
083 YP_009506270.1 -------------------NLINRMKRLGYIGEEPLKHWEKNQV------
084 NP_042513.1 ----------------------VTQLLEAICSQNPLDPQKNRNQ------
085 YP_006907834.1 G--------EKLRITQER-MVKIEELLEKVCSENPLDPEKSKGW------
086 YP_004442827.1 ------------------RKLMEEMKELGYIGDEPLKHWNNNKI------
087 YP_009408594.1 -----------------YRGTLDRLKEQGYIGDNPLQHWEKNQL------
088 YP_009508561.1 --------------------HTAQYAEISAVEFACKKASLLQGP------
089 NP_056907.1 --------------------NSAQKGELLALLSGIRAAKSWPS-------
090 YP_002308474.1 --------------------SSAQAGELFALLAALRECKPWSS-------
091 YP_007761644.1 AERGGHITQVSENSETKHWKMQVEEILLDTFSDNPLALYNPRHPKCQIEL
092 NP_040550.1 --------------------ASVQQLEARAVAMALLLWPTTP---------
093 NP_758808.1 -----------------FSPLMNELKAAGYIGEDPLKHWSKNKV------
094 YP_009345075.1 -----------------IQKALQQMEELNFIGPNPQIHWAKNRV------
095 NP_068729.1 ----------------------IEEQLEEVCSEHPLDETKNKNG------
096 NP_047255.1 --------------------TSAQRAELIALTQALKMAEGKK--------
097 YP_006273075.1 -----------------FRPLLEELKEAGFIGENPLQHWKKNGI------
098 YP_009508888.1 --------------------HTAQLAEVAAVEFACKQALKITGP------
099 NP_395469.1 ----------------------VNKILQERFSLDLLGE-KNKNK------
100 YP_009508546.1 --------------------HTAQMAEIAAVEFACKKALKIPGP------
101 NP_056762.1 -----------EIGNQSLITMVKELEALGFIGDDITKNRTTWVCD-----
102 YP_009508556.1 --------------------HTAQLAEIAAVEFACKKALKISGP------
103 NP_569153.1 -----------------FAGLIQDLKAQGYIGEEPMKYWAKNQV------
104 NP_043924.1 ----------------DFQKQLAEIMPQQDISHCTLAYFDKPSPEATKYH
105 YP_009243641.1 --------------------NSIQKNELLAIIQVLKDFKQD---------
106 YP_004300274.1 -------------------FSLVRNQIKTLFSENPLEFWDKHRT------
107 NP_056848.1 ----------QIESRNPIEEEINQILGTDIYGENPLEKWEKHKT------
108 YP_006495799.1 -----------------DHRLLAELKEQGFIGNDPMLHWAKNQV------
109 YP_009508409.1 --------------------------------------------------
110 NP_687035.1 --------------------TTNQQAELTAVKMALEDSGEN---------
111 NP_045937.2 --------------------ASAQTAELLALAAACHLATDKTVN------
112 NP_612577.1 D--------EMLYEIQISKFSAIEEMLERVSSENPIDPEKSKQW------
113 NP_619548.1 ---------EERYFLITEKYQKIEQLLDKVCSENPIDPIKSKQW------
114 NP_056880.1 --------------------NTAQQAEIVAVITAFEEVSQP---------
115 YP_009121747.1 -----------------FIPILKELKQQGXIGEEPLKHWRKNGE------
116 YP_009113237.2 -----------------IQPLIEELKEQGTVGENPLQHWERNRV------
117 YP_009508408.1 -----------------FGDLIQKLKEAGYVGEEPMKFWQQNKV------
118 NP_955579.1 --------------------TSAQRAELIALTQALKMAEGKK--------
119 YP_009229919.1 -----------------FKPLLERLKEQGYVGEEPMRHWSKNQV------

```
120 YP_004442833.1     ------------------MKIINDMKELGFIGEEPLKHWANNKI------
121 NP_862833.2        --------------------SSAQVVELFAVHQALLTVSTS---------
122 NP_040563.1        --------------------GTNQQAELKAICMALLDGPPK---------
123 YP_443922.1        --------------------ASAQLAEIIGLTRALTLAEGKT--------
124 NP_057933.2        --------------------TSAQRAELIALTQALKMAEGKK--------
125 YP_009116631.1     -----------------NHRLLQELREQGYIGEEPMRHWAKNGI------
126 YP_009513249.1     --------------------HTAQFAEIAAFEFALKKCLPLGGN------
127 YP_223871.1        --------------------TSAQKAELIALTKALEWSKDKS--------
128 NP_127504.1        ---------ELQQLEDVIKNQLCADSHVDFLSKCSHPLWLNQDF------
129 NP_044929.1        --------------------HTAQYAEIAAFEFAIRRATGIRGP------
130 NP_057860.1        --------------------KSAQRAELLGLLHGLSSARSWRC-------
131 NP_040973.1        --------------------GSNQKAEIQALLLALKAGSEE---------
132 NP_777317.1        -------------------RLIEEMKRLGFIGDEPLKHWRQNKV------
133 NP_054716.1        --------------------HTAQFAEIAAFEFALKQAIRKMGP------
134 YP_001856242.1     --------------------TTNQKTELQAIYLALQDSGLE---------
135 NP_954565.2        --------------------NSAQLVELQALIAVLSAFPNQP--------
136 NP_569150.1        -------------------DIIGKMKKLGYIGEEPLKHWEKNQV------
137 NP_659397.1        ---------EEKGFIEISKFREIEQLLEKVCSENPIDPEKSKGW------
138 YP_009513211.1     --------------------TSAQKAELIALTQALRLAEGKS--------
139 YP_233107.1        -------------------SEIDRLKRLGFIGEEPLRHWKRNQI------
140 YP_009109692.1     --------------------TSAQKAELIALTKALELSKGKK--------
141 NP_057849.4        --------------------TTNQKTELQAIYLALQDSGLE---------
142 NP_049560.1        --------------------KSAQRAELLGLLHGLSSAHSWRC-------
143 YP_009506267.1     -----------------YEELIVQLKNAGYIGENPMKFWAKNKV------
144 NP_039820.1        ------------------KDLLKEMKEMKYIGENPMEFWKNNKI------
145 NP_056790.1        --------------------TSAQKAELVALTQALRLAEGKN--------
146 YP_002321513.1     -----------------FTSTIEDLKAQGVIGDAPLQLWERNQV------
147 NP_955591.1        --------------------TSAQRAELIALTQALKMAEGKK--------
148 YP_001956722.2     --------------------HTAQLAEVAAVEFACKKALKIDGP------
149 NP_056902.1        --------------------VTHQVAERMAIQMALEDTRDKQ--------
150 NP_056886.1        --------------------ASVQQLEARAVAMALLLWPTTP--------
151 NP_041261.1        --------------------LSAQLVELYAILQVFTVLAHQP--------
```

```
001 YP_004442836.1     --KCRIEVKNPDMIIEDKPLKHVTPAMKETMAKHVKKLLELKVIRPS---
002 NP_056728.1        -MKASIKLSDPSKAIKVKP-MKYSPMDREEFDKQIKELLDLKVIKPS---
003 YP_009508537.1     -----ILVVTDSNYVAKAYNEELDVWASNGFVNNRKKPLKHISKWKS---
004 YP_004732983.2     --RCKLDIINPDITIEAKPPGHLTLEDKVKYQKHIDALLDLGVIRPS---
005 YP_009508407.1     -----TEAIDPPAVGYIKP-------------TEK---LALAVINLQ---
006 NP_040939.1        -----MNLVTDSRYAFEFLLRNWDEEVIKNPIQARIMEIAHKKDRIG---
007 YP_003864102.1     -----LNIYTDSAYLAHSIPLLETVAQIKHIS-DTAQLFLQCQQLIH---
008 YP_009002585.1     --LCYLDIKNPDMVIEDKPIKHVTPQMEESFRKHIKGLLELKVIRPS---
009 YP_003284237.1     --KCSLEIKNPDMIIEDRPLKHVTPKMKEQMKKHVDKLLELKVIRPS---
010 YP_009268869.1     -----MNIVTDSRYAYEFMRRNWDEEVIKNPIQARIMKLVHDKEQIG---
011 YP_009506251.1     --HCKLDIKNPDLIIEDKPIKHLTPAMEKQFAKHIKALLDIGVIRPS---
012 NP_955611.1        -----TNVVTDSAFVAKMLLKMGQEGVPSTAA------AFILEDALS---
013 YP_008567619.1     --TCKLEIKNPDLIIEDRPLKHVTPAMHASFLKHIDALLKIKVIRPS---
014 NP_149413.1        --KCKIELINPDLIIKDKPITMLTIQTKEAMRKHVDALLARGVIRKS---
015 YP_004442824.1     --RCRIEVKNPDLIIEDRPLKHVTPKMKEAMAVHIKKLLELKVIRPS---
016 YP_001931967.1     -LTASIKLIDPNKIIRVKP-IPYPPNIRQEFDIQIKELLAMNLIVPS---
017 YP_002455786.1     -----LNIFLDSKFLIRYLQSLASGAFQGSST----HHRLQASLPTL---
018 YP_009109694.1     -----ANIYTDSRYAFATAHTHGSIYERRGLLTSEGKEIKNKAEIIA---
019 NP_789739.1        --------------------------------------------------
020 NP_955619.1        -----LNIFLDSKYLYHYLRTLALGTFQGRSS----QAPFQALLPRL---
021 YP_224289.1        --KCKLELKNPDITIQDKPLKHVTAKMRETMRAHIDKLLQLKVIRPS---
022 YP_009182100.1     -TYADIQLKDPYALIRVKP-MPYSAEDEAEFERQLEEQLKLELIQPS---
023 NP_705927.1        -----VNIVTDSQYALGIIQAQ--PDQSESELVNQIIEQLIKKEKVY---
024 YP_009508577.1     -----VLIVTDSDYVARSVNEELPFWRSNGFVNNKKKPLKHISKWKN---
025 YP_004442830.1     --KCHIRIKNPELTIQDKPHKFVTPQMKEQMQKHMDELLKRKVIRPS---
026 NP_758887.1        -----VNIVTDSQYAMGILSAA--PDQSDNPIVREIIELMIHKEAVY---
027 YP_009508406.1     --------------------------------------------------
028 YP_610965.1        --TCQLEIINPDLTIQDKPLKHVTPLMEQQFKRHVEALLQLKVIRPS---
029 YP_001931961.1     -MKASIKLIDPKTVVKVKP-MRYNPQDVEEFAKQIKELLELKIIIPS---
030 YP_241114.1        -----FNLFTDSSYVVGALQMIETVPIIGTTSPEVFNLFTLIQQVLH---
031 YP_006732334.1     -MTASIKLADPKSVVKVKP-MVYSPQDRKEFEIQIKELLNLKVIIPS---
032 YP_009041481.1     --ICRLDIINPDITIQSQPLKHVTVEMERSFQTHVDGLLKLKVIRPS---
033 YP_001036293.1     --VCELNIINPDLTIQDKPLKHVTPAMEETFRKHIDALLKLKVIRAS---
034 YP_605811.1        --KCKIEIKNPDLIIEDRPLKHVTPAMKESMKKHIDKLLELKVIRPL---
035 NP_955577.1        -----LTVYTDSRYAFATTHVHGEIYRRRGLLTSEGKEIKNKNEILA---
036 YP_003208050.1     --KCRLEIINPDLTIQDKPLKHVTPQMEAQFKRHTDALLQLGVIRPS---
037 NP_789740.1        -----VNIVTDSQYALGIIQAQ--PDQSESELVNQIIEQLIKKEKVY---
038 YP_004442839.1     --RCRIEVKNPDLIIEDRPLKHVTPAMKESMKRHVDKLLELKVIRPS---
```

039 YP_009507791.1 -----MNIITDSQYIMNVLKQ--YPDSLSG-LWQKIIELLQSKIKIF---
040 YP_004581513.1 --YCKLQIKNTDLIIQDPPLKHVTPAAREFFKSQISDLLKAKLIRPS---
041 NP_043933.1 -FEAQIALLDTNKIIKCKP-MQYSPQDREEFKTQIEELLKLGIIRPS---
042 YP_009507248.1 -----LNRFHRQIHLTVVGDIKRRPIVTKPFESHTRIAIQLSESNLEQDV
043 YP_002117531.1 --KCKLDIINPNLTIEDKPLKHVTPAMKEQMRKHIDKLLELKVIRPS---
044 YP_004347415.1 --YAKIELKNPDDHIYKPP-INYQESDYKEFKMHIDEMVKEGFIEECKNL
045 NP_056891.1 -----LNIYTDSAYLAHSIPLLETVAQIKHIS-ETAKLFLQCQQLIY---
046 YP_009513242.1 -----VLIVTDSFYVAESANKELSYWKSNGFLNNKKKPLKHVSKWKS---
047 YP_009352866.1 --ICKLEIINPDITIQAQPLKHVTAEMEKSFKTQVDGLLKLRVIRPS---
048 YP_009165750.1 -MTASIKLKDPNTLVQVKP-MQYCPEDRKEFAVQIKELLDLKIIIPS---
049 NP_040840.1 -----MNIVTDSRYAYEFMLRNWDEEVIRNPIQARIMELVHNKEKIG---
050 YP_004222728.1 -----TNVVTDSAFVAKMLLKMGQKGVPSTAA------AFILEDALS---
051 YP_009506264.1 --KCRLEIKNPDMIIEDRPLKHVTPKLKEDMQKHIDQLLKLKVIRPS---
052 YP_567050.1 -----LNIFLDSKYLIKYLHSLAIGAFLGTST----HQSLYAHLPTL---
053 YP_009508571.1 -----VLIITDSFYVAESTNKELPYWKSNGFVNNKKKPLKHVSKWKS---
054 NP_056803.1 -----VLVITDSFYVAESANKELPYWKSNGFVNNKKEPLKHISKWKS---
055 NP_955564.1 -----FNLYTDSKYVTGLFPEIETATLSPRTK--IYTELKHLQRLIH---
056 YP_009508411.1 --MCKLDIINPNLIIQDKPLKHVTPAMTETFKKHIDALLELKVIRPS---
057 YP_009140788.1 --LCQLDIKNPDFIIEDRPLKSVTPQMKESFKRHVKALLDLKVIRPS---
058 YP_009508410.1 -----VESLEPPALGFIKP-------------ADFRAGLATASAAVQ---
059 NP_040333.1 -----LNVYTDSRYAFATAHIHGEIYRRRGLLTSEGKEIKNKDEILA---
060 YP_008992013.1 -FVKLPFKLNEDINPTKATHPGMSPSDLLLAQQECSQLLAQGLIEPT---
061 NP_041734.1 --LCRLDIKNPDLIIEDKPIKHLTPAMEKQFQKHIKALLDIGVIRPS---
062 NP_056895.1 -----LNIWVDSKYLYSLLRTLVLGAWLQPDP----VPSYALLYKSL---
063 YP_009109689.1 -----ANIYTDSRYAFATAHTHGSIYERRGLLTSEGKEIKNKAEIIA---
064 YP_002519387.1 -LLAEIKLINPATTVNVKS-MAYSPDDAVEINKQIQELLEIKVIRPS---
065 YP_595725.1 --KCKIELRNPDLIIKDKPQTLLNIQKKEAMRKHIDALLERKVIRPS---
066 YP_009508551.1 -----VLVITDSFYVAESANKELPYWKSNGFVNNKKKPLKHISKWKS---
067 YP_001497148.1 -----ANIYTDSRYAFATAHTHGSIYERRGLLTSEGKEIKNKAEIIA---
068 NP_041186.1 -----FNLFTDSSYVVGALQMIETVPIIGTTSPEVLNLFTLIQQVLH---
069 NP_663784.1 -----VNIIVDSQYVMGIVAGQ--PTESENRIVNQIIEEMIKKEAVY---
070 YP_009508443.1 --TCKLEIINPELTIQDKPLKHVTPSMEEAFRKHVQALLKLKVIRPS---
071 YP_003987465.1 --ICELQIKNPELTIEDRPLKHVTPALKEAMQKHVDKLLELKVIRPS---
072 NP_861410.1 -ELVKIELINPEKEVNVPNNIPYSLRDINEFSQECADLVRKGIIEES---
073 YP_009345071.1 --ECKLEIINPQLKIEDKPLKHITPQMQESFNKQVESLLKLGVIRPS---
074 YP_009508582.1 -----VLIVTDSDYVARSVNEELPFWRSNGFVNNKKKPLKHISKWKN---
075 NP_569141.1 --KVKIELIEENSIITQKP-LKYNFNDLTEFKIHIKDLLDNKYIQES---
076 YP_002916057.1 --ECTLRIKNPDLVIQDPPLKHVTPAAREFFQNQVSSLIKAQLIRPS---
077 NP_777384.2 -----LNIWVDSKYLYSLLRTLVLGAWLQPDP----VPSYALLYKSL---
078 YP_233110.1 --KCRIEIKNPDLIIEDRPLKHVTPAMKESMKKHVDKLLELKVIRPS---
079 NP_542258.1 -----LNIFLDSKFLIGHLRRMALGAFLGPST----QCDLHARLFPL---
080 YP_009130664.1 --TCQLEIKNPDLTIEDRPLKHISPQMEASYRRHTEAVLKLGTIRPS---
081 YP_009508566.1 -----VLIVTDSDYVARSVNNELNFWRSNGFVNNKKKPLKHISKWKS---
082 YP_006607892.1 -MTASIKLKDPNTIVKVKP-MQYNPDDRMEFSKQIKELLDLKIIIPS---
083 YP_009506270.1 --ECRIEVKNPDMIIEDEPLKHVTPAMKETMAKHVKKLLELKVIRPS---
084 NP_042513.1 -IIVHIDLIDPTKEVNVPNRIPYTQKDIDEFREETSKQIELGILRQS---
085 YP_006907834.1 -MQASIKLSDPTKVIKVKP-MKYSPMDREEFEKQIQELLDLKVIRPS---
086 YP_004442827.1 --KCHIRIKNPELTIQDKPQKMVTPQIKEQMKKHMDELLLRRVIRPS---
087 YP_009408594.1 --KCHIDIINPDITIQDPPLKHVTPALKETFQKHIDALLKLGVIRES---
088 YP_009508561.1 -----VLIVTDSDYVARSANKELPFWRSNGFLNNKKKPLKHISKWKN---
089 NP_056907.1 -----LNIFLDSKYLIKYLHSLAIGAFLGTST----HQSLYAHLPTL---
090 YP_002308474.1 -----LNIFLDSKFLVGQLRRLALGAFIGPST----QCDLHSQLLPL---
091 YP_007761644.1 LPQDQLDRLKLKCPVRCKP-IGANPIDMEEFHNQINELLRLKLIRKT---
092 NP_040550.1 -----TNVVTDSAFVAKMLLKMGQEGVPSTAA------AFILEDALS---
093 NP_758808.1 --TCKLDLKNTEITIQDKPLRHITPALEQSYGRHVNALLMLKVIQPS---
094 YP_009345075.1 --VCKIDIINPDLTIEDRPLKHITPEMKAQFSRHTEALLKLGVIRKS---
095 NP_068729.1 -LLIEIRLKDPLQEINVTNRIPYTIRDVQEFKEECEDLLKKGLIRES---
096 NP_047255.1 -----LTVYTDSRYAFATTHVHGEIYRRRGLLTSEGKEIKNKNEILA---
097 YP_006273075.1 --LCQLDIKNPDFIIEDRPLKNLTPQMKESFKKHIKVLLDLGVIRAS---
098 YP_009508888.1 -----VLIITDSFYVAESANKELPYWKSNGFVNNKKKPLKHVSKWKS---
099 NP_395469.1 -ELIEIKLKDPNAEIFVPNNIPYTQRDIEEFKEDMEDLINKGLIRPS---
100 YP_009508546.1 -----VLIITDSFYVAESANKELPYWKSNGFVNNKKKPLKHISKWKS---
101 NP_056762.1 -----FKIINPDINITCAT-IPYTPADKEVFEKQIKELLDNKLIKKADP-
102 YP_009508556.1 -----VLIVTDSFYVAESANKELSYWKSNGFLNNKKKPLRHVSKWKS---
103 NP_569153.1 --VCHLDIKNPDMVIEDRPLKHVTPQMEESFRKHVEALLKIGAIRPS---
104 NP_043924.1 DKIKPYLGKGQQLTLCDTYIGKEGAAILGQLRPDMQALHQAEGEVHVSLG
105 YP_009243641.1 -----INIIADSAYVVGVVKNIIG-AVINSLDKELTTLFKTLQVLLT---
106 YP_004300274.1 --EVKIELINPDSIVYQKP-LRWNFEDIEEFKLHIDELLKGGFIRPS---
107 NP_056848.1 --LAKIELKNETDNIYKPP-MLYQETDLPEFKMHIEEMIKEGFIEEKTNF
108 YP_006495799.1 --KCKLDIINPDITIQGKPPSTATPEIKDRYQRHIDALLSIGVIRPS---
109 YP_009508409.1 --------------------------------------------------
110 NP_687035.1 -----VNIVTDSQYVMNILTAC--PQESNSPLVEQIIQALMKKRQVY---
111 NP_045937.2 -------IYTDSRYAYGVVHDFGHLWMHRGFVTSAGTPIKNHKEIEYLLK

```
112 NP_612577.1        -MTATIELIDPKTVVKVKP-MSYSPSDREEFDRQIKELLELKVIKPS---
113 NP_619548.1        -MKASIKLIDPLKVIRVKP-MSYSPQDREGFAKQIKELLDLGLIIPS---
114 NP_056880.1        -----FNLYTDSKYVTGLFPEIETATLSPRTK--IYTELKHLQRLIH---
115 YP_009121747.1     --TCKLDIINPDITVQDKPLKHVTPALEASFKKXIEALLKLKVIRPS---
116 YP_009113237.2     --HCYLDIKNPDLTVQDKPLDQITPVQKEMYKKHIDALLQIGVIRRS---
117 YP_009508408.1     --TCKLDIINPELRIQDKPLKHVTPAMEETFRKHVQALLKLKVIRPS---
118 NP_955579.1        -----LTVYTDSRYAFATTHVHGEIYRRRGLLTSEGKEIKNKNEILA---
119 YP_009229919.1     --KCKLEIINPDITIQDKPMKHVTPAMKDQFQKHTQALLKLGVIRPS---
120 YP_004442833.1     --KCHIRIRNPELTIQDKPHKFVTPQMKEQMQFHMNELLKRKVIRPS---
121 NP_862833.2        -----FNLFTDSSYVVGALQMIETVPIIGTTSPEVLNLFTLIQQALH---
122 NP_040563.1        -----MNIITDSRYAYEGMREE-PETWAREGIWLEIAKILPFKQYVG---
123 YP_443922.1        -----VNIYTDSAYAHGAVHIDGPQWLRRNFTTTGNLPIKHKTQMEV---
124 NP_057933.2        -----LNVYTDSRYAFATAHIHGEIYRRRGLLTSEGKEIKNKDEILA---
125 YP_009116631.1     --KCKLDIKNPDIVISSKPPDSVSKETKAQYQRHIDALLKIGVIQPS---
126 YP_009513249.1     -----ILVVTDSNYVAKAYNEELDVWASNGFVNNRKKPLKHISKWKS---
127 YP_223871.1        -----VNIYTDSRYAFATLHVHGMIYRERGLLTAGGKAIKNAPEILA---
128 NP_127504.1        -FIQLPFKKNENINPTKASHSGMNPEHLQLAIKECDELQQFDLIEPS---
129 NP_044929.1        -----VLIVTDSNYVAKSYNEELPYWESNGFVNNKKKTLKHISKWKA---
130 NP_057860.1        -----LNIFLDSKYLYHYLRTLALGTFQGRSS----QAPFQALLPRL---
131 NP_040973.1        -----MNIITDSQYVINIILQ--QPDMMEG-IWQEVLEELEKKTAIF---
132 NP_777317.1        --TCKLEIKNPDLIIEDKPLKHVTPKMKEVMARHVTALLQSKVIRPS---
133 NP_054716.1        -----VLIVTDSDYVAKSYNQELDFWVSNGFVNNKKKPLKHVSKWKS---
134 YP_001856242.1     -----VNIVTDSQYALGIIQAQ--PDQSESELVNQIIEQLIKKEKVY---
135 NP_954565.2        -----LNIYTDSAYLAHSIPLLETVAQIKHIS-ETAKLFLQCQQLIY---
136 NP_569150.1        --KCRIEVKNPDMIIEDRPLKHVTPTMKETMAKHVQKLLELKVIRPS---
137 NP_659397.1        -MKAEIKLIKPETVVRVKP-MVYSPEDRKEFEIQIKELLDLKVIIPS---
138 YP_009513211.1     -----INIYTDSRYAFATAHVHGAIYKQRGLLTSAGKDIKNKEEILA---
139 YP_233107.1        --KCKLEIKNPDLIIEDRPLKHVTPAMKETMTKHVQRLLDIKVIRPS---
140 YP_009109692.1     -----ANIYTDSRYAFATAHTHGSIYERRGLLTSEGKEIKNKAEIIA---
141 NP_057849.4        -----VNIVTDSQYALGIIQAQ--PDQSESELVNQIIEQLIKKEKVY---
142 NP_049560.1        -----LNIFLDSKYLYHYLRTLALGTFQGRSS----QAPFQDPLPRL---
143 YP_009506267.1     --QCKLEIINPNLTIQDKPLKHVTPAMEATFKKHVDALLELKVIRPS---
144 NP_039820.1        --KCKLNIINPDIKIMGRPIKHVTPGDEEAMTRQINLLLQMKVIRPS---
145 NP_056790.1        -----INIYTDSRYAFATAHIHGAIYKQRGLLTSAGKDIKNKEEILA---
146 YP_002321513.1     --KCKLEVINPDITISDKPLKHVSIGLKQQFQNQLDPLLKMGLIRPS---
147 NP_955591.1        -----LNVYTDSRYAFATAHIHGEIYRRRGLLTSEGKEIKNKDEILA---
148 YP_001956722.2     -----VLIVTDSFYVAESVNKELPYWQSNGFFNNKKKPLKHVSKWKS---
149 NP_056902.1        -----VNIVTDSYYCWKNITEGLGLEGPQS-PWWPIIQNIREKEIVY---
150 NP_056886.1        -----TNVVTDSAFVAKMLLKMGQEGVPSTAA------AFILEDALS---
151 NP_041261.1        -----FNLYTDSAYIAQSVPLLETVPFIKSST-NATPLFSKLQQLIL---
```

```
001 YP_004442836.1     -QSKHRTTAMIVESGTEIDPVTGK--------------------------
002 NP_056728.1        -KSPHMAPAFLVNNEAEKR-------------------------------
003 YP_009508537.1     -VADFKKLRPDVVVTHEPG-------------------------------
004 YP_004732983.2     -KSRHRSAAFIVASGTSVDPKTGK--------------------------
005 YP_009508407.1     -KQNNTLTNLITQIFEKLT-------------------------------
006 NP_040939.1        -VHWVPGHKGIPQN---EEIDKYISEIFLAKEGEGILPKREEDAGYDLIC
007 YP_003864102.1     -NRSTPIYIGHVRA------------------------------------
008 YP_009002585.1     -TSKHRTTAFIVNSGTSVDPVTGK--------------------------
009 YP_003284237.1     -TSKHRTTAMIVESGTEIDPKTGQ--------------------------
010 YP_009268869.1     -VHWVPGHKGIPQN---EEIDKYISEIFLAREGSGILPKRAEDAGYDLIC
011 YP_009506251.1     -KSRHRTTAFIVESGTTIDPKTKK--------------------------
012 NP_955611.1        -QRSAMAAVLHVRS------------------------------------
013 YP_008567619.1     -KSRHRTCAFIVNSGTSIDPVTGK--------------------------
014 NP_149413.1        -NSPHRTNAFIVESGTSVDPVTKK--------------------------
015 YP_004442824.1     -TSKHRTTAMIVESGTEVDPVTGI--------------------------
016 YP_001931967.1     -KSPHMSPAFMVNKGAEQR-------------------------------
017 YP_002455786.1     -LQGKVVYLHHTRS------------------------------------
018 YP_009109694.1     -LLKALFLPQEVAIIHCPG-------------------------------
019 NP_789739.1        --------------------------------------------------
020 NP_955619.1        -LSRKVVYLHHVRS------------------------------------
021 YP_224289.1        -SSRHRTTAMIVESGTEVDPKTGQ--------------------------
022 YP_009182100.1     -KSPHSSPAFCVRNHAEIK-------------------------------
023 NP_705927.1        -LAWVPAHKGIGGN------------------------------------
024 YP_009508577.1     -ISDSLLLKRDIIIVHEPG-------------------------------
025 YP_004442830.1     -SSRHRTNAFIVNSGTTVDPLTKK--------------------------
026 NP_758887.1        -LAWVPAHKGIGGN------------------------------------
027 YP_009508406.1     --------------------------------------------------
028 YP_610965.1        -KSRHRTMAMIGNSGTSVDPTTGK--------------------------
029 YP_001931961.1     -KSPHQSPAFLVENEAERR-------------------------------
030 YP_241114.1        -CRQYPCFFGHIRA------------------------------------
```

```
031 YP_006732334.1  -KSQHMSPAFLVEKEAEKR-------------------------------
032 YP_009041481.1  -KSRHRTLAILVKSGTSIDPLTGK--------------------------
033 YP_001036293.1  -KSRHRTTAFIVYSGTTVDPVTGK--------------------------
034 YP_605811.1     -TSKHRTTAIIVQSGTEIDPLTGK--------------------------
035 NP_955577.1     -LLEALFLPKRLSIIHCPG-------------------------------
036 YP_003208050.1  -KSRHRTMAIMVQSGTTVDPATGK--------------------------
037 NP_789740.1     -LAWVPAHKGIGGN------------------------------------
038 YP_004442839.1  -ESKHRTTAIIVQSGTEIDPLTGK--------------------------
039 YP_009507791.1  -LDWVPGHKNIPGN---VEVDR-ICQENMIIEGDGIIDKREEDAGYDLIA
040 YP_004581513.1  -KSKHRTTAFMVESGTIVDPKTGK--------------------------
041 NP_043933.1     -KSPHSSPAFMVRNHAEIK-------------------------------
042 YP_009507248.1  ISAIKEYVNPKVKTALIIN-------------------------------
043 YP_002117531.1  -KSRHRTTAMIVQSGTEMVLQKDKNGKEKMV-------------------
044 YP_004347415.1  ENKKYSSPAFIVNKHSEIK-------------------------------
045 NP_056891.1     -NRSIPFYIGHVRA------------------------------------
046 YP_009513242.1  -IAECLQLKPDITIIHEKG-------------------------------
047 YP_009352866.1  -KSRHRTLALLVKSGTSIDPLTGK--------------------------
048 YP_009165750.1  -KSPHRSPAFLVENEAERR-------------------------------
049 NP_040840.1     -VHWVPGHKGIPQN---EEIDRYISEIFLAKEGRGILQKRAEDAGYDLIC
050 YP_004222728.1  -QRSAMAAVLHVRS------------------------------------
051 YP_009506264.1  -ASKHRTTAMLVESGTEVDPKTGL--------------------------
052 YP_567050.1     -LHNKVIYLHHIRS------------------------------------
053 YP_009508571.1  -IAECLSLKPDITIQHERG-------------------------------
054 NP_056803.1     -IAECLSIKPDITIQHEKG-------------------------------
055 NP_955564.1     -KRQEKFYIGHIRG------------------------------------
056 YP_009508411.1  -TSKHRTTAFIVHSGTTVDPETGL--------------------------
057 YP_009140788.1  -KSRHRTTAMLVNSGTTVDPKTGK--------------------------
058 YP_009508410.1  -KQNNTIIQLLLQLSEKVN-------------------------------
059 NP_040333.1     -LLKALFLPKRLSIIHCPG-------------------------------
060 YP_008992013.1  -TSQWACQAFYVEKHSEIV-------------------------------
061 NP_041734.1     -KSKHRTTAFIVESGTVIDPVTKK--------------------------
062 NP_056895.1     -LRHPAIFVGHVRS------------------------------------
063 YP_009109689.1  -LLKALFLPQEVAIIHCPG-------------------------------
064 YP_002519387.1  -RSPHSSPCFLVQNHNEIK-------------------------------
065 YP_595725.1     -KSPHRTNAFIVESGTSIDPKTGK--------------------------
066 YP_009508551.1  -IAECLSMKPDITIQHEKG-------------------------------
067 YP_001497148.1  -LLKALFLPRRVAIIHCPG-------------------------------
068 NP_041186.1     -CRQHPCFFGHIRA------------------------------------
069 NP_663784.1     -VAWVPAHKGIGGN------------------------------------
070 YP_009508443.1  -TSRHRTTAFIVHSGTTVDPKTGV--------------------------
071 YP_003987465.1  -TSRHRTTAMIVYSGTEVDPVTKK--------------------------
072 NP_861410.1     -KSPHSAPAFYVENHNEIK-------------------------------
073 YP_009345071.1  -KSQHRTTAFLVKSGTTVDPKTGK--------------------------
074 YP_009508582.1  -ISDSLLLKRDITIVHEPG-------------------------------
075 NP_569141.1     -NSKHTSPAFIVNKHSEQK-------------------------------
076 YP_002916057.1  -RSRHRTTAFMVESGTSVDPKTGK--------------------------
077 NP_777384.2     -LRHPAIFVGHVRS------------------------------------
078 YP_233110.1     -TSKHRTTAIIVQSGTEIDPLTGK--------------------------
079 NP_542258.1     -LQGKTVYVHHVRS------------------------------------
080 YP_009130664.1  -KSKHRTTAIIVNSGTTIDPITGK--------------------------
081 YP_009508566.1  -ISESLLLHKNITIVHEPG-------------------------------
082 YP_006607892.1  -KSPHQSPAFLVENEAERR-------------------------------
083 YP_009506270.1  -QSKHRTTAIIVQSGTEIDPVTGK--------------------------
084 NP_042513.1     -KSPHSAPAFYVENHNEIK-------------------------------
085 YP_006907834.1  -KSPHMAPAFLVNNEAEKR-------------------------------
086 YP_004442827.1  -TSRHRTNAFIVNSGTSIDPITMK--------------------------
087 YP_009408594.1  -SSRHRTMAMIVKSGTTVDPITGQ--------------------------
088 YP_009508561.1  -ISDSLLLKRNITIVHEPG-------------------------------
089 NP_056907.1     -LHNKVIYLHHIRS------------------------------------
090 YP_002308474.1  -LYNKTIYVHHVRS------------------------------------
091 YP_007761644.1  -NSPWSFQAFMVRNHAEIV-------------------------------
092 NP_040550.1     -QRSAMAAVLHVRS------------------------------------
093 NP_758808.1     -KSRHRTMAFLVNSGTTVT-ADGK--------------------------
094 YP_009345075.1  -SSRHRTNAMIVRSGTSVDPKTGE--------------------------
095 NP_068729.1     -QSPHSAPAFYVENHNEIK-------------------------------
096 NP_047255.1     -LLEALFLPKRLSIIHCPG-------------------------------
097 YP_006273075.1  -KSRHRTTAMLVNSGTTVDPKTGK--------------------------
098 YP_009508888.1  -IADCLSLKTGITIKHEKG-------------------------------
099 NP_395469.1     -KSPHSAPAFYVENHSEIK-------------------------------
100 YP_009508546.1  -IAECLSLKPDIVIQHEKG-------------------------------
101 NP_056762.1     -TCRHRTAAFIVRNHSEEVAQ-----------------------------
102 YP_009508556.1  -IAECLQLKPDITIIHEKG-------------------------------
103 NP_569153.1     -KSRHRTTAIIVNSGTSIDPITGK--------------------------
```

```
104 NP_043924.1        TRAGHCPQELGTMLTNLLKSTQER--------------------------
105 YP_009243641.1     -SRNARIFIVHIRS------------------------------------
106 YP_004300274.1     -NSKHSSPAFIVNKHSEQK-------------------------------
107 NP_056848.1        EDKKYSSPAFIVNKHSEQK-------------------------------
108 YP_006495799.1     -KSRHRTAAFITYSGTSVDPKTGE--------------------------
109 YP_009508409.1     --------------------------------------------------
110 NP_687035.1        -LQWVPAHKGIGGN------------------------------------
111 NP_045937.2        QIMKPKQVSVIKIEAHTKGVSMEVRG------------------------
112 NP_612577.1        -KSTHMSPAFLVENEAERR-------------------------------
113 NP_619548.1        -KSQHMSPAFLVENEAERR-------------------------------
114 NP_056880.1        -KRQEKFYIGHIRG------------------------------------
115 YP_009121747.1     -KSRHRTMAMIVNSGTTVDPATGK--------------------------
116 YP_009113237.2     -NSRHRTNAFIVHSGTTVDPRTGE--------------------------
117 YP_009508408.1     -TSKHRTTAFIVHSGTTIDPKTGV--------------------------
118 NP_955579.1        -LLEALFLPKRLSIIHCPG-------------------------------
119 YP_009229919.1     -KSRHRTMAMIVYSGTSVDEKTGK--------------------------
120 YP_004442833.1     -TSRHRTNAFIVNSGTTVDPITRK--------------------------
121 NP_862833.2        -CRQHPCFFGHIRA------------------------------------
122 NP_040563.1        -VGWVPAHKGIGGN---------------------------TEADEGVKKA
123 YP_443922.1        -LISAVALPKKVAIMKCKG-------------------------------
124 NP_057933.2        -LLKALFLPKRLSIIHCPG-------------------------------
125 YP_009116631.1     -KSKHRTAAFITHSGTSIDPITKK--------------------------
126 YP_009513249.1     -VADLKRLRPDVVVTHEPG-------------------------------
127 YP_223871.1        -LLTAVWLPKRVAVMHCKG-------------------------------
128 NP_127504.1        -DSQWACEAFYVNKRSEQV-------------------------------
129 NP_044929.1        -IAECKNLKADIHVIHEPG-------------------------------
130 NP_057860.1        -LSRKVVYLHHVRS------------------------------------
131 NP_040973.1        -IDWVPGHKGIPGN---EEVDK-LCQTMMIIEGDGILDKRSEDAGYDLLA
132 NP_777317.1        -TSKHRTTAIIVESGTEVDPITGK--------------------------
133 NP_054716.1        -IADCKKHKADIHVIHEPG-------------------------------
134 YP_001856242.1     -LAWVPAHKGIGGN------------------------------------
135 NP_954565.2        -NRSIPFYIGHVRA------------------------------------
136 NP_569150.1        -SSKHRTTAMIVESGTEVDPMTGK--------------------------
137 NP_659397.1        -KSQHMSPAFLVEKEAEKR-------------------------------
138 YP_009513211.1     -LLEAIHLPKRVAIIHCPG-------------------------------
139 YP_233107.1        -TSKHRTTAIMVNSGTEIDPITGA--------------------------
140 YP_009109692.1     -LLKALFLPQEVAIIHCPG-------------------------------
141 NP_057849.4        -LAWVPAHKGIGGN------------------------------------
142 NP_049560.1        -LSRKVVYLHHVRS------------------------------------
143 YP_009506267.1     -TSQHRTTAFIVYSGTTVDPATGK--------------------------
144 NP_039820.1        -ESKHRSTAFIVRSGTEIDPITGK--------------------------
145 NP_056790.1        -LLEAIHLPRRVAIIHCPG-------------------------------
146 YP_002321513.1     -TSRHRTMAMIINSGTTVDPVTGE--------------------------
147 NP_955591.1        -LLKALFLPKRLSIIHCPG-------------------------------
148 YP_001956722.2     -IADCIQLKPDIIIIHEKG-------------------------------
149 NP_056902.1        -FAWVPGHKGIYGNQLADEAAKIKEEIMLAYQGTQIKEKRDEDAGFDLCV
150 NP_056886.1        -QRSAMAAVLHVRS------------------------------------
151 NP_041261.1        -NRQHPFFIGHLRA------------------------------------
```

```
001 YP_004442836.1     ----------------------------------EKRGKERLVFNYKRLN
002 NP_056728.1        ------------------------------------RGKKRMVVNYKAMN
003 YP_009508537.1     ---------------------------------------HQKLDSSPHAY
004 YP_004732983.2     ----------------------------------ETRGKERMVIDYRMLN
005 YP_009508407.1     ----------------------------------------QIQTEVRSLK
006 NP_040939.1        PEEVTIEPGQVKCIPIELRLNLKKSQWAMIATKSSMAAKGVFTQGGIIDS
007 YP_003864102.1     ----------------------------------------HSGLPGPIAQ
008 YP_009002585.1     ----------------------------------ETKGKERMVFNYKRLN
009 YP_003284237.1     ----------------------------------EKRGKERLVFNYKRLN
010 YP_009268869.1     PQEVCIPAGQVRKIPINLRINLKEDQWAMVGTKSSFASKGVFVQGGIIDS
011 YP_009506251.1     ----------------------------------TIHGKERMVFNYKRLN
012 NP_955611.1        ----------------------------------------HSEVPGFFTE
013 YP_008567619.1     ----------------------------------EVKGKERLVFNYKRLN
014 NP_149413.1        ----------------------------------EIRGKPRLVFNYKRLN
015 YP_004442824.1     ----------------------------------EKRGKERLVFNYKRLN
016 YP_001931967.1     ------------------------------------RGKMRMVVNYKALN
017 YP_002455786.1     ----------------------------------------HTQLPDPIST
018 YP_009109694.1     ---------------------------------------HQKGQDP-VAV
019 NP_789739.1        --------------------------------------------------
020 NP_955619.1        ----------------------------------------HTNLPDPISR
021 YP_224289.1        ----------------------------------EKRGKERLVFNFKRLN
022 YP_009182100.1     ------------------------------------RQKARIVINYKELN
```

023 NP_705927.1 ---------------------------------------------EQVDK
024 YP_009508577.1 ---------------------------------------HKPSYTSIHTQ
025 YP_004442830.1 ----------------------------------EIKGKPRLVFNYKRLN
026 NP_758887.1 ---------------------------------------------EQVDK
027 YP_009508406.1 --------------------------------------------------
028 YP_610965.1 ----------------------------------EVKGKERMVFNYRSLN
029 YP_001931961.1 ------------------------------------RGKKRMVVNYKAIN
030 YP_241114.1 ----------------------------------------HSTLPGALVR
031 YP_006732334.1 ------------------------------------RGKKRMVVNYKKLN
032 YP_009041481.1 ----------------------------------EVKGKERMVYDYRQLN
033 YP_001036293.1 ----------------------------------ENKGKERMVFNYKRLN
034 YP_605811.1 ----------------------------------ERRGKERLVFNYKRLN
035 NP_955577.1 ---------------------------------------HQKGDSP-QAK
036 YP_003208050.1 ----------------------------------ETRGKERMVYNYRSLN
037 NP_789740.1 ---------------------------------------------EQVDK
038 YP_004442839.1 ----------------------------------EKRGKERLVFNYKRLN
039 YP_009507791.1 QEDVFLMKGEVRIVPINAKIMLPEGTWGLIIGKSSIGKLGLDVLGGVIDQ
040 YP_004581513.1 ----------------------------------EIRGKQRMVYNYKRLN
041 NP_043933.1 ------------------------------------RGKARMVINYKKLN
042 YP_009507248.1 ----------------------------------------PPLKMYSIIP
043 YP_002117531.1 ----------------------------------EKRGKERLVFNYKRLN
044 YP_004347415.1 ------------------------------------RGKSRMVIDYKDLN
045 NP_056891.1 ----------------------------------------HSGLPGPIAQ
046 YP_009513242.1 ---------------------------------------HQQPMTTLHTE
047 YP_009352866.1 ----------------------------------EVKGKERMVYDYRQLN
048 YP_009165750.1 ------------------------------------RGKKRMVVNYKKLN
049 NP_040840.1 PQEISIPAGQVKRIAIDLKINLKKDQWAMIGTKSSFANKGVFVQGGIIDS
050 YP_004222728.1 ----------------------------------------HSEVPGFFTE
051 YP_009506264.1 ----------------------------------EKKGKQRLVFNYKRLN
052 YP_567050.1 ----------------------------------------HTNLPDPIST
053 YP_009508571.1 ---------------------------------------HQPIYTSIHTE
054 NP_056803.1 ---------------------------------------HQPINTSIHTE
055 NP_955564.1 ----------------------------------------HTGLPGPLAQ
056 YP_009508411.1 ----------------------------------EKKGKERLVFNYKRLN
057 YP_009140788.1 ----------------------------------ETKGKERMVFNYKRLN
058 YP_009508410.1 ----------------------------------------SLQAEIRTLR
059 NP_040333.1 ---------------------------------------HQKGNRA-EAR
060 YP_008992013.1 ------------------------------------RGKKRLVIDYQPLN
061 NP_041734.1 ----------------------------------TIHGKERLVFNYKRLN
062 NP_056895.1 ----------------------------------------HSSASHPIAS
063 YP_009109689.1 ---------------------------------------HQKGQDP-VAV
064 YP_002519387.1 ------------------------------------RGKKRLVINYKALN
065 YP_595725.1 ----------------------------------EIRGKPRLVFNYKRLN
066 YP_009508551.1 ---------------------------------------HQPTNTSIHTE
067 YP_001497148.1 ---------------------------------------HQKGQDP-IAT
068 NP_041186.1 ----------------------------------------HSTLPGALVQ
069 NP_663784.1 ---------------------------------------------QEVDH
070 YP_009508443.1 ----------------------------------ETKGKERMVFNYRRLN
071 YP_003987465.1 ----------------------------------EKRGKERLVFNYKRLN
072 NP_861410.1 ------------------------------------RKKRRMVINYKALN
073 YP_009345071.1 ----------------------------------ETKGKERMVFNYQRLN
074 YP_009508582.1 ---------------------------------------HQPSHTSIHTQ
075 NP_569141.1 ------------------------------------RGKSRMVIDYRNLN
076 YP_002916057.1 ----------------------------------EVRGKQRMVLNYKRLN
077 NP_777384.2 ----------------------------------------HSSASHPIAS
078 YP_233110.1 ----------------------------------EKRGKERLVFNYKRLN
079 NP_542258.1 ----------------------------------------HTLLQDPISR
080 YP_009130664.1 ----------------------------------EVRGKERMVFNYKRLN
081 YP_009508566.1 ---------------------------------------HQPSSTSVHTQ
082 YP_006607892.1 ------------------------------------RGKKRMVVNYKKIN
083 YP_009506270.1 ----------------------------------EKRGKERLVFNYKRLN
084 NP_042513.1 ------------------------------------RGKRRLVINYK-MN
085 YP_006907834.1 ------------------------------------RGKKRMVVNYKAMN
086 YP_004442827.1 ----------------------------------EVKGKPRLVFNYKRLN
087 YP_009408594.1 ----------------------------------EQKGKERMVFNYRTLN
088 YP_009508561.1 ---------------------------------------HQPSKTSIHTL
089 NP_056907.1 ----------------------------------------HTNLPDPIST
090 YP_002308474.1 ----------------------------------------HTLLQDPISR
091 YP_007761644.1 ------------------------------------RGKARMVINYKPLN
092 NP_040550.1 ----------------------------------------HSEVPGFFTE
093 NP_758808.1 ----------------------------------EIKGKERMVFNYKALN
094 YP_009345075.1 ----------------------------------FRHGKERMVFDYRKLN
095 NP_068729.1 ------------------------------------RGKRRMVINYKKMN

```
096 NP_047255.1       ---------------------------------------HQKGDSP-QAK
097 YP_006273075.1    ----------------------------------EIKRKERMVFNYKRLN
098 YP_009508888.1    ---------------------------------------HQPSHTSVHTE
099 NP_395469.1       ------------------------------------RAKRRIVINYKAMN
100 YP_009508546.1    ---------------------------------------HQPMNTSIHTE
101 NP_056762.1       --------------------------------------KPRIVYNYKRLN
102 YP_009508556.1    ---------------------------------------HQQPMTTLHTE
103 NP_569153.1       ----------------------------------EVKGKERMVFNYKRLN
104 NP_043924.1       --------------------------------------IWQDPPVFKLHD
105 YP_009243641.1    ----------------------------------------HSKLPGPLVY
106 YP_004300274.1    ------------------------------------RGKSRMVIDYRNLN
107 NP_056848.1       ------------------------------------RGKTRMVIDYKDLN
108 YP_006495799.1    ----------------------------------EIRGKERMVFDYRALN
109 YP_009508409.1    --------------------------------------------------
110 NP_687035.1       ---------------------------------------------TEIDK
111 NP_045937.2       -----------------------------------NAAADEAAKNAVFLV
112 NP_612577.1       ------------------------------------RGKKRMVVNYKAMN
113 NP_619548.1       ------------------------------------RGKKRMVVNYKAIN
114 NP_056880.1       ----------------------------------------HTGLPGPLAQ
115 YP_009121747.1    ----------------------------------ETKGXXRMVFNYRTLN
116 YP_009113237.2    ----------------------------------ETKGKERMVFNYKRLN
117 YP_009508408.1    ----------------------------------EVKGKERLVFNYKRLN
118 NP_955579.1       ---------------------------------------HQKGDSP-QAK
119 YP_009229919.1    ----------------------------------EVKGKERMVFNYKTLN
120 YP_004442833.1    ----------------------------------ETKGKPRLVFNYKRLN
121 NP_862833.2       ----------------------------------------HSTLPGALVR
122 NP_040563.1       LEQMAPCSPPEAILLKPGEKQNLETGIYMQGLRPQSFLPRADLPVAITGT
123 YP_443922.1       ---------------------------------------HQVLNSR-ISK
124 NP_057933.2       ---------------------------------------HQKGHSA-EAR
125 YP_009116631.1    ----------------------------------EVRGKERMVFDYRSLN
126 YP_009513249.1    ---------------------------------------HQKLDSSPHAY
127 YP_223871.1       ---------------------------------------HQKDDAP-TST
128 NP_127504.1       ------------------------------------RGKLRLVINYQPLN
129 NP_044929.1       ---------------------------------------HQPAEASPHAQ
130 NP_057860.1       ----------------------------------------HTNLPDPISR
131 NP_040973.1       AKEIHLLPGEVKVIPTGVKLMLPKGYWGLIIGKSSIGSKGLDVLGGVIDE
132 NP_777317.1       ----------------------------------EKRGKERLVFNYKRLN
133 NP_054716.1       ---------------------------------------HQNDLQSPYAM
134 YP_001856242.1    ---------------------------------------------EQVDK
135 NP_954565.2       ----------------------------------------HSGLPGPIAQ
136 NP_569150.1       ----------------------------------ERRGKERLVFNYKRLN
137 NP_659397.1       ------------------------------------RGKKRMVVNYKKLN
138 YP_009513211.1    ---------------------------------------HQRGTDP-VAT
139 YP_233107.1       ----------------------------------EKKGKERLVFNYKRLN
140 YP_009109692.1    ---------------------------------------HQKGQDP-VAV
141 NP_057849.4       ---------------------------------------------EQVDK
142 NP_049560.1       ----------------------------------------HTNLPDPITR
143 YP_009506267.1    ----------------------------------EKKGKERMVFNYKRLN
144 NP_039820.1       ----------------------------------EKKGKERMVFNYKLLN
145 NP_056790.1       ---------------------------------------HQRGSNP-VAT
146 YP_002321513.1    ----------------------------------EKRGKERMVFNYKTLN
147 NP_955591.1       ---------------------------------------HQKGHSA-EAR
148 YP_001956722.2    ---------------------------------------HQPTASTFHTE
149 NP_056902.1       PYDIMIPVSDTKIIPTDVKIQVPPNSFGWVTGKSSMAKQGLLINGGIIDE
150 NP_056886.1       ----------------------------------------HSEVPGFFTE
151 NP_041261.1       ----------------------------------------HLNLPGPLAE
```

```
001 YP_004442836.1    DNTEKDQYSLPGINTIIKRIGNAKIYSKF---------------------
002 NP_056728.1       KATVGDAYNLPNKDELLTLIRGKKIFSSF---------------------
003 YP_009508537.1    GNNLADQLATQASFKVHTTK-NPKLDIEQIKAIQACQNNEKLPVGYPKQY
004 YP_004732983.2    DNCHKDQYSLPGITSIIKSLGQAKIFSKF---------------------
005 YP_009508407.1    SQASPVAASDELLDKVITKLGRLSIADKL---------------------
006 NP_040939.1       GYQGQIQVIMYNSNKIAVVIPQGRKFAQLILMDKKHGKLEPWG-------
007 YP_003864102.1    GNQLADLATKTIAF------------------------------------
008 YP_009002585.1    DLTEKDQYSLPGINTIMKRVGHAKIYSKF---------------------
009 YP_003284237.1    DNTEKDQYSLPGINTIIQRIGRSKIYSKF---------------------
010 YP_009268869.1    GYQGIIQVVVYNSNDKEVIIPQGRKFAQLILMPLIHEDLEAWG-------
011 YP_009506251.1    DNTEKDQYSLPGIHTILKRVGNKKIFSKF---------------------
012 NP_955611.1       GNDVADSQATFQA-------------------------------------
013 YP_008567619.1    DNTNKDQYSLPGINTIISKVGNSKVFSKF---------------------
014 NP_149413.1       DNTWPDQYSLPGINALLKNVARAKIFSKF---------------------
```

| | | |
|---|---|---|
| 015 | YP_004442824.1 | DNTEKDQYSLPGINTIIQRIGNAKIYSKF--------------------- |
| 016 | YP_001931967.1 | DATIGDAHNIPNRDSLMALISGKRIFSSF--------------------- |
| 017 | YP_002455786.1 | LNEYTDSLIVAPV------------------------------------- |
| 018 | YP_009109694.1 | GNRQADRVARQAAMAEVLTLATEPDNTS--HIT----------------- |
| 019 | NP_789739.1 | -------------------------------------------------- |
| 020 | NP_955619.1 | LNALTDALLITPV------------------------------------- |
| 021 | YP_224289.1 | DNTEKDQYSLPGINTIISRIGNAKVYSKF--------------------- |
| 022 | YP_009182100.1 | RVTKDDGYFLPNLETLVYQVAEARVFSKF--------------------- |
| 023 | NP_705927.1 | LVSAGIRKVL---------------------------------------- |
| 024 | YP_009508577.1 | GNNLADKLATQGSYTVNNIVKNPSLDAELEQLINGHSVK-----GYPSRY |
| 025 | YP_004442830.1 | DNTEKDQYSLPGINGLLKSVGNAKIYSKF--------------------- |
| 026 | NP_758887.1 | LVSRGVRQVLF--------------------------------------- |
| 027 | YP_009508406.1 | -------------------------------------------------- |
| 028 | YP_610965.1 | DNTHKDQYSLPGINTIIQKIGRATVYSKF--------------------- |
| 029 | YP_001931961.1 | TATIGDAHNLPNKDELLTLIRGKSIFSSF--------------------- |
| 030 | YP_241114.1 | GNHTADALTKQVF------------------------------------- |
| 031 | YP_006732334.1 | EVTIGDSHNLPNMQELITLLRGKSIFSSF--------------------- |
| 032 | YP_009041481.1 | NNTHKDQYSLPGINTIIQKVGRAKVYSKF--------------------- |
| 033 | YP_001036293.1 | DNTEKDQYSLPGINTILKRVGQSKIYSKF--------------------- |
| 034 | YP_605811.1 | DNTEKDQYSLPGINTIISRIGKSKIYSKF--------------------- |
| 035 | NP_955577.1 | GNRLADDTAKKAATETHSSLTVLPTELIEGPKRPPWEYDDSDL-----DL |
| 036 | YP_003208050.1 | DNTHKDQYSLPGINTILKKIGTSKVYSKF--------------------- |
| 037 | NP_789740.1 | LVSAGIRKVLF--------------------------------------- |
| 038 | YP_004442839.1 | DNTEKDQYSLPGINTIISRIGNSKIYSKF--------------------- |
| 039 | YP_009507791.1 | GYRGALNVILSNLTRHGVKISKGQKIAQLIILPYVTESLEKGK------- |
| 040 | YP_004581513.1 | DNTEKDQYSLPGINTIVSRISGKKIFSKF--------------------- |
| 041 | NP_043933.1 | DHTKGDGYLLPNKEQLLQRIGGKTFYSSF--------------------- |
| 042 | YP_009507248.1 | IIQKTFRSSSLNLVLTKVELENVKEYLRQ--------------------- |
| 043 | YP_002117531.1 | DNTEKDQYSLPGINTIIQRIGRSKIYSKF--------------------- |
| 044 | YP_004347415.1 | KKAKVIKHPIPNKDILINRGIKANYFSKF--------------------- |
| 045 | NP_056891.1 | GNQRADLATKIVAS------------------------------------ |
| 046 | YP_009513242.1 | GNNLADKLATQGSYVVHCNTT-PSLDAELDQLLQGHNPP-----GYPKQY |
| 047 | YP_009352866.1 | NNTHKDQYSLPGINTIIQKVGRAKVYSKF--------------------- |
| 048 | YP_009165750.1 | EVTLGDSHNLPNKDELLTLIRGKTIYSSF--------------------- |
| 049 | NP_040840.1 | GYQGTIQVVIYNSNNKEVVIPQGRKFAQLILMPLIHEELEPWG------- |
| 050 | YP_004222728.1 | GNDVADSQATFQA------------------------------------- |
| 051 | YP_009506264.1 | DNTEKDQYSLPGINTIIQRIGRSKIYSKF--------------------- |
| 052 | YP_567050.1 | LNEYTDSLIIAPL------------------------------------- |
| 053 | YP_009508571.1 | GNALADKLATQGSYVVNNNDKKPNLDAELDHLIQGKYPK-----GYPKQY |
| 054 | NP_056803.1 | GNALADKLATQGSYVVNCNTKKPNLDAELDQLLQGNNVK-----GYPKQY |
| 055 | NP_955564.1 | GNAYADSLTRILT------------------------------------- |
| 056 | YP_009508411.1 | DNTEKDQYSLPGINTILKKIGHSKVYSKF--------------------- |
| 057 | YP_009140788.1 | DITHKDQYSLPGINTILKRIGNSKIFSKF--------------------- |
| 058 | YP_009508410.1 | KEPE---LPDEILDQITSRFNKLSIAEAP--------------------- |
| 059 | NP_040333.1 | GNRMADQAAREVATRETPETSTLLIENSAPYTHEHFHYTVTDI-----KD |
| 060 | YP_008992013.1 | MFLQDDKFPLPRRQSMFTFLKNAQIFSKF--------------------- |
| 061 | NP_041734.1 | DNTEKDQYSLPGIQTILKRVGNKKVFSKF--------------------- |
| 062 | NP_056895.1 | LNNYVDQLLP---------------------------------------- |
| 063 | YP_009109689.1 | GNRQADRVARQAAMAEVLTLATEPDNTS--HITIEHTYTSEDQ-----EE |
| 064 | YP_002519387.1 | AATISDGYLLPNKETILTAIRGRKYFSTL--------------------- |
| 065 | YP_595725.1 | DNTWPDQYSLPGINALLKNVARAKIFSKF--------------------- |
| 066 | YP_009508551.1 | GNALADKLATQGSYVVNCNTKKPNLDAELDQLLQGHYIK-----GYPKQY |
| 067 | YP_001497148.1 | GNRQADQVARQVAVAETLTLTK-LEETN--LTTNKYAYTPEDQ-----EE |
| 068 | NP_041186.1 | GNHTADVLTKQVF------------------------------------- |
| 069 | NP_663784.1 | LVSQGIRQVLF--------------------------------------- |
| 070 | YP_009508443.1 | DNTEKDQYSLPGINTILKRIGQSVIYSKF--------------------- |
| 071 | YP_003987465.1 | DNTEKDQYSLPGISTILQKIGHSKIYSKF--------------------- |
| 072 | NP_861410.1 | KATIGNAHKLPRIDSILTKVKGSNWFSTL--------------------- |
| 073 | YP_009345071.1 | DNTEKDQYPLPGINTILQRIGKTKIYSKF--------------------- |
| 074 | YP_009508582.1 | GNNLADKLATQGSYNVNSIVKNPSLDAELEQLINGHSMK-----GYPSKY |
| 075 | NP_569141.1 | AKTKTYNYPIPNKILKIRQIQGYNYFSKF--------------------- |
| 076 | YP_002916057.1 | DNTEKDQYSLPGINTIISRVAGKKVFSKF--------------------- |
| 077 | NP_777384.2 | LNNYVDQLLP---------------------------------------- |
| 078 | YP_233110.1 | DNTEKDQYSLPGINTIISRIGKSKIYSKF--------------------- |
| 079 | NP_542258.1 | LNEATDALMLAPL------------------------------------- |
| 080 | YP_009130664.1 | DNTHKDQYSLPGINTIIQKVGNSKIYSKF--------------------- |
| 081 | YP_009508566.1 | GNALADKLAVQGSYTINNITIKPSLDTELRAVLEGKLPK-----GYPKNL |
| 082 | YP_006607892.1 | EATIGDSHNLPNKDELLTLIRGKTIYSSF--------------------- |
| 083 | YP_009506270.1 | DNTEKDQYSLPGINTIIRRIGNAKIYSKF--------------------- |
| 084 | NP_042513.1 | KATKGDAYNLDR---LYLTDRESNWFSTL--------------------- |
| 085 | YP_006907834.1 | DATVGDAYNLPNKDELLTLIRGKKIFSSF--------------------- |
| 086 | YP_004442827.1 | DNTEKDQYSLPGINALLKSVGNAKIYSKF--------------------- |
| 087 | YP_009408594.1 | DNTYKDQYSLPGINTILKKVGNSKIYSKF--------------------- |

```
088 YP_009508561.1    GNSLADKLAVQGSYSVNTINKIPSLDAELNQILEGNLPK-----GYPKQY
089 NP_056907.1       LNEYTDSLIIAPL-------------------------------------
090 YP_002308474.1    LNEATDALMLAPL-------------------------------------
091 YP_007761644.1    LRIRKNAYRIPNKDSLFLAIRESQFYSKF---------------------
092 NP_040550.1       GNDVADSQATFQA-------------------------------------
093 NP_758808.1       DNTYKDQYSLPNIQLILKKVINSTIYSKF---------------------
094 YP_009345075.1    QNTHKDQYSLPGINTIIKLVGNSKIYSKF---------------------
095 NP_068729.1       EATIGDSYKLPRKDFILEKIKGSLWFSSL---------------------
096 NP_047255.1       GNRLADDTAKKAATETHSSLTVLPTELIEGPKRPPWEYDDSDL-----DL
097 YP_006273075.1    DITHKDQYSLPGINTILKKVGNSKIFSKF---------------------
098 YP_009508888.1    GNALADKLATQGSYVVN-NIIKPSLDAELDQVLQGNLPK-----GYPKHF
099 NP_395469.1       EATIGTPKTLPRADYIMNRLKGKIWFSTL---------------------
100 YP_009508546.1    GNALADKLATQGSYVVNCNTKKPNLDAELDQLLQGQNVK-----GYPKQY
101 NP_056762.1       DNMHTDPFNIPHKISMINLIQKANIFSKF---------------------
102 YP_009508556.1    GNNLADKLATQGSYVVHCNTT-PSLDAELDQLLQGHNPP-----GYPKQY
103 NP_569153.1       DLTNKDQYSLPGIQTILQRLKGSTIFSKF---------------------
104 NP_043924.1       ETGKVQGYVIKTTLGMKTWIMNDHLVTQS---------------------
105 YP_009243641.1    GNEQVDKLAAFAS-------------------------------------
106 YP_004300274.1    AKTKTYNYPLPNKILRVRQVQGYNWFSKF---------------------
107 NP_056848.1       KKAKVVKYPIPNKDTLIHRSIQARYYSKF---------------------
108 YP_006495799.1    NNTHKDQYTLPGINSIVAAVGNAKIYSKF---------------------
109 YP_009508409.1    --------------------------------------------------
110 NP_687035.1       LVSKGIRQILF---------------------------------------
111 NP_045937.2       QRVLKKGDALASTDLVMEYSETDEKFTAG---------------------
112 NP_612577.1       KATKGDAHNLPNKDELLTLVRGKKIYSSF---------------------
113 NP_619548.1       QATIGDSHNLPNMQELLTLLRGKSIFSSF---------------------
114 NP_056880.1       GNAYADSLTRILT-------------------------------------
115 YP_009121747.1    DNTYKDXYSLPGINTLLKRIGNAKIFSKF---------------------
116 YP_009113237.2    DLTHKDQYSLPGIQGIIARVGRAKIFSKF---------------------
117 YP_009508408.1    DNTEKDQYSLPGINTILKRVGQSEIYSKF---------------------
118 NP_955579.1       GNRLADDTAKKAATETHSSLTVL---------------------------
119 YP_009229919.1    DNTFKDQYSLPGINTILQRIGKSKIYSKF---------------------
120 YP_004442833.1    DNTEKDQYSLPGINSLLNSVGNAKIYSKF---------------------
121 NP_862833.2       GNHTADVLTKQVF-------------------------------------
122 NP_040563.1       MVDSELQLQLLNIGTEHIRIQKDEVFMT----------------------
123 YP_443922.1       GNDAADQAAKKAGGYSPGQMVLRVDPPSTELTVEHIKELQQTAGPYEHSV
124 NP_057933.2       GNRMADQAARKAAITETPDTSTLLIENSSPYTSEHFHYTVTDI-----KD
125 YP_009116631.1    DNTHKDQYTLPGINTIISAIGNAKIFSKF---------------------
126 YP_009513249.1    GNNLADQLATQASFKVHMTK-NPKLDIEQIKAIQACQNNERLPVGYPKQY
127 YP_223871.1       GNRRADEVAREVAIRPLSTQATISDAPDMPDTETP-QYSNVEE-----AL
128 NP_127504.1       HFLQDDKFPIPNKLTLFSHLSKAKLFSKF---------------------
129 NP_044929.1       GNALADKQAVSGSYKVFSNELKPSLDAELEQVLSTGRPN---PQGYPNKY
130 NP_057860.1       LNALTDALLITPV-------------------------------------
131 NP_040973.1       GYRGEIGVIMINVSRKSITLMERQKIAQLIIILPCKHEVLEQGK------
132 NP_777317.1       DNTEKDQYSLPGINTIIKRIGTSKVYSKF---------------------
133 NP_054716.1       GNNAADKLAVKASYTVFSVQTLPSLDAELHQLLDKQTPN---PKGYPSKY
134 YP_001856242.1    LVSAGIRKVL----------------------------------------
135 NP_954565.2       GNQRADLATKIVAS------------------------------------
136 NP_569150.1       DNTEKDQYSLPGINTIIKRIGNAKIYSKF---------------------
137 NP_659397.1       EVTIGDSHNLPNMQELITLLRGKNIFSSF---------------------
138 YP_009513211.1    GNRKADEAAKQAAQSTRILTETTKNQEHFEPTRGKIKPRE----------
139 YP_233107.1       DNTEKDQYSLPGINTIIARISHSKIYSKF---------------------
140 YP_009109692.1    GNRQADRVARQAAMAEVLTLATEPDNTS--HITIEHTYTSEDQ-----EE
141 NP_057849.4       LVSAGIRKVLF---------------------------------------
142 NP_049560.1       LNALTDALLITPV-------------------------------------
143 YP_009506267.1    DNTEKDQYSLPGINTILKKVGNSKIYSKF---------------------
144 NP_039820.1       ENTESDQYSLPGINTIISKVGRSKIYSKF---------------------
145 NP_056790.1       GNRRADEAAKQAALSTRVLAGTTKPQEPIEPAQEKTRPRE----------
146 YP_002321513.1    DNTYRDPYSLPGINTIIQKVGRSKIYSKF---------------------
147 NP_955591.1       GNRMADQAARKAAITETPDTSTLLI-------------------------
148 YP_001956722.2    GNNLADKLATQGSYVVNINTT-PSLDAELDQLLQGQYPK-----GFPKHY
149 NP_056902.1       GYTGEIQVICTNIGKSNIKLIEGQKFAQLIILQHHSNSRQPWD-------
150 NP_056886.1       GNDVADSQATFQA-------------------------------------
151 NP_041261.1       GNALADAATQIFP-------------------------------------


001 YP_004442836.1    ----------------DLKSGFHQVAMDPESIPWTAFWAI-DGLYEWLVM
002 NP_056728.1       ----------------DCKSGFWQVLLDQESRPLTAFTCP-QGHYEWNVV
003 YP_009508537.1    TYELRNDKCMVLR--KDGWREIPPSRERYKLIKEAHD---ISHAGREAVL
004 YP_004732983.2    ----------------DLKSGFHQVMMEEESIPWTAFISP-AGLYEWLVM
005 YP_009508407.1    ----------------PEKQGK----------------------------
006 NP_040939.1       -------ESRKTERGEKGFGSTGMYW------IENIPLAEEDHTKWHQDA
```

007 YP_003864102.1 ------------------------------NLNTNLQNAQDTHALHHLNA
008 YP_009002585.1 ----------------DLKSGFHQVAMHPESIKWTAFWVP-DGLYEWLVM
009 YP_003284237.1 ----------------DLKSGFHQVAMEEASIPWTAFWAI-DGLYEWLVM
010 YP_009268869.1 -------ETRRTERGNQGFGSTGAYW------IENIPLAEEDHSKWHQDA
011 YP_009506251.1 ----------------DLKSGFHQVAMAEESIPWTAFWVP-QGLYEWLAM
012 NP_955611.1 ---------------------------------Y----------------
013 YP_008567619.1 ----------------DLKSGFHQVAMDPESIEWTAFSTP-NGLYEWLVM
014 NP_149413.1 ----------------DLKSGFHQVAMDPESIPLTAFTAY-NELYEWLVM
015 YP_004442824.1 ----------------DLKSGFHQVAMDPESIPWTAFWAI-DGLYEWLVM
016 YP_001931967.1 ----------------DCKSGFWQVLLDKPSQELTAFTCP-QGHYQWLVM
017 YP_002455786.1 ----------------------------------TPLKPEGLHALTHCNQ
018 YP_009109694.1 --------------------------------------------------
019 NP_789739.1 --------------------------------------------------
020 NP_955619.1 ----------------------------------LQLSPADLHSFTHCGQ
021 YP_224289.1 ----------------DLKSGFHQVAMDPESIPWTAFLAN-NELYEWLVM
022 YP_009182100.1 ----------------DCKSGYWQIKLTPESIPLTAFSTP-KGQYEWKVL
023 NP_705927.1 --------------------------------------------------
024 YP_009508577.1 KYILKEGQVFVLR--PEGEKIIPPKSDRLALVKIAHE---FSHAGREATV
025 YP_004442830.1 ----------------DLKSGFHQVAMEEESIEWTAFWAF-TGLYEWLVM
026 NP_758887.1 --------------------------------LEGIDKAQEEHDKYHNNW
027 YP_009508406.1 --------------------------------------------------
028 YP_610965.1 ----------------DLKSGFHQVAMSPESIEWTAFIVL-GGLYEWLVM
029 YP_001931961.1 ----------------DCKSGFWQVLLDEDSQLLTAFTCP-QGHYQWIVV
030 YP_241114.1 -------------------------------FQSAIDAARKSHDLHHQNS
031 YP_006732334.1 ----------------DCKSGFWQVFLDQESQKLTAFTCP-QGHFQWRVV
032 YP_009041481.1 ----------------DLKSGFHQVAMDEASIPWTAFLVP-GGLYEWLVM
033 YP_001036293.1 ----------------DLKSGFHQVAMAPQSVEWTAFLAP-GGLYEWLVM
034 YP_605811.1 ----------------DLKSGFHQVAMDPESIPWTAFWAI-DGLYEWLVM
035 NP_955577.1 VQKLEAHYEPKRGTWEYRGKTIMPEKYAKELISHLHKLTHLSARKMKTLL
036 YP_003208050.1 ----------------DLKSGFHQVAMDEESIPWTAFCVP-GGLYEWLVM
037 NP_789740.1 --------------------------------LDGIDKAQDEHEKYHSNW
038 YP_004442839.1 ----------------DLKSGFHQVAMDPESIPWTAFWAI-DGLYEWLVM
039 YP_009507791.1 -------MIMDSQRGEGGFGSTGAYVTQISSWMDNIEKAEDDHDKFHSDV
040 YP_004581513.1 ----------------DLKAGFHQIRMEEKSKPWTAFWTP-EGLYEFEVM
041 NP_043933.1 ----------------DCKSGFWQVRLAPETIQLTAFSCP-QGHYEWLVM
042 YP_009507248.1 -----------------------------QDIIRHYHDGKTNHRGINECY
043 YP_002117531.1 ----------------DLKSGFHQVAMDPASIEWTAFWAI-DGLYEWLVM
044 YP_004347415.1 ----------------DCKSGFYHIKLEEDSKKYTAFTVP-QGYYVWIVL
045 NP_056891.1 ------------------------------NINTNLESAQNAHTLHHLNA
046 YP_009513242.1 KYTLEDNKIIVER--PNGQRIVPPKSDREKIISMAHN---IAHTGRDATF
047 YP_009352866.1 ----------------DLKSGFHQVAMDEDSIPWTAFLVP-GGLYEWLVM
048 YP_009165750.1 ----------------DCKSGFWQVLLDQESQLLTAFTCP-TGHYQWKVV
049 NP_040840.1 -------ETRKTERGEQGFGSTGMYW------IENIPLAEEEHNKWHQDA
050 YP_004222728.1 ---------------------------------YPLREAKDLHTALHIGP
051 YP_009506264.1 ----------------DLKSGFHQVAMEAESIPWTAFWAI-DGLYEWLVM
052 YP_567050.1 ----------------------------------IPLTPQDLHKLTHCNS
053 YP_009508571.1 TYYMEDGKVKVNR--PEGTKIIPPSLERAGIVQKAHN---LAHTGREATL
054 NP_056803.1 TYYLEDGKVKVSR--PEGVKIIPPQSDRQKIVLQAHN---LAHTGREATL
055 NP_955564.1 ----------------------------------ALESAQESHALHHQNA
056 YP_009508411.1 ----------------DLKSGFHQVAMHPESIEWTAFLTT-AGLFEWLVM
057 YP_009140788.1 ----------------DLKSGFHQVAMHPDSIEWTAFWVP-DGLYEWLVM
058 YP_009508410.1 ----------------SEPRGR----------------------------
059 NP_040333.1 LTKLGATYDDAKKCWVYQGKPVMPDQFTFELLDFLHQLTHLSFSKTKALL
060 YP_008992013.1 ----------------DLKSGFWQLGIEPSERYKTAFCIP-NAHFQWTVL
061 NP_041734.1 ----------------DLKSGFHQVAMAEESIPWTAFWVP-QGLYEWLVM
062 NP_056895.1 -----------------------------------LETPEQWHKLTHCNS
063 YP_009109689.1 ARAIGATENKDTRNWEKEGKIVLPQKEALAMIQQMHAWTHLGNRKLKLLI
064 YP_002519387.1 ----------------DCKSGFWQIRLNENSKPLTAFSCP-MGQYEWNVV
065 YP_595725.1 ----------------DLKSGFHQVAMDEESIPLTAFSAY-NELYEWLVM
066 YP_009508551.1 TYFLEDGKVKVSR--PEGVKIIPPQSDRQKIVLQAHN---LAHTGREATL
067 YP_001497148.1 AKAIGAILNQDTKDWEKEGKIVLPRKEALAMIQQMHAWTHLSNRKLKSLI
068 NP_041186.1 -------------------------------FQSAIDAARKSHDLHHQNS
069 NP_663784.1 --------------------------------LEKIEPAQEEHEKYHSII
070 YP_009508443.1 ----------------DLKSGFHQVAMDPDSVQWTAFWVP-DGLYEWLVM
071 YP_003987465.1 ----------------DPKSGFHQVAMHPDSVPWTAFWAI-NGLYEWLVM
072 NP_861410.1 ----------------DAKSGYWQLRLHPQSKPLTAFSCPPQKHYQWNVL
073 YP_009345071.1 ----------------DLKSGFHQVAMAEESIPWTAFAIPGKGLFEWMVM
074 YP_009508582.1 KYILKEGQVFVLR--PEGEKIIPPKSDRPALVKVAHE---FSHAGREATV
075 NP_569141.1 ----------------DCKSGFYHLKLEDESKKLTAFTVP-QGFYEWNVL
076 YP_002916057.1 ----------------DLKSGFHQIRMSKESIPWTAFWTP-DGLYEFLVM
077 NP_777384.2 -----------------------------------LETPEQWHKLTHCNS
078 YP_233110.1 ----------------DLKSGFHQVAMDPESIPWTAFWAI-DGLYEWLVM
079 NP_542258.1 ----------------------------------LPLNPTTLHQITHCNP

080 YP_009130664.1 ----------------DLKSGFHQVAMHPYSIEWTAFWVP-QGLYEWLAM
081 YP_009508566.1 KYEYNSPNLIVIR--KEGQRIIPPLSDRPKLVKQAHE---LAHTGREATL
082 YP_006607892.1 ----------------DCKSGFWQVFLDQESQLLTAFTCP-SGHYQWRVV
083 YP_009506270.1 ----------------DLKSGFHQVAMDPESIPWTAFWAI-DGLYEWLVM
084 NP_042513.1 ----------------DAKSGFLQLRLDEETKPLTAFSCPPQMHLEYNVM
085 YP_006907834.1 ----------------DCKSGFWQVRLDEESKSLTAFTCP-QGHYEWNVV
086 YP_004442827.1 ----------------DLKSGFHQVAMEEESIEWTAFWAI-TGLYEWLVM
087 YP_009408594.1 ----------------DLKSGFHQIAMEKESIPWTAFIVP-QGLYEWLVM
088 YP_009508561.1 KYVLKNNELIVQR--PEGDKIIPPKADRLPLVKTAHE---LAHTGREATL
089 NP_056907.1 ----------------------------------IPLTPQDLHKLTHCNS
090 YP_002308474.1 ----------------------------------LPLSPATLHEITHCNP
091 YP_007761644.1 ----------------DCKSGFYQVPMEQDSIQLTAFSTP-IGSYEWLVM
092 NP_040550.1 ---------------------------------YPLREAKDLHTALHIGP
093 NP_758808.1 ----------------DLKSGFHQVAMDPDSVEWTAFLVP-QGLYEWLAM
094 YP_009345075.1 ----------------DLKSGFHQVAMEESSIPWTAFWVP-DGLYEWLVM
095 NP_068729.1 ----------------DAKSGYYQLRLHENTKPLTAFSCPPQKHYEWNVL
096 NP_047255.1 VQKLEAHYEPKRGTWEYRGKTIMPEKYAKELISHLHKLTHLSARKMKTLL
097 YP_006273075.1 ----------------DLKSGFHQVAMHPDSIEWTAFWVP-DGLYEWLVM
098 YP_009508888.1 VYTLEEGKVIVKR--PEGTKIIPPLADRKLLASQAHK---LSHSGREATL
099 NP_395469.1 ----------------DVKSAYWQLRLTEESKPLTAFSYPPPKHYEWNVL
100 YP_009508546.1 TYFLEDGKVKVSR--PEGVKVIPPYSDRQKIVLQAHN---LAHTGREATL
101 NP_056762.1 ----------------DLKAGFHHMKLKDDFKDWTTFTCS-EGLYTWNVC
102 YP_009508556.1 KYTLEDNKIIVER--PAGKRIVPPKADREKIISTAHN---IAHTGRDATF
103 NP_569153.1 ----------------DLKSGFHQVAMHPDSIEWTAFWVP-SGLYEWLVM
104 NP_043924.1 ----------------EQTEGRAKLSSTEGYALAQQYHHLYGHPSEESLR
105 YP_009243641.1 --------------------------------------PEEEHRYFHNNA
106 YP_004300274.1 ----------------DCKSGFYHLKLTEESKHLSAFNVP-QGFYEFNVL
107 NP_056848.1 ----------------DCKSGFYHIKLEEDSKKYTAFTVP-QGYYQWKVL
108 YP_006495799.1 ----------------DLKAGFHQVLMEESSIPWTAFITP-VGFYEWLVM
109 YP_009508409.1 --------------------------------------------------
110 NP_687035.1 --------------------------------LDRIEEAQDDHAKYHNNW
111 NP_045937.2 --------------AELHDGVFMRGDLIVPPLEMLHAILLAIHGVSHTHK
112 NP_612577.1 ----------------DCKSGLWQVLLDKESQLLTAFTCP-QGHYQWNVV
113 NP_619548.1 ----------------DCKSGFWQVVLDEESQKLTAFTCP-QGHFQWKVV
114 NP_056880.1 ----------------------------------ALESAQESHALHHQNA
115 YP_009121747.1 ----------------DLKSGFHQVAMEEESIPWTAFLIP-GGLYEWLVM
116 YP_009113237.2 ----------------DLKSGFHQVAMHPESIPWTAFWVP-QGLYEWLVM
117 YP_009508408.1 ----------------DLKSGFHQVAMDEGSIEWTAFWVP-DGLFEWLVM
118 NP_955579.1 --------------------------------------------------
119 YP_009229919.1 ----------------DLKSGFHQVAMDEESIPWTAFITP-EGLYEWLVM
120 YP_004442833.1 ----------------DLKSGFHQVAMEEESIEWTAFWAF-TGLYEWLVM
121 NP_862833.2 -------------------------------FQSAIDAARKSHNLHHQNS
122 NP_040563.1 ------------------------------CFLENIPSATEDHERWHTSP
123 YP_443922.1 WKQKGASQGPEGLWRCHDGRLVAPANLCPELIREAHGLTHEGKLKTLQRV
124 NP_057933.2 LTKLGAIYDKTKKYWVYQGKPVMPDQFTFELLDFLHQLTHLSFSKMKALL
125 YP_009116631.1 ----------------DLKSGFHQVLMDEESIPWTAFVTP-VGFYEWKVM
126 YP_009513249.1 TYELQNNKCMVLR--KDGWREIPPSRERYKLIKEAHN---ISHAGREAVL
127 YP_223871.1 GHRLRGTKDPAGWWHLPDGRLLLPRAVGRKVLEQTHRATHLGESKLTELV
128 NP_127504.1 ----------------DLKSGFWQLGIHPNERPKTGFCIP-DRHFQWKVM
129 NP_044929.1 EYKLVNGLCYVDRRGEEGLKIIPPKADRVKLCQLAHDGPGSAHLGRSALL
130 NP_057860.1 ----------------------------------LQLSPADLHSFTHCGQ
131 NP_040973.1 -------VVMDSERGDNGYGSTGVFS----SWVDRIEEAEINHEKFHSDP
132 NP_777317.1 ----------------DLKSGFHQVAMAEESIPWTAFWAI-DGLYEWLVM
133 NP_054716.1 EYTLRDGQVYVKR--TDGEKIIPSKDDRVKILELAHKGPGSGHLGKNTMY
134 YP_001856242.1 --------------------------------------------------
135 NP_954565.2 ------------------------------NINTNLESAQNAHTLHHLNA
136 NP_569150.1 ----------------DLKSGFHQVAMDPESIPWTAFWAI-DGLYEWLVM
137 NP_659397.1 ----------------DCKSGFWQVLLDDESQKLTAFTCP-QGHYQWRVV
138 YP_009513211.1 ----------------------LTPDQGREFIQRLHQLTHLGPDKLLQLV
139 YP_233107.1 ----------------DLKSGFHQVAMEEESIPWTAFWAI-NGLYEWLVM
140 YP_009109692.1 ARAIGATENKDTRNWEKEGKIVLPQKEALAMIQQMHAWTHLGNRKLKLLI
141 NP_057849.4 --------------------------------LDGIDKAQDEHEKYHSNW
142 NP_049560.1 ----------------------------------LQLSPAELHSLTHCGQ
143 YP_009506267.1 ----------------DLKSGFHQVAMHPDSIQWTAFCVP-GGLYEWLVM
144 NP_039820.1 ----------------DLKSGFWQVAMEEESVPWTAFLAG-NKLYEWLVM
145 NP_056790.1 ----------------------LTPDRGKEFIKRLHQLTHLGPEKLLQLV
146 YP_002321513.1 ----------------DLKSGFHQVAMDPESIPWTAFLTP-QGLFEWLVM
147 NP_955591.1 --------------------------------------------------
148 YP_001956722.2 QYQLENGQVMVTR--PNGKRIIPPKSDRPQIILQAHN---IAHTGRDSTF
149 NP_056902.1 -------ENKISQRGDKGFGSTGVFW------VENIQEAQDEHENWHTSP
150 NP_056886.1 ---------------------------------YPLREAKDLHTALHIGP
151 NP_041261.1 ------------------------------IISDPIHEATQAHTLHHLNA

```
001 YP_004442836.1    PFGLKNAPAIFQRKMDNCFRG-TEKFIAVYIDDILVFSET-VQQHKEHLK
002 NP_056728.1       PFGLKQAPSIFQRHMDEAFR-VFRKFCCVYVDDILVFSNN-EEDHLLHVA
003 YP_009508537.1    LKIQENYWWPKMKKDISSFLSTCNVCKMVNPLNLKPISPQA-IVHPTKPF
004 YP_004732983.2    PFGIQNAPAIFQRKMDECFKG-TEDFIAVYIDDILVFSNS-IREHEKHLQ
005 YP_009508407.1    -------------------------LLVFKDPVLIYQEE----------
006 NP_040939.1       RSLHLEFE--IPRTAAEDIVNQCEICKEARTPAVIR---GGNKRG----V
007 YP_003864102.1    QTLRLMFK--IPREQARQIIKQCPTC--VTYSPVPH---LGVNPRGLLPN
008 YP_009002585.1    PFGLKNAPAVFQRKMDNVFKG-TEAFIAVYIDDILVFSQS-EEEHIKHIR
009 YP_003284237.1    PFGLKNAPAVFQRKMDNCFRG-TEEFIAVYIDDILIFSES-PQQHVQHLK
010 YP_009268869.1    GSLHLDFG--IPRTAAEDIVQQCEVCQENKMPSTIR---GSNRRG----I
011 YP_009506251.1    PFGLKNAPAIFQRKMDVCFKG-TEDFIAVYIDDILVFSET-MEEHENHIS
012 NP_955611.1       --------------------------------------------------
013 YP_008567619.1    PFGLKNAPAVFQRKMDNCFKG-MEDFIAVYIDDILVFSEN-MRDHAQHLV
014 NP_149413.1       PFGLKNAPAVFQRKMDDCFRG-TERFIAVYIDDILVFSET-EEKHAEHLW
015 YP_004442824.1    PFGLKNAPAIFQRKMDHCFH---EDFIAVYIDDILVFSET-VQQHRQHLK
016 YP_001931967.1    PFGLKQAPAIFQRHMQIALN-EHSAYSCVYIDDILVFSEN-EKDHEIHVS
017 YP_002455786.1    QALVSHG---ATPAQAKQLVQACRTCQIINPQHHMP---RGHIRRGHFPN
018 YP_009109694.1    --------------------------------------------------
019 NP_789739.1       --------------------------------------------------
020 NP_955619.1       TALTLQG---ATTTEASNILRSCHACRKNNPQHQMP---QGHIRRGLLPN
021 YP_224289.1       PFGLKNAPAIFQRKMDTCFKG-TEAFIAVYIDDILVFSET-EQLHRDHLR
022 YP_009182100.1    PFGLKNAPNIFQRRMDKIFKDCY-QFCGVYVDDILVFSKD-KEEHLLHLD
023 NP_705927.1       --------------------------------------------------
024 YP_009508577.1    LRLQDKYWWPNMRKDVISHIRTCKPCLTTDGSNLTPIPPKP-QLRPEKPF
025 YP_004442830.1    PFGLKNAPAIFQRKMDNCFRG-TEKFITVYIDDILVFSET-EEEHAIHLT
026 NP_758887.1       RALAQDFC--IPNIVAKEIVAQCPKCQTK--GEPIH---GQVDTS----P
027 YP_009508406.1    ------MSAIWENSLKAYFE------------------------------
028 YP_610965.1       PFGLKNAPAVFQRKMDHCFAG-TEKFIAVYIDDILIFSAN-DEEHVEHLK
029 YP_001931961.1    PFGLKQAPSIFQRHMNNAFR-DFASYCCVYVDDILVFSNN-IKDHYAHVA
030 YP_241114.1       HSLRLQFK--ISREAARQIVKSCSTC--PQFFVLPQ---YGVNPRGLRPN
031 YP_006732334.1    PFGLKQAPSIFQRHMQNALR-GLEDYCLVYVDDIIVFSNS-EKEHYDHVL
032 YP_009041481.1    PFGLRNAPAIFQRKMDEVFAD-LKEFVSVYIDDILVFSET-YEEHAAHLK
033 YP_001036293.1    PFGLKNAPAVFQRKMDNVFRG-TEEFIAVYIDDILVFSET-EEEHLKHLR
034 YP_605811.1       PFGLKNAPAIFQRKMDNCFRG-TEEYIAVYIDDILIFSDN-VSDHRKHLS
035 NP_955577.1       EREETGFYLPNRDLHLRQVTESCRACAQINAGKIKFGPDV--RARGRRPG
036 YP_003208050.1    PFGLKNAPSVFQRKMDDCFKG-TEAFIAVYIDDILVFSKN-EEEHQKHLQ
037 NP_789740.1       RAMASDFN--LPPVVAKEIVASCDKCQLK--GEAMH---GQVDCS----P
038 YP_004442839.1    PFGLKNAPAIFQRKMDNCFRG-TEEFIAVYIDDILIFSDN-ISDHRKHLK
039 YP_009507791.1    VYLKQRFG--IPRQVAEEIIRKCPLCMIK--GEVSY---GQLKIG----E
040 YP_004581513.1    PFGLMNAPADFQRKMDNAFRG-TDAFIAVYIDDILVFSEN-EEEHEDHLL
041 NP_043933.1       PFGLKQAPAIFQRHMDESLSNMYPQFCAVYVDDIIVFSKT-EEEHLGHVK
042 YP_009507248.1    LALSKRYYWPRMKDQITKFINECTICGQAKYDRNPIR-----PQFNIVPP
043 YP_002117531.1    PFGLKNAPATFQRKMDMCFAG-TEHFIAVYIDDILVFSET-EEDHVKHLH
044 YP_004347415.1    PFGYHNSPSIYQQFMDGIFR-PYYDFILVYIDDILIFSKT-YEEHKIHLE
045 NP_056891.1       QTLRLMFN--IPREQARQIVKQCPIC--VTYLPVPH---LGVNPRGLFPN
046 YP_009513242.1    LKVSSKYWWPNLRKDVVKVIRQCKQCLVTNAANLTSPPILR-PEKPLKPF
047 YP_009352866.1    PFGLRNAPAIFQRKMDEVFGD-LKDFIAVYIDDILVFSET-HEQHAQHIK
048 YP_009165750.1    PFGLKQAPSIFQRHMNNAFR-EFEEFCCVYVDDILVFSKN-ETEHRKHVI
049 NP_040840.1       VSLHLEFG--IPRTAAEDIVQQCDVCQENKMPSTLR---GSNKRG----I
050 YP_004222728.1    RALSKACN--ISMQQAREVVQTCPHC---NSAPALE---AGVNPRGLGPL
051 YP_009506264.1    PFGLKNAPAVFQRKMDGCFKG-TEEFIAVYIDDILVFSET-PQQHVQHLR
052 YP_567050.1       RALVSSG---ATPQQAKSLLQTCYTCNIINSQHHMP---QGHIRRGLMPN
053 YP_009508571.1    LKIANLYWWPNMRKDVVRQLGRCQQCLVTNAFNQTSGPILR-PTRPLKPF
054 NP_056803.1       LKIANLYWWPNMRKDVVKQLGRCKQCLITNASNKTSGPILR-PDRPQKPF
055 NP_955564.1       AALRFQFH--ITREQAREIVKLCPNC--PDWGHAPQ---LGVNPRGLKPR
056 YP_009508411.1    PFGLKNAPAVFQRKMDNCFRG-MEDFLAVYIDDILVFSNS-EEEHMRHLE
057 YP_009140788.1    PFGLKNAPSIFQRKMDECFNG-TEEFIAVYIDDILVFSED-EKSHAKHLR
058 YP_009508410.1    -------------------------ILVHTDPVKIFEEE----------
059 NP_040333.1       ERNYCPYYMLNRDRTLKDITETCQACAQVNASKSAVKQGT--RVRGHRPG
060 YP_008992013.1    PFGLKTAPSIFQKSMIKIFQP-LLSNALIYIDDILLFSGS-HDDHHRLLN
061 NP_041734.1       PFGLKNAPAVFQRKMDQCFKG-TEEFIAVYIDDILVFSEN-MAEHTKHIG
062 NP_056895.1       RALSRWPNPRISAWDPRSPATLCETCQRLNPTGGGK---MRTIQRGWAPN
063 YP_009109689.1    EK--TDFLIPRASTLIEQVTSACKVCQQVNAGATRVPAGK--RTRGNRPG
064 YP_002519387.1    PFGLKQAPGLFQRFMDNSFK-EYSAFCAVYVDDILVFSKT-LDEHYDHLE
065 YP_595725.1       PFGLKNAPAIFQRKMDQCFRG-TEGFIAVYIDDILVFSED-EEQHAEHLW
066 YP_009508551.1    LKIANLYWWPNMRKDVVKQLGRCQQCLITNASNKASGPILR-PDRPQKPF
067 YP_001497148.1    EK--TDFLIPKAGTLIEQVTSACKVCQQVNAGATRVPEGK--RTRGNRPG
068 NP_041186.1       HSLRLQFK--ISREAARQIVKSCSTC--PQFFVLPQ---YGVNPRGLRPN
069 NP_663784.1       KELTHKFG--IPLLVARQIVNSCAQCQQK--GEAIH---GQVNAE----I
070 YP_009508443.1    PFGLKNAPAVFQRKMDNCFRG-TEDFIAVYIDDILVFSRS-QEEHAQHLK
```

071 YP_003987465.1 PFGLKNAPAVFQRKMDHCFRG-TEDFIAVYIDDILVFSET-PEQHKKHLE
072 NP_861410.1 PFGLKQAPGIYQNFMDKNLE-GLENFCLAYIDDILVFTNSSREEHLSKLL
073 YP_009345071.1 PFGLKNAPSVFQRKMDLCFTG-LEKFVAVYIDDVLIFSNS-EQEHLEHLD
074 YP_009508582.1 LRLQDKYWWPNMRKDVISHIRMCKPCLTTDSSNLTPIPPKP-QQRPKKPF
075 NP_569141.1 PFGYKNAPGRYQHFMDNYFN--QLENCIVYIDDILLYSRT-QDEHIKLLE
076 YP_002916057.1 PFGLVNAPADFQRKMDNAFRG-TEAFIAVYIDDILIFSET-EEDHEQHLL
077 NP_777384.2 RALSRWPNPRISAWDPRSPATLCETCQRLNPTGGGK---MRTIQRGWAPN
078 YP_233110.1 PFGLKNAPAIFQRKMDNCFRG-TEEFIAVYIDDILIFSDN-ISDHRKHLS
079 NP_542258.1 HALRNHG---ATASEAHAIVQACHTCKVINPQGRLP---QGYIRRGHAPN
080 YP_009130664.1 PFGLKNAPAVFQRKMDNCFKG-TENFIAVYIDDILVFSET-EREYARHLE
081 YP_009508566.1 LRLQNQYWWPKMRKDVSHCLRTCMPCLQTNSTNLTTTRPFQ-QIRPSKPF
082 YP_006607892.1 PFGLKQAPSIFQRHMQNAFR-SFEKFCCVYVDDILVFSDN-EQDHHRHVM
083 YP_009506270.1 PFGLKNAPAVFQRKMDSCFRG-TEEFIAIYIDDILVFSET-VQQHKEHLK
084 NP_042513.1 PMGLKQAPSQFQRFMDNNLR-GLEDISLAYIDDIIVFTKGTKDYHLKQVA
085 YP_006907834.1 PFGMKQAPSIFQRHMDEAFK-VFRKFCCIYVDDILVFSDN-EQNHQLHVA
086 YP_004442827.1 PFGLKNAPAIFQRKMDLCFQG-TESFIAVYIDDILIFSQN-EEEHAEHLY
087 YP_009408594.1 PFGLKNAPALFQRKMDNCFKG-TEDFIAVYIDDVLVFSNN-EQDHHRHLH
088 YP_009508561.1 LKLQTTHWWPNMRKDIITVLRQCKPCLQTDSTNLTPIPPVS-QPRPVKPF
089 NP_056907.1 RALVSSG---ATPQQAKSLLQTCYTCNIINSQHHMP---QGHIRRGLMPN
090 YP_002308474.1 PALCNHG---ATATETKAIVRACHTCKITNPQGRLP---QGHIRRGHAPN
091 YP_007761644.1 PFGLATAPSIFQAKMDNVLE-DHHDYCLVYIDDIIVFSRT-LEEHKIHVI
092 NP_040550.1 RALSKACN--ISMQQAREVVQTCPHC---NSAPALE---AGVNPRGLGPL
093 NP_758808.1 PFGLKNAPAVFQRKMDAVFKG-CEKFLAVYIDDILVFSNN-EEDHAKHLV
094 YP_009345075.1 PFGLKNAPAIFQRKMDNCFKG-TEEFIAVYIDDILVFSQT-EEEHVRHLQ
095 NP_068729.1 SFGLKQAPSIYQRFMDQSLK-GLEHICLAYIDDILIFTKGSKEQHVNDVR
096 NP_047255.1 EREETGFYLPNRDLHLRQVTESCRACAQINAGKIKFGPDV--RARGRRPG
097 YP_006273075.1 PFGLKNAPSVFQRKMDECFKG-TEDFIAVYIDDILVFSEN-EKDHAKHLK
098 YP_009508888.1 LKLSNTYWWPNMRKDVVKVIGQCQQCLVTNPSNLTSGPILR-PERPTKPF
099 NP_395469.1 PMGLKQAPGIFQEFMNRSLH-NLEHICLVYVDDIIIFSEKDKNDHLSKVL
100 YP_009508546.1 LKIANLYWWPNMRKDVVKQLGRCQQCLITNASNKASGPILR-PDRPQKPF
101 NP_056762.1 PFGIANAPCAFQRFMQESFG--DLKFALLYIDDILIASNN-EKEHIEHLK
102 YP_009508556.1 LKVSSKYWWPNLRKDVVKVIRQCKQCLVTNAANLTSPPILR-PEKPLKPF
103 NP_569153.1 PFGLKNAPAIFQRKMDHCFKG-TEAFIAVYIDDILVFSKT-EQDHEKHLQ
104 NP_043924.1 KVLTKRFVWEDMGQHCKEITNTCLTCAKYKVLRAGP--PMGVGRSAEGPC
105 YP_009243641.1 GSLHHRWR--ISWKQAKDILSNCQICK-PLHLRRTS---IGVNPRGLAPD
106 YP_004300274.1 MFGYKNAPGRYQCYMDSYFS--KLENCIVYIDDILLYSKT-KDEHETLLK
107 NP_056848.1 PFGYHNSPSIFQQFMDRIFR-PYYDFIIVYIDDILVFSKT-IEEHKIHIA
108 YP_006495799.1 PFGIANAPAVFQRKMDNCFHK-LREFVAVYIDDILVFSNS-LQEHESHLR
109 YP_009508409.1 ------MSVKWEKSVQAYFE------------------------------
110 NP_687035.1 RSMVQEFG--LPNIVAKEIVAACPKCQIR--GEPKH---GQVDAS----I
111 NP_045937.2 GGIMSYFSKFWTHPKASQTIDLILGHCQICLKHNPKYKSRLQGHRPLPSR
112 NP_612577.1 PFGLKQAPSIFPKTYANSHSNQYSKYCCVYVDDILVFSNTGRKEHYIHVL
113 NP_619548.1 PFGLKQAPSIFQRHMQTALN-GADKFCMVYVDDIIVFSNS-ELDHYNHVY
114 NP_056880.1 AALRFQFH--ITREQAREIVKLCPNC--PDWGHAPQ---LGVNPRGLKPR
115 YP_009121747.1 PFGLKNAPAIFQRKMDKCFKD-TEEFIAVYIDDILVYSNS-EADHERHLK
116 YP_009113237.2 PFGLKNAPAIFQRKMDNCFMG-TEEFIVVYIDDILVFSQN-EQDHERHIR
117 YP_009508408.1 PFGLKNAPAVFQRKMDNCFRG-MEDFVAVYIDDILVFSKT-PEEHAVHLE
118 NP_955579.1 --------------------------------------------------
119 YP_009229919.1 PFGLKNAPAIFQRKMDNCFRG-TEEFIAVYIDDILVFSET-EEQHAKHLQ
120 YP_004442833.1 PFGLKNAPAIFQRKMDNCFRG-TEAFIAVYIDDILIFSKD-PEEHAKHLE
121 NP_862833.2 HSLRLQFK--ISREAARQIVKSCSTC--PQFFVLPQ---YGVNPRGLRPN
122 NP_040563.1 DILVRQFH--LPKRIAKEIVARCQECKRT-TTSPVR---GTNPRG----R
123 YP_443922.1 SYTWWHPYMKEMTDLFCDNCTICGNYNPKKPYQTPMGHYP--VPNACFQD
124 NP_057933.2 ERSHSPYYMLNRDRTLKNITETCKACAQVNASKSAVKQGT--RVRGHRPG
125 YP_009116631.1 PFGLANAPAVFQRKMDQCFAG-TSEFIAVYIDDILVFSKT-LKEHEKHLS
126 YP_009513249.1 LKIQENYWWPKMKKDISSFLSTCNVCKMVNPLNLKPISPQA-IVHPTKPF
127 YP_223871.1 RK---HYLICGIYRAARDITTRCVACAQVNPGAAPVEKGLNSRIRGAAPG
128 NP_127504.1 PFGLKTAPSLFQKAMIKIFQP-ILFSALVYIDDILLFSET-LEDHIKLLN
129 NP_044929.1 LKLQQKYWWPRMHIDASRIVLNCTVCAQTNSTNQKPRPPLV-IPHDTKPF
130 NP_057860.1 TALTLQG---ATTTEASNILRSCHACRKNNPQHQMP---QGHIRRGLLPN
131 NP_040973.1 QYLRTEFN--LPKMVAEEIRRKCPVCRIR--GEQVG---GQLKIG----P
132 NP_777317.1 PFGLKNAPAIFQRKMDECFRG-MEEFIAVYVDDILIFSNS-VQDHKRHLQ
133 NP_054716.1 IKILNKYWWPNLIKDISKYIRTCTNCIITNTDNVPNKSYIV-QEKTGLPF
134 YP_001856242.1 --------------------------------------------------
135 NP_954565.2 QTLRLMFN--IPREQARQIVKQCPIC--VTYLPVPH---LGVNPRGLFPN
136 NP_569150.1 PFGLKNAPAIFQRKMDNCFRG-TEDFIAVYIDDILVFSET-IHQHKEHLK
137 NP_659397.1 PFGLKQAPSIFQRHMQDALR-GLEEFSLVYVDDIIVFSDN-KNDHQDHVM
138 YP_009513211.1 GR--TSFHIPNLQSVVREITSKCQVCAVTNAVTTYREPGR--RQRGDRPG
139 YP_233107.1 PFGLKNAPAIFQRKMDNCFRG-TEKFIAVYIDDILIFSDS-KEAHRTHLR
140 YP_009109692.1 EK--TDFLIPRASTLIEQVTSACKVCQQVNAGATRVPAGK--RTRGNRPG
141 NP_057849.4 RAMASDFN--LPPVVAKEIVASCDKCQLK--GEAMH---GQVDCS----P
142 NP_049560.1 AALTLQG---ATTTEASNILRSCHACRKNNPQHQMP---RGHIRRGLLPN
143 YP_009506267.1 PFGLKNAPSVFQRKMDNCFRG-TENFIAVYIDDILVFSRT-PEEHAEHLL

```
144 NP_039820.1        PFGLKNAPAIFQRKMDNVFKG-TEKFIAVYIDDILVFSET-AEQHSQHLY
145 NP_056790.1        NR--TSLLIPNLQSAVREVTSQCQACAMTNAVTTYRETGK--RQRGDRPG
146 YP_002321513.1     PFGLKNAPAIFQRKMDNCFSR-YSDFIAVYIDDILVFSES-ERDHEKHLR
147 NP_955591.1        --------------------------------------------------
148 YP_001956722.2     LKVSSKYWWPNLRKDVVKVIRQCKQCLVTNAATLAAPPILR-PERPVKPF
149 NP_056902.1        KILARNYK--IPLTVAKQITQECPHCTKQ--GSGPA---GCVMRS----P
150 NP_056886.1        RALSKACN--ISMQQAREVVQTCPHC---NSAPALE---AGVNPRGLGPL
151 NP_041261.1        HTLRLLYK--ITREQARDIVKACKQC--VVATPVPH---LGVNPRGLVPN


001 YP_004442836.1     KFMEICERNGLVLSPTKMKIGTRQVDFLGATIG-NSKIKLQPHIIQKIAE
002 NP_056728.1        MILQKCNQHGIILSKKKAQLFKKKINFLGLEID-EGTHKPQGHILEHINK
003 YP_009508537.1     DKFYMDYIGPLPPSEGYVHVLVVVDAATGFTWLYPTKAQTSKATIKVLT-
004 YP_004732983.2     RMLSICKEHGLVLSPTKMKIAVPGIDFLGAHIR-NSRVSLQPHIIKKIAD
005 YP_009508407.1     ----------------AEKLKRR---------------------------
006 NP_040939.1        NHWQVDYT-HYEN----IILLVWVETNSGLIYAEKVKGESGQEFRIKVMH
007 YP_003864102.1     MLWQMDIT-HCSEFNNLKYIHVSIDTFSGFILATLQTGEATKHVIAHLLH
008 YP_009002585.1     VMLEKCRENGLVLSPTKMKIAQRKVEFLGAILE-AGRIQLQPHIIKKVVN
009 YP_003284237.1     KFMEICEKNGLVLSPTKMKIGVSQVDFLGATIG-QSKIRLQPHIIKKIAD
010 YP_009268869.1     DHWQVDYT-HYED----KIILVWVETNSGLIYAERVKGETGQEFRIMTIR
011 YP_009506251.1     RMLEICKRHGLVLSPNKMSIAQEEIEFLGTIIS-KGRMKLQAHVIKKIVS
012 NP_955611.1        --------------------------------------------------
013 YP_008567619.1     AMLEVCKKNGLILSPTKMKIGLGTIDFLGATIG-NSKVKLQEHIVKKILD
014 NP_149413.1        KMLQICKKNGLILSPSKYKIGVKRVDFLGSTIG-DNQLAVQEHIVSKIAD
015 YP_004442824.1     VFFDICEKNGLVLSPTKMKIGTRQIDFLGATIG-DSRIKLQPHIIKKIAD
016 YP_001931967.1     KVLNRCINLGIILSKKKSQLFKETIDFLGISID-KGTHSPKPHILENIHN
017 YP_002455786.1     HTWQGDVT-HLKHKRTRYCLHVWVDTFSGAVSCVCKKKETSSDLIKTLLH
018 YP_009109694.1     --------------------------------------------------
019 NP_789739.1        --------------------------------------------------
020 NP_955619.1        HIWQGDIT-HFKYKNTLYRLHVWVDTFSGAISATQKRKETSSEAISSLLQ
021 YP_224289.1        KFLEISKANGLILSPTKMKIGVKTIDFLGASIG-NSKIKLQPHIIKKIAD
022 YP_009182100.1     IIINKIVQNGIIIGRTKYELVRESIDFLGVTIC-KGKIKLQPHILTHLME
023 NP_705927.1        --------------------------------------------------
024 YP_009508577.1     DKFFIDYIGPLPPSHGFVYVLVVVDAATGFTWLYPTKAPSTNATITSLN-
025 YP_004442830.1     KMLEICKRNGLILSPTKMKIGVKVIDFLGSTIG-QNQLDLQPHIVQKIVD
026 NP_758887.1        GTWQMDCT-HMEG----KVIIAAVHVASRYLEAEVIPTETGKETAHFLLK
027 YP_009508406.1     -------------------GRSTEPEYLNLYTV-DKPTNKHLATNLTYLN
028 YP_610965.1        VFCAIVEKHGLILSSNKMQLGKREIDFLGATLG-NRKIKLQAHIIKKIAA
029 YP_001931961.1     QVLRKCAELGIILSKKKAQLFKCRINFLGLDID-EGTHRPQNHILEHIHK
030 YP_241114.1        HLWQTDVT-HIPQFGRLKYVHVSIDTFSNFLMASLHTGESTRHCIQHLLF
031 YP_006732334.1     SVLRRVESLGIILSEKKANLFKEKINFLGLEID-RGTHTPQNHILEHLHG
032 YP_009041481.1     RMLQRCKKFGLVLSPTKMKIATREIDFLGATIK-DGRIKLQDHIIKKIST
033 YP_001036293.1     ILLQICQQHGLVLSPTKMKIGTKTIEFLGAVIG-NRKIKLQEHIIKKIAD
034 YP_605811.1        KFLEICKANGLVLSPTKMKIGAREIDFLGATIG-NSRIKLQPHIIKKIIE
035 NP_955577.1        THWEVDFTEIKPGMYGYKYLLVFIDTFSGWAEAYPAKHETAKVVAKKLLE
036 YP_003208050.1     KFLEIVEKEGLVLSPTKMKIAVPEVEFLGAIIG-NSTIKLQPHIIRKIAD
037 NP_789740.1        GIWQLDCT-HLEG----KVILVAVHVASGYIEAEVIPAETGQETAYFLLK
038 YP_004442839.1     KFLEICKANGLVLSPTKMKIGTPEIDFLGATIG-NSKIKLQPHIIKKVIE
039 YP_009507791.1     GIWQIDCT-HLEK----LILLVCIQTTSGFVVVYNLKREDAKETGLAFLN
040 YP_004581513.1     NLAQIVRREGLILSPTKMKIGVKEVDFLGIKIQ-GNKIQLQEHILKKIGD
041 NP_043933.1        IVLNRCKALGIVLSKKKAQLCKTTINFLGLVIE-RGNLKVQSHIGLHLVA
042 YP_009507248.1     ATKPLETVHMDLFTVQNEKYITFIDVFTKYGQAYHLRDGTAISILQALLR
043 YP_002117531.1     QMLHICEKEGLVLSPTKMKIGATSIDFLGATIG-RSKIRLQPHIIKKITE
044 YP_004347415.1     IFRNIIIKHGIVLSKKKAEIGKQKIEFLGVKIE-QGGIELQPHIIDKILE
045 NP_056891.1        MIWQMDVT-HYSEFGNLKYIHVSIDTFSGFLLATLQTGETTKHVITHLLH
046 YP_009513242.1     DKFYIDYIGPLPPSNGYLHVLVVVDGMTGFVWLYPTKAPSTSATVKALN-
047 YP_009352866.1     RMLLRCKKHGLVLSPSKMKIAQKEIEFLGATLT-GGQIKLQDHIVKKIAA
048 YP_009165750.1     AILEQCKNLGIILSKKKAHLFKTKINFLGLEID-QGSHKPQNHILEHIHK
049 NP_040840.1        DHWQVDYT-HYED----KIILVWVETNSGLIYAERVKGETGQEFRVQTMK
050 YP_004222728.1     QIWQTDFT-LEPRMAPRSWLAVTVDTASSAIVVTQHGRVTSVAAQHHWAT
051 YP_009506264.1     KFLEIVKKNGLVLSPTKMKIGVSQIDFLGATIG-QSRIKLQPHIIKKVTE
052 YP_567050.1        HIWQGDVT-HYKYKRNRYCLHVWVDTFSNAVSITCKTKETSSETVSALLH
053 YP_009508571.1     DKFFIDYIGPLPPSNGYLHVLVVVDSMTGFTWLYPTKAPTTNATVKALN-
054 NP_056803.1        DKFFIDYIGPLPPSQGYLYVLVIVDGMTGFTWLYPTKAPSTSATVKSLN-
055 NP_955564.1        VLWQMDVT-HVSEFGKLKYVHVTVDTYSHFTFATARTGEATKDVLQHLAQ
056 YP_009508411.1     QMLSVCEANGLVLSPTKMKIGTSNIEFLGARIG-NHKIQLQEHIIKKIAD
057 YP_009140788.1     IMLGICKRNGLVLSPTKMKIAVQEVEFLGAQIG-NQKIKLQPHVIKKIVE
058 YP_009508410.1     ----------------KLKL------------------------------
059 NP_040333.1        THWEIDFTEVKPGLYGYKYLLVFIDTFSGWVEAFPTKKETAKVVTKKLLE
060 YP_008992013.1     QFYDIVHSHGIMLSAKKSTIATDTIEFLGMIIK-DGHYQPGKHVAQELLH
061 NP_041734.1        IMLKICQENGLVLSPSKICLAQREIEFLGTVIS-QGQMKLQAHVIKKIVN
062 NP_056895.1        HIWQADIT-HYKYKQFTYALHVFVDTYSGATHASAKRGLTTQMTIEGLLE
```

| | |
|---|---|
| 063 YP_009109689.1 | VYWEIDFTEVKPHYAGYKYLLVFVDTFSGWVEAFPTRQETAHIVAKKILE |
| 064 YP_002519387.1 | TVLRKCIETGIILSKKKAEVAKTKINYLGFTIS-NGEIELQSHILENIKL |
| 065 YP_595725.1 | KMLQICKRNGLILSPSKYKIGVKKVDFLGSTIG-DNQLAVQEHIIKKIAE |
| 066 YP_009508551.1 | DKFFIDYIGPLPPSQGYLYVLVVVDGMTGFTWLYPTKAPSTSATVKSLN- |
| 067 YP_001497148.1 | VYWEIDFTEVKPHYAGYKYLLVFVDTFSGWVEAYPTRQETAHVVAKKILE |
| 068 NP_041186.1 | HLWQTDVT-HIPQFGRLKYVHVSIDTFSNFLMASLHTGESTRHCIQHLLF |
| 069 NP_663784.1 | GVWQMDYT-HLEG----KIIIVAVHVASGFIEAEVIPQESGRQTALFLLK |
| 070 YP_009508443.1 | QMLEICRRNGLVLSPTKMKIGTQVVEFLGAVIG-NRKIQLQSHIIKKIAD |
| 071 YP_003987465.1 | IFLQIARKNGLVLSPTKMKVGVQQVDFLGATIG-NSRIRLQPHIIQKVVQ |
| 072 NP_861410.1 | VVLERCKEKGLILSKKKAIIARQTIDFLGLTLQENGEIKLQPNVLEKLEL |
| 073 YP_009345071.1 | RFFERVQQHGLVLSPTKMKIGVRKVDFLGTVIS-NGLLHMQEHILKKIAA |
| 074 YP_009508582.1 | DKFFIDYIGPLPPSHGFAYVLVVVDAATGFTWLYPTKAPSTNATITSLN- |
| 075 NP_569141.1 | KFAHIIENSGISLSKTKAEIMKNQIEFLGIQID-KNGIKMQTHIVQKIIN |
| 076 YP_002916057.1 | KFAQIVEKNGLILSPTKMKIGVKSVDFLGVKIH-QNKVQLQEHILKKIGD |
| 077 NP_777384.2 | HIWQADIT-HYKYKQFTYALHVFVDTYSGATHASAKRGLTTQMTIEGLLE |
| 078 YP_233110.1 | KFLEICKANGLVLSPTKMKIGAKEIDFLGATIG-NSKIKLQPHIIKKIIE |
| 079 NP_542258.1 | VIWQGDVT-HLHYKRYKYCLLVWVDTYSGVVSVSCRRKETGSDCVASLLA |
| 080 YP_009130664.1 | IMLEICKKNGLILSNSKMKIAVSTIDFLGATIG-NRKIKLQEHIIKKIAD |
| 081 YP_009508566.1 | DKYYIDYIGPLPPSEGYSYVLVVVDSATGFCWLYPTKAPSTRATVKSLN- |
| 082 YP_006607892.1 | ALLQRCNQLGIILSKKKAQLFQEKINFLGLEID-QGSHKPQNHILEHIHK |
| 083 YP_009506270.1 | KFMEICERNGLVLSPTKMKIGTRQVDFLGATIG-NSKIKLQPHIIQKIID |
| 084 NP_042513.1 | RVLIQLGNHGVILSKEKAKIAFEEIEFLGLKILKNGFIEPQKHLLEKIAE |
| 085 YP_006907834.1 | MILQKCYQHGIILSKKKAQLFKERINFLGLEID-QGTHRPQSHILEHIQK |
| 086 YP_004442827.1 | KMMEICKKNGLILSPTKMKIGVKVVDFLGSTVG-ENHLDLQPHIVQKIVD |
| 087 YP_009408594.1 | VMLNICKQHGLILSPSKMMIAVPEVYFLGAVLG-KQKLKLQPHIIQKIAN |
| 088 YP_009508561.1 | DKFFIDYIGPLPPSHGFSYVLVIVDARTGFTWLYPTKAPSTNATINSLN- |
| 089 NP_056907.1 | HIWQGDVT-HYKYKRNRYCLHVWVDTFSNAVSITCKTKETSSETVSALLH |
| 090 YP_002308474.1 | TIWQGDVT-HLQYKKYKYCLLVWVDTYSGAVAVSCRRKETSSECVASLLA |
| 091 YP_007761644.1 | TIAKTLKKNGIVISKKKMELGLTKINFLGCEIE-NGRIILQNHVLENLSK |
| 092 NP_040550.1 | QIWQTDFT-LEPRMAPRSWLAVTVDTASSAIVVTQHGRVTSVAAQHHWAT |
| 093 NP_758808.1 | IMLQRCKEHGLVLSPTKMNIAVREVNFLGATIG-SRKVKLQENIIKKILD |
| 094 YP_009345075.1 | RFFQICQKHGLVLSKDKMAIAVPQIEFLGAIIG-NRKIKLQPHIIRKIAN |
| 095 NP_068729.1 | IVLQRIKEKGIIISKKKSKLIQQEIEYLGLKIQGNGEIDLSPHTQEKILQ |
| 096 NP_047255.1 | THWEVDFTEIKPGMYGYKYLLVFIDTFSGWAEAYPAKHETAKVVAKKLLE |
| 097 YP_006273075.1 | AMLEICKRNGLVLSPSKMKIAVQEVEFLGAQIG-NQRIRLQPHVIKKIVE |
| 098 YP_009508888.1 | DKFFIDYIGPLPPSNGYLHVLVVVDAMTGFVWLYPTKAPSANATVKALN- |
| 099 NP_395469.1 | QVLKRCEDEGIILSQPKAKIAHKEIDFFGLHIS-EGEIILQPHILEKLVL |
| 100 YP_009508546.1 | DKFFIDYIGPLPPSQGYLYVLVVVDGMTGFIWLYPTKAPSTSATVKSLN- |
| 101 NP_056762.1 | IFFNRVKEVGCVLSKKKSKMFLKEVEYLGVEIK-EGKISLQPHIVDKIKK |
| 102 YP_009508556.1 | DKFYIDYIGPLPPSNGYLHVLVVVDGMTGFVWLYPTKAPSTSATVKALN- |
| 103 NP_569153.1 | IMLAICQKNGLILSPTKMKIAQAEIEFLGAIIH-KGLIKLQPHIVQKLLT |
| 104 NP_043924.1 | QKLQVDHVGPLGPGTHGYRYLTTMVDVYTGWFWAKPCRGPTTGATIAALE |
| 105 YP_009243641.1 | ALWQMDVT-HIPEFGHLKYVHVVIDTYSRFIWATPQGGETVKHVIAHLLE |
| 106 YP_004300274.1 | KFYHIVKEAGVSLSEKKAIIGVNQIEFLGIEID-KSGVKMQNHIVTKIVQ |
| 107 NP_056848.1 | KFRDITLANGLIISKKKTELCKEKIDFLGVQIE-QGGIELQPHIINKILE |
| 108 YP_006495799.1 | QMLEVCRKNGLVLSPTKMKVAVTTVEFLGAIIG-NGKIKLQPHIVKKISE |
| 109 YP_009508409.1 | -------------------SHKPELEYLDLSSL-EKPPLKSIAHNLAYLN |
| 110 NP_687035.1 | ETWQMDCT-HLEG----KVIIVAVHVASGFIEAEVIPRETGKETAHFLLK |
| 111 NP_045937.2 | PFAHLQIDFVQMCVKKPMYALVIIDVFSKWPEIIPCNKEDAKTVCDILMK |
| 112 NP_612577.1 | NILRRCEKLGIILSKKKAQLFKEKINFLGLEID-QGTHCPQNHILEHIHK |
| 113 NP_619548.1 | AVLKIVEKYGIILSKKKANLFKEKINFLGLEID-KGTHCPQNHILENIHK |
| 114 NP_056880.1 | VLWQMDVT-HVSEFGKLKYVHVTVDTYSHFTFATARTGEATKDVLQHLAQ |
| 115 YP_009121747.1 | IMLGKCQENGLVLSPTKMKIAVPEVEFLGAIIG-RNKIKLQPHIIKKICD |
| 116 YP_009113237.2 | AMLKICKENGLILSPSKMKIGQTKVEFLGAIID-KGKIRLQPNVIKKVCD |
| 117 YP_009508408.1 | KMLAVCQRNGLVLSPTKMKIGTPTIEFLGATIG-NRKIKLQEHIIKKIAD |
| 118 NP_955579.1 | -------------------------------------------------- |
| 119 YP_009229919.1 | IMLEICEKNGLVLSANKMKIAVKEVEFLGAVIC-DRKIKLQPHIIKKITS |
| 120 YP_004442833.1 | EMLRICKANGLILSPTKYKIGVKVVDFLGSTIG-ENQLHLQPHIVQKIID |
| 121 NP_862833.2 | NLWQTDVT-HIPHFGRLKYVHASIDTFSNFLMASLHTGESTRHCIQHLLF |
| 122 NP_040563.1 | FLWQMDNT-HWNK----TIIWVAVETNSGLVEAQVIPEETALQVALCILQ |
| 123 YP_443922.1 | ISIDYTDMGADQVRGGKRYLLVMVDRFSRWVEAIPTAKEDAKSVIKWLQT |
| 124 NP_057933.2 | THWEIDFTEIKPGLYGYKYLLVFIDTFSGWIEAFPTKKETAKVVTKKLLE |
| 125 YP_009116631.1 | IMLGICRDNGLVLSPSKMKLAATEIDFLGATIG-DGRIKLQPHIIKKIAE |
| 126 YP_009513249.1 | DKFYMDYIGPLPPSEGYVHVLVVVDAATGFTWLYPTKAQTSKATIKVLN- |
| 127 YP_223871.1 | EHWEADFTEMITAKGGYKYLLVLVDTFSGWVEAYPAKRETSQVVIKHLIH |
| 128 NP_127504.1 | QFISLVKKFGVMLSAKKMILAQNKIQFLGMDFA-DGTFSPAGHISLELQK |
| 129 NP_044929.1 | QVWYMDYIGPLPPSNGYQHALVIVDAGTGFTWIYPTKAQTANATVKALT- |
| 130 NP_057860.1 | HIWQGDIT-HFKYKNTLYRLHVWVDTFSGAISATQKRKETSSEAISSLLQ |
| 131 NP_040973.1 | GIWQMDCT-HFDG----KIILVGIHVESGYIWAQIISQETADCTVKAVLQ |
| 132 NP_777317.1 | RFFEVCTKEGLVLSPTKMKIGVREVDFLGATIG-NSKIKLQPHIITKIID |
| 133 NP_054716.1 | QKYYMDYIGPLPPSDGYYHVLVIVDEGTGYTWLYPTKAQTANATVKALN- |
| 134 YP_001856242.1 | -------------------------------------------------- |
| 135 NP_954565.2 | MIWQMDVT-HYSEFGNLKYIHVSIDTFSGFLLATLQTGETTKHVITHLLH |

136 NP_569150.1 KFMTICEKNGLVLSPTKMKIGTRQIDFLGATIG-NSKIKLQPHIIKKIIE
137 NP_659397.1 KVLRRIESLGIILSKKKANLFKEKINFLGLEID-RGTHTPQNHILDHIHT
138 YP_009513211.1 VYWEVDFTEVKPGRYGNRYLLVFIDTFSGWVEAFPTKTETALTVCKKILE
139 YP_233107.1 QFITICEENGLVLSPTKMKIGVQQVDFLGATIG-DSKVRLQPHIVKKVLE
140 YP_009109692.1 VYWEIDFTEVKPHYAGYKYLLVFVDTFSGWVEAFPTRQETAHIVAKKILE
141 NP_057849.4 GIWQLDCT-HLEG----KVILVAVHVASGYIEAEVIPAETGQETAYFLLK
142 NP_049560.1 HIWQGDIT-HFKYKNTLYRLHVWVDTFSGAISATQKKKETSSEAISSLLQ
143 YP_009506267.1 KMLSICKEHGLVLSPTKMKIGTPTVEFLGATIG-NSKIKLQEHIIKKIAD
144 NP_039820.1 TMLQLCKENGLILSPTKMKIGTPEIDFLGASLG-CTKIKLQPHIISKICD
145 NP_056790.1 VYWEVDFTEIKPGRYGNKYLLVFIDTFSGWVEAFPTKTETALIVCKKILE
146 YP_002321513.1 VMLQVCQENGLVLSPTKMKVAVKTIEFLGAILG-NQCVQLQPHIIQKIVD
147 NP_955591.1 --------------------------------------------------
148 YP_001956722.2 DKFFIDYIGPLPPSNGYLHVLVVVDSMTGFVWLYPTKAPSTSATVKALN-
149 NP_056902.1 NHWQADCT-HLDN----KIILTFVESNSGYIHATLLSKENALCTSLAILE
150 NP_056886.1 QIWQTDFT-LEPRMAPRSWLAVTVDTASSAIVVTQHGRVTSVAVQHHWAT
151 NP_041261.1 AIWQMDVT-HFTPFGKQRFVHVTVDTFSGFILATPQTGEASKNVISHVIH

001 YP_004442836.1 TKDEELKEFRGLRRWLGILNYARS--YIPNLGKILGPLYSKTNPNGE--R
002 NP_056728.1 FP-DTLEDKKQLQRFLGILTYASD--YIPKLAQIRKPLQAKLKENVP--W
003 YP_009508537.1 -HLTGLAIPKVLHSDQGSAFTSEE------FAQWAKERNIQLEFSTP---
004 YP_004732983.2 KKDDELMTLKGLRSWLGVINYVRQ--YIPKCGTLLGPLYAKTSEHGD--R
005 YP_009508407.1 --------------------------------------------------
006 NP_040939.1 -WYALFG-PESLQSDNGPAFAAEP------TQLLMQYLGVKHTTGIP---
007 YP_003864102.1 -CFSILGPPRQLKTDNGPGYTSRN------FHDFCSKLNIKHTTGIP---
008 YP_009002585.1 FKDEDLQETKGLRSFLGLLNYARA--YIPQMGRLLSPLYAKVSPKGE--K
009 YP_003284237.1 FEDEKLKETKGLRSWLGILNYARN--YIPNLGKTLGPLYSKVSPTGE--K
010 YP_009268869.1 -WYGLFA-PKSLQSDNGPAFVAEP------TQLLMKYLGITHTTGIP---
011 YP_009506251.1 KAQMELSTTKGLRSFLGLLNYARI--YIPNLGRKLSPLYAKVSPTGE--K
012 NP_955611.1 --------------------------------------------------
013 YP_008567619.1 FNTDGLEDKKNLRSWLGILNYARA--YIPNLGRILGPLYAKVSPTGE--R
014 NP_149413.1 FDEERLKTKEGLKSWLATLNYARN--HIKDMGKLLGPLYPKTSEKGA--K
015 YP_004442824.1 IKDEELKEVRGLRKWLGILNYARN--YIPNLGKTLGPLYSKTSPNGE--R
016 YP_001931967.1 FP-ERFKDVNQCRKFLGIITYAMR--YIPELSRKRMFLQDKLKKNVP--W
017 YP_002455786.1 -AISVLGKPFSVNTDNGPAYLSQE------FHEFCTTLCIKHSTHIP---
018 YP_009109694.1 --------------------------------------------------
019 NP_789739.1 --------------------------------------------------
020 NP_955619.1 -AIAYLGKPSYINTDNGPAYISQD------FLNMCTSLAIRHTTHVP---
021 YP_224289.1 FDDHRLKETKGLRAWLGILNYARN--YIPNLGKTLGPLYSKISPNGE--K
022 YP_009182100.1 FP-DKLKDRLQVQQFLGCVNYAGK--FIQRLAELCSVLHTKLKKDHK--W
023 NP_705927.1 --------------------------------------------------
024 YP_009508577.1 -ILLGTAVPRVLHSDQGSAFTSST------FADWAKEKGIQLEFSTP---
025 YP_004442830.1 FSEEKLKTKEGLKSWLAILNYARG--HIKNLGRTLGPLYPKTSEKGE--R
026 NP_758887.1 -LAGRWP-VKHLHTDNGPNFTSEK------VATVCWWAQIEHTTGIP---
027 YP_009508406.1 HQ-EKLFSKVSLKHFYQLHEEIQQ--LKAENLKLRKDLVALTREVVE---
028 YP_610965.1 FPEAQLAEKKGLRSWLGILNYARS--YIPRLGILLGPLYQKTSPHGD--K
029 YP_001931961.1 FP-NKIEDKKQLQRFLGILTYASD--YIPQLASMRAPLQEKLKEDVP--W
030 YP_241114.1 -CFSTSGIPQTLKTDNGPGYTSRS------FQRFCLSFQIQHKTGIP---
031 YP_006732334.1 FP-DRLEDKKQLQRFLGVLTYADS--YIPKLAEKRKPLQVKLKKDQV--W
032 YP_009041481.1 VDEKSLETTKGLRSWLGIINYARG--YIPNCGTLLGPLYSKVGLHGD--K
033 YP_001036293.1 FSDDELKTTKGLRSWLGILNYARA--YMPNMGRILGPLYSKVSPNGE--K
034 YP_605811.1 TKDEELKETKGLRKWLGVLNYARA--YIPNLGKTLGPLYSKTSVNGE--K
035 NP_955577.1 EIFPRYGIPQVLGSDNGPAFISQV------SQSVATLLGINWKLHCA---
036 YP_003208050.1 IPEEQLKEKKGLRGWLGILNYART--YIPNLSTLLGPLYQKTSPHGD--K
037 NP_789740.1 -LAGRWP-VKTIHTDNGSNFTGAT------VRAACWWAGIKQEFGIP---
038 YP_004442839.1 TRDEELKEVKGLRKWLGVLNYARA--YIPNLGKTLGPLYSKTSGNGE--R
039 YP_009507791.1 -LFSQYY-VTEIQTDNGPNFANER------ITGICFHFGIKHKKGIP---
040 YP_004581513.1 FKEKDLLTKKGLRSWLGILNYARQ--YIPNLGKLLGPLYGKTSPTGE--I
041 NP_043933.1 FP-DQLSDRNALQRFLGLLNYISA--YFPKIANLRSPLQVKLKKEIT--W
042 YP_009507248.1 -FCTHHGLPITIVTDNGTEFSNQL------FSEFVRIHKIIHHKTLP---
043 YP_002117531.1 AKEEELQTTKGLRAWLGILNYARS--YIPNLGKTLGPLYSKVSPTGE--K
044 YP_004347415.1 KH-IKIKSKKELQSILGLVNQIRN--FLPNLSKILLPIQKKLKIKNEEVW
045 NP_056891.1 -CFSIIGLPKQIKTDNGPGYTSKN------FQEFCSTLQIKHITGIP---
046 YP_009513242.1 -MLTSIAIPKVLHSDQGAAFTSST------FADWAKEKGIQLEFSTP---
047 YP_009352866.1 VDDKSLQTTKGLRSWLGIINYTRA--YIPNCGTLLGPLYSKVGLHGD--K
048 YP_009165750.1 FP-DRLEDKKQLQRFLGILTYASD--YIPKLAQIRKPFQAKLKKDVT--W
049 NP_040840.1 -WYAMFA-PKSLQSDNGPAFVAES------TQLLMKYLGIEHTTGIP---
050 YP_004222728.1 -AIAVLGRPKAIKTDNGSCFTSKS------TREWLARWGIAHTTGIP---
051 YP_009506264.1 FQEEKLNNTKGLRQFLGILNYARN--YIPNLGKTLGPLYSKISPKGE--K
052 YP_567050.1 -AITILGKPLSINTDNGSAFLSQE------FQAFCASWHIKHSTHVP---
053 YP_009508571.1 -VLTSIAVPKVIHSDQGAAFTSST------FADWAKERGIQLEFSTP---
054 NP_056803.1 -VLTSIAIPKVIHSDQGAAFTSST------FAEWAKERGIHLEFSTP---

| | | |
|---|---|---|
| 055 | NP_955564.1 | -SFAYMGIPQKIKTDNAPAYVSRS------IQEFLARWKISHVTGIP--- |
| 056 | YP_009508411.1 | FPESELMTTKGMRAWLGILNYARS--YIPNLGKTLGPLYSKVSPNGE--K |
| 057 | YP_009140788.1 | FNEAELKEKKGMRSWLGILNYARA--YIPNLGRLLSPLYAKTSPTGD--K |
| 058 | YP_009508410.1 | -------------------------------------------------- |
| 059 | NP_040333.1 | EIFPRFGMPQVLGTDNGPAFVSKV------SQTVADLLGVDWKLHCA--- |
| 060 | YP_008992013.1 | FP-DQQLSKKQVQQFLGIINYIRD--FIPHVDHYTRHLSALLKKKPP--- |
| 061 | NP_041734.1 | KANIELETTKGLRSFLGLLNYARI--YIPNLGRKLSPLYAKTSPTGE--K |
| 062 | NP_056895.1 | -AIVHLGRPKKLNTDQGANYTSKT------FVRFCQQFGISLSHHVP--- |
| 063 | YP_009109689.1 | EIFPRFGLPKVIGSDNGPAFVSQV------SQGLARILGINWKLHCA--- |
| 064 | YP_002519387.1 | FP-SRIPDKKSLQRFLGILTYADQ--YIRKLAEWRKPLQRKLKKDTV--W |
| 065 | YP_595725.1 | FDDEKLKTKEGLKSWLATLNYARN--HIKDMGKLLGPLYPKTSEKGE--R |
| 066 | YP_009508551.1 | -VLTSIAIPKVIHSDQGAAFTSST------FAEWAKERGIHLEFSTP--- |
| 067 | YP_001497148.1 | EIFPRFGLPKVIGSDNGPAFVSQV------SQGLARTLGINWKLHCA--- |
| 068 | NP_041186.1 | -CFSTSGIPQTLKTDNGPGYTSRS------FQRFCLSFQIHHKTGIP--- |
| 069 | NP_663784.1 | -LASRWP-ITHLHTDNGPNFTSQE------VKMVAWWVGIEQSFGVP--- |
| 070 | YP_009508443.1 | FSDEEIKTTKGLRSWLGILNYARN--YIPNLGKILGPLYSKTSPSGE--K |
| 071 | YP_003987465.1 | FDNKDLQTTKGLRSFLGILNYARS--YIPQMGKLLGPLYSKVSPTGE--K |
| 072 | NP_861410.1 | FP-DAIEDRKQLQRFLGCLNYIADKGFLKEIAKETKNLYPKVSITNP--W |
| 073 | YP_009345071.1 | FGPEQYKSKKDLRSWLGLINYARN--HIPNVGRMLGPIYAKTSPQGE--P |
| 074 | YP_009508582.1 | -ILLGTAVPRVLHSDQGSAFTSSA------FADWAKEKGIQLEFSTP--- |
| 075 | NP_569141.1 | LD-ENIDTKKKLQSFLGIVNQVRE--YIPKLAENLKPLQKKLKKDVE--Y |
| 076 | YP_002916057.1 | FREEDLLTKKGLRSWLGILNYARQ--HIPNLGKMLGPLYGKTSPTGE--I |
| 077 | NP_777384.2 | -AIVHLGRPKKLNTDQGANYTSKT------FVRFCQQFGISLSHHVP--- |
| 078 | YP_233110.1 | TKDEELKETKGLRKWLGVLNYARA--YIPNLGKTLGPLYSKTSINGE--K |
| 079 | NP_542258.1 | -AISILGKPHSINTDNGTAYLSQE------FQQFCSSLSIKHSTHIP--- |
| 080 | YP_009130664.1 | YPEKELMNAKGLRSWLGILNYARS--YIPYLGKQLGPLYSKVSPTGE--R |
| 081 | YP_009508566.1 | -FLLGIAVPKILHSDQGSAFTSSD------FANWAKEKEITLEFSTP--- |
| 082 | YP_006607892.1 | FP-DKIEDKKQLQRFLGILTYASD--YIPKLAEIRKPLQSKLKKDVP--W |
| 083 | YP_009506270.1 | IKDEELKEVKGLRKWLGILNYARS--YIPKLGKILGPLYSKTSPNGE--R |
| 084 | NP_042513.1 | FP-DQLQDRKQIQKFLGCLNYIGEKGFFKELAKERKVLQKMLSEKLP--W |
| 085 | YP_006907834.1 | FP-DIIESKLQLQRFLGVLTYASD--YIPKLAQIRKPLQAKLKENVQ--W |
| 086 | YP_004442827.1 | FDEEKLKTKKGLKSWLAILNYARG--HIKNMGKILGPLYPKTSEKGE--K |
| 087 | YP_009408594.1 | FKDADLQEKKGLRSWLGILNYARS--YIPNLGSKLGPLYEKTSPHGD--K |
| 088 | YP_009508561.1 | -LLLGTAIPKVLHSDQGSAFTSAT------FAEWAKDRSIQLEFSTP--- |
| 089 | NP_056907.1 | -AITILGKPLSINTDNGSAFLSQE------FQAFCASWHIKHSTHVP--- |
| 090 | YP_002308474.1 | -AISILGKPHTINTDNGAAYLSQE------FQQFCTSLSIKHTTHVP--- |
| 091 | YP_007761644.1 | FP-SEIKDKKELQSFLGIINYAAS-HYSIEVTKLRVPLQKKLKKNYI--W |
| 092 | NP_040550.1 | -AIAVLGRPKAIKTDNGSCFTSKS------TREWLARWGIAHTTGIP--- |
| 093 | NP_758808.1 | FDTEKLQSKKGLRSFLGILNYARN--HIPNLGKIAGPLYSKTSIYGD--I |
| 094 | YP_009345075.1 | FPDDQLKEKKGLRSWLGILNYARN--YIPKLGQLLGPLYNKTSPMAD--K |
| 095 | NP_068729.1 | FP-DELEDRKQIQRFLGCINYIANEGFFKNLALERKHLQKKISVKNP--W |
| 096 | NP_047255.1 | EIFPRYGIPQVLGSDNGPAFISQV------SQSVATLLGINWKLHCA--- |
| 097 | YP_006273075.1 | FNEAELKEKKGMRSWLGILNYARA--YIPNLGRLLSPLYAKTSPTGD--K |
| 098 | YP_009508888.1 | -MLTSIAVPKVIHSDQGAAFTSST------FADWAKEKGIHLEYSTP--- |
| 099 | NP_395469.1 | FP-DEIENRKQLQRFLGNLNYISEKGFFKDFAKYRKELQKKVSEKVP--W |
| 100 | YP_009508546.1 | -VLTSIAIPKVIHSDQGAAFTSST------FAEWAKERGIHLEFSTP--- |
| 101 | NP_056762.1 | FDKNKLNTLKGLQAYLGLLNYARG--YIKDLSKLVGPLYKKTGKNGQ--R |
| 102 | YP_009508556.1 | -MLTSIAIPKVLHSDQGAAFTSST------FADWAKEKGIQLEFSTP--- |
| 103 | NP_569153.1 | FTNKQLEEVKGLRSWLGLLNYARS--YIPHMGRLLSPLYAKVSPTGE--R |
| 104 | NP_043924.1 | EHISIWGVPYSIQSDNGTAFTSKA------MQEWANTYGIEWKVGAI--- |
| 105 | YP_009243641.1 | -SFAVLGIPQTIKTDNGPAYSSNK------FKRFLTQYKIHHITGIE--- |
| 106 | YP_004300274.1 | CE-EVLDTKKKLQSFLGLINQVRE--YVPNIAKELLFLQKKLKKDVE--Y |
| 107 | NP_056848.1 | KH-TKIKNKTELQSILGLLNQIRH--FIPHLAQILLPIQKKLKIKDEEIW |
| 108 | YP_006495799.1 | VDDESLRTLKGLRSWLGIINYARN--YIPNCGTLLGPLYSKTSENGD--R |
| 109 | YP_009508409.1 | YQ-EKLQSKVLLKHLYTIQEKISE--LEAENKRLRKDLVSLTKEVVE--- |
| 110 | NP_687035.1 | -LLARWP-VKHLHTDNGPNFTSQN------VAAVCWWGNIEHTTGIP--- |
| 111 | NP_045937.2 | DIIPRWGLPDQIDSDQGTHFTAKIS------QELTHSIGVAWKLHCPG-- |
| 112 | NP_612577.1 | FP-DRIEDKKQLQRFLGILTYASD--YIPKLASIRKPLQSKLKEDST--W |
| 113 | NP_619548.1 | FP-DRLEDKKHLQRFLGVLTYAET--YIPKLAEIRKPLQVKLKKDVT--W |
| 114 | NP_056880.1 | -SFAYMGIPQKIKTDNAPAYVSRS------IQEFLARWKISHVTGIP--- |
| 115 | YP_009121747.1 | FDEEKLKTKAGLRSFLGILNYARN--YIPRLSILLGPLYEKTNPHGD--K |
| 116 | YP_009113237.2 | FKMEHLETKTGLRSWLGLLNYARP--YMPDLGKMLGPLYAKVSPNGE--R |
| 117 | YP_009508408.1 | FNDEELKTTKGLRSWLGILNYARN--YIPNLGKILGPLYSKVSPSGE--K |
| 118 | NP_955579.1 | -------------------------------------------------- |
| 119 | YP_009229919.1 | VPEDSLKEKKGLRSWLGILNYART--YIPKLGTLLGPLYEKTSPHGD--K |
| 120 | YP_004442833.1 | FDEEKLKTKKGLKSWLAILNYARG--HIKNMGRILGPLYPKTSEKGE--R |
| 121 | NP_862833.2 | -CFSISGIPQTLKTDNGPGYTSRS------FQRFCLSFQIHHKTGIP--- |
| 122 | NP_040563.1 | -LIQRYT-VLHLHSDNGPCFTAHR------IENLCKYLGITKTTGIP--- |
| 123 | YP_443922.1 | ELIPRYGVPRQIRSDNGSHFSNQH------LRQVEERFGIVHKFGSV--- |
| 124 | NP_057933.2 | EIFPRFGMPQVLGTDNGPAFVSKV------SQTVADLLGIDWKLHCA--- |
| 125 | YP_009116631.1 | VDDESLKTLKGLRSWLGVLNYARN--YIPKCGTLLGPLYSKTSEHGD--R |
| 126 | YP_009513249.1 | -HLTGLAIPKVLHSDQGSAFTSEE------FAQWAKERNIQLEFSTP--- |
| 127 | YP_223871.1 | DIIPRFGLPVQIGSDNGPAFVAKV------TQQLCEALNVSWKLHCA--- |

```
128 NP_127504.1        FP-DTNLSVKQIQQFLGIVNYIRD--FIPEVTEHISPLSDMLKKKPP---
129 NP_044929.1        -HLTGTAVPKVLHSDQGPAFTSSI------LADWAKDRGIQLEHSAP---
130 NP_057860.1        -AIAYLGKPSYINTDNGPAYISQD------FLNMCTSLAIRHTTHVP---
131 NP_040973.1        -LLSAHN-VTELQTDNGPNFKNQK------MEGVLNYMGVKHKFGIP---
132 NP_777317.1        MRDEEIKETRGLRKWLGILNYARS--YIPRLGQTLGPLYSKVSPNGE--K
133 NP_054716.1        -HLTGTAIPKVLHSDQGSAFTSAT------LVAWAKDKGIQMEYSSP---
134 YP_001856242.1     --------------------------------------------------
135 NP_954565.2        -CFSIIGLPKQIKTDNGPGYTSKN------FQEFCSTLQIKHITGIP---
136 NP_569150.1        MKDEELKEVKGLRKWLGILNYARS--YIPKLGKILGPLYAKTSPNGE--R
137 NP_659397.1        FP-DRIEDKKQLQRFLGVLTYADS--YIPKLAEKRKPLQVKLKKDQV--W
138 YP_009513211.1     EILPRFGIPKVLGSDNGPAFVAQV------SQGLATQLGIDWKLHCA---
139 YP_233107.1        TKEESLSETKALRRWLGILNYARA--YIPDLGKILGPLYSKTSGKGE--R
140 YP_009109692.1     EIFPRFGLPKVIGSDNGPAFVSQV------SQGLARILGINWKLHCA---
141 NP_057849.4        -LAGRWP-VKTIHTDNGSNFTGAT------VRAACWWAGIKQEFGIP---
142 NP_049560.1        -AIAYLGKPNYINTDNGPAYISQD------FLDMCTPLAIRHTTHVP---
143 YP_009506267.1     FKEEELKTTKGLRSWLGILNYARS--YIPNLGKTLGPLYSKISPTGE--R
144 NP_039820.1        FSDEKLATPEGMRSWLGILSYARN--YIQDIGKLVQPLRQKMAPTGD--K
145 NP_056790.1        EILPRFGIPKVLGSDNGPAFVAQV------SQGLATQLGINWKLHCA---
146 YP_002321513.1     FDEEHLQTKQGIRSWLGILNYART--YIPNIGTLLAPFYEKTSPHGD--K
147 NP_955591.1        --------------------------------------------------
148 YP_001956722.2     -MLTSIAVPKVIHSDQGAAFTSAT------FADWAKNKGIQLEFSTP---
149 NP_056902.1        -WARLFS-PKSLHTDNGTNFVAEP------VVNLLKFLKIAHTTGIP---
150 NP_056886.1        -AIAVLGRPKAIKTDNGSCFTSKS------TREWLARWGIAHTTGIP---
151 NP_041261.1        -CLATIGKPHTIKTDNGPGYTGKN------FQDFCQKLQIKHVTGIP---
```

```
001 YP_004442836.1     RMNRQDWVIVDKIKDLVKN-LPELELPPNNAMM----IIETDGCMEGWGG
002 NP_056728.1        RWTKEDTLYMQKVKKNLQG-FPPLHHPLPEEKL----IIETDASDDYWGG
003 YP_009508537.1     -YHPQSSGKVERKNSEIKKLLTKLLVGRPLKWYNL--ISSVQLALNNTHV
004 YP_004732983.2     RWHPKDWEIVRQIKKMVQS-LPDLELPPPHAVI----IIESDGCMEGWGG
005 YP_009508407.1     --------------------------------------------------
006 NP_040939.1        -WNPQSQAIVERAHQLLKSTLKKFQPQFV---------------AVESAI
007 YP_003864102.1     -YNPQGQGIVERAHLSLKTTIDKIKKGEW----------YPTKGTPRNIL
008 YP_009002585.1     KLNKEDWKLVRQIKEQVQQ-LPDLELPPQECFI----ILETDGCMEGWGG
009 YP_003284237.1     RMNHQDWVLINKVKRQVKE-LEDMELPPVEACI----VLETDGCMEGWGG
010 YP_009268869.1     -WNPQSQALVERTHQTLKNTIEKFVSMFA---------------SFDSAI
011 YP_009506251.1     KLNKQDWNLINEIKQMVQK-LPDLDIPPIESCI----VIESDGCMEGWGG
012 NP_955611.1        --------------------------------------------------
013 YP_008567619.1     KMNQQDWGIVAQIKKIIQE-LPELELPPEDCCI----VIETDGCMSGWGG
014 NP_149413.1        GLNSEDWKLISRIKTMVRS-LPSLTIPPEDAYI----IIETDACATGWGA
015 YP_004442824.1     RMNSQDWAIVRKIKEQVLN-LPDMELPPKEAVM----IIETDGCMEGWGG
016 YP_001931967.1     TWTSEDTRLLQKLKLSLKE-FPKLHIPQPGQQL----ILETDASQKYWGG
017 YP_002455786.1     -YNPTSSGLVERTNGILKTLLYKYFLDHP-------------DLPLESAV
018 YP_009109694.1     --------------------------------------------------
019 NP_789739.1        --------------------------------------------------
020 NP_955619.1        -YNPTSSGLVERSNGILKTLLYKYFTDKP-------------DLPMDNAL
021 YP_224289.1        RMNAQDWALVTQVKRQVQN-LPELELPPEKCKM----VIETDGCMEGWGG
022 YP_009182100.1     KFTEDDVKRVQLIKKVCAN-LPELTLPPPTWKI----VLQTDASDHHWAG
023 NP_705927.1        --------------------------------------------------
024 YP_009508577.1     -YHPQSSGMVERKNREIKRLITKLLVGRPTKWYPL--LPTIQLALNNTYS
025 YP_004442830.1     SLNSEDWRIIRRMKQEVQK-LPKLSIPPQDAYI----VIETDGSMNGWGG
026 NP_758887.1        -YNPQSQGVIEAKNHHLKQIIGQVRDQAE---------------KLETAV
027 YP_009508406.1     NRPLTEKKVQDLILKITEQ--PKEIEQQAVRLS------EDLKIKLERVE
028 YP_610965.1        RMKPHDWELVAQIKQMVQS-LPDLEVPPKDSFI----VLETDGCMEGWGG
029 YP_001931961.1     NWKHSDTEYVEEIKKSLTD-FPKLHHPATDEKL----IIECDASGKYWGG
030 YP_241114.1        -YNPQGQGIVERAHQRLKHQLLKQKKGNE----------LYSP-SPHNVL
031 YP_006732334.1     SWSQSDTDYIKKIKKGLIN-FPKLYLPKPEDSL----IIETDASDNFWGG
032 YP_009041481.1     RWKSSDWQLVRKVKAAVEN-LPELEIPPKNCHI----IIETDGCMEGWGG
033 YP_001036293.1     RLNAEDWRLIREVRTTIRA-LPDLELPPADSFI----VIEADGCMEGWGG
034 YP_605811.1        RMNSQDWKVVQLIKNQVQN-LPDLDIPPAGATM----VLETDGCMEGWGG
035 NP_955577.1        -YRPQSSGQVERMNRSIKETLTKLTLETGSKDWVL--LLPLVLYRVRNTP
036 YP_003208050.1     RLKAQDYALIRQIKALVQN-LPDLKIPPADSYI----VLETDGCMEGWGG
037 NP_789740.1        -YNPQSQGVVESMNKELKKIIGQVRDQAE---------------HLKTAV
038 YP_004442839.1     RMNSQDWKIVQNIKTQVQR-LPDLEIPPTSAII----ILETDGCMEGWGG
039 YP_009507791.1     -INPQSQALVENFNRTLKVWVDKFKEVTE---------------TLEAAV
040 YP_004581513.1     RFNAQDWKLVREIKRKVQQ-LPPLEIPPKDCCI----VLEADGCMEGWGA
041 NP_043933.1        SWTEKDTETVRKIKSLVKT-LPDLYNPSPEDKP----IIECDASDDHWGA
042 YP_009507248.1     -HSPSDNGNIERFHSTILEHIRILKLQHKDEPIVNLMPYAIIGYNSSIHS
043 YP_002117531.1     RMNHQDWALVKKVKDSVQN-LPDMELPPKQCCI----VIETDGCMDGWGG
044 YP_004347415.1     EWTKEDEQNIIKLKDYCKDNVIKMTYPVEEKDMNW--IIEVDASKEYYGN
045 NP_056891.1        -YNPQGQGIVERAHLSLKTTIEKIKKGEW----------YPRKGTPRNIL
046 YP_009513242.1     -YHPQSSGKVERKNSDIKRLLTKLLIGRPAKWYDL--LPVVQLALNNSYS
```

```
047 YP_009352866.1   RWKASDWQLVKQIKRLVQI-LPTLEIPPKDCHI----IIETDGCMEGWGG
048 YP_009165750.1   SWTDSDTSLMIKIKKGLKS-FPTLYHPKEEDNL----IIECDASDSFWGG
049 NP_040840.1      -WNPQSQALVERTHQTLKNTLEKLIPMFN---------------AFESAL
050 YP_004222728.1   -GNSQGQAMVERANRLLKDKIRVLAEGDG-----F--MKRIPASKQGELL
051 YP_009506264.1   RMNKQDWALVQQIKKQVAN-LPEMELPPRDAVM----ILETDGCMDGWGG
052 YP_567050.1      -YNSTSSGLVERTNGIVKALLNKYLLDSP-------------NLPLDNAI
053 YP_009508571.1   -YHPQSSGKVERKNSDIKRLLTKLLVGRPTKWYDL--LPVVQLALNNSYS
054 NP_056803.1      -YHPQSSGKVERKNSDIKRLLTKLLVGRPTKWYDL--LPVVQLALNNTYS
055 NP_955564.1      -YNPQGQAIVERTHQNIKAQLNKLQKAGK----------YYTP---HHLL
056 YP_009508411.1   RMNAQDWALVRKVKISIQN-LPELEIPPKNCVI----IIETDGCMEGWGG
057 YP_009140788.1   RMNSQDWKLVSQIKEEVQK-LPDLEIPPEDSFI----ILETDGCMTGWGG
058 YP_009508410.1   --------------------------------------------------
059 NP_040333.1      -YRPQSSGQVERMNRTIKETLTKLTLATGSRDWVL--LLPLALYRARNTP
060 YP_008992013.1   EWNDDHTNAVTTLKKIAQNPPPKHQTHPTKKIL------QTDASDQYWSA
061 NP_041734.1      RFNRQDWHLIKEIKDMVQK-LPNLAIPPARCYI----IIESDGCMEGWGA
062 NP_056895.1      -YNPTSSGLVERTNGLLKLLLSKYHLDEP-------------HLPMTQAL
063 YP_009109689.1   -YRPQSSGQVERMNRTIKETLTKLTLETGLKDWRR--LLSLALLRARNTP
064 YP_002519387.1   EWNDSDTQYVEKFKRNLKE-FPKLHHPLPDEYL----IIETDASHEHWAG
065 YP_595725.1      RLNSEDWKLINRIKTMVRT-LPNLTIPPEDAYI----IIETDACATGWGA
066 YP_009508551.1   -YHPQSSGKVERKNSDIKRLLTKLLVGRPTKWYDL--LPVVQLALNNTYS
067 YP_001497148.1   -YRPQSSGQVERMNRTIKETLTKLTLETGLKDWRR--LLSLALLRARNTP
068 NP_041186.1      -YNPQGQGIVERAHQRIKHQLLKQKKGNE----------LYSP-SPHNAL
069 NP_663784.1      -YNPQSQGVVEAMNHHLKNQISRIREQAN---------------TIETIV
070 YP_009508443.1   RMNAQDWELVRKVKKIVNE-LPDLIIPPEKCCI----IIETDGCMSGWGG
071 YP_003987465.1   RMNKQDWAIIEKIKQMVEQ-LPELELPPNGSVI----VIETDGCMEGWGG
072 NP_861410.1      HWSDLDSKLVNQIKKKCKD-LSPLYFPKPEDYL----IIETDASGDTWAG
073 YP_009345071.1   KMNAQDWKIIQDIQAQIHQ-LPTLEIPDKDAVI----IIETDGCMTGWGA
074 YP_009508582.1   -YHPQSSGMVERKNREIKRLITKLLVGRPTKWYPL--IPTIQLALNNTYS
075 NP_569141.1      SFDEKDKEQIKKIKILCKK-LPKLYFPDENKKFTY--IVETDSSNYSYGG
076 YP_002916057.1   RFNAQDWKLVREIKRKIQQ-LPPLEIPPKDCCI----VLEADGCMDGWGA
077 NP_777384.2      -YNPTSSGLVERTNGLLKLLLSKYHLDEP-------------HLPMTQAL
078 YP_233110.1      KMNSQDWKVVQLIKNQVQN-LPDLDIPPAEATM----VLETDGCMEGWGG
079 NP_542258.1      -YNPTSSGLVERTNGILKTLISKYLLDNH-------------HLPLETAI
080 YP_009130664.1   KMNQQDWAIVKQIKEKVQN-LPDLELPPPDCHI----VLETDGCMDGWGG
081 YP_009508566.1   -YHPQSSGKVERKNQEIKKLLTKLLVGRPAKWYPL--IPSVQLALNNTYS
082 YP_006607892.1   TWTHDDINYMSKIKKNLKS-FPTLYHPKLTDTL----IIETDASGSFWGG
083 YP_009506270.1   RMNSQDWKIVKEVKEIVAN-LPDLELPPEKAIM----IIETDGCMDGWGG
084 NP_042513.1      KWNDLATLAVKRLKQVCKN-LPRLYVAKPSDLL----ILTTDASDTTWGA
085 YP_006907834.1   RWTPEDTLYMKKVKKNLNG-FPPLHHPLPEEKL----IIETDASDNYWGG
086 YP_004442827.1   RLNSEDWKIIRKMKEEVRR-LPKMAIPPEDAYI----VIETDGSMNGWGG
087 YP_009408594.1   RFKPSDWALVKELKAQIQN-LPDLEIAPASAYI----VLETDGSMTGWGG
088 YP_009508561.1   -YHPQSSGMVERKNKEIKRLITKLLVGRPTKWYPL--LPTVQLALNNTYS
089 NP_056907.1      -YNSTSSGLVERTNGIVKALLNKYLLDSP-------------NLPLDNAI
090 YP_002308474.1   -YNPTSSGLVERTNGILKTLISKYLLDDH-------------HLPLDTAI
091 YP_007761644.1   SWTEQDKQIVEQIKTICQN-LPALELPKNGDKL----VLTTDASDKHWAG
092 NP_040550.1      -GNSQGQAMVERANRLLKDKIRVLAEGDG-----F--MKRIPTSKQGELL
093 NP_758808.1      RFSASDWKLINEIKAIVEK-LPPLDYPPEQAYI----IIESDGCMEGWGA
094 YP_009345075.1   RMTASDWNLVKKIKEKVQA-LPDMELPPKDAFV----IIETDGCTEGWGA
095 NP_068729.1      KWDTIDTKMVQSIKGKIQS-LPKLYNASIQDFL----IVETDASQHSWSG
096 NP_047255.1      -YRPQSSGQVERMNRSIKETLTKLTLETGSKDWVL--LLPLVLYRVRNTP
097 YP_006273075.1   RMNAQDWKLVAQIKEEVQK-LPDLELPPEDCFI----ILETDGCMTGWGG
098 YP_009508888.1   -YHPQSSGKVERKNSDIKRLLTKLLVGRPTKWYDL--LSTVQLALNNAYS
099 NP_395469.1      KWTSYDTTQVQALKTLCQR-LPKLYNAKESDLL----IIATDASNGHWGA
100 YP_009508546.1   -YHPQSSGKVERKNSDIKRLLTKLLVGRPTKWYDL--LPVVQLALNNTYS
101 NP_056762.1      IFNKEDWNIIFKIEREVSK-IKPLERPKETDYI----IIETDASEEGWGA
102 YP_009508556.1   -YHPQSSGKVERKNSDIKRLLTKLLIGRPAKWYDL--LPVVQLALNNSYS
103 NP_569153.1      RMNRQDWALIDKIRAQVQN-LPALELPPADCFI----IIETDGCMDGWGG
104 NP_043924.1      -YHPQSQGKVERKHRLLKDRLKRATHEGKNWVQALPSILLFINSMHPRDQ
105 YP_009243641.1   -YNPQGQAIIERAHGTLKLYIKKLKKRGERELELP--PLIALQNNPRAVL
106 YP_004300274.1   HFDSQDQEKLKKIKEKCSN-LPKLLFPDETKQFDW--IVETDASEISYGG
107 NP_056848.1      TWTKEDEEKIKLIQDYSKNLVIKMKYPINKEDMNW--IIEVDASNNAYGS
108 YP_006495799.1   RMSPQDWKLVRKVKELVKS-LPDLELPPAGAYV----VIETDGCMEGWGG
109 YP_009508409.1   NRPLTERKVQELIVNITSQ--PKEIEQQAVKLS------KDLQLKLDRVE
110 NP_687035.1      -YNPQSQGSVESMNRQLKEIISQIRDDCE---------------RLETAV
111 NP_045937.2      --HPRSSGIVERTNRTLKSKIIKAQEQLQLSKWTE--VLPYVLLEMRATP
112 NP_612577.1      TWNDTDSQYMAKIKKNLKS-FPKLYHPEPNDKL----VIETDASEEFWGG
113 NP_619548.1      NWTQSDSDYVKKIKKNLGS-FPKLYLPKPEDHL----IIETDASDSFWGG
114 NP_056880.1      -YNPQGQAIVERTHQNIKAQLNKLQKAGK----------YYTP---HHLL
115 YP_009121747.1   RMKPSDYEIVRKIKAKVQQ-LPDLQIPPDDAYI----TIETDGCMDGWGG
116 YP_009113237.2   RFNREDRELLRKLTEKIRN-LPDLELPPPNAYI----IIESDGCMTGWGG
117 YP_009508408.1   RMNGQDWELVRKVKQLVQK-LPDLEIPPKDCCI----LIETDGCMDGWGG
118 NP_955579.1      --------------------------------------------------
119 YP_009229919.1   RMKPSDWELVKKIKAQVQN-LPDMEIPPEKAHI----VIETDGSMEGWGA
```

```
120 YP_004442833.1     RLNSEDWKIIRGMKQEVKN-LPKLAIPPENAYI----VIETDGSMNGWGG
121 NP_862833.2        -YNPQGQGIVERAHQRLKHQLLKQKKGSD----------LYSP-SPHNAL
122 NP_040563.1        -YNPQSQGVVERAHRDLKDRLAAYQGDCE---------------TVEAAL
123 YP_443922.1        -YKPQSQGLVERCNQTLKAKIAKVCAGTKLTWVEA--LPLALMAMRSSPG
124 NP_057933.2        -YRPQSSGQVERMNRTIKETLTKLTLATGSRDWVL--LLPLALYRARNTP
125 YP_009116631.1     RWHASDWALVKRIKGLVQN-LPDLKLPTEEAYM----IIETDGCMEGWGG
126 YP_009513249.1     -YHPQSSGKVERKNSEIKKLLTKLLVGRPLKWYNL--ISSVQLALNNTHV
127 YP_223871.1        -YRPQSSGQVERMNRTLKETIAKLRKETGG-DWVS--LLPQALLRARCTP
128 NP_127504.1        AWGKCQDNAVKQLKQLAQQVKSLHIPSEGKKIL------QTDASDQYWSA
129 NP_044929.1        -YHPQSSGKVERKNSEIKRLLTKLLAGRPTKWYPL--IPIVQLALNNTPN
130 NP_057860.1        -YNPTSSGLVERSNGILKTLLYKYFTDKP-------------DLPMDNAL
131 NP_040973.1        -GNPQSQALVENVNHTLKVWIRKFLPETT---------------SLDNAL
132 NP_777317.1        RMNSQDWAIVKKIKQQVQN-LPDLELPPQEAIM----VIEADGCMEGWGG
133 NP_054716.1        -YHPQSSGKVERKNSEIKRLLTKLLVGRPTKWYPL--IPTVQLALNNTPN
134 YP_001856242.1     --------------------------------------------------
135 NP_954565.2        -YNPQGQGIVERAHLSLKTTIEKIKKGEW----------YPRKGTPRNIL
136 NP_569150.1        RMNTQDWKIVKEVKEVVAN-LPELELPPEKAIM----IIETDGCMEGWGG
137 NP_659397.1        IWTQSDTDYVKKIKKGLIN-FPKLYLPKKEDSL----IIETDASDHFWGG
138 YP_009513211.1     -YRPQSSGQVERMNRTIKETLTKLALETGGKDWVT--LLPLALLRARNTP
139 YP_233107.1        KLNHQDMKIIHQIKEKVKN-LPELEVPPPESII----LIETDGCMDGWGG
140 YP_009109692.1     -YRPQSSGQVERMNRTIKETLTKLTLETGLKDWRR--LLSLALLRARNTP
141 NP_057849.4        -YNPQSQGVVESMNKELKKIIGQVRDQAE---------------HLKTAV
142 NP_049560.1        -YNPTSSGLVERSNGILKTLLYKYFTDKP-------------DLPMDNAL
143 YP_009506267.1     RMNDQDWDLIRKIKRQIHN-LPDLEIPPEDAFI----ILEVDGCMEGWGG
144 NP_039820.1        RMNPETWKMVRQIKEKVKN-LPDLQLPPKDSFI----IIETDGCMTGWGA
145 NP_056790.1        -YRPQSSGQVERMNRTIKETLTKLALETGGKDWVT--LLPLALLRARNTP
146 YP_002321513.1     RLKPSDRELVKKIKEMVQH-LPPLELPPPDAYI----ILETDGSMEGWGG
147 NP_955591.1        --------------------------------------------------
148 YP_001956722.2     -YHPQSSGKVERKNSDIKRLLTKLLVGRPAKWYDL--LPVVQLALNNSYS
149 NP_056902.1        -YHPESQGIVERANRTLKEKIQSHRDNTQ---------------TLEAAL
150 NP_056886.1        -GNSQGQAMVERANRLLKDRIRVLAEGDG-----F--MKRIPTSKQGELL
151 NP_041261.1        -YNPQGQGVVERAHQTLKNALNRLARSPL----------GFSMQQPRNLL
```

```
001 YP_004442836.1     VCKWKEQSGQPRWSEK----------------------------------
002 NP_056728.1        MLKAIKINEGTNT-------------------------------------
003 YP_009508537.1     VSTKYTPHQLMFG-------------------------------------
004 YP_004732983.2     ICKWKNSKGESKGKER----------------------------------
005 YP_009508407.1     --------------------------------------------------
006 NP_040939.1        A-------------------------------------------------
007 YP_003864102.1     N-------------------------------------------------
008 YP_009002585.1     VCKWKKKEYDPKNTER----------------------------------
009 YP_003284237.1     VCKWKLPGQN-KASEK----------------------------------
010 YP_009268869.1     A-------------------------------------------------
011 YP_009506251.1     ICKWKKAKEDSRTTGR----------------------------------
012 NP_955611.1        --------------------------------------------------
013 YP_008567619.1     VCKWKPMACDPKNTER----------------------------------
014 NP_149413.1        VCKWKKSKADPRSSEL----------------------------------
015 YP_004442824.1     VCKWKKNISQPRLEER----------------------------------
016 YP_001931967.1     ILKAEVIHSNN-EI------------------------------------
017 YP_002455786.1     S-------------------------------------------------
018 YP_009109694.1     --------------------------------------------------
019 NP_789739.1        --------------------------------------------------
020 NP_955619.1        S-------------------------------------------------
021 YP_224289.1        VCKWTTVG---KAQEK----------------------------------
022 YP_009182100.1     TIGAISPDKKEE--------------------------------------
023 NP_705927.1        --------------------------------------------------
024 YP_009508577.1     VHYKKTPHQLLFG-------------------------------------
025 YP_004442830.1     ICCWKKSKNDPRSTEQ----------------------------------
026 NP_758887.1        Q-------------------------------------------------
027 YP_009508406.1     TLLKKIEGSVL---------------------------------------
028 YP_610965.1        ICKWKQAKKDPRGKEK----------------------------------
029 YP_001931961.1     ILKAIHQSE-----------------------------------------
030 YP_241114.1        N-------------------------------------------------
031 YP_006732334.1     VLKAQTTEGQ----------------------------------------
032 YP_009041481.1     VCKWCPPNKRTKGEER----------------------------------
033 YP_001036293.1     VCKWKKLKKDFRRNER----------------------------------
034 YP_605811.1        VCKWKLHPSDTRLAEK----------------------------------
035 NP_955577.1        GPHGLTPFEILYGAPPPMAHFF----------------------------
036 YP_003208050.1     VCKWKPNKHSPKSQEK----------------------------------
037 NP_789740.1        Q-------------------------------------------------
038 YP_004442839.1     ICKWKAKPSDTRLNEK----------------------------------
```

```
039 YP_009507791.1    Q-------------------------------------------------
040 YP_004581513.1    VCKWKQSAYDPRSKER----------------------------------
041 NP_043933.1       ILKAKLPEGKE---------------------------------------
042 YP_009507248.1    FTKCRPFDLLNGHFDPRD--------------------------------
043 YP_002117531.1    VCKWKLTRHDSKTTEK----------------------------------
044 YP_004347415.1    CLKYK--KDKIEY-------------------------------------
045 NP_056891.1       N-------------------------------------------------
046 YP_009513242.1    PSSKYTPHQLLFG-------------------------------------
047 YP_009352866.1    ICKWCPIRKKAKKEER----------------------------------
048 YP_009165750.1    ILKAKTIDEDK---------------------------------------
049 NP_040840.1       A-------------------------------------------------
050 YP_004222728.1    A-------------------------------------------------
051 YP_009506264.1    ICKWKMPGAT-KASEK----------------------------------
052 YP_567050.1       S-------------------------------------------------
053 YP_009508571.1    PSLKHTPHQLLFG-------------------------------------
054 NP_056803.1       PVLKYTPHQLLFG-------------------------------------
055 NP_955564.1       A-------------------------------------------------
056 YP_009508411.1    VCKWKLFAKDPKATEK----------------------------------
057 YP_009140788.1    VCKWKPKRNDPRRSER----------------------------------
058 YP_009508410.1    --------------------------------------------------
059 NP_040333.1       GPHGLTPYEILYGAPPPLVNFP----------------------------
060 YP_008992013.1    VLLEEHNGKRH---------------------------------------
061 NP_041734.1       VCKWKLAKEDSRTTEK----------------------------------
062 NP_056895.1       S-------------------------------------------------
063 YP_009109689.1    NRFGLTPYEILYGGPPPLSTLL----------------------------
064 YP_002519387.1    VLKSRGTDN-----------------------------------------
065 YP_595725.1       VCKWKKNKADPRNTEQ----------------------------------
066 YP_009508551.1    PVLKYTPHQLLFG-------------------------------------
067 YP_001497148.1    NRFGLTPYEILYGGPPPLSTLL----------------------------
068 NP_041186.1       N-------------------------------------------------
069 NP_663784.1       L-------------------------------------------------
070 YP_009508443.1    VCKWKKFLKDPRVTEQ----------------------------------
071 YP_003987465.1    ICKWKFPGAP-RNQEK----------------------------------
072 NP_861410.1       CLKAAEL-------------------------------------------
073 YP_009345071.1    ICKWKKKEKDPKSTEK----------------------------------
074 YP_009508582.1    VHYKKTPHQLLFG-------------------------------------
075 NP_569141.1       VLKYRYNKEKIEH-------------------------------------
076 YP_002916057.1    ICKWKQFAYDPRSKER----------------------------------
077 NP_777384.2       S-------------------------------------------------
078 YP_233110.1       VCKWKLHPSDTRLAEK----------------------------------
079 NP_542258.1       S-------------------------------------------------
080 YP_009130664.1    VCKWKPQKCDGRQMEK----------------------------------
081 YP_009508566.1    PKIKLTPHQLLFG-------------------------------------
082 YP_006607892.1    ILKARTENSDS---------------------------------------
083 YP_009506270.1    VCKWKDNSSQPRWSEK----------------------------------
084 NP_042513.1       VLMAVPNAYEEFLSFVTNNRPSDLKTFLKQKEGSSRYAEKSVPEKLNSMH
085 YP_006907834.1    ILKAIHIDLSTNES------------------------------------
086 YP_004442827.1    VCYWKKSKADPRSTEQ----------------------------------
087 YP_009408594.1    VCKWKKAKGDPRSTER----------------------------------
088 YP_009508561.1    PRTKLTPHKLLFG-------------------------------------
089 NP_056907.1       S-------------------------------------------------
090 YP_002308474.1    S-------------------------------------------------
091 YP_007761644.1    VLQFYRKIEQEVFEK-----------------------------------
092 NP_040550.1       A-------------------------------------------------
093 NP_758808.1       ICKWKLAEYDPKSSEQ----------------------------------
094 YP_009345075.1    VAKWRPFKGASRGTEK----------------------------------
095 NP_068729.1       CLRALPKGKQKIGLDEFGIPTADLCTGS-------------------SSA
096 NP_047255.1       GPHGLTPFEILYGAPPPMAHFF----------------------------
097 YP_006273075.1    ICKWKPKRNDPRRMEK----------------------------------
098 YP_009508888.1    PILKHTPHQLLFG-------------------------------------
099 NP_395469.1       VMTAVTP----VHITNYGISLEDLF-------------------------
100 YP_009508546.1    PVLKHTPHQLLFG-------------------------------------
101 NP_056762.1       VLVCKPDKYSGKDTEK----------------------------------
102 YP_009508556.1    PSSKYTPHQLLFG-------------------------------------
103 NP_569153.1       VCKWKVAQYDPRSSER----------------------------------
104 NP_043924.1       FSAYELMTGRVPHLGG----------------------------------
105 YP_009243641.1    S-------------------------------------------------
106 YP_004300274.1    VLKYKYHQDKIEY-------------------------------------
107 NP_056848.1       CLKYKPKNSKIEY-------------------------------------
108 YP_006495799.1    VCKWKHKKGESASAEK----------------------------------
109 YP_009508409.1    DILRRLEGAVT---------------------------------------
110 NP_687035.1       Q-------------------------------------------------
111 NP_045937.2       KKHGLSPHEIVMG-------------------------------------
```

```
112 NP_612577.1        ILKAIHNSH-----------------------------------------
113 NP_619548.1        VLKARALDGV----------------------------------------
114 NP_056880.1        A-------------------------------------------------
115 YP_009121747.1     ICKWKKKKEDPRRDEK----------------------------------
116 YP_009113237.2     VCKWKTGESDPRTTER----------------------------------
117 YP_009508408.1     VCKWKAFTHDPRKDER----------------------------------
118 NP_955579.1        --------------------------------------------------
119 YP_009229919.1     VCKWKPNQSDPKSTER----------------------------------
120 YP_004442833.1     VCLWKKSKNDPKSTEQ----------------------------------
121 NP_862833.2        N-------------------------------------------------
122 NP_040563.1        S-------------------------------------------------
123 YP_443922.1        AGTHLSPHEIMTGRVMPGPPRE----------------------------
124 NP_057933.2        GPHGLTPYEILYGAPPPLVNFP----------------------------
125 YP_009116631.1     VCKWKPMKADSASKEE----------------------------------
126 YP_009513249.1     VSTKYTPHQLMFG-------------------------------------
127 YP_223871.1        GREGLSPFEILYGLKPPVVPRV----------------------------
128 NP_127504.1        VLLEEHNGKRK---------------------------------------
129 NP_044929.1        TRQKYTPHQLMYG-------------------------------------
130 NP_057860.1        S-------------------------------------------------
131 NP_040973.1        S-------------------------------------------------
132 NP_777317.1        ICKWKIFG--PRTSEK----------------------------------
133 NP_054716.1        AKIGKTPHQLMYG-------------------------------------
134 YP_001856242.1     --------------------------------------------------
135 NP_954565.2        N-------------------------------------------------
136 NP_569150.1        VCKWKTDSLQPRWSEK----------------------------------
137 NP_659397.1        VLKAQTTEGE----------------------------------------
138 YP_009513211.1     GQFGLTPYEILHGGPPPVLASG----------------------------
139 YP_233107.1        ICKWKLNKGEPRSAEK----------------------------------
140 YP_009109692.1     NRFGLTPYEILYGGPPPLSTLL----------------------------
141 NP_057849.4        Q-------------------------------------------------
142 NP_049560.1        S-------------------------------------------------
143 YP_009506267.1     VCKWRPKKNDPKSTEK----------------------------------
144 NP_039820.1        VCKWKMSKHDPRSTER----------------------------------
145 NP_056790.1        GRFGLTPYEILYGGPPPILESG----------------------------
146 YP_002321513.1     ICKWKHKKNDPRSSER----------------------------------
147 NP_955591.1        --------------------------------------------------
148 YP_001956722.2     PSSKYTPHQLLFG-------------------------------------
149 NP_056902.1        Q-------------------------------------------------
150 NP_056886.1        A-------------------------------------------------
151 NP_041261.1        S-------------------------------------------------
```

```
001 YP_004442836.1     ----------------------------------ICAYASGKFNPIKSTI
002 NP_056728.1        --------------------------------ELICRYASGSFKAAEKNY
003 YP_009508537.1     ---------------------------------IDCNLPFANKDTLDWTR
004 YP_004732983.2     ----------------------------------ICAYASGKFPTVKSTI
005 YP_009508407.1     --------------------------------------------------
006 NP_040939.1        ------------------------------------------AALVAINI
007 YP_003864102.1     ------------------------------------------HALFILNF
008 YP_009002585.1     ----------------------------------ICAYASGKFHPIKSTI
009 YP_003284237.1     ----------------------------------ICAYASGKFSPIKSTI
010 YP_009268869.1     ------------------------------------------AALITLNI
011 YP_009506251.1     ----------------------------------ICAYASGKFGVIKSTI
012 NP_955611.1        --------------------------------------------------
013 YP_008567619.1     ----------------------------------VCAYCSGKFDPPKSTI
014 NP_149413.1        ----------------------------------ICRYASGKFDKPKGTC
015 YP_004442824.1     ----------------------------------VCAYASGKFTPIKSTI
016 YP_001931967.1     -------------------------------TEEICCYASGTFKQAELNY
017 YP_002455786.1     ------------------------------------------KALWTINH
018 YP_009109694.1     --------------------------------------------------
019 NP_789739.1        --------------------------------------------------
020 NP_955619.1        ------------------------------------------IALWTINH
021 YP_224289.1        ----------------------------------VCAYASGKFTPIKSTI
022 YP_009182100.1     ---------------------------------VICAYRSGTFKPAEQNY
023 NP_705927.1        --------------------------------------------------
024 YP_009508577.1     ---------------------------------VDGNVPFANQDTLDLTR
025 YP_004442830.1     ----------------------------------ICRYASGKFAKPKSTI
026 NP_758887.1        ------------------------------------------MAVLIHNF
027 YP_009508406.1     ----------------------------------L---------------
028 YP_610965.1        ----------------------------------VAAYASGRFPVPKSTI
029 YP_001931961.1     --------------------------------ERICRYTSGSFKKAELNY
030 YP_241114.1        ------------------------------------------HALYVLNF
```

```
031 YP_006732334.1  --------------------------------ELIC---------STVQE
032 YP_009041481.1  ----------------------------------VCAYVSGKFPTVKSTI
033 YP_001036293.1  ----------------------------------VCAYASGKFDPPKSTI
034 YP_605811.1     ----------------------------------VCAYASGRYHPIKSTI
035 NP_955577.1     --------------------------------DTDISSFATSPT-MQAHL
036 YP_003208050.1  ----------------------------------VAAYASGKFPVVKSTI
037 NP_789740.1     ------------------------------------------MAVFIHNF
038 YP_004442839.1  ----------------------------------ICAYASGSYNPIKSTI
039 YP_009507791.1  ------------------------------------------LAVHALNH
040 YP_004581513.1  ----------------------------------IAAYASGKFQPIKSTI
041 NP_043933.1     ---------------------------------VICRYASGTFKPAEKNY
042 YP_009507248.1  --------------------------------PLDIDLTEHILQQYAQNH
043 YP_002117531.1  ----------------------------------TCAYASGKFSPVKSTI
044 YP_004347415.1  ----------------------------------ICRYNSGTFKEHEKNY
045 NP_056891.1     ------------------------------------------HALFILNF
046 YP_009513242.1  ---------------------------------VDSNTPFANSDTLDLSR
047 YP_009352866.1  ----------------------------------ICAYVSGKFLTVKSTI
048 YP_009165750.1  --------------------------------EYICRYTSGSFKQAELNY
049 NP_040840.1     ------------------------------------------GTLITLNI
050 YP_004222728.1  ------------------------------------------KAMYALNH
051 YP_009506264.1  ----------------------------------VCAYASGRFPAVKSTI
052 YP_567050.1     ------------------------------------------KALWTLNQ
053 YP_009508571.1  ---------------------------------IDSNTPFANQDTLDLTR
054 NP_056803.1     ---------------------------------IDSNTPFANQDTLDLTR
055 NP_955564.1     ------------------------------------------HALFVLNH
056 YP_009508411.1  ----------------------------------VCAYASGKFNPPKSTI
057 YP_009140788.1  ----------------------------------ICAYASGKFNPVKSTI
058 YP_009508410.1  --------------------------------------------------
059 NP_040333.1     --------------------------------DPDMAKVTHNPS-LQAHL
060 YP_008992013.1  ----------------------------------FCAHASGQFKHSEKNY
061 NP_041734.1     ----------------------------------ICAYASGKFGVVKSTI
062 NP_056895.1     ------------------------------------------RALWTHNQ
063 YP_009109689.1  --------------------------------N-SFSPSNSKTD-LQARL
064 YP_002519387.1  -------------------------------LERLCRYTSGSFKPAEINY
065 YP_595725.1     ----------------------------------ICRYASGKFDKPKGTC
066 YP_009508551.1  ---------------------------------IDSNTPFANQDTLDLTR
067 YP_001497148.1  --------------------------------N-SFSPSDPKTD-LQARL
068 NP_041186.1     ------------------------------------------HALYVLNF
069 NP_663784.1     ------------------------------------------MAVHCMNF
070 YP_009508443.1  ----------------------------------VCAYASGRFDPPKSTI
071 YP_003987465.1  ----------------------------------VCAYASGRFQPIKSTI
072 NP_861410.1     --------------------LFPKGTKNKVVER-LCKYTSGIFSSAEQKY
073 YP_009345071.1  ----------------------------------ICAYASGPYNPLKSTI
074 YP_009508582.1  ---------------------------------VDGNVPFANQDTLDLTR
075 NP_569141.1     ----------------------------------HCRYYSGSYTEPQEKW
076 YP_002916057.1  ----------------------------------ITAYASGKFQPIKSTI
077 NP_777384.2     ------------------------------------------RALWTHNQ
078 YP_233110.1     ----------------------------------VCAYASGRYHPIKSTI
079 NP_542258.1     ------------------------------------------KSLWTINH
080 YP_009130664.1  ----------------------------------ICAYASGKFNPPKSTI
081 YP_009508566.1  ---------------------------------VDGNIPFANSDTLDLKR
082 YP_006607892.1  --------------------------------ELICRYTSGSFKAAELNY
083 YP_009506270.1  ----------------------------------ICAYASGKFTPIKSTI
084 NP_042513.1     DKSHGEKAEKSVSRQTSQGRSSSQSQQSYIGSFMITKWSSGTFKPAEENY
085 YP_006907834.1  -------------------------------IELVCRYASGSFKPAEQNY
086 YP_004442827.1  ----------------------------------VCRYASGKFNKPKSTI
087 YP_009408594.1  ----------------------------------VCAYANGKFPTVKSSI
088 YP_009508561.1  ---------------------------------VDGNVPFANQDTLDLTR
089 NP_056907.1     ------------------------------------------KALWTLNQ
090 YP_002308474.1  ------------------------------------------KTLWTINH
091 YP_007761644.1  -------------------------------DLRVSRYCSGTWNQTEQNW
092 NP_040550.1     ------------------------------------------KAMYALNH
093 NP_758808.1     ----------------------------------ICAYASGKFSPIKSTI
094 YP_009345075.1  ----------------------------------AFAYASGRFPALQSTI
095 NP_068729.1     SSDNSPAEIDKCHSASKQDTHVASKIKKLENELLLCKYVSGTFTDTETRY
096 NP_047255.1     --------------------------------DTDISSFATSPT-MQAHL
097 YP_006273075.1  ----------------------------------VCAYASGKFSPIKSTI
098 YP_009508888.1  ---------------------------------VDANIPFANQDTLDLTR
099 NP_395469.1     -----PKEQHTAQALSSQYQFFG--TKDFVQKELLTKYASGTFTDTEKRY
100 YP_009508546.1  ---------------------------------IDSNTPFANQDTLDLTR
101 NP_056762.1     -----------------------------------IAGYASGNFGEKKTW
102 YP_009508556.1  ---------------------------------VDSNTPFANSDTLDLSR
103 NP_569153.1     ----------------------------------VCAYASGKFNPPKSTI
```

```
104 NP_043924.1          ------------------------------YHPHPLETAKEEEVRIFLRA
105 YP_009243641.1       ------------------------------------------QVLFTLNF
106 YP_004300274.1       ----------------------------------HCRYYSGTFKDNEKNW
107 NP_056848.1          ----------------------------------LCRYNSGTFKENEQKY
108 YP_006495799.1       ----------------------------------VCAYASGKFPAIKSTI
109 YP_009508409.1       ----------------------------------V---------------
110 NP_687035.1          ------------------------------------------MATHIHNF
111 NP_045937.2          ---------------------------------RPMKTTYLSDMSPLWAT
112 NP_612577.1          --------------------------------EYICRYASGSFKAAERNY
113 NP_619548.1          --------------------------------ELICRYSSGSFKQAEKNY
114 NP_056880.1          ------------------------------------------HALFVLNH
115 YP_009121747.1       ----------------------------------ICAYASGKFKVVQSTI
116 YP_009113237.2       ----------------------------------VCAFASGTFSPIKSTI
117 YP_009508408.1       ----------------------------------ICAYASGKFDPPKSTI
118 NP_955579.1          --------------------------------------------------
119 YP_009229919.1       ----------------------------------PCAYASGKFPAVKSTI
120 YP_004442833.1       ----------------------------------ICRYASGKFAKPKSTI
121 NP_862833.2          ------------------------------------------HALYVLNF
122 NP_040563.1          ------------------------------------------LALVSLN-
123 YP_443922.1          --------------------------------GGHMPALDVHQIGMTDYV
124 NP_057933.2          --------------------------------DPDMTRVTNSPS-LQAHL
125 YP_009116631.1       ----------------------------------ICAYASGKFPTVKSTI
126 YP_009513249.1       ---------------------------------IDCNLPFANKDTLDWTR
127 YP_223871.1          --------------------------------GCDKLASITNQT-LLKSL
128 NP_127504.1          ----------------------------------ICGFASGKFKVSEQHY
129 NP_044929.1          ---------------------------------ADCNLPFENLDTLDLTR
130 NP_057860.1          ------------------------------------------IALWTINH
131 NP_040973.1          ------------------------------------------LAVHSLNF
132 NP_777317.1          ----------------------------------VCAYASGKFNPIKSTI
133 NP_054716.1          ---------------------------------VDCNLPFQDLSTLDLTR
134 YP_001856242.1       --------------------------------------------------
135 NP_954565.2          ------------------------------------------HALFILNF
136 NP_569150.1          ----------------------------------ICAYASGKFTPIKSTI
137 NP_659397.1          --------------------------------ELICRYSSGTFKPAELNY
138 YP_009513211.1       --------------------------------EVVGSNGDFFPV-LFTHL
139 YP_233107.1          ----------------------------------ICAYASGRFNPIKGAI
140 YP_009109692.1       --------------------------------N-SFSPSNSKTD-LQARL
141 NP_057849.4          ------------------------------------------MAVFIHNF
142 NP_049560.1          ------------------------------------------IALWTINH
143 YP_009506267.1       ----------------------------------ICAYASGKFDIPKSTI
144 NP_039820.1          ----------------------------------ICAYASGSFNPIKSTI
145 NP_056790.1          --------------------------------ETLGPDDRFLPV-LFTHL
146 YP_002321513.1       ----------------------------------VCDYASGKFPVVKSSI
147 NP_955591.1          --------------------------------------------------
148 YP_001956722.2       ---------------------------------IDSNTPFANSDTLDLSR
149 NP_056902.1          ------------------------------------------LALITCNK
150 NP_056886.1          ------------------------------------------KAMYALNH
151 NP_041261.1          ------------------------------------------HALFQLNF
```

```
001 YP_004442836.1       D---AEIQAVINSLDKFKIYYLDKKELIIRTDSQAIVSFYKKSSDHKPSR
002 NP_056728.1          HSNDKETLAVINTIKKFSIYLTPVHFLIRTDNTHFKS-FVNLNYKGDSKL
003 YP_009508537.1       EEELALLQEIRESLQHPVRPP---TCSGWSPYVGQLVQERVYRPSQLRPK
004 YP_004732983.2       D---AEIYAVMASLENFKIYYLDKREITIRTDCQAIISFYDKTAIKKPSR
005 YP_009508407.1       --------------------------------------------------
006 NP_040939.1          KRKGGLG-TSPMDIFIYNKEQKRINNKYNKNSQKIQFCYYRIRKRGHQES
007 YP_003864102.1       LNLDD-QGKSAADRFWHS--------------DPKKQLAMVKWKDPLDNS
008 YP_009002585.1       D---AEIHAVMNSLEAFKIHYLDKPSLTIRTDCQAIIAFFDTISNHKPSR
009 YP_003284237.1       D---AEIQAVINSLDKFKIYYLDKKKIIIRTDCQAIVAFYSKIAQNKPSR
010 YP_009268869.1       KRKGGLG-TSPMDIFIFNKEQQRIQQQSTRNQSKFRFCYYRVRKRGHPGE
011 YP_009506251.1       D---AEIYALIKALDAFKIFYLDKGHLIVRTDCQAIVTFYNKTNTHKPSR
012 NP_955611.1          --------------------------------------------------
013 YP_008567619.1       D---AEIHAVMNSLEKLKIYYLDKREIIIRTDCQAIISFFNKSSVNKPSR
014 NP_149413.1          D---AEIYGVMNGLEKMRLFYLDKREITVRTDSAAIERFYNKSVEHKPSE
015 YP_004442824.1       D---AEVQAVINSLEKFKIYYLDKKELIIRTDSQAIVTFYKKISDHKPSR
016 YP_001931967.1       HSNEKEILAVIRSIQSFPVYLTPVEFIVRTDNKTMEH-FLTSKFELGTKS
017 YP_002455786.1       LNVMRPCGKTRWQLHHTPPLPPISESIQT--TPTRLHWYYYKTPGLTNQR
018 YP_009109694.1       --------------------------------------------------
019 NP_789739.1          --------------------------------------------------
020 NP_955619.1          LNVLTNCHKTRWQLHHSPRLQPIPETHSL--SNKQTHWYYFKLPGLNSRQ
021 YP_224289.1          D---AEVQAVINSLDKFKIYYLDKKELLIRTDCEAIVRFYKSTAQNKPSR
022 YP_009182100.1       STQEKEVLAIIRTIQKAKIFVLKP-FLVRTDSKFAAL-FLDKKINESLAR
```

023 NP_705927.1 --------------------------------------------------
024 YP_009508577.1 EEELSLLSEVRTSLLPPSTP--PASKRAWLPSVGLLVQERVARPSQLRPK
025 YP_004442830.1 E---AEITGVLQSLEKFRLYYIDKAEITVRTDSSAIERFYNKSTEHKPSE
026 NP_758887.1 KRKGGIGEYSPGERIVDIIATD-LLTTKLQHNIQKIQNFRVYYREGRDQQ
027 YP_009508406.1 --------------------------------------------------
028 YP_610965.1 D---AEIYACTETLSALKIHYLDRKEIILRTDCQAIISFYNKSSVNKPSR
029 YP_001931961.1 HSNEKEILAVIRVIAKFTIYLTPLEFLIRTDNKNFTF-FMNTNVKGDYKQ
030 YP_241114.1 LTLDA-EGNSAAQRFWGE--------------RSSCKKPLVRWKDPLTNL
031 YP_006732334.1 HSNQQ----------------KPITIVMKKNY------------------
032 YP_009041481.1 D---AEIFAVMESLSALKIFFLDKEEITVRTDCQAIIAFYNKQAQNKPSR
033 YP_001036293.1 D---AELFAVMNSMDKFKIHYLDRKELLIRSDCQALISFYNKSSENKPSR
034 YP_605811.1 D---AEVHAVINSLEKFKIYYLDKKELIIRTDSQAIVAFYKKQADHKPSR
035 NP_955577.1 RALQLVQEEIQRPLAAAYREKLETPVVPHPFKPGDSVWVRRHQTKNLEPR
036 YP_003208050.1 D---AEIHACMGTLSALKIHYLDRKEITLRTDCQAIVSFHNKMAQNKPSR
037 NP_789740.1 KRKGGIGGYSAGERIVDIIATD-IQTKELQKQITKIQNFRVYYRDSRNPL
038 YP_004442839.1 D---AEVHAVINSLEKFKIYYLDKKELIIRTDSQAIVAFYKKQADHKPSR
039 YP_009507791.1 KRKGGLGGITPYELYVQ-QESIRIQELYSSIPHKFLKDSWIYYKDRKDKL
040 YP_004581513.1 D---AEIFAIMNAMEAFKIYYLDKKEMIIRTDCEAIVSFFNKSASNKPSR
041 NP_043933.1 HSNEKEILSIIKAIKAFRAYILPYKFLVRTDNTNAAY-FVRTNIAGDYKQ
042 YP_009507248.1 RQQMKQVYEIINETSLANRTALIESRNKTRESEVEYIPQQQVFIKNPLAS
043 YP_002117531.1 D---AEIQAVICSLDKFKIFYLDKQEVVIRTDCQAIISFFDKTSQNKPSR
044 YP_004347415.1 DINRKELIAIYKGLEHYAIFTTQGKKLVRTDNSQAYYWIKNSKIKNSIDM
045 NP_056891.1 LNLDD-QNKSAADRFWHN--------------NPKKQFAMVKWKDPLDNT
046 YP_009513242.1 EEELSLLQEIRSSLHQPTSP--PASSRSWSPSVGQLVQERVARPASLRPR
047 YP_009352866.1 D---AEIFAVMESLSALKIYFLDKEEITVRTDCQAIISFYNKQAQNKPSR
048 YP_009165750.1 HSNEKEILAAMNTIKKFSGYLTPVKFLIRTDNKNFTF-FLNTNHKGDYKQ
049 NP_040840.1 KRKGGLG-TSPMDIFIFNKEQQRIQQQSKSKQEKIRFCYYRTRKRGHPGE
050 YP_004222728.1 FERGE-NTKTPIQKHWRP--------------TILTEGPPVKIRIETG-E
051 YP_009506264.1 D---AEIQAVINSLDKFKIYYLDKKALTIRTDCQAIVSFYAKIAQNKPSR
052 YP_567050.1 LNVMSSQWKTRWQLHHGPRLRPIPETPRP--SRAPTNWYYYKLPGLTNQR
053 YP_009508571.1 EEELSLLQEIRSSLHQPSTP--PASTRSWSPIGGQIVQERVPRPASLRPR
054 NP_056803.1 EEELSLLQEIRASLYQPSTP--PASSRSWSPVVGQLVQERVARPASLRPR
055 NP_955564.1 VNMDN-QGHTAAERHWGP--------------ISADPKPMVMWKDLLTGS
056 YP_009508411.1 D---AEIFAVMNSLEKFKIHYLDKEHIVIRTDCQAIISFYNKSVDNKPSR
057 YP_009140788.1 D---AEIHAVMNSLEKLKIYYLDKRELIIRTDCQAIISFFNKSAMNKPSR
058 YP_009508410.1 --------------------------------------------------
059 NP_040333.1 QALYLVQHEVWRPLAAAYQEQLDRPVVPHPFRVGDTVWVRRHQTKNLEPR
060 YP_008992013.1 HVIYKEILTVKYGIKKFEFHLISHTFLINMDNSSFPRIFDFKNKLLLDKQ
061 NP_041734.1 D---AEIYALIKALESFKIFYLDKKHLVVRTDCQAIVTFYNKTSTHKPSR
062 NP_056895.1 INLLP-ILKTRWELHHSPPLAVISEGGET--PKGSDKLFLYKLPGQNNRR
063 YP_009109689.1 KGLQAVQAQIWAPLAELYRP--GHSQTSHPFQVGDSVYVRRHRSQGLEPR
064 YP_002519387.1 HSNEKEVLAVKRTITKFKGYLASSEFLVRTDNKYFTY-FLRTSIKDDYKQ
065 YP_595725.1 D---AEIYGVMNGLEKMRLFYLDKREITVRTDSSAIERFYNKSAEHKPSE
066 YP_009508551.1 EEELSLLQEIRTSLYHPSTP--PASSRSWSPVVGQLVQERVARPASLRPR
067 YP_001497148.1 KGLQAVQAQIWTPLAELYRP--GHPQTSHPFQVGDSVYVRRHRSQGLEPR
068 NP_041186.1 LTLDT-EGNSAAQRFWGE--------------RSSCKKPLVRWKDPLTNL
069 NP_663784.1 KRRGGIGDMTPAERLINMITTE-QEIQFLQRKNSNFKNFQVYYREGRDQL
070 YP_009508443.1 D---AEIHAVMNSLDKFKIHYLDKEEIIIRTDCQAIISFYNKSAENKPSR
071 YP_003987465.1 D---AEIQAVINSLDKFKIYYLNQKELVVRTDCQAIVSFYEKMANNKPSR
072 NP_861410.1 TVHEKETLAALKTMRKWKAELLPKEFTLRTDSSYVTG-FARHNLKANYNQ
073 YP_009345071.1 D---AEISAVLHALETFKIFYLSQPTIIIRTDCQAIISFYNKSFNHKPSR
074 YP_009508582.1 EEELSLLSEVRTSLLPPSTP--PASRRSWLPSVGLLVQERVARPSQLRPK
075 NP_569141.1 EINRKELFALYKCLLAFEPYIVYTRFIVRTDNTQVKWWITR-KVQDSVTT
076 YP_002916057.1 D---AEIFAIMNAMEAFKIYYLDKKEMVIRTDCQAIVSFFNKSASNKPSR
077 NP_777384.2 INLLP-ILKTRWELHHSPPLAVISEGGET--PKGSDKLFLYKLPGQNNRR
078 YP_233110.1 D---AEVHAVINSLEKFKIYYLDKKELIIRTDSQAIVAFYKKQADHKPSR
079 NP_542258.1 LNVLPSCQKTRWQLHQAQPLPSISENTLP--PRASPKWYYYKIPGLTNPR
080 YP_009130664.1 D---AEIHAVMNSLNSFKIYYLDKKELLIRTDCQAIISFFNKSAKNKPSR
081 YP_009508566.1 EEELALLSEIRTTLSTVSPEPFPSTAKTWTPSVGLLVQERVYRPSQLRPK
082 YP_006607892.1 HSNEKEILAVMNTIKKFTGYLTPVKFLIRTDNKNFTH-FLKINLKGDYKQ
083 YP_009506270.1 D---AEIQAVINSLDKFKIYYLDKKELVIRTDSQAIVSFYKKSSDHKPSR
084 NP_042513.1 TVHEKELLAFVNALKKFQIDLRPVKFIFQTDNSWVQG-FLVNKLKETYNR
085 YP_006907834.1 HSNDKETLAVIRTIQKFSIYLTPVRFLVRTDNTHFKY-FLNINYKGDSKM
086 YP_004442827.1 E---AEIFGVLNSLEKFRLYYIDKAEITVRTDSSAIEKFYNKSTEHKPSE
087 YP_009408594.1 D---AEIFACMETLDALKIHYLDKEEITLRTDCQAIISFYNKSATNKPSR
088 YP_009508561.1 EEELSLLAELRSSFLPSASP--PASSRSWRPSVGLLVQERVARPSQLRPK
089 NP_056907.1 LNVMSSQWKTRWQLHHGPRLRPIPETPRP--SRAPTNWYYYKLPGLTNQR
090 YP_002308474.1 LNVLSSCQKTRWQLHQAQPLPPVPENLPL--PEPVPKWYYYKIPGLTSSR
091 YP_007761644.1 STFGKELRAIKLALQKFKLFLFEP-FTLYSDNLAVIN-FLKKDLNEKRSQ
092 NP_040550.1 FERGE-NTKTPIQKHWRP--------------TVLTEGPPVKIRIETG-E
093 NP_758808.1 D---AEITAAMEGLEAFKIHYLDKQKITLRTDCQAIISFCNKTSVNKPSR
094 YP_009345075.1 D---AEISALINGLESFKIHYLDKGEIMVRTDCHAIVSFYNKLHANKPSR
095 NP_068729.1 PIAELEVLAGVKVLEKWRIDLLQTRFLLRTDSKYFAG-FCRYNIKTDYRN

```
096 NP_047255.1        RALQLVQEEIQRPLAAAYREKLETPVVPHPFKPGDSVWVRRHQTKNLEPR
097 YP_006273075.1     D---AEIHAVMNTLEKLKIYYLDKKELIIRTDCQAIISFFNKSALNKPSR
098 YP_009508888.1     EEELSLLQEIRESLIQPPSP--PASSRSWTPAVGLLVQERVARPASLRPR
099 NP_395469.1        PIHELETLAVLQTFRKWKVDLLSKPFILKTDSKYVTG-FLRYKIKANYNQ
100 YP_009508546.1     EEELSLLQEIRTSLYQPSTP--PASVRSWSPVVGQLVQERVARPASLRPR
101 NP_056762.1        TSLDYEIEAINEALNKFQIY--LDKDFTIRTDCEAIVKGIKTEDYKKRSK
102 YP_009508556.1     EEELSLLQEIRSSLHQPSSP--PASSRSWSPSVGQLVQERVARPASLRPR
103 NP_569153.1        D---AEIHAVMNSLNNFKIYYLDKSSLCLRTDCQAIISFFNKSNVNKPSR
104 NP_043924.1        THDCMQNKKWEEQLEKVTKAEAHSHWTQRSPNLEPGCIVLVRKFTGDAFS
105 YP_009243641.1     MNLPQGDIQTTAERHFEE--------------KSEPLIHEDVWVRPLMET
106 YP_004300274.1     EINRKELLSVFKCLYAFEPYIVYNKFILRTDNTQVKYWLTG-KLDNSVTT
107 NP_056848.1        DINRKELIAVYQGLQSYSLFTCEGNKLVRTDNSQVYYWIKNDTNKKSIEF
108 YP_006495799.1     D---AEMHGVMNSLEKFQIYFMDKGEVTIRTDCQAIVAFYEKLNANKPSR
109 YP_009508409.1     --------------------------------------------------
110 NP_687035.1        KRKGGIGGISSAERLVNMLTTQ-LELNTLQNQIQKILNFKVYYREGRDPV
111 NP_045937.2        DTLVTYMNKLTRQLSAYHQQVVDQWPSTSLPPGPEPGSWCMLRNPKKSSN
112 NP_612577.1        HSNEKELLAVIRVIKKFSIYLTPSRFLIRTDNKNFTH-FVNINLKGDRKQ
113 NP_619548.1        HSNDKELLAVKQVITKFSAYLTPVRFTVRTDNKNFTY-FLRINLKGDSKQ
114 NP_056880.1        VNMDN-QGHTAAERHWGP--------------ISADPKPMVMWKDLLTGS
115 YP_009121747.1     D---AEINACINSLEKLKIYYLDKKEITLRTDCQAIISFYNKTNSNKASR
116 YP_009113237.2     D---AEIHAVMNSLVKFKLYYIGRQKITIRTDCQAIISFVNKSANSKPSR
117 YP_009508408.1     D---AEIHAVMNSLDKFKIHYLDKREILIRTDCQAIISFYNKSMENKPSR
118 NP_955579.1        --------------------------------------------------
119 YP_009229919.1     D---AEIFAVMEGLKNFKIYYLDRGAVTIRTDCQAIISFYNKSAQNKPSR
120 YP_004442833.1     E---AEISGVLCSLEKFRLYYLDKAEITVRTDSGAIERFYNKSTEHKPSE
121 NP_862833.2        LTLDA-EGNSAAQRFWGE--------------RSSCKKPLVRWKDPLTNL
122 NP_040563.1        KKRGGIGGHTPYEIYLESEHTKYQDQLEQQFSKQKIEKWCYVRNRRKE--
123 YP_443922.1        RKLTELSAALSIQIQRVAEGELTDDQDQPRVKVGDWVRIKVHKRKWADPR
124 NP_057933.2        QALYLVQHEVWRPLAAAYQEQLDRPVVPHPYRVGDTVWVRRHQTKNLEPR
125 YP_009116631.1     D---AEIFAVMESLEKFKIFYMNKDEVTIRTDCQAIITFYEKLNAKKPSR
126 YP_009513249.1     EEELALLQEIRESLQHPVQPP---TCSGWSPYVGQLVQERVYRPSQLRPK
127 YP_223871.1        QALQATRSLARATLRDQLPQKEAQQDRTPLFQPGDLVFVKKHDFQQLGPR
128 NP_127504.1        HSTFKEILAVKNGIKKFNFFLIHTNFLVEMDMRAFPKMIRLNPKIVPNSQ
129 NP_044929.1        EEQLAVLKEVRDGLLDLYPSPSQTTARSWTPSPGLLVQERVARPAQLRPK
130 NP_057860.1        LNVLTNCHKTRWQLHHSPRLQPIPETHSL--SNKQTHWYYFKLPGLNSRQ
131 NP_040973.1        KRRGRIGGMAPYELLAQ-QESLRIQDYFSAIPQK-LQAQWIYYKDQKDKK
132 NP_777317.1        D---AEIQAVINSLDKFKIYYLDKKELVIRTDSQAIVTFYKRMADHKPSR
133 NP_054716.1        EEQLAVLQEIRTALEQPAQHP---TLPKWTPCPGLLVQERVNRPAQLRPK
134 YP_001856242.1     --------------------------------------------------
135 NP_954565.2        LNLDD-QNKSAADRFWHN--------------NPKKQFAMVKWKDPLDNT
136 NP_569150.1        D---AEIQAVINSLDKFKIYYLDKKELIIRTDSQAIVSFYKKSSDHKPSR
137 NP_659397.1        HSNEKELLAVKQVITKFSIYLTPVCFTVRTDNVNLLKGFMNKKITGDSKQ
138 YP_009513211.1     KALEVVRTQIWDQIKEAYRP--GTVAIPHPFQVGDRVLVRRHRSGSLEPR
139 YP_233107.1        D---AEIQAVIYSLEKFKIYYLDKRELILRTDSKAIVRFYEKCSEHKPSR
140 YP_009109692.1     KGLQAVQAQIWAPLAELYRP--GHSQTSHPFQVGDSVYVRRHRSQGLEPR
141 NP_057849.4        KRKGGIGGYSAGERIVDIIATD-IQTKELQKQITKIQNFRVYYRDSRNPL
142 NP_049560.1        LNVLTHCHKTRWQLHHSPRLQPIPETRSL--SNKQTHWYYFKLPGLNSRQ
143 YP_009506267.1     D---AEIHAVMNCMDKFKIHYLDKKELLIRTDCHAIMAFYNKSSCNKPSR
144 NP_039820.1        D---AEIQAAIHGLDKFKIYYLDKKELIIRSDCEAIIKFYNKTNENKPSR
145 NP_056790.1        KALEIVRTQIWDQIKEVYKP--GTVTIPHPFQVGDQVLVRRHRPSSLEPR
146 YP_002321513.1     D---AEIHAAMFTLEKLKIYYLDKEEIILRTDCQAIISYYNKSNSNKPSR
147 NP_955591.1        --------------------------------------------------
148 YP_001956722.2     EEELSLLQEIRSSLYLPSTP--PASIRAWSPSVGQLVQERVARPASLRPR
149 NP_056902.1        GRE-SMGGQTPWEVFITNQAQVIHEKLLLQQAQS--SKKFCFYKIPGEHD
150 NP_056886.1        FERGE-NTKTPIQKHWRP--------------TVLTEGPPVKIRIETG-E
151 NP_041261.1        LQLDS-QGRSAADRLWHP--------------QTSQQHATVMWRDPLTSV
```

```
001 YP_004442836.1     VRWLAFTDYITGTGLDVKFEHIDGKDNVLADTLSRLVK------------
002 NP_056728.1        GRNIRWQAWLSHYS--FDVEHIKGTDNHFADFLSREFNKVNS--------
003 YP_009508537.1     WRKPTKVLEILNPRTVIIVDHLGQRKPVSIDNLKPTAHQHNG--------
004 YP_004732983.2     VRWINFCDYITNTGIKVQFEHIKGQDNQLADQLSR---------------
005 YP_009508407.1     --------------------------------------------------
006 NP_040939.1        GKDQPRYCGKGKEPIVVKDIESEKYLVIPYKDAKFIPPPTKEKE------
007 YP_003864102.1     WNGPDPVLIWGRGSVCVYSQAHDAARWLPERLVRQINNENQ---SRESYS
008 YP_009002585.1     LRWINFTDYVTGLGIPVNFQHVKGEDNRLADALSR---------------
009 YP_003284237.1     VRWLTFSDFISGLGVQVSFEHIDGKDNTIANTLSRLVH------------
010 YP_009268869.1     WLGPTQVLWEGEGAIVIKDKNLEKYLVIAKKDVKFIPQPKEIQTE-----
011 YP_009506251.1     IRWITFLDYITGLGVSVTIEHIDGKDNHLADTLSR---------------
012 NP_955611.1        --------------------------------------------------
013 YP_008567619.1     VRWIGFTDYITGLGIDVKFQHIDGKENSLADSLSR---------------
014 NP_149413.1        IRWIRFMDYITGAGPTIVIEHIKGKHNGLADILSRLKA------------
```

015 YP_004442824.1 VRWLAFVDYITGTGLNIRFEHIDGKDNILADTLSRLVQ------------
016 YP_001931967.1 GRLVRWQMWFKHYN--FKVEHIKGTSNFLADYLSREYNEI----------
017 YP_002455786.1 WKGPVQSLQEAAG-AALLQVSDGSPQWIPWRLLKKTVCPKPDDPEPAGHV
018 YP_009109694.1 --------------------------------------------------
019 NP_789739.1 --------------------------------------------------
020 NP_955619.1 WKGPQEALQEAAG-AALIPVSASSAQWIPWRLLKRAACPRPVG-GPADPK
021 YP_224289.1 VRWLMLTDFISGTGLEIKFEHINGCENILADSLSRLVQ------------
022 YP_009182100.1 GRLIRWQIMLRQYY--LDIEHIPGVSNYLPNALTREMAGCNIIYQIKMVN
023 NP_705927.1 --------------------------------------------------
024 YP_009508577.1 WKKPTPILEVVNDRTVVILDNQGQRRTVSIDNLKLTPHQD----------
025 YP_004442830.1 VRWIHFMDVISGAGPEIKFEHIKGKENTLADLLSRLN-------------
026 NP_758887.1 WKGPAELIWKGEGAVVIKEGTDLKVVPRRKAKIIRDYG--KTVDSDPNVE
027 YP_009508406.1 --------------------------------------------------
028 YP_610965.1 VRWLAFTDFITGLGIDVQFEHIKGEQNVLADTLSRLTFVLL---------
029 YP_001931961.1 GRLVRWQQWLSRYS--FKVEHITGVKNIFADFLTREFQSKNSIEL-----
030 YP_241114.1 WYGPDPVLIWGRGHVCVFPQDAEAPRWIPERLVRAAEELPDISNATHDIE
031 YP_006732334.1 --------------------------------------------------
032 YP_009041481.1 VRWLAFCDYINGNGLSVKFEHIKGEENVLADQLSR---------------
033 YP_001036293.1 VRWIAFNDFVTGLGIDVTFEHIDGKDNQLADSLSR---------------
034 YP_605811.1 TRWLMLIDYITGLGINVKFEHIDGKENVLADTLSRLVQ------------
035 NP_955577.1 WKGP--HIVLLTTPTALKVDG--VAAWIHASHVKAAGPTTNQDLSDSPSS
036 YP_003208050.1 VRWISFTDFITGLGIKVNIEHIDGKDNVLADTLSRLIFKIQ---------
037 NP_789740.1 WKGPAKLLWKGEGAVVIQDNSDIKVVPRRKAKIIRDYG--KQMAGDDCVA
038 YP_004442839.1 TRWLMLIDYITGLGINVKFEHIDGKENVLADTLSRLVQ------------
039 YP_009507791.1 WKGPTQVYYWGEGAVLIKDEN-NKYLLIPRRRIRRVPAPKDLTQEDGFNQ
040 YP_004581513.1 ARWISFTDYITGTGIKIRIEHIDGKDNILADYLSR---------------
041 NP_043933.1 SRLVRWQMALREYS--FDIEHVSGQKNVLADIMTRELAGKT---------
042 YP_009507248.1 RQKVAPRYTQDTVLADLPIHIYTSKKRGPVAKARLKRVPKGN--------
043 YP_002117531.1 VRWLTFTDFITGIGIPVRFEHISGKDNLLADTLSRLVL------------
044 YP_004347415.1 KNVKGILAKIIMYD--FDIEIIDGKTNIVADFLSRNGTDDATITG-----
045 NP_056891.1 WHGPDPVLIWGRGSVCVYSQTYDAARWLPERLVRQVSNNNQ---SRE---
046 YP_009513242.1 WHKPTSILEVVNPRTVVILDHLGNRRTVSVDNLKLTAHQN----------
047 YP_009352866.1 VRWLAFCDYINGNGLCVKFEHIKGEENFLADHLSR---------------
048 YP_009165750.1 GRLIRWQMWFSRYS--FTVEHLPGNKNVFADFLTREFNN-----------
049 NP_040840.1 WQGPTQVLWGGDGAIVVKDRGTDRYLVIANKDVKFIPPPKEIQKE-----
050 YP_004222728.1 WEKGWNVLVWGRGYAAVKNRDTDKVIWVPSRKVKPDITQKDEVTKKDEAS
051 YP_009506264.1 VRWLTFSDFITGLGVVVTFEHIDGKDNVLADTLSRLVV------------
052 YP_567050.1 WKGPLQSLQEAAG-AALLSID-GSPQWIPWRLLKKTVCPRPDGSELVAHA
053 YP_009508571.1 WHKPSRIVDILNERTVVIVDHLGNNRTVSIDNLKLTPHQ-----------
054 NP_056803.1 WHKPSTVLEVLNPRTVVILDHLGNNRTVSIDNLKPTSHQ-----------
055 NP_955564.1 WKGPDVLITAGRGYACVFPQDAETPIWVPDRFIRPFTERKEATPTPGTAE
056 YP_009508411.1 VRWVAFNDYLTGLGIEVKIEHIQGVHNTLADALSR---------------
057 YP_009140788.1 VRWISFTDFITGLGINVEFEHIDGKDNVLADSLSR---------------
058 YP_009508410.1 --------------------------------------------------
059 NP_040333.1 WKGP--YTVLLTTPTALKVDG--IAAWIHAAHVKAADTRI--------EP
060 YP_008992013.1 LLNLKTWFAK----YDFTVQHIKGNQNLIPDFLTRPAINKPSLISSIQTI
061 NP_041734.1 IRWITFSDYITGLGVPVTIEHIDGKENQLADTLSR---------------
062 NP_056895.1 WLGPLPALVEASG-GALLATN--PPVWVPWRLLKAFKCPKNDGPEDAHNR
063 YP_009109689.1 WKGP--YIVLLTTPTAIKVDG--IATWIHASHAKAAPGTP--------GP
064 YP_002519387.1 GRLIRWQQWFSHYK--FNVEHLAGTHNFTADSLTREFANKPP--------
065 YP_595725.1 IRWIRFMDYITGAGPEIVIEHIKGKSNGLADILSRLKA------------
066 YP_009508551.1 WHKPSTVLKVLNPRTVVILDHLGNNRTVSIDNLKPTSHQ-----------
067 YP_001497148.1 WKGP--YIVLLTTPTAIKVDG--IAAWIHASHAKAAPKTP--------GP
068 NP_041186.1 WYGPDPVLIWGRGHVCVFPQDAEAPRWIPERLVRAAEELPDASDATHDPE
069 NP_663784.1 WKGPGELLWKGEGAVIVKVGTDIKVVPRRKAKIIRDYGGRQELDSSPHLE
070 YP_009508443.1 VRWIAFNDFITGLGISVHFEHIDGKHNQLADALSR---------------
071 YP_003987465.1 VRWLTFSDFITGIGVPVKFEHIDGEDNLLADTLSRLVV------------
072 NP_861410.1 GRLVRWQLEFLQYP--ARVEYIKGEKNSLADTLTREWKQQ----------
073 YP_009345071.1 LRWIRLSDWLTGTGVAYKFEHIKGELNNLADHLSRNPEIALT--------
074 YP_009508582.1 WRKPTPILEVVNDRTVVILDNQGQRRTVSIDNLKLTPHQD----------
075 NP_569141.1 KEIRRLVLNILNFT--FTIEIINTNKNVVADYLSRQSYPN----------
076 YP_002916057.1 ARWISFTDYITGTGIKIRIEHIDGKDNTLADYLSR---------------
077 NP_777384.2 WLGPLPALVEASG-GALLATN--PPVWVPWRLLKAFKCPKNDGPEDAHNR
078 YP_233110.1 TRWLMLIDYITGLGINVKFEHIDGKENVLADTLSRLVQ------------
079 NP_542258.1 WSGPVQSLKEAAG-AALIPVG-GSHLWIPWRLLKRGICPRPESNAVADPE
080 YP_009130664.1 VRWVSFTDFITGTGLEIRFQHIDEKDNALADALSR---------------
081 YP_009508566.1 WKKPTPILEVLNERTVVIDNN-GQRRTVSVDNLKYTPHQKD---------
082 YP_006607892.1 GRLVRWQMWFSRYV--FEVEHLSGDKNVFADFLTREFHY-----------
083 YP_009506270.1 VRWLAFTDYITGTGLEVKFEHIDGKDNVLADTLSRLVK------------
084 NP_042513.1 GRLARWSMFIQQFD--FISLHIAGKENYLADTLTREWKTSSR--------
085 YP_006907834.1 GRNIRWQGWLQNYV--FDVDHIKGTNNCLADFLSREFNGA----------
086 YP_004442827.1 IRWIKFMDFISGAGPEIKFEHIKGKDNTLADLLSRLN-------------
087 YP_009408594.1 VRWLNFADFITGTGVKIVFEHIDGRQNVLADSLSRLITYMS---------

```
088 YP_009508561.1    WKKPVIICELINDRTVVIVDKAGNKRTVSIDNLKLTPHQKNS--------
089 NP_056907.1       WKGPLQSLQEAAG-AALLSID-GSPQWIPWRLLKKTVCPRPDGSELVAHA
090 YP_002308474.1    WSGPVQSVKEAAG-AALIPVG-TRHIWIPWRLLKRGACPRPGDSVTTESK
091 YP_007761644.1    -REIRDKLDILQYQGWMTLKHIPGTKNVLADALTRGLSN-----------
092 NP_040550.1       WEKGWNVLVWGRGYAAVKNRDTDKVIWVPSRKVKPDVTQKDEVTKKDEAS
093 NP_758808.1       VRWLKFIDYITNTGIDVKFEHIDAKNNVLADTLSRLVN-TLQDLP--WLD
094 YP_009345075.1    LRWAKLLDYLTGTGVQITLEHIDGRDNILADQLSRLSSYIS---------
095 NP_068729.1       GRLIRWQLRLQAYQ--PYVELIKSENNPFADTLTREWSKPSSS-------
096 NP_047255.1       WKGP--HIVLLTTPTALKVDG--VAAWIHASHVKAAGPTTNQDLSDSPSS
097 YP_006273075.1    VRWVAFTDFITGIGINVEFEHIDGRDNVLADSLSR---------------
098 YP_009508888.1    WHKPVKILEVLNPRTVVILDHLGNNRTVSVDNLKLTANQ-----------
099 NP_395469.1       GRFIRWQLELSQFN--YRTFYIKGSENYGPDTLTREWKEL----------
100 YP_009508546.1    WHKPSTVLEVLNPRTVVILDHLGNNRTVSIDNLKPTSHQ-----------
101 NP_056762.1       TRWIKLRDNLLKDGYKPTFEHIKGNKNFLPNFLSREGDFILKCLQNPDST
102 YP_009508556.1    WHRPTTVLEVVNPRTVIILDHLGNRRTVSVDNLKLTAHQD----------
103 NP_569153.1       VRWIAFTDFLTGLGIPVNIEHIDGKNNHLADALSR---------------
104 NP_043924.1       PKWEGPYVITETTKYAAKVQAMSDKVTQHSGWIHRTHLVLFPSQNKRWAD
105 YP_009243641.1    EWVPAQLLVKGKGYGLVLLQNGK-QQWLPACRIQVRRTTGQETIATASAV
106 YP_004300274.1    KEIRRLVVKINCYN--FDVVVIKSKDNCFADYLSREVKK-----------
107 NP_056848.1       RNIKYLLAKIAVYN--FEIQLIDGKTNIIADYLSRYNSSDTDGRYDEANT
108 YP_006495799.1    VRWLNFCDYITNTGVKVVFEHIKGKDNVLADTLSR---------------
109 YP_009508409.1    --------------------------------------------------
110 NP_687035.1       WKGPARLIWKGEGAVVIKEGEDIKVVPRRKAKIIKDYGERKTMDSEGSME
111 NP_045937.2       WEGPFLILLSTPTAVKVEGRPTWIHLDHCKLLRSSLSSSLG---------
112 NP_612577.1       GRLVRWQMWLSQYD--FDVEHIAGTKNVFADFLQENTLTNYV--------
113 NP_619548.1       GRLVRWQNWFSKYQ--FDVEHLEGVKNVLADCLTRDFNA-----------
114 NP_056880.1       WKGPDVLITAGRGYACVFPQDAETPIWVPDRFIRPFTERKEATPTPGTAE
115 YP_009121747.1    VRWLKFSDIITGTGVTINMEHIDGKNNVLADSLSRMVKLCFTGCT--EQE
116 YP_009113237.2    VRWISFTDFITGCGSEVRFEHIEGKENQLADHLSR---------------
117 YP_009508408.1    IRWIAFSDFISGLGISVHFEHIEGRHNQLADALSR---------------
118 NP_955579.1       --------------------------------------------------
119 YP_009229919.1    VRWLGFLDFITGSGVDVTFEHVEGKNNMLADALSRLTSNLCYAECPENQK
120 YP_004442833.1    VRWLKFMDFISGAGPEIKFEHIKGKDNTLADLLSRLN-------------
121 NP_862833.2       WYGPDPVLIWGRGHVCVFPQDAEAPRWIPERLVRAAEEFPDTSNASNDTE
122 NP_040563.1       WKGPYKVLWDGDGAAVIEEEGKTALYPHRHMRFIPPPDSDIQDGSS----
123 YP_443922.1       WTGP--YEVKEVTSHSVQVKGKSGAPWHHLTHCTPAPAPDR---------
124 NP_057933.2       WKGP--YTVLLTTPTALKVDG--IAAWIHAAHVKAADPGG--------GP
125 YP_009116631.1    VRWLAFCDYITNSGVRMKFEHIKGKDNQLADNLSR---------------
126 YP_009513249.1    WRKPTKVLEILNPRTVIIVDHLGQRKSVSIDNLKPTAHQHNG--------
127 YP_223871.1       WDGP--YTVVLSTPTAVKVAG--KTPWIHYSRLKKAPDNQ----------
128 NP_127504.1       LLRWAQWFSP----YQFEVKHLKGKDNILADFLSRPHEFSQRLKNSPKVL
129 NP_044929.1       WRKPTPIKKVLNERTVIIDHLG-QDKVVSIDNLKPAAHQKLA--------
130 NP_057860.1       WKGPQEALQEAAG-AALIPVSASSAQWIPWRLLKRAACPRPVG-GPADPK
131 NP_040973.1       WKGPMRVEYWGQGSVLLKDEE-KGYFLIPRRHIRRVPEPCALPEGDE---
132 NP_777317.1       VRWLAFTDYITGTGLDIKFEHIDGKENILADTLSRLVN------------
133 NP_054716.1       WKKPTPILKVLNPKTVVIAGPGGQERIVSIDNLKKTPHHDTS--------
134 YP_001856242.1    --------------------------------------------------
135 NP_954565.2       WHGPDPVLIWGRGSVCVYSQTYDAARWLPERLVRQVSNNNQ---SRE---
136 NP_569150.1       VRWLAFTDYITGTGLEIKFEHIDGKDNVLADTLSRLVK------------
137 NP_659397.1       GRLIRWQMWFSHYT--FKVDHLKGEQNVLADYLTREFHNGNVIHDD----
138 YP_009513211.1    WKGP--YLVLLTTPTAVKVDG--IAAWVHASHLKPAPPGAP---------
139 YP_233107.1       VRWMTLTDYISGCGVKVYFEHIDGKDNTLADELSRLVQA-----------
140 YP_009109692.1    WKGP--YIVLLTTPTAIKVDG--IATWIHASHAKAAPGTP--------GP
141 NP_057849.4       WKGPAKLLWKGEGAVVIQDNSDIKVVPRRKAKIIRDYG--KQMAGDDCVA
142 NP_049560.1       WKGPQEALQEAAG-PALIPVIASSAQWIPWRLLKRAACPRPVG-APPIPK
143 YP_009506267.1    VRWIAFNDFLTGLGIPVTIEHIDGKNNVLADALSR---------------
144 NP_039820.1       VRWLTFSDFLTGLGITVTFEHIDGKHNGLADALSRMIN------------
145 NP_056790.1       WKGP--YLVLLTTPTAVKVDG--IAAWVHASHLKPAPPSAP---------
146 YP_002321513.1    ARWIAFTDYITNTGTNIKFEHIEGKLNVLADALSR---------------
147 NP_955591.1       --------------------------------------------------
148 YP_001956722.2    WHKPTPVLEVINPRAVVILDHLGNRRTVSVDNLKLTAYQK----------
149 NP_056902.1       WKGPTRVLWKGDGAVVVNDEG-KGIIAVPLTRTKLLIKPN----------
150 NP_056886.1       WEKGWNVLVWGRGYAAVKNRDTDKVIWVPSRKVKPDITQKDEVTKKDEAS
151 NP_041261.1       WKGPDPVLIWGRGSACIYDQKEDGPRWLPERLIRHINNQTAPLCDRPSNP
```

```
001 YP_004442836.1    ---IICHKEKHPSETILINVAEEILQKGSIGAKRKLGEMISGYEAWMTR-
002 NP_056728.1       --------------------------------------------------
003 YP_009508537.1    --TRTCDDPEGMDGMECSQTTT----------------------------
004 YP_004732983.2    -----LAQNVCAIQVIPESA--------HEALSIILEQD-----------
005 YP_009508407.1    --------------------------------------------------
006 NP_040939.1       --------------------------------------------------
```

```
007 YP_003864102.1    P-------------------------------------------------
008 YP_009002585.1    -----LISVIFEAGQQWQE---------ESTRMQHLTQLIGE--------
009 YP_003284237.1    ---ILIKEEDQELINKIMVLWEKKTE------------------------
010 YP_009268869.1    --------------------------------------------------
011 YP_009506251.1    -----LVWSQFP------------------IEPPEEEPLKSQ--------
012 NP_955611.1       --------------------------------------------------
013 YP_008567619.1    -----LTCSLIR-QWHHLEP---------VITTMEAALVQEQ--------
014 NP_149413.1       ---KLAEA-PSEEVVLLAKALKEIAYYPDHPQVPKLIEWGKQILDPFPKF
015 YP_004442824.1    ---IIIHKEAHPAMSILIDAAEEVVRKYNPHSAARLGRMIEAVKDWQDNQ
016 YP_001931967.1    --------------------------------------------------
017 YP_002455786.1    ETDHQHHG------------------------------------------
018 YP_009109694.1    --------------------------------------------------
019 NP_789739.1       --------------------------------------------------
020 NP_955619.1       EKDHQHHG------------------------------------------
021 YP_224289.1       ---TLLQGWQHQHLNGILLALEELYQKPNPEVAKKIGQIIMKVLE-----
022 YP_009182100.1    HGKAPRMDLRDVITSKARARENQQLENEDIHWKQHLASTSRSFAKRKDVP
023 NP_705927.1       --------------------------------------------------
024 YP_009508577.1    ---GTSNEPDGMDPLEQKKKYDNHDI------------------------
025 YP_004442830.1    ---KVLKVEASTEMITLARALREIDYDQDHPAFKKIHEYSQKVKWP----
026 NP_758887.1       A-------------------------------------------------
027 YP_009508406.1    --------------------------------------------------
028 YP_610965.1       -QVPDPFSPPITELCQALKEIQATSTTPVGVWN---RFQTQICKT-----
029 YP_001931961.1    --------------------------------------------------
030 YP_241114.1       --------------------------------------------------
031 YP_006732334.1    --------------------------------------------------
032 YP_009041481.1    -----ITALLFVQEWPTQEE--------VEELGTFLTAIDDT--------
033 YP_001036293.1    -----LVTSLVRHEGKWQQQ---------RKALVIIEEMMQK--------
034 YP_605811.1       ---VLITRVHHPAETQLVEAVMEVLSNPKKEALDKVNHFIFLTQQWIAEQ
035 NP_955577.1       DDPSRWKVQRTQNPLKIRLSRGT---------------------------
036 YP_003208050.1    -EEGALMALERRPEWEPRPPPRVDNNSPMNIIYGLIHQDQQELIN-----
037 NP_789740.1       SRQDED--------------------------------------------
038 YP_004442839.1    ---VLVTKIHHPAEEQLVDAVLEVLKNPKEVAIKKVNHFILLTERWIAGF
039 YP_009507791.1    TEQESNNA------------------------------------------
040 YP_004581513.1    -----LVFSLIIAEWKTQ-EKSIAPLQAPRITLTKXSCSKQQEPLL----
041 NP_043933.1       --------------------------------------------------
042 YP_009507248.1    ---TLLQDSAATDNTCDASSRDKT--------------------------
043 YP_002117531.1    ---CMVQGWTHDGIKAITPAITAIKDDSDSARVAAIIMKVLP--------
044 YP_004347415.1    --------------------------------------------------
045 NP_056891.1       --------------------------------------------------
046 YP_009513242.1    --NGTTNDSGTMAPVEEDESGSPSS-------------------------
047 YP_009352866.1    -----ITALLVTGPWPTEEE--------AEELGTFLAVADDT--------
048 YP_009165750.1    --------------------------------------------------
049 NP_040840.1       --------------------------------------------------
050 YP_004222728.1    PLFAGISDWAPWKSEQEGL-------------------------------
051 YP_009506264.1    ---MFLQED------KYDDLIQKALK------------------------
052 YP_567050.1       ATDHQHHG------------------------------------------
053 YP_009508571.1    --DGTSNVSSTMDHLE----------------------------------
054 NP_056803.1       --NGTTNDTATMDHLEQNEQSS----------------------------
055 NP_955564.1       KTPPRDEKDQQESPKNESSPHQREDGLATSAGVDLRSGGGP---------
056 YP_009508411.1    -----LVGSLVFHTEAWNSR---------KEALMVAEACINK--------
057 YP_009140788.1    -----LINSFFLTEWN--------------QLKEQSMMELEE--------
058 YP_009508410.1    --------------------------------------------------
059 NP_040333.1       PSESTWRVQRSQNPLKIRLTRGTS--------------------------
060 YP_008992013.1    PVIALNRQLPFKALTQRHFPMNISFQSAYQLQ------------------
061 NP_041734.1       -----LVYTTWNQS--------------QTHQPEEEELEKSQ--------
062 NP_056895.1       SSDG----------------------------------------------
063 YP_009109689.1    TSSGTWRLRRSEDPLKIRLSRT----------------------------
064 YP_002519387.1    --------------------------------------------------
065 YP_595725.1       ---KLAQNEPTEEMILLTQAIREVIPYPDHPYTEQLREWGNKILDPFPTF
066 YP_009508551.1    --NGTTNDTATMDHLEQNE-------------------------------
067 YP_001497148.1    ETPKTWKLRRSENPLKIRLSRV----------------------------
068 NP_041186.1       --------------------------------------------------
069 NP_663784.1       GAREDGEMACPCQVPEIQNKRPRGGALCSPPQGGMGMVDLQQGNIPTTRK
070 YP_009508443.1    -----LVGALIQNSEKWSQR---------THIIRAMGLAIEQ--------
071 YP_003987465.1    ---MMLHEEAYTEPLQRILPLLSKKE------------------------
072 NP_861410.1       --------------------------------------------------
073 YP_009345071.1    ----KLVSTLISEWQDLPEP--------IYHLLSETLSMGEE--------
074 YP_009508582.1    ---GTSNEPNGVDPLEQEKEHDNHDI------------------------
075 NP_569141.1       --------------------------------------------------
076 YP_002916057.1    -----LVFSLIIAEWKTQGRRSTAHHQAQSMAQTIGSCSKQQEPLL----
077 NP_777384.2       SSDG----------------------------------------------
078 YP_233110.1       ---VLITKVHHPAETQLVEAVMEVISNPKKEALDKVNHFIFLTQQWIAER
079 NP_542258.1       TKDHQLHG------------------------------------------
```

```
080 YP_009130664.1    -----LVHGLVSKPDFHSDQ---------TILLVELAIAEAQ--------
081 YP_009508566.1    ---GETYDSS----------------------------------------
082 YP_006607892.1    --------------------------------------------------
083 YP_009506270.1    ---IIFHQEKHQSEEILINAVEKTLKKGDASTRQKINDLVKRYESWMNTG
084 NP_042513.1       --------------------------------------------------
085 YP_006907834.1    --------------------------------------------------
086 YP_004442827.1    ---RALKAEATIELVTLAQALKEIDYDQDHIAFEKIKRYAENIKWP----
087 YP_009408594.1    --------TEWQTLEEADQAAQAQLSIQTNLLL-----------------
088 YP_009508561.1    --HGSTPDITGMDVMEQEEEPGNMD-------------------------
089 NP_056907.1       ATDHQHHG------------------------------------------
090 YP_002308474.1    HKDLQLHG------------------------------------------
091 YP_007761644.1    --------------------------------------------------
092 NP_040550.1       PLFAGISDWIPWEDEQEGLQGETASNKQERPGEDTLAANES---------
093 NP_758808.1       EPHQDQTVSLMQEIEDAPLEIKQR----SLTCLQRLICRSFMEDST--EE
094 YP_009345075.1    -KEGKLVTSQQQQALKVLDQVYQAKDHQRWKDPALIACLHQLFPX-----
095 NP_068729.1       --------------------------------------------------
096 NP_047255.1       DDPSRWKVQRTQNPLKIRLSRGT---------------------------
097 YP_006273075.1    -----LINSFFLTEWS--------------LKKEEAVAELEK--------
098 YP_009508888.1    --NGSTNDSTTMAGLGQNAESQ----------------------------
099 NP_395469.1       --------------------------------------------------
100 YP_009508546.1    --NGTTNDITTMDHLEQNE-------------------------------
101 NP_056762.1       ESYSIDSSESIPLYIDSKESHSIESDDSIPLYRDKLLPLVERLKEKSA--
102 YP_009508556.1    --NGTTNDSGTMAPVEKDESGPSSS-------------------------
103 NP_569153.1       -----LVTGFVFAEPQCQDKFQDDLGKLEAALQEKKEAPQAM--------
104 NP_043924.1       PGNPGNQPDEDCTGKKGTVSTADMSPTTSTTRDRGINMGTEQTHRRRSPR
105 YP_009243641.1    --------------------------------------------------
106 YP_004300274.1    --------------------------------------------------
107 NP_056848.1       --------------------------------------------------
108 YP_006495799.1    -----LTQTLAAVREMPAEQ--------EEILRQALNNTEVQ--------
109 YP_009508409.1    --------------------------------------------------
110 NP_687035.1       GVREANKQMEGDSDLQDQE-------------------------------
111 NP_045937.2       ---GPVNQLLS---------------------------------------
112 NP_612577.1       --------------------------------------------------
113 NP_619548.1       --------------------------------------------------
114 NP_056880.1       KTPPRDEKDQQESPKNESSPHQREDGLATSAGVDLRSGGGP---------
115 YP_009121747.1    TEVLEKGLAAMGEIFEENVKLSNEKNGYEEIIQISSYCQKFIEDNSRLKN
116 YP_009113237.2    -----LINTLILYDEHCTQR--------QELNLLIPAIEENK--------
117 YP_009508408.1    -----LVLSLVQVPDEWHHQ---------QGLTSKIEDALEQ--------
118 NP_955579.1       --------------------------------------------------
119 YP_009229919.1    EELATLAEKAWNELSDGTKAGYNTAKLTMELAHLMMTWMDELSMSSREST
120 YP_004442833.1    ---AALKAEPTQELITLAKALKEIEYDEDNPVFKKIKEYSEKIKWP----
121 NP_862833.2       --------------------------------------------------
122 NP_040563.1       --------------------------------------------------
123 YP_443922.1       -TLTEVRSDLIASNLAINSESPDN--------------------------
124 NP_057933.2       SSRLTWRVQRSQNPLKIRLTREAP--------------------------
125 YP_009116631.1    -----LTQLITFVKWLPTEL--------KDLAAELTRKDDGT--------
126 YP_009513249.1    --TRTCDDPEGMDGMECSQTTT----------------------------
127 YP_223871.1       ---EEWTVSPTSDPLRVKLTRRAKP-------------------------
128 NP_127504.1       MFQRRTRSSSTKSKADSSQSTGSSYKLSHNLPENPPEVFNLDYPWDTSVF
129 NP_044929.1       --QTPDSAEICPSATPCPPNTSLWYDLDTGTWTCQRCGYQCPDKYHQPQC
130 NP_057860.1       EKDHQHHG------------------------------------------
131 NP_040973.1       --------------------------------------------------
132 NP_777317.1       ---FLCFAGTNEEVKELAVQAITQEQK-NPMGLKRLGQILQHKEERVHYL
133 NP_054716.1       --NDSTRMDAVEVPTECQ--------------------------------
134 YP_001856242.1    --------------------------------------------------
135 NP_954565.2       --------------------------------------------------
136 NP_569150.1       ---IILHPEKHQSEGVLINAVEEVFHKGNTDAKQRVNDVVKRYEDWLSKG
137 NP_659397.1       --------------------------------------------------
138 YP_009513211.1    --DESWELEKTDHPLKLRVRRRNESTA-----------------------
139 YP_233107.1       ---ILINKEESPIILSLIKATTEVLQKENPISRSRLALCISRALG-----
140 YP_009109692.1    TSSGTWRLRRSEDPLKIRLSRT----------------------------
141 NP_057849.4       SRQDED--------------------------------------------
142 NP_049560.1       KKTTNTMGKFLTTLILFFQFCPLILGDYSPSCCTLTIGVSSYHSKPCNPA
143 YP_009506267.1    -----LVCSIIKNPVYWQSK---------EELLKQTEEALEQ--------
144 NP_039820.1       ----FIVEKNDESPYRFTSSVEDALKVCN---------------------
145 NP_056790.1       --DESWELEKTDHPLKLRIRRRRDESAK----------------------
146 YP_002321513.1    -----ITCYLCTAGSQGCLAPEE-----DDHMKVMETLPEEIN-------
147 NP_955591.1       --------------------------------------------------
148 YP_001956722.2    --DGTPNESAAVVAMEKDE-------------------------------
149 NP_056902.1       --------------------------------------------------
150 NP_056886.1       PLFAGISDWIPWEDEQEGLQGETASNKQERPGEDTLAANES---------
151 NP_041261.1       NTAPGPKGSP----------------------------------------
```

001 YP_004442836.1 IQEHKIKTLTLIEK------------------------PIFKCGCR----
002 NP_056728.1 --------------------------------------------------
003 YP_009508537.1 ----ETSVDSS---------------------------------------
004 YP_004732983.2 ----CTAQELMAQ-----------------------FNSMLQANLR----
005 YP_009508407.1 --------------------------------------------------
006 NP_040939.1 --------------------------------------------------
007 YP_003864102.1 --------------------------------------------------
008 YP_009002585.1 ---HYKFEASGKK-------------------HHQLKETLTALITS----
009 YP_003284237.1 ------DVMHINDN------------------------PLLVCGCG----
010 YP_009268869.1 --------------------------------------------------
011 YP_009506251.1 ---QLNYWE-------------------------QVTHTAWPMMAS----
012 NP_955611.1 --------------------------------------------------
013 YP_008567619.1 ---QNPTPGSTKA-------------------LKQALHQANQWLSS----
014 NP_149413.1 KKDMFERTEHIMMA---------------------TQEPTLLCGCR----
015 YP_004442824.1 MH-PKVFMYRYIKE------------------------PKFKCGCG----
016 YP_001931967.1 --------------------------------------------------
017 YP_002455786.1 --------------------------------------------------
018 YP_009109694.1 --------------------------------------------------
019 NP_789739.1 --------------------------------------------------
020 NP_955619.1 --------------------------------------------------
021 YP_224289.1 -KPAGIQINMITEG------------------------PKLRCACG----
022 YP_009182100.1 MPSPIPPPFEYKTYRRLTEEQKSQLDWLTSSWHTVDQDTFFDVLKAMAKE
023 NP_705927.1 --------------------------------------------------
024 YP_009508577.1 --------------------------------------------------
025 YP_004442830.1 QVTAKEVICMADVK---------------------EIAP--LCNCS----
026 NP_758887.1 --------------------------------------------------
027 YP_009508406.1 --------------------------------------------------
028 YP_610965.1 ---YASMASSRSS-----------------------PTLDDDLWCQ----
029 YP_001931961.1 --------------------------------------------------
030 YP_241114.1 --------------------------------------------------
031 YP_006732334.1 --------------------------------------------------
032 YP_009041481.1 ---SSSSTLKTED-----------------------KESLIQLSGI----
033 YP_001036293.1 ---ALSKEIRS---------------------------QVAHQISA----
034 YP_605811.1 RKEHMVNTLLQLEE------------------------PQLHCGCKDHTT
035 NP_955577.1 --------------------------------------------------
036 YP_003208050.1 ---RHKEALWQST-----------------------PKYEGPTGLM----
037 NP_789740.1 --------------------------------------------------
038 YP_004442839.1 KDEHPVNVLTLDEE------------------------TRLRCSCQDPIT
039 YP_009507791.1 --------------------------------------------------
040 YP_004581513.1 ---LRELSMKRPL-------------------EDKEDQGPWSILLL----
041 NP_043933.1 --------------------------------------------------
042 YP_009507248.1 --------------------------------------------------
043 YP_002117531.1 --QERIFHLTEVEG------------------------PALKCACN----
044 YP_004347415.1 --------------------------------------------------
045 NP_056891.1 --------------------------------------------------
046 YP_009513242.1 --------------------------------------------------
047 YP_009352866.1 ---SISSTLTRET-----------------------KESLIQLSGT----
048 YP_009165750.1 --------------------------------------------------
049 NP_040840.1 --------------------------------------------------
050 YP_004222728.1 --------------------------------------------------
051 YP_009506264.1 ------HIKVEEEA------------------------YMLAWPC-----
052 YP_567050.1 --------------------------------------------------
053 YP_009508571.1 --------------------------------------------------
054 NP_056803.1 --------------------------------------------------
055 NP_955564.1 --------------------------------------------------
056 YP_009508411.1 ---LLTHPSHE---------------------------ATMKGVKL----
057 YP_009140788.1 ---LMKEEQFKEK-------------------LPSLINKMIGCFSN----
058 YP_009508410.1 --------------------------------------------------
059 NP_040333.1 --------------------------------------------------
060 YP_008992013.1 ----------------------------------------DFTKKFL---
061 NP_041734.1 ---HLSFAG-------------------------LAIPIAWPMMGS----
062 NP_056895.1 --------------------------------------------------
063 YP_009109689.1 --------------------------------------------------
064 YP_002519387.1 --------------------------------------------------
065 YP_595725.1 KKDMFERTEQAFML---------------------TEEPVLLCACR----
066 YP_009508551.1 --------------------------------------------------
067 YP_001497148.1 --------------------------------------------------
068 NP_041186.1 --------------------------------------------------
069 NP_663784.1 KSSRNTGILEPNTRKRMALLSCSKINLVYRKVLDRCYPRLCRHPNT----
070 YP_009508443.1 ---VTSRPHAE---------------------------AMKKLATM----

```
071 YP_003987465.1   ------AVHVLTHK------------------------PILKCGCG----
072 NP_861410.1      --------------------------------------------------
073 YP_009345071.1   ---TQALMTSYLN-------------------TRSRSKTISKHAPN----
074 YP_009508582.1   --------------------------------------------------
075 NP_569141.1      --------------------------------------------------
076 YP_002916057.1   ---LREPPLKQEQ-------------------EG-TGEGPWGIEQL----
077 NP_777384.2      --------------------------------------------------
078 YP_233110.1      KEEHTVNTLLQLEE------------------------PQLHCGCRNYET
079 NP_542258.1      --------------------------------------------------
080 YP_009130664.1   ---AKPQPGKN--------------------------FQLTKLISS----
081 YP_009508566.1   --------------------------------------------------
082 YP_006607892.1   --------------------------------------------------
083 YP_009506270.1   FN-HHVNVLTIKEE------------------------PVFKCGCN----
084 NP_042513.1      --------------------------------------------------
085 YP_006907834.1   --------------------------------------------------
086 YP_004442827.1   SYEKIEICCMTETN---------------------ELAP--KCNCG----
087 YP_009408594.1   --------YCGLQ-----------------------PIIETGLKTS----
088 YP_009508561.1   --------------------------------------------------
089 NP_056907.1      --------------------------------------------------
090 YP_002308474.1   --------------------------------------------------
091 YP_007761644.1   --------------------------------------------------
092 NP_040550.1      --------------------------------------------------
093 NP_758808.1      AIHFLEDDKIEPT------------------AESSTPITLDEFSRK----
094 YP_009345075.1   ICLKTTSGICNKP-----------------------ATYHRTCNST----
095 NP_068729.1      --------------------------------------------------
096 NP_047255.1      --------------------------------------------------
097 YP_006273075.1   ---LMEDKQCRKQ-------------------LLPLINKMTNCFNN----
098 YP_009508888.1   --------------------------------------------------
099 NP_395469.1      --------------------------------------------------
100 YP_009508546.1   --------------------------------------------------
101 NP_056762.1      --------------------------------------------------
102 YP_009508556.1   --------------------------------------------------
103 NP_569153.1      ---HVEYVSLLIR-------------------SADRITRSLCFMRD----
104 NP_043924.1      FQRGSDSPSGGQ--------------------------------------
105 YP_009243641.1   --------------------------------------------------
106 YP_004300274.1   --------------------------------------------------
107 NP_056848.1      --------------------------------------------------
108 YP_006495799.1   ---PKERRILMDH-----------------------ICGMLEAQAQ----
109 YP_009508409.1   --------------------------------------------------
110 NP_687035.1      --------------------------------------------------
111 NP_045937.2      --------------------------------------------------
112 NP_612577.1      --------------------------------------------------
113 NP_619548.1      --------------------------------------------------
114 NP_056880.1      --------------------------------------------------
115 YP_009121747.1   QVLCIDTIKSEPTPPQYALKNQVSTQELPNNTKEKPPNKPSELCAT----
116 YP_009113237.2   ---KLKSKWSTEK-------------------IKELINLLIYNIKN----
117 YP_009508408.1   ---VMSRPNQA---------------------------AMTKLAEL----
118 NP_955579.1      --------------------------------------------------
119 YP_009229919.1   LAYHCIQGPSHKS-----------------------SSTESSLASS----
120 YP_004442833.1   EYKKLEVVCMAQTA---------------------DEVSPLLCNCE----
121 NP_862833.2      --------------------------------------------------
122 NP_040563.1      --------------------------------------------------
123 YP_443922.1      --------------------------------------------------
124 NP_057933.2      --------------------------------------------------
125 YP_009116631.1   ---P-AKKEVQEE-----------------------ISCFLEAALR----
126 YP_009513249.1   ----ETSVDSS---------------------------------------
127 YP_223871.1      --------------------------------------------------
128 NP_127504.1      LERRTFYELQVFKKYGGSILRPFGVDPEYPFAHIFIPNPTDFSEDLLWMF
129 NP_044929.1      TWSCEDRCGHRWKECGNCIPQDGSSDDASAVAAVEI--------------
130 NP_057860.1      --------------------------------------------------
131 NP_040973.1      --------------------------------------------------
132 NP_777317.1      EDENTTRVGQTYSVGTED-----------------LIQPTLNCACK----
133 NP_054716.1      ----QDQRGHT---------------------------------------
134 YP_001856242.1   --------------------------------------------------
135 NP_954565.2      --------------------------------------------------
136 NP_569150.1      YRLHQINVLTLSEE------------------------PVFKCGCN----
137 NP_659397.1      --------------------------------------------------
138 YP_009513211.1   --------------------------------------------------
139 YP_233107.1      -NKYQVNFMTWEQP-------------------------QLKCACG----
140 YP_009109692.1   --------------------------------------------------
141 NP_057849.4      --------------------------------------------------
142 NP_049560.1      QPVCSWTLDLLALSADQALQPPCPNLVSYSSYHATYSLYLFPHWIKKPNR
143 YP_009506267.1   ---IMKKPHQG---------------------------AIARIATL----
```

```
144 NP_039820.1         --------------------------------------------------
145 NP_056790.1         --------------------------------------------------
146 YP_002321513.1      ---SLKDTSLQIE----------------------LQESFLKKIYY----
147 NP_955591.1         --------------------------------------------------
148 YP_001956722.2      --------------------------------------------------
149 NP_056902.1         --------------------------------------------------
150 NP_056886.1         --------------------------------------------------
151 NP_041261.1         --------------------------------------------------



001 YP_004442836.1      ---KPARLHTSRTSRNPGREFYSCENK--ACFTWVWKDQIDE---YVQEV
002 NP_056728.1         --------------------------------------------------
003 YP_009508537.1      --------------------------------------------------
004 YP_004732983.2      ---LNHGR-----------PNTTWYSRTKPKKSKARKQAQVQLRFDVSND
005 YP_009508407.1      --------------------------------------------------
006 NP_040939.1         --------------------------------------------------
007 YP_003864102.1      --------------------------------------------------
008 YP_009002585.1      ---TMNSWTEKSLSSTQKEALLSDTRITSTNKWPLLKDKPVHKQNKHSN-
009 YP_003284237.1      ---IPAVQRESRTSRNPNRKFVTCRDNR--CRCWWWSDNIDDYTRRMIDL
010 YP_009268869.1      --------------------------------------------------
011 YP_009506251.1      ---CSNEGIPLHMAPSPWQLNMPSPDSSIRSRKKLPRKRSTHYET-----
012 NP_955611.1         --------------------------------------------------
013 YP_008567619.1      ---ISSTKMPLKDSQGLTAPAHENGGTTCASSKSLKEKPPKKPKKRWKSS
014 NP_149413.1         ---KPAIMLTSGTRLNPRRRFYKCAMNI--CHCWYWADLLEE---YVQER
015 YP_004442824.1      ---RKAEEVVSHTQKNPDRKFYRCKRS--NCHTWVWKDQIDN---YVQNY
016 YP_001931967.1      --------------------------------------------------
017 YP_002455786.1      --------------------------------------------------
018 YP_009109694.1      --------------------------------------------------
019 NP_789739.1         --------------------------------------------------
020 NP_955619.1         --------------------------------------------------
021 YP_224289.1         ---KDAEIAVSHTSRNPDRPFYKCQRNL--CHIWIWKDLVDD---YFQNL
022 YP_009182100.1      FEGLNKLKNGNPAKRVKTSREAKLARWKAQPKKFTFFEEWRTILLRRLRN
023 NP_705927.1         --------------------------------------------------
024 YP_009508577.1      --------------------------------------------------
025 YP_004442830.1      ---KPAMLRTSMTSRSPNRQFWNCAEKR--CHAWWWYDNLEG---YIKNE
026 NP_758887.1         --------------------------------------------------
027 YP_009508406.1      --------------------------------------------------
028 YP_610965.1         ---EQSKKAPLSTHCRGTGEHSTSLRPTNGEPCGPSSALQSSS--RNSRW
029 YP_001931961.1      --------------------------------------------------
030 YP_241114.1         --------------------------------------------------
031 YP_006732334.1      --------------------------------------------------
032 YP_009041481.1      ---VMDNWCALK-----SRHSSTALDSPGAQLKGKACPSKNMKSTDAAWP
033 YP_001036293.1      ---LLK-------IIDKGKSI-----------------------------
034 YP_605811.1         GERRNAVLLQSHTSANPYRWFYKCAQD--RCHTWIWKDILDQ---YAEDY
035 NP_955577.1         --------------------------------------------------
036 YP_003208050.1      ---IPLNEAQEWRAFARR-MRIAQAQKVDKELLGLLQMAQDKA--QYIED
037 NP_789740.1         --------------------------------------------------
038 YP_004442839.1      KQGMFAALLQSHTATNPGRWFYKCERN--RCHLWIWKDLLDQ---YANEY
039 YP_009507791.1      --------------------------------------------------
040 YP_004581513.1      ---TQPIEHLLKGSRNGQDRSKPITETGSMTSLKIALMILDLSPEVIIST
041 NP_043933.1         --------------------------------------------------
042 YP_009507248.1      --------------------------------------------------
043 YP_002117531.1      ---KEVLIKTSHTWRNPDRKFCVCPDGR--CHIWYWWDLLED---YVQQR
044 YP_004347415.1      --------------------------------------------------
045 NP_056891.1         --------------------------------------------------
046 YP_009513242.1      --------------------------------------------------
047 YP_009352866.1      ---VMETWCSQR-----LQLNSTAQNSQGQQQK-KVYPSKQQLNIGQGCP
048 YP_009165750.1      --------------------------------------------------
049 NP_040840.1         --------------------------------------------------
050 YP_004222728.1      --------------------------------------------------
051 YP_009506264.1      ----------------PKPNLLCSQDS--------------------LDI
052 YP_567050.1         --------------------------------------------------
053 YP_009508571.1      --------------------------------------------------
054 NP_056803.1         --------------------------------------------------
055 NP_955564.1         --------------------------------------------------
056 YP_009508411.1      ---LNQLMNITDSAIASEESEGLQLSRKKTKREDFSPC------------
057 YP_009140788.1      ---TRISQMKDE-PFIMKEYLSPMNKKSESSLELLKLSQQRKQSKLLRNI
058 YP_009508410.1      --------------------------------------------------
059 NP_040333.1         --------------------------------------------------
060 YP_008992013.1      --------------------------------------------------
061 NP_041734.1         ---YNKRRTPLLTGQSLWQRNKPSQHSSTASKSRQPRKHYWPYVT-----
062 NP_056895.1         --------------------------------------------------
```

```
063 YP_009109689.1   --------------------------------------------------
064 YP_002519387.1   --------------------------------------------------
065 YP_595725.1      ---KPAIQLVSRTSANPGRKFFKCAMNK--CHCWYWADLIEE---HIQDR
066 YP_009508551.1   --------------------------------------------------
067 YP_001497148.1   --------------------------------------------------
068 NP_041186.1      --------------------------------------------------
069 NP_663784.1      --------------------------------------------------
070 YP_009508443.1   ---IQG-------VLDSTHPRE----------------------------
071 YP_003987465.1   ---KTAIRKMSRTSRNPNRHYYCCEQEK--CHCWWWED-------HLLQF
072 NP_861410.1      --------------------------------------------------
073 YP_009345071.1   ---QGQWTNSSNLGPKSRTRSTNCTPKCVKNSQKYTQSLNSTGTQLDPRQ
074 YP_009508582.1   --------------------------------------------------
075 NP_569141.1      --------------------------------------------------
076 YP_002916057.1   ---TLPLERLLKGLKTGPYKNKLIIGTGLMNSLKIALRILDLSPEVIIST
077 NP_777384.2      --------------------------------------------------
078 YP_233110.1      GERRNAILLQSHTSANPNRWFYKCAEN--KCHIWIWKDILDQ---YAEDY
079 NP_542258.1      --------------------------------------------------
080 YP_009130664.1   ---LAG-----DGKRGRQLLQNTKGNGTASKTHDIVA-------------
081 YP_009508566.1   --------------------------------------------------
082 YP_006607892.1   --------------------------------------------------
083 YP_009506270.1   ---KPARLKTSRTSRNPNRQFYSCETK--SCFTWVWKDQIDT---FV---
084 NP_042513.1      --------------------------------------------------
085 YP_006907834.1   --------------------------------------------------
086 YP_004442827.1   ---QPAVLKISQTSKNPGRRFWTCEQRK--CHAWWWDDHLED---YIHAR
087 YP_009408594.1   ---LRGP------------LRQINMTRSPNGLEQLTRSPKMLSWQHWKKC
088 YP_009508561.1   --------------------------------------------------
089 NP_056907.1      --------------------------------------------------
090 YP_002308474.1   --------------------------------------------------
091 YP_007761644.1   --------------------------------------------------
092 NP_040550.1      --------------------------------------------------
093 NP_758808.1      ---RFQEHTDLLEEFQLTLLQINLLEAS---LHERLMKCQ-SYATRDNFW
094 YP_009345075.1   ---ASMKK-----------EQXTACRTNSXRYQNSCRNSEPWSTDHWPTI
095 NP_068729.1      --------------------------------------------------
096 NP_047255.1      --------------------------------------------------
097 YP_006273075.1   ---TRTLLRKEKGSFDQEEYLLPETMNSENNSGRLKKNLQEEPLKPWNNT
098 YP_009508888.1   --------------------------------------------------
099 NP_395469.1      --------------------------------------------------
100 YP_009508546.1   --------------------------------------------------
101 NP_056762.1      --------------------------------------------------
102 YP_009508556.1   --------------------------------------------------
103 NP_569153.1      ---SSHSRIYSCRPGKEPMKALICEQKSCQSKGDLGNTRTVHSKSAFNQQ
104 NP_043924.1      --------------------------------------------------
105 YP_009243641.1   --------------------------------------------------
106 YP_004300274.1   --------------------------------------------------
107 NP_056848.1      --------------------------------------------------
108 YP_006495799.1   ---KAS-------------LHQHGLSP-----------------------
109 YP_009508409.1   --------------------------------------------------
110 NP_687035.1      --------------------------------------------------
111 NP_045937.2      --------------------------------------------------
112 NP_612577.1      --------------------------------------------------
113 NP_619548.1      --------------------------------------------------
114 NP_056880.1      --------------------------------------------------
115 YP_009121747.1   ---SRQFWTSKPKSVLQSHQQITIGETIGPMLRNRISKPENSYQNLRPYA
116 YP_009113237.2   ---SSTRSRPRWPQEECHLASMTNCEETEMSWNHEQKLKQSGHCRPIALS
117 YP_009508408.1   ---IYCSE--NIIKLQISSKEDTLRLQNKMKEDAYSPY------------
118 NP_955579.1      --------------------------------------------------
119 YP_009229919.1   ---MKTNWNEPRTAWPHCYQWPISIWSEFNEKRPRTTGPQIVSPRRRKRS
120 YP_004442833.1   ---QPAGRQMSKTSRNPGRWFWSCVQRK--CHAWWWDDHLED---YIETQ
121 NP_862833.2      --------------------------------------------------
122 NP_040563.1      --------------------------------------------------
123 YP_443922.1      --------------------------------------------------
124 NP_057933.2      --------------------------------------------------
125 YP_009116631.1   ---RAKRSVTTHQSEPRHVLWQKWQNPEGWLYCDERRSSTALPNTSATRS
126 YP_009513249.1   --------------------------------------------------
127 YP_223871.1      --------------------------------------------------
128 NP_127504.1      WYLLNHFHILMKFRCSKFSKIDQVNPWMLKFLLWFNNHNYWASLFKCMKG
129 NP_044929.1      --------------------------------------------------
130 NP_057860.1      --------------------------------------------------
131 NP_040973.1      --------------------------------------------------
132 NP_777317.1      ---IKAQKLISRTTRNPDRAYYRCGLSPPKCYTWIWEDILEA---YVTER
133 NP_054716.1      --------------------------------------------------
134 YP_001856242.1   --------------------------------------------------
135 NP_954565.2      --------------------------------------------------
```

```
136 NP_569150.1        ---KPAKLKISRTSRNPDREFYSCETN--TCFTWVWKDQIDS---FVQEK
137 NP_659397.1        --------------------------------------------------
138 YP_009513211.1     --------------------------------------------------
139 YP_233107.1        ---ENAVLLTSHTSRNPGRRFYRCGTN--TCHVWYWADLIED---YIAQL
140 YP_009109692.1     --------------------------------------------------
141 NP_057849.4        --------------------------------------------------
142 NP_049560.1        NGGGYYSASYSDPCSLKCPYLGCQSWTCPYTGAISSPYWKFQQDVNFTQE
143 YP_009506267.1     ---VKK----MLEDVPTFPPRDA---------------------------
144 NP_039820.1        --------------DDHGRNLISAVIN---------------------DI
145 NP_056790.1        --------------------------------------------------
146 YP_002321513.1     ---LTPALECIDIPYIGGTMEMLSQPMLQQGDGYNTPGMSCLEETSLKET
147 NP_955591.1        --------------------------------------------------
148 YP_001956722.2     --------------------------------------------------
149 NP_056902.1        --------------------------------------------------
150 NP_056886.1        --------------------------------------------------
151 NP_041261.1        --------------------------------------------------


001 YP_004442836.1     MTWNDQVSQ----------------------------------------L
002 NP_056728.1        --------------------------------------------------
003 YP_009508537.1     --------------------------------------------------
004 YP_004732983.2     D-------------------------------------------------
005 YP_009508407.1     --------------------------------------------------
006 NP_040939.1        --------------------------------------------------
007 YP_003864102.1     --------------------------------------------------
008 YP_009002585.1     -NWKEYKDK-----------------------------------------
009 YP_003284237.1     AQEEADYDEAM--------------------------------------R
010 YP_009268869.1     --------------------------------------------------
011 YP_009506251.1     --YKGSYIT-----------------------------------------
012 NP_955611.1        --------------------------------------------------
013 YP_008567619.1     STSTN---------------------------------------------
014 NP_149413.1        IEDFMVENFDKKA--------------------------KLDEPSSSNVH
015 YP_004442824.1     INWLKESEEPEDEGYDPINRPRSLDSSPWSFDEPGSSQHQKEEEDERLLC
016 YP_001931967.1     --------------------------------------------------
017 YP_002455786.1     --------------------------------------------------
018 YP_009109694.1     --------------------------------------------------
019 NP_789739.1        --------------------------------------------------
020 NP_955619.1        --------------------------------------------------
021 YP_224289.1        TAWNRISEEHR--------------------------------------R
022 YP_009182100.1     WTNGATIMDEELYQRIINTPCRAQLQWWSQLNKLWQYQPPEQSFICKATC
023 NP_705927.1        --------------------------------------------------
024 YP_009508577.1     --------------------------------------------------
025 YP_004442830.1     VDKRLQR-----------------------------------------IL
026 NP_758887.1        --------------------------------------------------
027 YP_009508406.1     --------------------------------------------------
028 YP_610965.1        RHVTGSERV-----------------------------------------
029 YP_001931961.1     --------------------------------------------------
030 YP_241114.1        --------------------------------------------------
031 YP_006732334.1     --------------------------------------------------
032 YP_009041481.1     ASKTRNTKSSWYK-----------------------------------PP
033 YP_001036293.1     --------------------------------------------------
034 YP_605811.1        AAYTRIGLE-----------------------------------------
035 NP_955577.1        --------------------------------------------------
036 YP_003208050.1     -QRAGNNWA-----------------------------------------
037 NP_789740.1        --------------------------------------------------
038 YP_004442839.1     --YINMGLN-----------------------------------------
039 YP_009507791.1     --------------------------------------------------
040 YP_004581513.1     YGLKEGSLE-----------------------------------------
041 NP_043933.1        --------------------------------------------------
042 YP_009507248.1     --------------------------------------------------
043 YP_002117531.1     VTAAQLAECKN--------------------------------------F
044 YP_004347415.1     --------------------------------------------------
045 NP_056891.1        --------------------------------------------------
046 YP_009513242.1     --------------------------------------------------
047 YP_009352866.1     PLRTRNTRNSLPK-----------------------------------LQ
048 YP_009165750.1     --------------------------------------------------
049 NP_040840.1        --------------------------------------------------
050 YP_004222728.1     --------------------------------------------------
051 YP_009506264.1     ANDD----------------------------------------------
052 YP_567050.1        --------------------------------------------------
053 YP_009508571.1     --------------------------------------------------
054 NP_056803.1        --------------------------------------------------
```

```
055 NP_955564.1       -------------------------------------------------
056 YP_009508411.1    -------------------------------------------------
057 YP_009140788.1    SESMQSKHW----------------------------------------
058 YP_009508410.1    -------------------------------------------------
059 NP_040333.1       -------------------------------------------------
060 YP_008992013.1    -------------------------------------------------
061 NP_041734.1       --YRAYSTS----------------------------------------
062 NP_056895.1       -------------------------------------------------
063 YP_009109689.1    -------------------------------------------------
064 YP_002519387.1    -------------------------------------------------
065 YP_595725.1       IDEFLKN--------------------------------LEVLKTGGVQ
066 YP_009508551.1    -------------------------------------------------
067 YP_001497148.1    -------------------------------------------------
068 NP_041186.1       -------------------------------------------------
069 NP_663784.1       -------------------------------------------------
070 YP_009508443.1    -------------------------------------------------
071 YP_003987465.1    AAERGALEQEA-------------------------------------L
072 NP_861410.1       -------------------------------------------------
073 YP_009345071.1    QGIMLGETI----------------------------------------
074 YP_009508582.1    -------------------------------------------------
075 NP_569141.1       -------------------------------------------------
076 YP_002916057.1    YKLKEGYSE----------------------------------------
077 NP_777384.2       -------------------------------------------------
078 YP_233110.1       ATYTRIGLE----------------------------------------
079 NP_542258.1       -------------------------------------------------
080 YP_009130664.1    -------------------------------------------------
081 YP_009508566.1    -------------------------------------------------
082 YP_006607892.1    -------------------------------------------------
083 YP_009506270.1    -------------------------------------------------
084 NP_042513.1       -------------------------------------------------
085 YP_006907834.1    -------------------------------------------------
086 YP_004442827.1    IKMKLED----------------------------------------IQ
087 YP_009408594.1    PKYWRPRKT----------------------------------------
088 YP_009508561.1    -------------------------------------------------
089 NP_056907.1       -------------------------------------------------
090 YP_002308474.1    -------------------------------------------------
091 YP_007761644.1    -------------------------------------------------
092 NP_040550.1       -------------------------------------------------
093 NP_758808.1       GDWLPEARR----------------------------------------
094 YP_009345075.1    PASRQKTTM----------------------------------------
095 NP_068729.1       -------------------------------------------------
096 NP_047255.1       -------------------------------------------------
097 YP_006273075.1    GRFTHLKQP----------------------------------------
098 YP_009508888.1    -------------------------------------------------
099 NP_395469.1       -------------------------------------------------
100 YP_009508546.1    -------------------------------------------------
101 NP_056762.1       -------------------------------------------------
102 YP_009508556.1    -------------------------------------------------
103 NP_569153.1       DNWWPSTST----------------------------------------
104 NP_043924.1       -------------------------------------------------
105 YP_009243641.1    -------------------------------------------------
106 YP_004300274.1    -------------------------------------------------
107 NP_056848.1       -------------------------------------------------
108 YP_006495799.1    -------------------------------------------------
109 YP_009508409.1    -------------------------------------------------
110 NP_687035.1       -------------------------------------------------
111 NP_045937.2       -------------------------------------------------
112 NP_612577.1       -------------------------------------------------
113 NP_619548.1       -------------------------------------------------
114 NP_056880.1       -------------------------------------------------
115 YP_009121747.1    SSWAKSQSKPKES----------------------------------KC
116 YP_009113237.2    --DKQSRTG----------------------------------------
117 YP_009508408.1    -------------------------------------------------
118 NP_955579.1       -------------------------------------------------
119 YP_009229919.1    KGWSRPTRR----------------------------------------
120 YP_004442833.1    VELRLAK----------------------------------------IF
121 NP_862833.2       -------------------------------------------------
122 NP_040563.1       -------------------------------------------------
123 YP_443922.1       -------------------------------------------------
124 NP_057933.2       -------------------------------------------------
125 YP_009116631.1    SSPESTLRQQRPE----------------------------------QP
126 YP_009513249.1    -------------------------------------------------
127 YP_223871.1       -------------------------------------------------
```

```
128 NP_127504.1         IKKYVVIWFYRPVNYYQGKLCALPHSSIVKWNHVSVLNDEDEYSELQRFI
129 NP_044929.1         --------------------------------------------------
130 NP_057860.1         --------------------------------------------------
131 NP_040973.1         --------------------------------------------------
132 NP_777317.1         IIRTRQEQE-------------------------------------IDIL
133 NP_054716.1         --------------------------------------------------
134 YP_001856242.1      --------------------------------------------------
135 NP_954565.2         --------------------------------------------------
136 NP_569150.1         IRWEKKLEE----------------------------------------I
137 NP_659397.1         --------------------------------------------------
138 YP_009513211.1      --------------------------------------------------
139 YP_233107.1         SNLQNLDSG----------------------------------------Q
140 YP_009109692.1      --------------------------------------------------
141 NP_057849.4         --------------------------------------------------
142 NP_049560.1         VSRLTINLHFSKCGFPFSLLIDAPGYDPIWFLNTEPSQLPPTAPPLLPHS
143 YP_009506267.1      --------------------------------------------------
144 NP_039820.1         ITVLRR--------------------------------------------
145 NP_056790.1         --------------------------------------------------
146 YP_002321513.1      SQPALSTFTLG---------------------------------------
147 NP_955591.1         --------------------------------------------------
148 YP_001956722.2      --------------------------------------------------
149 NP_056902.1         --------------------------------------------------
150 NP_056886.1         --------------------------------------------------
151 NP_041261.1         --------------------------------------------------


001 YP_004442836.1      SE-----------EPEGHNEGC-------------------------TIE
002 NP_056728.1         --------------------------------------------------
003 YP_009508537.1      --------------------------------------------------
004 YP_004732983.2      --------------------------------------------------
005 YP_009508407.1      --------------------------------------------------
006 NP_040939.1         --------------------------------------------------
007 YP_003864102.1      --------------------------------------------------
008 YP_009002585.1      --------RRTTSGSQRQETTG-------------------------PVT
009 YP_003284237.1      RYHEELQEKKLDLMQINEALEA-------------------------AED
010 YP_009268869.1      --------------------------------------------------
011 YP_009506251.1      --------SATTCQALQQGTTG-------------------------RLT
012 NP_955611.1         --------------------------------------------------
013 YP_008567619.1      --------------------------------------------------
014 NP_149413.1         HDDYEEHRSSVIDRPRPTDDHFRPWGDVTYWLNKEAEECHTRGDNVEGAE
015 YP_004442824.1      KETMGEDYNESEQEVIRNNQAT-------------------------AIE
016 YP_001931967.1      --------------------------------------------------
017 YP_002455786.1      --------------------------------------------------
018 YP_009109694.1      --------------------------------------------------
019 NP_789739.1         --------------------------------------------------
020 NP_955619.1         --------------------------------------------------
021 YP_224289.1         EMAREEGQNLEEEDYWENVFNE-------------------------VFD
022 YP_009182100.1      FDNNHTMLKLKCPPINKLTPEFMDWFENGFIQSFALKRTSDYSLLPTDFI
023 NP_705927.1         --------------------------------------------------
024 YP_009508577.1      --------------------------------------------------
025 YP_004442830.1      AEQD-EEFEELCEIYKELEQQE--------------------ILRNQATD
026 NP_758887.1         --------------------------------------------------
027 YP_009508406.1      --------------------------------------------------
028 YP_610965.1         --------IMHGATHTPTCGTS-------------------------MNS
029 YP_001931961.1      --------------------------------------------------
030 YP_241114.1         --------------------------------------------------
031 YP_006732334.1      --------------------------------------------------
032 YP_009041481.1      QLTRSSARNWRPPGDAPVDETM-------------------------PTG
033 YP_001036293.1      --------------------------------------------------
034 YP_605811.1         ALSLEDWFEEPEPDPPDPVDRQ-------------------------KIE
035 NP_955577.1         --------------------------------------------------
036 YP_003208050.1      --------TDAWPNYLEDVNVI-------------------------HDV
037 NP_789740.1         --------------------------------------------------
038 YP_004442839.1      SLLLNMDWPDEESDPPDSPSPA-------------------------RTE
039 YP_009507791.1      --------------------------------------------------
040 YP_004581513.1      --------KACQEMWVHNSKQA-------------------------DRP
041 NP_043933.1         --------------------------------------------------
042 YP_009507248.1      --------------------------------------------------
043 YP_002117531.1      RQQLDNIKRQLDSDKRNRAHSI----------------------------
044 YP_004347415.1      --------------------------------------------------
045 NP_056891.1         --------------------------------------------------
046 YP_009513242.1      --------------------------------------------------
```

```
047 YP_009352866.1   QLTPSSVKSWKPPKAVQVAGII-------------------------TTE
048 YP_009165750.1   --------------------------------------------------
049 NP_040840.1      --------------------------------------------------
050 YP_004222728.1   --------------------------------------------------
051 YP_009506264.1   --------------------------------------------------
052 YP_567050.1      --------------------------------------------------
053 YP_009508571.1   --------------------------------------------------
054 NP_056803.1      --------------------------------------------------
055 NP_955564.1      --------------------------------------------------
056 YP_009508411.1   --------------------------------------------------
057 YP_009140788.1   --------NANHGVHQDETEII-------------------------GPT
058 YP_009508410.1   --------------------------------------------------
059 NP_040333.1      --------------------------------------------------
060 YP_008992013.1   --------------------------------------------------
061 NP_041734.1      --------RETIWPLLPLETTG-------------------------LAT
062 NP_056895.1      --------------------------------------------------
063 YP_009109689.1   --------------------------------------------------
064 YP_002519387.1   --------------------------------------------------
065 YP_595725.1      T--MEEELMKEVTKLKIEEQEFEEY------------QATPRAMSPVAAE
066 YP_009508551.1   --------------------------------------------------
067 YP_001497148.1   --------------------------------------------------
068 NP_041186.1      --------------------------------------------------
069 NP_663784.1      --------------------------------------------------
070 YP_009508443.1   --------------------------------------------------
071 YP_003987465.1   RMEELFDRLQFN---TNEPHGG-------------------------PLE
072 NP_861410.1      --------------------------------------------------
073 YP_009345071.1   --------SPTSKEKPKPPTIT-------------------------SVS
074 YP_009508582.1   --------------------------------------------------
075 NP_569141.1      --------------------------------------------------
076 YP_002916057.1   --------MGSQEMSVHNLQLI-------------------------DSP
077 NP_777384.2      --------------------------------------------------
078 YP_233110.1      ALNLEDWFEEPEPDPPNPVDRQ-------------------------RIE
079 NP_542258.1      --------------------------------------------------
080 YP_009130664.1   --------------------------------------------------
081 YP_009508566.1   --------------------------------------------------
082 YP_006607892.1   --------------------------------------------------
083 YP_009506270.1   --------------------------------------------------
084 NP_042513.1      --------------------------------------------------
085 YP_006907834.1   --------------------------------------------------
086 YP_004442827.1   KEQNPYDFEDWDDLLKET-------------------------QEEYKLE
087 YP_009408594.1   --------IFKCRQ------------------------------------
088 YP_009508561.1   --------------------------------------------------
089 NP_056907.1      --------------------------------------------------
090 YP_002308474.1   --------------------------------------------------
091 YP_007761644.1   --------------------------------------------------
092 NP_040550.1      --------------------------------------------------
093 NP_758808.1      --------DLLQIQLAKEIIEK-------------------------VRE
094 YP_009345075.1   --------LGTTKKHSNMHRVR-------------------------F--
095 NP_068729.1      --------------------------------------------------
096 NP_047255.1      --------------------------------------------------
097 YP_006273075.1   --------NVHHGVHPEGMGII-------------------------GLI
098 YP_009508888.1   --------------------------------------------------
099 NP_395469.1      --------------------------------------------------
100 YP_009508546.1   --------------------------------------------------
101 NP_056762.1      --------------------------------------------------
102 YP_009508556.1   --------------------------------------------------
103 NP_569153.1      --------NSLTSEAKLQGTTH-------------------------MPI
104 NP_043924.1      --------------------------------------------------
105 YP_009243641.1   --------------------------------------------------
106 YP_004300274.1   --------------------------------------------------
107 NP_056848.1      --------------------------------------------------
108 YP_006495799.1   --------------------------------------------------
109 YP_009508409.1   --------------------------------------------------
110 NP_687035.1      --------------------------------------------------
111 NP_045937.2      --------------------------------------------------
112 NP_612577.1      --------------------------------------------------
113 NP_619548.1      --------------------------------------------------
114 NP_056880.1      --------------------------------------------------
115 YP_009121747.1   PGAHAWDEEIARRQQLHNRSMK-------------------------YNQ
116 YP_009113237.2   --------VQGTHPGITSMVTH-------------------------YPP
117 YP_009508408.1   --------------------------------------------------
118 NP_955579.1      --------------------------------------------------
119 YP_009229919.1   --------SQGTSPSSEEKDVR-------------------------DDV
```

```
120 YP_004442833.1   KEQGTQEYEESMEKLVIKDNSPLPYQE-----NPEDAYQADFIQEDYQPE
121 NP_862833.2      --------------------------------------------------
122 NP_040563.1      --------------------------------------------------
123 YP_443922.1      --------------------------------------------------
124 NP_057933.2      --------------------------------------------------
125 YP_009116631.1   GTTGMLTSHQHWNDEPPQHGSS-------------------------WPL
126 YP_009513249.1   --------------------------------------------------
127 YP_223871.1      --------------------------------------------------
128 NP_127504.1      FQENKCIPKEIWPGSSGSWNYGNSDHPHGQWIRDALREYREMNDYFQDAQ
129 NP_044929.1      --------------------------------------------------
130 NP_057860.1      --------------------------------------------------
131 NP_040973.1      --------------------------------------------------
132 NP_777317.1      SDDNEAGINDLQNTTYDPGDDN-------------------------NWE
133 NP_054716.1      --------------------------------------------------
134 YP_001856242.1   --------------------------------------------------
135 NP_954565.2      --------------------------------------------------
136 NP_569150.1      SEDSLWEELLREQENLRAEQEY-------------------------LIE
137 NP_659397.1      --------------------------------------------------
138 YP_009513211.1   --------------------------------------------------
139 YP_233107.1      ADDEGWAYQTEDLINPEDLANS-------------------------DID
140 YP_009109692.1   --------------------------------------------------
141 NP_057849.4      --------------------------------------------------
142 NP_049560.1      NLDHILEPSIPWKSKLLTPVQLTLQSTNYTCIVCIDRASLSTWHVLYFPN
143 YP_009506267.1   --------------------------------------------------
144 NP_039820.1      --------------------------------------------------
145 NP_056790.1      --------------------------------------------------
146 YP_002321513.1   ---------KTEFVKPEKPSSS-------------------------YVT
147 NP_955591.1      --------------------------------------------------
148 YP_001956722.2   --------------------------------------------------
149 NP_056902.1      --------------------------------------------------
150 NP_056886.1      --------------------------------------------------
151 NP_041261.1      --------------------------------------------------
```

```
001 YP_004442836.1   DAFDLLDVSNDDQWARS---------------------------------
002 NP_056728.1      --------------------------------------------------
003 YP_009508537.1   --------------------------------------------------
004 YP_004732983.2   --------------------------------------------------
005 YP_009508407.1   --------------------------------------------------
006 NP_040939.1      --------------------------------------------------
007 YP_003864102.1   --------------------------------------------------
008 YP_009002585.1   GYHPP---------------------------------------------
009 YP_003284237.1   QDRREGVISDADFEPNVIRDD-----------------------------
010 YP_009268869.1   --------------------------------------------------
011 YP_009506251.1   DSH-----------------------------------------------
012 NP_955611.1      --------------------------------------------------
013 YP_008567619.1   --------------------------------------------------
014 NP_149413.1      DAVDLTDVSNDDQWRRS---------------------------------
015 YP_004442824.1   DALDLLDVSNDDQWGRS---------------------------------
016 YP_001931967.1   --------------------------------------------------
017 YP_002455786.1   --------------------------------------------------
018 YP_009109694.1   --------------------------------------------------
019 NP_789739.1      --------------------------------------------------
020 NP_955619.1      --------------------------------------------------
021 YP_224289.1      HEEITEFYPDGGDPG-----------------------------------
022 YP_009182100.1   KVIEQIWAIKDIGNTLFIMMYSAPGEWNYKDCSYYYPYHLVRIYDRVPSV
023 NP_705927.1      --------------------------------------------------
024 YP_009508577.1   --------------------------------------------------
025 YP_004442830.1   DPLDWNEVSNDD--------------------------------------
026 NP_758887.1      --------------------------------------------------
027 YP_009508406.1   --------------------------------------------------
028 YP_610965.1      SY-ASLDNSKS---------------------------------------
029 YP_001931961.1   --------------------------------------------------
030 YP_241114.1      --------------------------------------------------
031 YP_006732334.1   --------------------------------------------------
032 YP_009041481.1   TSSL----------------------------------------------
033 YP_001036293.1   --------------------------------------------------
034 YP_605811.1      DILDLQDVSNDD--------------------------------------
035 NP_955577.1      --------------------------------------------------
036 YP_003208050.1   RR-KIQNITRH---------------------------------------
037 NP_789740.1      --------------------------------------------------
038 YP_004442839.1   DIFDLANVSNDD--------------------------------------
```

```
039 YP_009507791.1    --------------------------------------------------
040 YP_004581513.1    YCMRRPRRPGEA--------------------------------------
041 NP_043933.1       --------------------------------------------------
042 YP_009507248.1    --------------------------------------------------
043 YP_002117531.1    --------------------------------------------------
044 YP_004347415.1    --------------------------------------------------
045 NP_056891.1       --------------------------------------------------
046 YP_009513242.1    --------------------------------------------------
047 YP_009352866.1    ICSQPWKKGKG---------------------------------------
048 YP_009165750.1    --------------------------------------------------
049 NP_040840.1       --------------------------------------------------
050 YP_004222728.1    --------------------------------------------------
051 YP_009506264.1    --------------------------------------------------
052 YP_567050.1       --------------------------------------------------
053 YP_009508571.1    --------------------------------------------------
054 NP_056803.1       --------------------------------------------------
055 NP_955564.1       --------------------------------------------------
056 YP_009508411.1    --------------------------------------------------
057 YP_009140788.1    TFQMSRDQTRISKGSFTISEKNCLMP------------------------
058 YP_009508410.1    --------------------------------------------------
059 NP_040333.1       --------------------------------------------------
060 YP_008992013.1    --------------------------------------------------
061 NP_041734.1       DCQLPNKTQPP---------------------------------------
062 NP_056895.1       --------------------------------------------------
063 YP_009109689.1    --------------------------------------------------
064 YP_002519387.1    --------------------------------------------------
065 YP_595725.1       DVLDLQDVSNDD--------------------------------------
066 YP_009508551.1    --------------------------------------------------
067 YP_001497148.1    --------------------------------------------------
068 NP_041186.1       --------------------------------------------------
069 NP_663784.1       --------------------------------------------------
070 YP_009508443.1    --------------------------------------------------
071 YP_003987465.1    DDDQASLLDAVGL-------------------------------------
072 NP_861410.1       --------------------------------------------------
073 YP_009345071.1    --------------------------------------------------
074 YP_009508582.1    --------------------------------------------------
075 NP_569141.1       --------------------------------------------------
076 YP_002916057.1    YCMKRPRRPGEAS-------------------------------------
077 NP_777384.2       --------------------------------------------------
078 YP_233110.1       DILDLLNVSNDD--------------------------------------
079 NP_542258.1       --------------------------------------------------
080 YP_009130664.1    --------------------------------------------------
081 YP_009508566.1    --------------------------------------------------
082 YP_006607892.1    --------------------------------------------------
083 YP_009506270.1    --------------------------------------------------
084 NP_042513.1       --------------------------------------------------
085 YP_006907834.1    --------------------------------------------------
086 YP_004442827.1    DALDLDDFMNDDQWRRA---------------------------------
087 YP_009408594.1    --------------------------------------------------
088 YP_009508561.1    --------------------------------------------------
089 NP_056907.1       --------------------------------------------------
090 YP_002308474.1    --------------------------------------------------
091 YP_007761644.1    --------------------------------------------------
092 NP_040550.1       --------------------------------------------------
093 NP_758808.1       KLHSI---------------------------------------------
094 YP_009345075.1    --------------------------------------------------
095 NP_068729.1       --------------------------------------------------
096 NP_047255.1       --------------------------------------------------
097 YP_006273075.1    ACQALGDMTQD---------------------------------------
098 YP_009508888.1    --------------------------------------------------
099 NP_395469.1       --------------------------------------------------
100 YP_009508546.1    --------------------------------------------------
101 NP_056762.1       --------------------------------------------------
102 YP_009508556.1    --------------------------------------------------
103 NP_569153.1       GYPHAIGTTSNCVKWSSY--------------------------------
104 NP_043924.1       --------------------------------------------------
105 YP_009243641.1    --------------------------------------------------
106 YP_004300274.1    --------------------------------------------------
107 NP_056848.1       --------------------------------------------------
108 YP_006495799.1    --------------------------------------------------
109 YP_009508409.1    --------------------------------------------------
110 NP_687035.1       --------------------------------------------------
111 NP_045937.2       --------------------------------------------------
```

```
112 NP_612577.1          --------------------------------------------------
113 NP_619548.1          --------------------------------------------------
114 NP_056880.1          --------------------------------------------------
115 YP_009121747.1       ALWLIDESWGPPLSMPRYCKL-----------------------------
116 YP_009113237.2       CCTGMNNSPPTRML------------------------------------
117 YP_009508408.1       --------------------------------------------------
118 NP_955579.1          --------------------------------------------------
119 YP_009229919.1       KYPKTSGMTKNI--------------------------------------
120 YP_004442833.1       DVLDLDDFTNDDQWRRS---------------------------------
121 NP_862833.2          --------------------------------------------------
122 NP_040563.1          --------------------------------------------------
123 YP_443922.1          --------------------------------------------------
124 NP_057933.2          --------------------------------------------------
125 YP_009116631.1       TRNSPRVRM-----------------------------------------
126 YP_009513249.1       --------------------------------------------------
127 YP_223871.1          --------------------------------------------------
128 NP_127504.1          DPYPAYSKVDLTQEELNTLRITRSYGSSSEDADMVKRSIYTVQSNIVKDS
129 NP_044929.1          --------------------------------------------------
130 NP_057860.1          --------------------------------------------------
131 NP_040973.1          --------------------------------------------------
132 NP_777317.1          DAVDISAYDSIDAFWDAHT-------------------------------
133 NP_054716.1          --------------------------------------------------
134 YP_001856242.1       --------------------------------------------------
135 NP_954565.2          --------------------------------------------------
136 NP_569150.1          DALDLLDISNDD--------------------------------------
137 NP_659397.1          --------------------------------------------------
138 YP_009513211.1       --------------------------------------------------
139 YP_233107.1          DPPEDSGLFHRHDD------------------------------------
140 YP_009109692.1       --------------------------------------------------
141 NP_057849.4          --------------------------------------------------
142 NP_049560.1          VSVPSSPSTPLLYPSLALPAPHLTLPFNWTHCFNPQIQAIVSSPCHNSLI
143 YP_009506267.1       --------------------------------------------------
144 NP_039820.1          --------------------------------------------------
145 NP_056790.1          --------------------------------------------------
146 YP_002321513.1       TMRKK---------------------------------------------
147 NP_955591.1          --------------------------------------------------
148 YP_001956722.2       --------------------------------------------------
149 NP_056902.1          --------------------------------------------------
150 NP_056886.1          --------------------------------------------------
151 NP_041261.1          --------------------------------------------------
```

```
001 YP_004442836.1       --------------------------------------------------
002 NP_056728.1          --------------------------------------------------
003 YP_009508537.1       --------------------------------------------------
004 YP_004732983.2       --------------------------------------------------
005 YP_009508407.1       --------------------------------------------------
006 NP_040939.1          --------------------------------------------------
007 YP_003864102.1       --------------------------------------------------
008 YP_009002585.1       --------------------------------------------------
009 YP_003284237.1       --------------------------------------------------
010 YP_009268869.1       --------------------------------------------------
011 YP_009506251.1       --------------------------------------------------
012 NP_955611.1          --------------------------------------------------
013 YP_008567619.1       --------------------------------------------------
014 NP_149413.1          --------------------------------------------------
015 YP_004442824.1       --------------------------------------------------
016 YP_001931967.1       --------------------------------------------------
017 YP_002455786.1       --------------------------------------------------
018 YP_009109694.1       --------------------------------------------------
019 NP_789739.1          --------------------------------------------------
020 NP_955619.1          --------------------------------------------------
021 YP_224289.1          --------------------------------------------------
022 YP_009182100.1       TRDTILEGEIFPHLRSQEQIQRDEEMGMYEPLYCPIDDWLSRDPRHVSKW
023 NP_705927.1          --------------------------------------------------
024 YP_009508577.1       --------------------------------------------------
025 YP_004442830.1       --------------------------------------------------
026 NP_758887.1          --------------------------------------------------
027 YP_009508406.1       --------------------------------------------------
028 YP_610965.1          --------------------------------------------------
029 YP_001931961.1       --------------------------------------------------
030 YP_241114.1          --------------------------------------------------
```

```
031 YP_006732334.1    --------------------------------------------------
032 YP_009041481.1    --------------------------------------------------
033 YP_001036293.1    --------------------------------------------------
034 YP_605811.1       --------------------------------------------------
035 NP_955577.1       --------------------------------------------------
036 YP_003208050.1    --------------------------------------------------
037 NP_789740.1       --------------------------------------------------
038 YP_004442839.1    --------------------------------------------------
039 YP_009507791.1    --------------------------------------------------
040 YP_004581513.1    --------------------------------------------------
041 NP_043933.1       --------------------------------------------------
042 YP_009507248.1    --------------------------------------------------
043 YP_002117531.1    --------------------------------------------------
044 YP_004347415.1    --------------------------------------------------
045 NP_056891.1       --------------------------------------------------
046 YP_009513242.1    --------------------------------------------------
047 YP_009352866.1    --------------------------------------------------
048 YP_009165750.1    --------------------------------------------------
049 NP_040840.1       --------------------------------------------------
050 YP_004222728.1    --------------------------------------------------
051 YP_009506264.1    --------------------------------------------------
052 YP_567050.1       --------------------------------------------------
053 YP_009508571.1    --------------------------------------------------
054 NP_056803.1       --------------------------------------------------
055 NP_955564.1       --------------------------------------------------
056 YP_009508411.1    --------------------------------------------------
057 YP_009140788.1    --------------------------------------------------
058 YP_009508410.1    --------------------------------------------------
059 NP_040333.1       --------------------------------------------------
060 YP_008992013.1    --------------------------------------------------
061 NP_041734.1       --------------------------------------------------
062 NP_056895.1       --------------------------------------------------
063 YP_009109689.1    --------------------------------------------------
064 YP_002519387.1    --------------------------------------------------
065 YP_595725.1       --------------------------------------------------
066 YP_009508551.1    --------------------------------------------------
067 YP_001497148.1    --------------------------------------------------
068 NP_041186.1       --------------------------------------------------
069 NP_663784.1       --------------------------------------------------
070 YP_009508443.1    --------------------------------------------------
071 YP_003987465.1    --------------------------------------------------
072 NP_861410.1       --------------------------------------------------
073 YP_009345071.1    --------------------------------------------------
074 YP_009508582.1    --------------------------------------------------
075 NP_569141.1       --------------------------------------------------
076 YP_002916057.1    --------------------------------------------------
077 NP_777384.2       --------------------------------------------------
078 YP_233110.1       --------------------------------------------------
079 NP_542258.1       --------------------------------------------------
080 YP_009130664.1    --------------------------------------------------
081 YP_009508566.1    --------------------------------------------------
082 YP_006607892.1    --------------------------------------------------
083 YP_009506270.1    --------------------------------------------------
084 NP_042513.1       --------------------------------------------------
085 YP_006907834.1    --------------------------------------------------
086 YP_004442827.1    --------------------------------------------------
087 YP_009408594.1    --------------------------------------------------
088 YP_009508561.1    --------------------------------------------------
089 NP_056907.1       --------------------------------------------------
090 YP_002308474.1    --------------------------------------------------
091 YP_007761644.1    --------------------------------------------------
092 NP_040550.1       --------------------------------------------------
093 NP_758808.1       --------------------------------------------------
094 YP_009345075.1    --------------------------------------------------
095 NP_068729.1       --------------------------------------------------
096 NP_047255.1       --------------------------------------------------
097 YP_006273075.1    --------------------------------------------------
098 YP_009508888.1    --------------------------------------------------
099 NP_395469.1       --------------------------------------------------
100 YP_009508546.1    --------------------------------------------------
101 NP_056762.1       --------------------------------------------------
102 YP_009508556.1    --------------------------------------------------
103 NP_569153.1       --------------------------------------------------
```

```
104 NP_043924.1        --------------------------------------------------
105 YP_009243641.1     --------------------------------------------------
106 YP_004300274.1     --------------------------------------------------
107 NP_056848.1        --------------------------------------------------
108 YP_006495799.1     --------------------------------------------------
109 YP_009508409.1     --------------------------------------------------
110 NP_687035.1        --------------------------------------------------
111 NP_045937.2        --------------------------------------------------
112 NP_612577.1        --------------------------------------------------
113 NP_619548.1        --------------------------------------------------
114 NP_056880.1        --------------------------------------------------
115 YP_009121747.1     --------------------------------------------------
116 YP_009113237.2     --------------------------------------------------
117 YP_009508408.1     --------------------------------------------------
118 NP_955579.1        --------------------------------------------------
119 YP_009229919.1     --------------------------------------------------
120 YP_004442833.1     --------------------------------------------------
121 NP_862833.2        --------------------------------------------------
122 NP_040563.1        --------------------------------------------------
123 YP_443922.1        --------------------------------------------------
124 NP_057933.2        --------------------------------------------------
125 YP_009116631.1     --------------------------------------------------
126 YP_009513249.1     --------------------------------------------------
127 YP_223871.1        --------------------------------------------------
128 NP_127504.1        PRKRKGKAKSRSSTRSEKRRAKNKCKYRSLHGEDWWIELGYSTKPSTPSW
129 NP_044929.1        --------------------------------------------------
130 NP_057860.1        --------------------------------------------------
131 NP_040973.1        --------------------------------------------------
132 NP_777317.1        --------------------------------------------------
133 NP_054716.1        --------------------------------------------------
134 YP_001856242.1     --------------------------------------------------
135 NP_954565.2        --------------------------------------------------
136 NP_569150.1        --------------------------------------------------
137 NP_659397.1        --------------------------------------------------
138 YP_009513211.1     --------------------------------------------------
139 YP_233107.1        --------------------------------------------------
140 YP_009109692.1     --------------------------------------------------
141 NP_057849.4        --------------------------------------------------
142 NP_049560.1        LPPFSLSPVPTLGSRSRRAVPVAVWLVSALAMGAGMAGGITGSMSLASGR
143 YP_009506267.1     --------------------------------------------------
144 NP_039820.1        --------------------------------------------------
145 NP_056790.1        --------------------------------------------------
146 YP_002321513.1     --------------------------------------------------
147 NP_955591.1        --------------------------------------------------
148 YP_001956722.2     --------------------------------------------------
149 NP_056902.1        --------------------------------------------------
150 NP_056886.1        --------------------------------------------------
151 NP_041261.1        --------------------------------------------------


001 YP_004442836.1     --------------------------------------------------
002 NP_056728.1        --------------------------------------------------
003 YP_009508537.1     --------------------------------------------------
004 YP_004732983.2     --------------------------------------------------
005 YP_009508407.1     --------------------------------------------------
006 NP_040939.1        --------------------------------------------------
007 YP_003864102.1     --------------------------------------------------
008 YP_009002585.1     --------------------------------------------------
009 YP_003284237.1     --------------------------------------------------
010 YP_009268869.1     --------------------------------------------------
011 YP_009506251.1     --------------------------------------------------
012 NP_955611.1        --------------------------------------------------
013 YP_008567619.1     --------------------------------------------------
014 NP_149413.1        --------------------------------------------------
015 YP_004442824.1     --------------------------------------------------
016 YP_001931967.1     --------------------------------------------------
017 YP_002455786.1     --------------------------------------------------
018 YP_009109694.1     --------------------------------------------------
019 NP_789739.1        --------------------------------------------------
020 NP_955619.1        --------------------------------------------------
021 YP_224289.1        --------------------------------------------------
022 YP_009182100.1     RAFALMEIQDLINEDKGFLPRWHYIHSNQRFIFEVEGISMTRACESYSLF
```

023 NP_705927.1 --------------------------------------------------
024 YP_009508577.1 --------------------------------------------------
025 YP_004442830.1 --------------------------------------------------
026 NP_758887.1 --------------------------------------------------
027 YP_009508406.1 --------------------------------------------------
028 YP_610965.1 --------------------------------------------------
029 YP_001931961.1 --------------------------------------------------
030 YP_241114.1 --------------------------------------------------
031 YP_006732334.1 --------------------------------------------------
032 YP_009041481.1 --------------------------------------------------
033 YP_001036293.1 --------------------------------------------------
034 YP_605811.1 --------------------------------------------------
035 NP_955577.1 --------------------------------------------------
036 YP_003208050.1 --------------------------------------------------
037 NP_789740.1 --------------------------------------------------
038 YP_004442839.1 --------------------------------------------------
039 YP_009507791.1 --------------------------------------------------
040 YP_004581513.1 --------------------------------------------------
041 NP_043933.1 --------------------------------------------------
042 YP_009507248.1 --------------------------------------------------
043 YP_002117531.1 --------------------------------------------------
044 YP_004347415.1 --------------------------------------------------
045 NP_056891.1 --------------------------------------------------
046 YP_009513242.1 --------------------------------------------------
047 YP_009352866.1 --------------------------------------------------
048 YP_009165750.1 --------------------------------------------------
049 NP_040840.1 --------------------------------------------------
050 YP_004222728.1 --------------------------------------------------
051 YP_009506264.1 --------------------------------------------------
052 YP_567050.1 --------------------------------------------------
053 YP_009508571.1 --------------------------------------------------
054 NP_056803.1 --------------------------------------------------
055 NP_955564.1 --------------------------------------------------
056 YP_009508411.1 --------------------------------------------------
057 YP_009140788.1 --------------------------------------------------
058 YP_009508410.1 --------------------------------------------------
059 NP_040333.1 --------------------------------------------------
060 YP_008992013.1 --------------------------------------------------
061 NP_041734.1 --------------------------------------------------
062 NP_056895.1 --------------------------------------------------
063 YP_009109689.1 --------------------------------------------------
064 YP_002519387.1 --------------------------------------------------
065 YP_595725.1 --------------------------------------------------
066 YP_009508551.1 --------------------------------------------------
067 YP_001497148.1 --------------------------------------------------
068 NP_041186.1 --------------------------------------------------
069 NP_663784.1 --------------------------------------------------
070 YP_009508443.1 --------------------------------------------------
071 YP_003987465.1 --------------------------------------------------
072 NP_861410.1 --------------------------------------------------
073 YP_009345071.1 --------------------------------------------------
074 YP_009508582.1 --------------------------------------------------
075 NP_569141.1 --------------------------------------------------
076 YP_002916057.1 --------------------------------------------------
077 NP_777384.2 --------------------------------------------------
078 YP_233110.1 --------------------------------------------------
079 NP_542258.1 --------------------------------------------------
080 YP_009130664.1 --------------------------------------------------
081 YP_009508566.1 --------------------------------------------------
082 YP_006607892.1 --------------------------------------------------
083 YP_009506270.1 --------------------------------------------------
084 NP_042513.1 --------------------------------------------------
085 YP_006907834.1 --------------------------------------------------
086 YP_004442827.1 --------------------------------------------------
087 YP_009408594.1 --------------------------------------------------
088 YP_009508561.1 --------------------------------------------------
089 NP_056907.1 --------------------------------------------------
090 YP_002308474.1 --------------------------------------------------
091 YP_007761644.1 --------------------------------------------------
092 NP_040550.1 --------------------------------------------------
093 NP_758808.1 --------------------------------------------------
094 YP_009345075.1 --------------------------------------------------
095 NP_068729.1 --------------------------------------------------

```
096 NP_047255.1        --------------------------------------------------
097 YP_006273075.1     --------------------------------------------------
098 YP_009508888.1     --------------------------------------------------
099 NP_395469.1        --------------------------------------------------
100 YP_009508546.1     --------------------------------------------------
101 NP_056762.1        --------------------------------------------------
102 YP_009508556.1     --------------------------------------------------
103 NP_569153.1        --------------------------------------------------
104 NP_043924.1        --------------------------------------------------
105 YP_009243641.1     --------------------------------------------------
106 YP_004300274.1     --------------------------------------------------
107 NP_056848.1        --------------------------------------------------
108 YP_006495799.1     --------------------------------------------------
109 YP_009508409.1     --------------------------------------------------
110 NP_687035.1        --------------------------------------------------
111 NP_045937.2        --------------------------------------------------
112 NP_612577.1        --------------------------------------------------
113 NP_619548.1        --------------------------------------------------
114 NP_056880.1        --------------------------------------------------
115 YP_009121747.1     --------------------------------------------------
116 YP_009113237.2     --------------------------------------------------
117 YP_009508408.1     --------------------------------------------------
118 NP_955579.1        --------------------------------------------------
119 YP_009229919.1     --------------------------------------------------
120 YP_004442833.1     --------------------------------------------------
121 NP_862833.2        --------------------------------------------------
122 NP_040563.1        --------------------------------------------------
123 YP_443922.1        --------------------------------------------------
124 NP_057933.2        --------------------------------------------------
125 YP_009116631.1     --------------------------------------------------
126 YP_009513249.1     --------------------------------------------------
127 YP_223871.1        --------------------------------------------------
128 NP_127504.1        TQDSSSEPCV----------------------------------------
129 NP_044929.1        --------------------------------------------------
130 NP_057860.1        --------------------------------------------------
131 NP_040973.1        --------------------------------------------------
132 NP_777317.1        --------------------------------------------------
133 NP_054716.1        --------------------------------------------------
134 YP_001856242.1     --------------------------------------------------
135 NP_954565.2        --------------------------------------------------
136 NP_569150.1        --------------------------------------------------
137 NP_659397.1        --------------------------------------------------
138 YP_009513211.1     --------------------------------------------------
139 YP_233107.1        --------------------------------------------------
140 YP_009109692.1     --------------------------------------------------
141 NP_057849.4        --------------------------------------------------
142 NP_049560.1        SLLHEVDKDISQLTQAIVKNHKNLLKIAQYAAQNRRGLDLLFWEQGGLCK
143 YP_009506267.1     --------------------------------------------------
144 NP_039820.1        --------------------------------------------------
145 NP_056790.1        --------------------------------------------------
146 YP_002321513.1     --------------------------------------------------
147 NP_955591.1        --------------------------------------------------
148 YP_001956722.2     --------------------------------------------------
149 NP_056902.1        --------------------------------------------------
150 NP_056886.1        --------------------------------------------------
151 NP_041261.1        --------------------------------------------------
```

```
001 YP_004442836.1     --------------------------------------------------
002 NP_056728.1        --------------------------------------------------
003 YP_009508537.1     --------------------------------------------------
004 YP_004732983.2     --------------------------------------------------
005 YP_009508407.1     --------------------------------------------------
006 NP_040939.1        --------------------------------------------------
007 YP_003864102.1     --------------------------------------------------
008 YP_009002585.1     --------------------------------------------------
009 YP_003284237.1     --------------------------------------------------
010 YP_009268869.1     --------------------------------------------------
011 YP_009506251.1     --------------------------------------------------
012 NP_955611.1        --------------------------------------------------
013 YP_008567619.1     --------------------------------------------------
014 NP_149413.1        --------------------------------------------------
```

```
015 YP_004442824.1   --------------------------------------------------
016 YP_001931967.1   --------------------------------------------------
017 YP_002455786.1   --------------------------------------------------
018 YP_009109694.1   --------------------------------------------------
019 NP_789739.1      --------------------------------------------------
020 NP_955619.1      --------------------------------------------------
021 YP_224289.1      --------------------------------------------------
022 YP_009182100.1   QSLSLKVPMNYDTQLEIDCIQTLDAAQSEDPCDEPYDESDSYWQDMDDET
023 NP_705927.1      --------------------------------------------------
024 YP_009508577.1   --------------------------------------------------
025 YP_004442830.1   --------------------------------------------------
026 NP_758887.1      --------------------------------------------------
027 YP_009508406.1   --------------------------------------------------
028 YP_610965.1      --------------------------------------------------
029 YP_001931961.1   --------------------------------------------------
030 YP_241114.1      --------------------------------------------------
031 YP_006732334.1   --------------------------------------------------
032 YP_009041481.1   --------------------------------------------------
033 YP_001036293.1   --------------------------------------------------
034 YP_605811.1      --------------------------------------------------
035 NP_955577.1      --------------------------------------------------
036 YP_003208050.1   --------------------------------------------------
037 NP_789740.1      --------------------------------------------------
038 YP_004442839.1   --------------------------------------------------
039 YP_009507791.1   --------------------------------------------------
040 YP_004581513.1   --------------------------------------------------
041 NP_043933.1      --------------------------------------------------
042 YP_009507248.1   --------------------------------------------------
043 YP_002117531.1   --------------------------------------------------
044 YP_004347415.1   --------------------------------------------------
045 NP_056891.1      --------------------------------------------------
046 YP_009513242.1   --------------------------------------------------
047 YP_009352866.1   --------------------------------------------------
048 YP_009165750.1   --------------------------------------------------
049 NP_040840.1      --------------------------------------------------
050 YP_004222728.1   --------------------------------------------------
051 YP_009506264.1   --------------------------------------------------
052 YP_567050.1      --------------------------------------------------
053 YP_009508571.1   --------------------------------------------------
054 NP_056803.1      --------------------------------------------------
055 NP_955564.1      --------------------------------------------------
056 YP_009508411.1   --------------------------------------------------
057 YP_009140788.1   --------------------------------------------------
058 YP_009508410.1   --------------------------------------------------
059 NP_040333.1      --------------------------------------------------
060 YP_008992013.1   --------------------------------------------------
061 NP_041734.1      --------------------------------------------------
062 NP_056895.1      --------------------------------------------------
063 YP_009109689.1   --------------------------------------------------
064 YP_002519387.1   --------------------------------------------------
065 YP_595725.1      --------------------------------------------------
066 YP_009508551.1   --------------------------------------------------
067 YP_001497148.1   --------------------------------------------------
068 NP_041186.1      --------------------------------------------------
069 NP_663784.1      --------------------------------------------------
070 YP_009508443.1   --------------------------------------------------
071 YP_003987465.1   --------------------------------------------------
072 NP_861410.1      --------------------------------------------------
073 YP_009345071.1   --------------------------------------------------
074 YP_009508582.1   --------------------------------------------------
075 NP_569141.1      --------------------------------------------------
076 YP_002916057.1   --------------------------------------------------
077 NP_777384.2      --------------------------------------------------
078 YP_233110.1      --------------------------------------------------
079 NP_542258.1      --------------------------------------------------
080 YP_009130664.1   --------------------------------------------------
081 YP_009508566.1   --------------------------------------------------
082 YP_006607892.1   --------------------------------------------------
083 YP_009506270.1   --------------------------------------------------
084 NP_042513.1      --------------------------------------------------
085 YP_006907834.1   --------------------------------------------------
086 YP_004442827.1   --------------------------------------------------
087 YP_009408594.1   --------------------------------------------------
```

```
088 YP_009508561.1     --------------------------------------------------
089 NP_056907.1        --------------------------------------------------
090 YP_002308474.1     --------------------------------------------------
091 YP_007761644.1     --------------------------------------------------
092 NP_040550.1        --------------------------------------------------
093 NP_758808.1        --------------------------------------------------
094 YP_009345075.1     --------------------------------------------------
095 NP_068729.1        --------------------------------------------------
096 NP_047255.1        --------------------------------------------------
097 YP_006273075.1     --------------------------------------------------
098 YP_009508888.1     --------------------------------------------------
099 NP_395469.1        --------------------------------------------------
100 YP_009508546.1     --------------------------------------------------
101 NP_056762.1        --------------------------------------------------
102 YP_009508556.1     --------------------------------------------------
103 NP_569153.1        --------------------------------------------------
104 NP_043924.1        --------------------------------------------------
105 YP_009243641.1     --------------------------------------------------
106 YP_004300274.1     --------------------------------------------------
107 NP_056848.1        --------------------------------------------------
108 YP_006495799.1     --------------------------------------------------
109 YP_009508409.1     --------------------------------------------------
110 NP_687035.1        --------------------------------------------------
111 NP_045937.2        --------------------------------------------------
112 NP_612577.1        --------------------------------------------------
113 NP_619548.1        --------------------------------------------------
114 NP_056880.1        --------------------------------------------------
115 YP_009121747.1     --------------------------------------------------
116 YP_009113237.2     --------------------------------------------------
117 YP_009508408.1     --------------------------------------------------
118 NP_955579.1        --------------------------------------------------
119 YP_009229919.1     --------------------------------------------------
120 YP_004442833.1     --------------------------------------------------
121 NP_862833.2        --------------------------------------------------
122 NP_040563.1        --------------------------------------------------
123 YP_443922.1        --------------------------------------------------
124 NP_057933.2        --------------------------------------------------
125 YP_009116631.1     --------------------------------------------------
126 YP_009513249.1     --------------------------------------------------
127 YP_223871.1        --------------------------------------------------
128 NP_127504.1        --------------------------------------------------
129 NP_044929.1        --------------------------------------------------
130 NP_057860.1        --------------------------------------------------
131 NP_040973.1        --------------------------------------------------
132 NP_777317.1        --------------------------------------------------
133 NP_054716.1        --------------------------------------------------
134 YP_001856242.1     --------------------------------------------------
135 NP_954565.2        --------------------------------------------------
136 NP_569150.1        --------------------------------------------------
137 NP_659397.1        --------------------------------------------------
138 YP_009513211.1     --------------------------------------------------
139 YP_233107.1        --------------------------------------------------
140 YP_009109692.1     --------------------------------------------------
141 NP_057849.4        --------------------------------------------------
142 NP_049560.1        ALQEQCCFLNITNSHVSILQERPPLENRVLTGWGLNWDLGLSQWAREALQ
143 YP_009506267.1     --------------------------------------------------
144 NP_039820.1        --------------------------------------------------
145 NP_056790.1        --------------------------------------------------
146 YP_002321513.1     --------------------------------------------------
147 NP_955591.1        --------------------------------------------------
148 YP_001956722.2     --------------------------------------------------
149 NP_056902.1        --------------------------------------------------
150 NP_056886.1        --------------------------------------------------
151 NP_041261.1        --------------------------------------------------


001 YP_004442836.1      --------------------------------------------
002 NP_056728.1         --------------------------------------------
003 YP_009508537.1      --------------------------------------------
004 YP_004732983.2      --------------------------------------------
005 YP_009508407.1      --------------------------------------------
006 NP_040939.1         --------------------------------------------
```

```
007 YP_003864102.1     --------------------------------------------
008 YP_009002585.1     --------------------------------------------
009 YP_003284237.1     --------------------------------------------
010 YP_009268869.1     --------------------------------------------
011 YP_009506251.1     --------------------------------------------
012 NP_955611.1        --------------------------------------------
013 YP_008567619.1     --------------------------------------------
014 NP_149413.1        --------------------------------------------
015 YP_004442824.1     --------------------------------------------
016 YP_001931967.1     --------------------------------------------
017 YP_002455786.1     --------------------------------------------
018 YP_009109694.1     --------------------------------------------
019 NP_789739.1        --------------------------------------------
020 NP_955619.1        --------------------------------------------
021 YP_224289.1        --------------------------------------------
022 YP_009182100.1     WHNIENMMEG----------------------------------
023 NP_705927.1        --------------------------------------------
024 YP_009508577.1     --------------------------------------------
025 YP_004442830.1     --------------------------------------------
026 NP_758887.1        --------------------------------------------
027 YP_009508406.1     --------------------------------------------
028 YP_610965.1        --------------------------------------------
029 YP_001931961.1     --------------------------------------------
030 YP_241114.1        --------------------------------------------
031 YP_006732334.1     --------------------------------------------
032 YP_009041481.1     --------------------------------------------
033 YP_001036293.1     --------------------------------------------
034 YP_605811.1        --------------------------------------------
035 NP_955577.1        --------------------------------------------
036 YP_003208050.1     --------------------------------------------
037 NP_789740.1        --------------------------------------------
038 YP_004442839.1     --------------------------------------------
039 YP_009507791.1     --------------------------------------------
040 YP_004581513.1     --------------------------------------------
041 NP_043933.1        --------------------------------------------
042 YP_009507248.1     --------------------------------------------
043 YP_002117531.1     --------------------------------------------
044 YP_004347415.1     --------------------------------------------
045 NP_056891.1        --------------------------------------------
046 YP_009513242.1     --------------------------------------------
047 YP_009352866.1     --------------------------------------------
048 YP_009165750.1     --------------------------------------------
049 NP_040840.1        --------------------------------------------
050 YP_004222728.1     --------------------------------------------
051 YP_009506264.1     --------------------------------------------
052 YP_567050.1        --------------------------------------------
053 YP_009508571.1     --------------------------------------------
054 NP_056803.1        --------------------------------------------
055 NP_955564.1        --------------------------------------------
056 YP_009508411.1     --------------------------------------------
057 YP_009140788.1     --------------------------------------------
058 YP_009508410.1     --------------------------------------------
059 NP_040333.1        --------------------------------------------
060 YP_008992013.1     --------------------------------------------
061 NP_041734.1        --------------------------------------------
062 NP_056895.1        --------------------------------------------
063 YP_009109689.1     --------------------------------------------
064 YP_002519387.1     --------------------------------------------
065 YP_595725.1        --------------------------------------------
066 YP_009508551.1     --------------------------------------------
067 YP_001497148.1     --------------------------------------------
068 NP_041186.1        --------------------------------------------
069 NP_663784.1        --------------------------------------------
070 YP_009508443.1     --------------------------------------------
071 YP_003987465.1     --------------------------------------------
072 NP_861410.1        --------------------------------------------
073 YP_009345071.1     --------------------------------------------
074 YP_009508582.1     --------------------------------------------
075 NP_569141.1        --------------------------------------------
076 YP_002916057.1     --------------------------------------------
077 NP_777384.2        --------------------------------------------
078 YP_233110.1        --------------------------------------------
079 NP_542258.1        --------------------------------------------
```

080 YP_009130664.1 --------------------------------------------
081 YP_009508566.1 --------------------------------------------
082 YP_006607892.1 --------------------------------------------
083 YP_009506270.1 --------------------------------------------
084 NP_042513.1 --------------------------------------------
085 YP_006907834.1 --------------------------------------------
086 YP_004442827.1 --------------------------------------------
087 YP_009408594.1 --------------------------------------------
088 YP_009508561.1 --------------------------------------------
089 NP_056907.1 --------------------------------------------
090 YP_002308474.1 --------------------------------------------
091 YP_007761644.1 --------------------------------------------
092 NP_040550.1 --------------------------------------------
093 NP_758808.1 --------------------------------------------
094 YP_009345075.1 --------------------------------------------
095 NP_068729.1 --------------------------------------------
096 NP_047255.1 --------------------------------------------
097 YP_006273075.1 --------------------------------------------
098 YP_009508888.1 --------------------------------------------
099 NP_395469.1 --------------------------------------------
100 YP_009508546.1 --------------------------------------------
101 NP_056762.1 --------------------------------------------
102 YP_009508556.1 --------------------------------------------
103 NP_569153.1 --------------------------------------------
104 NP_043924.1 --------------------------------------------
105 YP_009243641.1 --------------------------------------------
106 YP_004300274.1 --------------------------------------------
107 NP_056848.1 --------------------------------------------
108 YP_006495799.1 --------------------------------------------
109 YP_009508409.1 --------------------------------------------
110 NP_687035.1 --------------------------------------------
111 NP_045937.2 --------------------------------------------
112 NP_612577.1 --------------------------------------------
113 NP_619548.1 --------------------------------------------
114 NP_056880.1 --------------------------------------------
115 YP_009121747.1 --------------------------------------------
116 YP_009113237.2 --------------------------------------------
117 YP_009508408.1 --------------------------------------------
118 NP_955579.1 --------------------------------------------
119 YP_009229919.1 --------------------------------------------
120 YP_004442833.1 --------------------------------------------
121 NP_862833.2 --------------------------------------------
122 NP_040563.1 --------------------------------------------
123 YP_443922.1 --------------------------------------------
124 NP_057933.2 --------------------------------------------
125 YP_009116631.1 --------------------------------------------
126 YP_009513249.1 --------------------------------------------
127 YP_223871.1 --------------------------------------------
128 NP_127504.1 --------------------------------------------
129 NP_044929.1 --------------------------------------------
130 NP_057860.1 --------------------------------------------
131 NP_040973.1 --------------------------------------------
132 NP_777317.1 --------------------------------------------
133 NP_054716.1 --------------------------------------------
134 YP_001856242.1 --------------------------------------------
135 NP_954565.2 --------------------------------------------
136 NP_569150.1 --------------------------------------------
137 NP_659397.1 --------------------------------------------
138 YP_009513211.1 --------------------------------------------
139 YP_233107.1 --------------------------------------------
140 YP_009109692.1 --------------------------------------------
141 NP_057849.4 --------------------------------------------
142 NP_049560.1 TGITLVALLPLVILAGPCILRQLRHLPSRVRHPHYSLINPESCL
143 YP_009506267.1 --------------------------------------------
144 NP_039820.1 --------------------------------------------
145 NP_056790.1 --------------------------------------------
146 YP_002321513.1 --------------------------------------------
147 NP_955591.1 --------------------------------------------
148 YP_001956722.2 --------------------------------------------
149 NP_056902.1 --------------------------------------------
150 NP_056886.1 --------------------------------------------
151 NP_041261.1 --------------------------------------------

| 1 | 2 | 3 | 4 | 5 | 6 | 7 | 8 | 9 |
|---|---|---|---|---|---|---|---|---|

Variable Average Conserved

X - Insufficient data - the calculation for this site was performed on less than 10% of the sequences.
